# Eye structure shapes neuron function in *Drosophila* motion vision

**DOI:** 10.1101/2022.12.14.520178

**Authors:** Arthur Zhao, Eyal Gruntman, Aljoscha Nern, Nirmala A. Iyer, Edward M. Rogers, Sanna Koskela, Igor Siwanowicz, Marisa Dreher, Miriam A. Flynn, Connor W. Laughland, Henrique D.F. Ludwig, Alex G. Thomson, Cullen P. Moran, Bruck Gezahegn, Davi D. Bock, Michael B. Reiser

## Abstract

Many animals rely on vision to navigate through their environment. The pattern of changes in the visual scene induced by self-motion is the *optic flow*^1^, which is first estimated in local patches by directionally selective (DS) neurons^2–4^. But how should the arrays of DS neurons, each responsive to motion in a preferred direction at a specific retinal position, be organized to support robust decoding of optic flow by downstream circuits? Understanding this global organization is challenging because it requires mapping fine, local features of neurons across the animal’s field of view^3^. In *Drosophila*, the asymmetric dendrites of the T4 and T5 DS neurons establish their preferred direction, making it possible to predict DS responses from anatomy^4,5^. Here we report that the preferred directions of fly DS neurons vary at different retinal positions and show that this spatial variation is established by the anatomy of the compound eye. To estimate the preferred directions across the visual field, we reconstructed hundreds of T4 neurons in a full brain EM volume^6^ and discovered unexpectedly stereotypical dendritic arborizations that are independent of location. We then used whole-head μCT scans to map the viewing directions of all compound eye facets and found a non-uniform sampling of visual space that explains the spatial variation in preferred directions. Our findings show that the organization of preferred directions in the fly is largely determined by the compound eye, exposing an intimate and unexpected connection between the peripheral structure of the eye, functional properties of neurons deep in the brain, and the control of body movements.

## Main

By moving through an environment, seeing animals can determine the physical layout and estimate their own path using visual motion detection (Fig. 1A)^1^, analogous to solving the *structure from motion* problem in Computer Vision^7^. However, biological vision does not provide perfect geometric measurements. Instead, the global structure is synthesized using arrays of DS neurons that report relative motion in small regions of the scene. Insects are famously skilled at rapid flight maneuvers that depend on optic flow–the global structure of visual motion^8,9^. Recent progress *in Drosophila* has elucidated key aspects of the circuits computing motion detection as well as the visual control of navigation. Nevertheless, the intervening logic by which local motion detectors are spatially organized for reliable, behaviorally relevant optic flow estimation, remains unclear.

**Figure 1:**
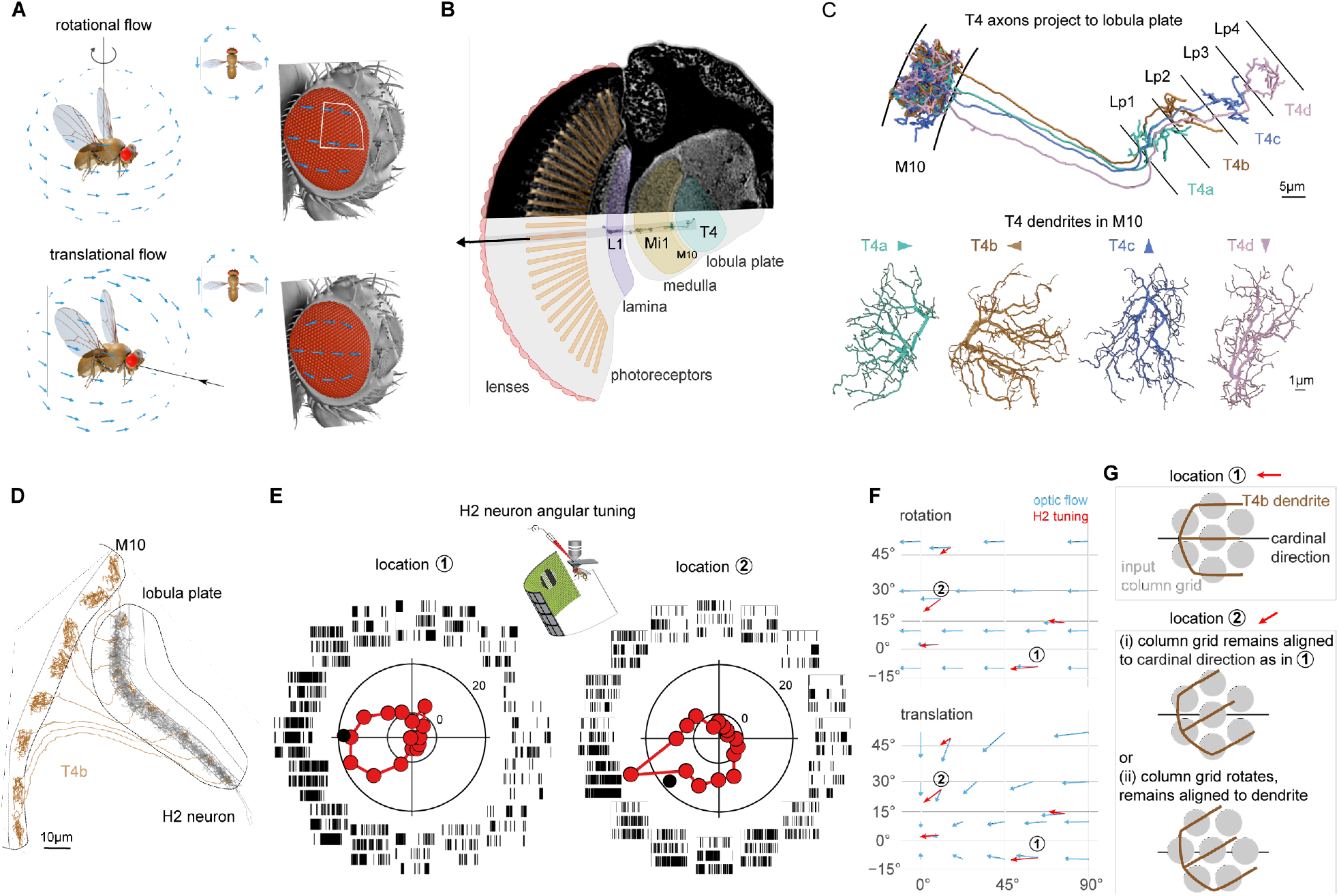
Non-cardinal direction preference by DS neurons. **A**. Ideal optic flow fields induced by yaw rotation or backwards translation, projected onto a model fly’s right eye. The local structure of the optic flow is similar near the eye’s equator, but different away from it. **B**. Columnar architecture of the fly’s compound eye. Top half: cross section of a μCT image stack, with neuropils of the visual system indicated. Bottom half: schematic drawing overlaid with EM reconstructed neuron skeletons in one column. The arrow illustrates that ommatidium’s viewing direction, and the long gray rectangle schematizes the corresponding, single column. **C**. Four subtypes of the direction selective (DS) T4 cells, receive inputs in the proximal medulla layer (M10), and project to one of the 4 lobula plate layers (Lp1-4). A T4 cell’s preferred direction (PD, arrowheads) of motion is roughly opposite to the primary orientation of its dendrites^4^. **D**. EM reconstruction of a wide-field H2 neuron’s dendrite that receives inputs from T4b cells across nearly the entire layer 2 (Lp2) of the lobula plate. The complete morphology of the H2 neuron is shown in Extended Data Fig. 1B. **E**. Electrophysiology recordings of H2 angular tuning. Raster plots show the spiking activity in response to local stimulation with square gratings moving in 16 directions at 2 different retina locations. Polar plots show average spiking response rate (in Hz). The black dot marks the PD, and the inset shows the experimental setup. **F**. Ideal optic flow fields overlaid with H2 local direction tuning in a Mercator projection. The plotted area corresponds to the region outlined in white on the eye in (**A**). **G**. Two potential mechanisms for how the dendrites of T4 neurons could establish different PDs at location ➁: (i) location-dependent changes in how the T4 dendrite samples the input column grid or (ii) consistent dendritic orientation (with respect to local input columnar grid) but visual space is non-uniformly represented.

Each fruit fly eye is composed ∼750 columnar units called ommatidia, arranged on a hemisphere to maximize the field of view^10^. Each ommatidium houses photoreceptors and collects light from a small area of visual space^10,11^. Along the motion pathway, columnar neurons such as L1 and Mi1, receive, modify, and transmit photoreceptor signals, preserving retinotopy^4,12,13^ (Fig. 1B). T4 neurons are the local ON-DS cells^14,15^, sensitive to bright edge movement (analogous T5 neurons are the OFF-DS cells^5,16–18^). T4s integrate columnar inputs along their dendrites, whose principal anatomical orientation corresponds to the neurons’ preferred direction (PD) of motion^4,5^ (Fig. 1C). There are four T4 subtypes, each with a distinct dendritic orientation, and an axon terminal projecting to one of four layers in the lobula plate^2,19^. These neurons are best understood near the center of the eye, where the PDs of each subtype align to one of four orthogonal, cardinal directions (forward, backward, up, and down)^2,4^. It is unclear how well this relationship holds for T4s away from the center. Indeed, due to the spherical geometry of the compound eye, the PDs cannot be globally aligned with the cardinal directions while also maintaining orthogonality between subtypes (Extended Data Fig. 1A). Since wide-field neurons in the lobula plate integrate from large ensembles of T4 neurons^19,20^, the directional tuning of T4s across the eye directly shapes global optic flow processing.

### Non-cardinal direction preference by DS neurons

To survey the directional preference of T4s across visual space, we measured the local PD of H2, a large, wide-field neuron that integrates from T4bs throughout the 2^nd^ layer of the lobula plate^19,21^ (Fig. 1D, Extended Data Fig. 1B). We used whole-cell electrophysiology to record H2 responses to gratings moving in 16 directions, at several locations on the eye (Fig. 1E,F). We find that near the eye’s equator, H2’s PD is aligned with cardinal, back-to-front motion, as previously reported^21–23^. However, at more dorso-frontal locations, the PD shows a prominent downward component (Fig. 1E,F; consistently across animals, Extended Data Fig. 1C,D). Surprisingly, these responses resembles a translational optic flow field (Fig. 1F), rather than a purely rotational one, as expected for H2 (blowflies^23^). This shift in the local PD of H2 implies that T4 neurons are not globally tuned to cardinal motion directions, a prediction that agrees well with a recent imaging study of T4/T5 axons^24^. But what causes T4 cells to change their directional preference across the eye?

Two parsimonious mechanisms could account for how T4 dendrites are differentially oriented with respect to each other at different retinotopic locations. Either T4 dendritic orientations vary with respect to their retinotopic inputs (Fig. 1G (i)) throughout the eye, or T4s dendrites employ a conserved integration strategy, but the representation of space by the array of input neurons is non-uniform (Fig. 1G (ii)). To distinguish between these two hypotheses, we reconstructed the morphology of hundreds of T4 neurons to determine the spatial integration pattern in the medulla. We then established a new, high-resolution map, detailing the spatial sampling by each ommatidium in the eye. By combining these datasets, we map T4s’ preferred directions into visual space, thereby revealing the mechanism underlying the non-cardinal motion sensitivity. Finally, our global analysis of the fly eye reveals principal axes of body movements that are most efficiently measured via optic flow.

### EM reconstruction of T4 dendrites across the eye reveals stereotypical arborization pattern

To compare T4 neurons’ arborization pattern across the entire medulla, we manually reconstructed all 779 Mi1 neurons on the right side of the Full Adult Fly Brain (FAFB) volume^6^ to establish a neuroanatomical coordinate system. Mi1s are columnar cells that are a major input to T4 neurons^4,15^ (Fig. 2A,B, Extended Data Fig. 2A). Their reconstruction was essential for propagating retinotopic coordinates from the more regular, distal layers to M10, where Mi1s synapse onto T4 dendrites (Fig. 2C). All Mi1s in M10 were then mapped into a 2D regular grid with the orthogonal +h and +v axes (Fig. 2D). Because the rows of Mi1s are not generally straight (Fig. 2C), capturing the global grid structure (Fig. 2D) enables the direct comparison of T4 neurons’ arborization pattern across the eye. We note two special rows of points that serve as global landmarks: the “equator” (Fig. 2D, in orange) is derived from the equatorial region in Fig. 2C, which is located via the corresponding lamina cartridges with additional photoreceptors (see Methods and Extended Data Fig. 2 B-D), and the “central meridian” (Fig. 2D, in black) that divides the points into roughly equal halves and coincides well with the first optic chiasm (Extended Data Fig. 2E). This regular grid mapping required access to the complete medulla and lamina neuropils in the EM volume, and further tracing of columnar neurons can extend this coordinate system into deeper neuropils, like the lobula (Extended Data Fig. 2F).

**Figure 2:**
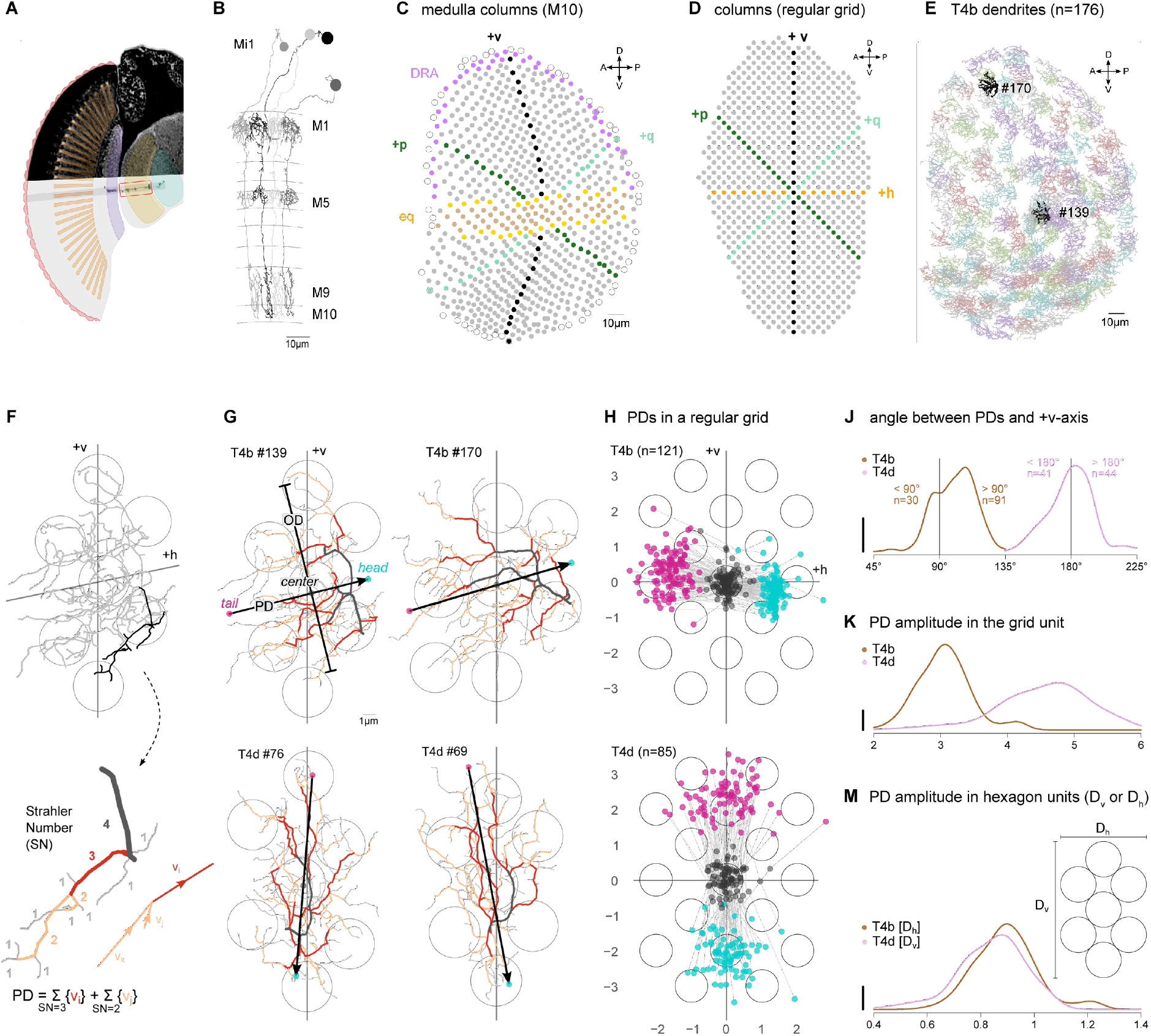
EM reconstruction of T4 dendrites across the eye reveals stereotypical arborization pattern. **A**. *Drosophila* visual system schematic highlighting an Mi1 and a T4 cell in medulla. **B**. EM reconstruction of 4 Mi1 cells arborizing primarily in medulla layer M1, M5 and M9/10. **C**. Medulla columns identified by the center-of-mass of the M10 arbors of Mi1 cells. Magenta dots at the top denote dorsal rim area columns^51^. The belt in the middle denotes the equatorial region, where there are 7 (yellow) or 8 (tan) photoreceptors in the corresponding lamina cartridges (Extended Data Fig. 2B). Black dots denote the “central meridian,” separating the points into approximately equal halves. Empty circles denote medulla columns with no R7/8 inputs, which presumably have no corresponding ommatidia^68^. **D**. Medulla columns mapped onto a 2D regular grid, with orthogonal +h and +v axes defined by the equatorial region and central meridian. Also noted +p and +q axes for consistency with prior work^28,70^. **E**. The dendritic arbors of 176 T4b cells in M10. Two highlighted example neurons are shown in (**G**). **F**. An example T4b (#139) dendrite. Bold branch (upper) is color coded by Strahler number (SN; lower). Arrows represent the direction vectors of SN = {2, 3} branches. The preferred direction (PD) of the dendrite is defined as the vector sum of all SN = {2, 3} branches. **G**. Example T4b and T4d cells’ dendrites, with preferred directions (PD) and orthogonal directions (OD). Branches are colored by their SN (> 3 in black). Seven circles in each plot represent the home column and its 6 nearest neighbors. **H**. Mapping PDs to a regular grid using 19 neighboring columns (see Methods, the tail, center, and head of each PD vector is indicated as in (**G**)). **J**. Distribution of the angles between T4 PDs and +v-axis. The scale bar in histogram plots spans from zero to the height of a uniform distribution. **K**. Distribution of PD amplitudes in units of the regular grid for T4b and T4d cells (significantly different, Wilcoxon rank test, p-value < 2.2e-16). **M**. Distribution of PD amplitudes normalized by respective hexagon length units defined in inset (overlapping, though significantly different, Wilcoxon rank test, p-value=0.015).

Since the orientation of T4 dendrites corresponds to their PD (Fig. 1C), we reconstructed the complete dendritic morphology of 176 horizontal-motion-sensitive T4b cells (Fig. 2E), and 114 vertical-motion-sensitive T4d cells (Extended Data Fig. 2G). We applied branching analysis developed for river networks^25^ to each T4’s dendritic tree to capture the primary orientation (Fig. 2F, Extended Data Fig. 3A) as an anatomical PD estimate. This estimate yields a PD vector that is represented as an arrow going through the dendrite’s center-of-mass, with a length corresponding to the spatial extent of the dendrite along the PD (Fig. 2G).

While the dendritic tree of each T4 neuron is idiosyncratic in its fine features, many conserved characteristics, such as the size and dominant branch orientation, suggest these neurons may be more stereotyped than expected from visual inspection of their morphology. To examine potential stereotypy, we transformed all T4 PD vectors (and their orthogonal directions, ODs) into the regular grid of Mi1s (Fig. 2D,H), using kernel regression that maintains the spatial relationships between each PD and its neighboring Mi1s (excluding edge T4s, see methods). Once transformed, the PD vectors for both T4 subtypes show a high degree of similarity. First, the centers of mass for all T4 dendrites fall within a “home” column. Second, the heads and tails of the PD vectors are each localized to a small area (the standard deviation of the head and tail positions is less than ½ the inter-column distance). Third, the dendrites of both subtypes roughly span a single unit hexagon (1 home + 6 nearest columns). T4b’s and T4d’s PD vectors are mostly aligned with the +h and -v axes, respectively (Fig. 2J). The bias (>90º) in the T4b distribution is mostly accounted for by neurons below the equator (Extended Data Fig. 3B,C). The PD vector lengths between the subtypes are notably different (Fig. 2K and Extended Data Fig. 3D). However, the unit hexagon is anisotropic since its height is greater than the width. Assuming that columns are space-filing, we defined new unit distances, “hexagon unit,” as the edge-to-edge span: 3 horizontal columns for T4b (D_h_) and 5 vertical columns (D_v_) for T4d (Fig. 2M inset). When we normalize the PD length by these new unit distances for each subtype separately, we find that T4b’s and T4d’s are now highly overlapping (Fig. 2M, Extended Data Fig. 3E). Since we identified the T4 subtypes based on lobula plate layer innervation, the striking within-subtype similarity of the PDs, does not support further divisions based on morphology^19,24,26^. Our analysis has thus revealed that T4 neurons share a universal sampling strategy—throughout the eye they innervate a unit hexagon of columns, while establishing a preferred direction by aligning their dendrites mostly in one direction, parallel to either the horizontal or vertical axes of the hexagonal grid.

### Non-uniform sampling of visual space established by μCT of the *Drosophila* eye

Having established that T4’s PD is governed by a simple local rule that is conserved throughout the medulla (strong evidence against the hypothesis in Fig. 1G(i)), understanding the global PD organization now reduces to understanding how visual space, sampled by the compound eye, maps onto the array of medulla columns (required to evaluate the hypothesis in Fig. 1G(ii)). Since the EM volume did not contain the eye, we instead imaged whole fly heads with approximately the same number of ommatidia. We first explored confocal imaging (Extended Data Fig. 4A), but ultimately used micro computed tomography (μCT; Fig. 3A,B). The isotropic ∼1 μm resolution of the μCT data allowed us to define the viewing direction of each ommatidium (as the vector connecting the ‘tip’ of the photoreceptors to the center of each corneal lens, Fig. 3B,C, Extended Data Fig. 4B), and to locate the eye’s equator (using the chirality of the photoreceptor arrangement^27^, Fig. 3D).

**Figure 3:**
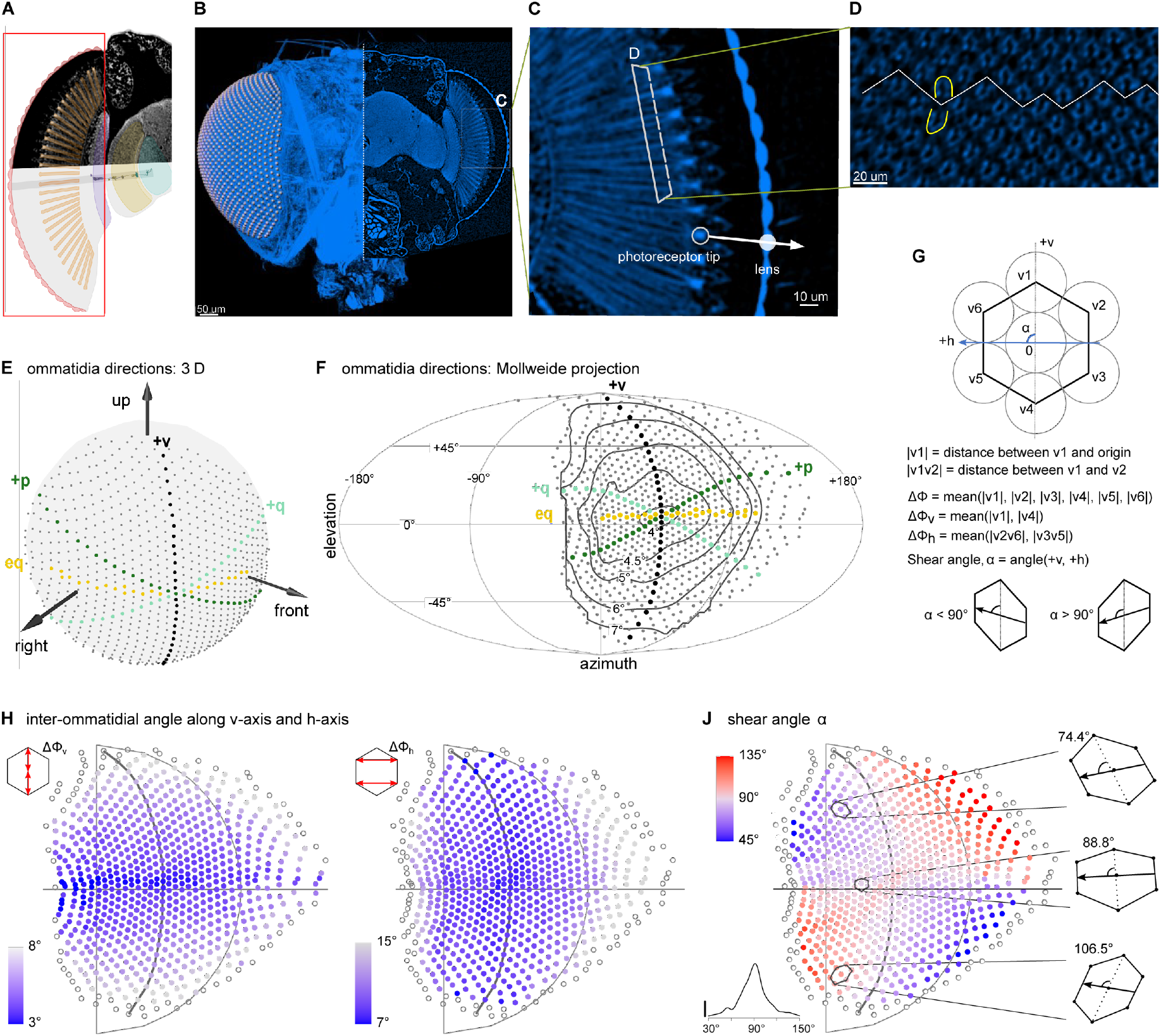
Non-uniform sampling of visual space established by μCT of the *Drosophila* eye. **A**. *Drosophila* visual system schematic highlighting the retina. **B**. Maximal intensity projection of a whole fly head scanned with μCT. Lenses on the right eye (left half) are labeled by white spheres. **C**. A zoomed-in cross section showing lenses and photoreceptors, with an example tip-lens pair that defines the viewing direction of that ommatidium. **D**. Photoreceptors in each ommatidium are arranged in an “n” or “u” shape above or below the equator, respectively^27^. **E**. Right eye ommatidia directions represented by points on a unit sphere. The “eq” row of points is based on (**D**) and “+v” separates the points into approximately equal halves. **F**. Mollweide projection of 3D ommatidia directions, and the inter-ommatidial angles (IOA, ΔΦ, averaged over 6 neighbors). Contour lines label iso-IOA levels. **G**. A schematic unit hexagon containing 7 columns (home column at the origin plus v1-6). The +h-axis is the line from the center of 2 right neighbors to that of 2 left neighbors, and the +v-axis as the line from the bottom neighbor to the top one. We define the 6-neighbor IOA ΔΦ, vertical IOA ΔΦ_v_, horizontal IOA ΔΦ_h_, and shear angle α. Because of the small angle approximation, we determine the IOAs using the Euclidean distance (|·|) of points on the unit sphere in (**E**). **H**. Spatial distribution of ΔΦ_v_ and ΔΦ_h_. Points represent ommatidia directions as in (**F**). **J**. Distribution of shear angles across the eye, with 3 example unit hexagons from the same vertical grid line. The inset plot is the histogram of all shear angles. In (**H**) and (**J)**, points lacking the complete set of neighbors for each calculation are displayed as empty circles. Also note that, compared with (**F**), points not matched to medulla columns (see Fig. 4A) are excluded. The 3 examples are aligned with the meridian lines through the home column.

We represent the ommatidia directions in eye coordinates on a unit sphere (Fig. 3E) or with a geographic projection (Mollweide projection, Fig. 3F; Mercator projection, Extended Data Fig. 5F-J). The field of view of this eye spans from directly above to directly below (−90° to 90°) in elevation, and in azimuth, from ∼20° into the opposite hemisphere in front to 160° behind, with a binocular overlap of ∼40° (Extended Data Fig. 4C,D). The maps of ommatidia directions produced from 3 different females are quite consistent (Extended Data Fig. 4E), and show greater binocular overlap than prior data based on a coarser, optical method^28^. The ommatidia directions are well described by a hexagonal grid that we then aligned to the medulla column grid using the equator (+h) and central meridian (+v) as global landmarks (Extended Data Fig. 5A, Fig. 4A).

**Figure 4:**
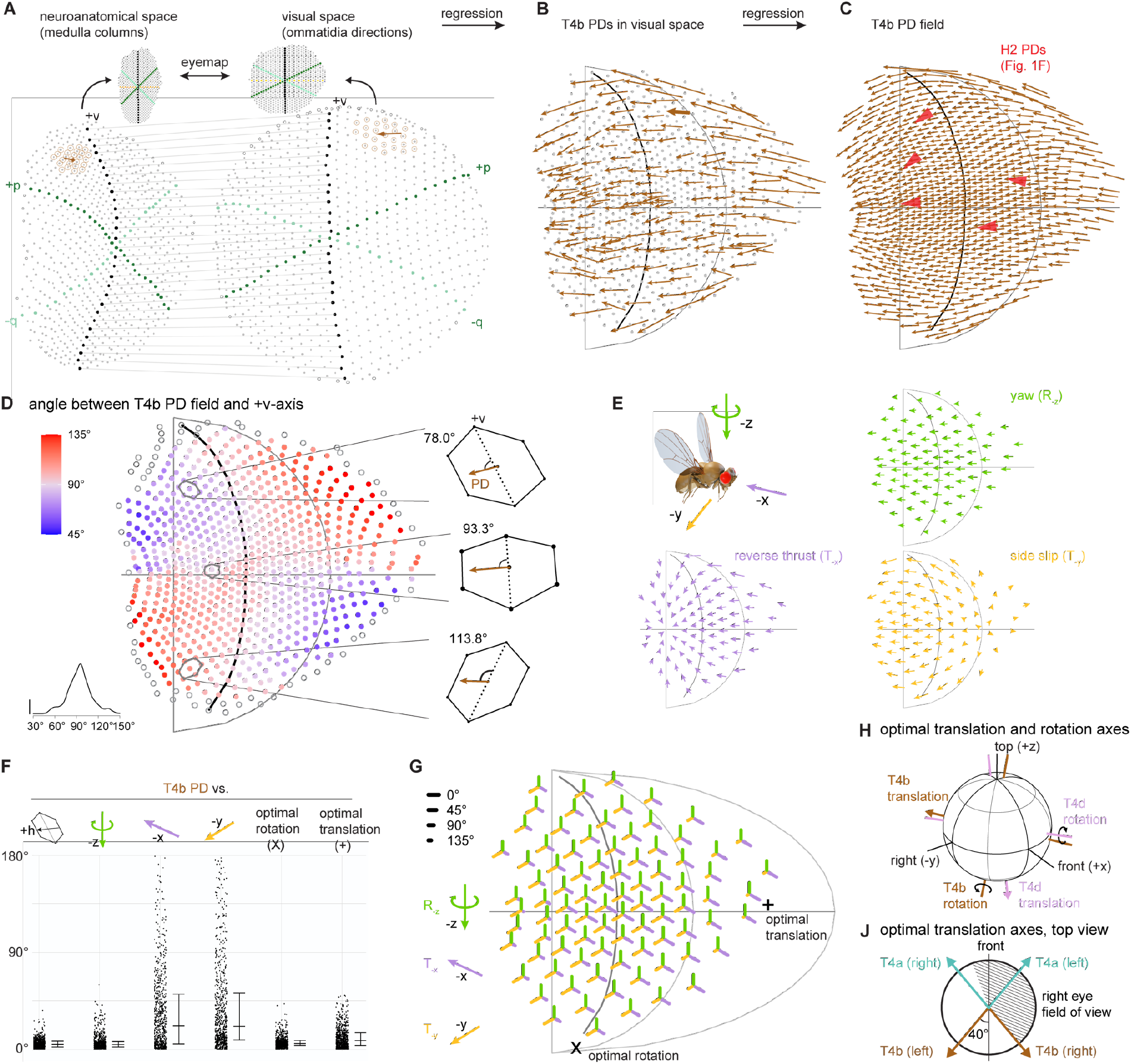
Mapping neuroanatomical space into visual space explains global organization of DS neuron preferred directions. **A**. 1-to-1 mapping between medulla columns, from the EM data set, and ommatidia directions, from the μCT data set, via mapping to regular grids. Unmatched columns on the periphery are denoted by empty circles. A T4b PD vector is transformed from the medulla to visual space. The neighboring columns used for kernel regression are highlighted with brown circles. **B**. 176 reconstructed T4b PDs mapped to visual space. The example vector in A is bolded. **C**. T4b PD field interpolated (see Methods) from (**B)**, assigning one T4b PD vector to each ommatidia direction (length re-scaled from (**B**) by 50%). For comparison, the PDs recorded from an H2 neuron are replotted from Fig. 1F as red arrowheads. **D**. Angular difference between T4b PD field in (**C**) and the +v-axis. This structure of the PD field matches features of ommatidial shearing (Fig. 3J). **E**. Ideal optic flow fields induced by yaw rotation, reverse thrust, and side slip. The number of ommatidia directions is down-sampled by a factor of 9 (keeping every third row or column). **F**. Angular differences between T4b PD field, +h-axis, three cardinal self-motion optic flow fields, and optimized self-motion flow fields (see Extended Data Fig. 6E). The horizontal bars represent 25%, 50% and 75% quantiles. **G**. Spatial distribution of the angular differences for the comparison with the 3 cardinal self-motions optic flow fields. The angular difference at each ommatidia direction (also down-sampled by a factor of 9) is represented with 3 line segments, with color matched to the cardinal self-motions and length given by the angular difference. The symbols “X” and “+” indicate the optimal rotation and translation axes, respectively. **H**. Optimal rotation and translation axes for T4b and T4d PD fields in the fly’s eye coordinates. **J**. Top view of the optimal translation axes for both T4a and T4b in both eyes represented along with the field of view at the equator.

The hexagonal arrangement is a dense spatial packing that maximizes the resolving power of the eye^10^. However, many unit hexagons are irregular, as illustrated by the inter-ommatidial angles (ΔΦ, Fig. 3G-H) and the shear angles (α, Fig. 3J). ΔΦ is smallest near the equator and the central meridian, and increases in size away from this region (Fig. 3F). When calculated separately for vertical (ΔΦ_v_) and horizontal (ΔΦ_h_) neighbors (Fig. 3H), we find that the vertical visual acuity is highest (lowest ΔΦ_v_) along the equator (a typical feature of flying insect eyes^29,30^, not previously reported for *Drosophila melanogaster*^28^), and the horizontal acuity is highest in the central part of the eye, though the effect of photoreceptor pooling (neural superposition^11^) on these acuity differences is unclear. These acuity differences are consistent with the aspect ratio changes of the unit hexagons across the eye (Extended Data Fig. 5C). Furthermore, the shear angle of the hexagons systematically changes, with the most regular hexagons (α ≈ 90°) found near the equator and the central meridian, and sheared hexagons with α < 90° in the fronto-dorsal and posterior-ventral quadrants, and α > 90° in the other quadrants (Fig. 3J). This μCT scan of the full fly head, provides the most detailed description of how the compound eye samples visual space. Our analysis reveals an irregular arrangement of ommatidia directions with spatially varying aspect ratios, inter-ommatidial angles, and shear angles, that shape the inputs to visual pathways. Could this non-uniform sampling explain the global structure of T4 PDs?

### Mapping neuroanatomical space into visual space explains the global organization of DS neuron preferred directions

We now have all the data required to map T4 PDs from their neuronal coordinates into the visual coordinates of the eye. We used the regular grids established for medulla columns (Fig. 2D) and ommatidia directions (Extended Data Fig. 5A) to construct a 1-to-1 mapping between them, matching from the center outward (Fig. 4A, Supplementary Video 1 and 2). We used kernel regression to transform the T4b PDs into eye coordinates (Fig. 4B). Finally, the T4b PDs were estimated for all ommatidia directions (Fig. 4C, Extended Data Fig. 6A,B; T4d in Extended Data Fig. 7A-D). Since T4a/b and T4c/d are mostly anti-parallel (Extended Data Fig. 6C,D), these estimates can be directly extended to all T4 subtypes. The stereotypical alignment of T4b PDs in the medulla (Fig. 2H) suggests that the PD field in eye coordinates should follow the ommatidia shearing (Fig. 3J), which is indeed the case (Fig. 4D). Remarkably, the T4b PDs are well-aligned to the spatially registered H2 responses (red arrowheads in Fig. 4C). It is noteworthy that both show a downward component in dorso-frontal PDs, which in our anatomical analysis, could only have originated from the non-uniform sampling of visual space. This global pattern has features of a translational optic flow field (Fig. 1A,F), that can be readily seen in the Mercator projection comparing the PD field with the eye coordinate parallel lines of constant elevation (Extended Data Fig. 6A). Since T4b provides substantial input to H2^19^, this agreement provides strong evidence for the mechanism hypothesized in Fig. 1G(ii) and validates our anatomy based PD prediction and mapping into visual coordinates. Taken together, these results demonstrate that the non-uniform sampling of the eye powerfully shapes the organization of PDs available for optic flow processing.

Is the T4b PD field (Fig. 4C) optimized for the optic flow induced by cardinal motion along body axes (Fig. 4E), as was found for mice DS neurons^3^? The distribution of angular differences between the T4b PD field, the eye’s +h-axis, and several optic flow fields, shows that the PDs are best aligned with the eye axis and yaw rotation (Fig. 4F). In contrast, there is a large spread in the differences between the PD field and reverse-thrust or side-slip optic flow, suggesting substantial regional variations. The spatial distribution shows that central eye PDs agree well with all three flow fields, while frontal PDs are more sensitive to side-slip, posterior PDs to reverse-thrust motion, and yaw rotation is well matched throughout (Fig. 4G, similar analysis for T4d in Extended Data Fig. 7E,F). Consequently, all neurons that integrate from most of a lobula plate layer, like H2 (Extended Data Fig. 1B), will inherit this eye-derived sensitivity. However, by selectively integrating from regional patches, lobula plate neurons can encode diverse features of optic flow, providing an expansive set of motion patterns for behavioral control.

Which body-movement-generated flow fields are most efficiently encoded by the T4b population? We searched and found the optimal rotation axis (by minimizing angular differences, see methods) quite close to the yaw axis, and a translation axis approximately half-way between reverse-thrust and side-slip, near the posterior boundary of the eye’s field of view (rightmost distributions in Fig. 4F, locations denoted with symbols in Fig. 4G, and complete error map in Extended Data Fig. 6E). Comparing to the optimal axes for T4d PDs (Extended Data Fig. 7F) we find a remarkable agreement between these principal axes—with the body yaw axis matching T4ds’ optimal translation axis, and T4bs’ non-canonical optimal translation axis matching T4ds’ optimal rotation axis (Fig. 4H). Since optic flow is a direct consequence of movement, it is likely that these principal axes of maximal motion sensitivity are fundamental for controlling body movements. Intriguingly, the optimal translation axes for the left and right T4a populations are near the eye’s equator and approximately ±40º from the midline (Fig. 4J), precisely where we predict T4s exhibit their highest acuity (Extended Data Fig. 6B). We note a striking resemblance between the optimal translation axes for T4a/b (Fig. 4J) and the tuning of optic flow sensitive inputs to the central complex^31^, from which the transformations between body-centered and world-centered coordinates are built^32^. This unexpected correspondence of maximal motion sensitivities exposes a deep link between the structure of the eye and the coordinate systems organizing goal-directed navigation in the central brain.

## Discussion

Our analysis of the eye-derived pattern of spatial integration by the T4 directionally selective neurons, unifies two rich perspectives on fly motion vision—the recent discoveries about the local circuit mechanism for computing directional selectivity in *Drosophila*^14,15,17,33^ together with groundbreaking work in larger flies on the sensing of global optic flow patterns by wide-field lobula plate neurons^9,20,22,23^. Consequently, our study reconciles multiple previous findings. Behavioral studies using precise, localized visual stimulation described maximal responses to motion directions aligned with rows on the eye^34,35^, and work in larger flies noted that the local PDs of several Lobula Plate Tangential Cells^36^ reflected the orientation of the hexagonal grid in frontal eye regions^37^. A recent study of the looming-sensitive LPLC2 cells in *Drosophila* found this neuron was most sensitive to non-cardinal, diagonal movement directions in the dorso-frontal eye regions, and found that LPi interneurons had shifting PDs across the field of view^38^. Finally, a recent study found T4/T5 axonal responses that resembled a translation-like pattern with smoothly varying PDs across lobula plate layers^24^. Our study provides a mechanistic explanation for these observations—the missing link between the arrangement of eye facets and local PDs measured in the lobula plate, is the universal sampling rule we discovered for T4 neurons (Fig. 2) that adheres closely to the coordinate system of the eye. Based on our anatomical analysis of the dendritic orientation of T4 neurons, identified by their targeted lobula plate layer, we find no evidence for additional subtypes of T4 neurons. However, recent transcriptomic studies^26^ provide evidence for additional subtype diversity, and functional studies^24^ suggest that local PDs may be regionally modified through as-yet-undescribed connectivity differences, an important question for future EM studies. Finally, our analysis of global optic flow patterns (Fig. 4, Extended Data Figs. 6,7) provides a simple explanation for the observation that HS and VS cell responses simultaneously represent information about both self-rotations and self-translations^39,40^, since the eye-derived PDs are most sensitive to different cardinal self-motions in different eye regions.

The computation of directional selectivity depends on asymmetric wiring in the dendrites of T4 and T5 neurons. Each subtype connects to different cell types at different locations along the dendrite^5^, but the developmental mechanisms establishing this wiring asymmetry are not known^41^. Our discovery of a universal sampling of medulla columns by T4 dendrites suggests that the core developmental mechanisms may be identical across the medulla (and lobula for T5s) and all subtypes, acting together with a process that established the subtype-specific dendritic orientation. Supporting this proposal is evidence from recent RNA-Seq studies showing that all 8 T4 and T5 subtypes are transcriptionally very similar, including during development^41–43^. The realization that in the appropriate reference frame, all T4 neurons are quite similar greatly simplifies the scope for a required explanatory mechanism.

Arthropods with compound eyes, which comprise the majority of described animal species, show a remarkable diversity of anatomical specializations, reflecting their diverse visual ecology^30^. Since many features of optic lobe anatomy, including key cell types involved in motion vision, are conserved across flies^44^ and comparable neurons and brain regions are found across arthropods^45^, the insights uncovered in *Drosophila* may apply broadly. Extrapolating from our work, we wonder whether detailed eye maps would make strong predictions about the motion directions sensed by the animal, and thus its behavior and natural history. This correspondence between the structure of the sensory system and an animal’s behavioral repertoire^46^ is an important demonstration that neural computations cannot be considered in isolation, as evolution jointly sculpts the function of the nervous system and the structure of the body.

## Methods

### Anatomical data

#### EM reconstruction

All reconstructions in this manuscript are from a serial section transmission EM volume of a *Drosophila melanogaster* full adult fly brain (FAFB)^6^. We manually reconstructed neuron skeletons in the CATMAID environment^47^ (in which 27 labs were collaboratively building connectomes for specific circuits, mostly outside of the optic lobe) following established practices^48^. We also used two recent auto-segmentations of the same data set, FAFB-FFN1^49^ and FlyWire^50^ to quickly examine many auto-segmented fragments for neurons of interest. Once a fragment of interest was found, it was imported to CATMAID, followed by manual tracing and identity confirmation.

For the data reported here, we identified and reconstructed a total of 780 Mi1, 38 T4a, 176 T4b, 22 T4c, 114 T4d, 63 TmY5a and 1 H2 cells. All the columnar neurons could be reliably matched to well-established morphology from golgi-stained neurons^13^. This reconstruction is based on approximately 1.35 million manually placed nodes. (1) Mi1: we traced the main branches of the M5 and M9/10 arbors such that the centers-of-mass of the arbors formed a visually identifiable grid. We used the auto-segmentation to accelerate the search for Mi1 cells wherever there appeared to be a missing point in the grid. After an extensive process, we believe that we have found all Mi1 cells in the right optic lobe (Fig. 2C,D). One Mi1 near the neuropil boundary was omitted in later analysis because its center-of-mass was clearly “off the grid” established by neighboring Mi1 cells despite a complete arbor morphology. (2) T4: we traced their axon terminals in the lobula plate for subtype identification (each subtypes innervating a specific depth in the lobula plate^19^) and manually reconstructed their complete dendritic morphology to determine their anatomical preferred direction. To sample T4 morphology across the whole eye with a reasonable amount of time and effort, we focused on the T4b (Fig. 2E) and T4d (Extended Data Fig. 2G) subtypes with sufficient density to allow us to interpolate the PDs at each column position. In addition, we chose 4 locations on the eye: medial (M), anterior dorsal (AD), anterior ventral (AV) and lateral ventral (LV), where we reconstructed 3 ∼ 4 sets of T4 cells and confirmed that the PDs were mostly anti-parallel between T4a and T4b, as well as between T4c and T4d (Extended Data Fig. 6C,D). (3) TmY5a: we searched for cells along the equator and central meridian of the medulla and traced out their main branches to be able to extend (with further interpolation) the columnar structure of the medulla to the lobula (Extended Data Fig. 2F). (4) H2: The neuron was found during a survey (unpublished) of the LPTCs in the right side of the FAFB brain and was completely reconstructed, including all fine branches in the lobula plate (Fig. 1D, Extended Data Fig. 1B).

In addition, we identified several lamina monopolar cells and photoreceptor cells. (5) Lamina cells, mainly L1, L2, L3 and outer photoreceptor cells (R1-6) were reconstructed, often making some use of auto-segmented data, to allow for their identification. This helped us locate the equatorial columns in medulla that have different numbers of photoreceptor inputs in the corresponding lamina cartridge (Fig. 2C, Extended Data Fig. 2B-D). (6) Inner photoreceptor cells R7/8: we searched for R7/8 cells throughout the eye, initially as part of a focused study on the targets of these photoreceptors^51^. We extended these reconstructions to complete the medulla map in Fig. 2. We searched for R7/8 corresponding to each Mi1 cells near the boundary of the medulla. Mi1 cells in columns lacking inner photoreceptors were identified and excluded from further analysis (Fig. 2C). Furthermore, we reconstructed several cells near the central meridian and used their axons’ shape to identify the location of the chiasm (Extended Data Fig. 2E).

### Generation and imaging of split-GAL4 driver lines

We used split-GAL4 driver lines SS00809^15^ and SS01010 to drive reporter expression in Mi1 and H2 neurons, respectively. Driver lines and representative images of their expression patterns are available at https://splitgal4.janelia.org/. SS01010 (newly reported here; 32A11-p65ADZp in attP40; 81E05-ZpGdbd in attP2) was identified and constructed using previously described methods and hemidriver lines^52,53^. We used MCFO^54^ for multicolor stochastic labeling. Sample preparation and imaging, performed by the Janelia FlyLight Project Team, were as in previous studies^53,54^. Detailed protocols are available online (https://www.janelia.org/project-team/flylight/protocols under “IHC - MCFO”). Images were acquired on Zeiss LSM 710 or 780 confocal microscopes with 63x 1.4 NA objectives at 0.19×0.19×0.38 μm^3^ voxel size. The reoriented views shown in Extended Data Fig. 1B and Extended Data Fig. 2A,B were displayed using VVDviewer (https://github.com/JaneliaSciComp/VVDViewer). This involved manual editing to exclude labeling outside of approximately medulla layers M9/10 (Extended Data Fig. 2A,B) or to only show a single H2 neuron (Extended Data Fig. 1B).

### Confocal imaging of a whole fly eye

#### Sample preparation

Flies were anesthetized with CO2 and briefly washed with 70% ethanol. Heads were isolated, proboscis removed under 2% paraformaldehyde/PBS/0.1% triton X-100 (PBS-T) and fixed in this solution overnight at 4°C. After washing with PBS-T, the heads were bisected along the midline with fine scissors and incubated in PBS with 1% triton X-100, 3% normal goat serum, 0.5% DMSO and Escin (0.05 mg/ml, Sigma-Aldrich, #E1378) containing chicken anti-GFP (1:500; Abcam #ab 13970), mouse anti-nc82 (1:50; Developmental Studies Hybridoma Bank) and rabbit anti-RFP (1:1000; TaKaRa Bio USA, #632496) at room temperature with agitation for 2 days. After a series of three ∼1h-long washes in PBS-T the sections were incubated for another 24h in the above buffer containing secondary antibodies: Alexa Fluor 488 goat anti-chicken (1:1000; Thermo Fisher #A11039), Alexa Fluor 633 goat anti-mouse (1:1000; Thermo Fisher #A21050) and Alexa Fluor 568 goat anti-rabbit (1:1000; Thermo Fisher #A11011). The samples were then washed in PBS/1% triton (4 × 1 h) and post-fixed for 4 h in PBS-T/2% paraformaldehyde. To avoid artefacts caused by osmotic shrinkage of soft tissue, samples were gradually dehydrated in glycerol (2-80%) and then ethanol (20 to 100%)^55^ and mounted in methyl salicylate (Sigma-Aldrich #M6752) for imaging.

### Imaging and rendering

Serial optical sections were obtained at 1 μm intervals on a Zeiss 710 confocal microscopes with a LD-LCI 25x/0.8 NA objective using 488, 560 and 630 lasers, respectively. The image in Extended Data Fig. 4A is a reoriented substack projections, processed in Imaris v9.5 (Oxford Instruments).

### μCT imaging of whole fly heads

μCT is an x-ray imaging technique that is similar to medical CT scanners, but with much higher resolution more suitable for smaller samples^56^. A 3D data volume set is reconstructed from a series of 2D x-ray images of the physical sample at different angles. The advantage of this method for determining the ommatidia directions (Fig. 3) is that internal details of the eye, such as individual rhabdoms, distinguishable ‘tips’ of the photoreceptors at the boundary between the pseudocone and the neural retina^57^, and the chirality of the outer photoreceptors, can be resolved across the entire intact fly head with isotropic resolution, which is an important requirement for preserving the geometry of the eye.

### Sample preparation

Based on previously published fixation and staining protocols for a variety of biological models^58^, we undertook extensive testing of fixatives and stains in addition to mounting/ immobilizing steps for μCT scanning. Fixatives tested were Bouins fluid, alcoholic Bouins, 70% ethanol. We tested staining with Phospho-tungstic acid in water and in ethanol, Phosphomolybdic acid in water and in ethanol, Lugol’s Iodine solution, 1% Iodine metal dissolved in 100% ethanol. Various combinations of fixatives and stains were tried along with variations in times for each. Fixing and staining samples in aqueous solutions and then scanning these samples in an aqueous environment, despite efforts to immobilize the head, yielded blurry images and poor resolution. Drying the samples using hexamethyldisilazane (HMDS) did not yield images with the resolution achievable with critical point dried samples^58^. The protocol that worked best involved fixing and staining in ethanol-based solutions followed by critical point drying giving good contrast, high resolution images with excellent reproducibility.

6 − 7 day old female D. melanogaster flies were anesthetized with CO_2_ and immersed in 70% ethanol. The heads were dissected out from the body at the thorax region just below the neck to allow for a larger surface area of fixative absorption. Samples were fixed in 70% ethanol at room temperature for 3 days in a 1.5 ml Eppendorf tube with rotation. The ethanol was then replaced with staining solution of 0.5% Phospho-tungstic acid in 70% ethanol. Samples remained in the staining solution for 5-6 days at room temperature with rotation. The heads were rinsed 3×10 min with 70% ethanol at room temperature to remove the staining solution followed by dehydration in 90% and 100% ethanol for 30 min each. The samples were then critical point dried (Tousimis supercritical autosamdri 931.GL). The stasis mode protocol was used with 3 stasis cycles lasting 90 minutes each. Next, the fly head was mounted on the tip of a toothpick using a tiny drop of superglue on the remaining thorax region. We confirmed that no glue got on to the head region.

### Imaging and reconstruction

The samples were scanned with Zeiss Xradia Versa XRM 500 microCT scanner. The scanning was carried out at a voltage of 40kV, current of 72μA (power 2.9W) at 20x magnification with 10 sec exposures and a total of 1601 projections. Images had a pixel size of 1.0343 μm with camera binning at 2 and reconstruction binning at 1. The Zeiss XRM reconstruction software was used to generate TIFF stacks of the tomographs. Image segmentation and annotation (lenses and photoreceptor tips) were done in Imaris v9.5 (Oxford Instruments).

### Whole cell recordings of labeled H2 neurons

#### Electrophysiology

All the flies used in electrophysiological recordings were from a single genotype: pJFRC28-10XUAS-IVS-GFP-p10^59^ in attP2 crossed to the H2 driver line SS01010 (see section ‘Generation and imaging of split-GAL4 driver lines’). Flies were reared at a 16 light:8 dark light cycle at 24°C. To perform the recordings, 2-3 days old female *Drosophila melanogaster* were anesthetized on ice and glued to a custom-built PEEK platform, with their heads tilted down, using a UV cured glues (Loctite 3972) and a high-power UV curing LED system (Thorlabs CS2010). To reduce brain motion, the two front legs were removed, the folded proboscis was glued in its socket, and muscle 16^60^ was removed from between the antennae. The cuticle was removed from the posterior part of the head capsule using a hypodermic needle (BD Precisionglide 26G X ½’’) and fine forceps. Manual peeling of the perineural sheath using the forceps seemed to damage the stability of the recordings and, therefore, the sheath was removed using collagenase (following prior method^61^). To prevent contamination, the pipette holder was replaced after collagenase application.

The brain was continuously perfused with an extracellular saline containing (in mM): 103 NaCl, 3 KCl, 1.5 CaCl_2_ 2H_2_O, 4 MgCl_2_ 6H_2_O, 1 NaH_2_PO_4_ H2O, 26 NaHCO_3_, 5 N-Tris (hydroxymethyl) methyl-2-aminoethane-sulfonic acid, 10 Glucose, and 10 Trehalose with Osmolarity adjusted to 275mOsm and bubbled with carbogen throughout the experiment. Patch clamp electrodes were pulled (Sutter P97), pressure polished (ALA CPM2) and filled with an intracellular saline containing (in mM): 140 Kasp, 10 HEPES, 1 EGTA, 1 KCl, 0.1 CaCl2, 4 MgATP, 0.5 NaGTP, and 5 Glutathione^62^. 250μM Alexa 594 Hydrazide was added to the intracellular saline prior to each experiment, to reach a final osmolarity of 265mOsm, with a pH of 7.3.

Recordings were obtained using a Sutter SOM microscope with a 60X water-immersion objective (60X Nikon CFI APO NIR Objective, 1.0 NA, 2.8 mm WD). Contrast was generated using oblique illumination from an 850nm LED connected to a light guide positioned behind the fly’s head. Images were acquired using Micro-Manager^63^, to allow for automatic contrast adjustment. All recordings were obtained from the left side of the brain. To block visual input from the contralateral side, the right eye was painted with miniature paint (MSP Bones grey primer followed by dragon black). Current clamp recordings were sampled at 20KHz and low-pass filtered at 10KHz using Axon multiClamp 700B amplifier (National Instrument PCIe-7842R LX50 Multifunction RIO board) using custom LabView (2013 v.13.0.1f2; National Instruments) and MATLAB (Mathworks, Inc.) software.

#### Visual stimuli

The display was a G4 LED arena^64^ with a manual instead of a motorized rotation axis. The arena covered slightly more than one half of a cylinder (216° in azimuth and ∼72° in elevation) of the fly’s visual field, with the diameter of each pixel subtending at most 2.25° on the fly eye. Visual stimuli were generated using custom written MATLAB code. Presented stimuli were:

1. Moving grating: square wave grating with a constant spatial frequency (7 pixels ON / 7 pixels OFF) were presented in a ∼22° circular window over an intermediate intensity background. Gratings moved at 1.78Hz (40ms steps) and were presented at 16 different orientations. Grating were presented for 3 full cycles (1.68 sec) with 3 repetitions for each stimulus condition. The H2 responses to these trials are the basis for Fig. 1E,F and Extended Data Fig. 1C,D.
2. Moving bars: This stimulus was used to detect the extent of the field of view of the inputs to the H2 cell. Moving bars were presented in both contrasts (ON and OFF) and both preferred and non-preferred directions for H2 cells (back to front and front to back respectively). Bars were 7 pixels wide and 21 pixels high and moved with 40ms steps (∼56°/sec). Bars moved within a 21 pixels window that was centered around different positions in the arena. The H2 responses to these trials are not shown.

The local preferred direction for the H2 cells were determined using the responses to the 16 directions of the moving gratings. Spikes were extracted from the recorded data and summed per trial, then averaged across repeated presentations of each stimulus. The polar plots (Fig. 1E) represent these averages (relative to baseline firing rate), while the vector sum over all 16 directions is represented by the black marker in Fig. 1E and the red arrows in Fig. 1F. The subthreshold responses of H2 were also be used to determine the local preferred direction of the neuron, showing excellent agreement to the directions based on the neuron’s spiking responses (Extended Data Fig. 1C,D).

#### Determining head orientation

First, the camera (Point grey Flea 3 with an 8X CompactTL™ telecentric lens with in-line illumination, Edmund Optics) was aligned to a platform holder using a custom-made target. This allowed us to adjust the camera and platform holder such that when the holder is centered in the camera’s view, both yaw and roll angles are zero. Next, after the fly was glued to the platform, but before the dissection, images were taken from the front to check for yaw and roll angle of head orientation. If the deviation of the head away from a ‘straight ahead’ orientation was greater than 2°, then that fly was discarded. Finally, to measure pitch angle, the holder was rotated ±90°, and images of the fly’s eye were taken on both sides. Head orientation was then measured as previously described^65^.

### Data analysis

#### Mapping medulla columns

We based our map of medulla columns on the principal, columnar cell type Mi1 that is found as one per column. Mi1 neurons nearly resemble columns with processes that do not spread far from the main ‘trunk’ of the neuron. They have a stereotypical arborization pattern in medulla layer M1, M5, and M9/10. For each Mi1 cell, we calculated the centers-of-mass of its arbors in both M5 and M10 and used them as column markers (Fig. 2B,C). The medulla columns do not form a perfectly regular grid— the column arrangement is squeezed along the anterior-posterior direction and the dorsal and ventral portions shift towards anterior. Nevertheless, we were able to map all column positions onto a regular grid via visual inspection (Fig. 2D). This was much clearer based on the positions of the M5 column markers, which are more regular, and were used as the basis for our grid assignment. For occasional ambiguous cases, we compared the whole cells (across layers) in a neighborhood to confirm our assignment. We then propagated the grid assignment to M10 column markers and use them throughout the paper since T4 cells received inputs in layer M10.

Establishing a global reference that could be used to compare the medulla map (Fig. 2C) to the eye map (Fig. 3F) was essential, and so we endeavored to find the “equator” of the eye in both the EM and uCT data sets. Lamina cartridges in the equatorial region receive more outer photoreceptor inputs (7-8 compared to the normal 6)^11,66^. We traced hundreds of lamina monopolar cells (L1s or L3s), with at least one input to each of ∼100 Mi1 cells near the equator region and counted the number of photoreceptor cells in each corresponding lamina cartridge (Extended Data Fig. 2B-D). This allowed us to locate the equatorial region of the medulla (Fig. 2C). The equator in uCT is identified by the chirality of the outer photoreceptors (Fig. 3D). We further identified the “central meridian, +v” row, which is roughly the vertical midline. Note that there is some ambiguity in defining +h as the equator in Fig. 2D since there are 4 rows of ommatidia with 8 photoreceptors (points in tan). We opted for one of the middle two rows that intersects with +v. We also identified the chiasm region based on the twisting of R7/8 photoreceptor cells (Extended Data Fig. 2E), which very nearly aligned with the central meridian.

### T4 preferred direction (PD)

Strahler number (SN) was first developed in hydrology to define the hierarchy of tributaries of a river^25^, and has since been adapted to analyze the branching pattern of a tree graph (Fig. 2F). A dendrite of a neuron can be considered as a tree graph. The smallest branches (leaves of a tree) are assigned with SN = 1. When two branches of SN = a and SN = b merge into a larger branch, the latter is assigned with SN = max(a, b) if a ≠ b, or with SN = a + 1 if a = b.

We used SN = {2, 3} branches to define the PD because they are the most consistently directional (Extended Data Fig. 3A). SN = 1 branches have a relatively flat angular distribution. Most T4 cells we reconstructed have few SN = 4 branches (which are also directional, but too few to be relied upon), and rarely SN = 5 branches. Each branch is represented by a 3D vector. Vector sums are calculated for all SN = {2, 3} branches which define the directions of the PD vectors (Fig. 2F). We also assigned an amplitude to the PD, in addition to its direction. To generate a mass distribution for each T4 dendrite, we re-sampled the neuron’s skeleton such that the nodes are positioned roughly equidistantly (not so after manual tracing). Then all dendrite nodes were projected onto the PD axis. We define the length of the PD vector using a robust estimator, the distance between the 1^st^ and 99^th^ percentiles of this distribution. The orthogonal direction (OD) is a segment orthogonal to the PD vector, with its length similarly defined as PD and without a direction (Fig. 2G).

### Mapping T4 PDs into the regular grid in medulla (Fig. 2H) and the eye coordinates (Fig. 4B, C) using kernel regression

Given a point set P in space A and a second point set Q in space B, and a 1-to-1 mapping between P and Q, one can map a new point x in A to a location y in B based on the relationships of x with respect to P. In our case, because the mapping from A to B is not a rigid coordinate transformation, we applied kernel regression to map the new points. Intuitively, this method takes into account the spatial relationships between x and all the points in P, but gives more weights to the points that are closer to x. The weight is assigned using a Gaussian kernel, hence the name kernel regression.

We used this method to map PDs from local medulla space to a regular grid in Fig. 2H, and to map PDs from medulla space to visual space in Fig. 4B. For mapping to a regular grid, we defined a 2D reference grid with 19 points, which represented the home column (+1) and the 2^nd^ (+6) and 3^rd^ (+12) closest neighboring columns in a hexagonal grid. For a given T4 neuron, we searched for the same set of its neighboring medulla columns. We flattened these columns and the T4’s PD locally by projecting them onto a 2D plane that is given by principal component analysis, that is, the plane is perpendicular to the 3^rd^ principal axis. Finally, we used kernel regression to map the PD from the locally flattened 2D medulla space to the 2D reference grid. The difference in mapping to the visual space (Fig. 4B, Extended Data Fig. 7A) is that the regression is from the locally flattened 2D medulla space to a unit sphere in 3D (the space of ommatidia directions).

Kernel regression can also be used as an interpolation method, which is equivalent to mapping from a space to its scalar or vector field, that is, to assign a value to a new location based on existing values in a neighborhood. This is how we calculated the PD fields in Fig. 4C and Extended Data Fig. 7B.

In practice, we used the np package in R^67^, in particular, the “npregbw” function which determines the width of the Gaussian kernel. Most parameters of the npregbw function are set to default except: (1) We used the local-linear estimator, “regtype = ‘ll’”, which performs better near boundaries. (2) We used fixed bandwidth, “bwtype= ‘fixed’”, for interpolation and adaptive nearest neighbor method, “bwtype= ‘adaptive_nn’”, for mapping between 2 different spaces (eg. from medulla to ommatidia). Further details can be found in our Github repository and the np package manual for more details.

### Ommatidia directions

We analyzed the μCT volumes in Imaris v9.5 (Oxford Instruments). We manually segmented the lenses and photoreceptor tips, so they could be analyzed separately. We then used the “spot detection” (based on contrast) algorithm in Imaris to locate the centers of individual lenses and photoreceptor tips, and quality-controlled by visual inspection and manual editing. The lens positions are extremely regular and can be readily mapped onto a regular hexagonal grid (Extended Data Fig. 5A, directly comparable to the medulla grid, Fig. 2D). With our optimized μCT data, it is also straightforward to match all individual lenses to all individual photoreceptor “tips” in a one-to-one manner, and consequently to compute the ommatidia viewing directions. These directional vectors can be represented as points on a unit sphere (Fig. 3E). We then performed a local weighted smoothing for points with at least 5 neighbors: the position of the point itself weights 50% while its neighbors’ average position accounts for the remining 50%. This gentle smoothing only impacts the positions in the bulk of the eye while leaving the boundary points alone.

Assuming left-right symmetry, we used the lens positions from both eyes to define the frontal midline (sagittal plane) of the visual field. Together with the equator, identified by the inversion in the chirality of the outer photoreceptors (Fig. 3C,D), we could then define an eye coordinate system for the fly’s visual space – represented for one eye in Fig. 3E and F. Note that the z = 0 plane (“z” is “up” in Fig. 3E) in the coordinate system is defined by lens positions, hence the “equator” ommatidia directions do not necessarily lie in this plane (more easily seen in Fig. 3F). In addition, we defined the “central meridian” line of points (“+v” in Fig. 2E,F and Extended Data Fig. 5A) that divides the whole grid into roughly equal halves. Because this definition is based on the grid structure, this “central meridian” does not lie on a geographic meridian line in the eye coordinates.

### Eyemap: 1-to-1 mapping between medulla columns and ommatidia directions

With both medulla columns and ommatidia directions mapped to regular grid (Fig. 2D and Extended Data Fig. 5A), and equators and central meridians defined, it’s straightforward to match these two points sets, starting from the center outwards. Because the medulla columns are from a fly imaged with EM and ommatidia directions from a different fly imaged with μCT, we don’t expect these two point sets to match exactly, but we endeavored to use flies with a very similar total number of ommatidia (and of the same genotype). By matching the points from center outwards and relying on anatomical features such as the equator, we expect to minimize the column receptive field discrepancies, especially in the interior of eye. The matching at the boundary is somewhat complicated by the existence of medulla columns with no inner photoreceptor (R7/8) inputs (Fig. 2C)^68^. In the eyemap in Fig. 4A, we denoted unmatched points with empty circles, all of which solely lie on the boundaries (which is why the ommatidia directions in Fig. 3F contain additional points). Any consideration of these medulla columns and ommatidia directions should be done with caution. In addition, we also noted those boundary points that do not have enough neighbors for computing the inter-ommatidial angles, the shear angles, or the aspect ratios in Fig. 3H, J, and Extended Data Fig. 5C,D. Importantly, our primary discoveries about the universal sampling of medulla columns (Fig. 2), and the strong relationship between T4 PDs and shear angle of ommatidia hexagons (comparing Fig. 3J to Fig. 4D) are well supported by the anatomy of the bulk of the eye and do not depend on perfect matching across data sets or the particular fly used to construct the eye map (see Extended Data Fig. 4E,F).

### Grid convention: regular vs irregular, hexagonal vs square

Facet lenses of the fly’s eye are arranged in an almost regular hexagonal grid. However, the medulla columns are squeezed along the anterior-posterior direction and more closely resemble a square grid tilted at 45º (Extended Data Fig. 5E). This difference can also be seen by comparing the aspect ratios (Extended Data Fig. 5C,D). To preserve these anatomical features, we mapped the medulla columns and T4 PDs onto a regular square grid (tilted by 45º, e.g. Fig. 2D, H), and the ommatidia directions onto a regular hexagonal grid (Extended Data Fig. 5A).

### Mercator and Mollweide projections

For presenting spherical data, the Mercator projection is more common, but we prefer the Mollweide projection since it produces only small distortion near the poles, and because of these smaller distortions it provides a more intuitive representation of spatial coverage. On the other hand, the Mercator projection preserves all the angular relationships and is more convenient for reading out angular distributions, which is why we use it for presenting the H2 data (Fig. 1F, Extended Data Fig. 1D). Otherwise, we used the Mollweide projections in the main figures and provide the Mercator version for some plots (Extended Data Figs. 5F-J, 6A, 7C).

### Ideal optic flow fields

Following the classic framework for the geometry of optic flow^69^, we calculate the optic flow field for a spherical sampling of visual space under the assumption that all objects are at an equal distance from the animal (only relevant for translational movements). With ommatidia directions represented by unit vectors in 3D, the optic flow field induced by translation is computed as the component of the inverse of the translation vector (since motion and optic flow are “opposite”) perpendicular to the ommatidia directions (also known as a “vector rejection”). The flow field induced by rotation is computed as the cross product between the ommatidia directions and the rotation vector. Since the motion perceived by the animal would be the opposite of the induced motion, the flow field is the reverse of the ones described above (Fig. 4E). The angles between T4 PDs and ideal optic flows at each ommatidia direction are computed for subsequent comparisons between various optic flow fields (Fig. 4F,G, Extended Data Fig. 7E,F).

To determine the optimal axis of movement (minimal average errors) for a given PD field, we performed a grid search. We defined 10356 axes on the unit sphere (roughly 1º sampling) and generated optic flow fields induced by translations and rotations along these axes. We compared all these optic flows fields and the PD fields for T4b and T4d to determine the axes with minimal average angular differences (Extended Data Fig. 6E). These are the optimal axes in Fig. 4F-J, Extended Data Fig. 6E, Extended Data Fig. 7F.

### Data analysis and plotting conventions

All histograms are smoothed as a kernel density estimation. To set the scale of each histogram plot, we show a scale bar on the left-hand side that spans from zero at the bottom to the height of a uniform distribution. All 2D projections (Mollweide or Mercator) are such that the right half (azimuth > 0) represent the right-side visual field of the fly (looking from inside out). Top half (elevation > 0) represents the dorsal visual field.

## Supporting information

Supplementary Files, T4 galleries

## Data and Code Availability

Data analyses were carried out with custom code in R using open-source packages, mainly “natverse”, “tidyverse”, and “np”. Animations were created using Blender and the Python package “navis”. We will make the data and code used to produce the major results of this study available at the time of publication. We will provide the most updated materials to correspond to the final version of the manuscript. EM reconstructed neurons will be uploaded to a public CATMAID server: https://catmaid.virtualflybrain.org. Flylight images will be available on the FlyLight website: https://splitgal4.janelia.org/cgi-bin/splitgal4.cgi. The μCT and some of the confocal stacks will be uploaded to FigShare. Analysis and plotting code will be available on github: https://github.com/reiserlab.

## Acknowledgements

The authors thank Ruchi Parekh for managing the Connectome Annotation Team, Gerry Rubin for support of Aljosha Nern and for sharing the H2 split-GAL4 line, Tess Oram and Gwyneth Card for sharing collagenase, the Janelia Fly Core for fly care, Janelia FlyLight Project Team for help with preparation and imaging of light microscopy samples, and members of the Reiser lab for helpful feedback. We thank the FAFB tracing community for supportive and open sharing of methods and data, especially Greg Jefferis, Marta Costa, Philipp Schlegel, and the “FAFB optic lobe working group”, especially the groups of Mathias Wernet, Eugenia Chiappe, Marion Silies, and Rudy Behnia for collaborations and feedback. Development and administration of the FAFB tracing environment and analysis tools were funded in part by National Institutes of Health BRAIN Initiative grant 1RF1MH120679-01 to Davi Bock and Greg Jefferis, with software development effort and administrative support provided by Tom Kazimiers (Kazmos GmbH) and Eric Perlman (Yikes LLC). We thank Peter Li for sharing his automatic segmentation (https://doi.org/10.1101/605634). We thank the Princeton FlyWire team and members of the Mala Murthy and Sebastian Seung labs for development and maintenance of FlyWire (supported by BRAIN Initiative grant MH117815 to Mala Murthy and Sebastian Seung). This work made use of VVDviewer, based on software funded by the NIH: Fluorender: An Imaging Tool for Visualization and Analysis of Confocal Data as Applied to Zebrafish Research, R01-GM098151-01. This work is funded by the Howard Hughes Medical Institute through its support of the Janelia Research Campus. This article is subject to HHMI’s Open Access to Publications policy. HHMI lab heads have previously granted a nonexclusive CC BY 4.0 license to the public and a sublicensable license to HHMI in their research articles. Pursuant to those licenses, the author-accepted manuscript of this article can be made freely available under a CC BY 4.0 license immediately upon publication.

---

**Extended Data Figure 1:**
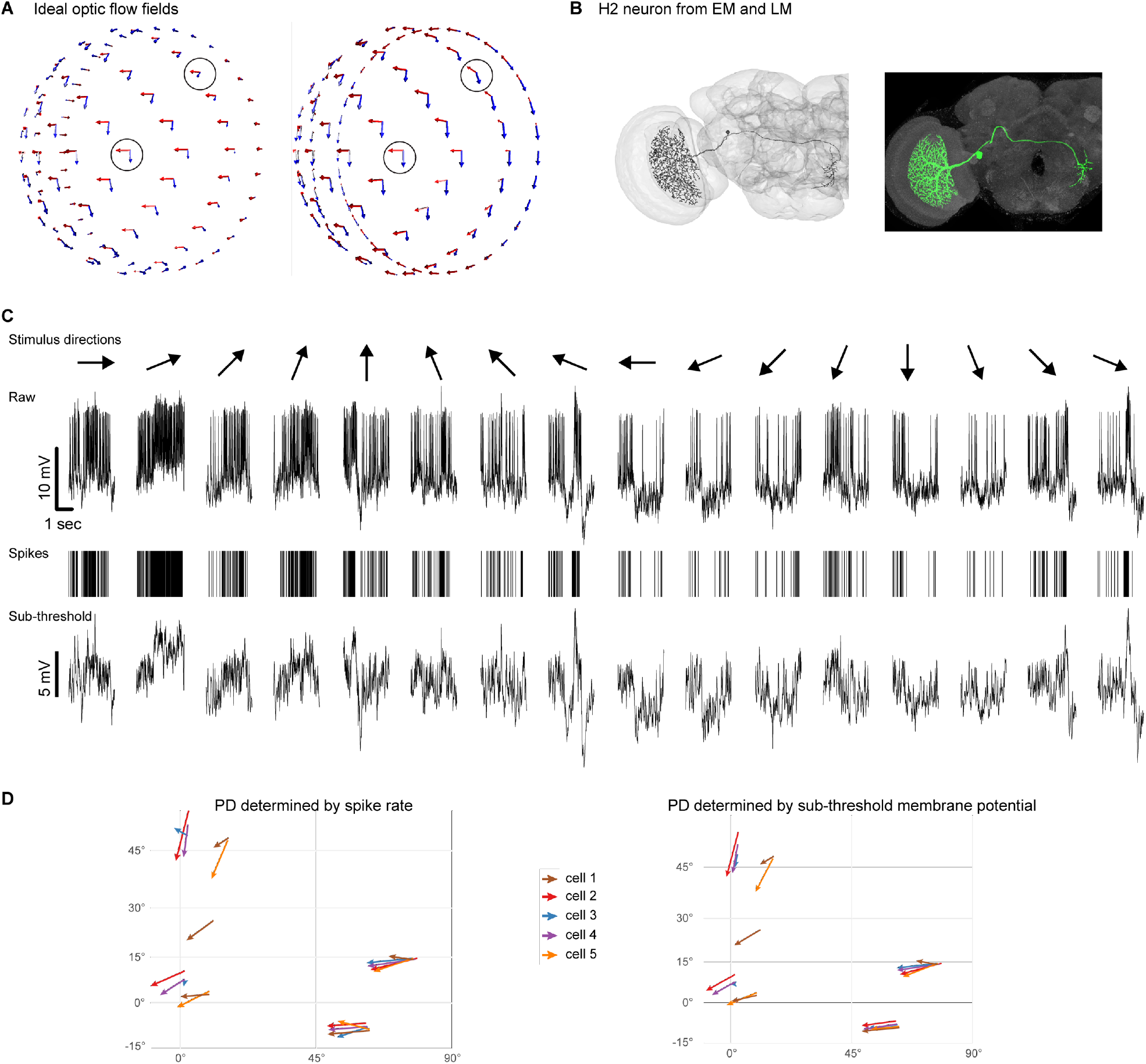
Global optic flow patterns and H2 neuron results, related to Fig. 1. **A**. Left: two optic flow fields on a hemisphere (representing the right eye of a fly) induced by a yaw rotation (red) and a pitch rotation (blue). Right: two optic flow fields induced by a thrust (red) and a lift (blue). Note in both cases, the local red and blue optic flow components are orthogonal near the center of the eye while forming skewed (yet different) angles near the boundary. **B**. EM reconstruction and light microscopy image of an H2 neuron (see Methods). **C**. An H2 response to moving gratings stimuli along 16 directions at location ➁ in Fig. 1F. Raw, filtered spike, sub-threshold time series are shown. **D**. H2 angular tuning at various visual locations, from recordings of 5 flies. The vector locations for each animal are based on our procedure for aligning each fly’s head.

**Extended Data Figure 2:**
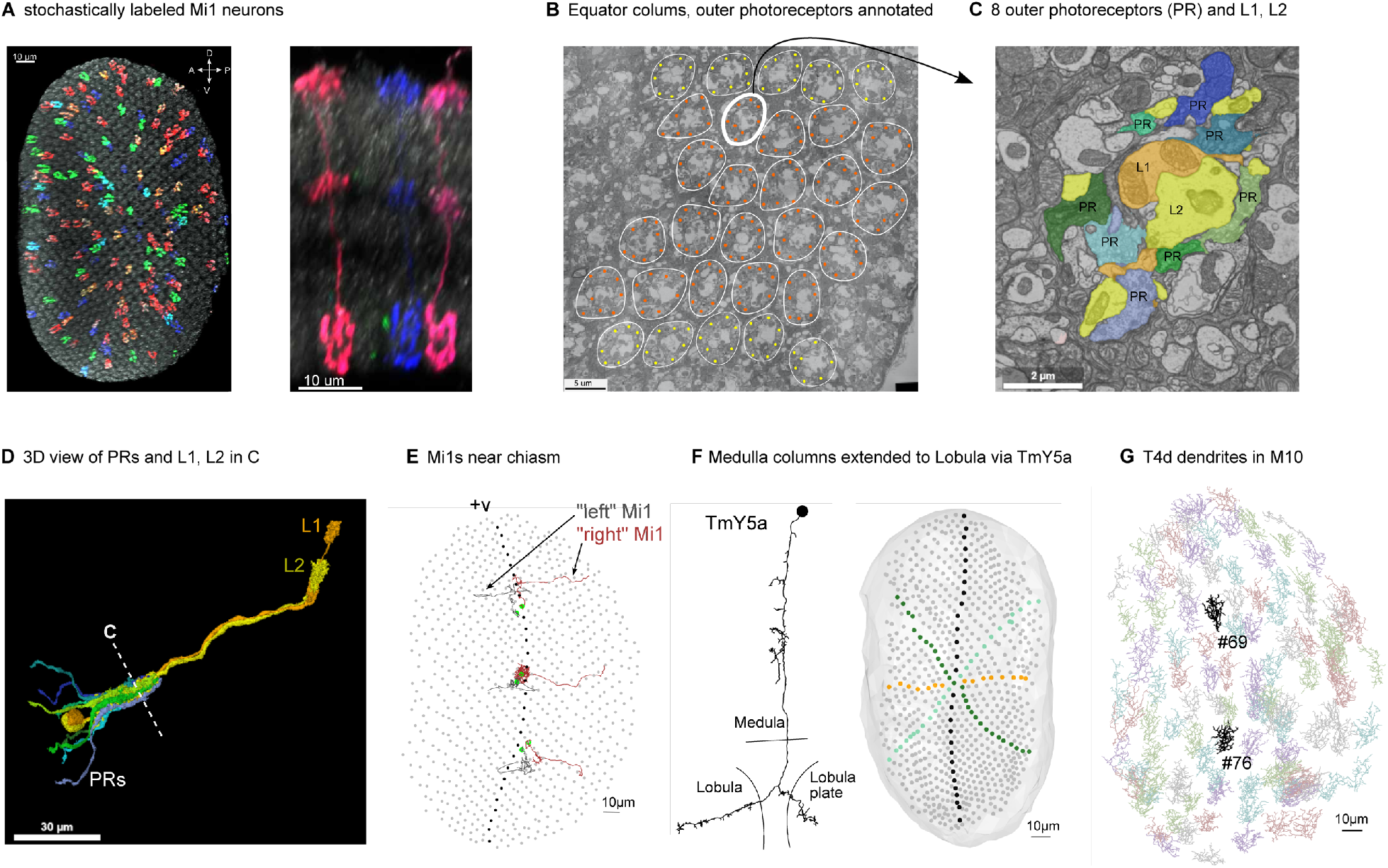
Anatomical considerations for mapping visual neurons throughout the medulla, related to Fig. 2. A. Left: light microscopy of stochastically labeled Mi1 cells in the background of nc82 staining, only showing the M10 portion of the medulla. Right: side view of stochastically labeled Mi1 cells (3 out of 4 columns have visible cells). **B**. A cross section of the equatorial region in the lamina, identified by cartridges (white contour) with 8 (orange) or fewer (yellow) outer photoreceptors. **C**. A zoomed-in view of a single cartridge showing L1/2 cells and 8 photoreceptor cells. L1 cells receive input from photoreceptor cells (R1-6) and output to Mi1 cells. **D**. 3D rendering of the same cartridge. **E**. Chiasm medulla columns (green) with corresponding Mi1 cells at 3 vertical locations identified by the twisting of R7/8 photoreceptor axons (not shown). For comparison, the central meridian is indicated in black. **F**. Extension of medulla column map to the lobula via interpolation of the positions of 63 reconstructed TmY5a neurons. **G**. The dendritic arbors of 114 reconstructed T4d cells in M10. The 2 highlighted neurons are the examples in Fig. 2G.

**Extended Data Figure 3:**
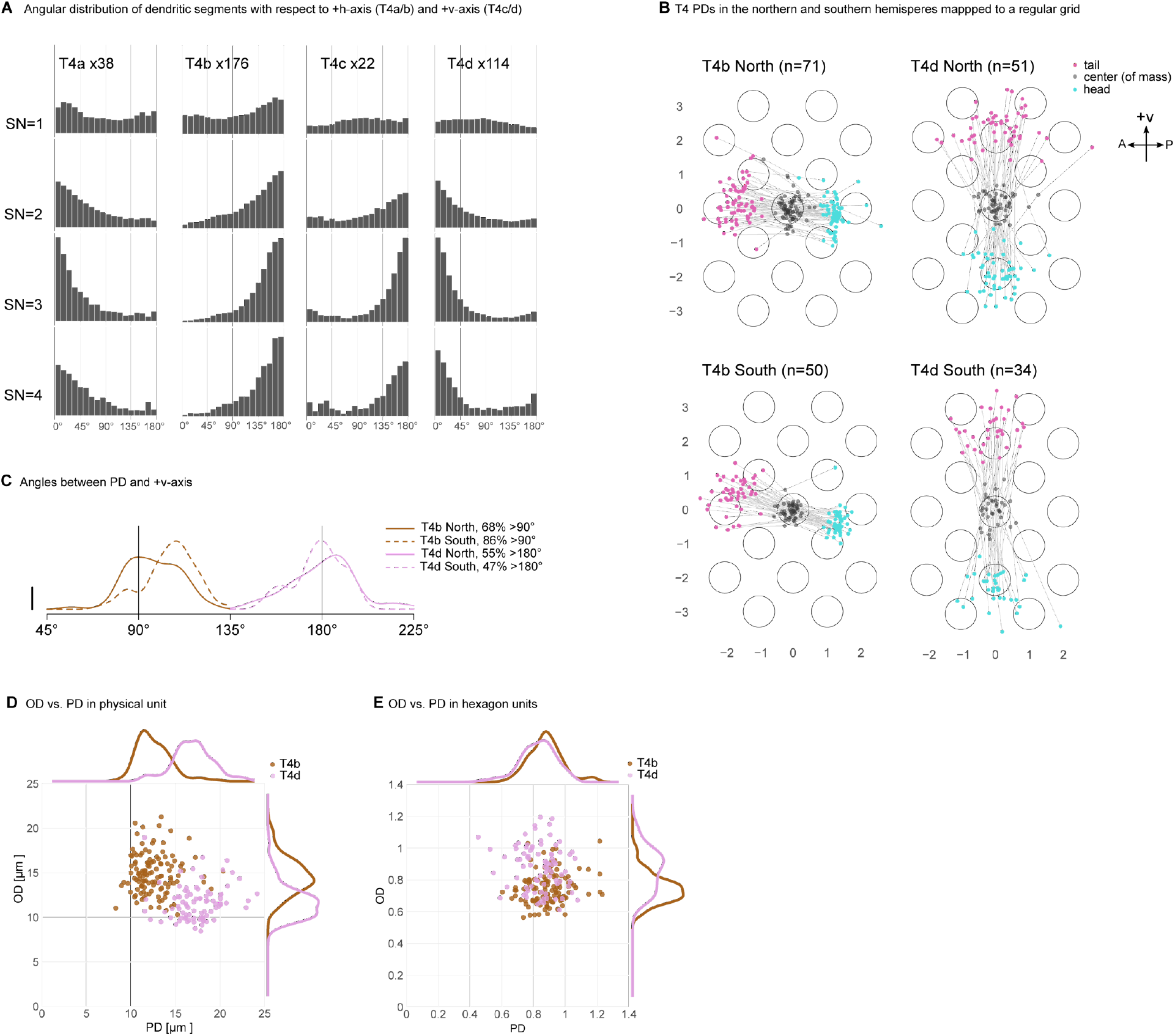
Analysis of dendritic morphology of T4 neurons, related to Fig. 2. **A**. The angular distribution of dendritic segments, treating each segment as a vector, and calculating the angle formed with respect to the grid axes. For T4a/b subtypes, the angle was computed with respect to the +h-axis, normalized by solid angle and grouped by Strahler number (SN). For T4c/d, the angle was computed with respect to the -v-axis. To compute the PD vector, we use SN = {2, 3} because these segments are abundant (compared to SN = 4) and have more consistent directions (compared to SN = 1). **B**. PD vectors mapped to a regular grid for T4b/d cells above (North) and below (South) the equator. **C**. Angles between the PDs and +v-axis above and below the equator. Wilcoxon rank test for the null-hypothesis that the distributions in the northern and southern hemispheres are the same yields a p-value = 0.00034 for T4b and 0.67 for T4d. **D**. OD vs. PD length in raw physical units [μm] in medulla. **E**. OD vs. PD length: T4b PDs and T4d ODs are normalized by the horizontal hexagon unit D_h_, while T4b ODs and T4d PDs are normalized by the vertical hexagon unit D_v_.

**Extended Data Figure 4:**
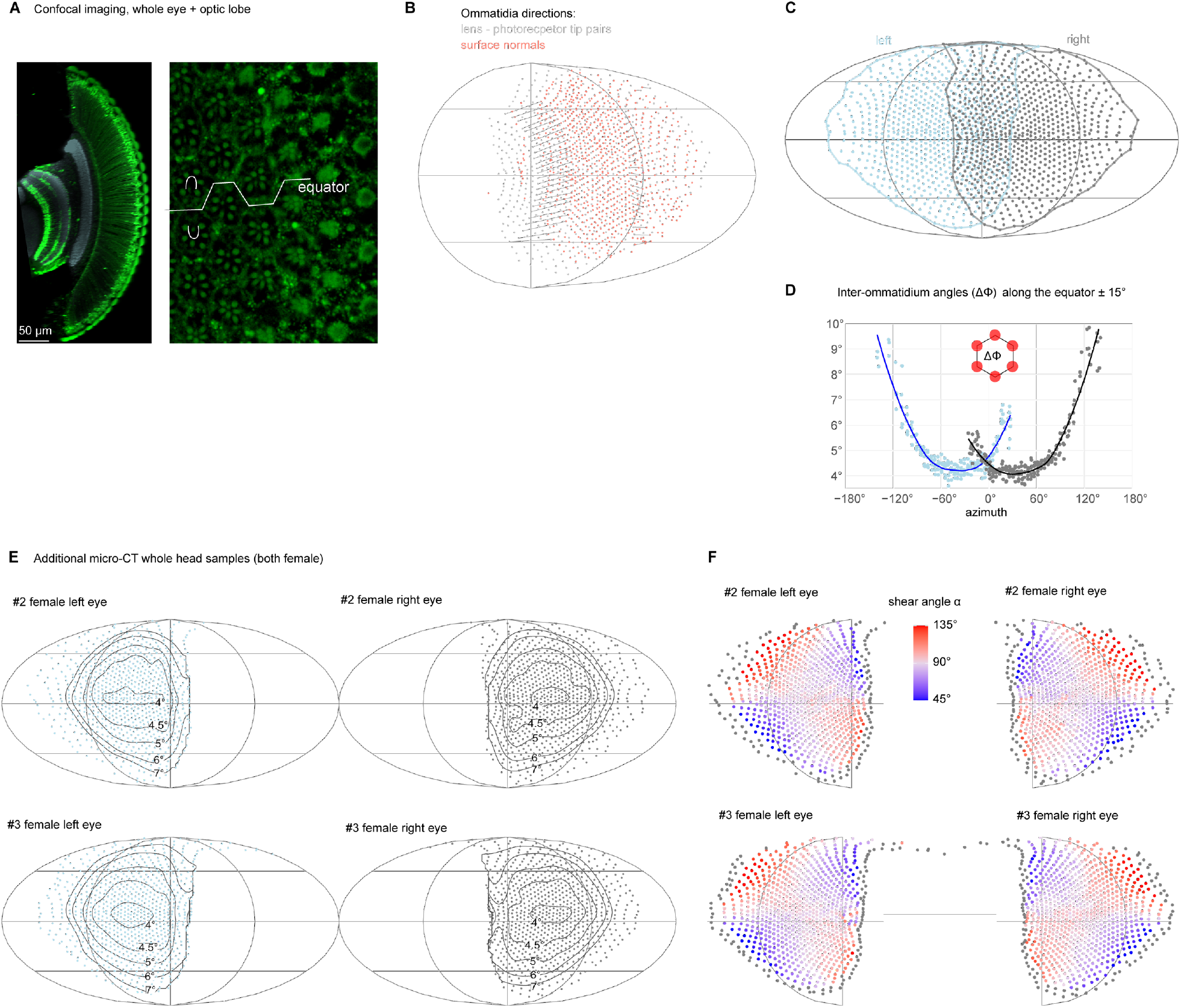
Ommatidia viewing directions in *Drosophila* compound eye maps, related to Fig. 3. **A**. Left: a high-resolution confocal image showing auto fluorescence (green) from ommatidia and many cells in the medulla. Lamina and medulla neuropils are visible (grey) due to nc82 antibody staining. Right: a different cross section showing the arrangement of individual photoreceptors near the equator. 6 dots (R1-R6) arranged an “n” or “u” shape can be readily seen in each ommatidium. The 7th smaller dot (R7+R8) in the center is often visible as well. **B**. Comparison of ommatidia directions defined by lens-photoreceptor tip pairs (gray, used in this study) and by surface normal (red). The surface normal is a typical approximation for the viewing direction, but this estimate differs substantially from that based on the high-resolution structure of each ommatidium. Two corresponding rows are connected with gray lines to illustrate the differences in different eye regions. Notably these differences are small near the center of the eye, and very large towards the front of the eye. **C**. Ommatidia directions and field of views (contours) for both eyes for the same fly as in Fig. 3. **D**. Six-neighbor inter-ommatidial angle (ΔΦ) along the equator (+/-15° elevation) for this same fly. **E**. Ommatidia directions and ΔΦ for 2 additional female flies. **F**. Shear angles for these 2 female flies, plotted as in Fig. 3J.

**Extended Data Figure 5:**
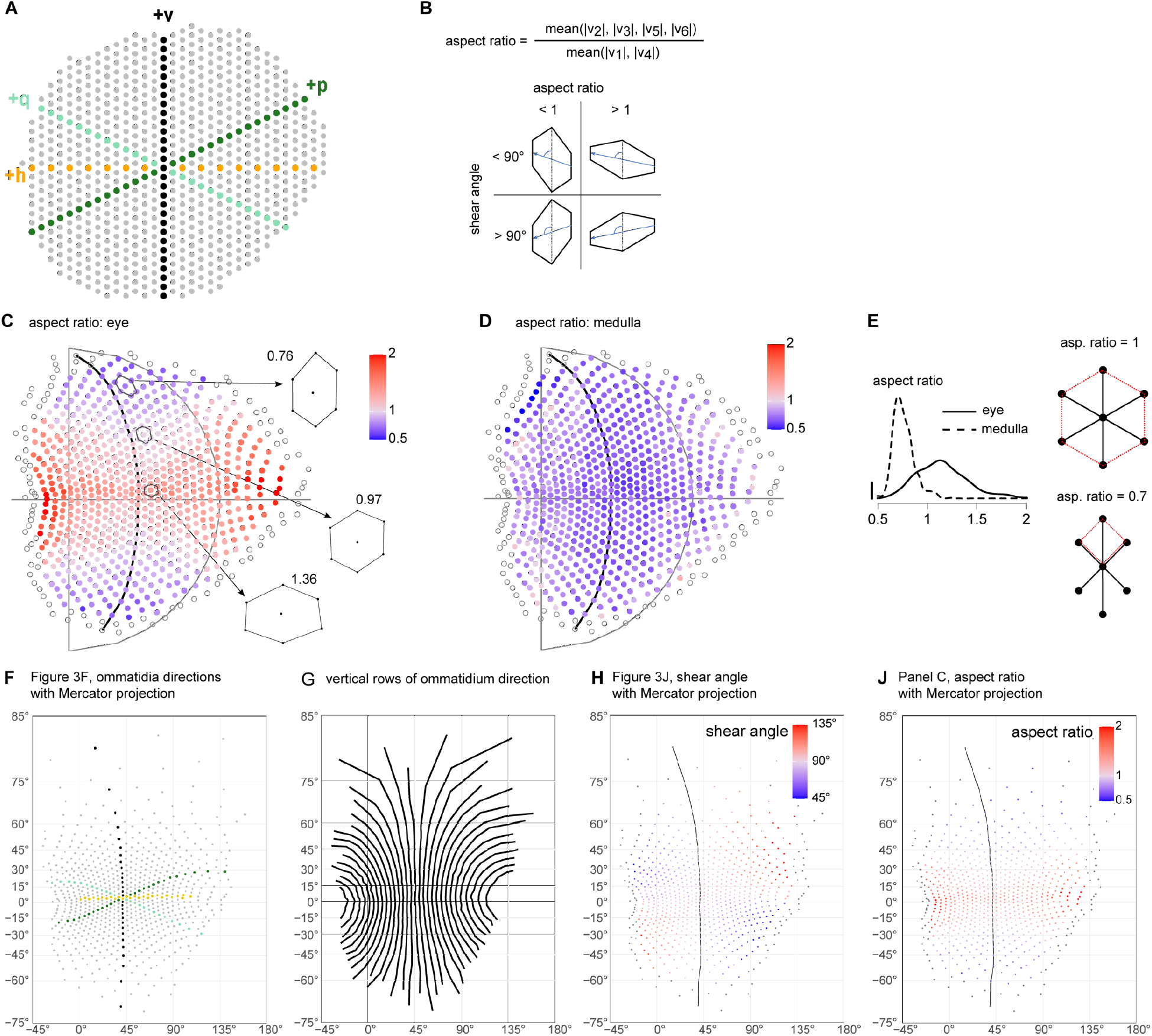
Quantification of the ommatidial viewing direction grids, related to Fig. 3. **A**. Ommatidia directions mapped onto a regular hexagonal grid. **B**. Definitions of aspect ratio and shear angle for a unit hexagon with examples. **C**. Aspect ratios calculated from ommatidia directions. **D**. Aspect ratios calculated from medulla columns. **E**. Distributions of aspect ratio for ommatidia and medulla columns. Comparison with the aspect ratios for a regular hexagonal grid and a regular square grid shows that the arrangement of ommatidia directions is more hexagonal while the arrangement of medulla columns is more square-like. **F**. Fig. 2F replotted using Mercator projection. **G**. Vertical rows of ommatidia directions given by the grid structure, shown using Mercator projection. **H**. Fig. 3J replotted using Mercator projection. **J**. The aspect ratio map in **C** replotted using Mercator projection.

**Extended Data Figure 6:**
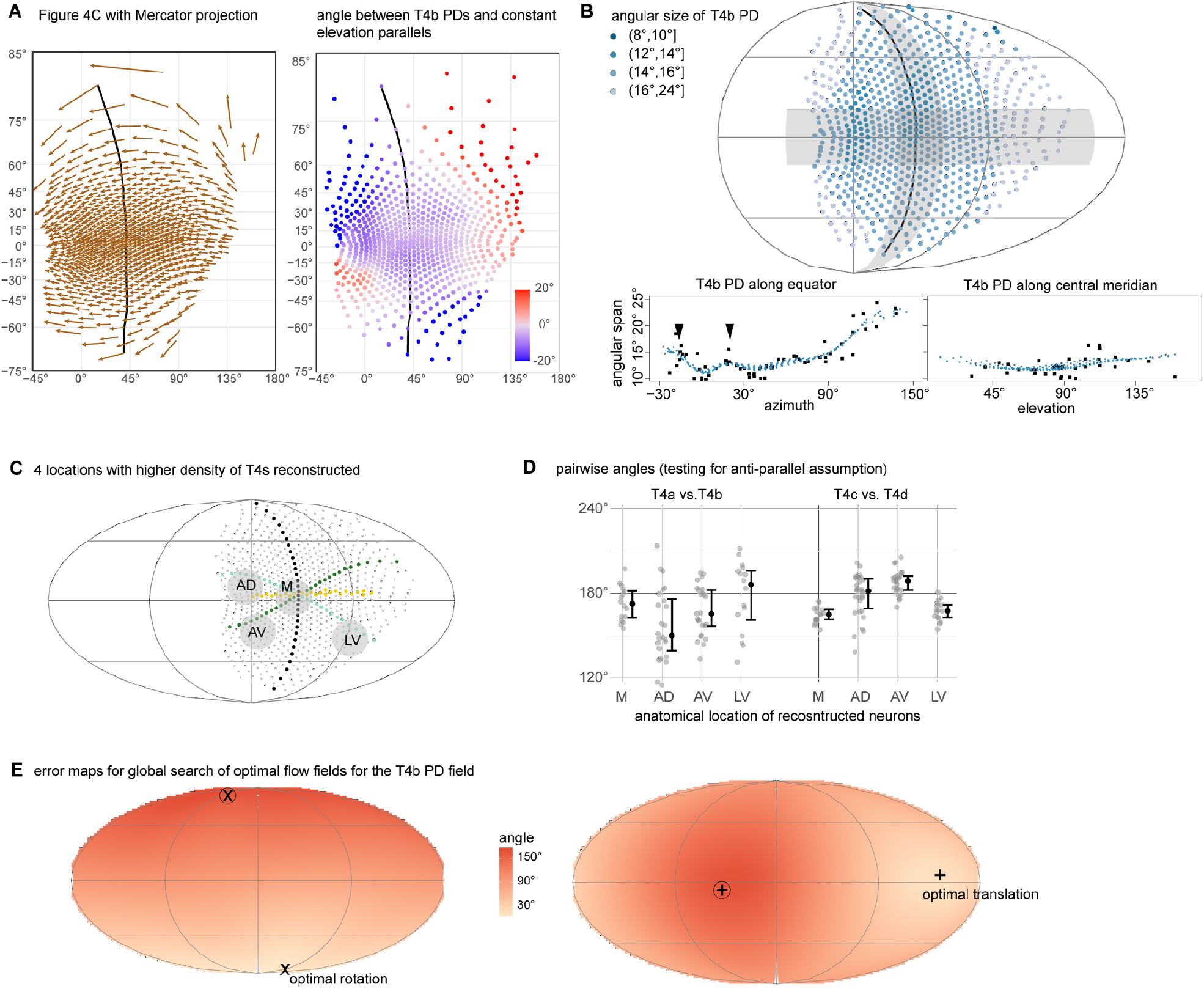
Further quantification of T4b PD distribution, related to Fig. 4. **A**. Left: Fig. 4C replotted using Mercator projection; right: map of angle between T4b PD field and the constant elevation parallels. **B**. Visual angles subtended by T4b PD vectors (i.e., angular size). Scatter plots show the reconstructed T4b PDs (black dots, also in Fig. 4B) and interpolated ones (blue dots, also in Fig. 4C) along the equator (+/-15° horizontal shaded band) and the central meridian (+/-15° vertical shaded crescent). Most T4b PDs span between 10º-15º degrees, but there are almost 2-fold differences found across the eye, with larger spans towards the rear and smaller spans near the equatorial higher-acuity zone and front (black arrowheads). **C**. In an early pilot study we reconstructed all T4 subtypes (16∼20 cells) at each of these four locations. **D**. We first mapped these T4s’ PD vectors to the regular grid in Fig. 2H. Then at each location, we computed the angles between all T4a vs T4b pairs. Similarly, for T4c vs T4d. **E**. Global search for optimal optic flow fields yielded these error maps, showing the average angular differences between the T4b PD field and the optic flow field induced by a rotation (left) or translation (right) along that direction (see Methods: Ideal optic flow). Symbols “+” and “X” denote the axes of translational and rotational motion with minimal angular difference, respectively. Symbols ⨁ and ⨂ denote those with maximal differences (minimum and maximum are antipodal).

**Extended Data Figure 7:**
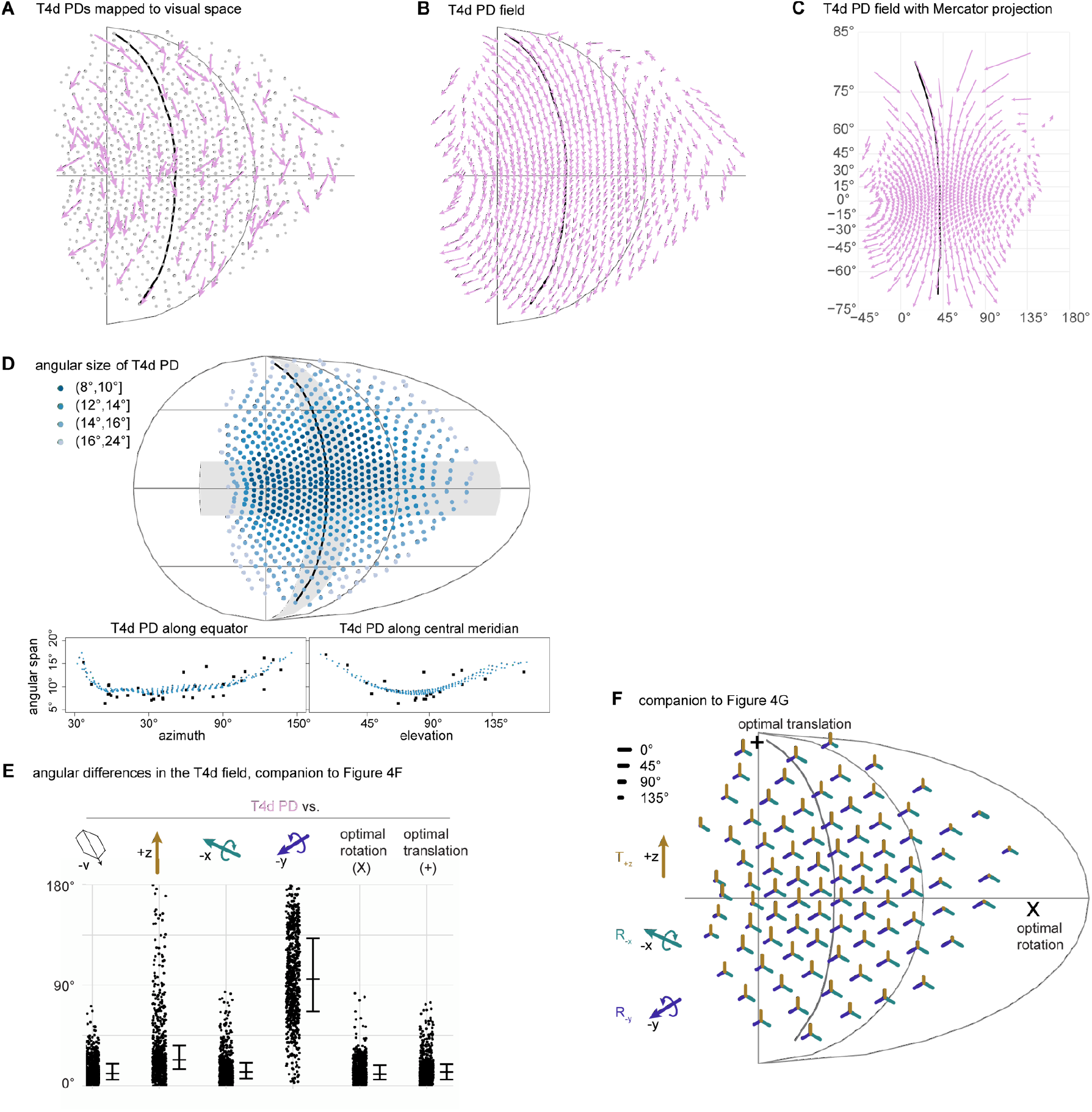
T4d neuron analysis, related to Fig. 4. **A**. T4d PDs mapped to eye coordinates. **B**. Interpolated T4d PD field (arrows are re-scaled to 50% of length in **A**). **C**. The T4d PD field from **B** replotted using Mercator projection. **D**. Visual angles subtended by T4d PD vectors (i.e., angular size). Scatter plots show the reconstructed T4d PDs (black dots, also in (A)) and interpolated ones (blue dots, also in (B)) along the equator (+/-15° horizontal shaded band) and the central meridian (+/-15° vertical shaded crescent). **E**. Agular differences between T4d PD field, -v-axis, three cardinal self-motion optic flow fields (lift, leftward roll, and upward pitch), and optimized self-motion flow fields. The horizontal bars represent 25%, 50% and 75% quantiles. **F**. Spatial distribution of angular differences between T4b PD field and the three cardinal self-motion optic flow fields.

## Supplementary Information

**Supplementary Video 1: Summary of eyemap, enabling the projection of the compound eye’s visal space into the neural circuits of the optic lobe**

Whole-head μCT scan with overlaid EM reconstructed neurons, showing the columnar structure of the compound eye and optic lobe. Ommatidia directions were determined by the lens-photoreceptor tip pairs. Medulla columns were defined as the Mi1 cells’ arbor in layer M10. Finally, we established an eyemap: a 1-to-1 mapping between ommatidia directions and medulla columns.

**Supplementary Video 2: Illustration of how the dendritic orientation of T4 neurons facilitates motion detection in different directions**

There are 4 subtypes of T4 cells, innervating 4 distinct layers in lobula plate. A T4 cell’s preferred direction (PD) is computed based on its dendritic arborization pattern. PDs can be mapped to eye coordinates using the eyemap defined in Supplementary Video 1. In the central region of the eye, the four T4 subtypes are well aligned with directions of motion in the 4 cardinal directions (forward, backward, up, and down).

**Supplementary Data files 1-4: galleries of T4 neurons with PDs**

All T4 neurons reconstructed in the FAFB data set: 38 T4a, 176 T4b, 22 T4c, 114 T4d, are plotted in a similar fashion as in Fig. 2G. Using the eyemap established in Fig. 4A, we include the position (elevation and azimuth angles) in the eye coordinate. The angle between T4’s PD and the local meridian line is computed, instead of using the +v-axis as the reference as in Fig. 2G. The meridian line is defined as the direction line going from south pole to north pole in the eye reference frame (often close to the +v-axis). The cell and surrounding columns are also aligned such that the vertical direction in the plot coincide with the meridian direction. A summary of the Strahler number (SN) analysis for each cell is included.

## Notes

### Competing Interest Statement

The authors have declared no competing interest.

