## Supplementary Files, T4 galleries for "Eye structure shapes neuron function in *Drosophila* motion vision": Supplementary_Data_File1_T4a.pdf

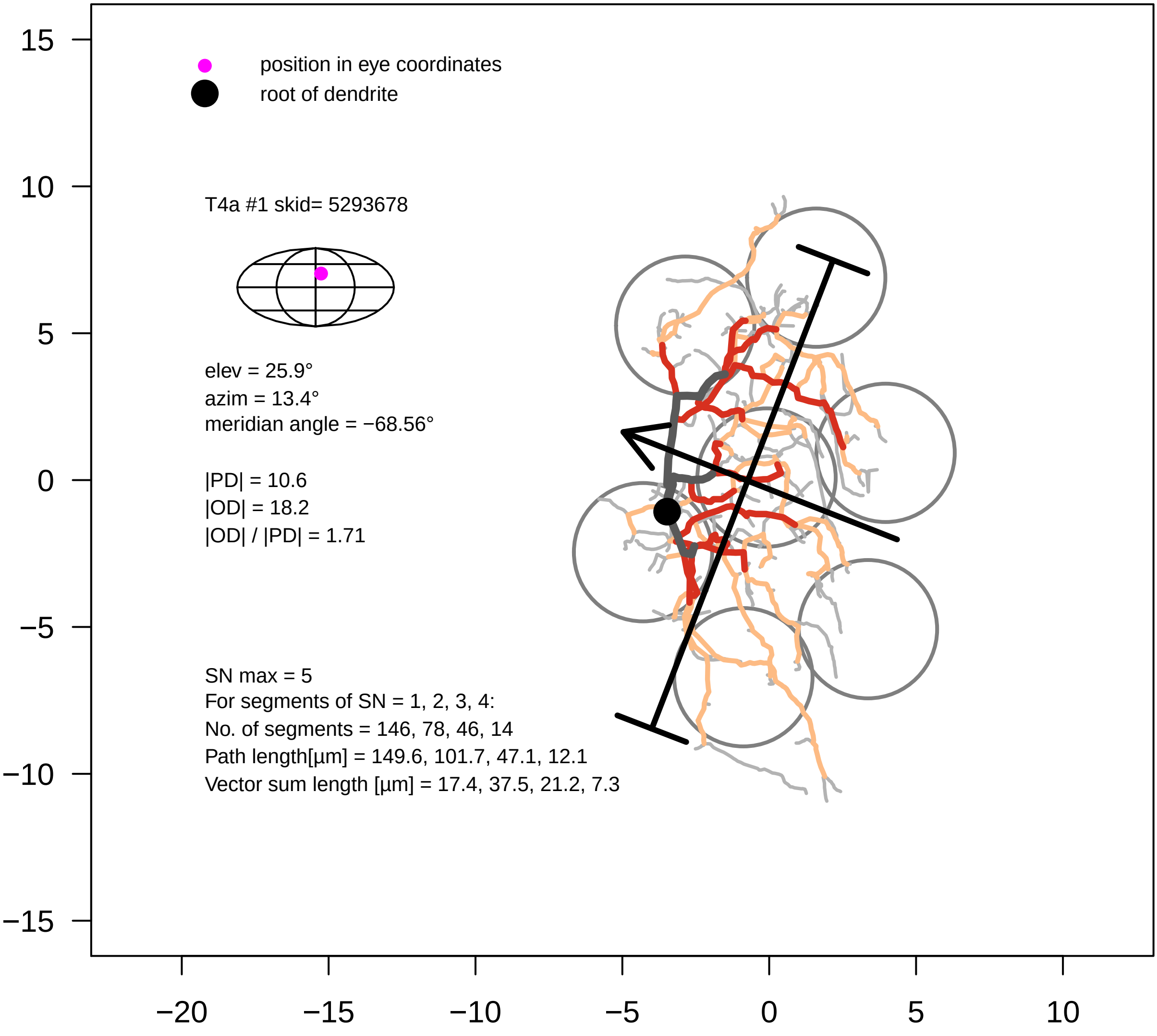

position in eye coordinates
root of dendrite
T4a #1 skid= 5293678
elev = 25.9°
azim = 13.4°
meridian angle = −68.56°
|PD| = 10.6
|OD| = 18.2
|OD| / |PD| = 1.71
SN max = 5
For segments of SN = 1, 2, 3, 4:
No. of segments = 146, 78, 46, 14
Path length[µm] = 149.6, 101.7, 47.1, 12.1
Vector sum length [µm] = 17.4, 37.5, 21.2, 7.3
15
10
5
0
−5
−10
−15
−20
−15
−10
−5
0
5
10

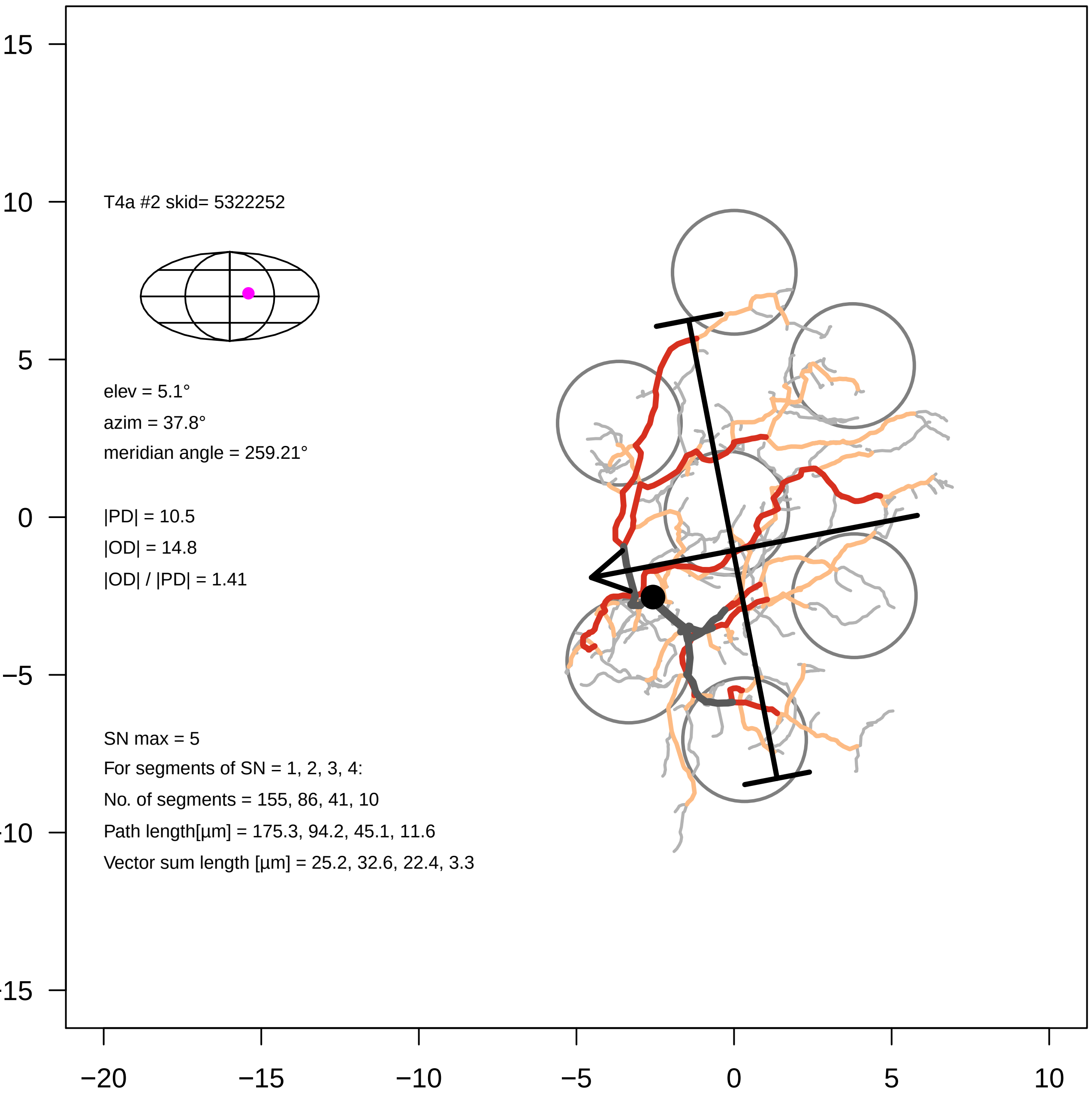
T4a #2 skid= 5322252
elev = 5.1°
azim = 37.8°
meridian angle = 259.21°
|PD| = 10.5
|OD| = 14.8
|OD| / |PD| = 1.41
SN max = 5
For segments of SN = 1, 2, 3, 4:
No. of segments = 155, 86, 41, 10
Path length[µm] = 175.3, 94.2, 45.1, 11.6
Vector sum length [µm] = 25.2, 32.6, 22.4, 3.3
15
10
5
0
−5
−10
−15
−20
−15
−10
−5
0
5
10

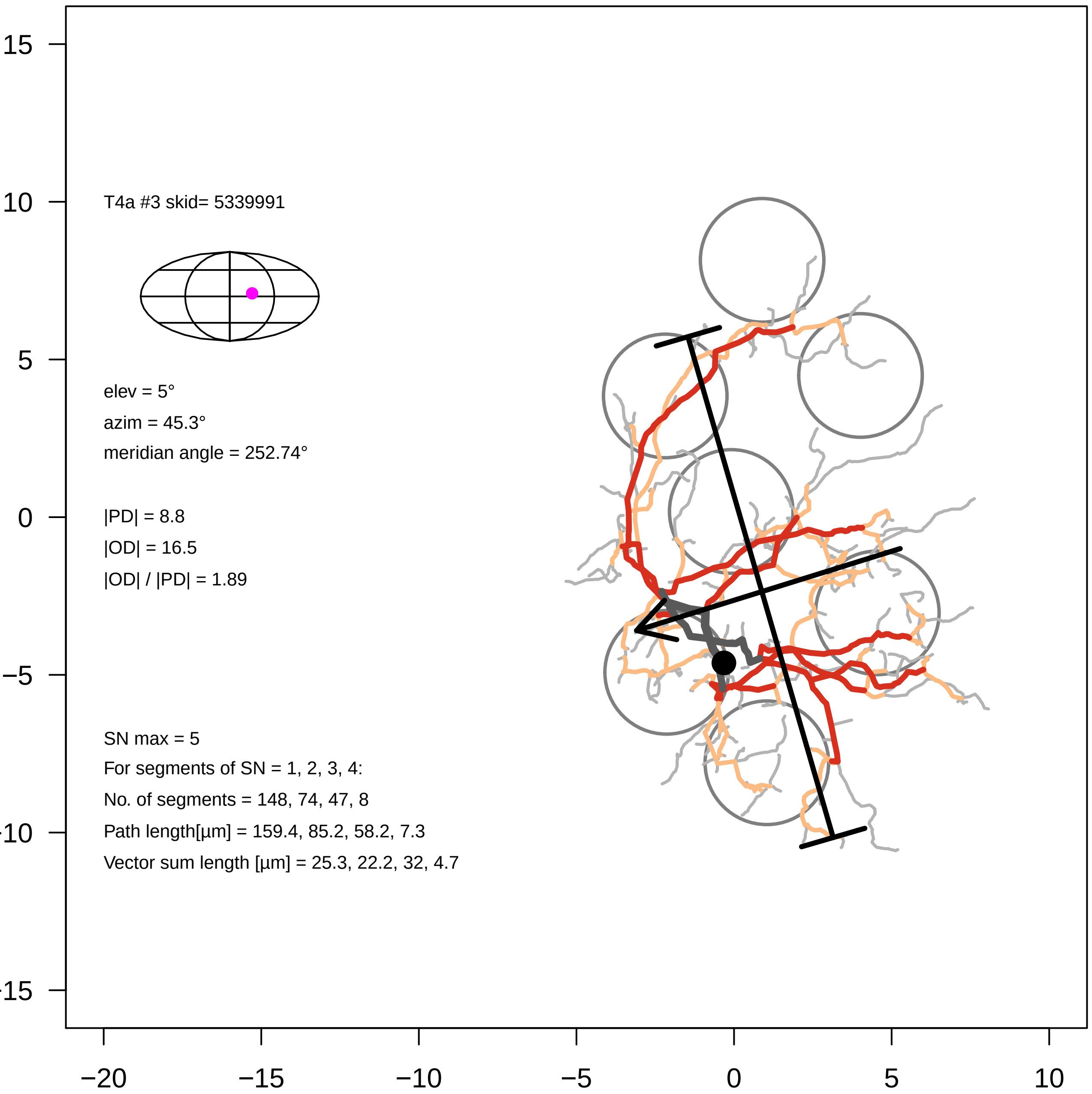
T4a #3 skid= 5339991
elev = 5°
azim = 45.3°
meridian angle = 252.74°
|PD| = 8.8
|OD| = 16.5
|OD| / |PD| = 1.89
SN max = 5
For segments of SN = 1, 2, 3, 4:
No. of segments = 148, 74, 47, 8
Path length[µm] = 159.4, 85.2, 58.2, 7.3
Vector sum length [µm] = 25.3, 22.2, 32, 4.7
15
10
5
0
−5
−10
−15
−20
−15
−10
−5
0
5
10

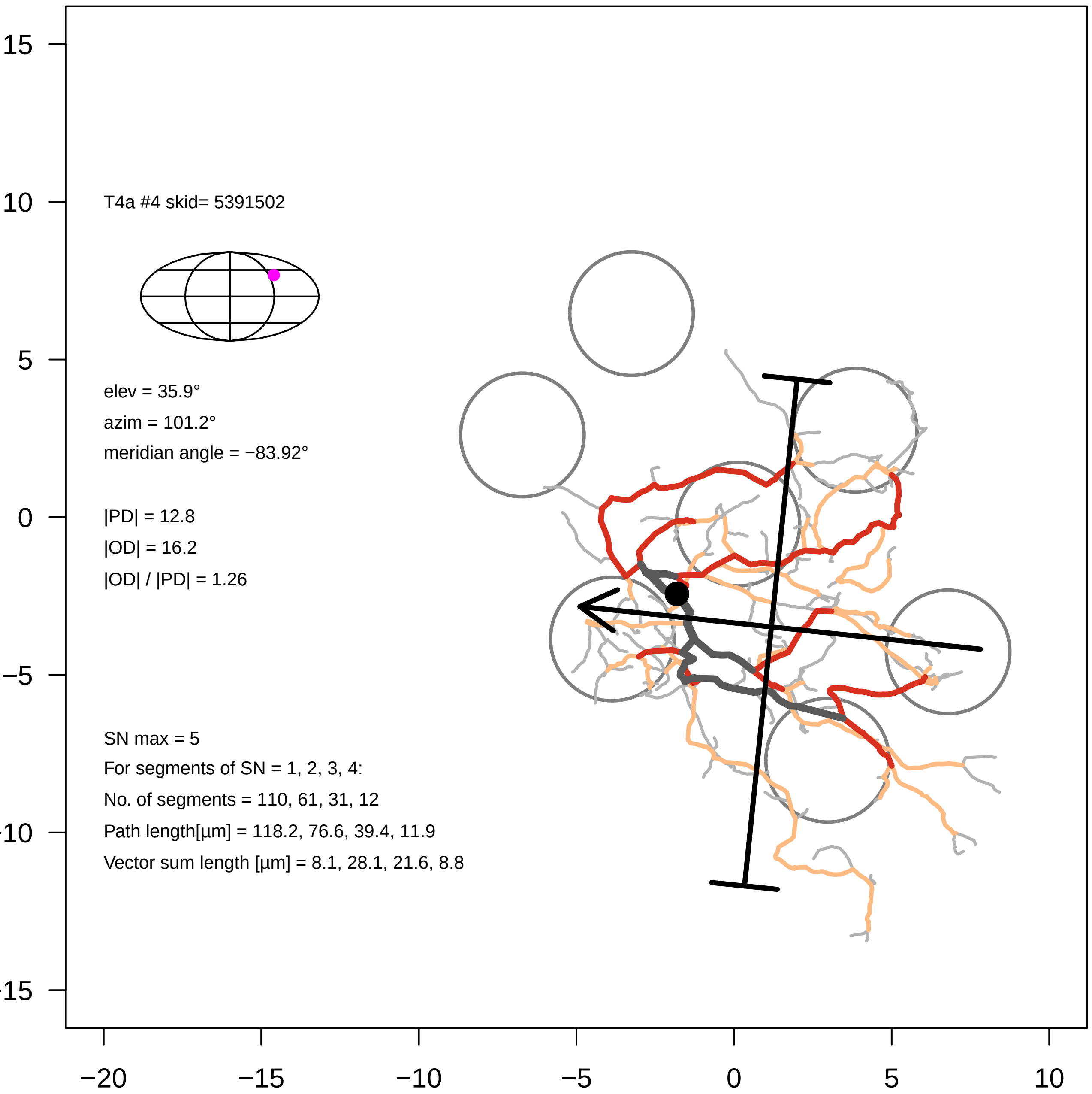

T4a #4 skid= 5391502
elev = 35.9°
azim = 101.2°
meridian angle = −83.92°
|PD| = 12.8
|OD| = 16.2
|OD| / |PD| = 1.26
SN max = 5
For segments of SN = 1, 2, 3, 4:
No. of segments = 110, 61, 31, 12
Path length[µm] = 118.2, 76.6, 39.4, 11.9
Vector sum length [µm] = 8.1, 28.1, 21.6, 8.8
15
10
5
0
−5
-10
-15
−20
−15
−10
−5
0
5
10

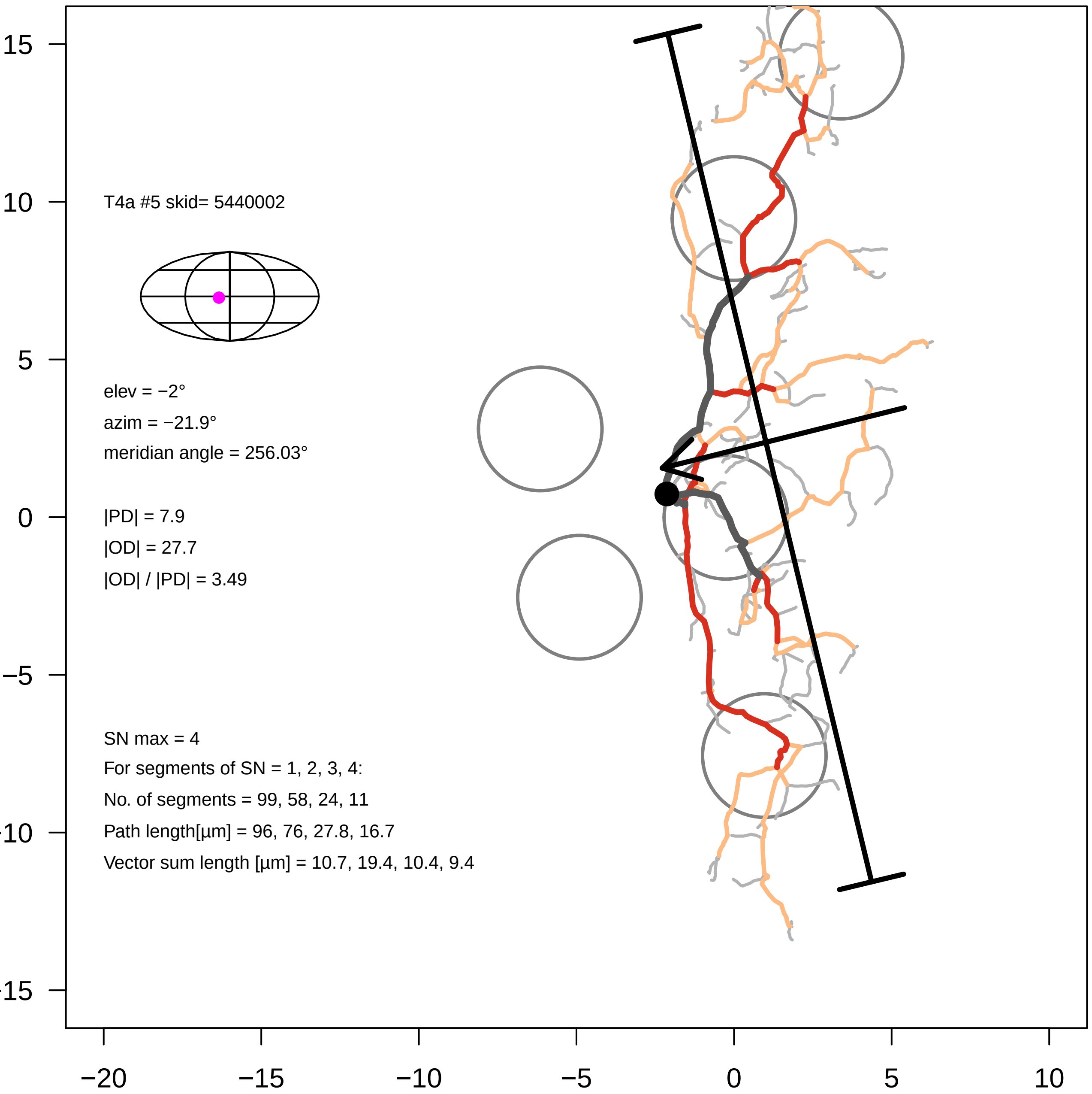

T4a #5 skid= 5440002
elev = −2°
azim = −21.9°
meridian angle = 256.03°
|PD| = 7.9
|OD| = 27.7
|OD| / |PD| = 3.49
SN max = 4
For segments of SN = 1, 2, 3, 4:
No. of segments = 99, 58, 24, 11
Path length[µm] = 96, 76, 27.8, 16.7
Vector sum length [µm] = 10.7, 19.4, 10.4, 9.4
15
10
5
0
−5
−10
−15
−20
−15
−10
−5
0
5
10

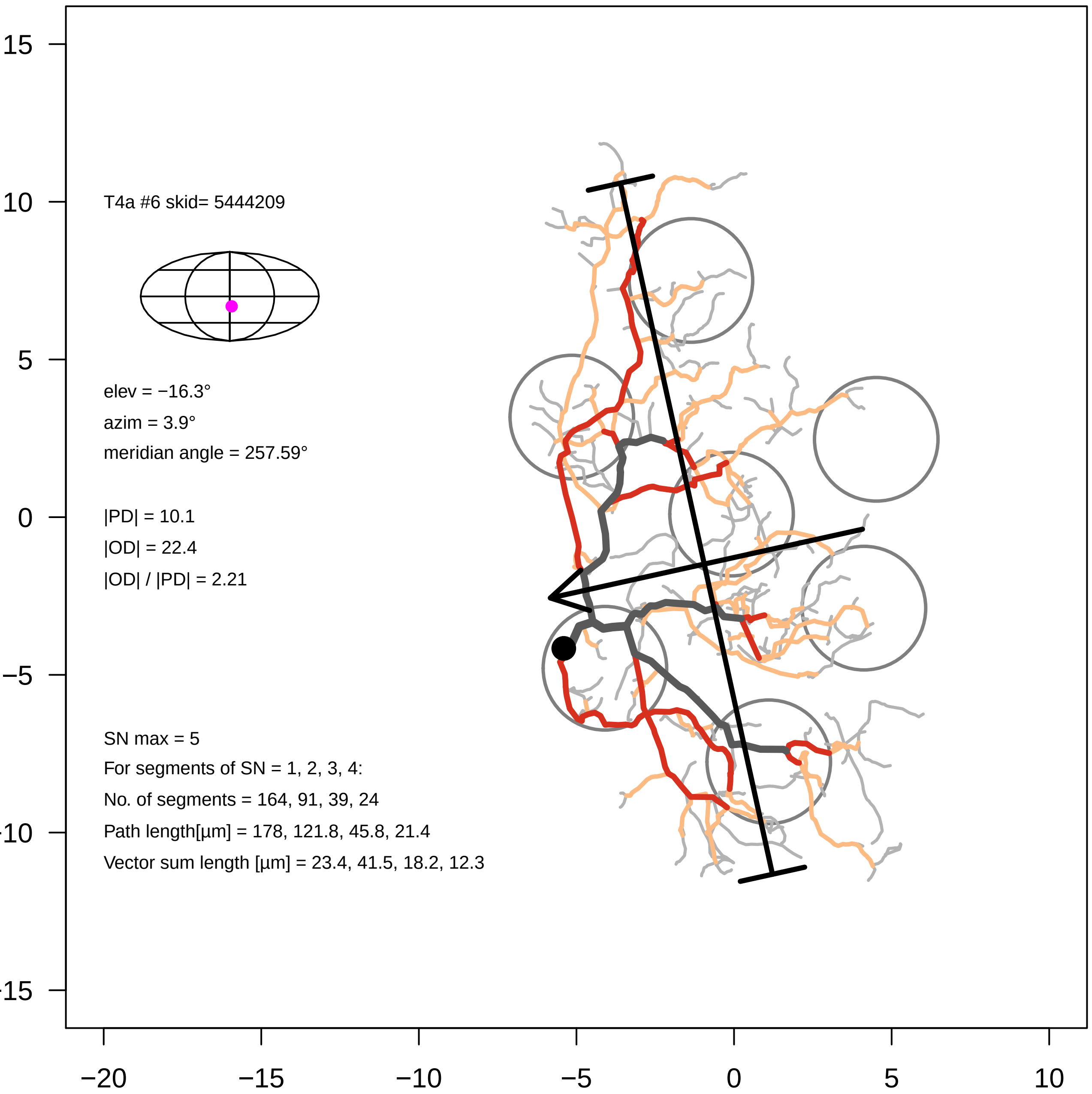

T4a #6 skid= 5444209
elev = −16.3°
azim = 3.9°
meridian angle = 257.59°
|PD| = 10.1
|OD| = 22.4
|OD| / |PD| = 2.21
SN max = 5
For segments of SN = 1, 2, 3, 4:
No. of segments = 164, 91, 39, 24
Path length[µm] = 178, 121.8, 45.8, 21.4
Vector sum length [µm] = 23.4, 41.5, 18.2, 12.3
15
10
5
0
−5
−10
−15
−20
−15
−10
−5
0
5
10

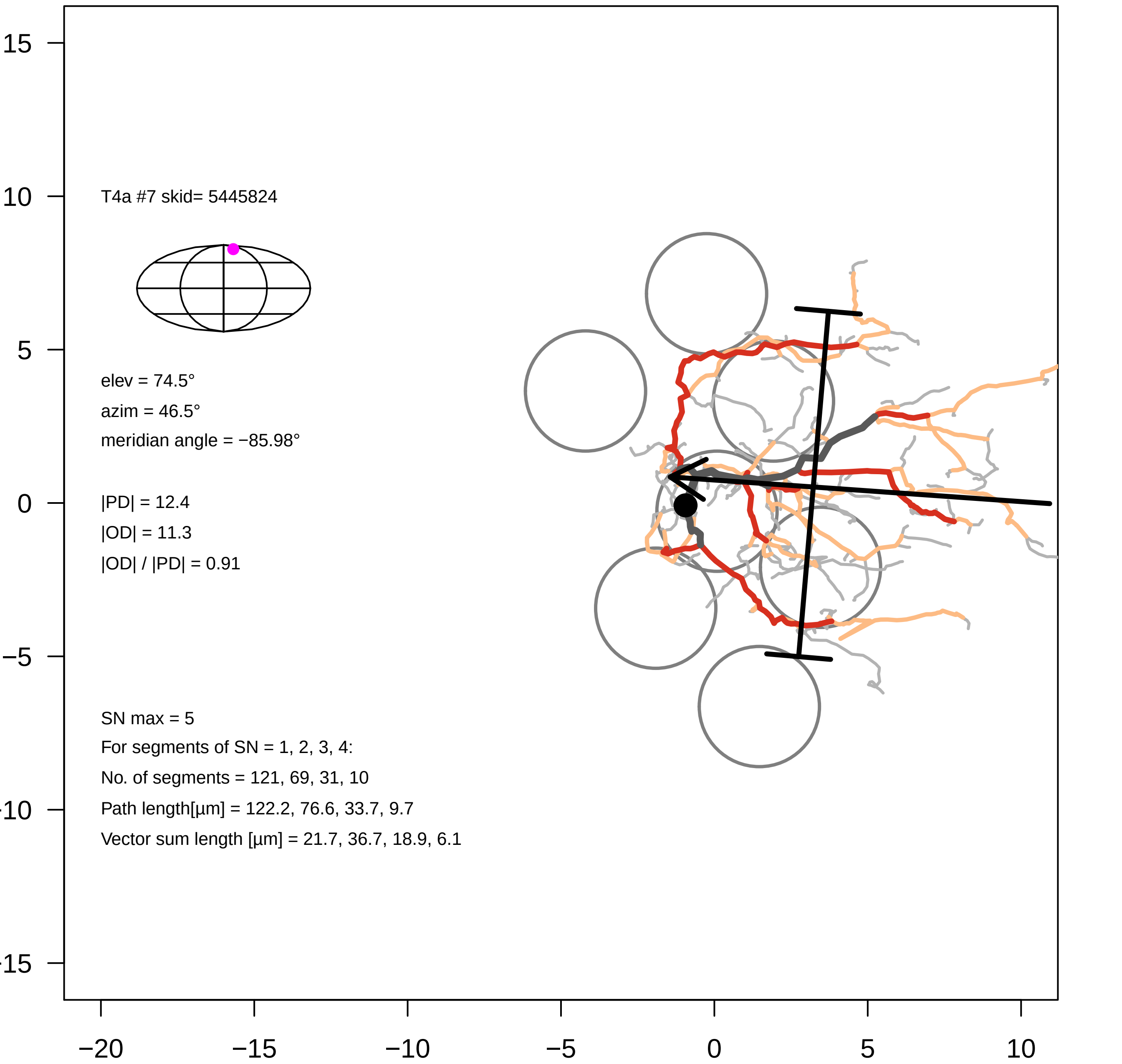

T4a #7 skid= 5445824
elev = 74.5°
azim = 46.5°
meridian angle = −85.98°
|PD| = 12.4
|OD| = 11.3
|OD| / |PD| = 0.91
SN max = 5
For segments of SN = 1, 2, 3, 4:
No. of segments = 121, 69, 31, 10
Path length[µm] = 122.2, 76.6, 33.7, 9.7
Vector sum length [µm] = 21.7, 36.7, 18.9, 6.1
15
10
5
0
−5
−10
−15
−20
−15
−10
−5
0
5
10

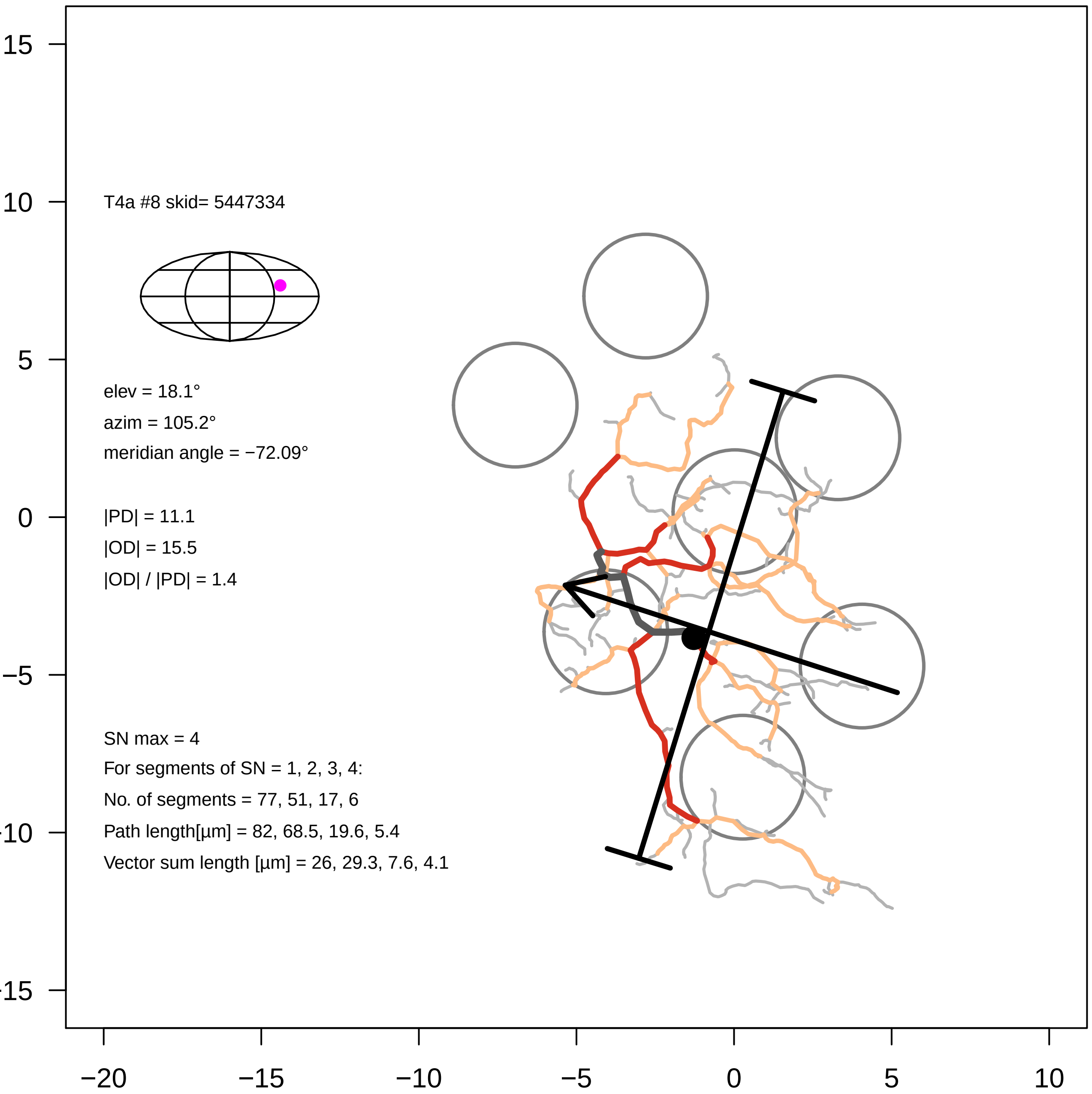

T4a #8 skid= 5447334
elev = 18.1°
azim = 105.2°
meridian angle = −72.09°
|PD| = 11.1
|OD| = 15.5
|OD| / |PD| = 1.4
SN max = 4
For segments of SN = 1, 2, 3, 4:
No. of segments = 77, 51, 17, 6
Path length[µm] = 82, 68.5, 19.6, 5.4
Vector sum length [µm] = 26, 29.3, 7.6, 4.1
15
10
5
0
−5
−10
−15
−20
−15
−10
−5
0
5
10

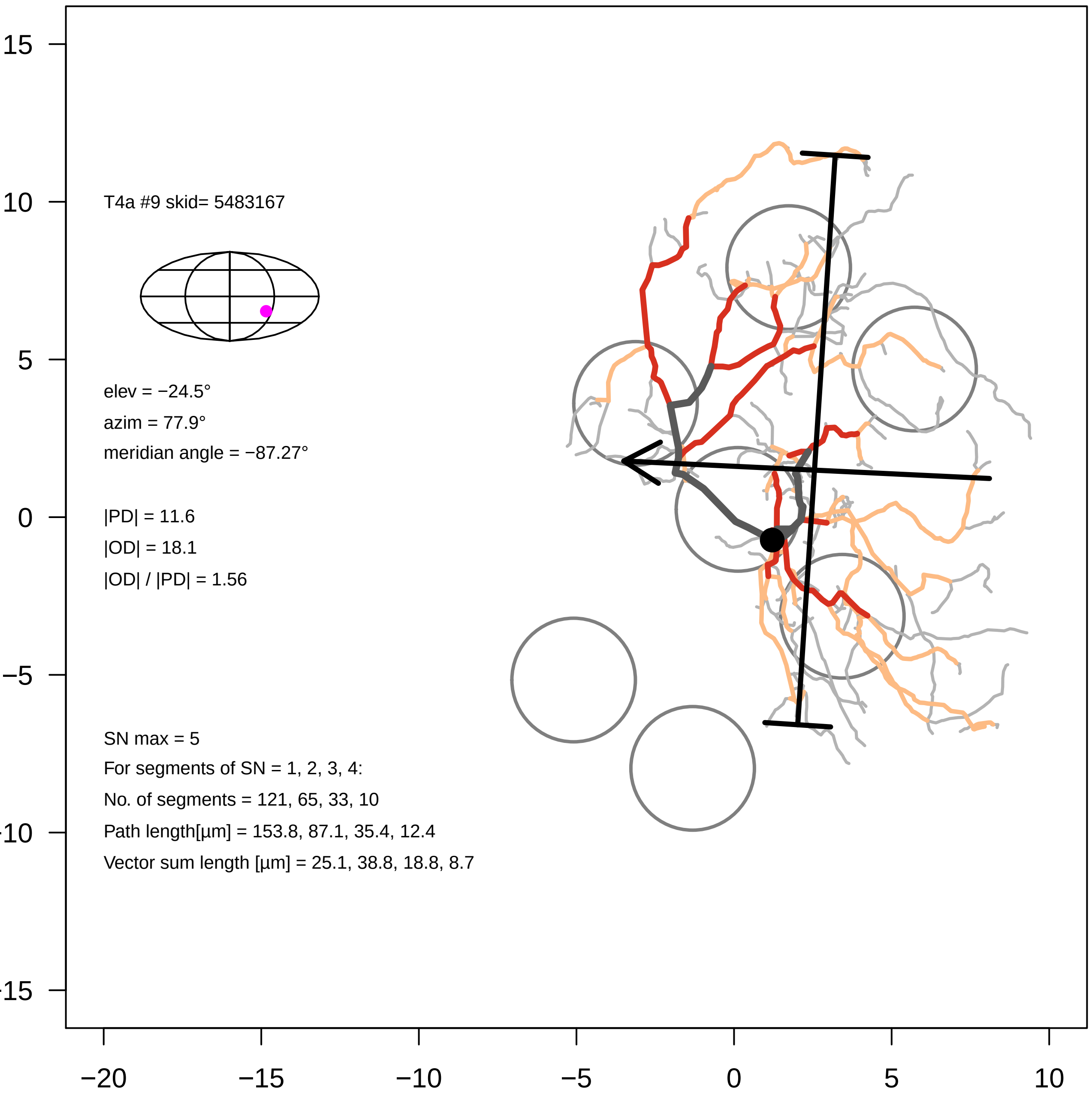

T4a #9 skid= 5483167
elev = −24.5°
azim = 77.9°
meridian angle = −87.27°
|PD| = 11.6
|OD| = 18.1
|OD| / |PD| = 1.56
SN max = 5
For segments of SN = 1, 2, 3, 4:
No. of segments = 121, 65, 33, 10
Path length[µm] = 153.8, 87.1, 35.4, 12.4
Vector sum length [µm] = 25.1, 38.8, 18.8, 8.7
15
10
5
0
−5
−10
−15
−20
−15
−10
−5
0
5
10

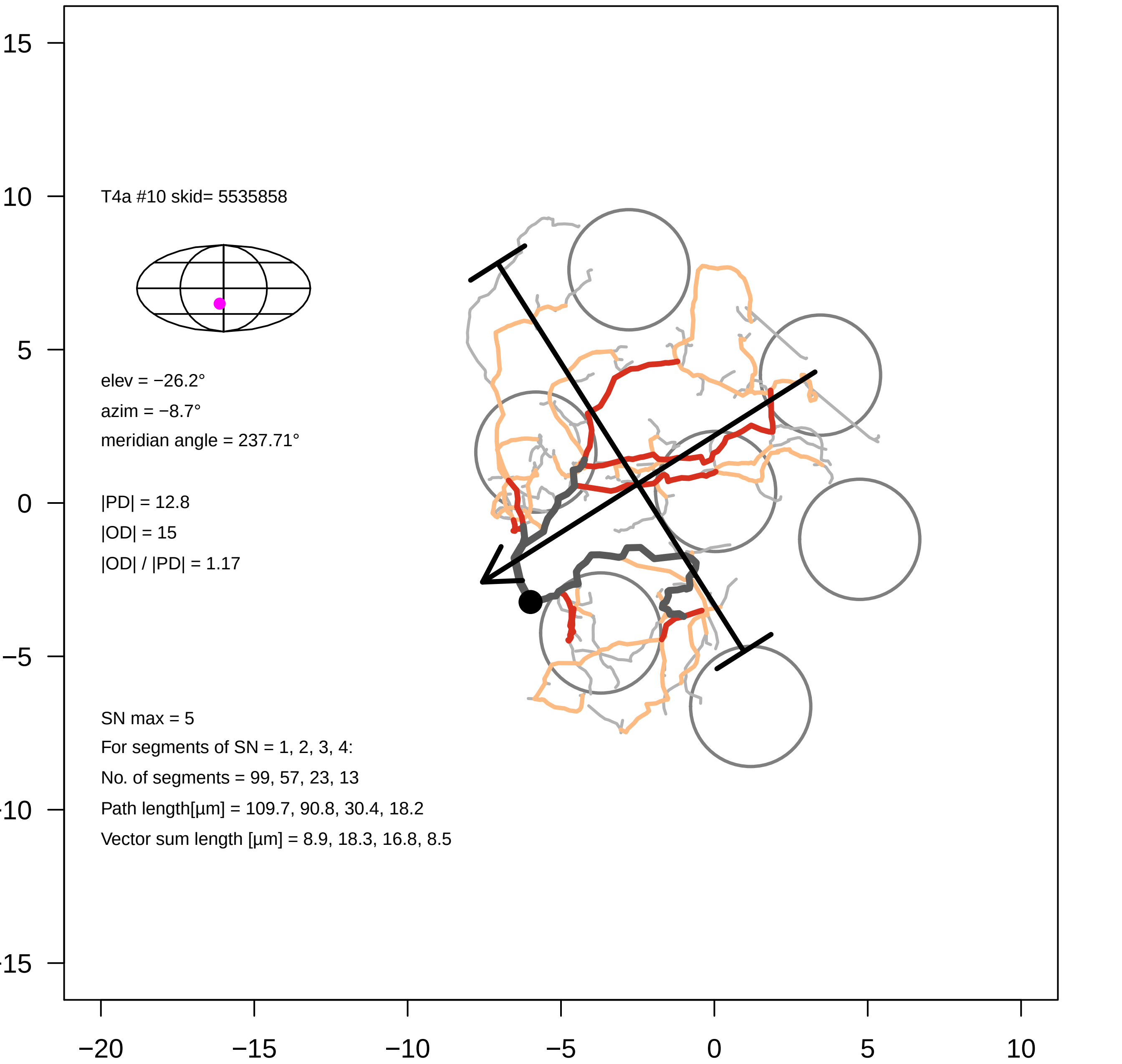
T4a #10 skid= 5535858
elev = −26.2°
azim = −8.7°
meridian angle = 237.71°
|PD| = 12.8
|OD| = 15
|OD| / |PD| = 1.17
SN max = 5
For segments of SN = 1, 2, 3, 4:
No. of segments = 99, 57, 23, 13
Path length[µm] = 109.7, 90.8, 30.4, 18.2
Vector sum length [µm] = 8.9, 18.3, 16.8, 8.5
15
10
5
0
−5
−10
−15
−20
−15
−10
−5
0
5
10

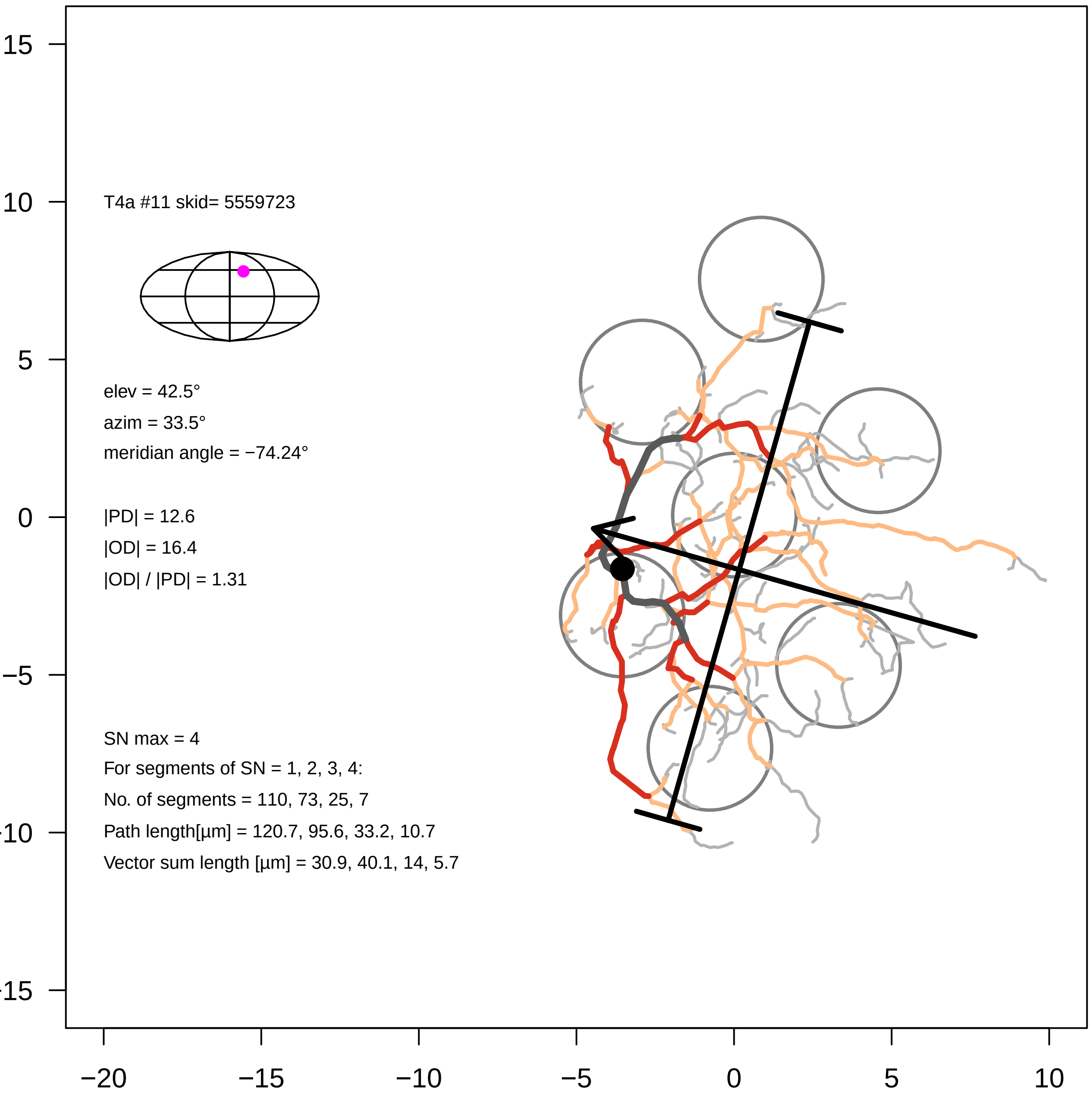

T4a #11 skid= 5559723
elev = 42.5°
azim = 33.5°
meridian angle = −74.24°
|PD| = 12.6
|OD| = 16.4
|OD| / |PD| = 1.31
SN max = 4
For segments of SN = 1, 2, 3, 4:
No. of segments = 110, 73, 25, 7
Path length[µm] = 120.7, 95.6, 33.2, 10.7
Vector sum length [µm] = 30.9, 40.1, 14, 5.7
15
10
5
0
−5
−10
−15
−20
−15
−10
−5
0
5
10

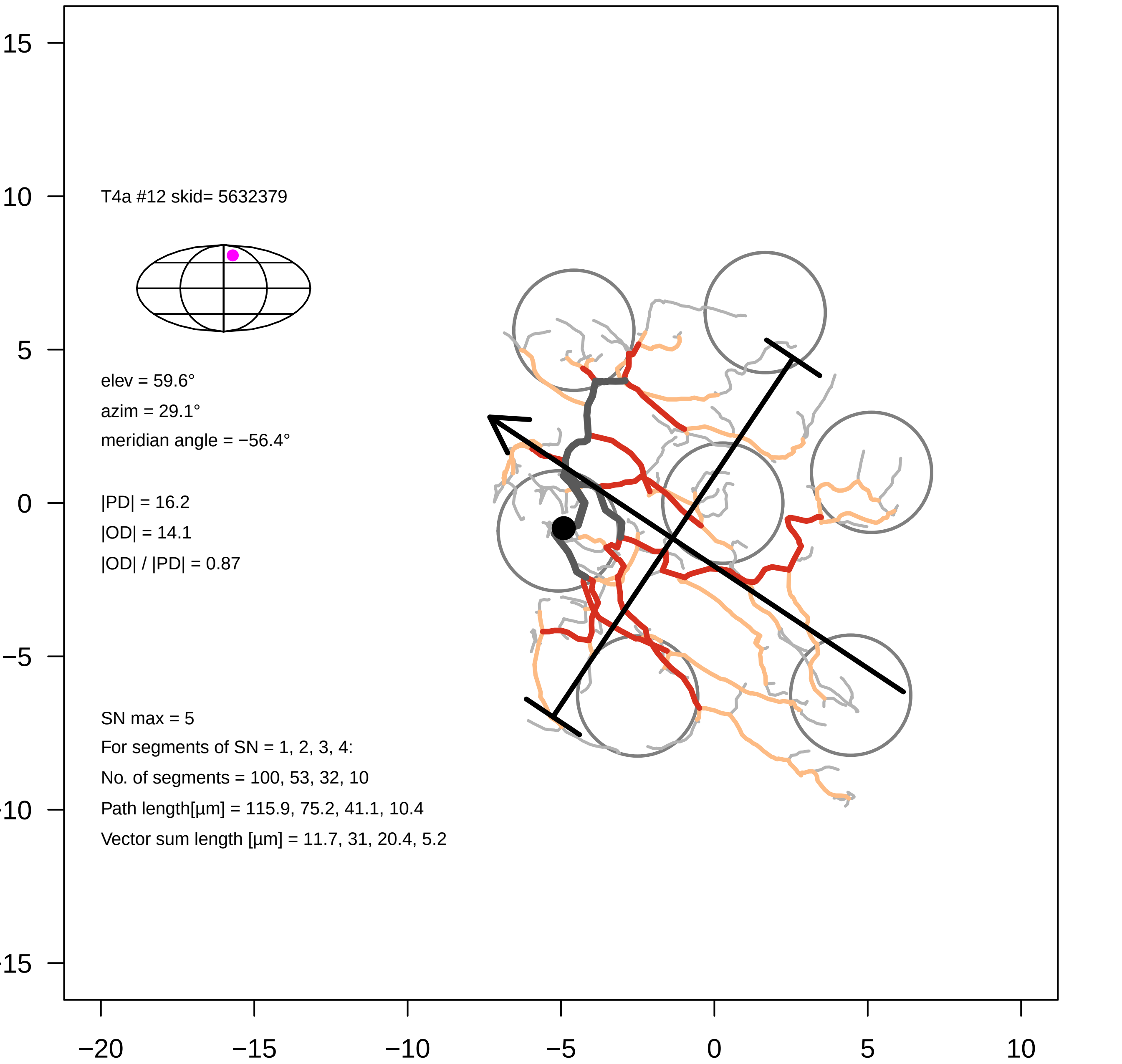

T4a #12 skid= 5632379
elev = 59.6°
azim = 29.1°
meridian angle = −56.4°
|PD| = 16.2
|OD| = 14.1
|OD| / |PD| = 0.87
SN max = 5
For segments of SN = 1, 2, 3, 4:
No. of segments = 100, 53, 32, 10
Path length[µm] = 115.9, 75.2, 41.1, 10.4
Vector sum length [µm] = 11.7, 31, 20.4, 5.2
15
10
5
0
−5
−10
−15
−20
−15
−10
−5
0
5
10

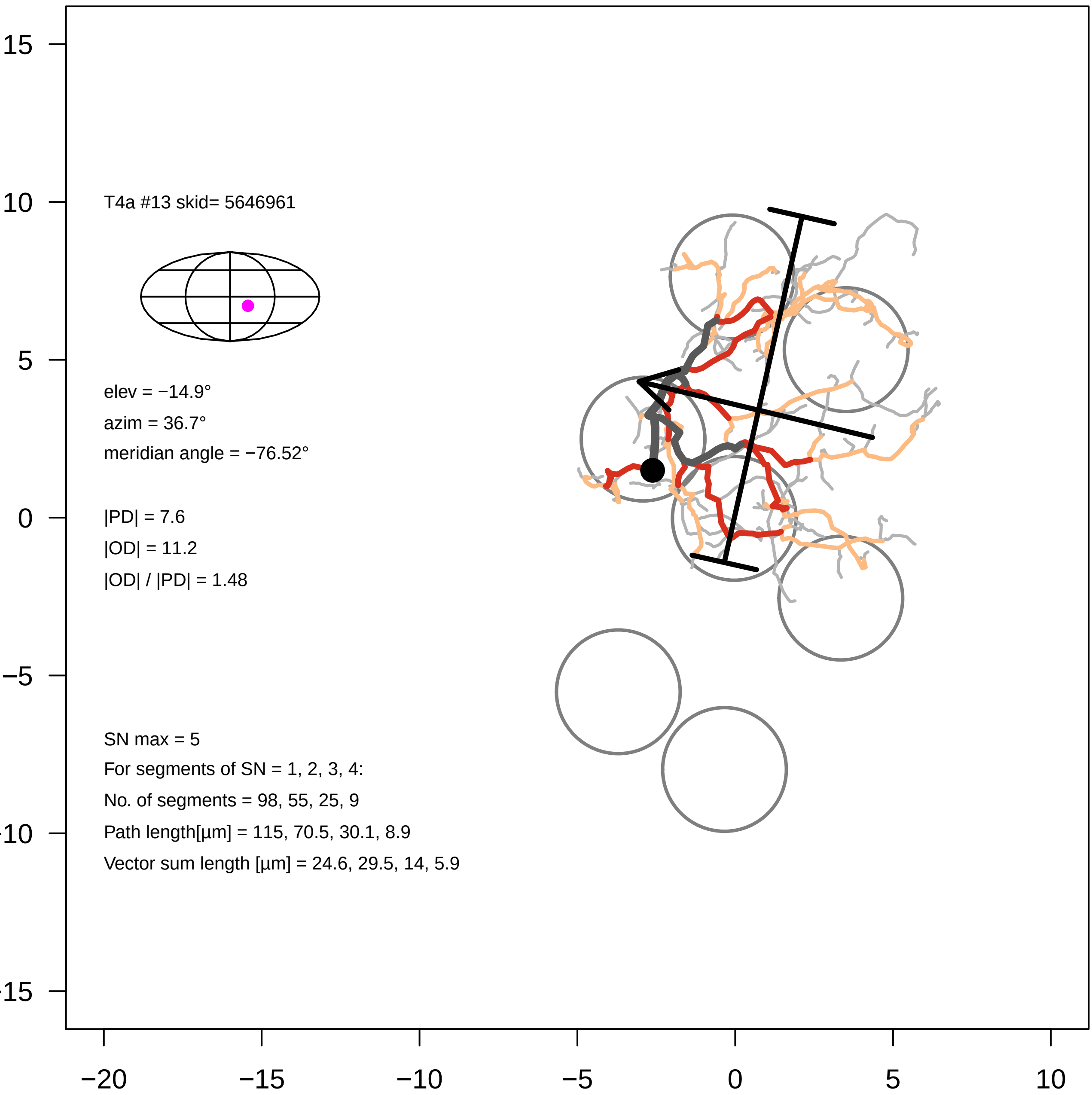
T4a #13 skid= 5646961
elev = −14.9°
azim = 36.7°
meridian angle = −76.52°
|PD| = 7.6
|OD| = 11.2
|OD| / |PD| = 1.48
SN max = 5
For segments of SN = 1, 2, 3, 4:
No. of segments = 98, 55, 25, 9
Path length[µm] = 115, 70.5, 30.1, 8.9
Vector sum length [µm] = 24.6, 29.5, 14, 5.9
15
10
5
0
−5
−10
−15
−20
−15
−10
−5
0
5
10

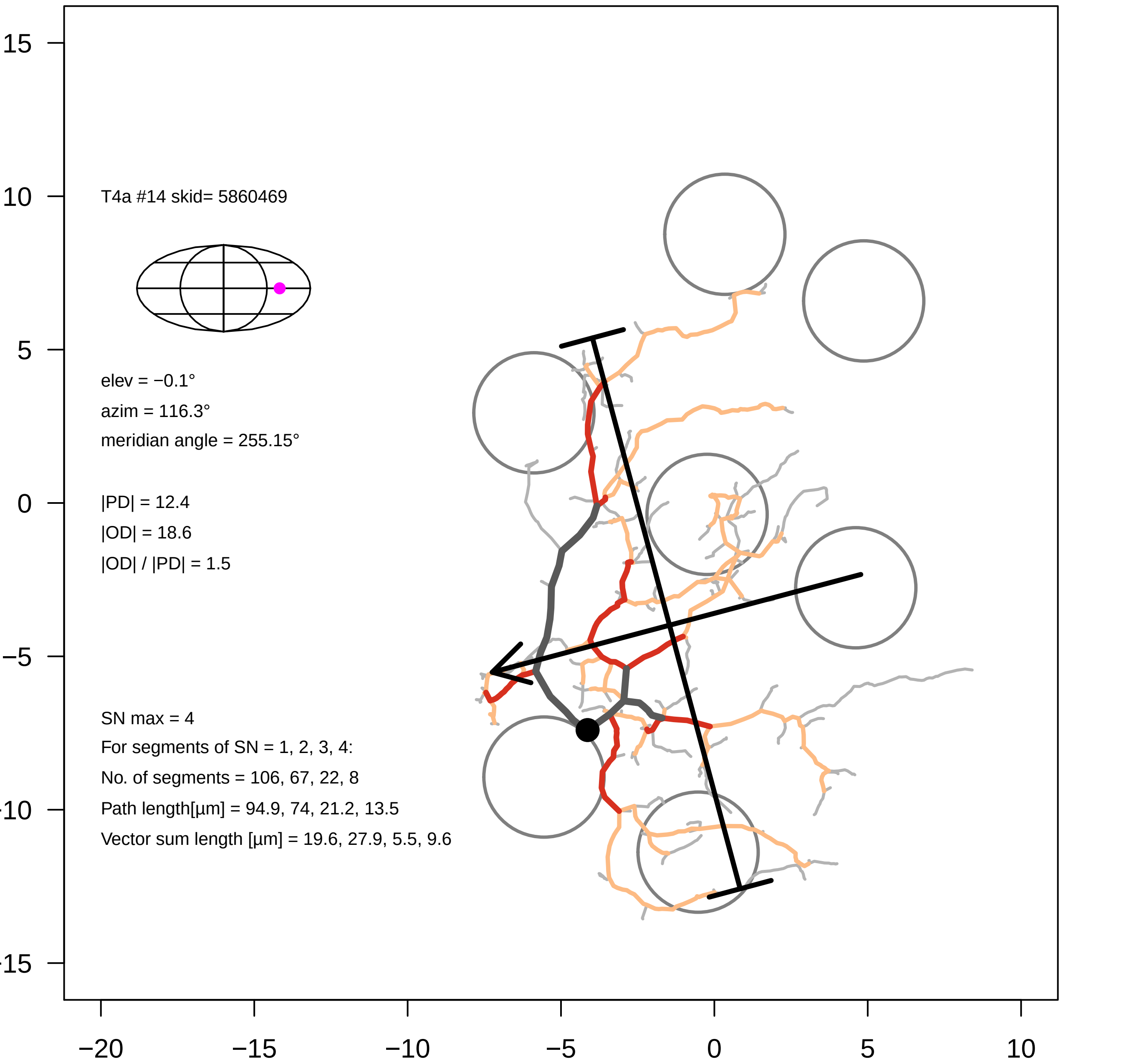

T4a #14 skid= 5860469
elev = −0.1°
azim = 116.3°
meridian angle = 255.15°
|PD| = 12.4
|OD| = 18.6
|OD| / |PD| = 1.5
SN max = 4
For segments of SN = 1, 2, 3, 4:
No. of segments = 106, 67, 22, 8
Path length[µm] = 94.9, 74, 21.2, 13.5
Vector sum length [µm] = 19.6, 27.9, 5.5, 9.6
15
10
5
0
−5
−10
−15
−20
−15
−10
−5
0
5
10

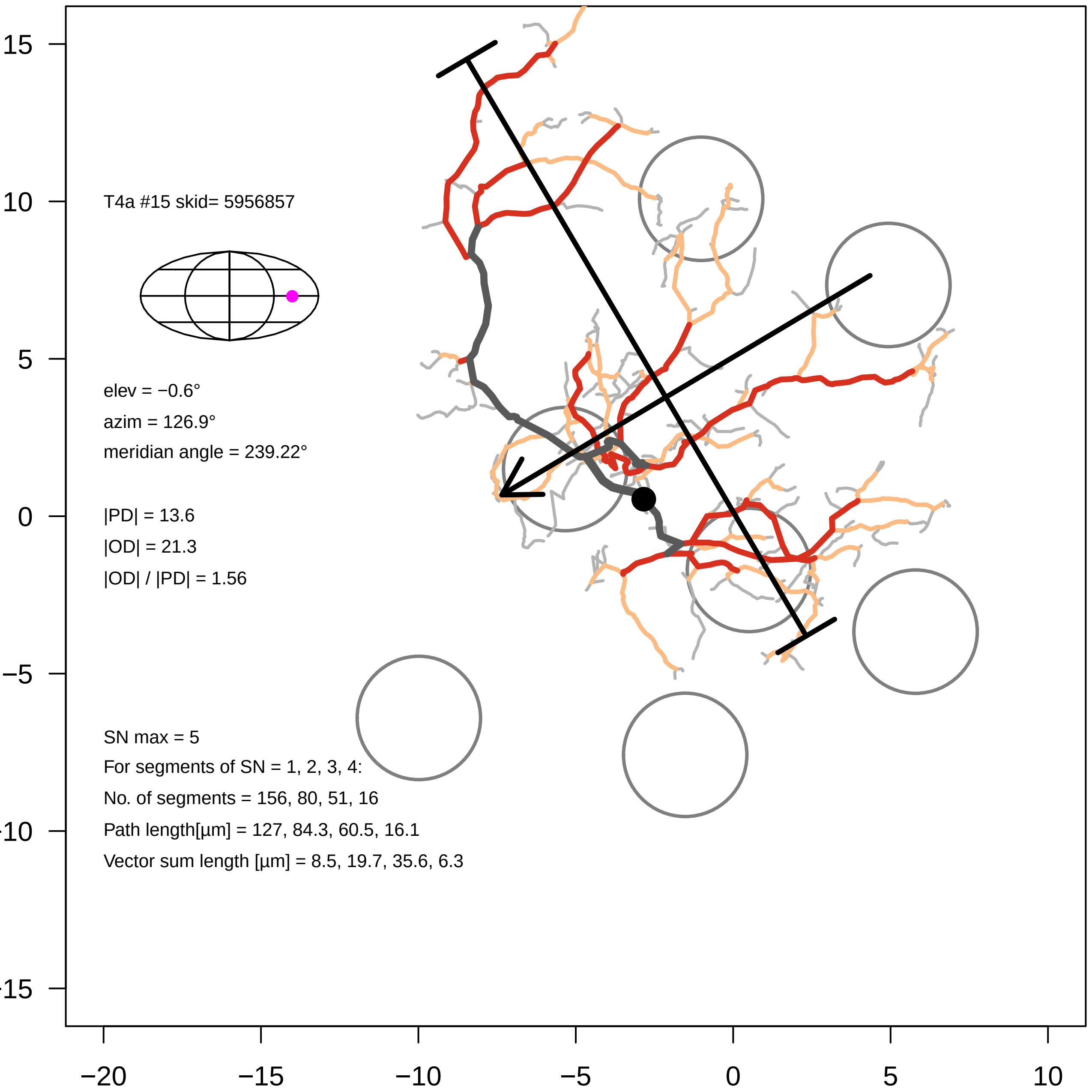
T4a #15 skid= 5956857
elev = −0.6°
azim = 126.9°
meridian angle = 239.22°
|PD| = 13.6
|OD| = 21.3
|OD| / |PD| = 1.56
SN max = 5
For segments of SN = 1, 2, 3, 4:
No. of segments = 156, 80, 51, 16
Path length[µm] = 127, 84.3, 60.5, 16.1
Vector sum length [µm] = 8.5, 19.7, 35.6, 6.3
15
10
5
0
−5
−10
−15
−20
−15
−10
−5
0
5
10

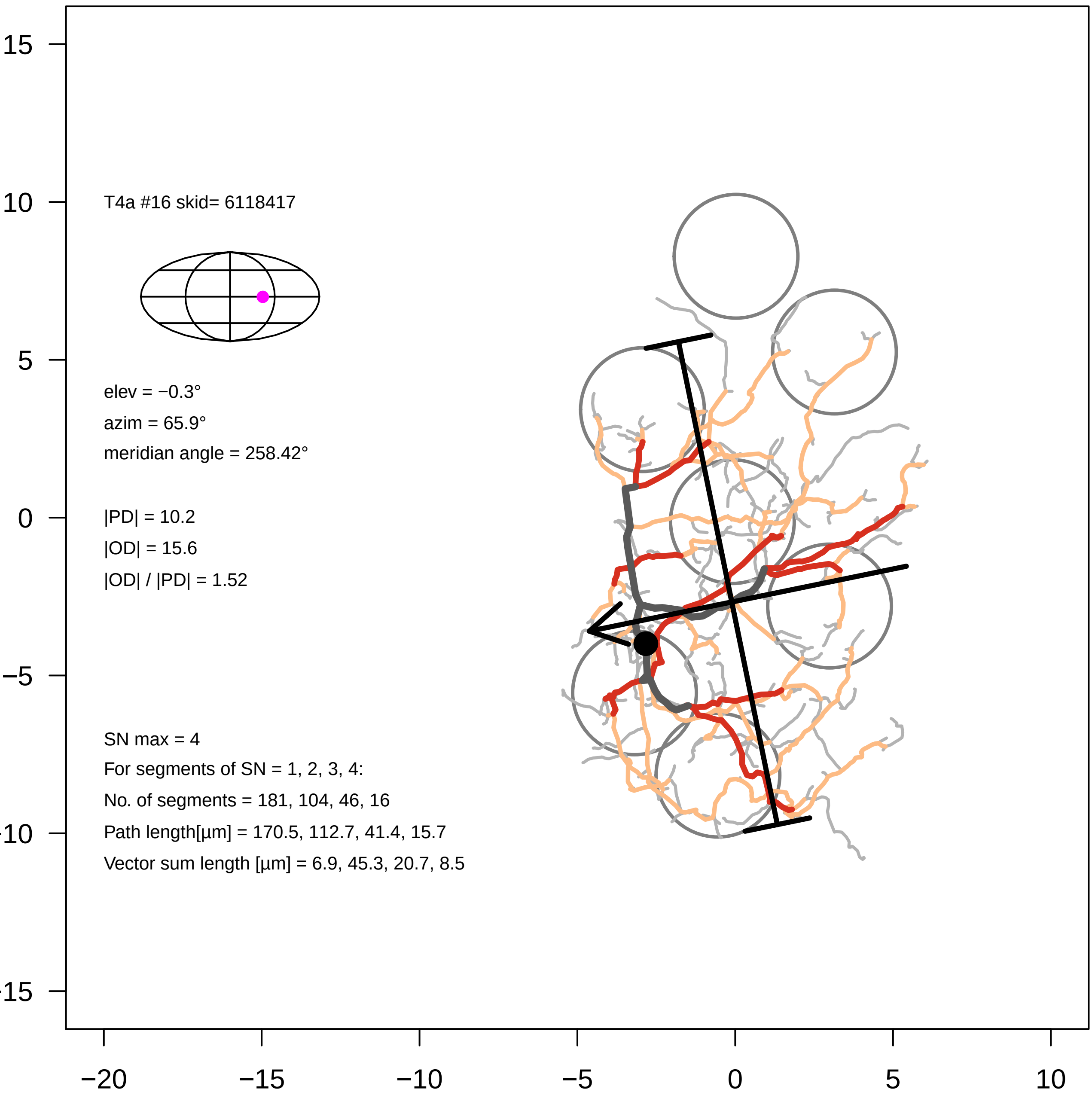

T4a #16 skid= 6118417
elev = −0.3°
azim = 65.9°
meridian angle = 258.42°
|PD| = 10.2
|OD| = 15.6
|OD| / |PD| = 1.52
SN max = 4
For segments of SN = 1, 2, 3, 4:
No. of segments = 181, 104, 46, 16
Path length[µm] = 170.5, 112.7, 41.4, 15.7
Vector sum length [µm] = 6.9, 45.3, 20.7, 8.5
15
10
5
0
−5
−10
−15
−20
−15
−10
−5
0
5
10

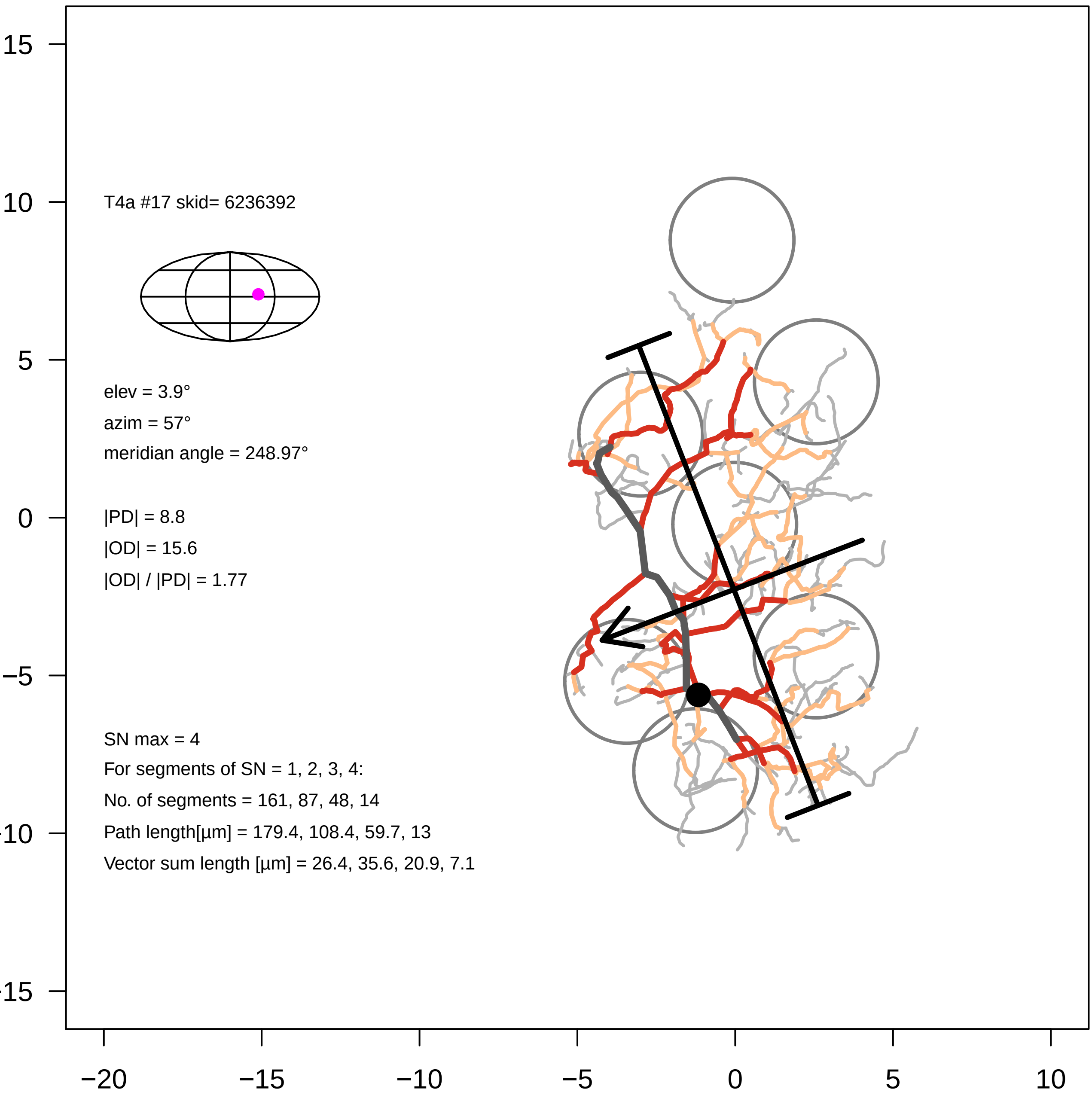

T4a #17 skid= 6236392
elev = 3.9°
azim = 57°
meridian angle = 248.97°
|PD| = 8.8
|OD| = 15.6
|OD| / |PD| = 1.77
SN max = 4
For segments of SN = 1, 2, 3, 4:
No. of segments = 161, 87, 48, 14
Path length[µm] = 179.4, 108.4, 59.7, 13
Vector sum length [µm] = 26.4, 35.6, 20.9, 7.1
15
10
5
0
−5
−10
−15
−20
−15
−10
−5
0
5
10

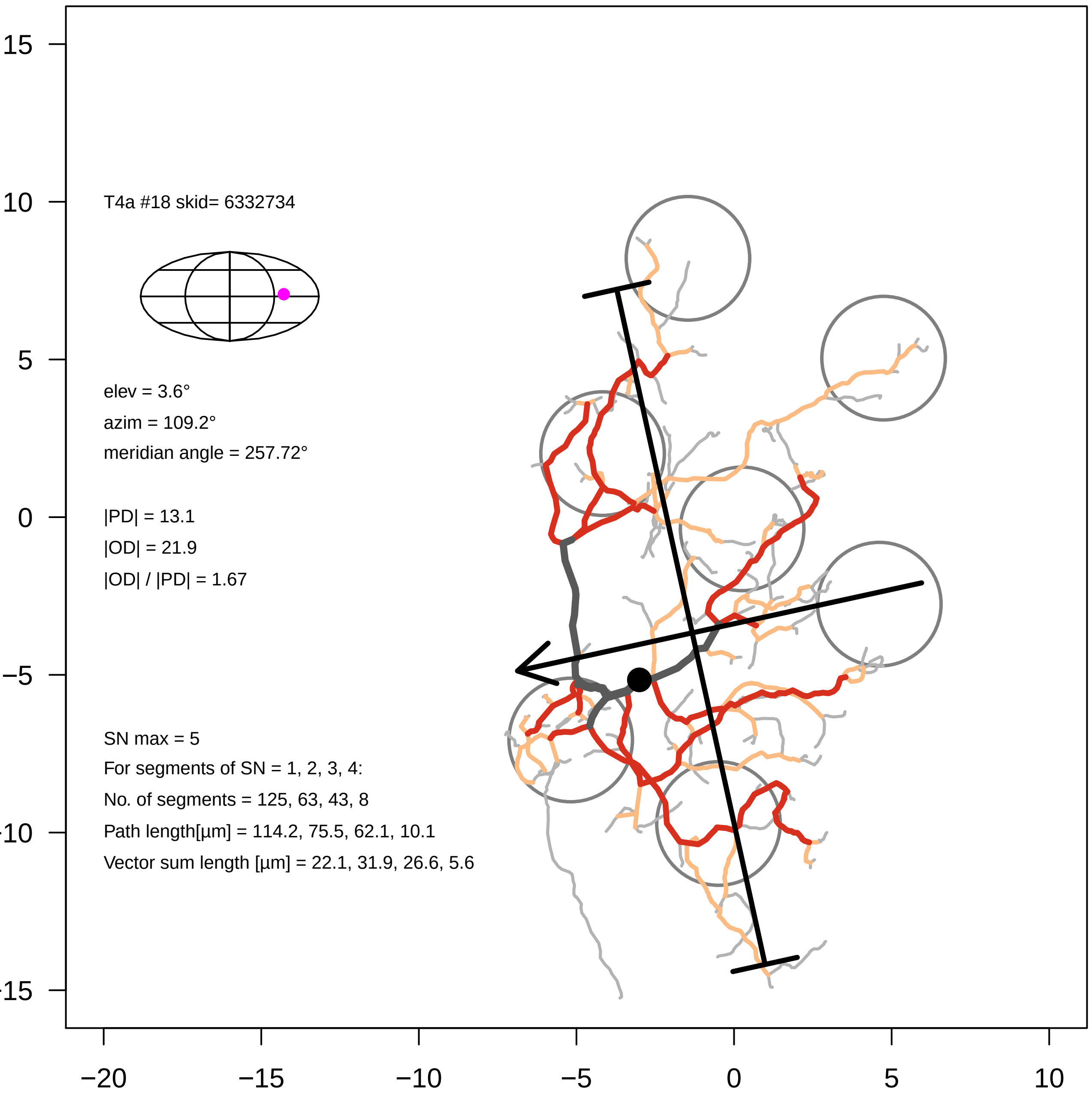

T4a #18 skid= 6332734
elev = 3.6°
azim = 109.2°
meridian angle = 257.72°
|PD| = 13.1
|OD| = 21.9
|OD| / |PD| = 1.67
SN max = 5
For segments of SN = 1, 2, 3, 4:
No. of segments = 125, 63, 43, 8
Path length[µm] = 114.2, 75.5, 62.1, 10.1
Vector sum length [µm] = 22.1, 31.9, 26.6, 5.6
15
10
5
0
−5
−10
−15
−20
−15
−10
−5
0
5
10

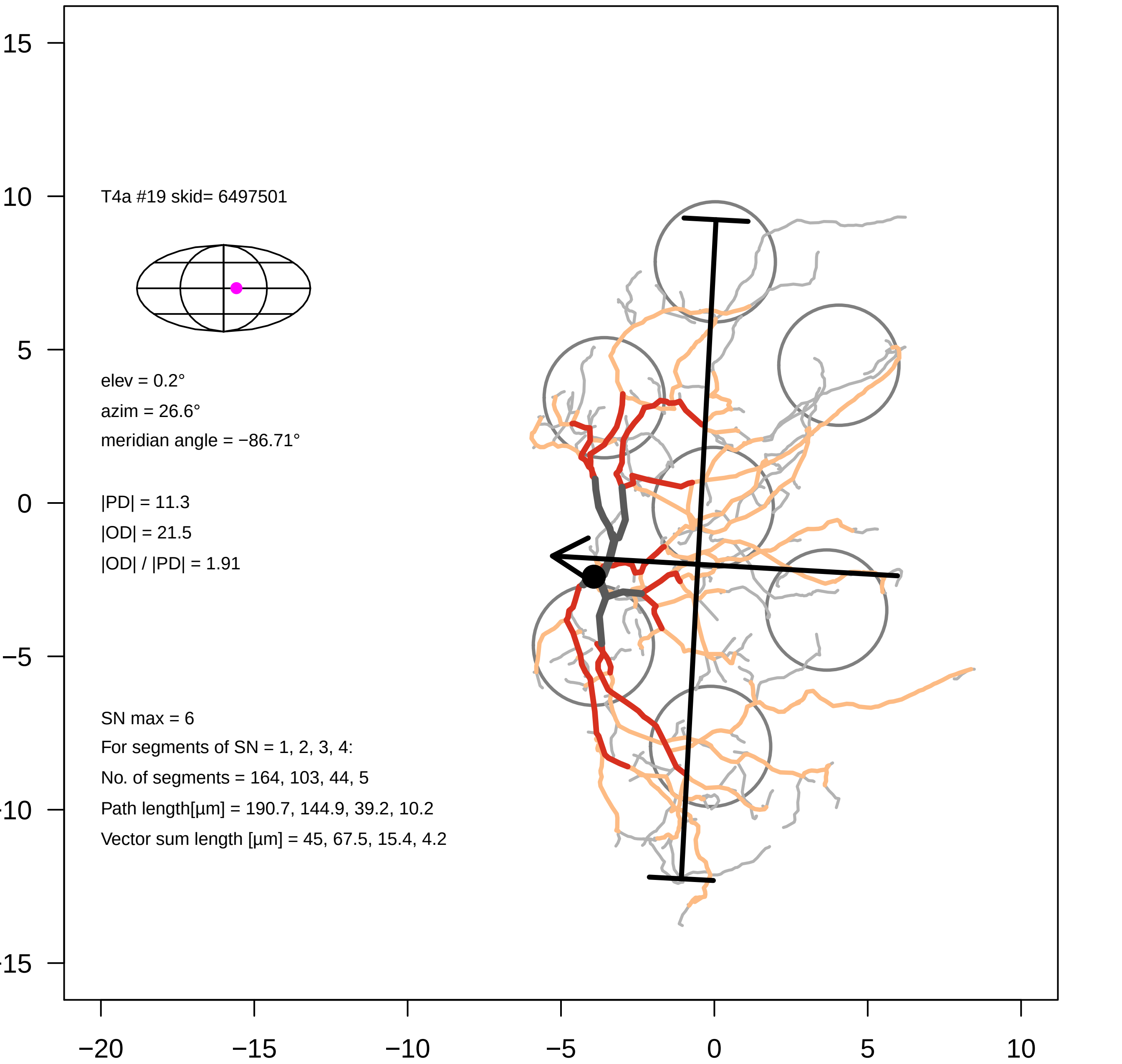
T4a #19 skid= 6497501
elev = 0.2°
azim = 26.6°
meridian angle = −86.71°
|PD| = 11.3
|OD| = 21.5
|OD| / |PD| = 1.91
SN max = 6
For segments of SN = 1, 2, 3, 4:
No. of segments = 164, 103, 44, 5
Path length[µm] = 190.7, 144.9, 39.2, 10.2
Vector sum length [µm] = 45, 67.5, 15.4, 4.2
15
10
5
0
−5
−20
−15
−10
−5
0
5
10

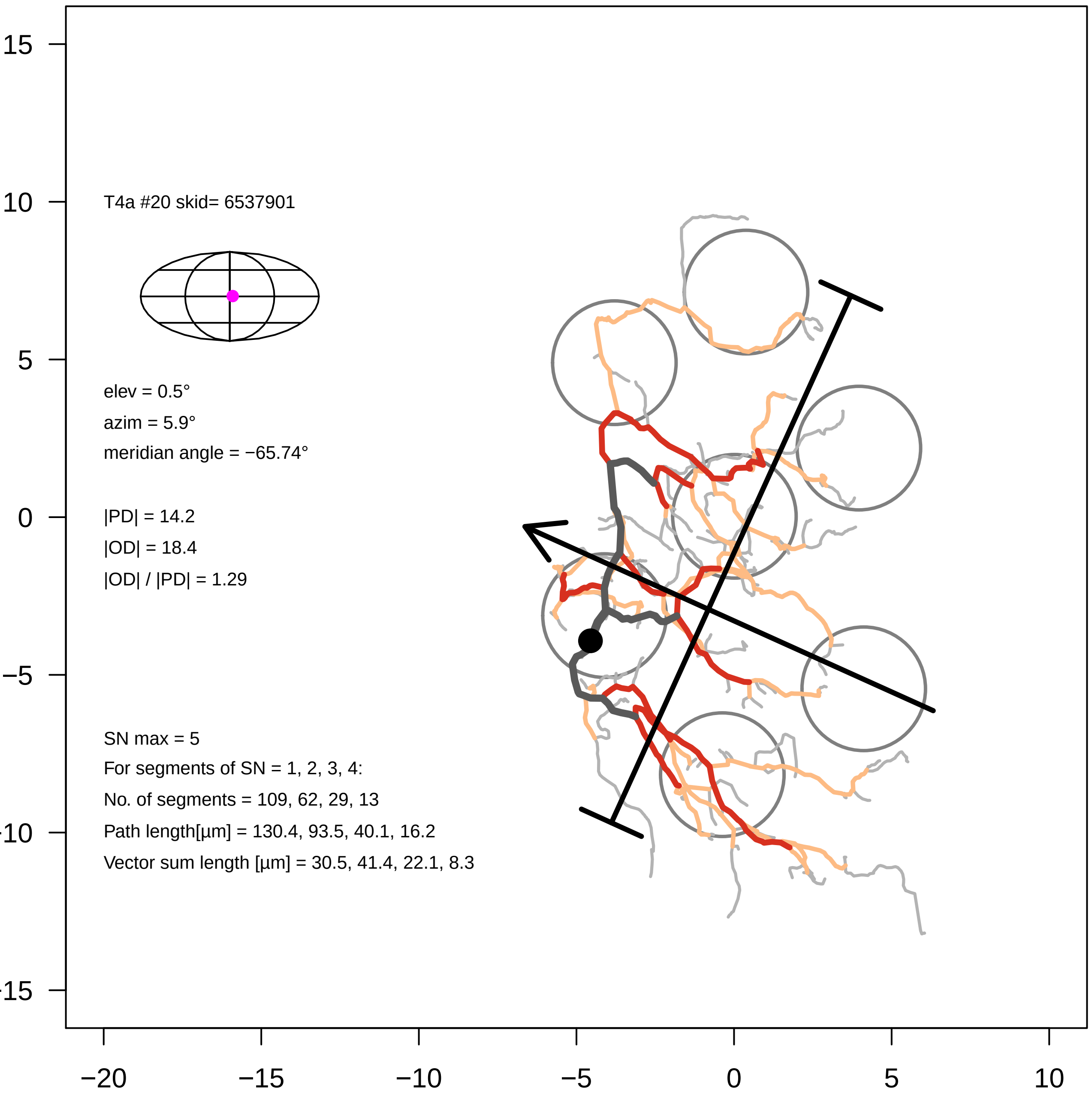

T4a #20 skid= 6537901
elev = 0.5°
azim = 5.9°
meridian angle = −65.74°
|PD| = 14.2
|OD| = 18.4
|OD| / |PD| = 1.29
SN max = 5
For segments of SN = 1, 2, 3, 4:
No. of segments = 109, 62, 29, 13
Path length[µm] = 130.4, 93.5, 40.1, 16.2
Vector sum length [µm] = 30.5, 41.4, 22.1, 8.3
15
10
5
0
−5
−10
−15
−20
−15
−10
−5
0
5
10

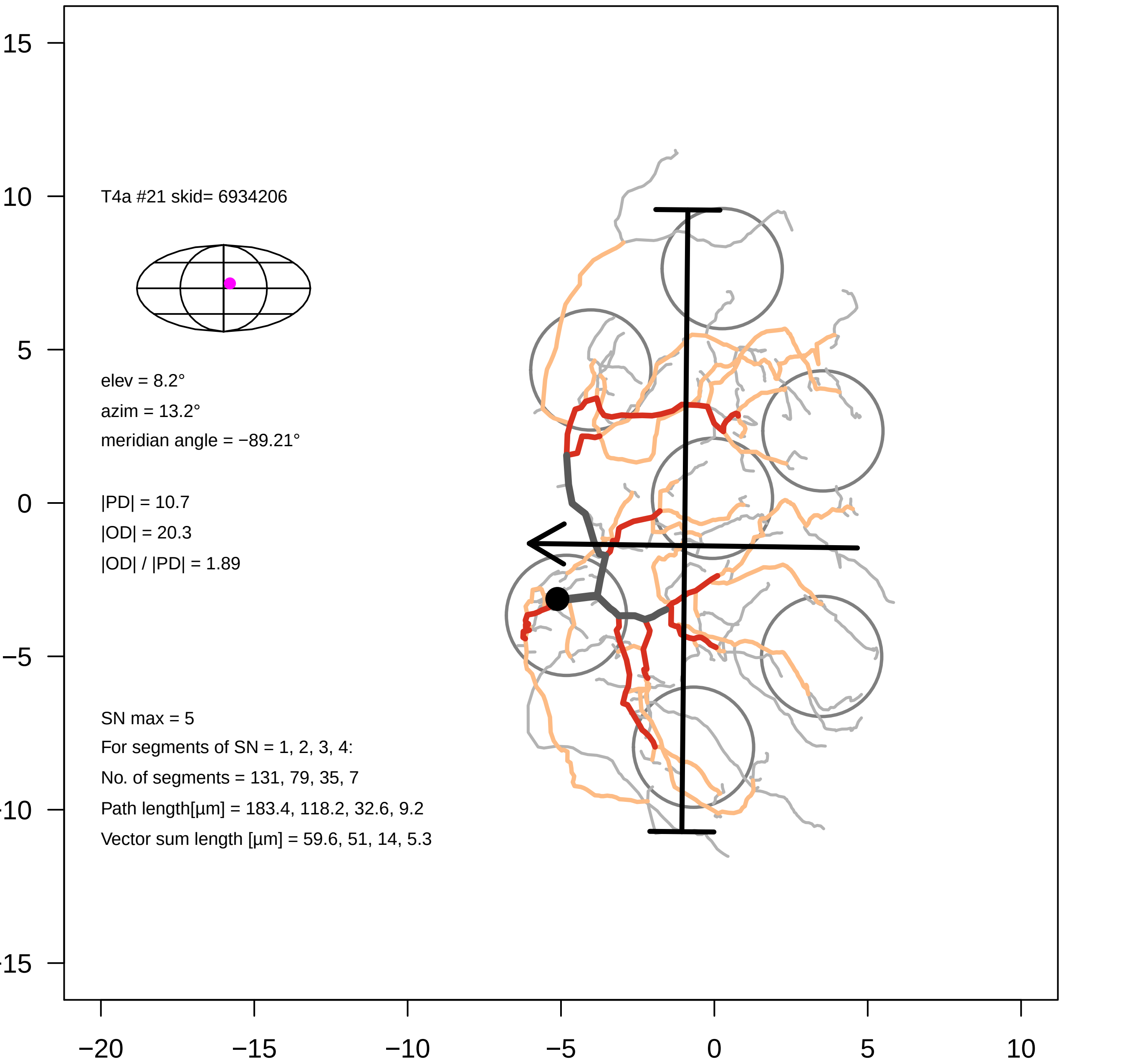
T4a #21 skid= 6934206
elev = 8.2°
azim = 13.2°
meridian angle = −89.21°
|PD| = 10.7
|OD| = 20.3
|OD| / |PD| = 1.89
SN max = 5
For segments of SN = 1, 2, 3, 4:
No. of segments = 131, 79, 35, 7
Path length[µm] = 183.4, 118.2, 32.6, 9.2
Vector sum length [µm] = 59.6, 51, 14, 5.3
15
10
5
0
−5
−10
−15
−20
−15
−10
−5
0
5
10

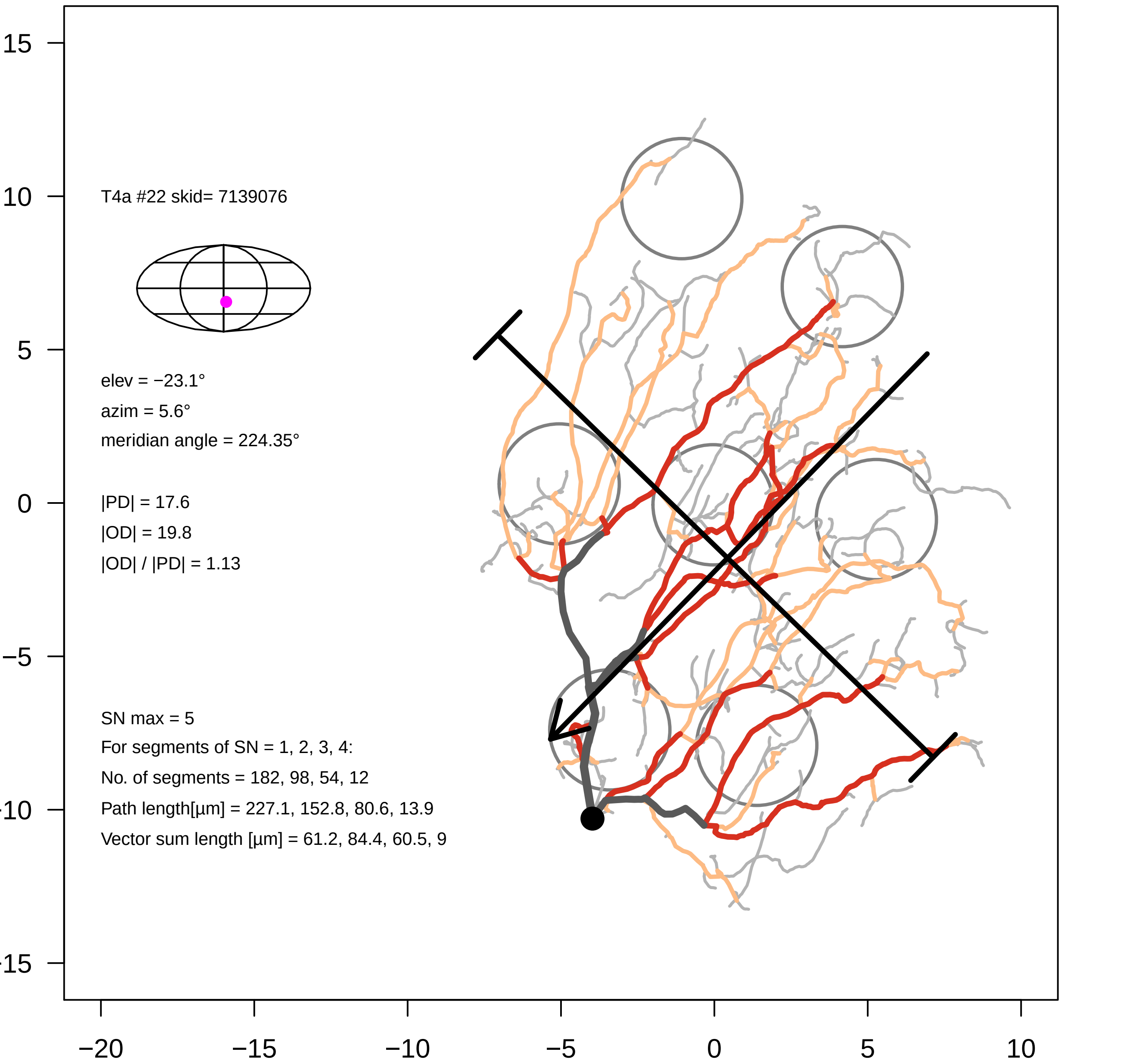
T4a #22 skid= 7139076
elev = −23.1°
azim = 5.6°
meridian angle = 224.35°
|PD| = 17.6
|OD| = 19.8
|OD| / |PD| = 1.13
SN max = 5
For segments of SN = 1, 2, 3, 4:
No. of segments = 182, 98, 54, 12
Path length[µm] = 227.1, 152.8, 80.6, 13.9
Vector sum length [µm] = 61.2, 84.4, 60.5, 9
15
10
5
0
−5
−10
−15
−20
−15
−10
−5
0
5
10

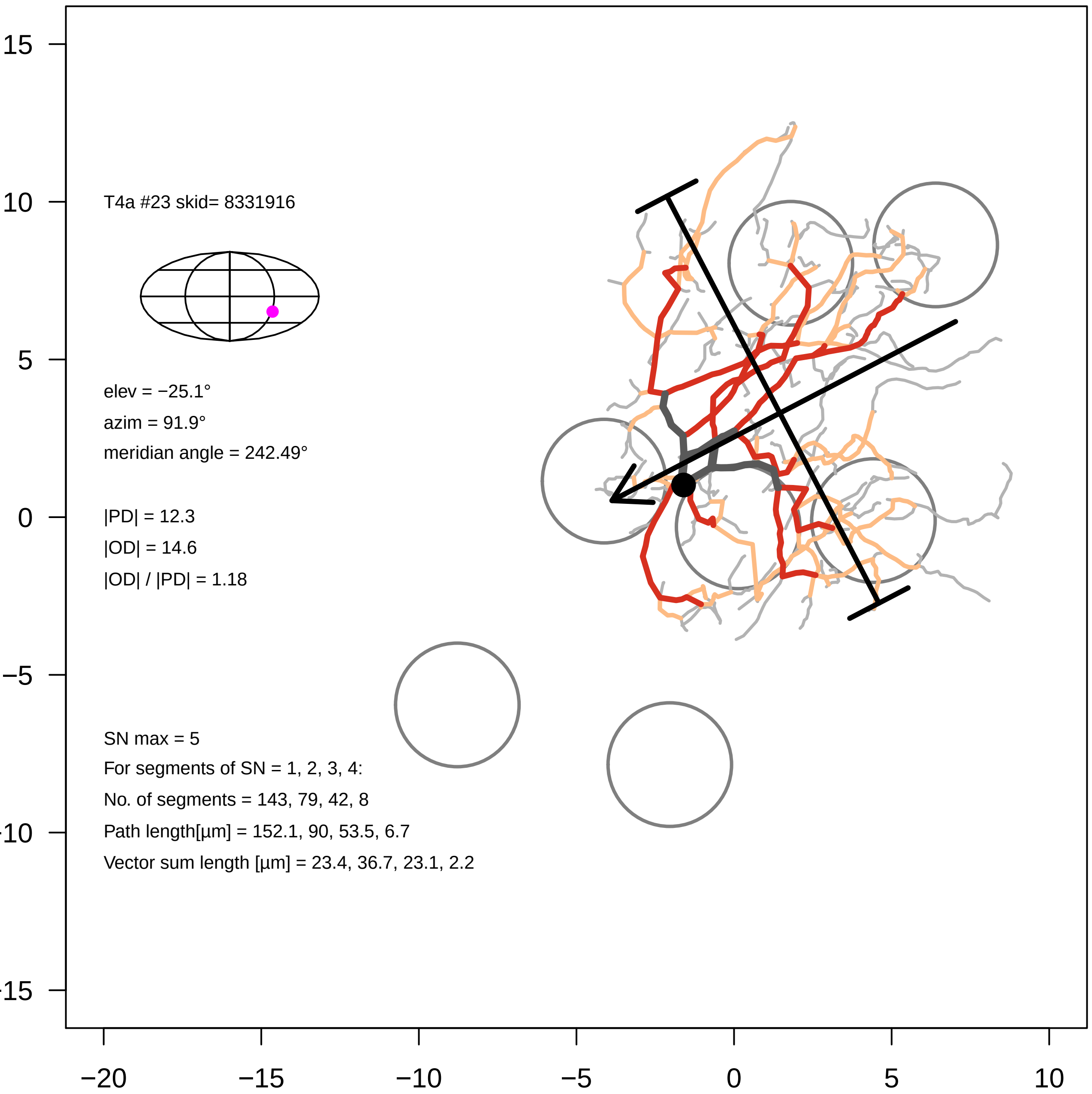
T4a #23 skid= 8331916
elev = −25.1°
azim = 91.9°
meridian angle = 242.49°
|PD| = 12.3
|OD| = 14.6
|OD| / |PD| = 1.18
SN max = 5
For segments of SN = 1, 2, 3, 4:
No. of segments = 143, 79, 42, 8
Path length[µm] = 152.1, 90, 53.5, 6.7
Vector sum length [µm] = 23.4, 36.7, 23.1, 2.2
15
10
5
0
−5
−10
−15
−20
−15
−10
−5
0
5
10

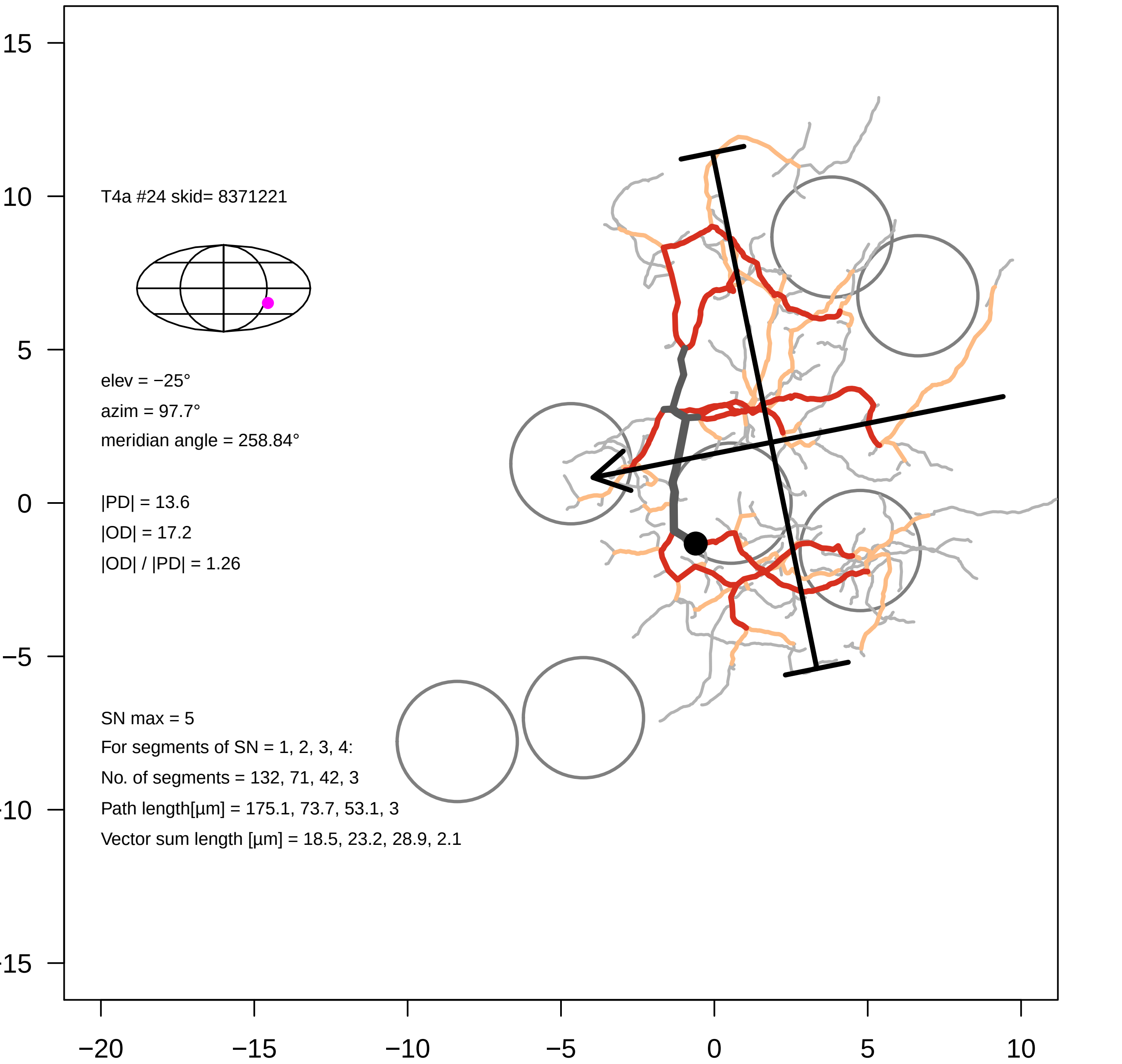
T4a #24 skid= 8371221
elev = −25°
azim = 97.7°
meridian angle = 258.84°
|PD| = 13.6
|OD| = 17.2
|OD| / |PD| = 1.26
SN max = 5
For segments of SN = 1, 2, 3, 4:
No. of segments = 132, 71, 42, 3
Path length[µm] = 175.1, 73.7, 53.1, 3
Vector sum length [µm] = 18.5, 23.2, 28.9, 2.1
15
10
5
0
−5
−10
−15
−20
−15
−10
−5
0
5
10

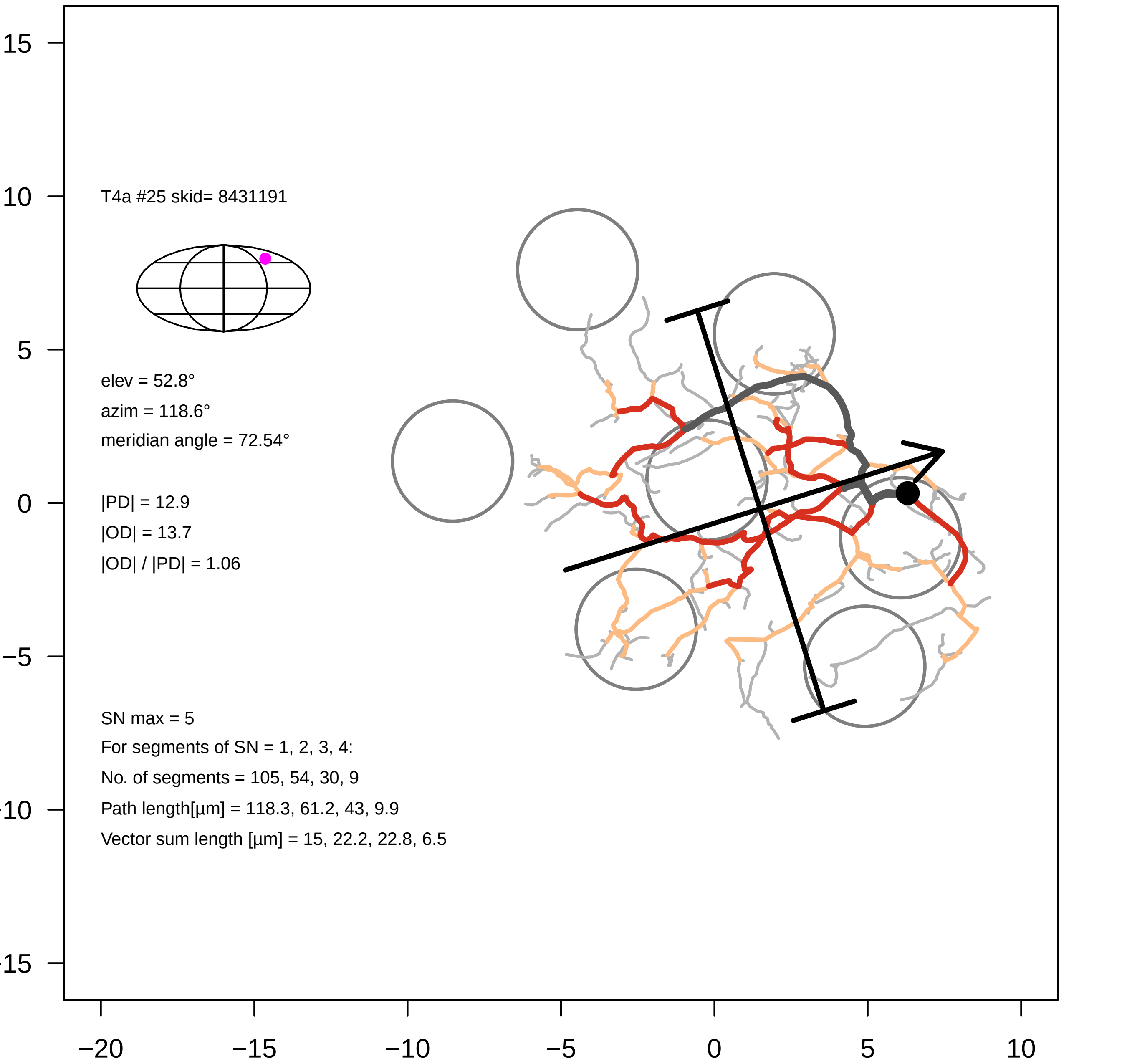

T4a #25 skid= 8431191
elev = 52.8°
azim = 118.6°
meridian angle = 72.54°
|PD| = 12.9
|OD| = 13.7
|OD| / |PD| = 1.06
SN max = 5
For segments of SN = 1, 2, 3, 4:
No. of segments = 105, 54, 30, 9
Path length[µm] = 118.3, 61.2, 43, 9.9
Vector sum length [µm] = 15, 22.2, 22.8, 6.5
15
10
5
0
−5
−10
−15
−20
−15
−10
−5
0
5
10

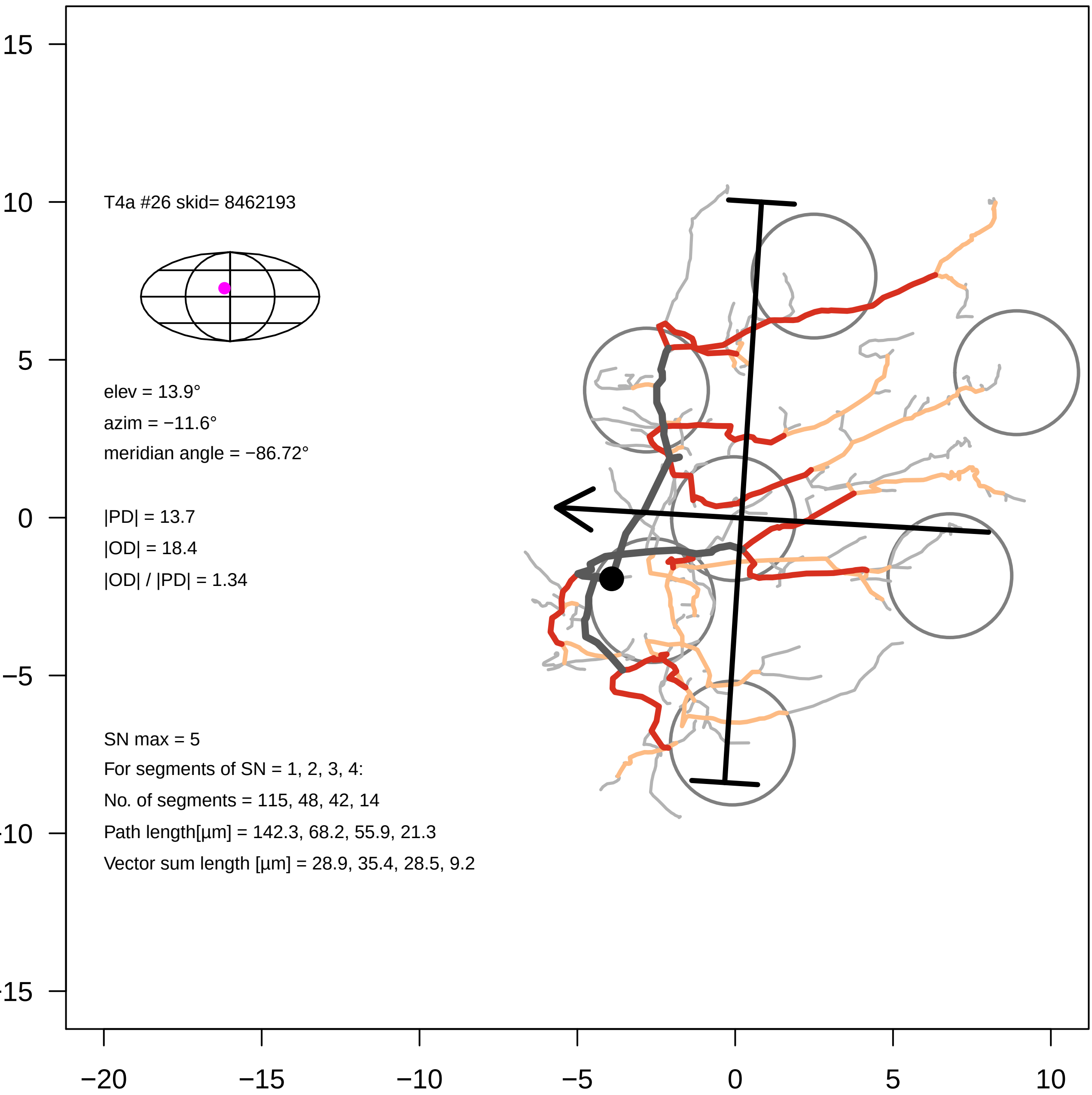

T4a #26 skid= 8462193
elev = 13.9°
azim = −11.6°
meridian angle = −86.72°
|PD| = 13.7
|OD| = 18.4
|OD| / |PD| = 1.34
SN max = 5
For segments of SN = 1, 2, 3, 4:
No. of segments = 115, 48, 42, 14
Path length[µm] = 142.3, 68.2, 55.9, 21.3
Vector sum length [µm] = 28.9, 35.4, 28.5, 9.2
15
10
5
0
−5
−10
−15
−20
−15
−10
−5
0
5
10

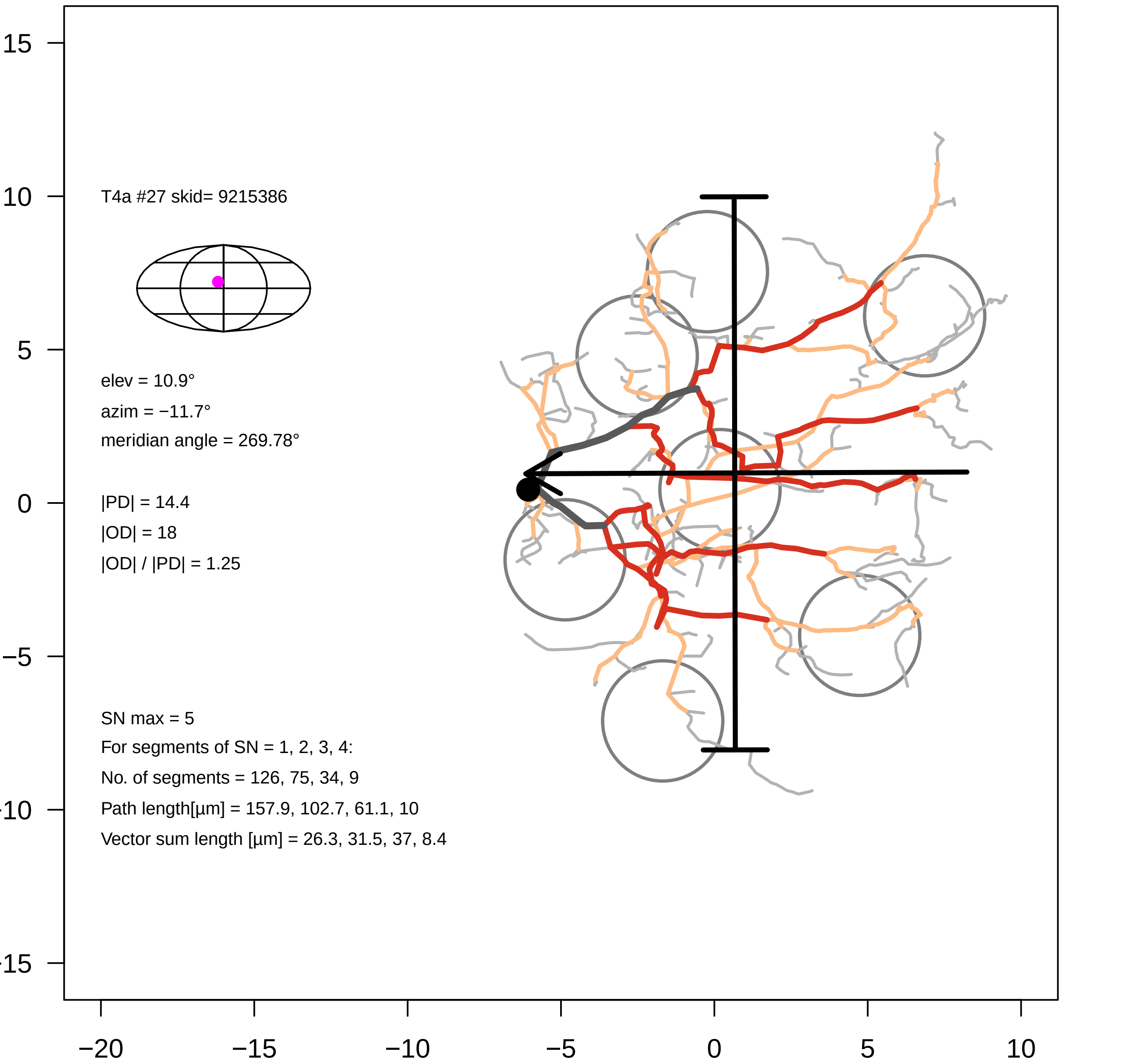

T4a #27 skid= 9215386
elev = 10.9°
azim = −11.7°
meridian angle = 269.78°
|PD| = 14.4
|OD| = 18
|OD| / |PD| = 1.25
SN max = 5
For segments of SN = 1, 2, 3, 4:
No. of segments = 126, 75, 34, 9
Path length[µm] = 157.9, 102.7, 61.1, 10
Vector sum length [µm] = 26.3, 31.5, 37, 8.4
15
10
5
0
−5
−10
−15
−20
−15
−10
−5
0
5
10

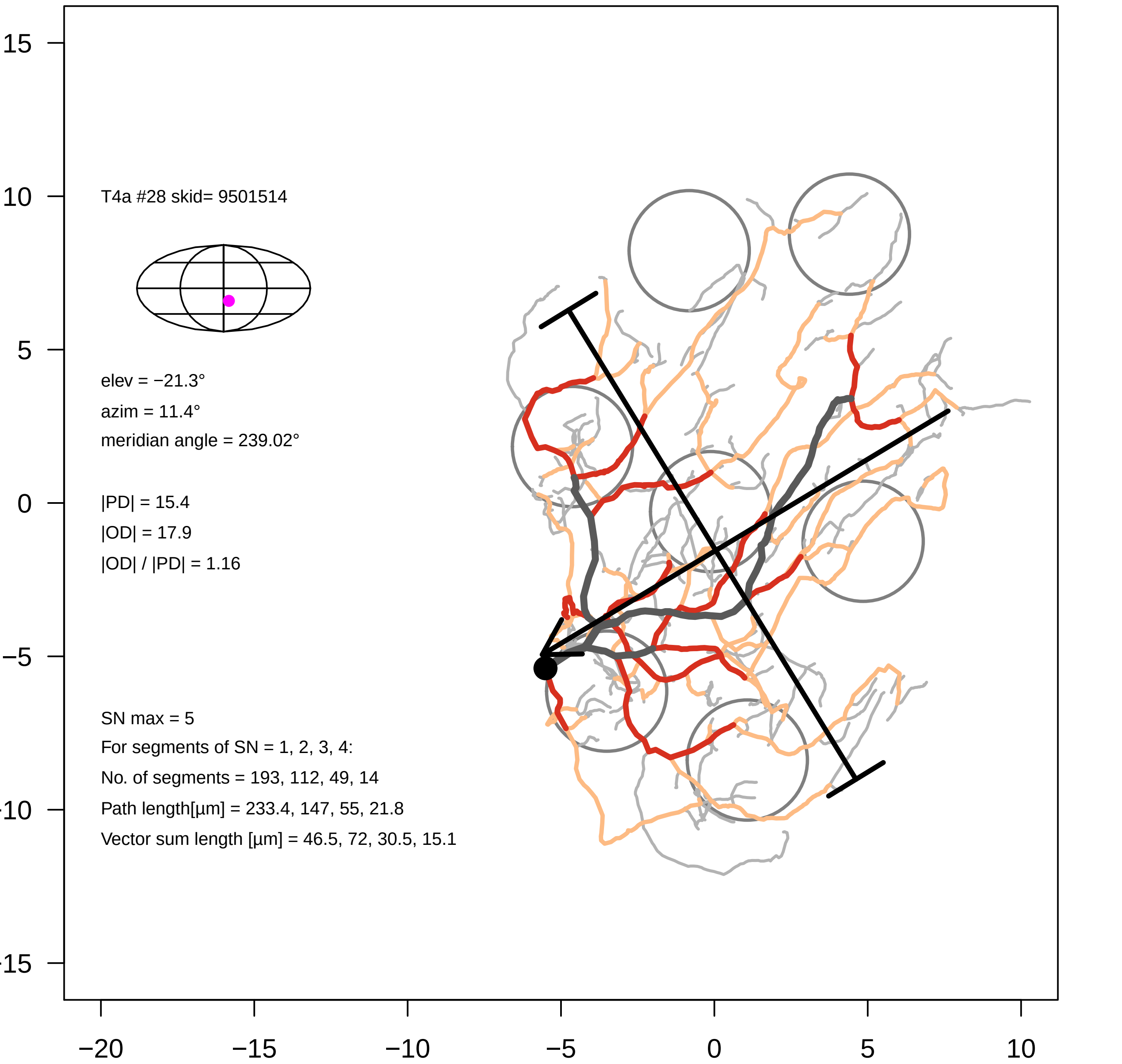
T4a #28 skid= 9501514
elev = −21.3°
azim = 11.4°
meridian angle = 239.02°
|PD| = 15.4
|OD| = 17.9
|OD| / |PD| = 1.16
SN max = 5
For segments of SN = 1, 2, 3, 4:
No. of segments = 193, 112, 49, 14
Path length[μm] = 233.4, 147, 55, 21.8
Vector sum length [μm] = 46.5, 72, 30.5, 15.1
15
10
5
0
−5
−10
−15
−20
−15
−10
−5
0
5
10

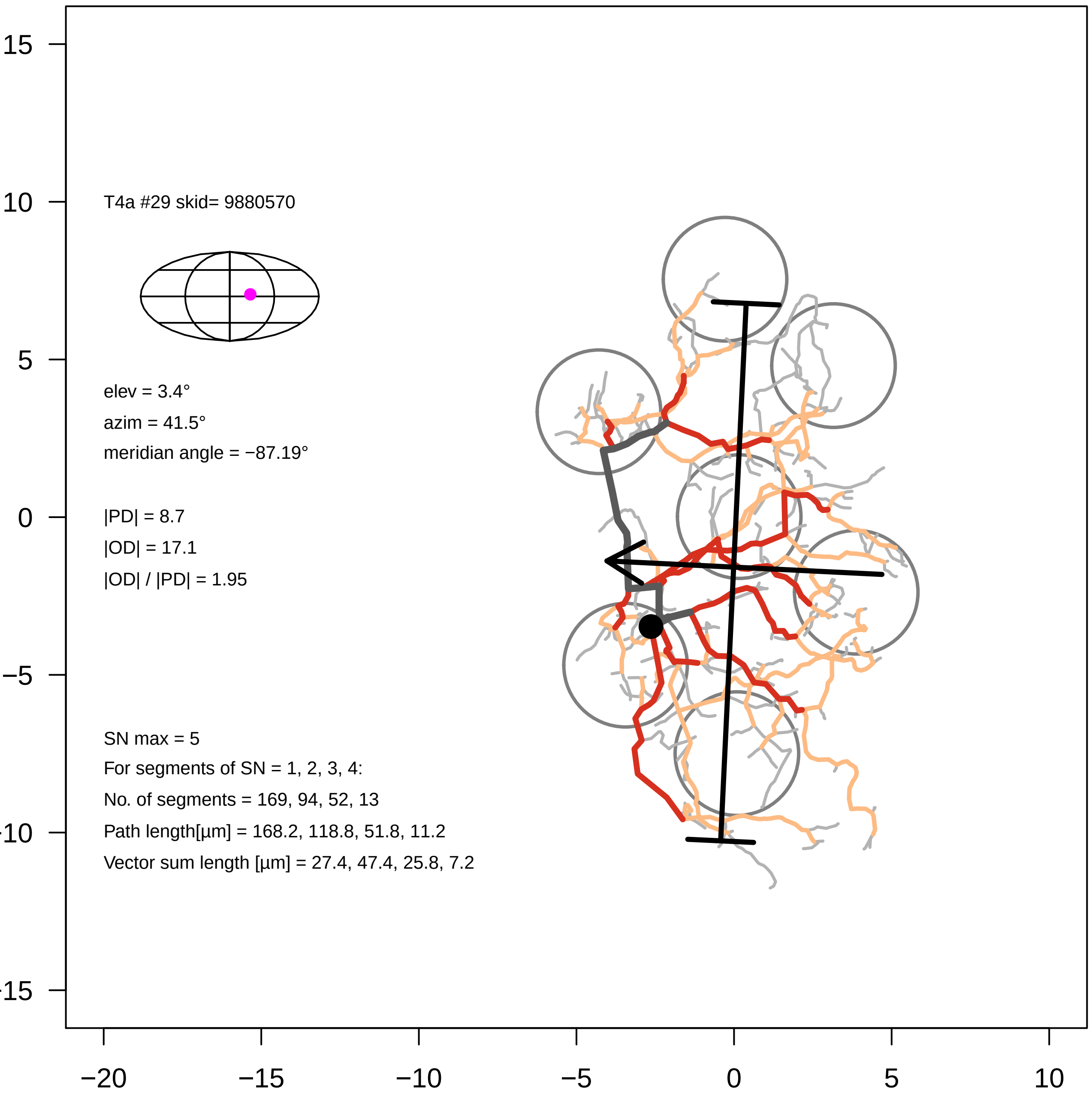

T4a #29 skid= 9880570
elev = 3.4°
azim = 41.5°
meridian angle = −87.19°
|PD| = 8.7
|OD| = 17.1
|OD| / |PD| = 1.95
SN max = 5
For segments of SN = 1, 2, 3, 4:
No. of segments = 169, 94, 52, 13
Path length[µm] = 168.2, 118.8, 51.8, 11.2
Vector sum length [µm] = 27.4, 47.4, 25.8, 7.2
15
10
5
0
−5
−10
−15
−20
−15
−10
−5
0
5
10

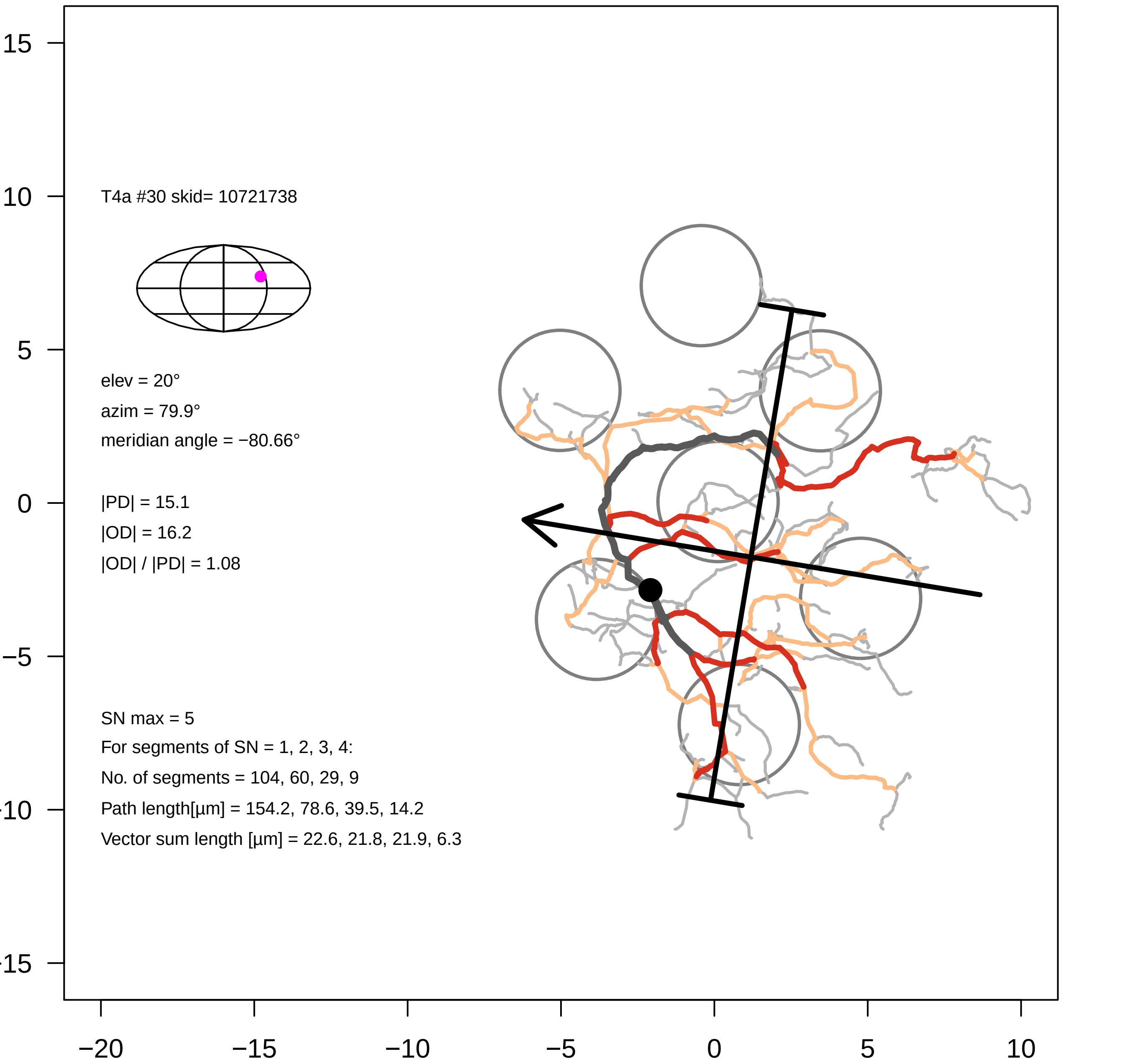

T4a #30 skid= 10721738
elev = 20°
azim = 79.9°
meridian angle = −80.66°
|PD| = 15.1
|OD| = 16.2
|OD| / |PD| = 1.08
SN max = 5
For segments of SN = 1, 2, 3, 4:
No. of segments = 104, 60, 29, 9
Path length[µm] = 154.2, 78.6, 39.5, 14.2
Vector sum length [µm] = 22.6, 21.8, 21.9, 6.3
15
10
5
0
−5
−10
−15
−20
−15
−10
−5
0
5
10

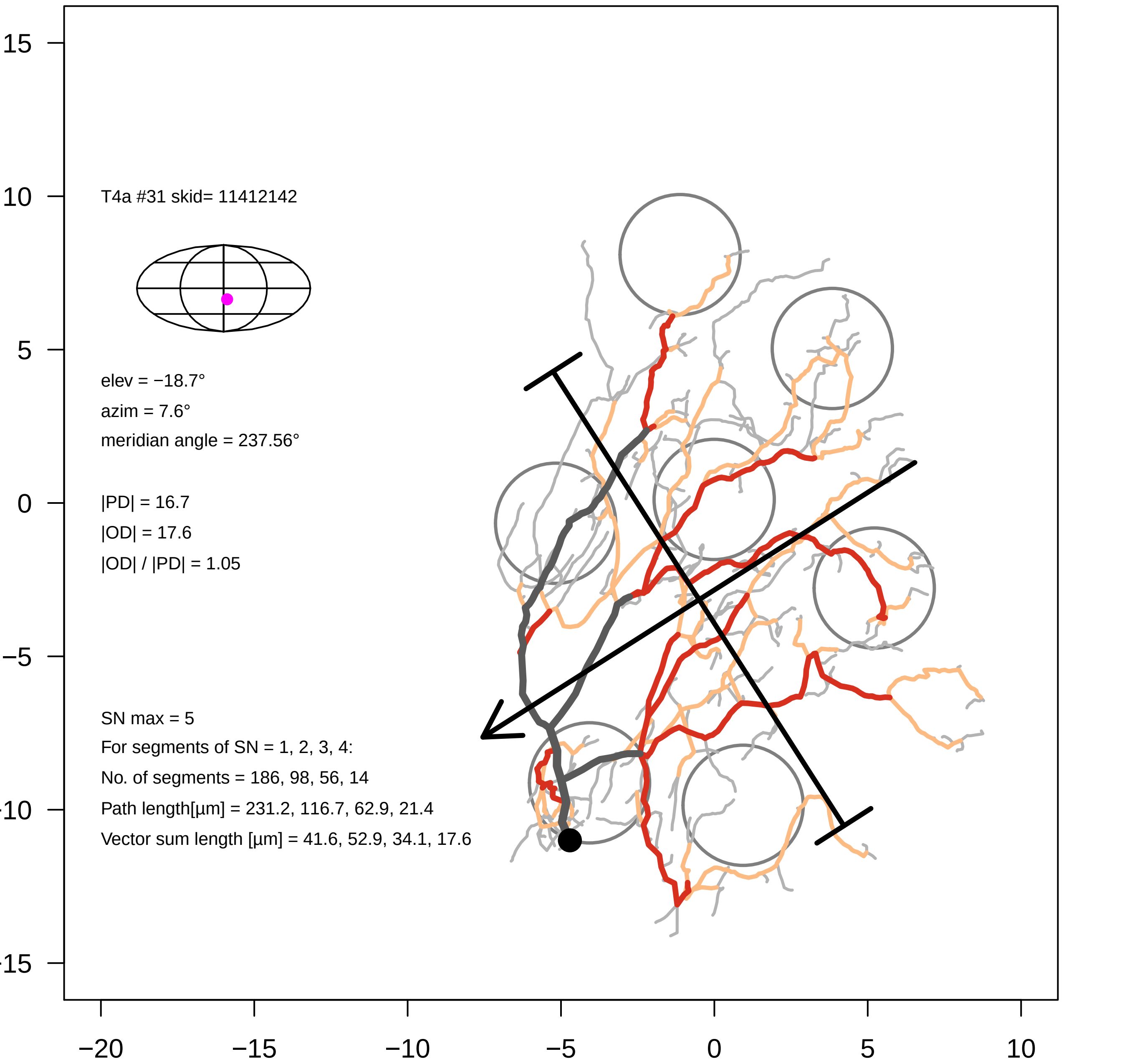
T4a #31 skid= 11412142
elev = −18.7°
azim = 7.6°
meridian angle = 237.56°
|PD| = 16.7
|OD| = 17.6
|OD| / |PD| = 1.05
SN max = 5
For segments of SN = 1, 2, 3, 4:
No. of segments = 186, 98, 56, 14
Path length[µm] = 231.2, 116.7, 62.9, 21.4
Vector sum length [µm] = 41.6, 52.9, 34.1, 17.6
15
10
5
0
−5
−10
−15
−20
−15
−10
−5
0
5
10

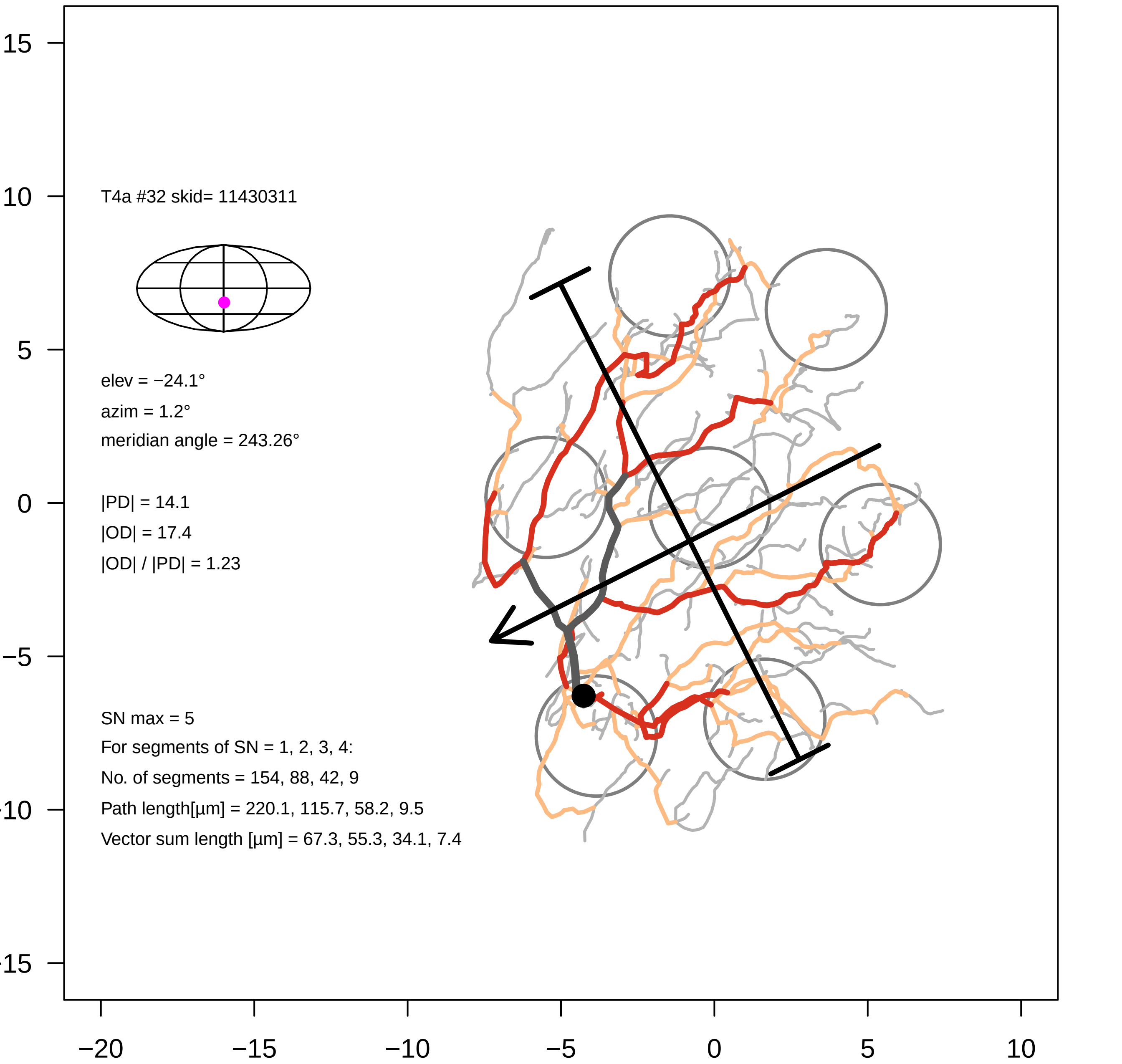

T4a #32 skid= 11430311
elev = −24.1°
azim = 1.2°
meridian angle = 243.26°
|PD| = 14.1
|OD| = 17.4
|OD| / |PD| = 1.23
SN max = 5
For segments of SN = 1, 2, 3, 4:
No. of segments = 154, 88, 42, 9
Path length[µm] = 220.1, 115.7, 58.2, 9.5
Vector sum length [µm] = 67.3, 55.3, 34.1, 7.4
15
10
5
0
−5
-10
-15
−20
−15
−10
−5
0
5
10

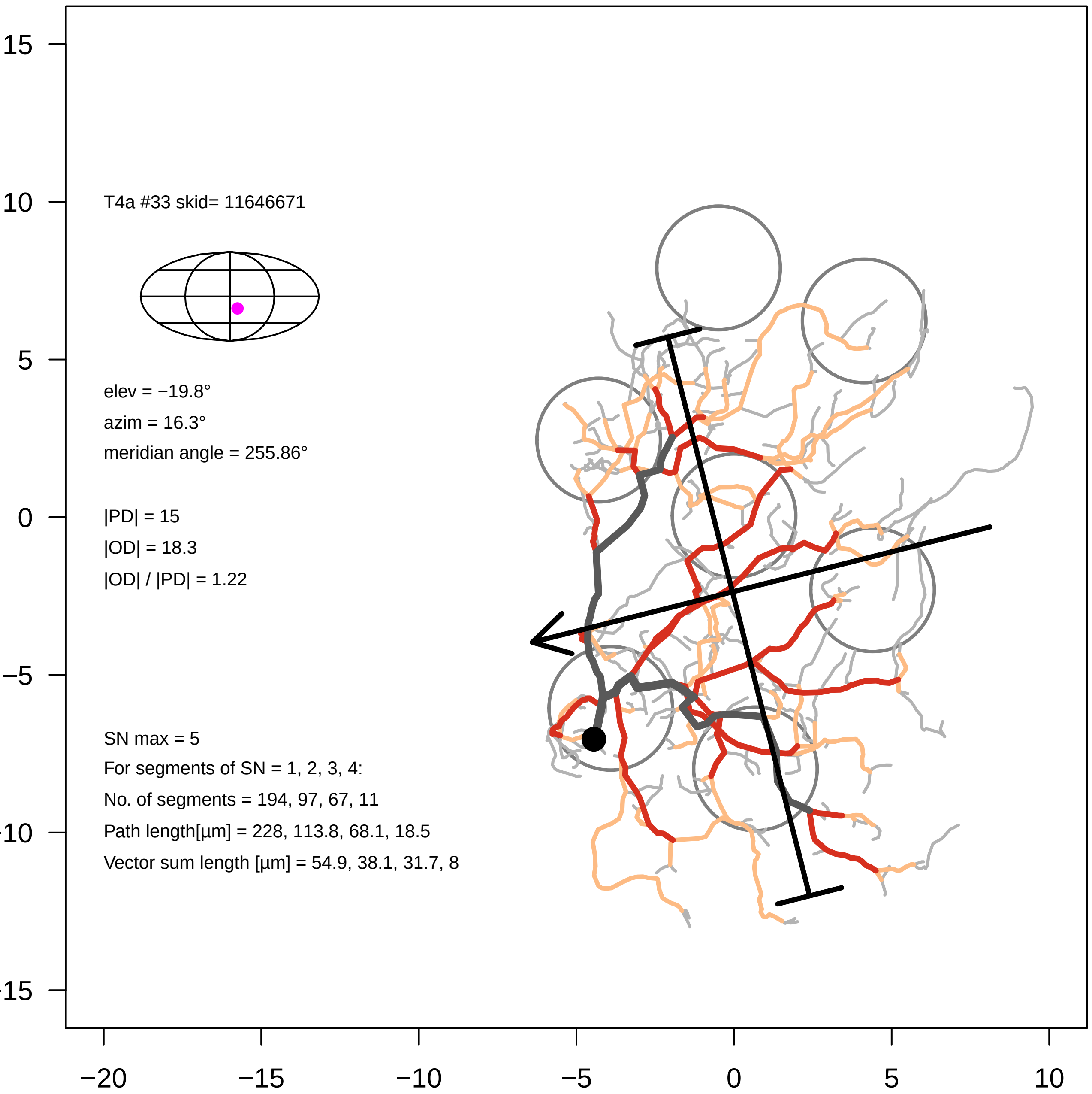
T4a #33 skid= 11646671
elev = −19.8°
azim = 16.3°
meridian angle = 255.86°
|PD| = 15
|OD| = 18.3
|OD| / |PD| = 1.22
SN max = 5
For segments of SN = 1, 2, 3, 4:
No. of segments = 194, 97, 67, 11
Path length[µm] = 228, 113.8, 68.1, 18.5
Vector sum length [µm] = 54.9, 38.1, 31.7, 8
15
10
5
0
−5
−10
−15
−20
−15
−10
−5
0
5
10

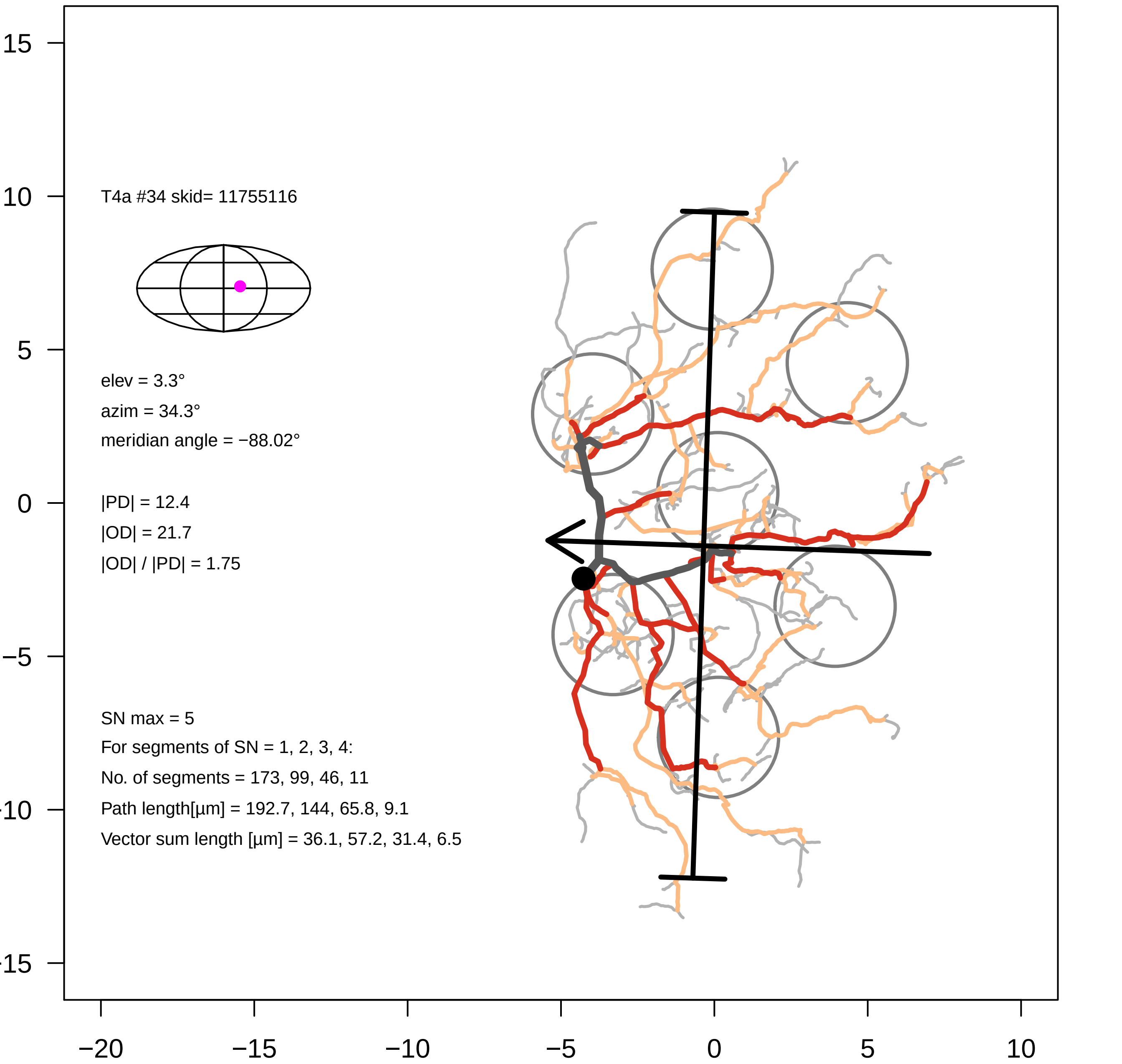
T4a #34 skid= 11755116
elev = 3.3°
azim = 34.3°
meridian angle = −88.02°
|PD| = 12.4
|OD| = 21.7
|OD| / |PD| = 1.75
SN max = 5
For segments of SN = 1, 2, 3, 4:
No. of segments = 173, 99, 46, 11
Path length[µm] = 192.7, 144, 65.8, 9.1
Vector sum length [µm] = 36.1, 57.2, 31.4, 6.5
15
10
5
0
−5
−10
−15
−20
−15
−10
−5
0
5
10

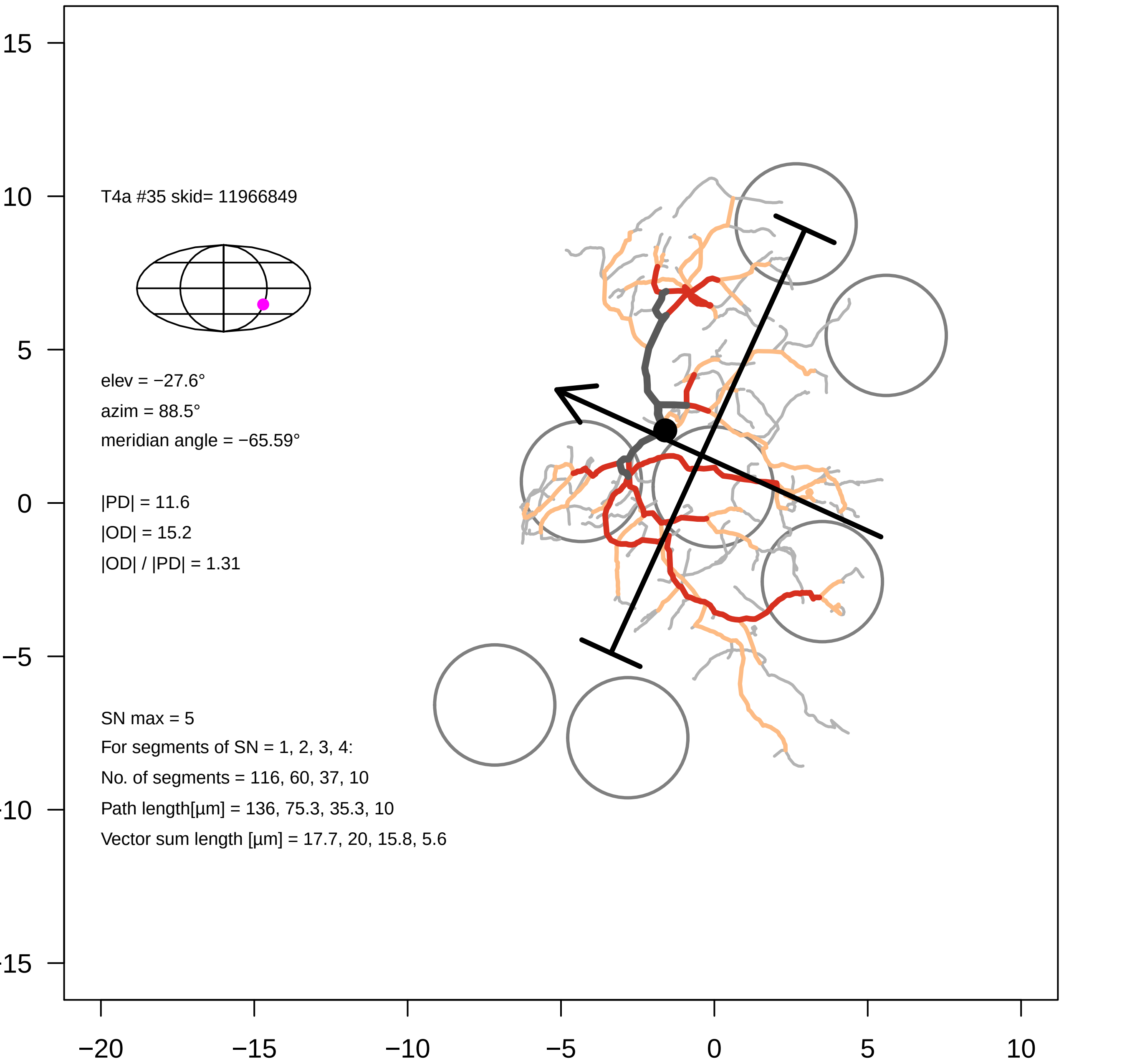
T4a #35 skid= 11966849
elev = −27.6°
azim = 88.5°
meridian angle = −65.59°
|PD| = 11.6
|OD| = 15.2
|OD| / |PD| = 1.31
SN max = 5
For segments of SN = 1, 2, 3, 4:
No. of segments = 116, 60, 37, 10
Path length[µm] = 136, 75.3, 35.3, 10
Vector sum length [µm] = 17.7, 20, 15.8, 5.6
15
10
5
0
−5
−10
−15
−20
−15
−10
−5
0
5
10

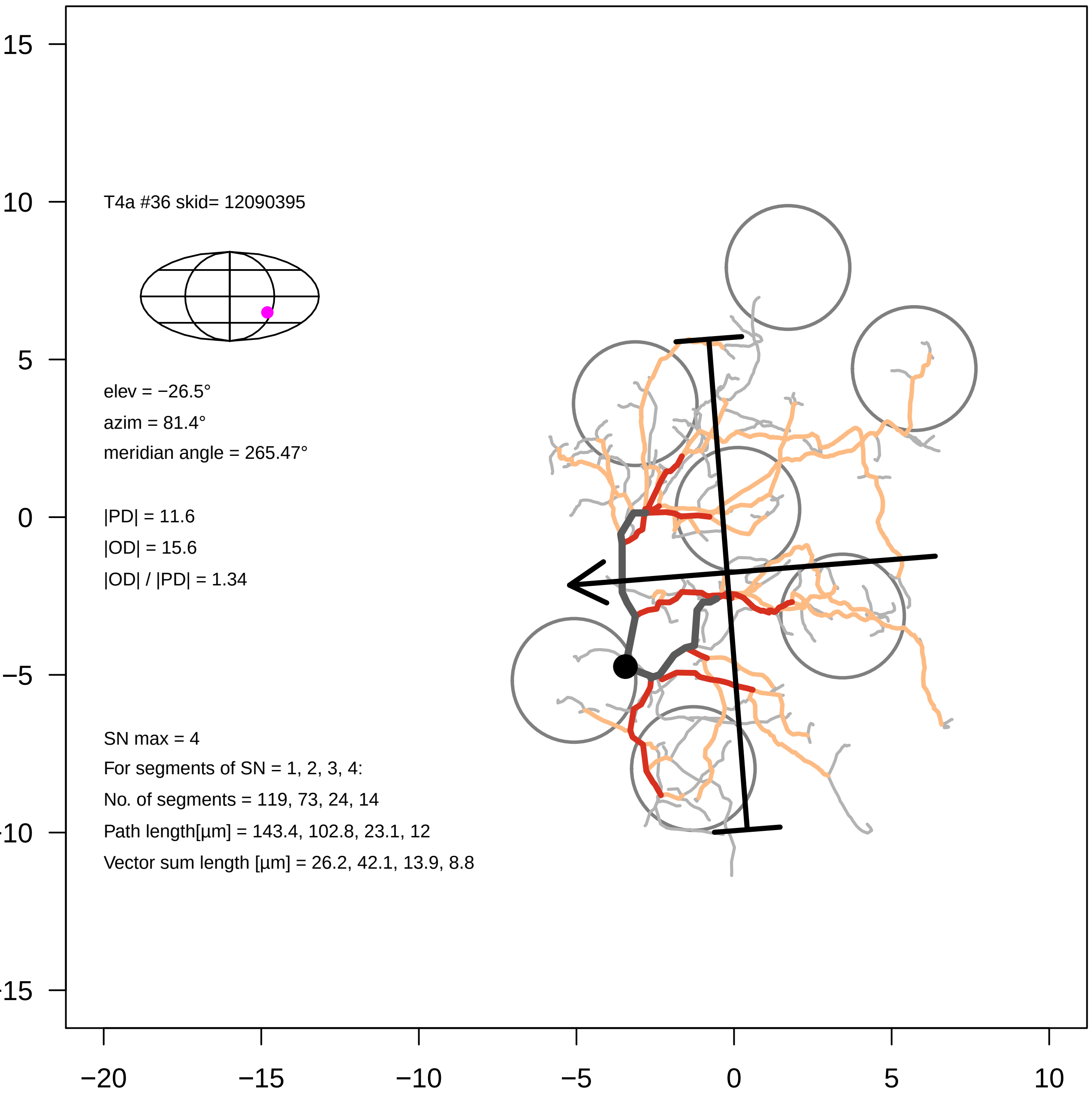

T4a #36 skid= 12090395
elev = −26.5°
azim = 81.4°
meridian angle = 265.47°
|PD| = 11.6
|OD| = 15.6
|OD| / |PD| = 1.34
SN max = 4
For segments of SN = 1, 2, 3, 4:
No. of segments = 119, 73, 24, 14
Path length[µm] = 143.4, 102.8, 23.1, 12
Vector sum length [µm] = 26.2, 42.1, 13.9, 8.8
15
10
5
0
−5
−10
−15
−20
−15
−10
−5
0
5
10

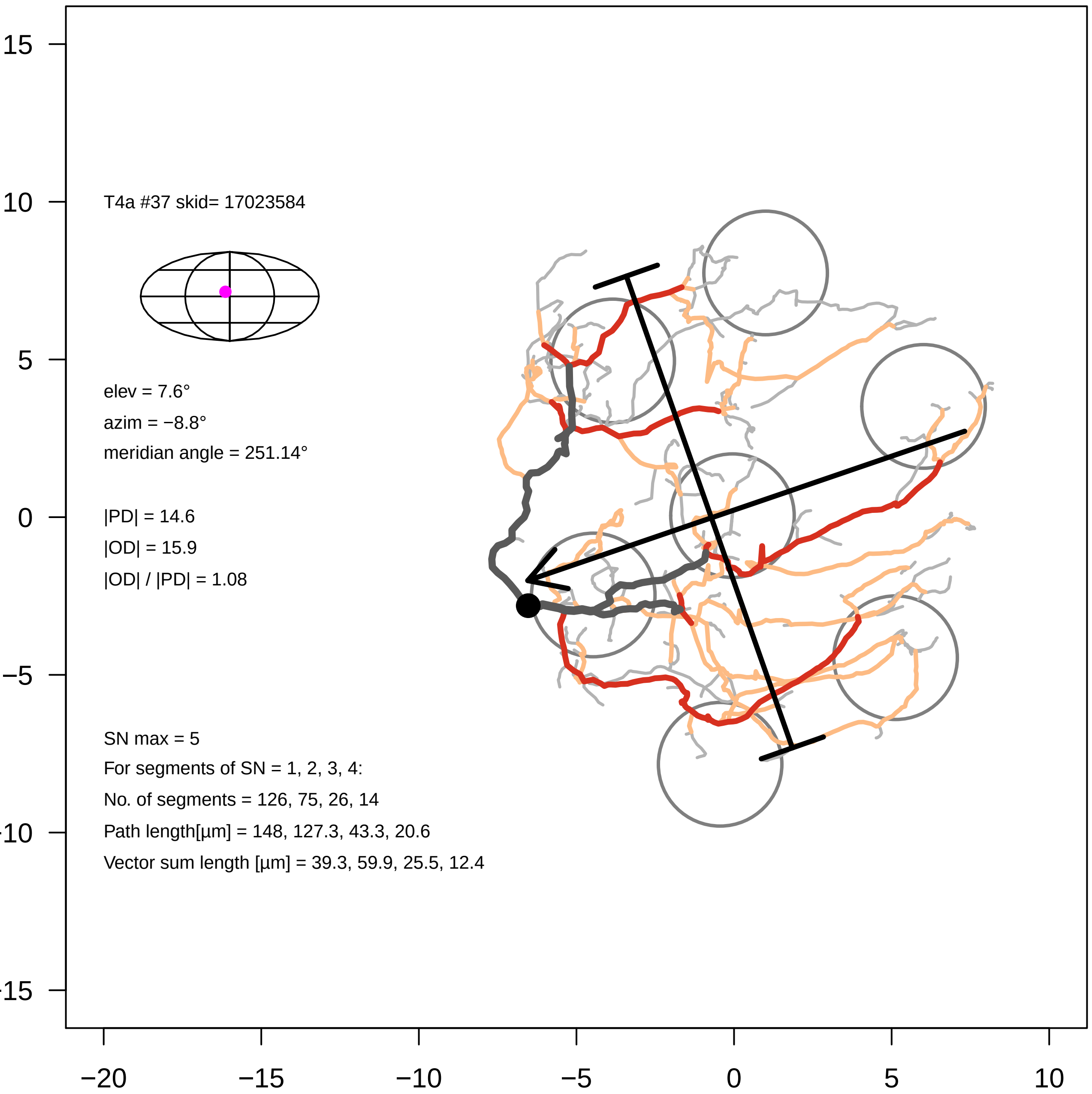

T4a #37 skid= 17023584
elev = 7.6°
azim = −8.8°
meridian angle = 251.14°
|PD| = 14.6
|OD| = 15.9
|OD| / |PD| = 1.08
SN max = 5
For segments of SN = 1, 2, 3, 4:
No. of segments = 126, 75, 26, 14
Path length[µm] = 148, 127.3, 43.3, 20.6
Vector sum length [µm] = 39.3, 59.9, 25.5, 12.4
15
10
5
0
−5
−10
−15
−20
−15
−10
−5
0
5
10

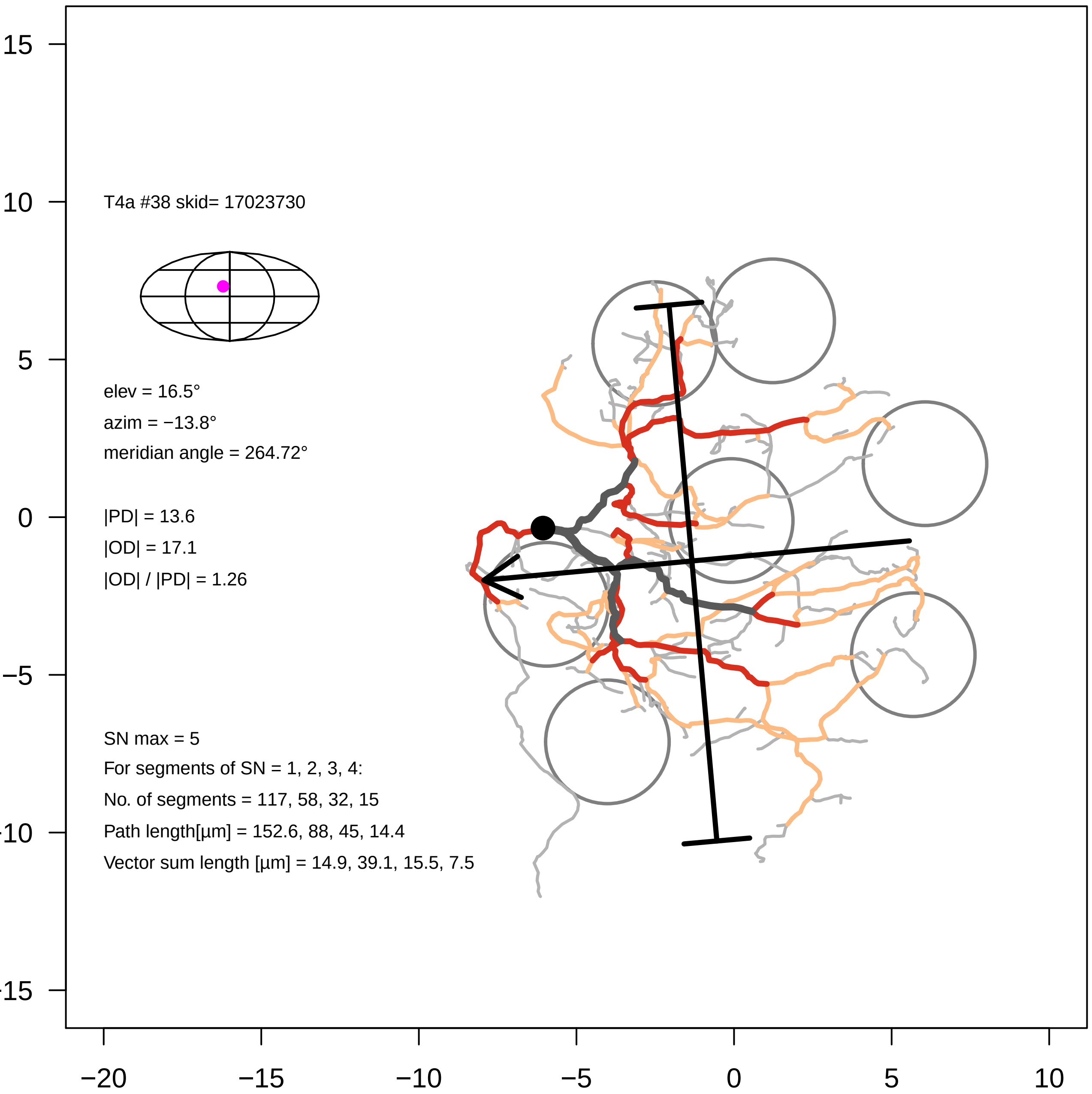
T4a #38 skid= 17023730
elev = 16.5°
azim = −13.8°
meridian angle = 264.72°
|PD| = 13.6
|OD| = 17.1
|OD| / |PD| = 1.26
SN max = 5
For segments of SN = 1, 2, 3, 4:
No. of segments = 117, 58, 32, 15
Path length[µm] = 152.6, 88, 45, 14.4
Vector sum length [µm] = 14.9, 39.1, 15.5, 7.5
−20
−15
−10
−5
0
5
10
15
