## Supplementary Files, T4 galleries for "Eye structure shapes neuron function in *Drosophila* motion vision": Supplementary_Data_File2_T4b.pdf

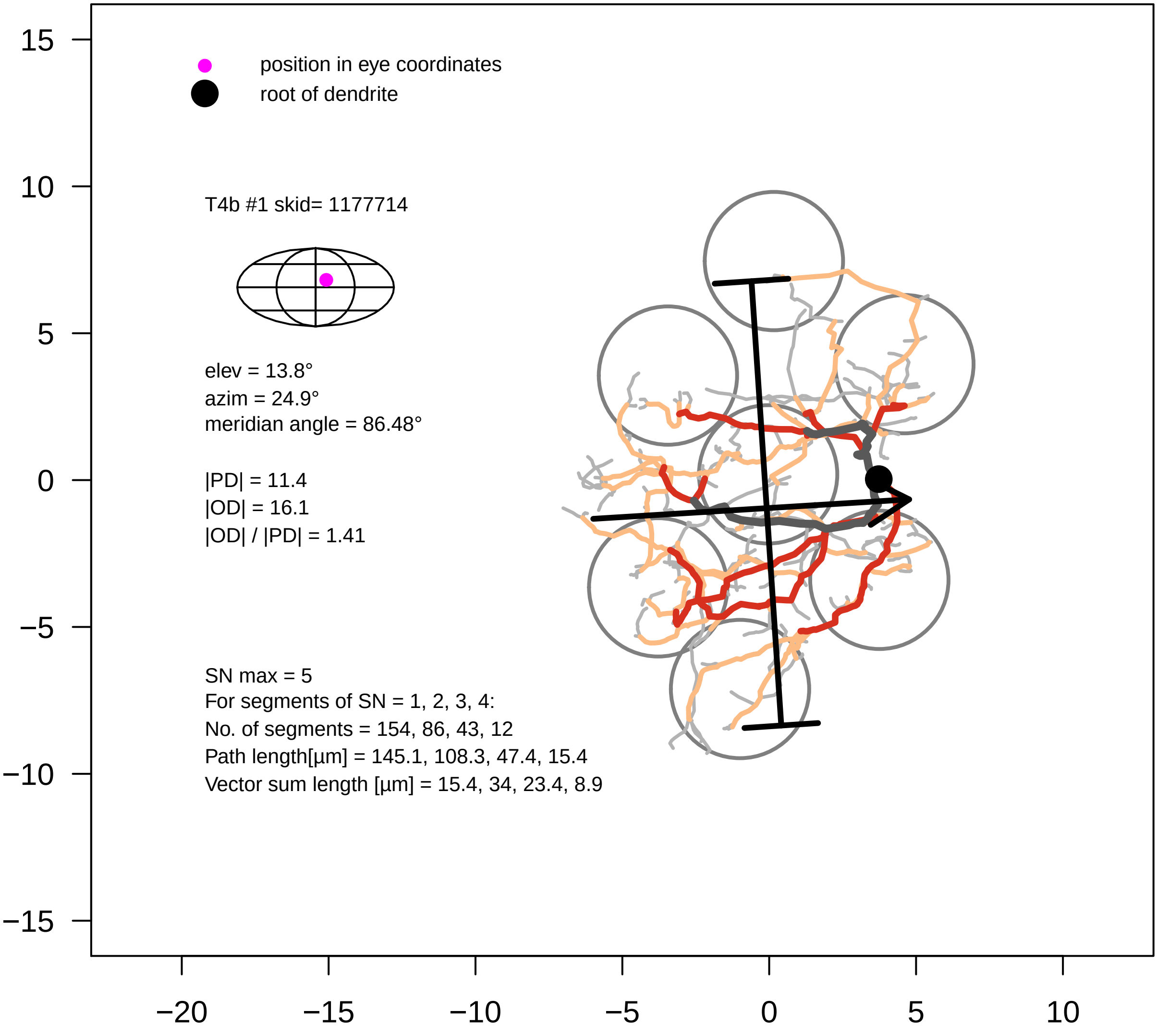
position in eye coordinates
root of dendrite
T4b #1 skid= 1177714
elev = 13.8°
azim = 24.9°
meridian angle = 86.48°
|PD| = 11.4
|OD| = 16.1
|OD| / |PD| = 1.41
SN max = 5
For segments of SN = 1, 2, 3, 4:
No. of segments = 154, 86, 43, 12
Path length[µm] = 145.1, 108.3, 47.4, 15.4
Vector sum length [µm] = 15.4, 34, 23.4, 8.9
15
10
5
0
−5
−10
−15
−20
−15
−10
−5
0
5
10

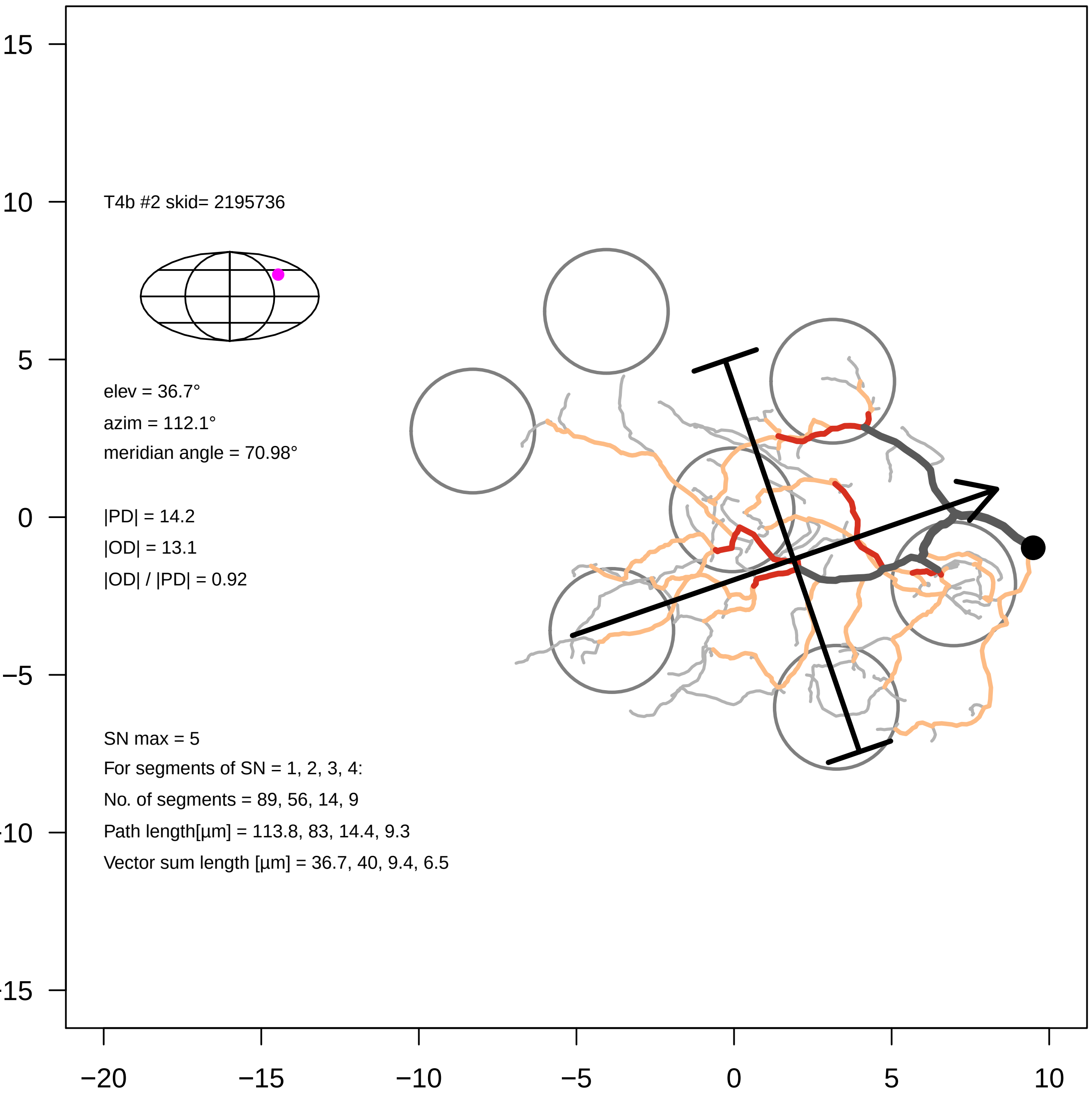
T4b #2 skid= 2195736
elev = 36.7°
azim = 112.1°
meridian angle = 70.98°
|PD| = 14.2
|OD| = 13.1
|OD| / |PD| = 0.92
SN max = 5
For segments of SN = 1, 2, 3, 4:
No. of segments = 89, 56, 14, 9
Path length[µm] = 113.8, 83, 14.4, 9.3
Vector sum length [µm] = 36.7, 40, 9.4, 6.5
15
10
5
0
−5
−10
−15
−20
−15
−10
−5
0
5
10

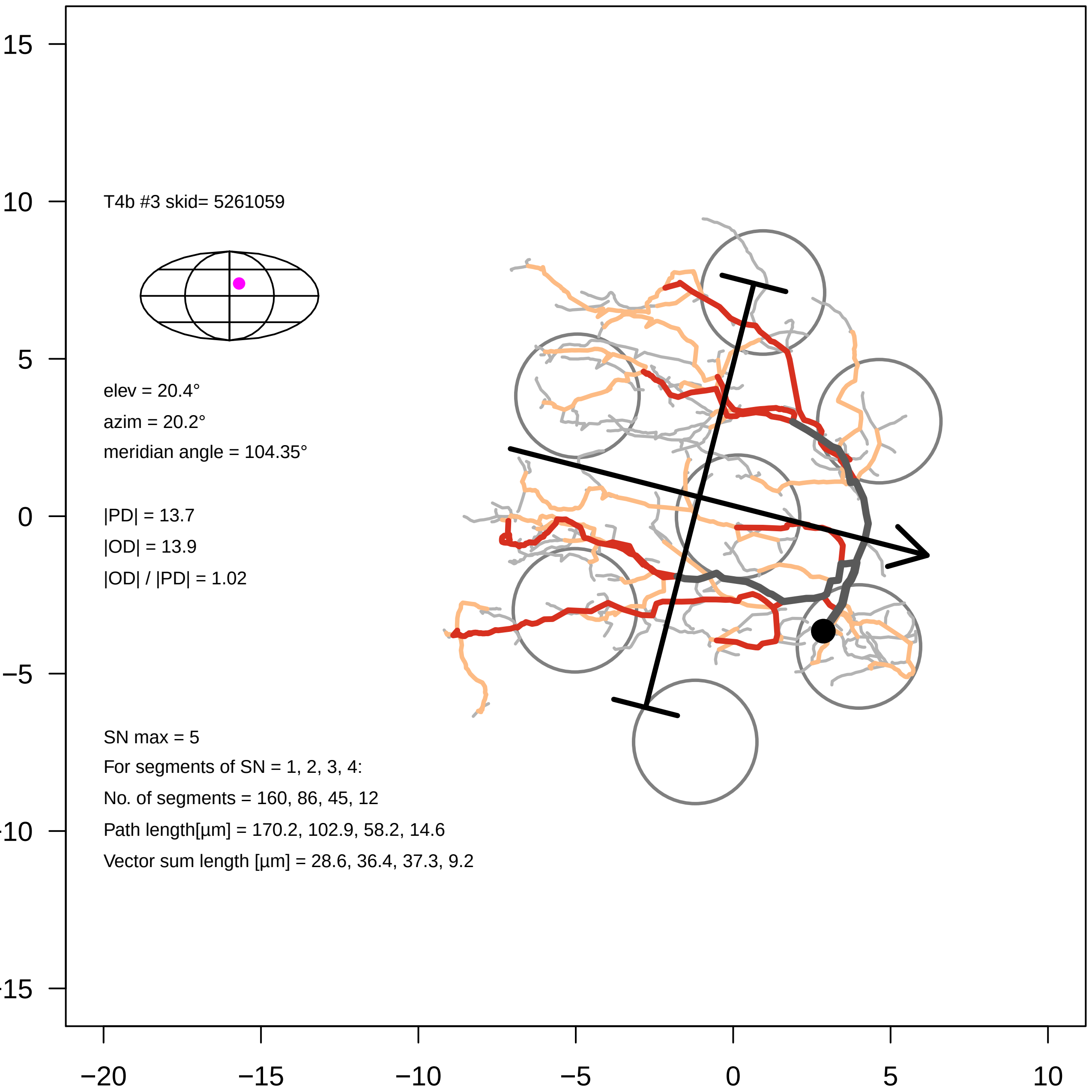

T4b #3 skid= 5261059
elev = 20.4°
azim = 20.2°
meridian angle = 104.35°
|PD| = 13.7
|OD| = 13.9
|OD| / |PD| = 1.02
SN max = 5
For segments of SN = 1, 2, 3, 4:
No. of segments = 160, 86, 45, 12
Path length[µm] = 170.2, 102.9, 58.2, 14.6
Vector sum length [µm] = 28.6, 36.4, 37.3, 9.2
15
10
5
0
−5
−10
−15
−20
−15
−10
−5
0
5
10

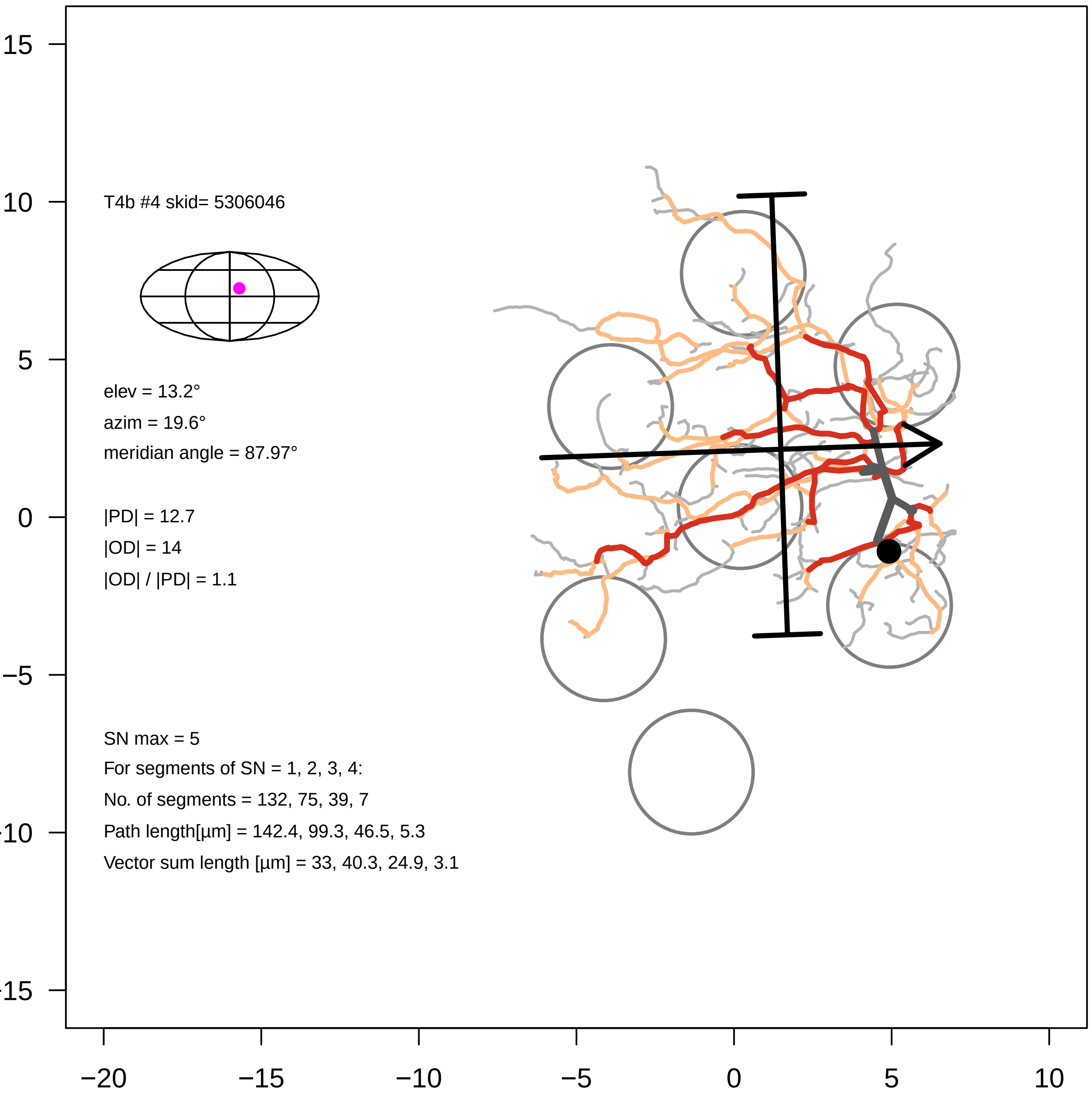
T4b #4 skid= 5306046
elev = 13.2°
azim = 19.6°
meridian angle = 87.97°
|PD| = 12.7
|OD| = 14
|OD| / |PD| = 1.1
SN max = 5
For segments of SN = 1, 2, 3, 4:
No. of segments = 132, 75, 39, 7
Path length[µm] = 142.4, 99.3, 46.5, 5.3
Vector sum length [µm] = 33, 40.3, 24.9, 3.1
15
10
5
0
−5
−10
−15
−20
−15
−10
−5
0
5
10

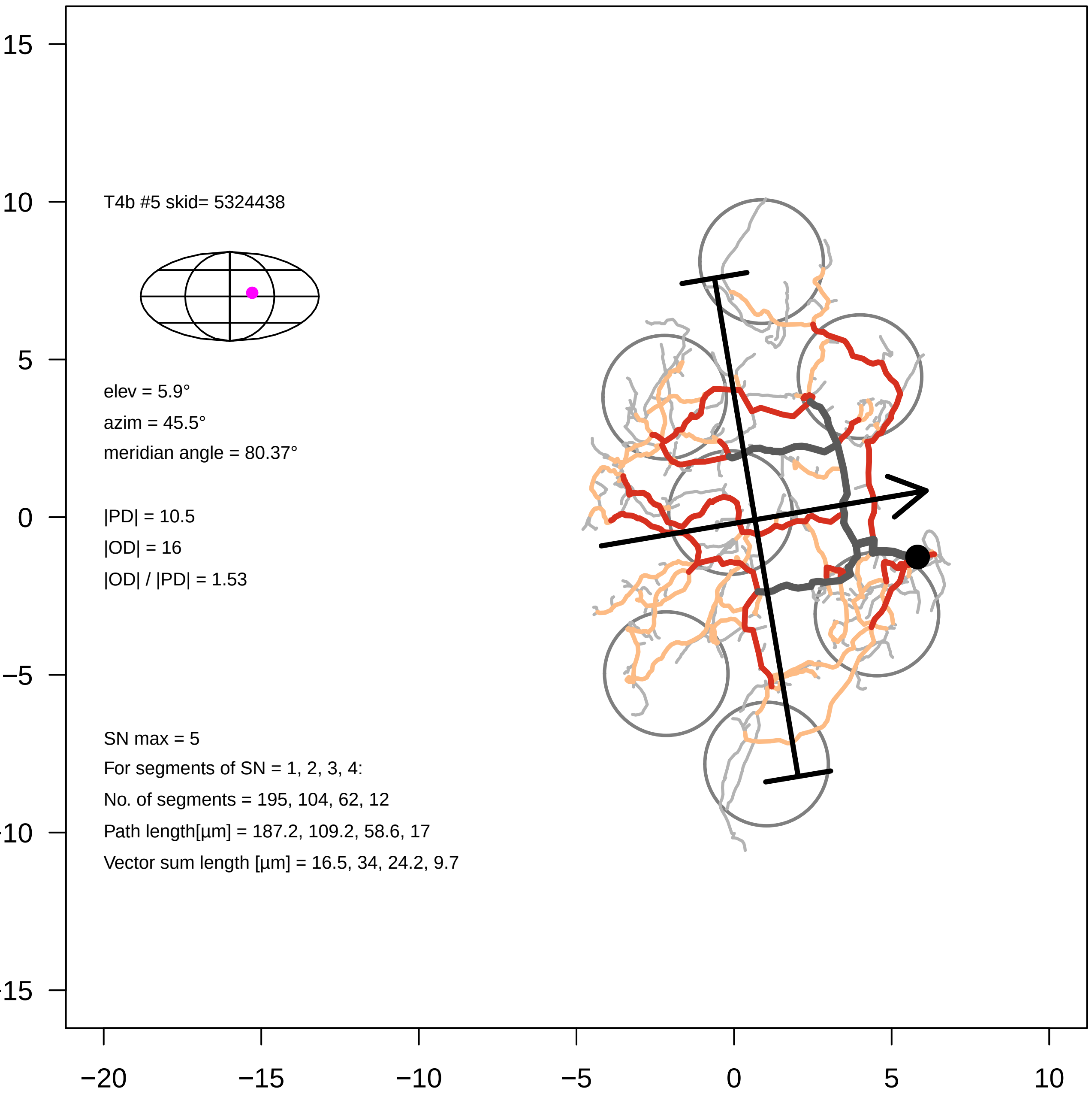
T4b #5 skid= 5324438
elev = 5.9°
azim = 45.5°
meridian angle = 80.37°
|PD| = 10.5
|OD| = 16
|OD| / |PD| = 1.53
SN max = 5
For segments of SN = 1, 2, 3, 4:
No. of segments = 195, 104, 62, 12
Path length[µm] = 187.2, 109.2, 58.6, 17
Vector sum length [µm] = 16.5, 34, 24.2, 9.7
15
10
5
0
−5
−10
−15
−20
−15
−10
−5
0
5
10

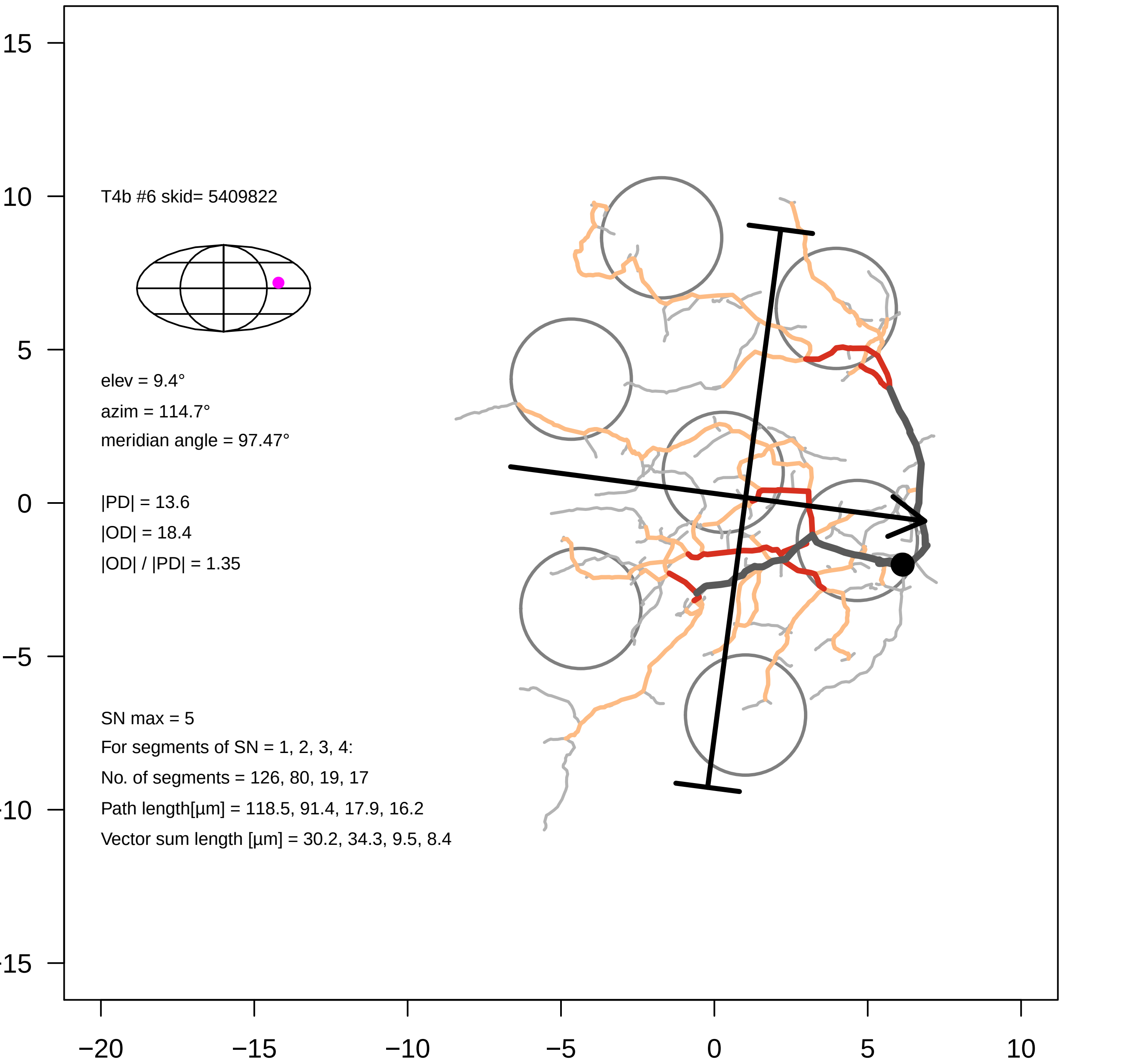

T4b #6 skid= 5409822
elev = 9.4°
azim = 114.7°
meridian angle = 97.47°
|PD| = 13.6
|OD| = 18.4
|OD| / |PD| = 1.35
SN max = 5
For segments of SN = 1, 2, 3, 4:
No. of segments = 126, 80, 19, 17
Path length[µm] = 118.5, 91.4, 17.9, 16.2
Vector sum length [µm] = 30.2, 34.3, 9.5, 8.4
15
10
5
0
−5
−10
−15
−20
−15
−10
−5
0
5
10

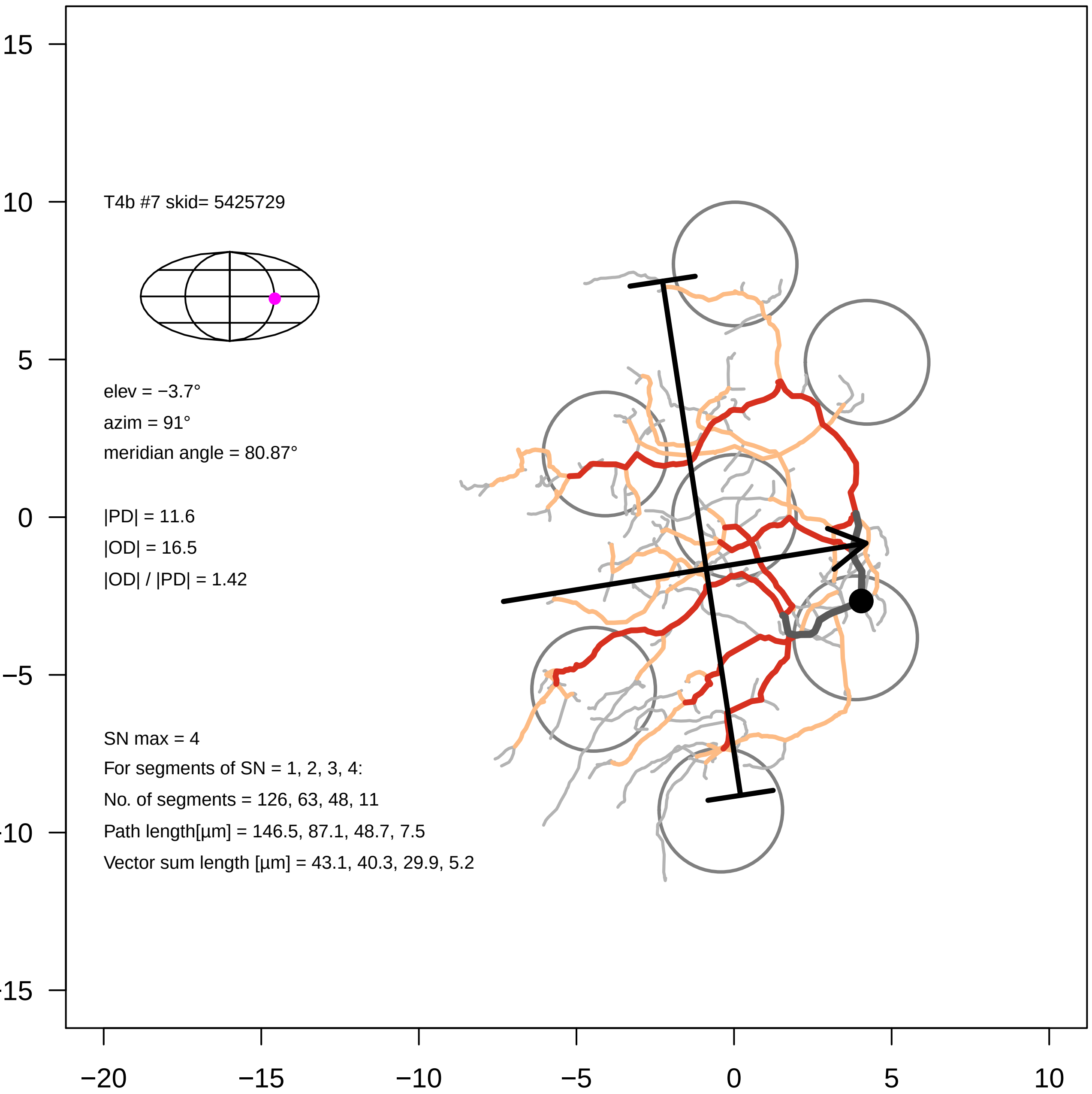
T4b #7 skid= 5425729
elev = −3.7°
azim = 91°
meridian angle = 80.87°
|PD| = 11.6
|OD| = 16.5
|OD| / |PD| = 1.42
SN max = 4
For segments of SN = 1, 2, 3, 4:
No. of segments = 126, 63, 48, 11
Path length[µm] = 146.5, 87.1, 48.7, 7.5
Vector sum length [µm] = 43.1, 40.3, 29.9, 5.2
15
10
5
0
−5
−10
−15
−20
−15
−10
−5
0
5
10

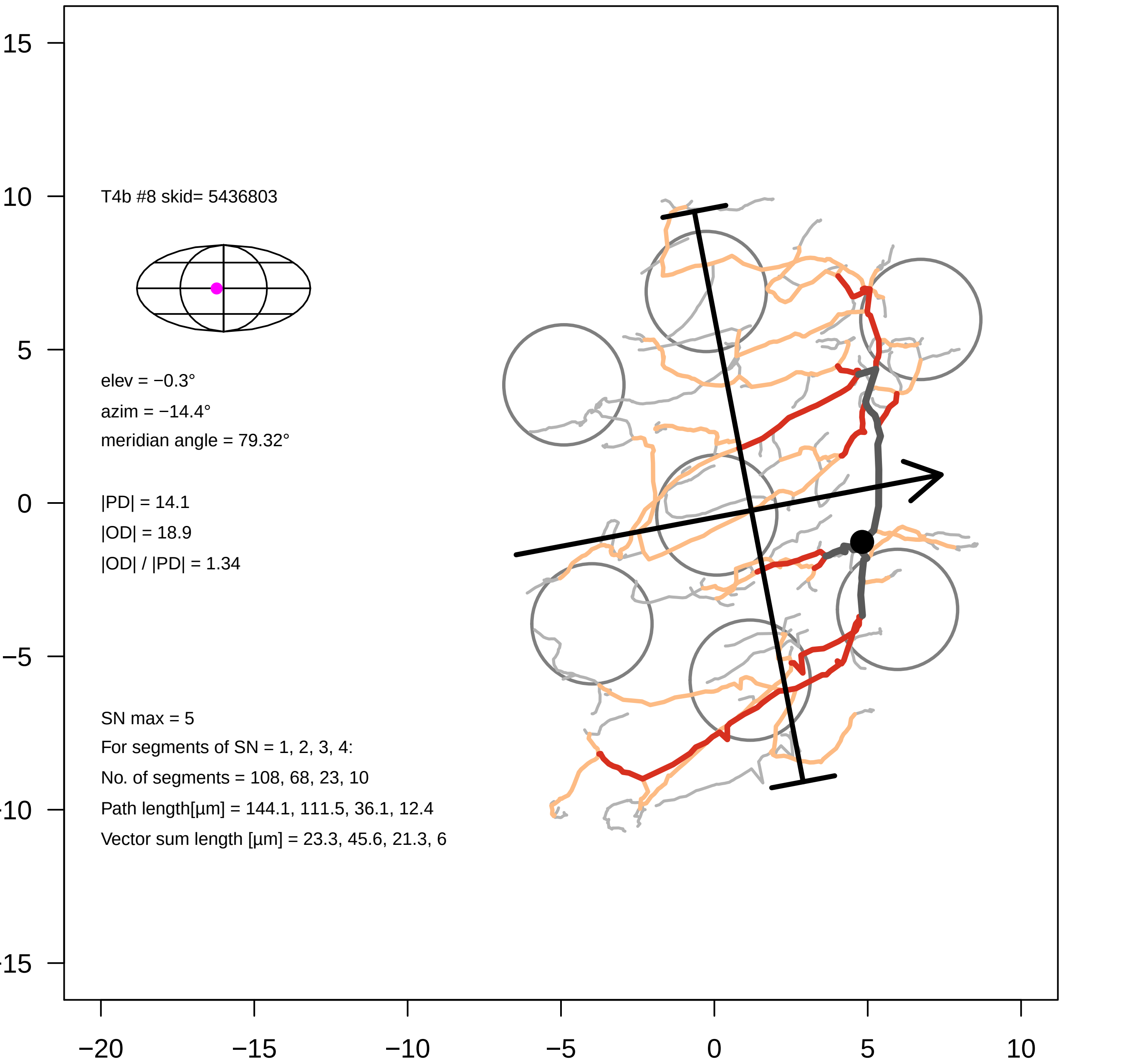

T4b #8 skid= 5436803
elev = −0.3°
azim = −14.4°
meridian angle = 79.32°
|PD| = 14.1
|OD| = 18.9
|OD| / |PD| = 1.34
SN max = 5
For segments of SN = 1, 2, 3, 4:
No. of segments = 108, 68, 23, 10
Path length[µm] = 144.1, 111.5, 36.1, 12.4
Vector sum length [µm] = 23.3, 45.6, 21.3, 6
15
10
5
0
−5
−10
−15
−20
−15
−10
−5
0
5
10

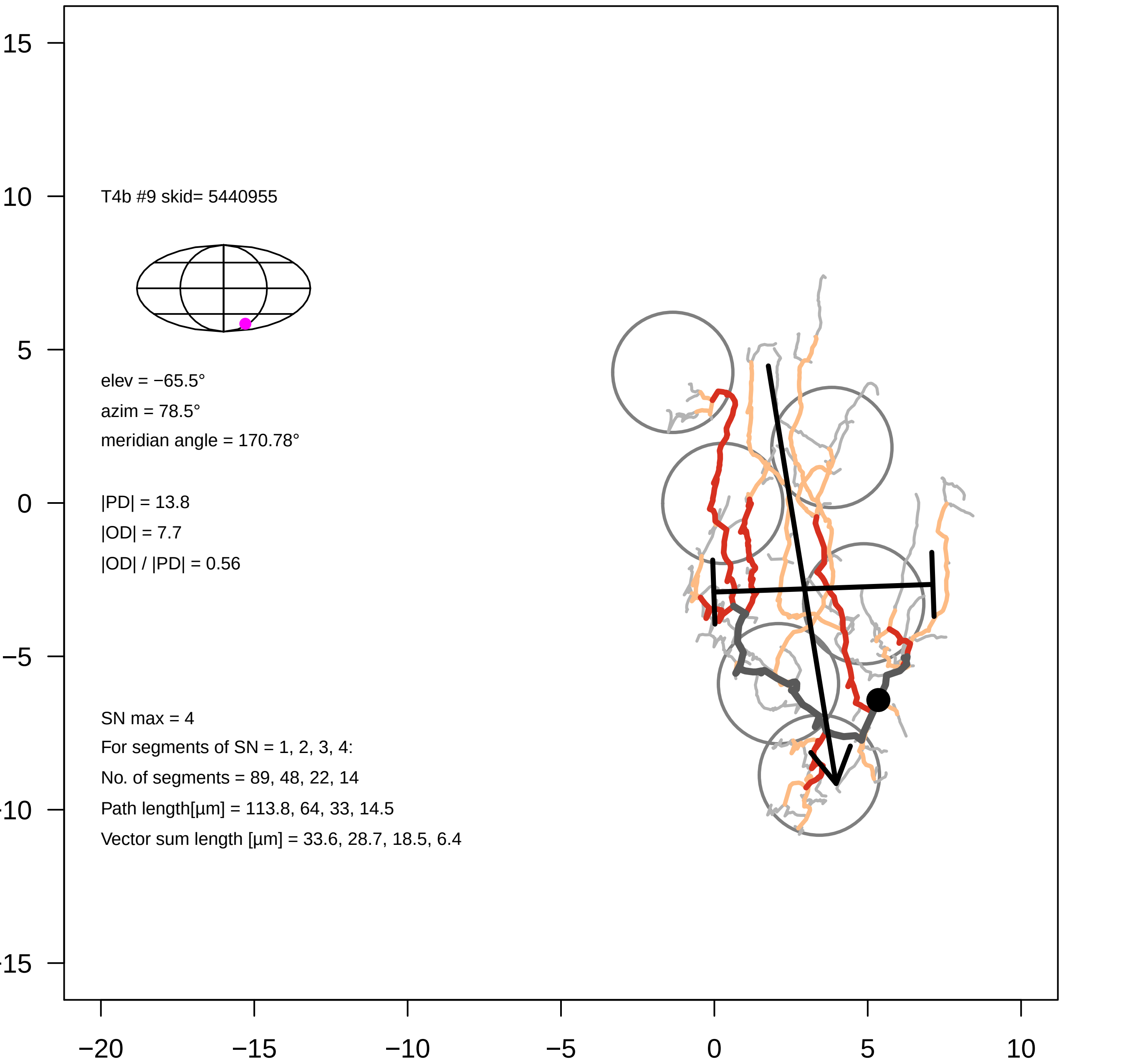
T4b #9 skid= 5440955
elev = −65.5°
azim = 78.5°
meridian angle = 170.78°
|PD| = 13.8
|OD| = 7.7
|OD| / |PD| = 0.56
SN max = 4
For segments of SN = 1, 2, 3, 4:
No. of segments = 89, 48, 22, 14
Path length[µm] = 113.8, 64, 33, 14.5
Vector sum length [µm] = 33.6, 28.7, 18.5, 6.4
15
10
5
0
−5
−15
−20
−15
−10
−5
0
5
10

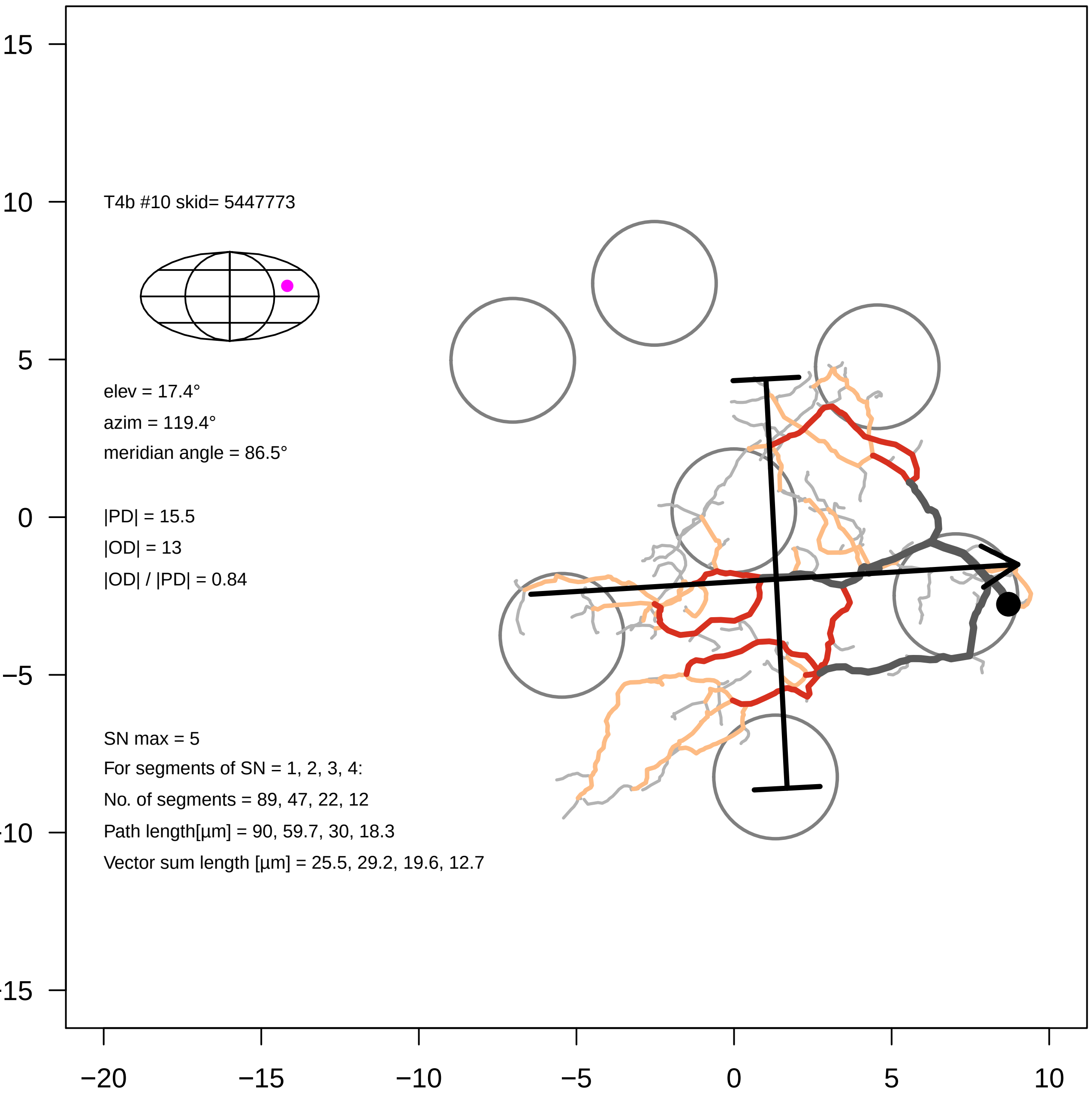
T4b #10 skid= 5447773
elev = 17.4°
azim = 119.4°
meridian angle = 86.5°
|PD| = 15.5
|OD| = 13
|OD| / |PD| = 0.84
SN max = 5
For segments of SN = 1, 2, 3, 4:
No. of segments = 89, 47, 22, 12
Path length[µm] = 90, 59.7, 30, 18.3
Vector sum length [µm] = 25.5, 29.2, 19.6, 12.7
15
10
5
0
−5
−20
−15
−10
−5
0
5
10

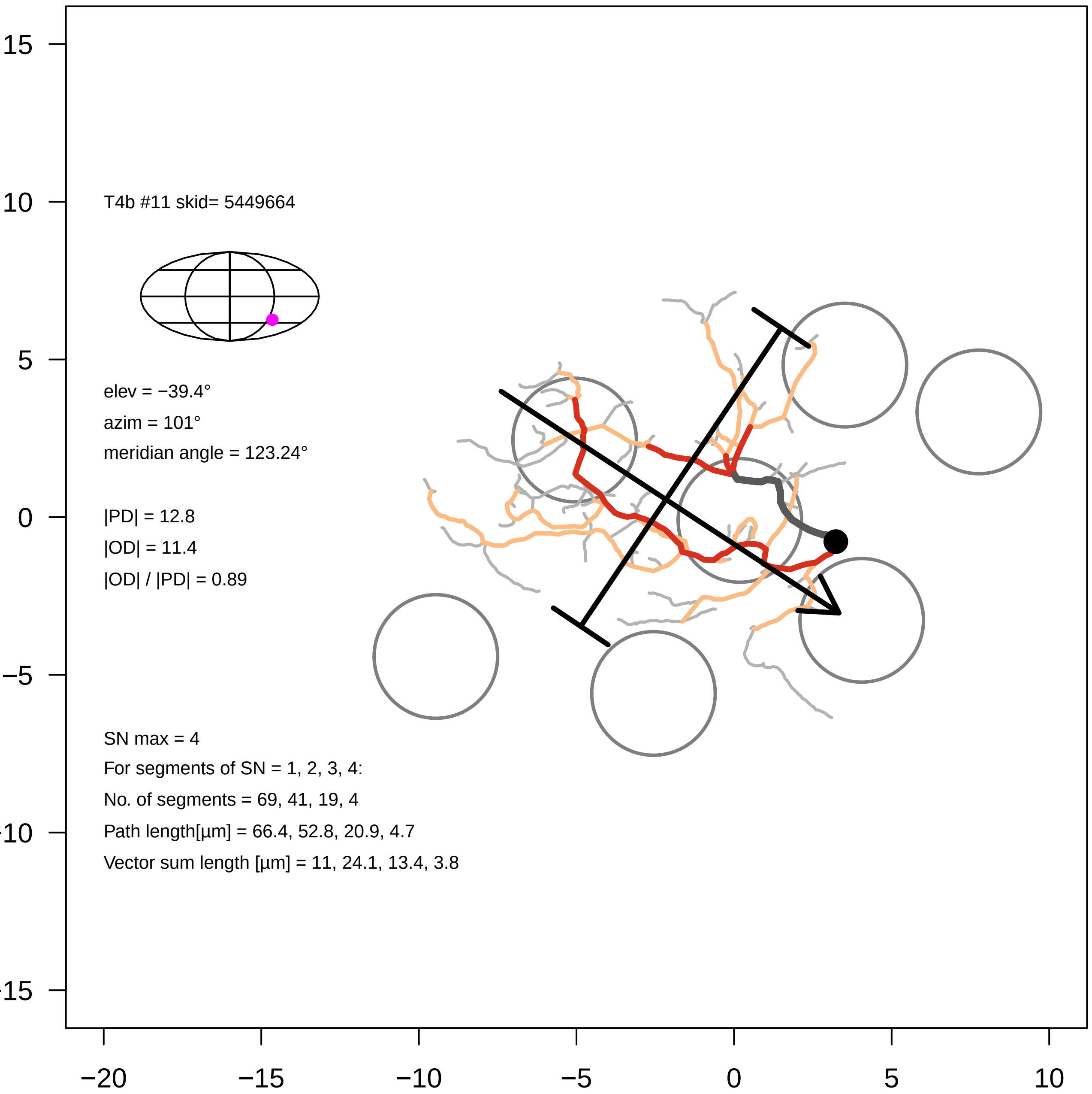
T4b #11 skid= 5449664
elev = −39.4°
azim = 101°
meridian angle = 123.24°
|PD| = 12.8
|OD| = 11.4
|OD| / |PD| = 0.89
SN max = 4
For segments of SN = 1, 2, 3, 4:
No. of segments = 69, 41, 19, 4
Path length[µm] = 66.4, 52.8, 20.9, 4.7
Vector sum length [µm] = 11, 24.1, 13.4, 3.8
15
10
5
0
−5
−10
−15
−20
−15
−10
−5
0
5
10

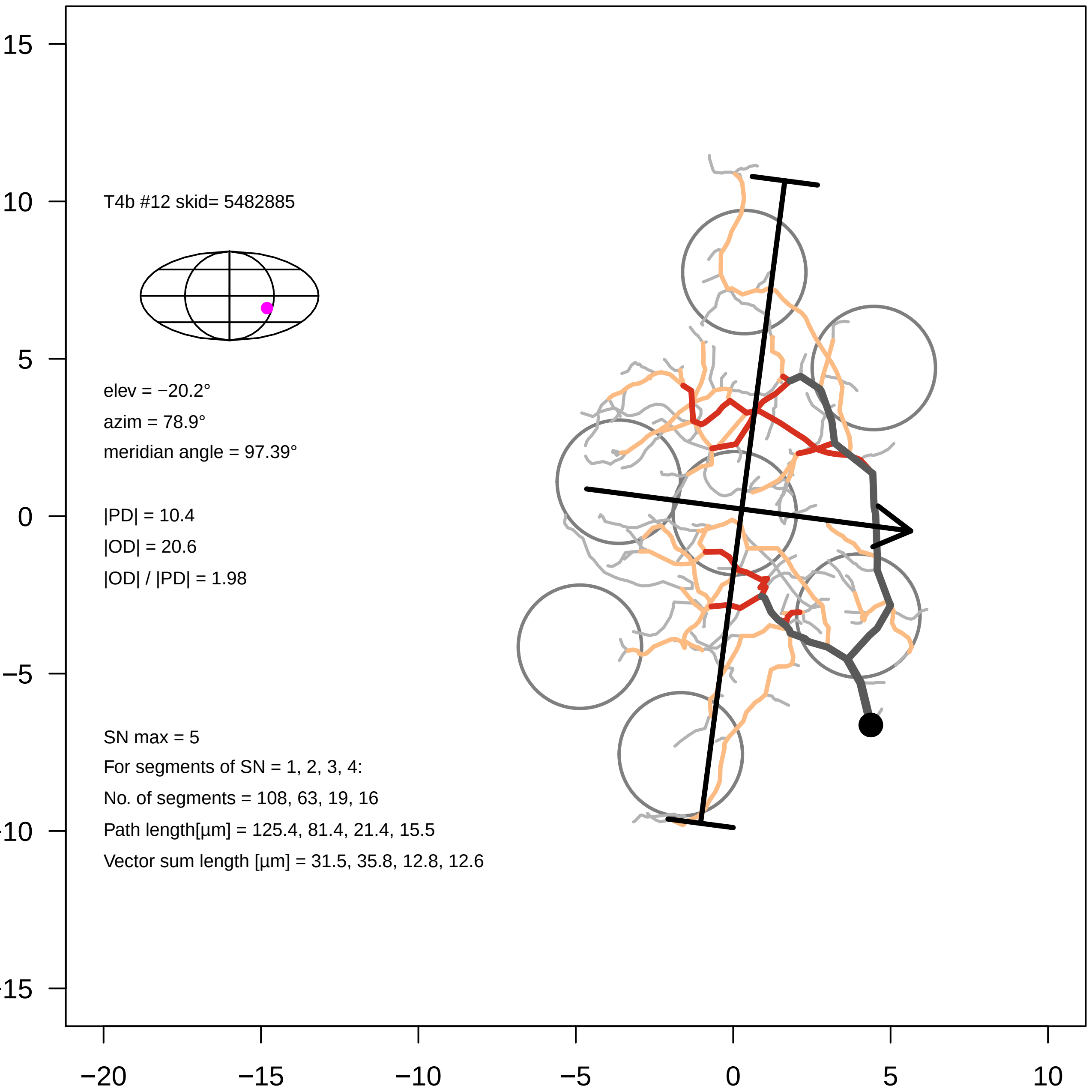

T4b #12 skid= 5482885
elev = −20.2°
azim = 78.9°
meridian angle = 97.39°
|PD| = 10.4
|OD| = 20.6
|OD| / |PD| = 1.98
SN max = 5
For segments of SN = 1, 2, 3, 4:
No. of segments = 108, 63, 19, 16
Path length[µm] = 125.4, 81.4, 21.4, 15.5
Vector sum length [µm] = 31.5, 35.8, 12.8, 12.6
15
10
5
0
−5
−20
−15
−10
−5
0
5
10

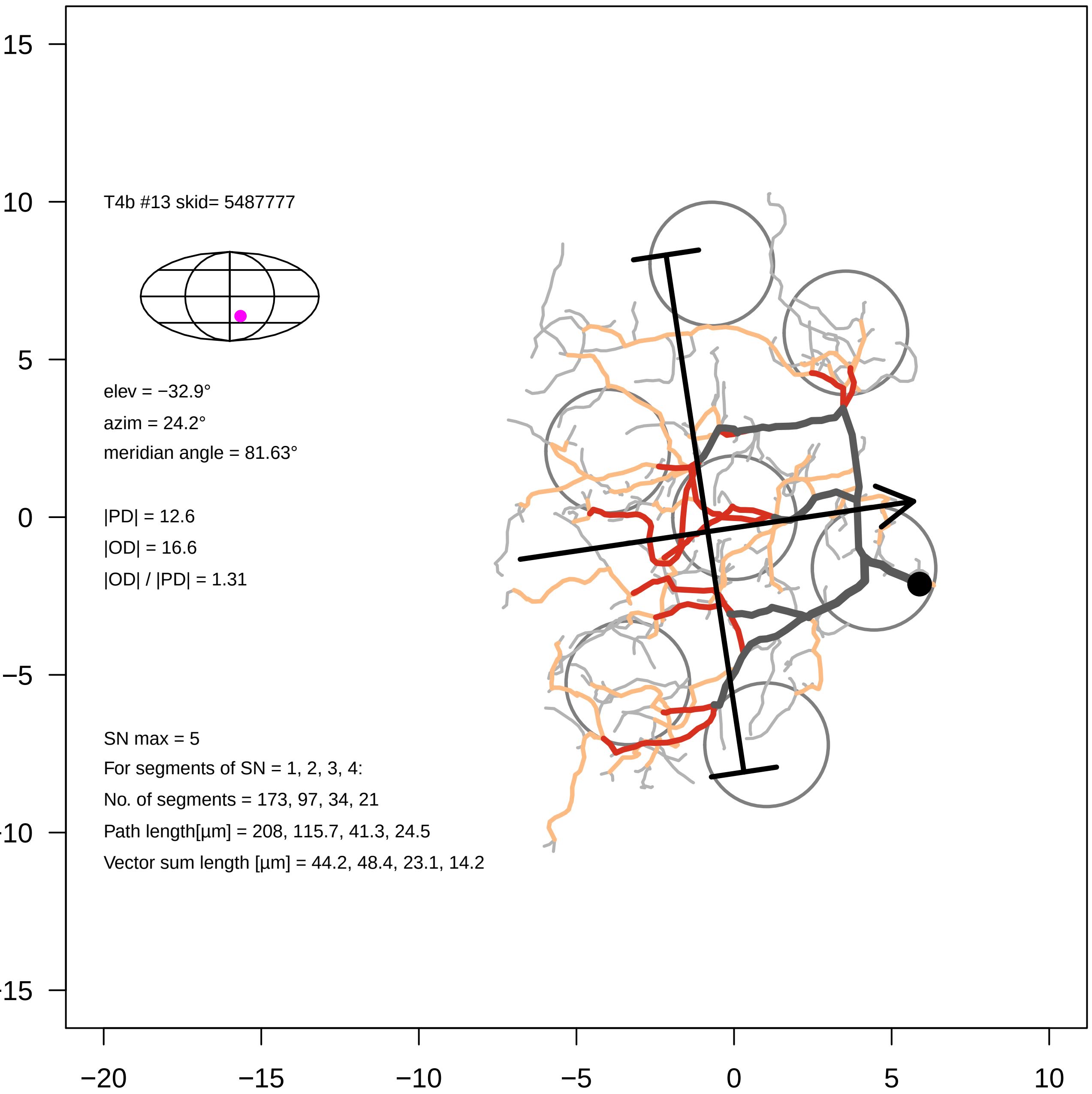

T4b #13 skid= 5487777
elev = −32.9°
azim = 24.2°
meridian angle = 81.63°
|PD| = 12.6
|OD| = 16.6
|OD| / |PD| = 1.31
SN max = 5
For segments of SN = 1, 2, 3, 4:
No. of segments = 173, 97, 34, 21
Path length[µm] = 208, 115.7, 41.3, 24.5
Vector sum length [µm] = 44.2, 48.4, 23.1, 14.2
15
10
5
0
−5
−10
−15
−20
−15
−10
−5
0
5
10

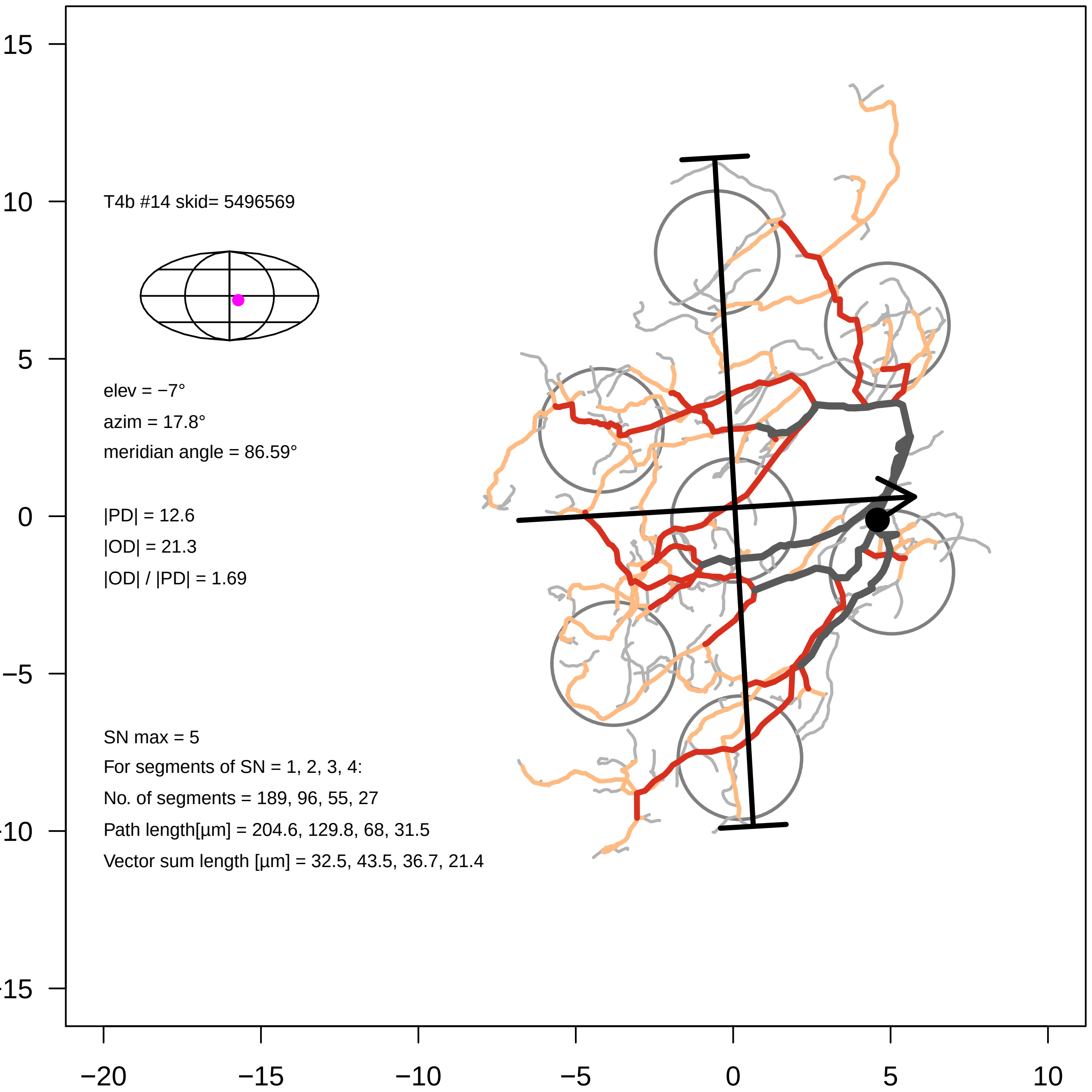
T4b #14 skid= 5496569
elev = −7°
azim = 17.8°
meridian angle = 86.59°
|PD| = 12.6
|OD| = 21.3
|OD| / |PD| = 1.69
SN max = 5
For segments of SN = 1, 2, 3, 4:
No. of segments = 189, 96, 55, 27
Path length[µm] = 204.6, 129.8, 68, 31.5
Vector sum length [µm] = 32.5, 43.5, 36.7, 21.4
15
10
5
0
−5
−20
−15
−10
−5
0
5
10

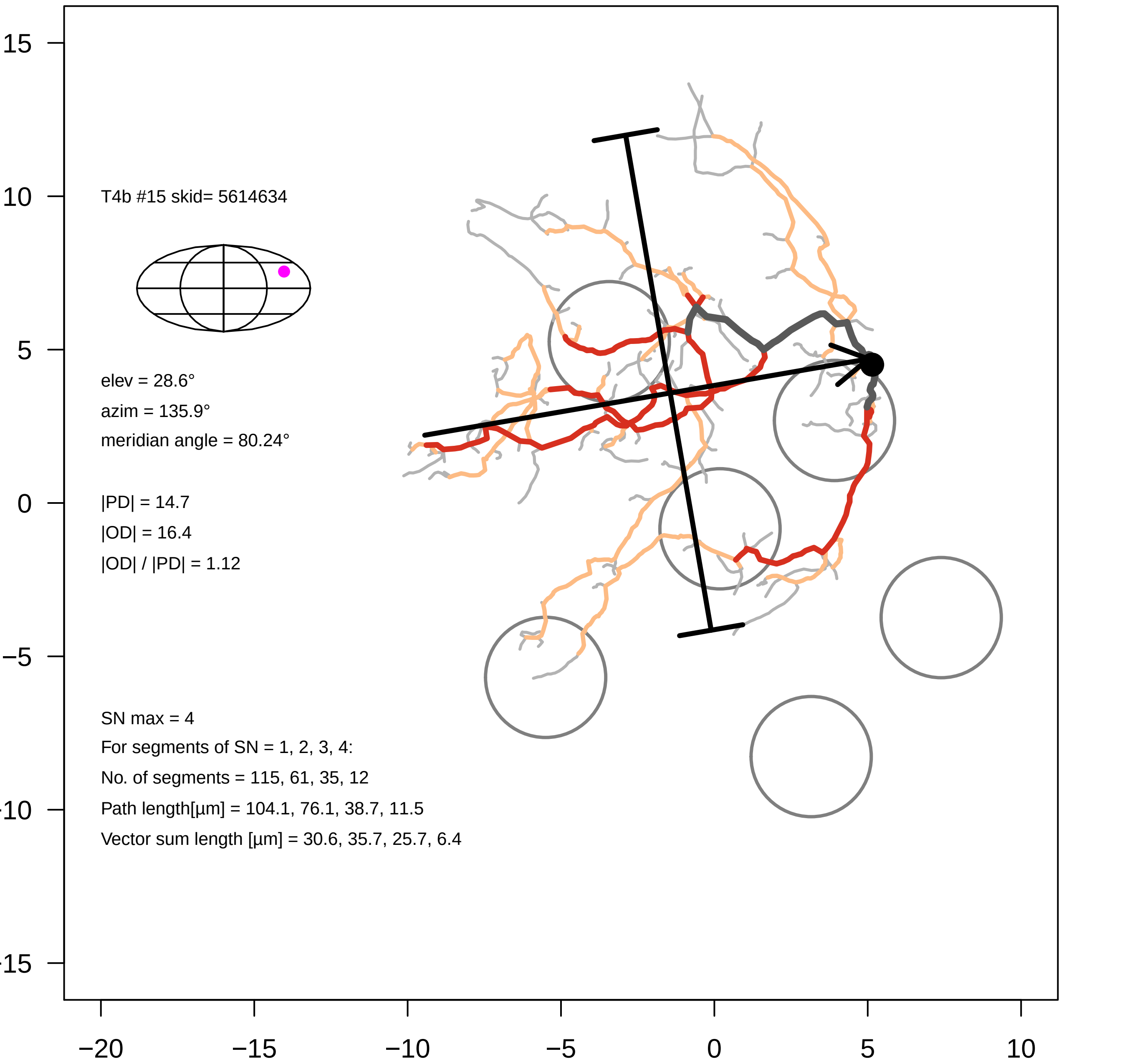
T4b #15 skid= 5614634
elev = 28.6°
azim = 135.9°
meridian angle = 80.24°
|PD| = 14.7
|OD| = 16.4
|OD| / |PD| = 1.12
SN max = 4
For segments of SN = 1, 2, 3, 4:
No. of segments = 115, 61, 35, 12
Path length[µm] = 104.1, 76.1, 38.7, 11.5
Vector sum length [µm] = 30.6, 35.7, 25.7, 6.4
15
10
5
0
−5
−10
−15
−20
−15
−10
−5
0
5
10

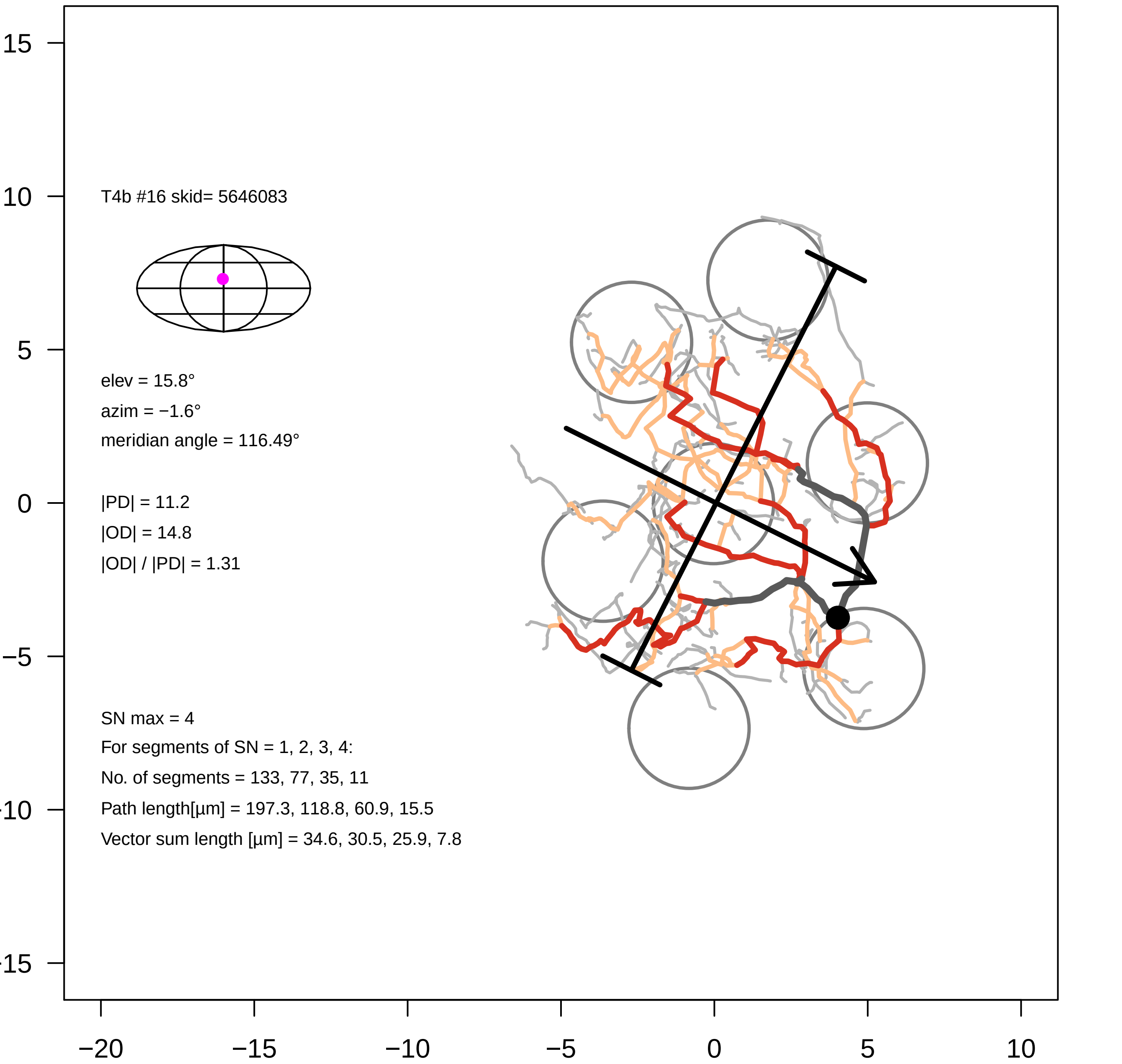
T4b #16 skid= 5646083
elev = 15.8°
azim = −1.6°
meridian angle = 116.49°
|PD| = 11.2
|OD| = 14.8
|OD| / |PD| = 1.31
SN max = 4
For segments of SN = 1, 2, 3, 4:
No. of segments = 133, 77, 35, 11
Path length[µm] = 197.3, 118.8, 60.9, 15.5
Vector sum length [µm] = 34.6, 30.5, 25.9, 7.8
15
10
5
0
−5
−10
−15
−20
−15
−10
−5
0
5
10

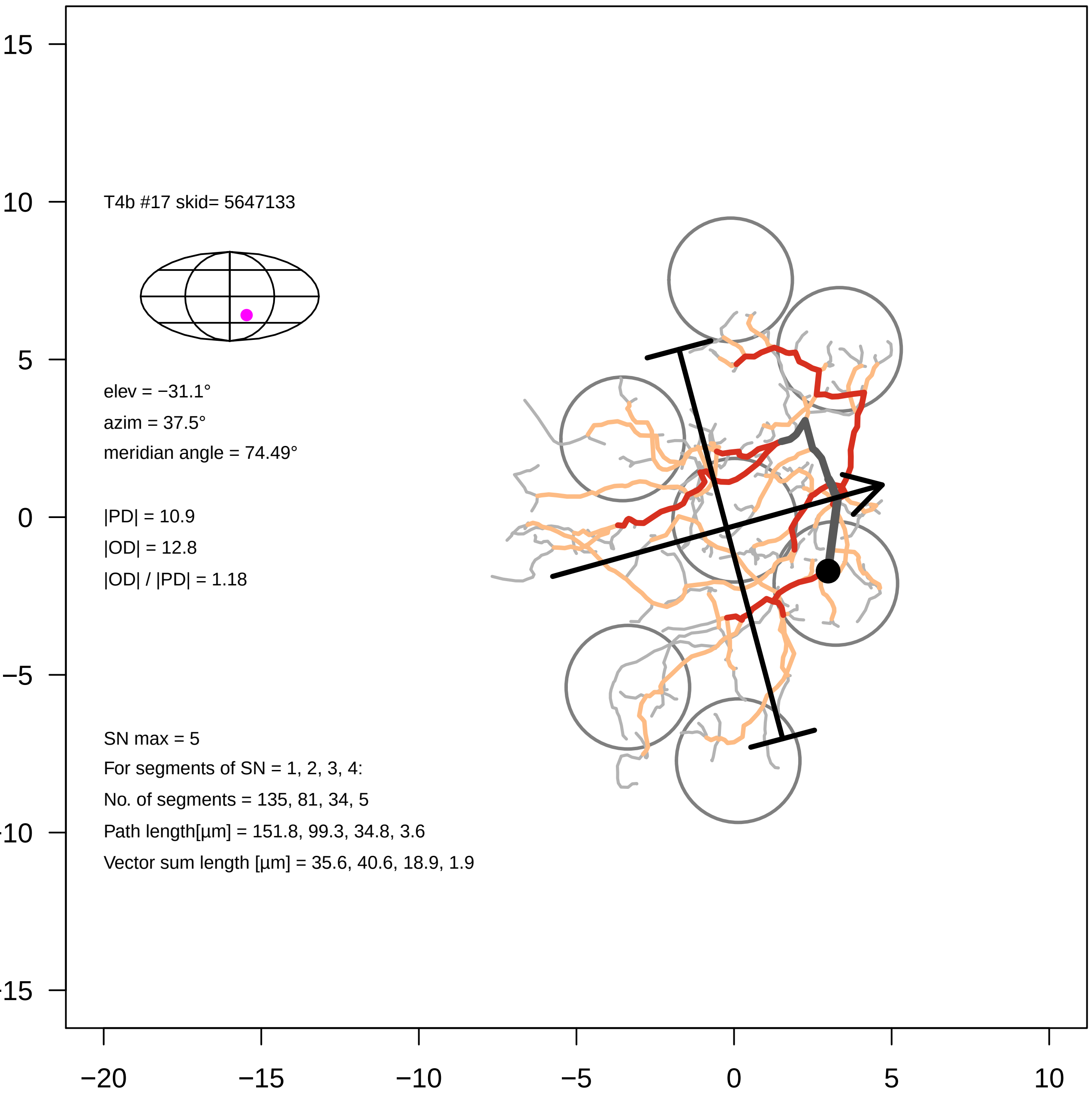
T4b #17 skid= 5647133
elev = −31.1°
azim = 37.5°
meridian angle = 74.49°
|PD| = 10.9
|OD| = 12.8
|OD| / |PD| = 1.18
SN max = 5
For segments of SN = 1, 2, 3, 4:
No. of segments = 135, 81, 34, 5
Path length[µm] = 151.8, 99.3, 34.8, 3.6
Vector sum length [µm] = 35.6, 40.6, 18.9, 1.9
15
10
5
0
−5
−10
−15
−20
−15
−10
−5
0
5
10

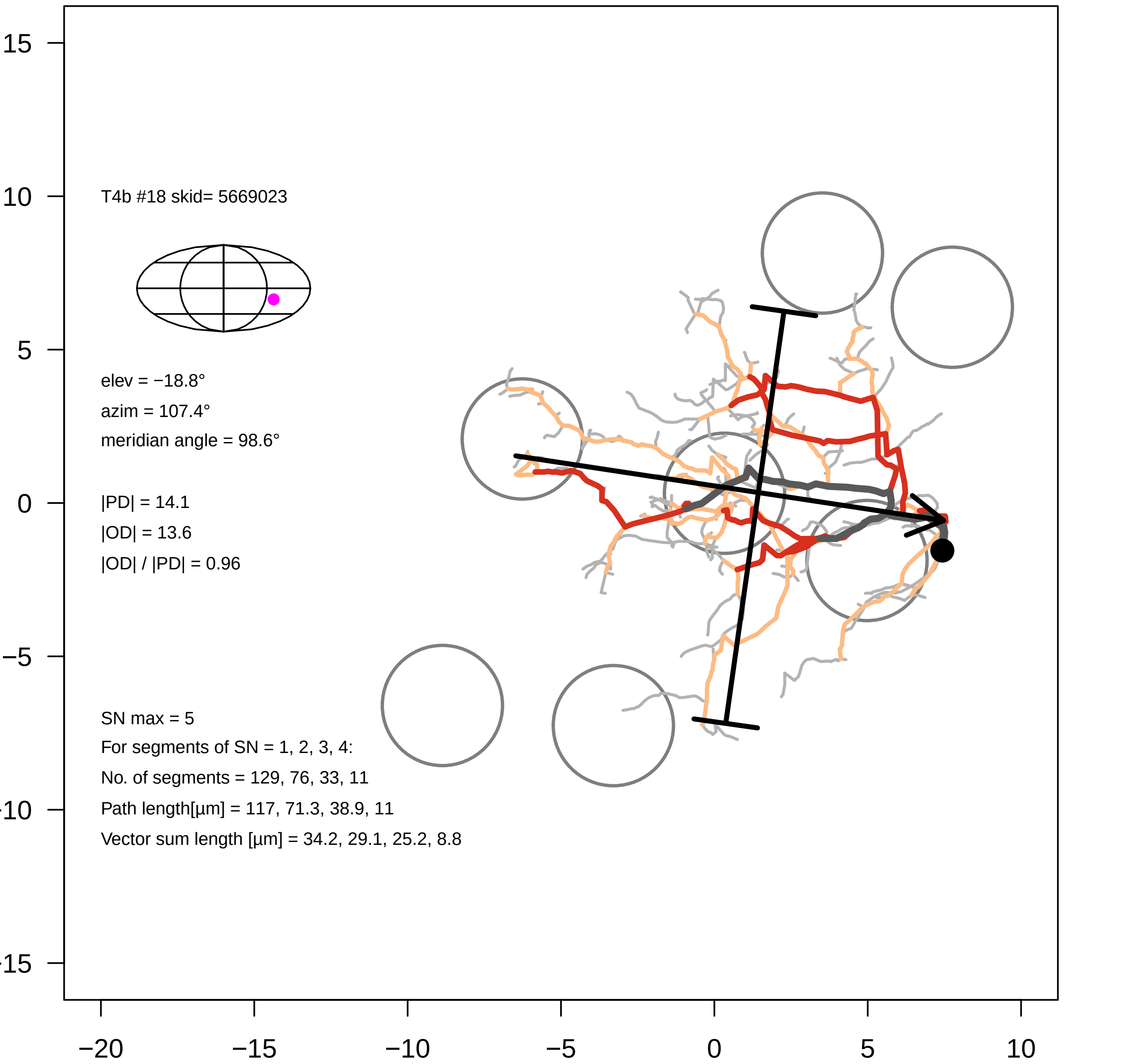

T4b #18 skid= 5669023
elev = −18.8°
azim = 107.4°
meridian angle = 98.6°
|PD| = 14.1
|OD| = 13.6
|OD| / |PD| = 0.96
SN max = 5
For segments of SN = 1, 2, 3, 4:
No. of segments = 129, 76, 33, 11
Path length[µm] = 117, 71.3, 38.9, 11
Vector sum length [µm] = 34.2, 29.1, 25.2, 8.8
15
10
5
0
−5
-10
-15
−20
−15
−10
−5
0
5
10

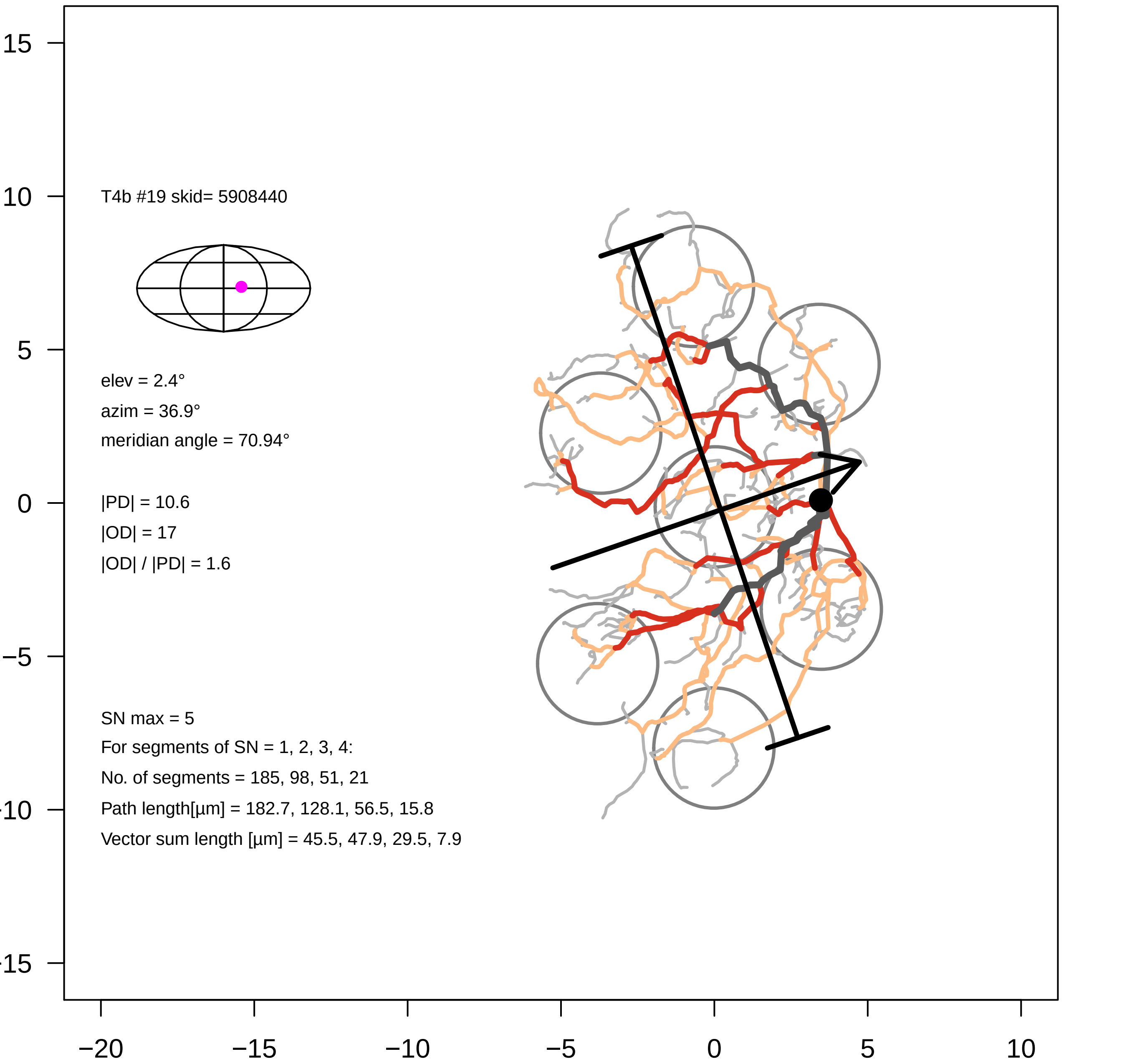

T4b #19 skid= 5908440
elev = 2.4°
azim = 36.9°
meridian angle = 70.94°
|PD| = 10.6
|OD| = 17
|OD| / |PD| = 1.6
SN max = 5
For segments of SN = 1, 2, 3, 4:
No. of segments = 185, 98, 51, 21
Path length[µm] = 182.7, 128.1, 56.5, 15.8
Vector sum length [µm] = 45.5, 47.9, 29.5, 7.9
15
10
5
0
−5
−10
−15
−20
−15
−10
−5
0
5
10

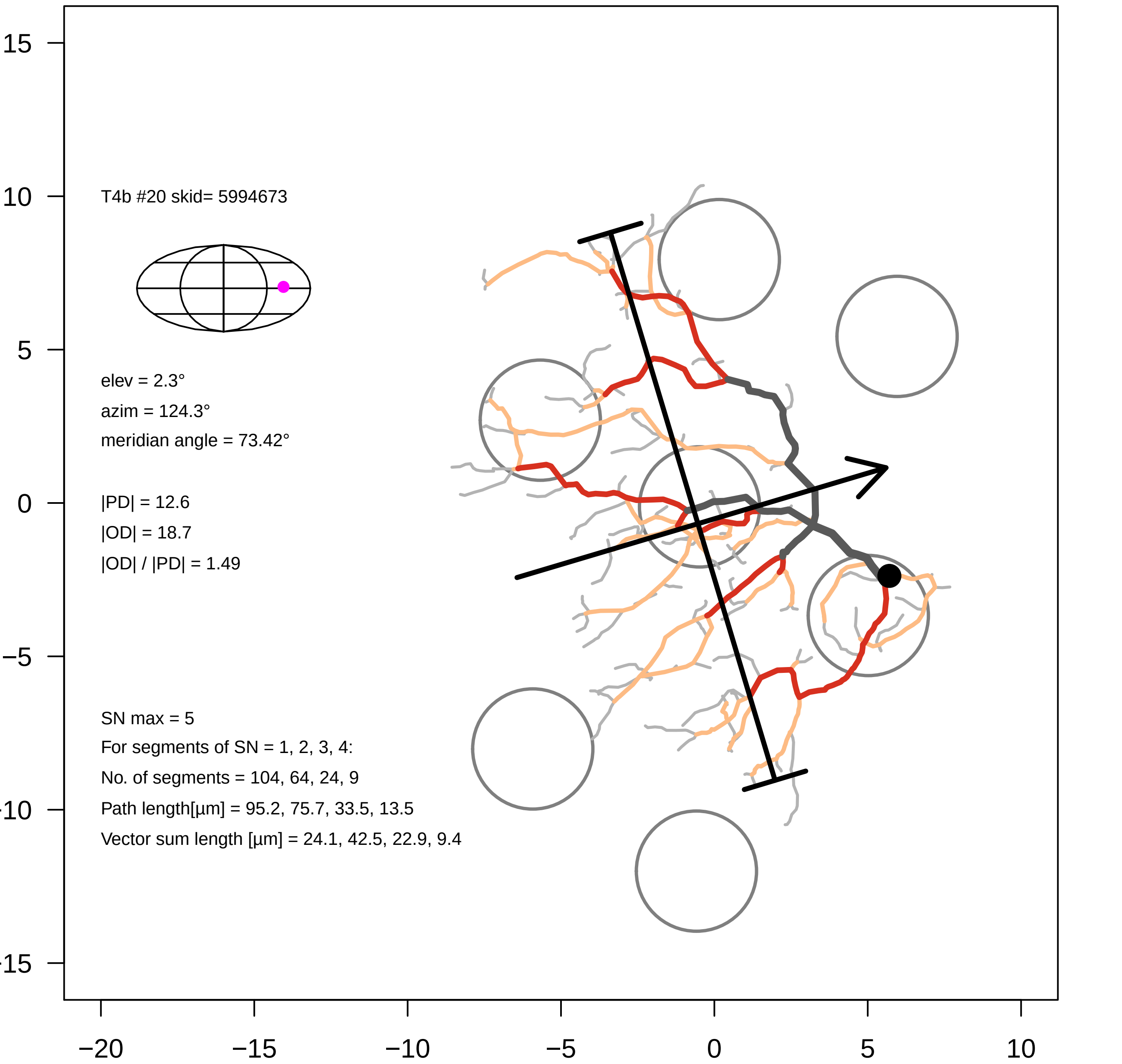
T4b #20 skid= 5994673
elev = 2.3°
azim = 124.3°
meridian angle = 73.42°
|PD| = 12.6
|OD| = 18.7
|OD| / |PD| = 1.49
SN max = 5
For segments of SN = 1, 2, 3, 4:
No. of segments = 104, 64, 24, 9
Path length[µm] = 95.2, 75.7, 33.5, 13.5
Vector sum length [µm] = 24.1, 42.5, 22.9, 9.4
−20
−15
−10
−5
0
5
10
15

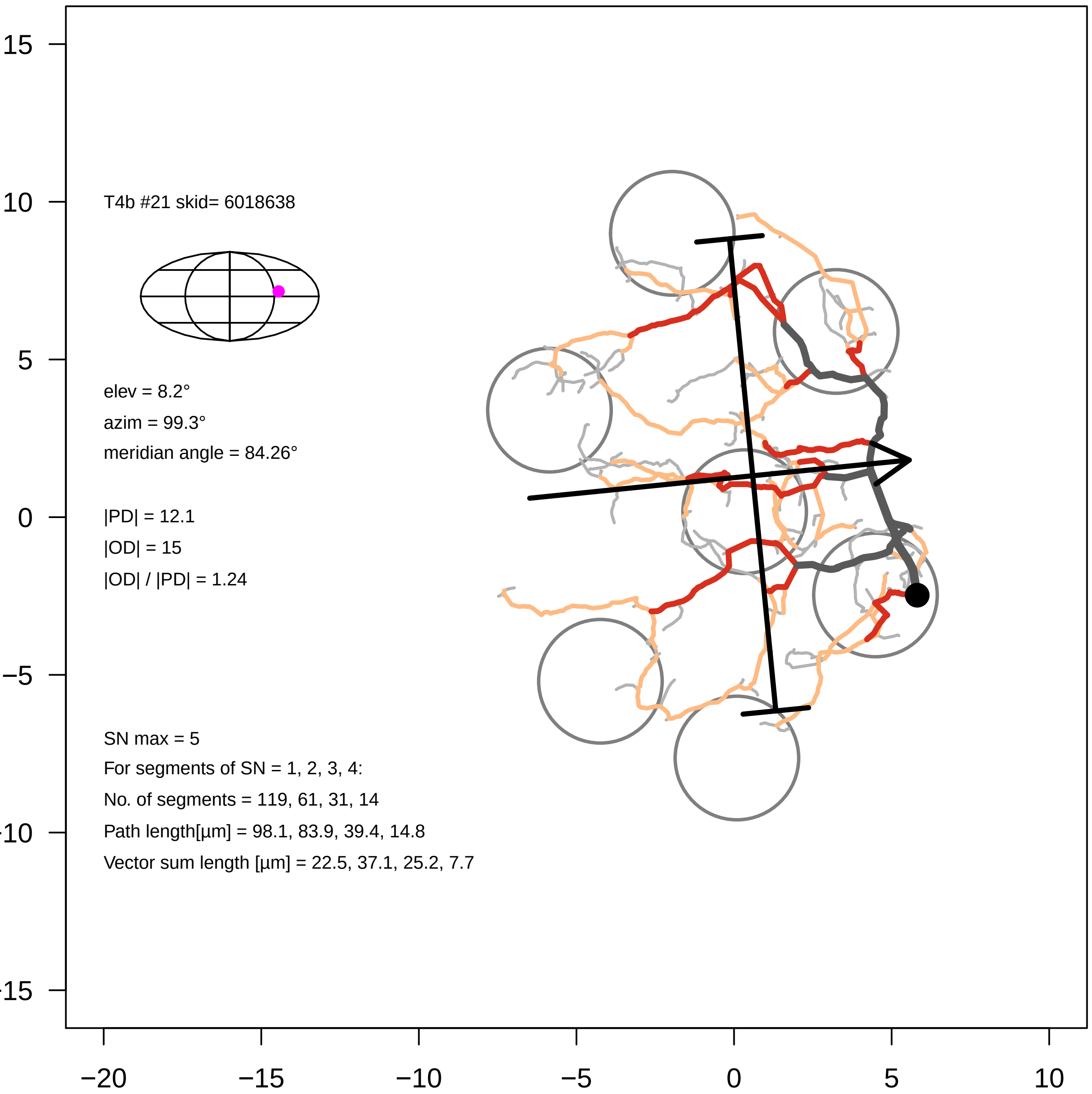
T4b #21 skid= 6018638
elev = 8.2°
azim = 99.3°
meridian angle = 84.26°
|PD| = 12.1
|OD| = 15
|OD| / |PD| = 1.24
SN max = 5
For segments of SN = 1, 2, 3, 4:
No. of segments = 119, 61, 31, 14
Path length[µm] = 98.1, 83.9, 39.4, 14.8
Vector sum length [µm] = 22.5, 37.1, 25.2, 7.7
15
10
5
0
−5
−10
−15
−20
−15
−10
−5
0
5
10

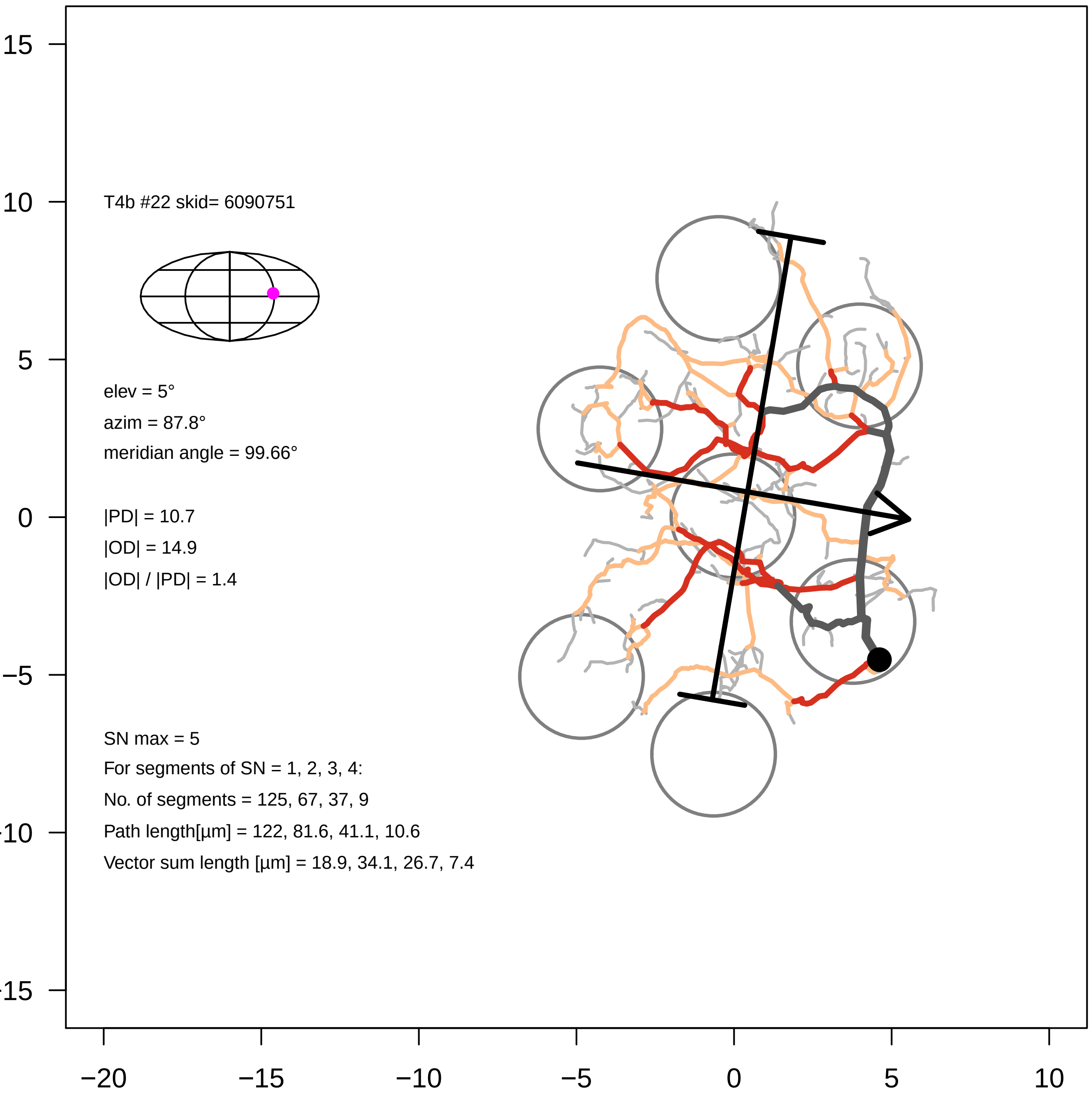

T4b #22 skid= 6090751
elev = 5°
azim = 87.8°
meridian angle = 99.66°
|PD| = 10.7
|OD| = 14.9
|OD| / |PD| = 1.4
SN max = 5
For segments of SN = 1, 2, 3, 4:
No. of segments = 125, 67, 37, 9
Path length[µm] = 122, 81.6, 41.1, 10.6
Vector sum length [µm] = 18.9, 34.1, 26.7, 7.4
15
10
5
0
−5
−10
−15
−20
−15
−10
−5
0
5
10

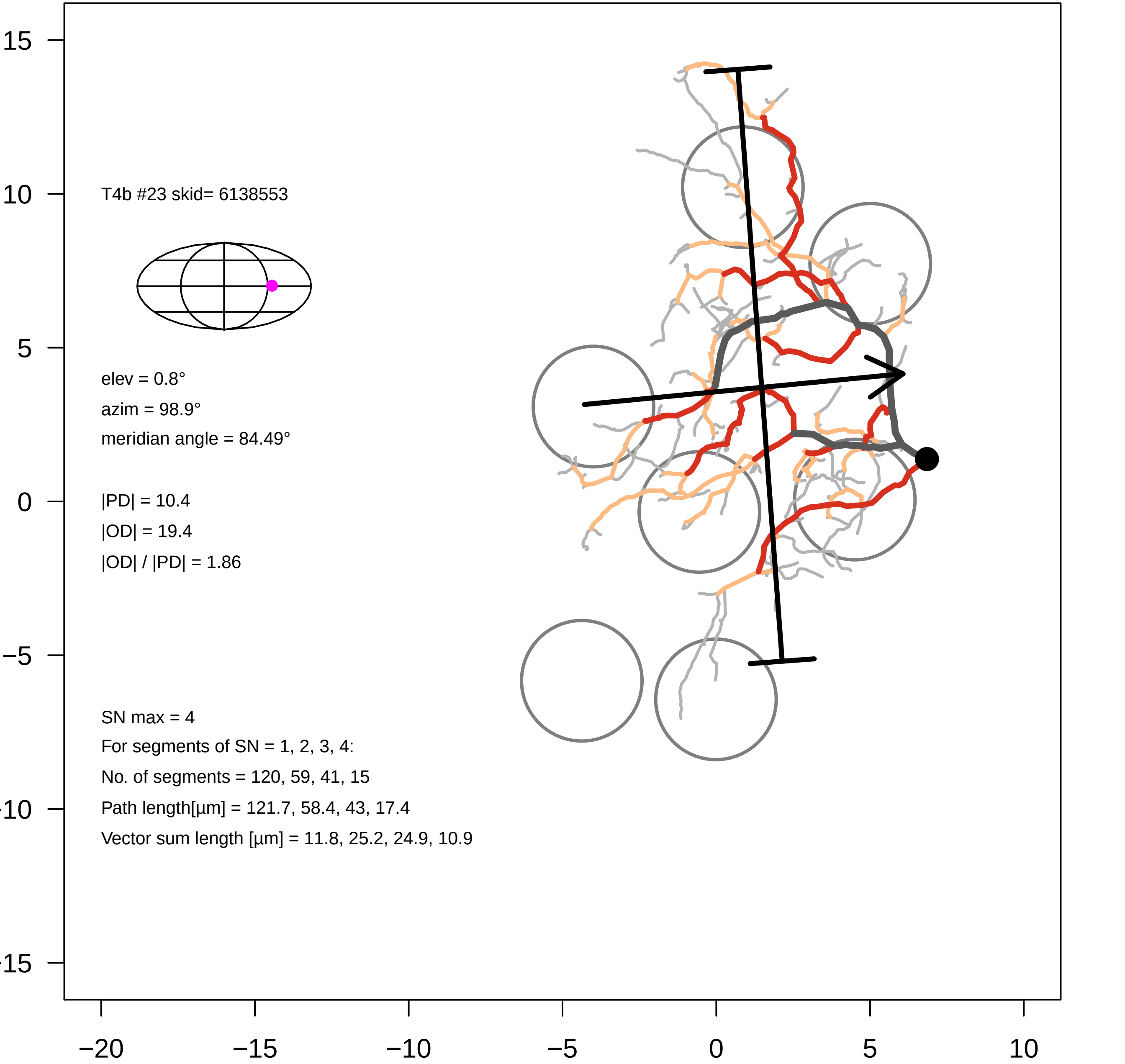
T4b #23 skid= 6138553
elev = 0.8°
azim = 98.9°
meridian angle = 84.49°
|PD| = 10.4
|OD| = 19.4
|OD| / |PD| = 1.86
SN max = 4
For segments of SN = 1, 2, 3, 4:
No. of segments = 120, 59, 41, 15
Path length[µm] = 121.7, 58.4, 43, 17.4
Vector sum length [µm] = 11.8, 25.2, 24.9, 10.9
15
10
5
0
−5
−10
−15
−20
−15
−10
−5
0
5
10

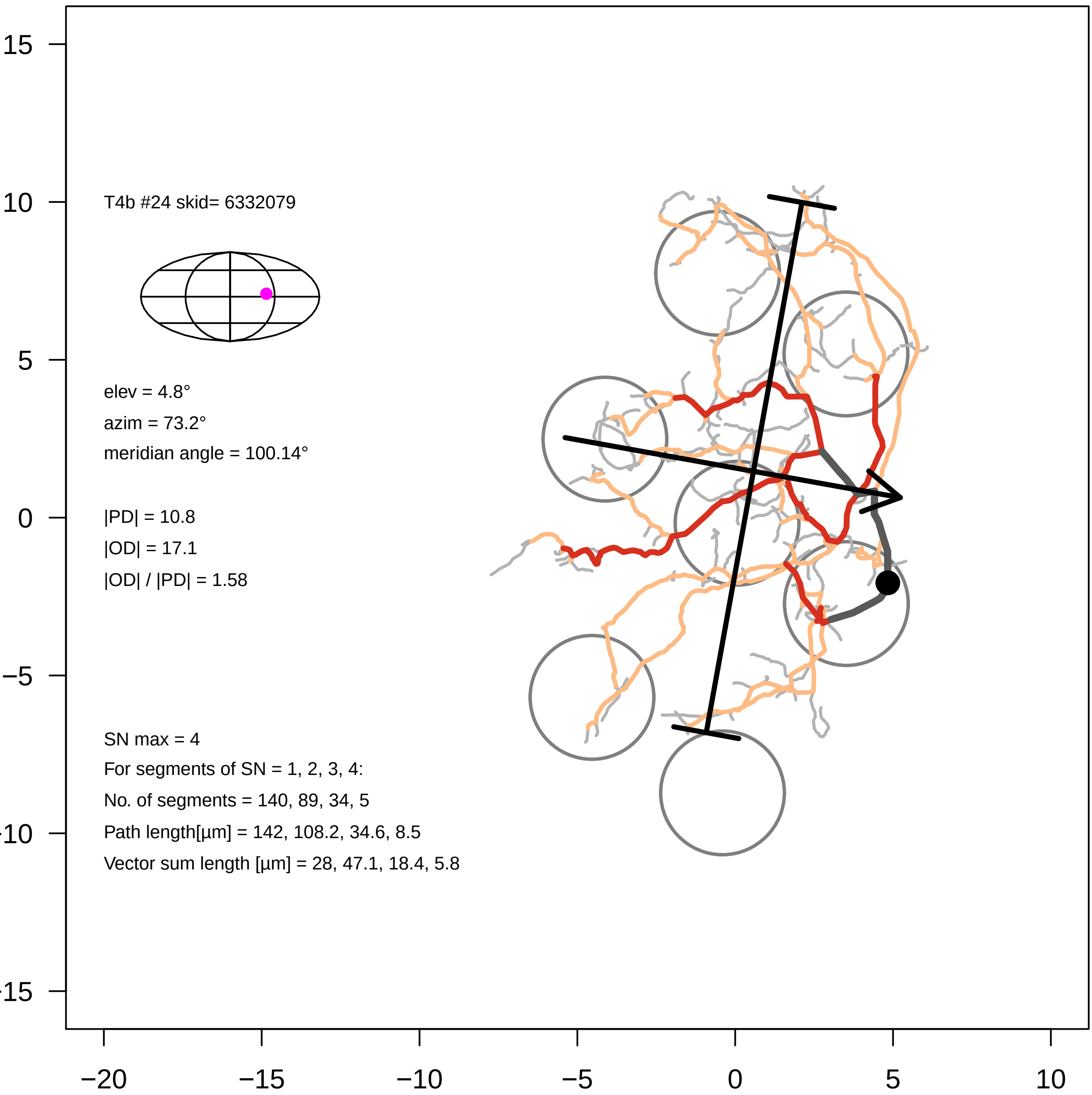
T4b #24 skid= 6332079
elev = 4.8°
azim = 73.2°
meridian angle = 100.14°
|PD| = 10.8
|OD| = 17.1
|OD| / |PD| = 1.58
SN max = 4
For segments of SN = 1, 2, 3, 4:
No. of segments = 140, 89, 34, 5
Path length[µm] = 142, 108.2, 34.6, 8.5
Vector sum length [µm] = 28, 47.1, 18.4, 5.8
15
10
5
0
−5
−10
−15
−20
−15
−10
−5
0
5
10

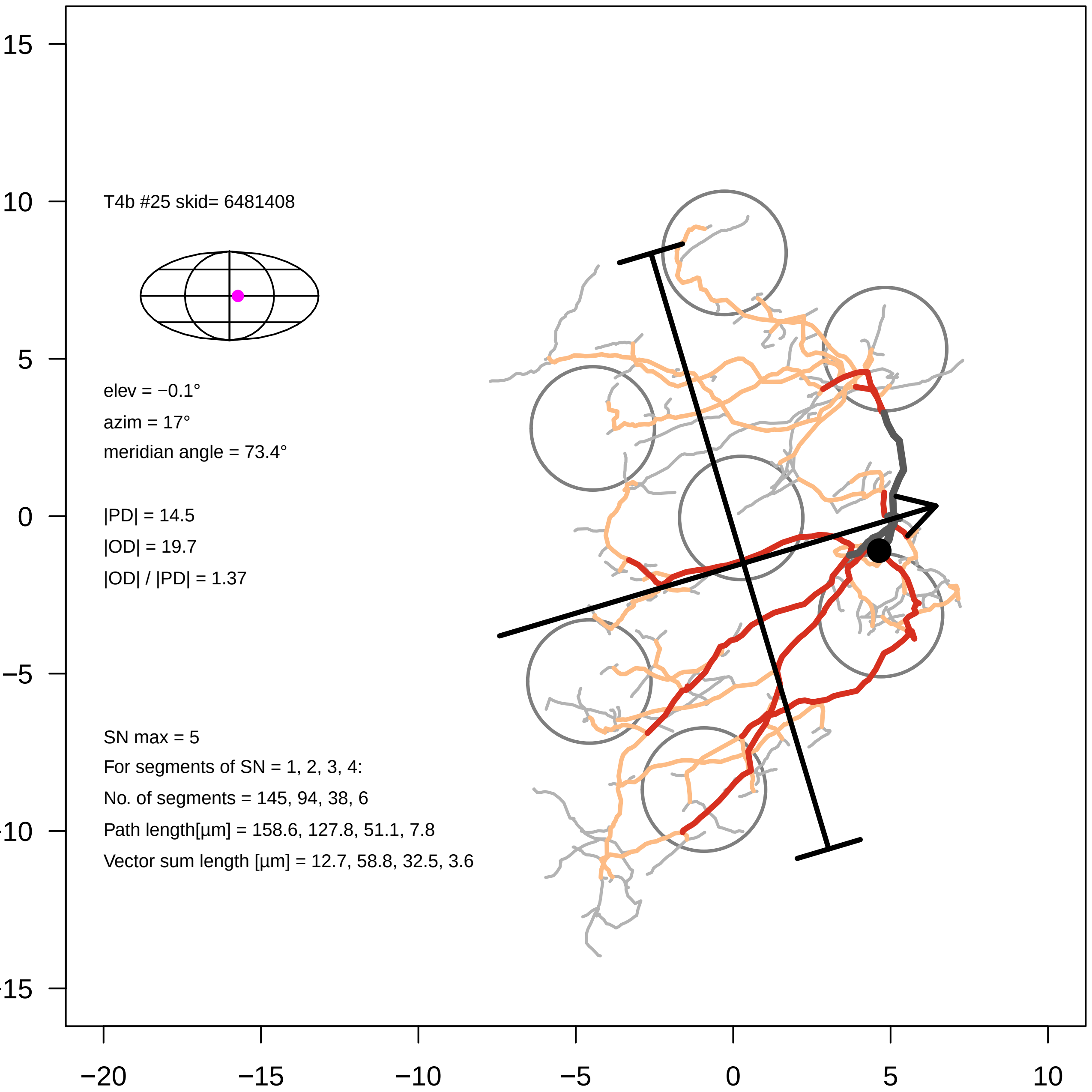
T4b #25 skid= 6481408
elev = −0.1°
azim = 17°
meridian angle = 73.4°
|PD| = 14.5
|OD| = 19.7
|OD| / |PD| = 1.37
SN max = 5
For segments of SN = 1, 2, 3, 4:
No. of segments = 145, 94, 38, 6
Path length[µm] = 158.6, 127.8, 51.1, 7.8
Vector sum length [µm] = 12.7, 58.8, 32.5, 3.6
15
10
5
0
−5
−10
−15
−20
−15
−10
−5
0
5
10

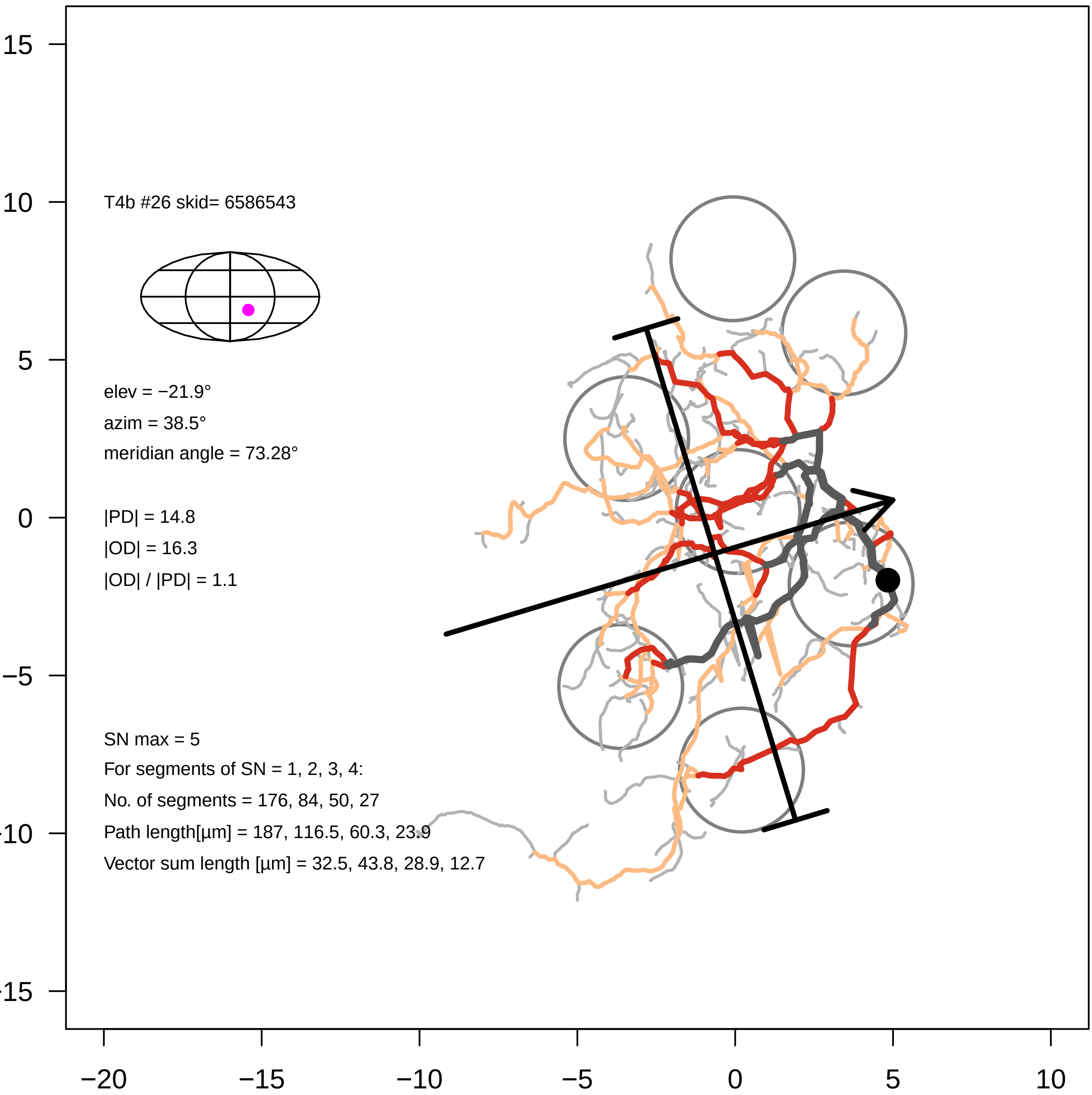

15
10
5
0
−5
-10
-15
−20
−15
−10
−5
0
5
10
T4b #26 skid= 6586543
elev = −21.9°
azim = 38.5°
meridian angle = 73.28°
|PD| = 14.8
|OD| = 16.3
|OD| / |PD| = 1.1
SN max = 5
For segments of SN = 1, 2, 3, 4:
No. of segments = 176, 84, 50, 27
Path length[µm] = 187, 116.5, 60.3, 23.9
Vector sum length [µm] = 32.5, 43.8, 28.9, 12.7

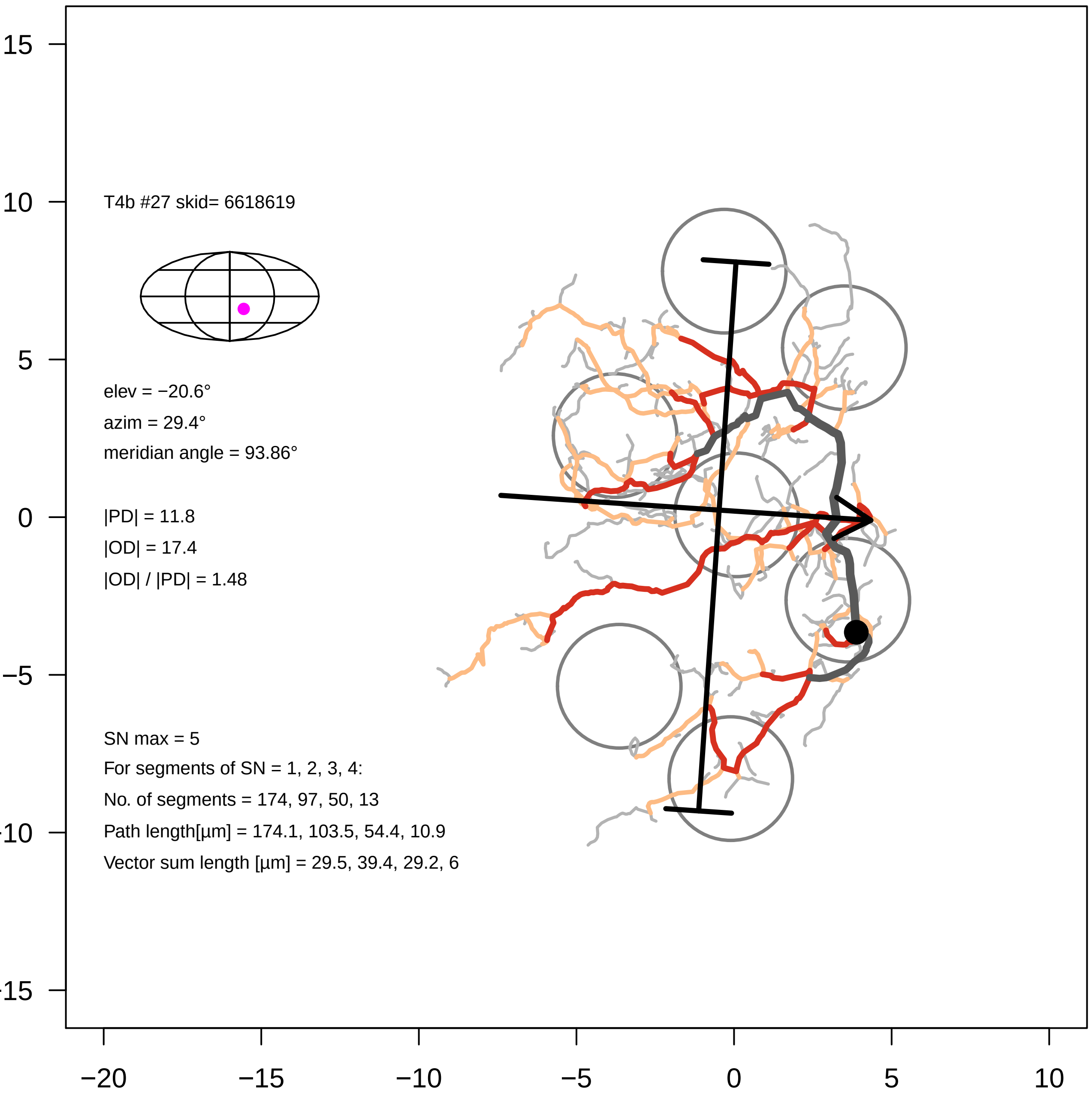

T4b #27 skid= 6618619
elev = −20.6°
azim = 29.4°
meridian angle = 93.86°
|PD| = 11.8
|OD| = 17.4
|OD| / |PD| = 1.48
SN max = 5
For segments of SN = 1, 2, 3, 4:
No. of segments = 174, 97, 50, 13
Path length[µm] = 174.1, 103.5, 54.4, 10.9
Vector sum length [µm] = 29.5, 39.4, 29.2, 6
15
10
5
0
−5
−10
−15
−20
−15
−10
−5
0
5
10

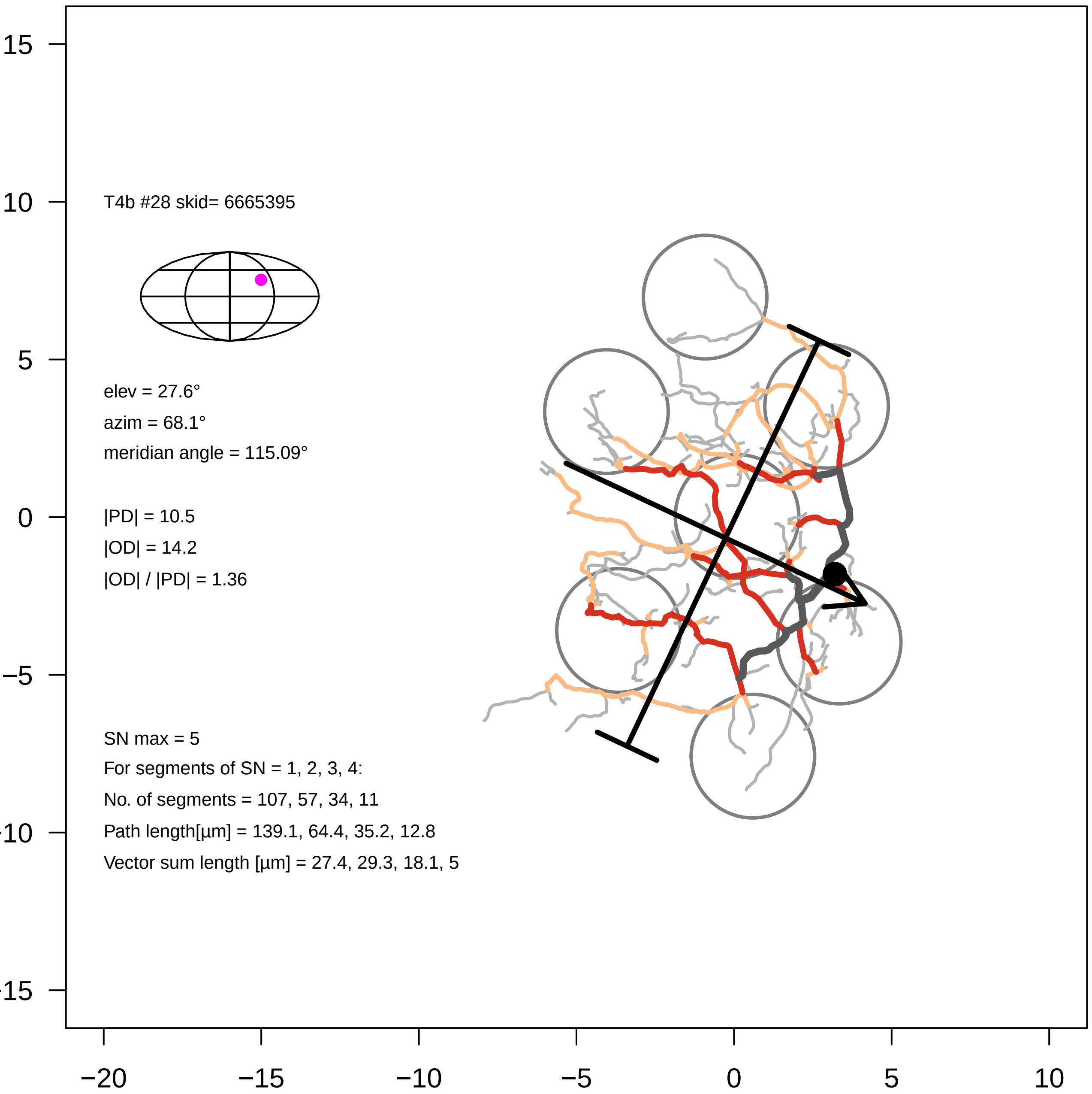

T4b #28 skid= 6665395
elev = 27.6°
azim = 68.1°
meridian angle = 115.09°
|PD| = 10.5
|OD| = 14.2
|OD| / |PD| = 1.36
SN max = 5
For segments of SN = 1, 2, 3, 4:
No. of segments = 107, 57, 34, 11
Path length[µm] = 139.1, 64.4, 35.2, 12.8
Vector sum length [µm] = 27.4, 29.3, 18.1, 5
15
10
5
0
−5
−10
−15
−20
−15
−10
−5
0
5
10

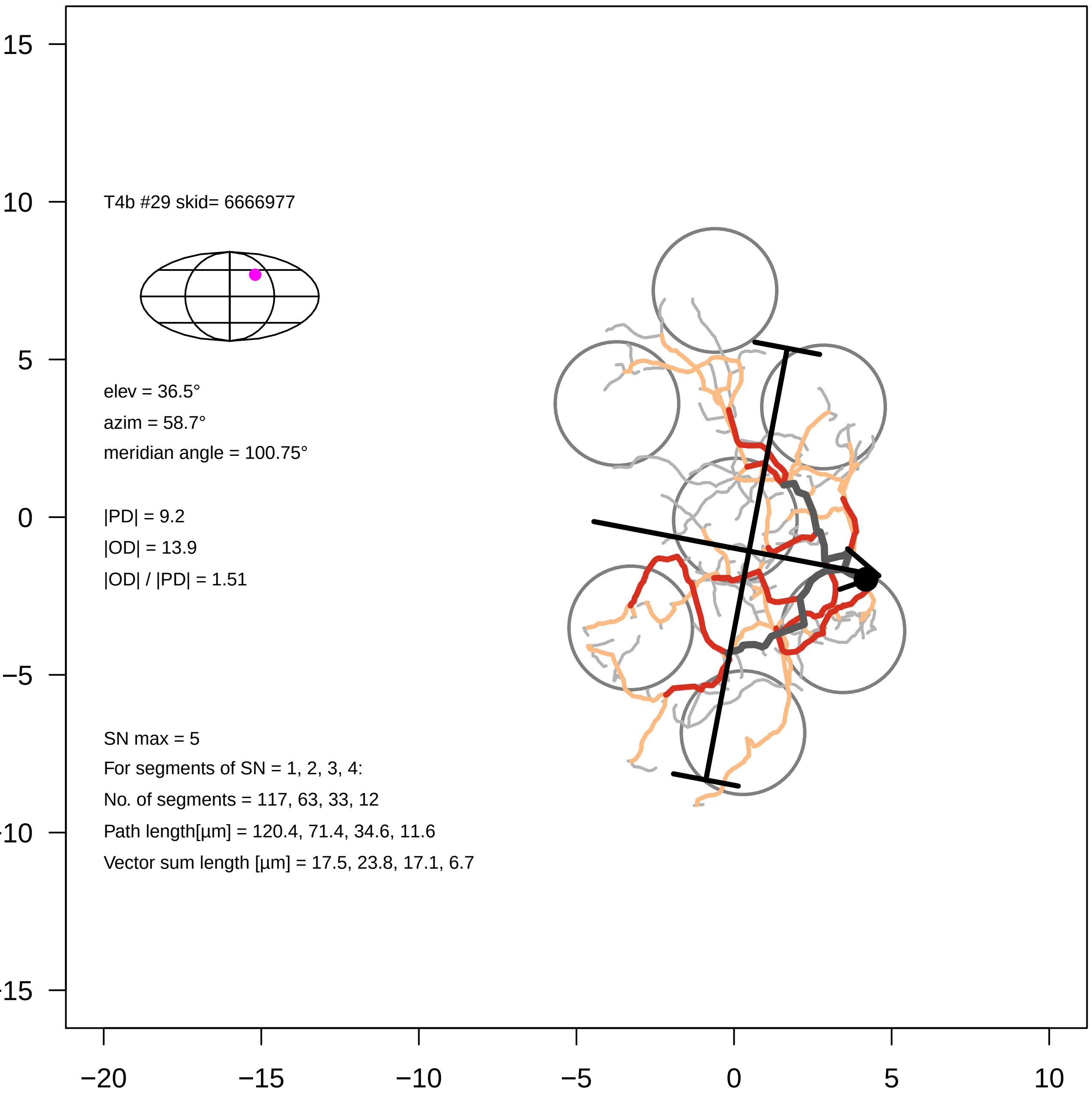
T4b #29 skid= 6666977
elev = 36.5°
azim = 58.7°
meridian angle = 100.75°
|PD| = 9.2
|OD| = 13.9
|OD| / |PD| = 1.51
SN max = 5
For segments of SN = 1, 2, 3, 4:
No. of segments = 117, 63, 33, 12
Path length[µm] = 120.4, 71.4, 34.6, 11.6
Vector sum length [µm] = 17.5, 23.8, 17.1, 6.7
15
10
5
0
−5
−10
−15
−20
−15
−10
−5
0
5
10

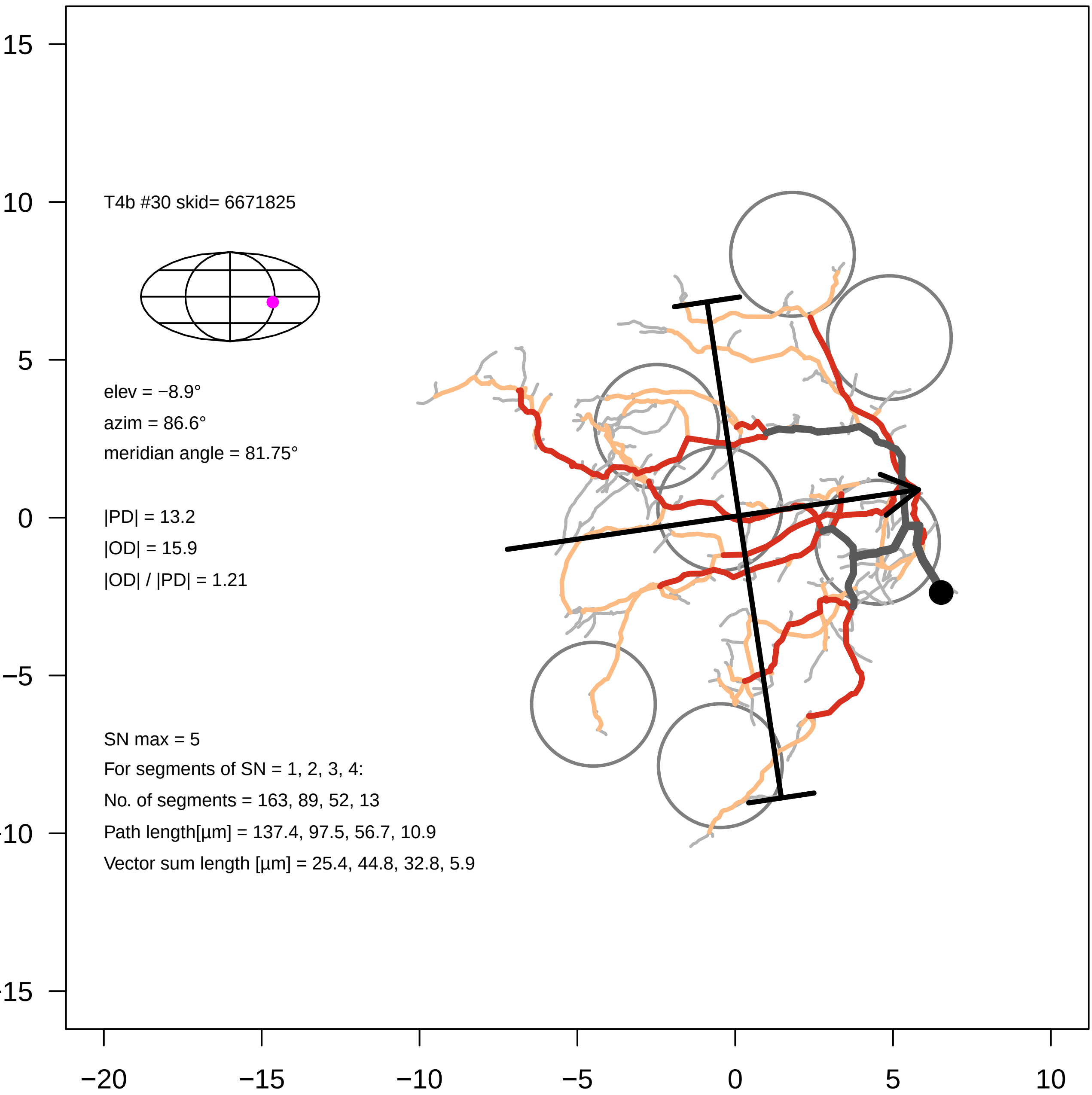

T4b #30 skid= 6671825
elev = −8.9°
azim = 86.6°
meridian angle = 81.75°
|PD| = 13.2
|OD| = 15.9
|OD| / |PD| = 1.21
SN max = 5
For segments of SN = 1, 2, 3, 4:
No. of segments = 163, 89, 52, 13
Path length[µm] = 137.4, 97.5, 56.7, 10.9
Vector sum length [µm] = 25.4, 44.8, 32.8, 5.9
15
10
5
0
−5
−10
−15
−20
−15
−10
−5
0
5
10

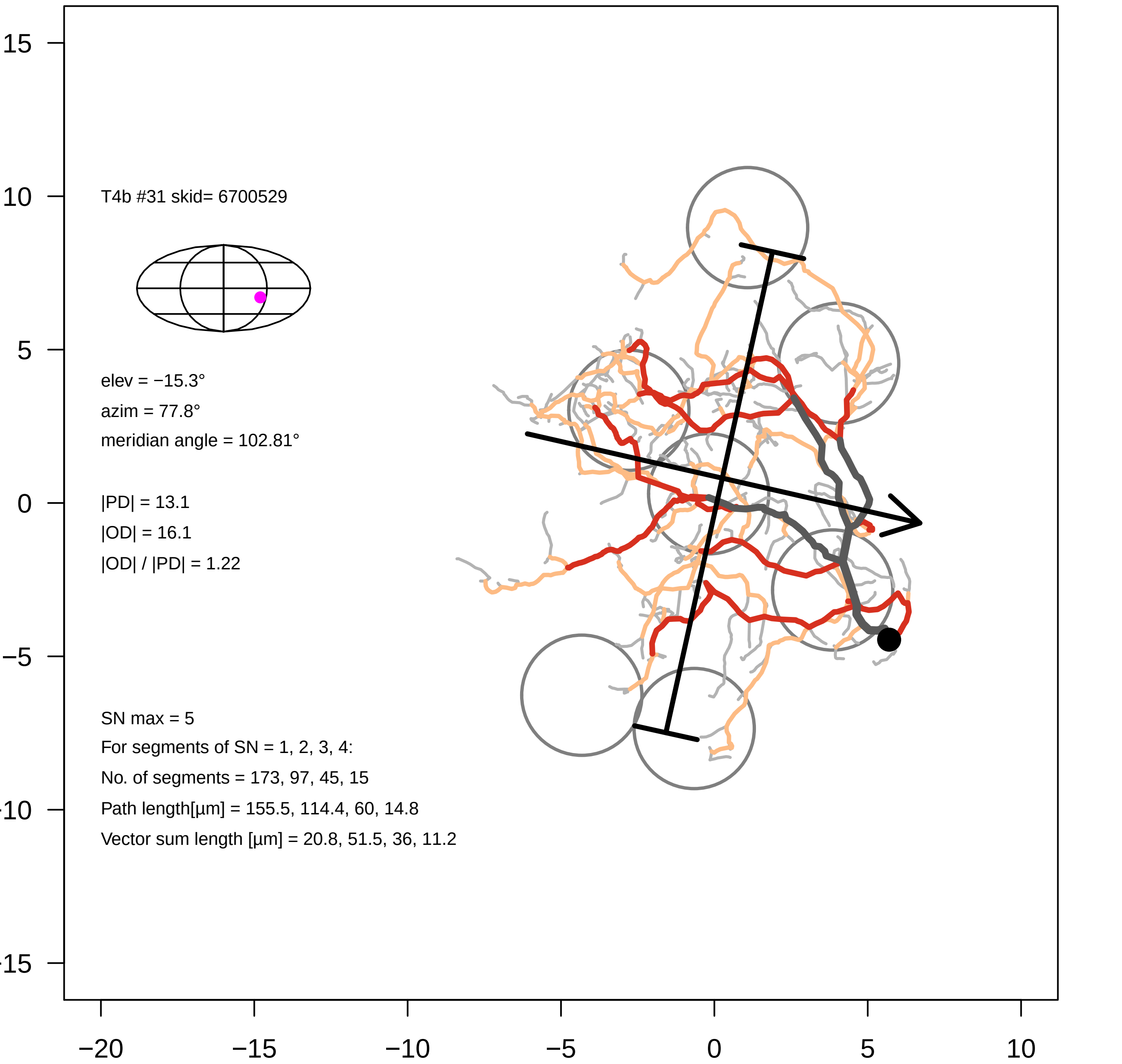
T4b #31 skid= 6700529
elev = −15.3°
azim = 77.8°
meridian angle = 102.81°
|PD| = 13.1
|OD| = 16.1
|OD| / |PD| = 1.22
SN max = 5
For segments of SN = 1, 2, 3, 4:
No. of segments = 173, 97, 45, 15
Path length[µm] = 155.5, 114.4, 60, 14.8
Vector sum length [µm] = 20.8, 51.5, 36, 11.2
15
10
5
0
−5
−10
−15
−20
−15
−10
−5
0
5
10

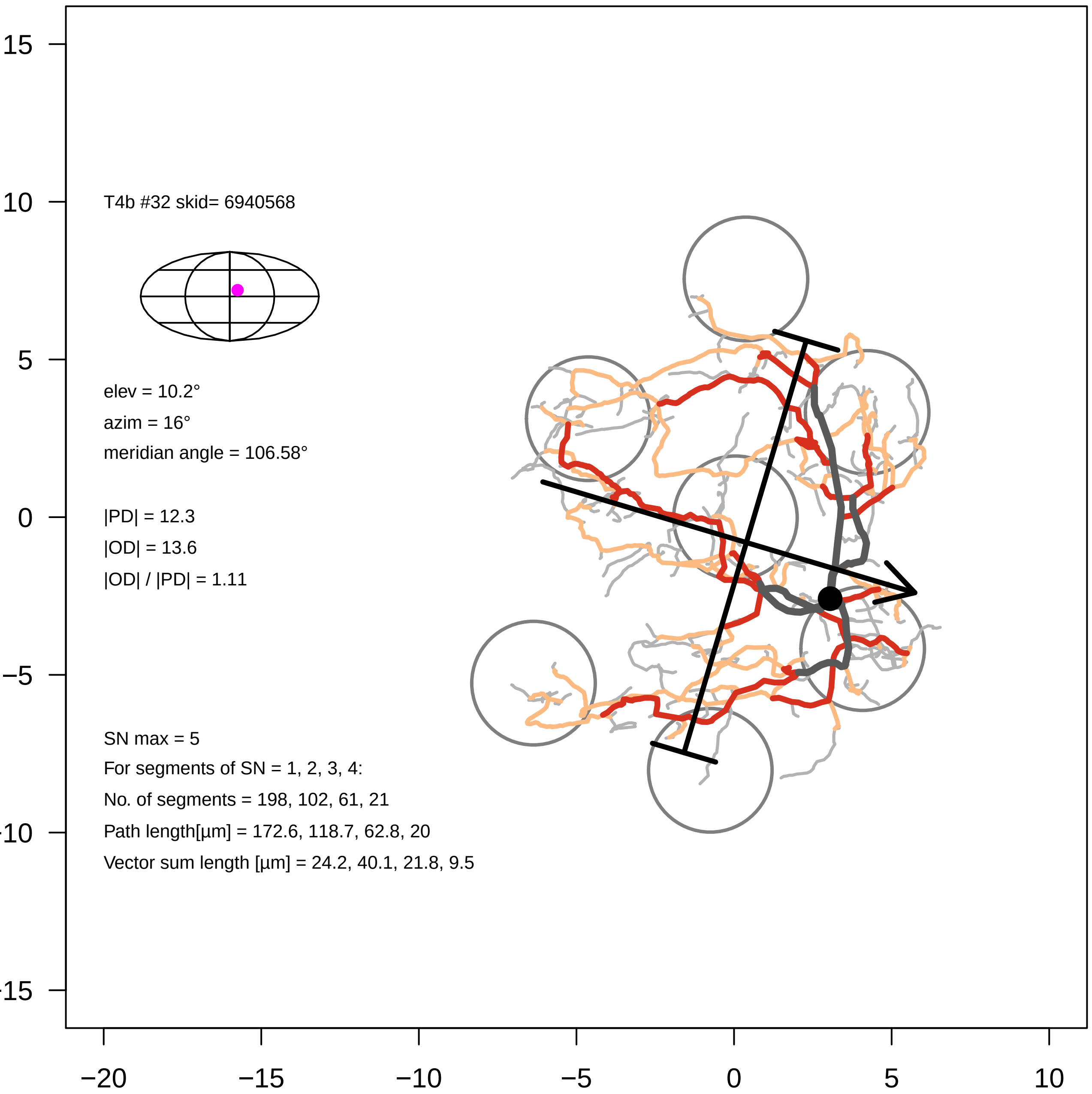
T4b #32 skid= 6940568
elev = 10.2°
azim = 16°
meridian angle = 106.58°
|PD| = 12.3
|OD| = 13.6
|OD| / |PD| = 1.11
SN max = 5
For segments of SN = 1, 2, 3, 4:
No. of segments = 198, 102, 61, 21
Path length[µm] = 172.6, 118.7, 62.8, 20
Vector sum length [µm] = 24.2, 40.1, 21.8, 9.5
15
10
5
0
−5
−10
−15
−20
−15
−10
−5
0
5
10

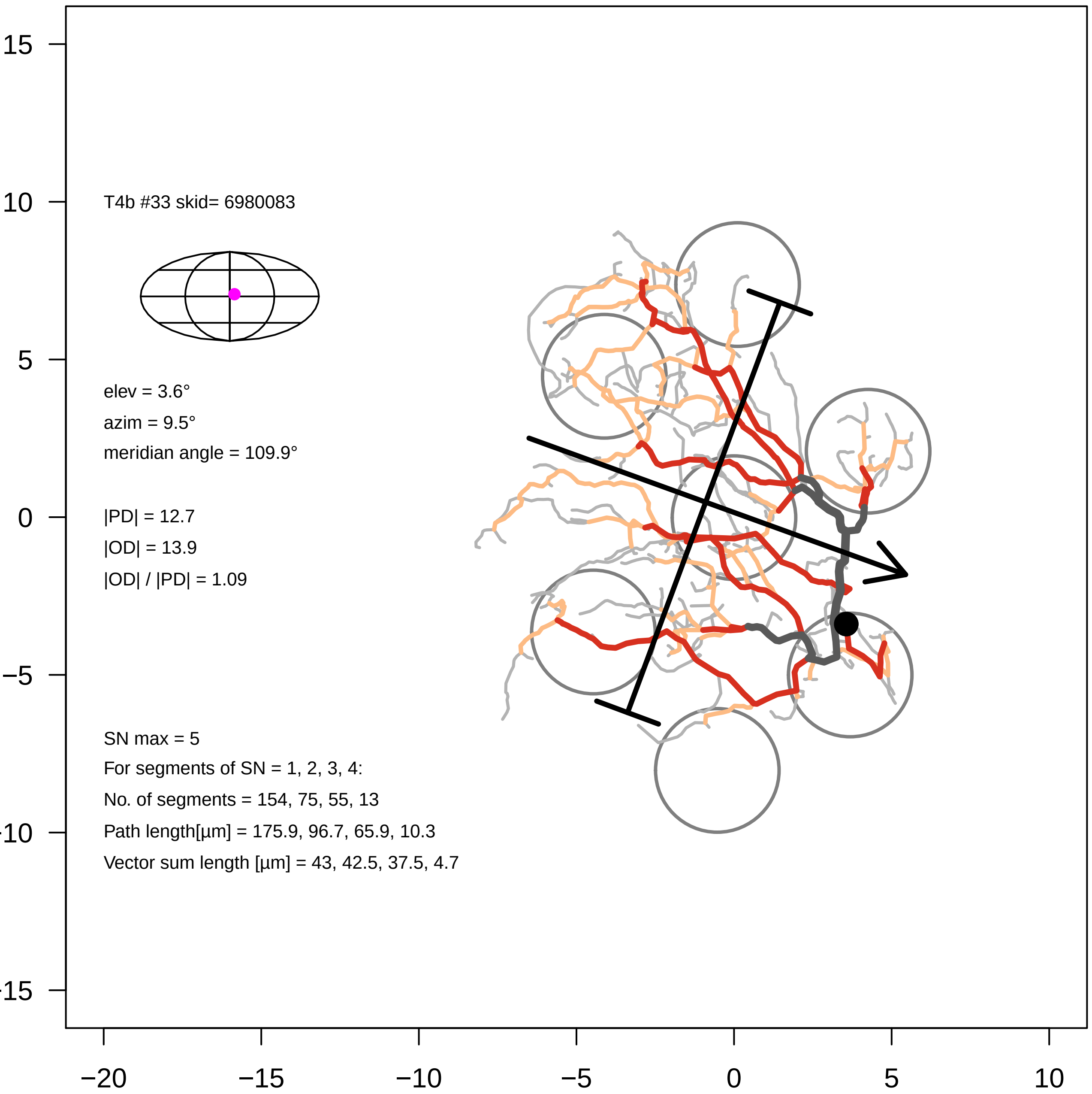

T4b #33 skid= 6980083
elev = 3.6°
azim = 9.5°
meridian angle = 109.9°
|PD| = 12.7
|OD| = 13.9
|OD| / |PD| = 1.09
SN max = 5
For segments of SN = 1, 2, 3, 4:
No. of segments = 154, 75, 55, 13
Path length[µm] = 175.9, 96.7, 65.9, 10.3
Vector sum length [µm] = 43, 42.5, 37.5, 4.7
15
10
5
0
−5
−20
−15
−10
0
5
10

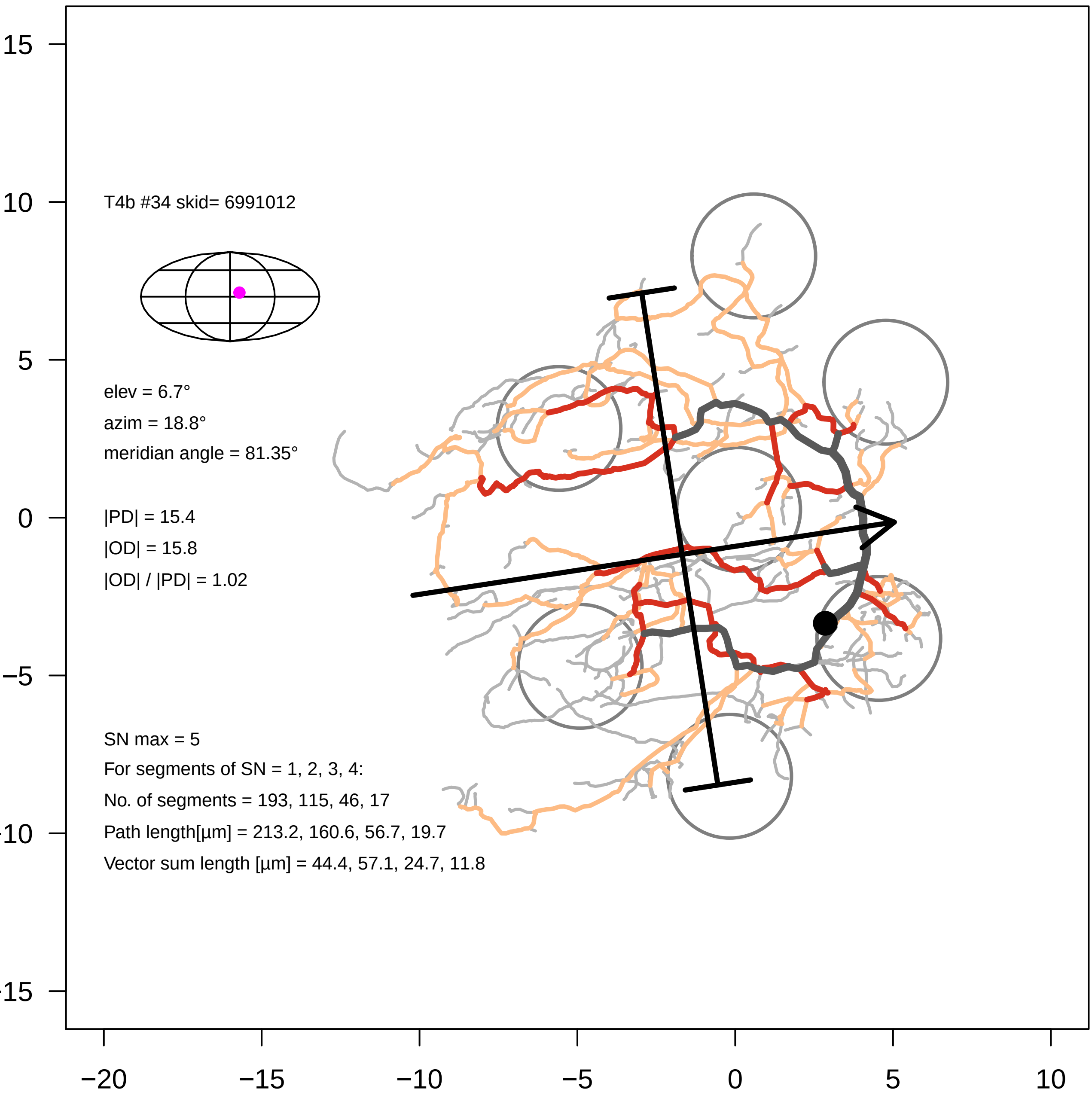

T4b #34 skid= 6991012
elev = 6.7°
azim = 18.8°
meridian angle = 81.35°
|PD| = 15.4
|OD| = 15.8
|OD| / |PD| = 1.02
SN max = 5
For segments of SN = 1, 2, 3, 4:
No. of segments = 193, 115, 46, 17
Path length[µm] = 213.2, 160.6, 56.7, 19.7
Vector sum length [µm] = 44.4, 57.1, 24.7, 11.8
15
10
5
0
−5
−10
−15
−20
−15
−10
−5
0
5
10

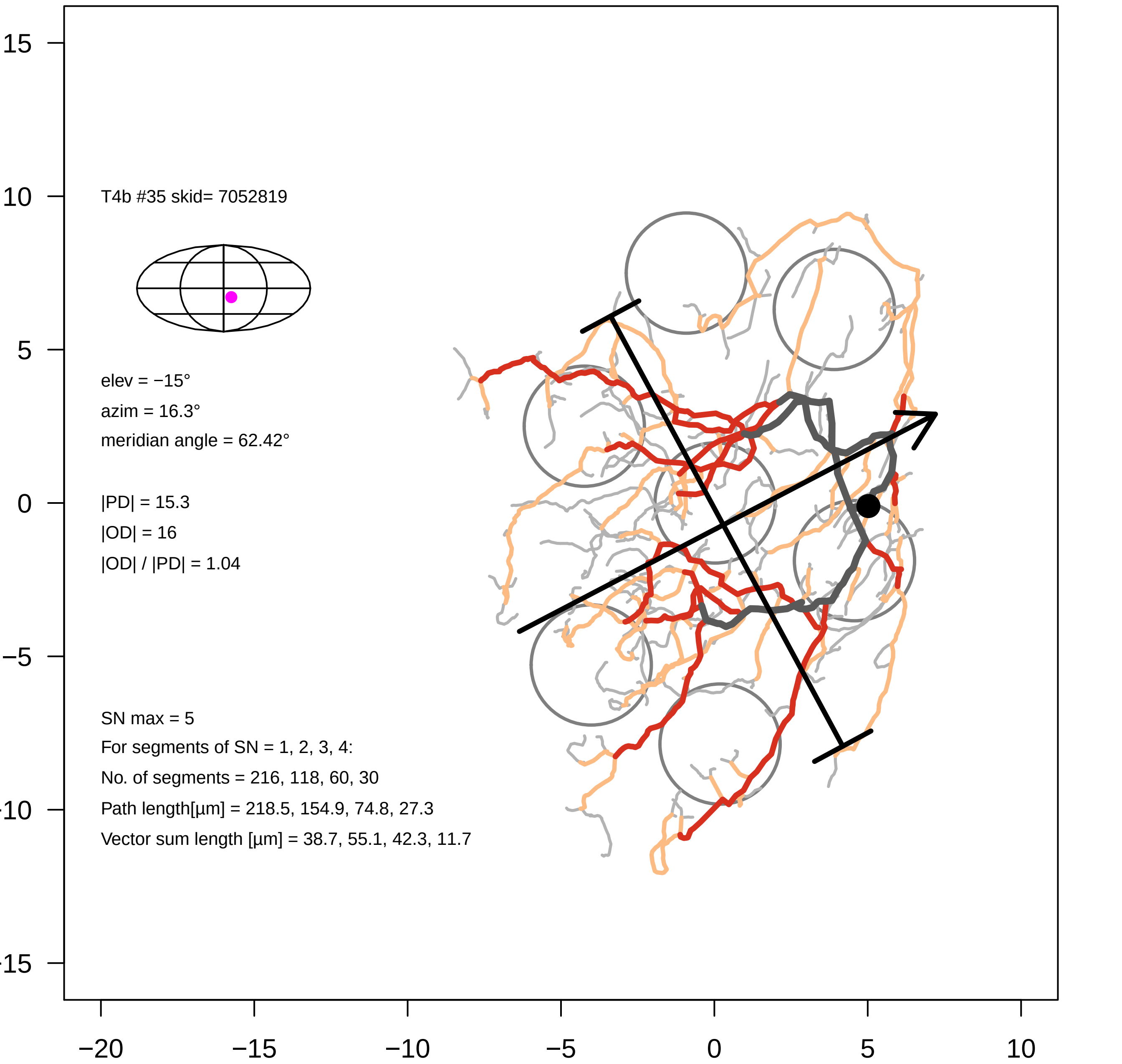

T4b #35 skid= 7052819
elev = −15°
azim = 16.3°
meridian angle = 62.42°
|PD| = 15.3
|OD| = 16
|OD| / |PD| = 1.04
SN max = 5
For segments of SN = 1, 2, 3, 4:
No. of segments = 216, 118, 60, 30
Path length[µm] = 218.5, 154.9, 74.8, 27.3
Vector sum length [µm] = 38.7, 55.1, 42.3, 11.7
15
10
5
0
−5
−10
−15
−20
−15
−10
−5
0
5
10

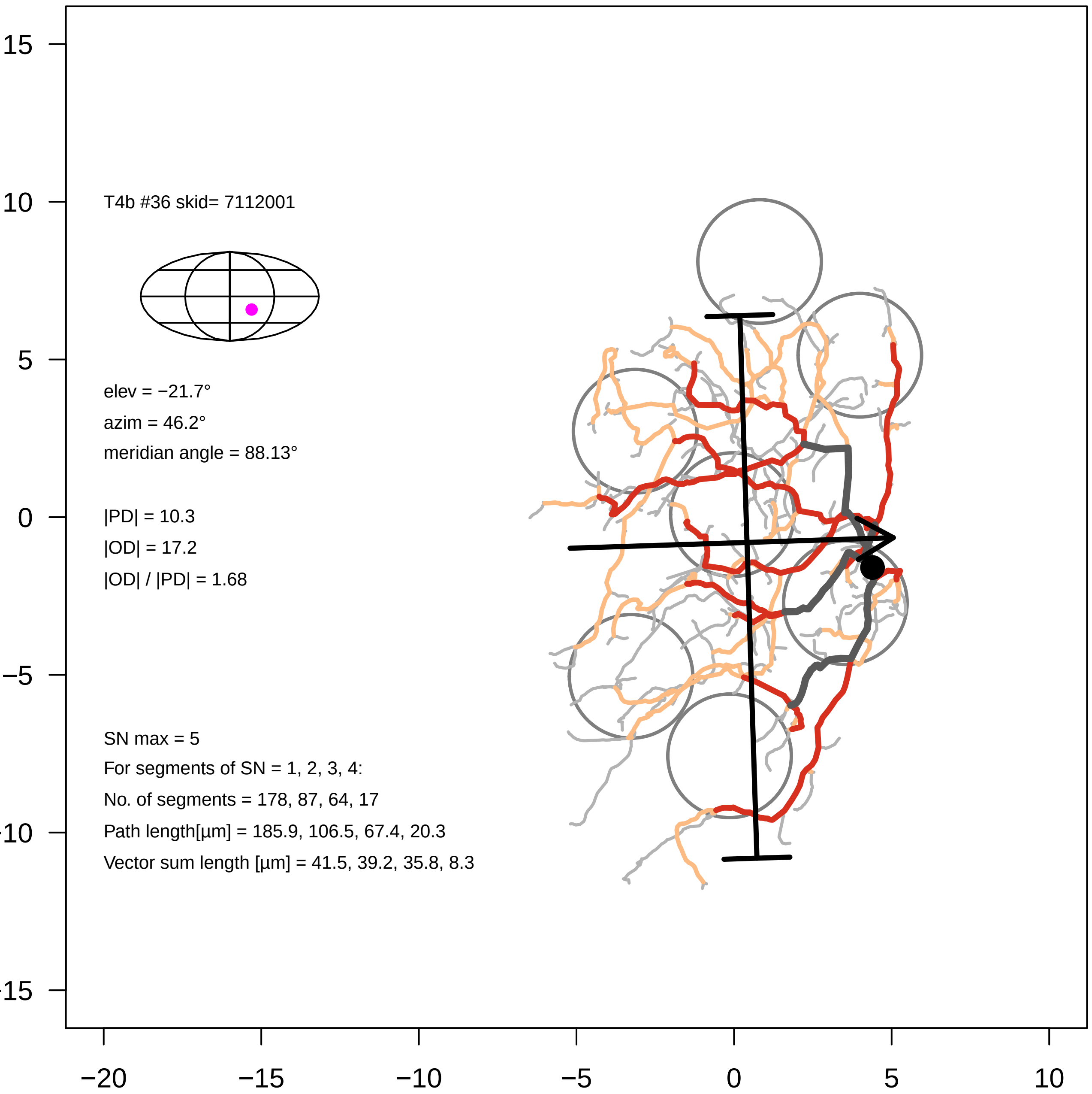

T4b #36 skid= 7112001
elev = −21.7°
azim = 46.2°
meridian angle = 88.13°
|PD| = 10.3
|OD| = 17.2
|OD| / |PD| = 1.68
SN max = 5
For segments of SN = 1, 2, 3, 4:
No. of segments = 178, 87, 64, 17
Path length[µm] = 185.9, 106.5, 67.4, 20.3
Vector sum length [µm] = 41.5, 39.2, 35.8, 8.3
15
10
5
0
−5
−10
−15
−20
−15
−10
−5
0
5
10

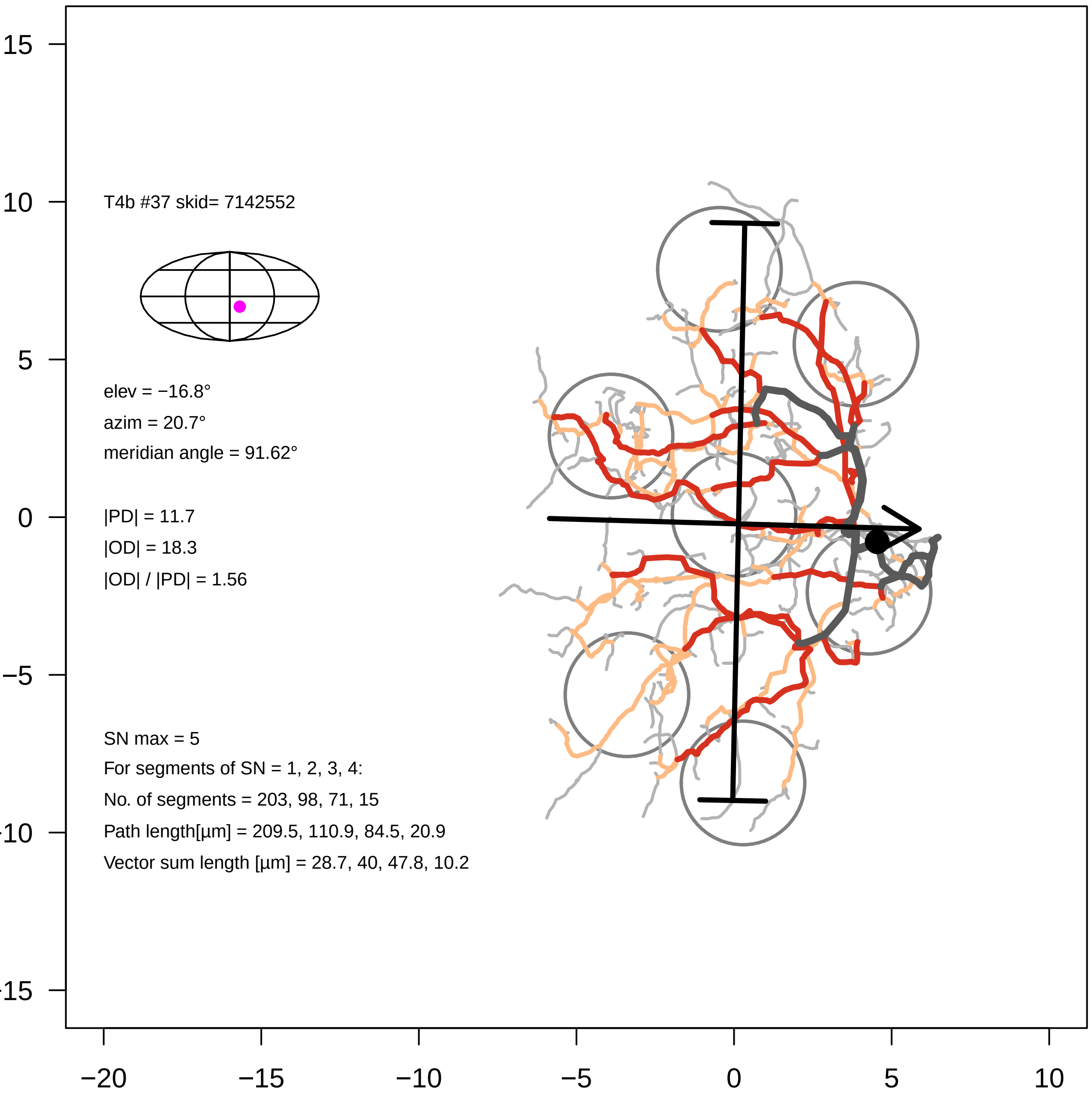

T4b #37 skid= 7142552
elev = −16.8°
azim = 20.7°
meridian angle = 91.62°
|PD| = 11.7
|OD| = 18.3
|OD| / |PD| = 1.56
SN max = 5
For segments of SN = 1, 2, 3, 4:
No. of segments = 203, 98, 71, 15
Path length[µm] = 209.5, 110.9, 84.5, 20.9
Vector sum length [µm] = 28.7, 40, 47.8, 10.2
15
10
5
0
−5
−10
−15
−20
−15
−10
−5
0
5
10

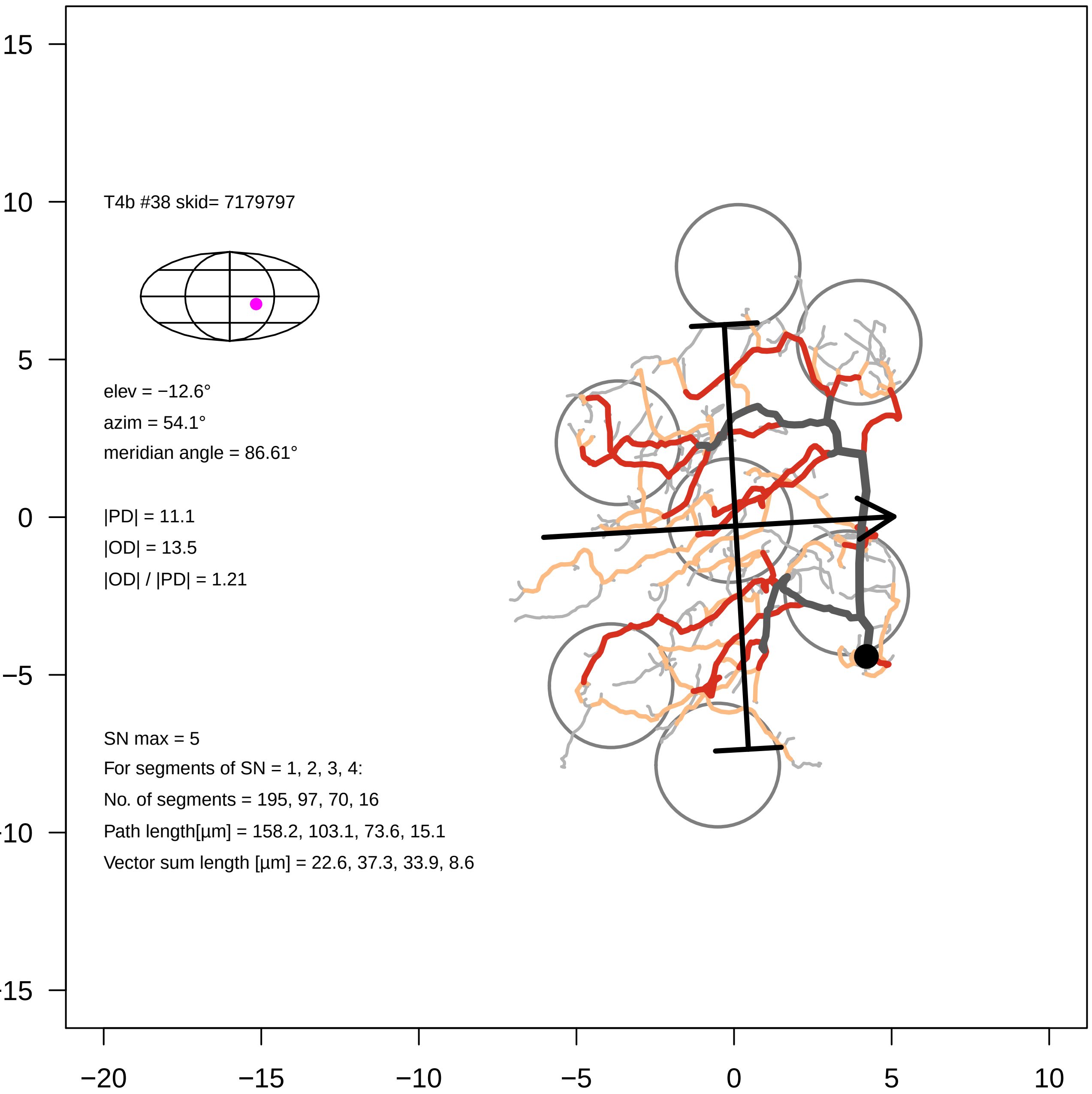

T4b #38 skid= 7179797
elev = −12.6°
azim = 54.1°
meridian angle = 86.61°
|PD| = 11.1
|OD| = 13.5
|OD| / |PD| = 1.21
SN max = 5
For segments of SN = 1, 2, 3, 4:
No. of segments = 195, 97, 70, 16
Path length[µm] = 158.2, 103.1, 73.6, 15.1
Vector sum length [µm] = 22.6, 37.3, 33.9, 8.6
15
10
5
0
−5
−10
−15
−20
−15
−10
−5
0
5
10

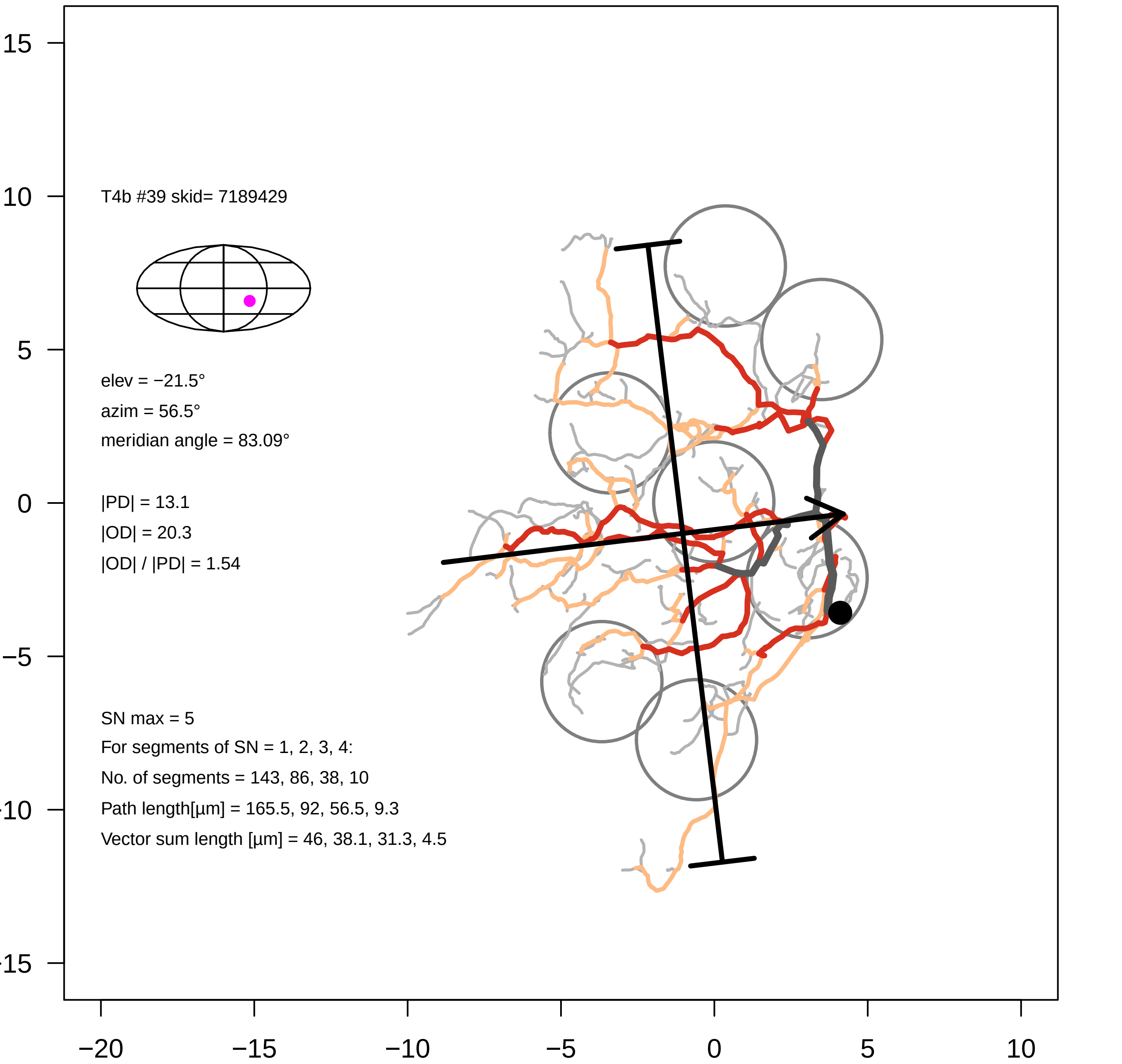

T4b #39 skid= 7189429
elev = −21.5°
azim = 56.5°
meridian angle = 83.09°
|PD| = 13.1
|OD| = 20.3
|OD| / |PD| = 1.54
SN max = 5
For segments of SN = 1, 2, 3, 4:
No. of segments = 143, 86, 38, 10
Path length[µm] = 165.5, 92, 56.5, 9.3
Vector sum length [µm] = 46, 38.1, 31.3, 4.5
15
10
5
0
−5
−10
−15
−20
−15
−10
−5
0
5
10

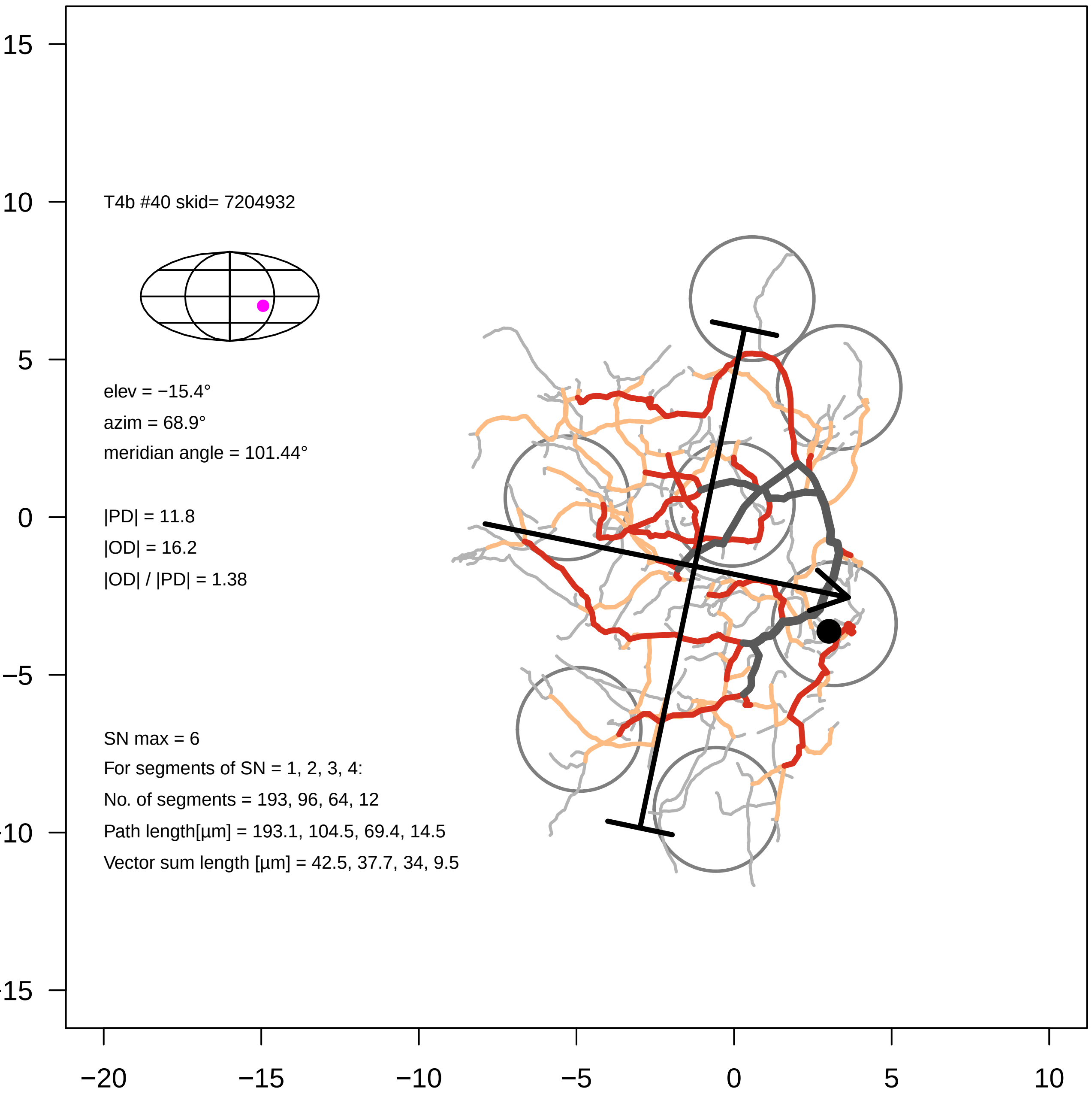
15
10
5
0
-5
-10
-15
−20
−15
−10
−5
0
5
10
T4b #40 skid= 7204932
elev = −15.4°
azim = 68.9°
meridian angle = 101.44°
|PD| = 11.8
|OD| = 16.2
|OD| / |PD| = 1.38
SN max = 6
For segments of SN = 1, 2, 3, 4:
No. of segments = 193, 96, 64, 12
Path length[µm] = 193.1, 104.5, 69.4, 14.5
Vector sum length [µm] = 42.5, 37.7, 34, 9.5

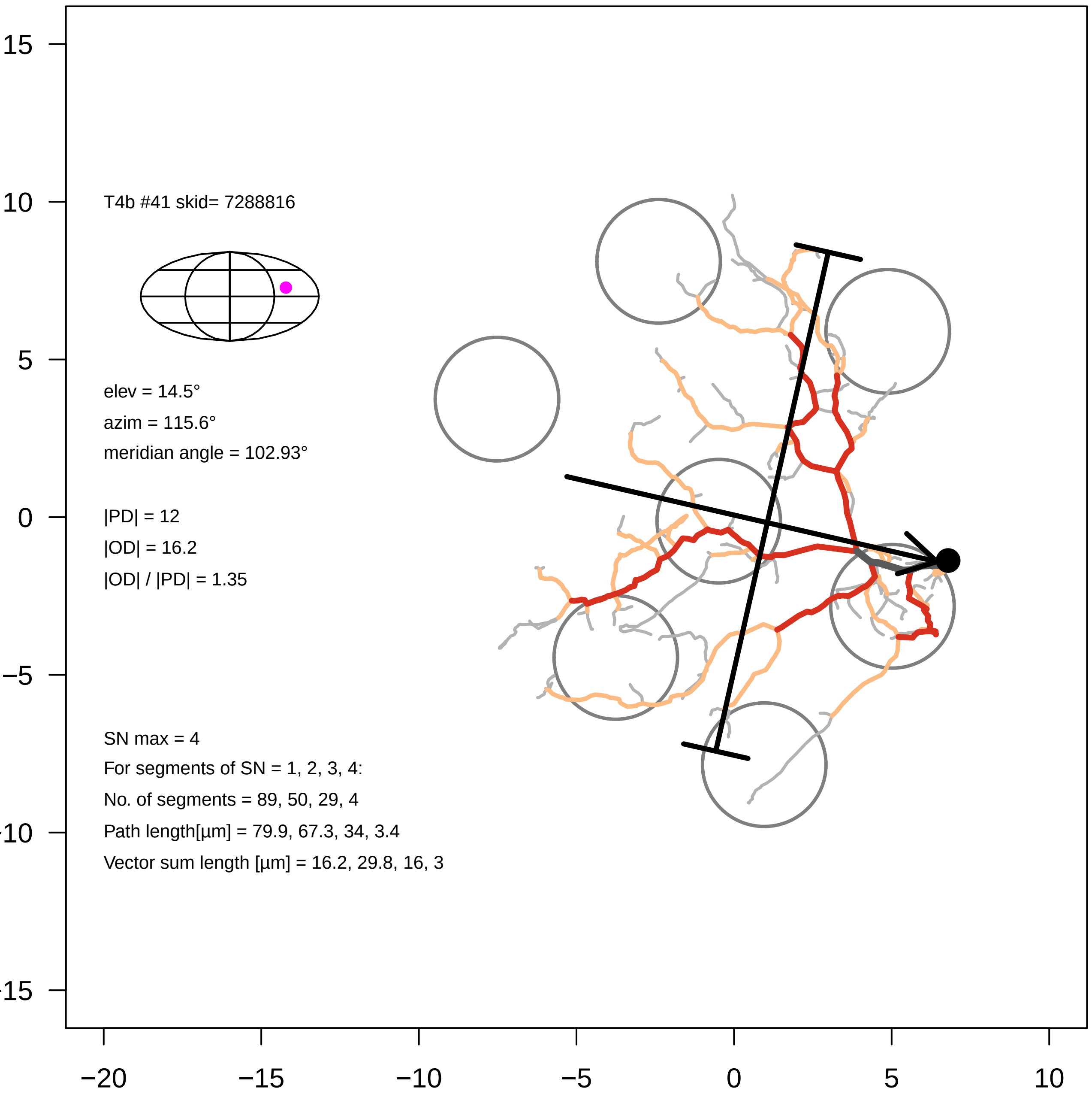

T4b #41 skid= 7288816
elev = 14.5°
azim = 115.6°
meridian angle = 102.93°
|PD| = 12
|OD| = 16.2
|OD| / |PD| = 1.35
SN max = 4
For segments of SN = 1, 2, 3, 4:
No. of segments = 89, 50, 29, 4
Path length[µm] = 79.9, 67.3, 34, 3.4
Vector sum length [µm] = 16.2, 29.8, 16, 3
15
10
5
0
−5
−10
−15
−20
−15
−10
−5
0
5
10

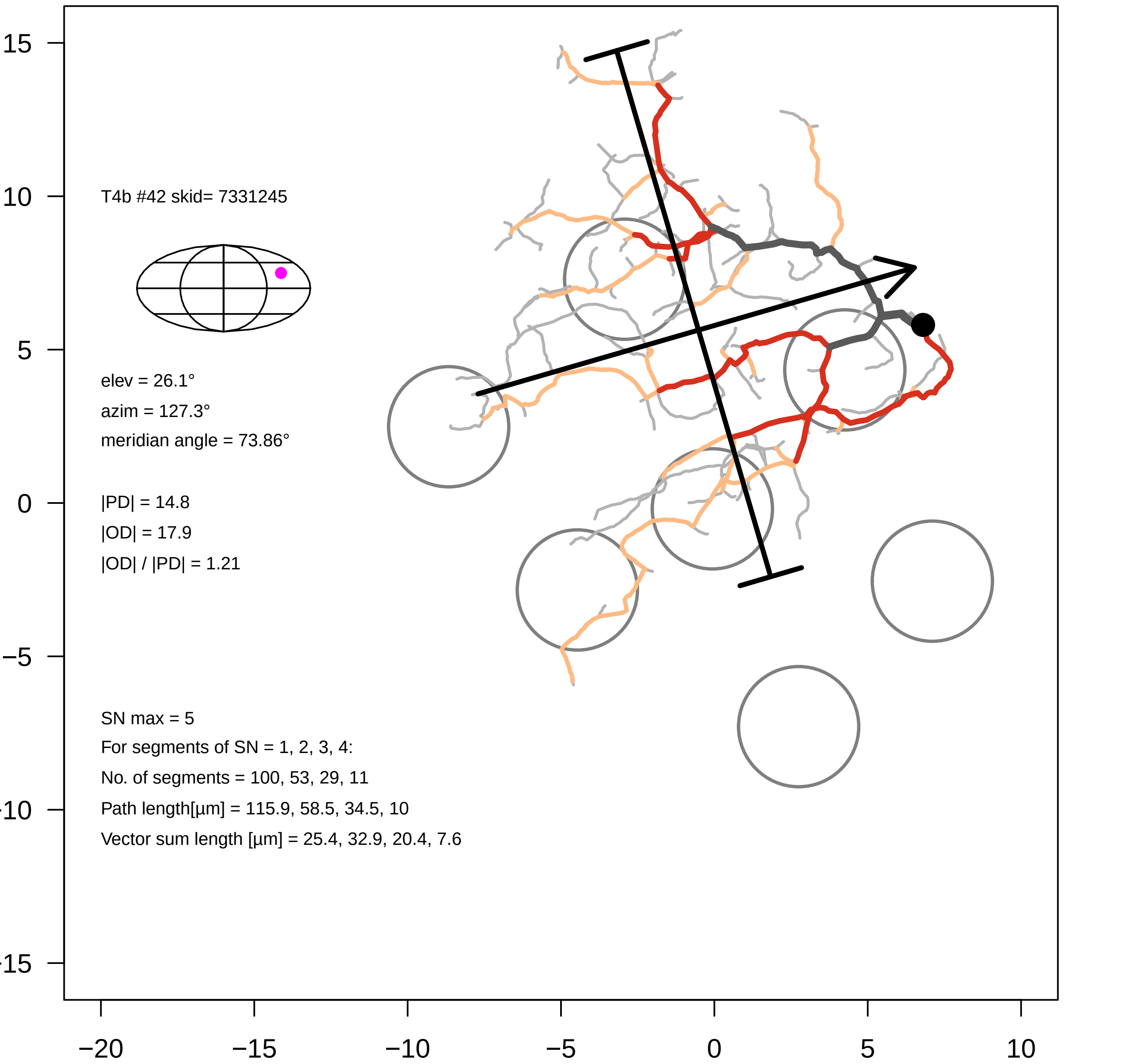
T4b #42 skid= 7331245
elev = 26.1°
azim = 127.3°
meridian angle = 73.86°
|PD| = 14.8
|OD| = 17.9
|OD| / |PD| = 1.21
SN max = 5
For segments of SN = 1, 2, 3, 4:
No. of segments = 100, 53, 29, 11
Path length[µm] = 115.9, 58.5, 34.5, 10
Vector sum length [µm] = 25.4, 32.9, 20.4, 7.6
15
10
5
0
−5
−10
−15
−20
−15
−10
−5
0
5
10

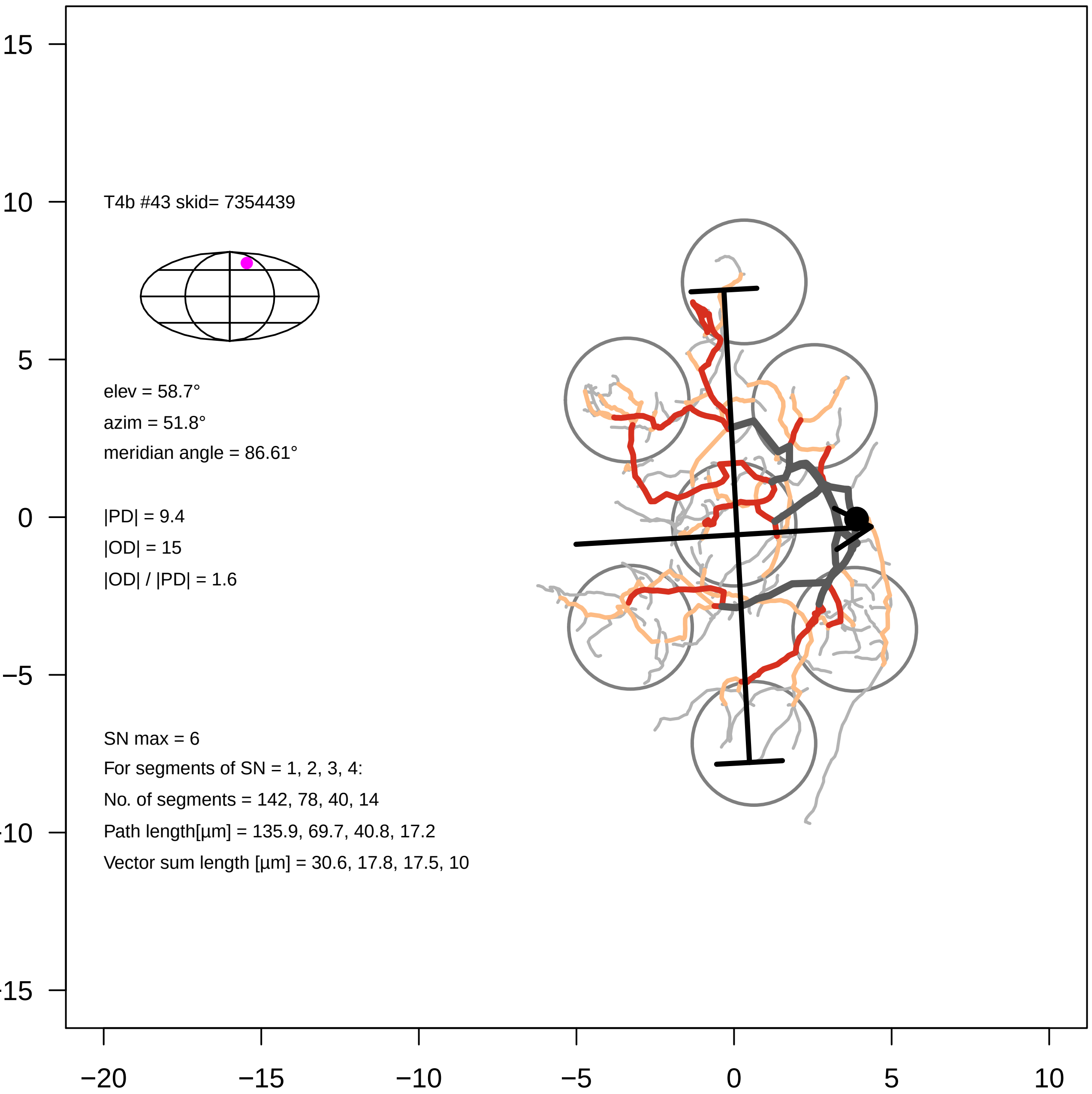
T4b #43 skid= 7354439
elev = 58.7°
azim = 51.8°
meridian angle = 86.61°
|PD| = 9.4
|OD| = 15
|OD| / |PD| = 1.6
SN max = 6
For segments of SN = 1, 2, 3, 4:
No. of segments = 142, 78, 40, 14
Path length[µm] = 135.9, 69.7, 40.8, 17.2
Vector sum length [µm] = 30.6, 17.8, 17.5, 10
15
10
5
0
−5
−10
−15
−20
−15
−10
−5
0
5
10

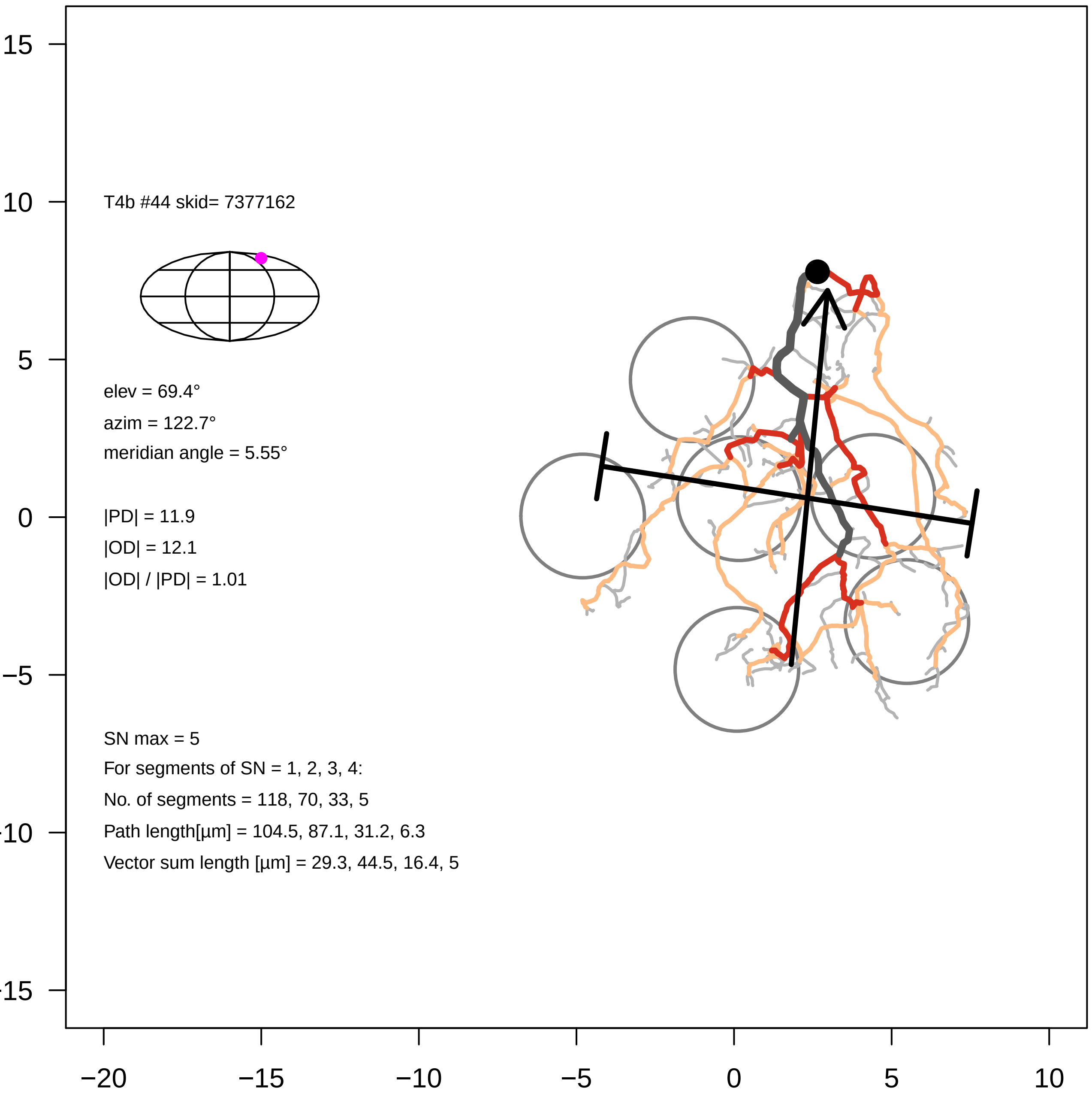

T4b #44 skid= 7377162
elev = 69.4°
azim = 122.7°
meridian angle = 5.55°
|PD| = 11.9
|OD| = 12.1
|OD| / |PD| = 1.01
SN max = 5
For segments of SN = 1, 2, 3, 4:
No. of segments = 118, 70, 33, 5
Path length[µm] = 104.5, 87.1, 31.2, 6.3
Vector sum length [µm] = 29.3, 44.5, 16.4, 5
15
10
5
0
−5
−10
−15
−20
−15
−10
−5
0
5
10

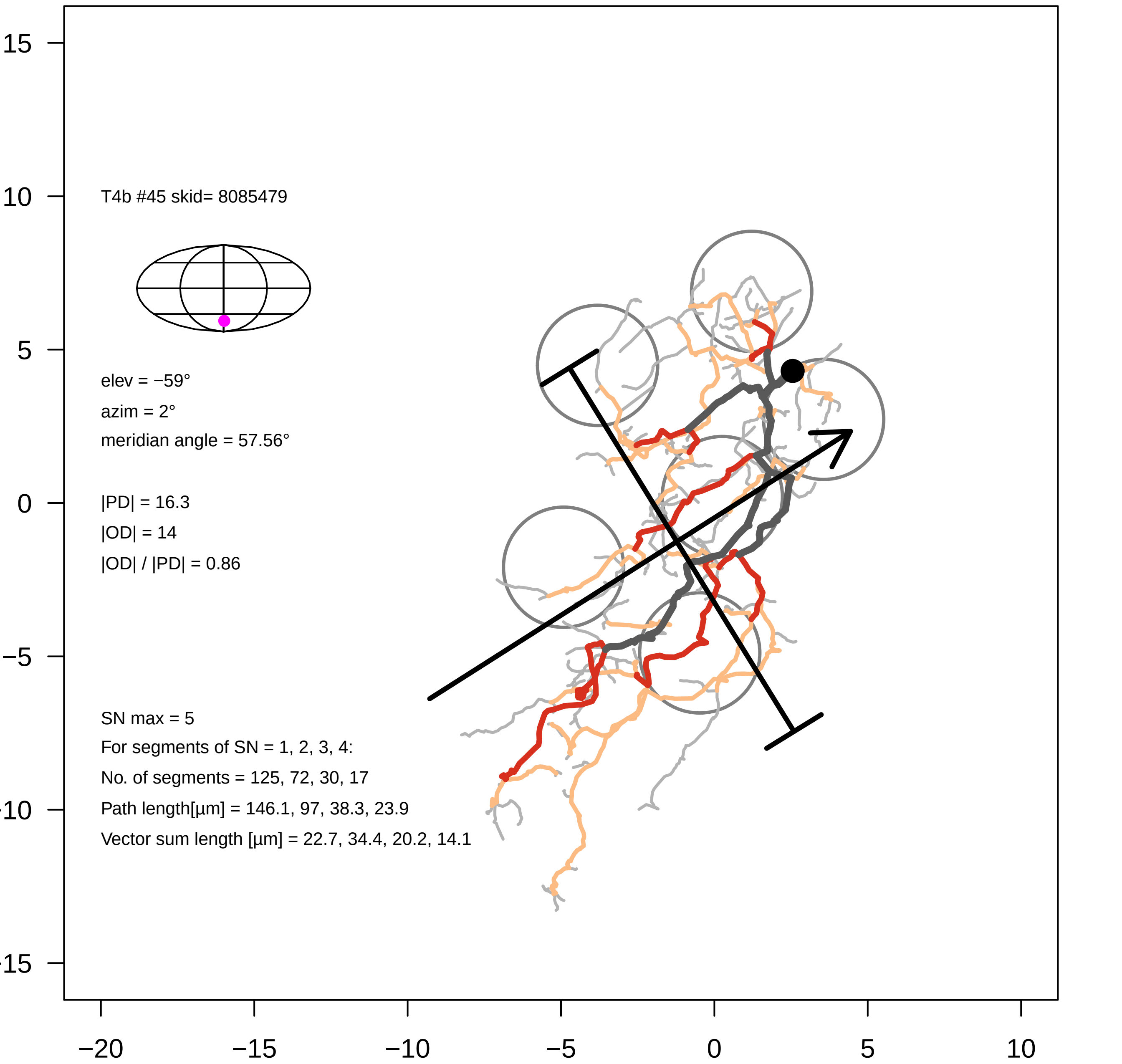
T4b #45 skid= 8085479
elev = −59°
azim = 2°
meridian angle = 57.56°
|PD| = 16.3
|OD| = 14
|OD| / |PD| = 0.86
SN max = 5
For segments of SN = 1, 2, 3, 4:
No. of segments = 125, 72, 30, 17
Path length[µm] = 146.1, 97, 38.3, 23.9
Vector sum length [µm] = 22.7, 34.4, 20.2, 14.1
15
10
5
0
−5
−10
−15
−20
−15
−10
−5
0
5
10

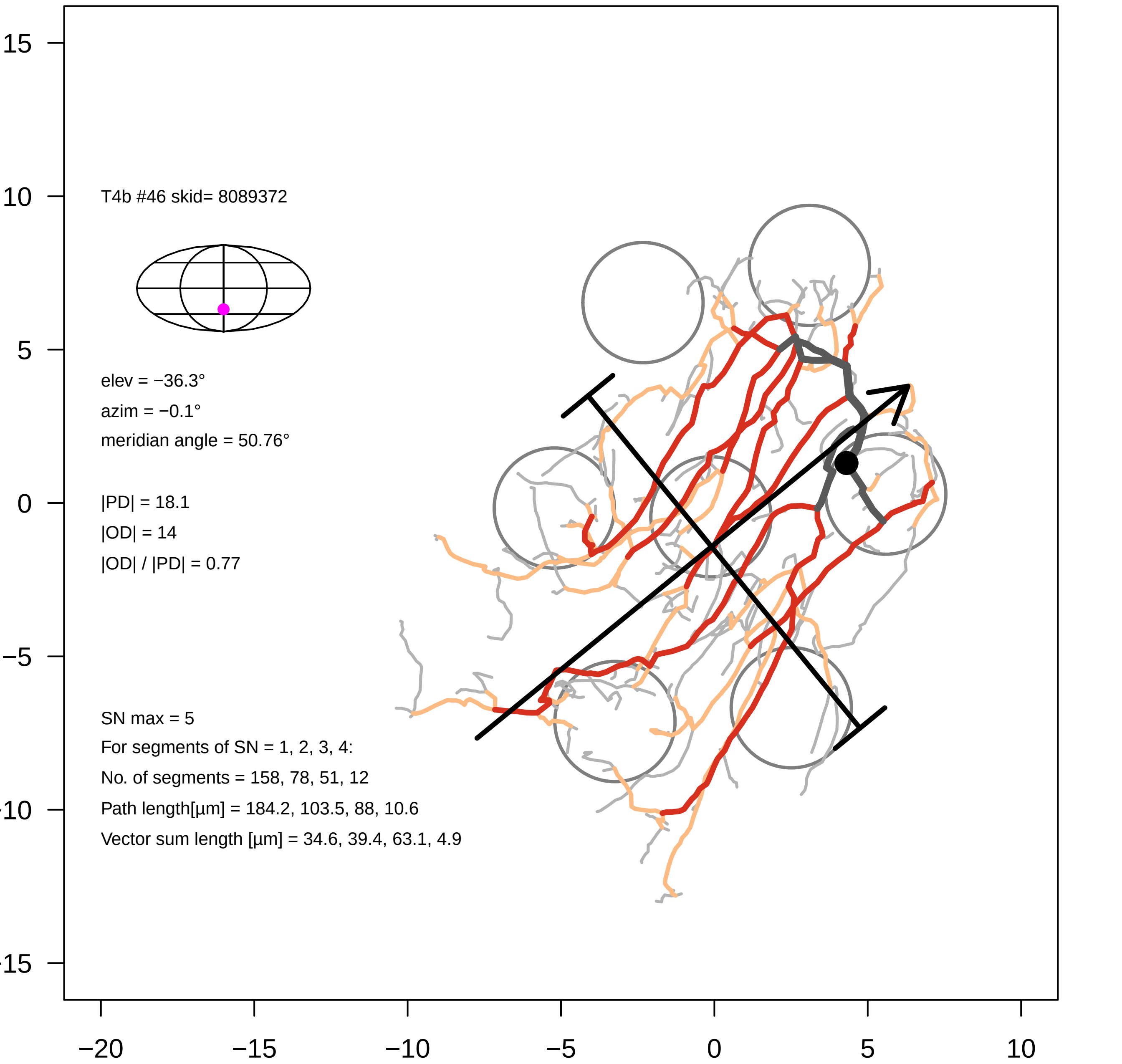

T4b #46 skid= 8089372
elev = −36.3°
azim = −0.1°
meridian angle = 50.76°
|PD| = 18.1
|OD| = 14
|OD| / |PD| = 0.77
SN max = 5
For segments of SN = 1, 2, 3, 4:
No. of segments = 158, 78, 51, 12
Path length[µm] = 184.2, 103.5, 88, 10.6
Vector sum length [µm] = 34.6, 39.4, 63.1, 4.9
15
10
5
0
−5
−10
−15
−20
−15
−10
−5
0
5
10

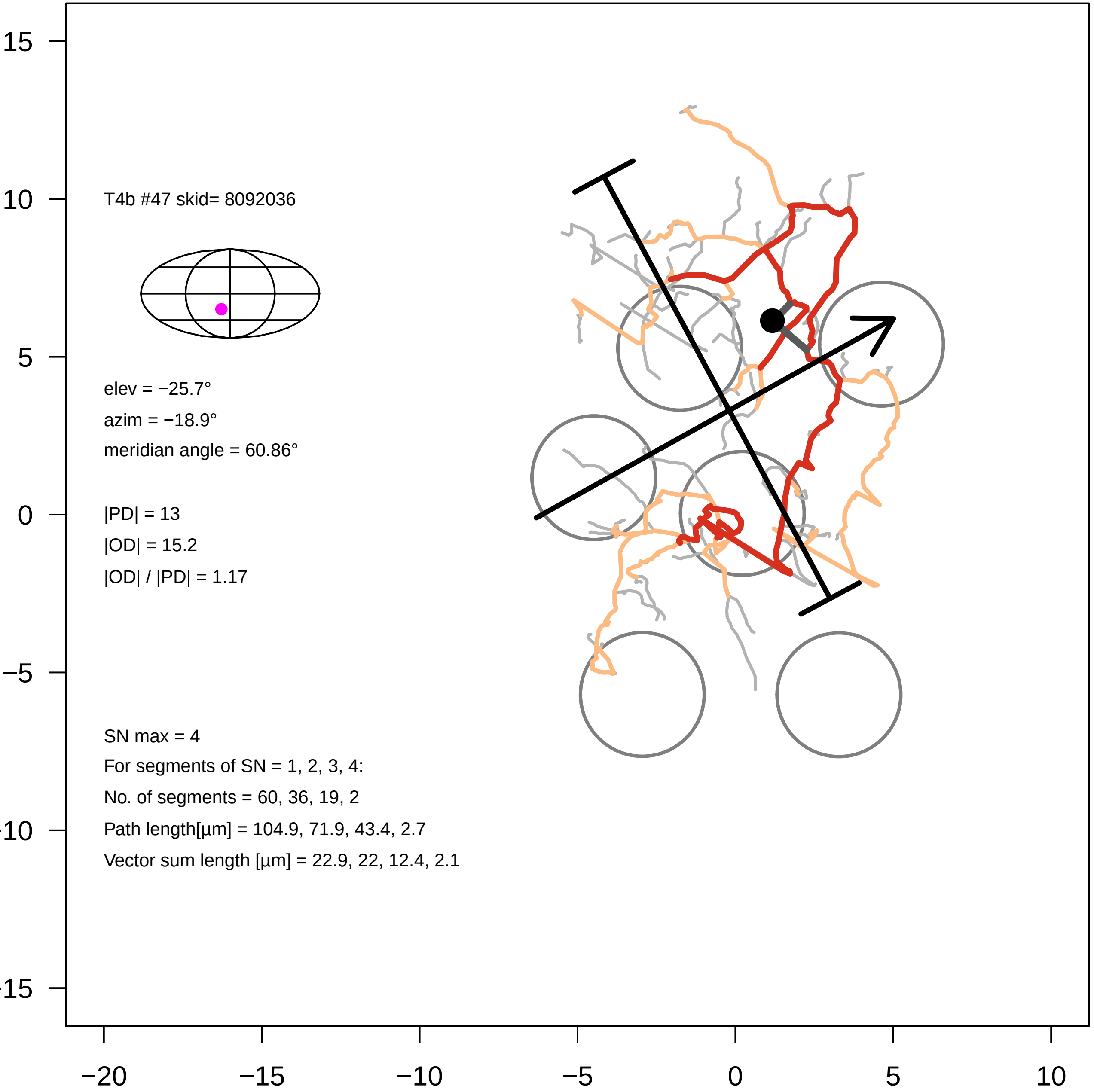
T4b #47 skid= 8092036
elev = −25.7°
azim = −18.9°
meridian angle = 60.86°
|PD| = 13
|OD| = 15.2
|OD| / |PD| = 1.17
SN max = 4
For segments of SN = 1, 2, 3, 4:
No. of segments = 60, 36, 19, 2
Path length[µm] = 104.9, 71.9, 43.4, 2.7
Vector sum length [µm] = 22.9, 22, 12.4, 2.1
15
10
5
0
−5
−10
−15
−20
−15
−10
−5
0
5
10

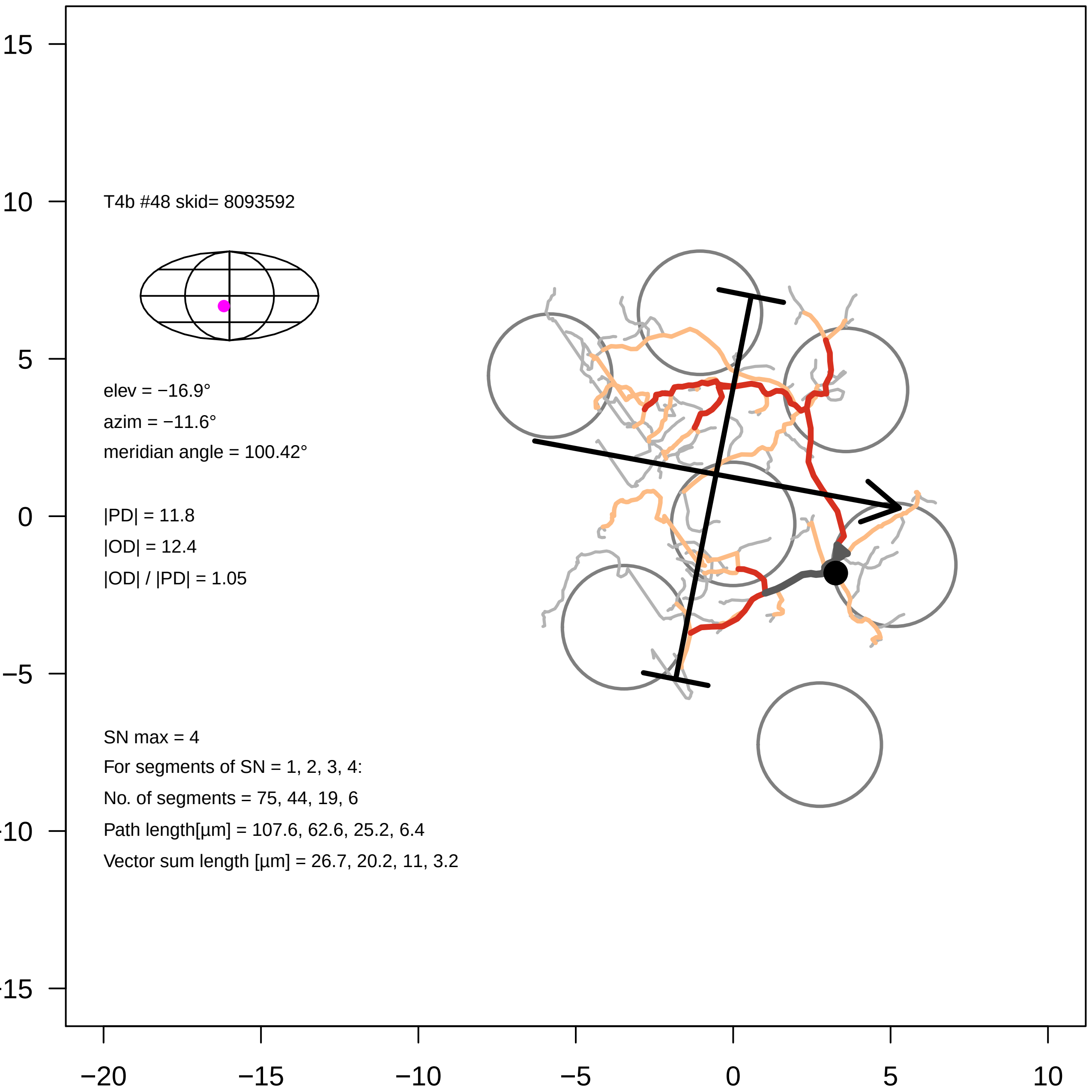
T4b #48 skid= 8093592
elev = −16.9°
azim = −11.6°
meridian angle = 100.42°
|PD| = 11.8
|OD| = 12.4
|OD| / |PD| = 1.05
SN max = 4
For segments of SN = 1, 2, 3, 4:
No. of segments = 75, 44, 19, 6
Path length[µm] = 107.6, 62.6, 25.2, 6.4
Vector sum length [µm] = 26.7, 20.2, 11, 3.2
15
10
5
0
−5
−10
−15
−20
−15
−10
−5
0
5
10

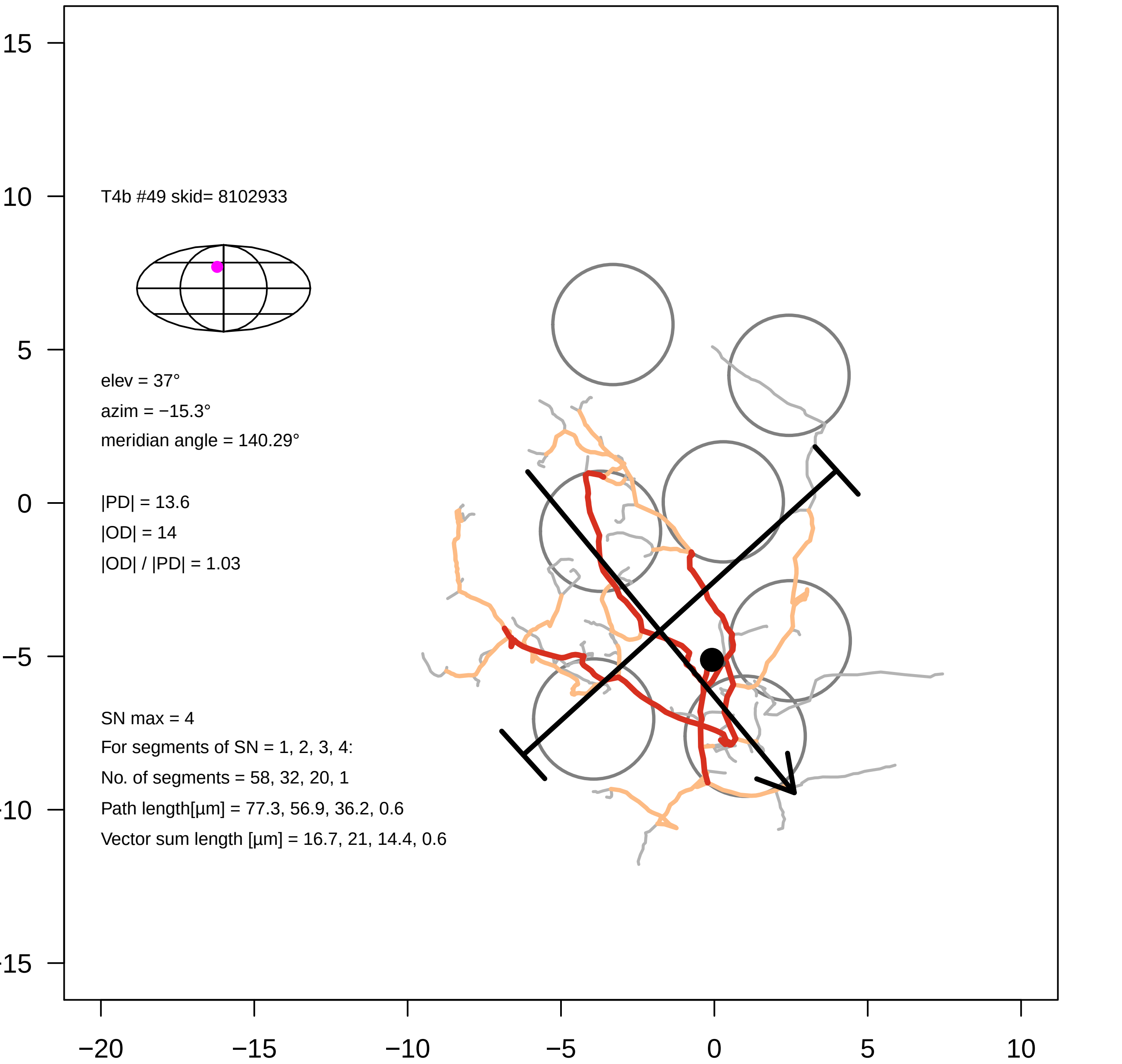

T4b #49 skid= 8102933
elev = 37°
azim = −15.3°
meridian angle = 140.29°
|PD| = 13.6
|OD| = 14
|OD| / |PD| = 1.03
SN max = 4
For segments of SN = 1, 2, 3, 4:
No. of segments = 58, 32, 20, 1
Path length[µm] = 77.3, 56.9, 36.2, 0.6
Vector sum length [µm] = 16.7, 21, 14.4, 0.6
15
10
5
0
−5
−10
−15
−20
−15
−10
−5
0
5
10

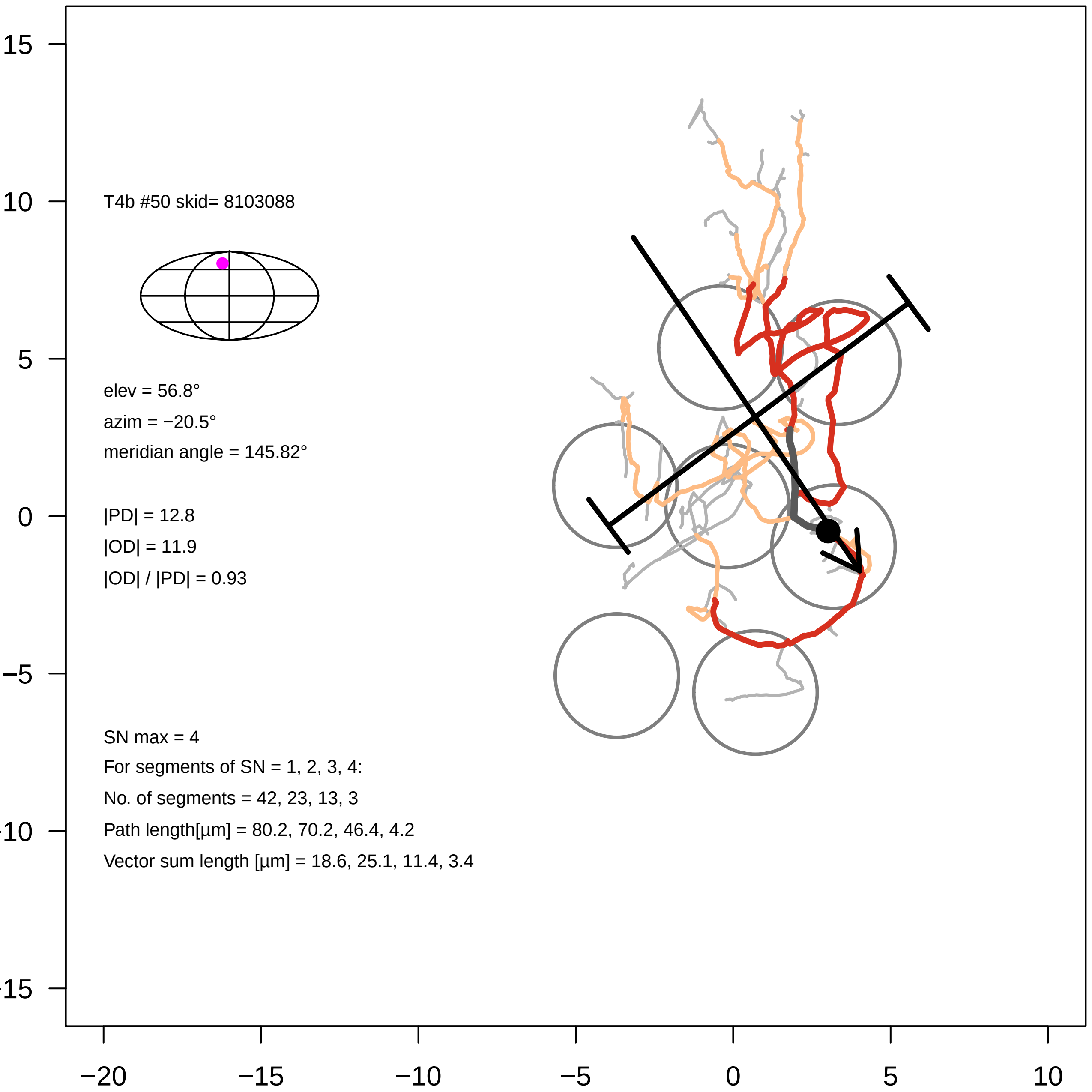
T4b #50 skid= 8103088
elev = 56.8°
azim = −20.5°
meridian angle = 145.82°
|PD| = 12.8
|OD| = 11.9
|OD| / |PD| = 0.93
SN max = 4
For segments of SN = 1, 2, 3, 4:
No. of segments = 42, 23, 13, 3
Path length[µm] = 80.2, 70.2, 46.4, 4.2
Vector sum length [µm] = 18.6, 25.1, 11.4, 3.4
15
10
5
0
−5
−10
−15
−20
−15
−10
−5
0
5
10

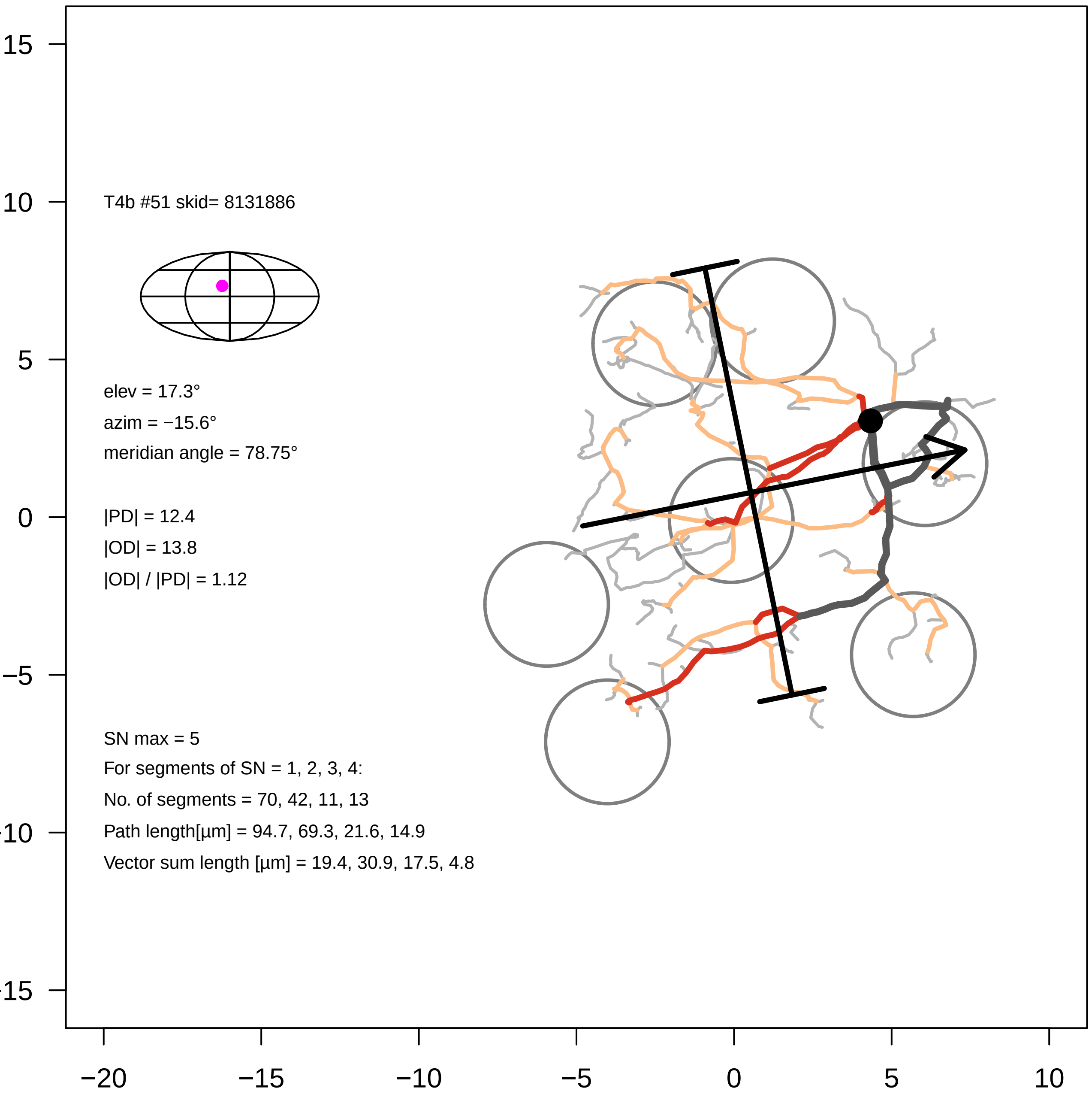

T4b #51 skid= 8131886
elev = 17.3°
azim = −15.6°
meridian angle = 78.75°
|PD| = 12.4
|OD| = 13.8
|OD| / |PD| = 1.12
SN max = 5
For segments of SN = 1, 2, 3, 4:
No. of segments = 70, 42, 11, 13
Path length[µm] = 94.7, 69.3, 21.6, 14.9
Vector sum length [µm] = 19.4, 30.9, 17.5, 4.8
15
10
5
0
−5
−10
−15
−20
−15
−10
−5
0
5
10

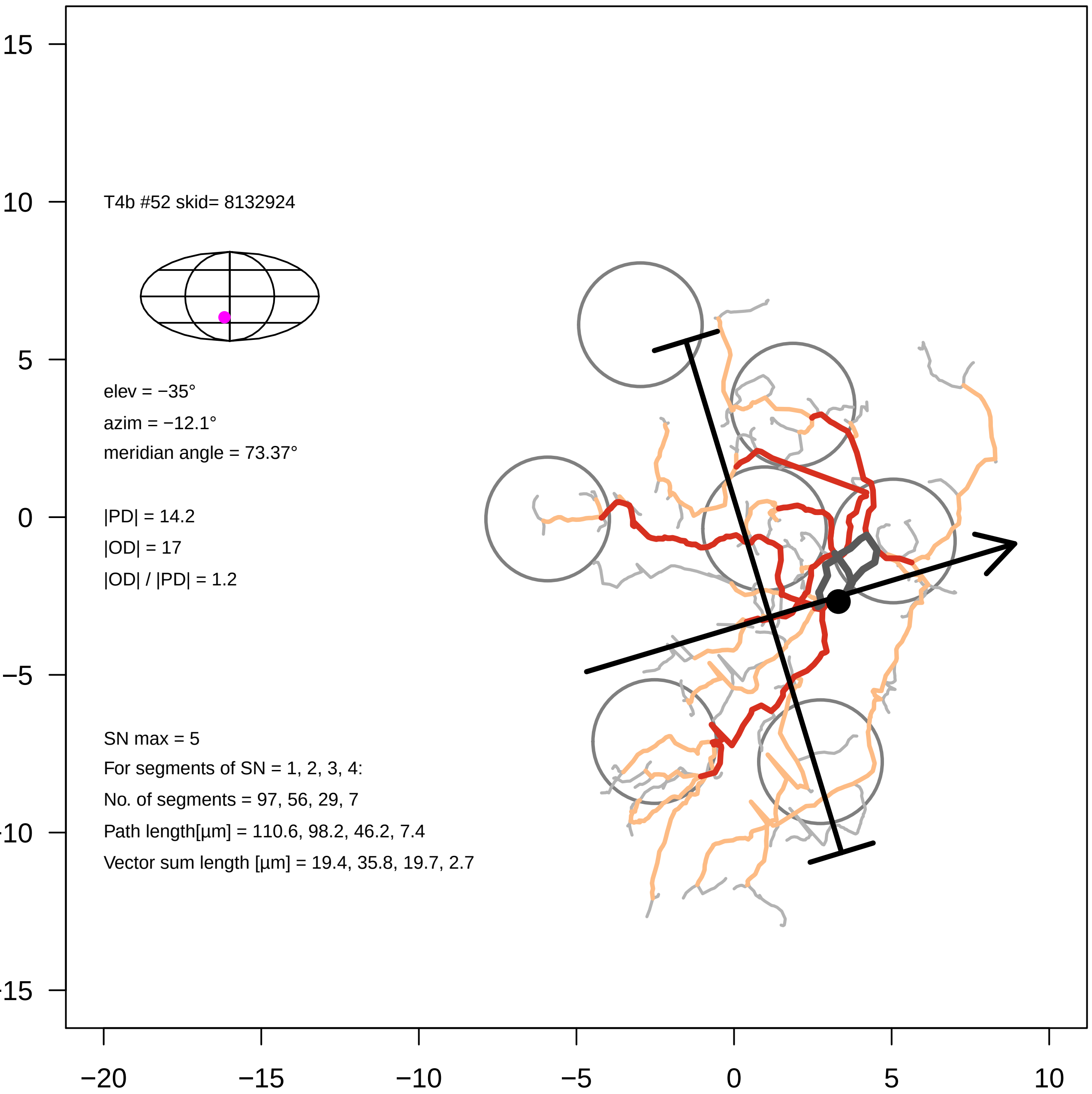
T4b #52 skid= 8132924
elev = −35°
azim = −12.1°
meridian angle = 73.37°
|PD| = 14.2
|OD| = 17
|OD| / |PD| = 1.2
SN max = 5
For segments of SN = 1, 2, 3, 4:
No. of segments = 97, 56, 29, 7
Path length[µm] = 110.6, 98.2, 46.2, 7.4
Vector sum length [µm] = 19.4, 35.8, 19.7, 2.7
15
10
5
0
−5
-10
-15
−20
−15
−10
−5
0
5
10

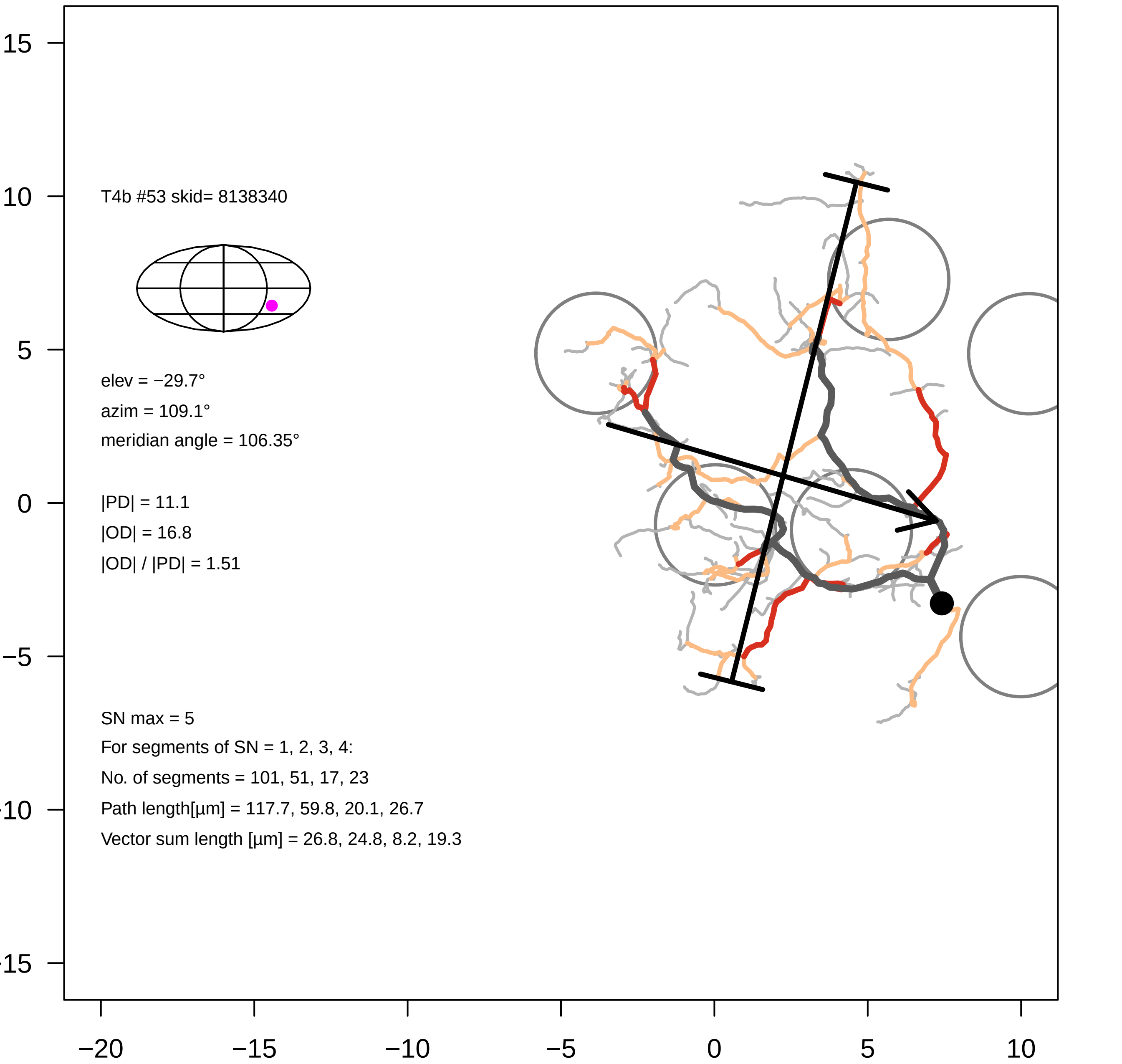
T4b #53 skid= 8138340
elev = −29.7°
azim = 109.1°
meridian angle = 106.35°
|PD| = 11.1
|OD| = 16.8
|OD| / |PD| = 1.51
SN max = 5
For segments of SN = 1, 2, 3, 4:
No. of segments = 101, 51, 17, 23
Path length[µm] = 117.7, 59.8, 20.1, 26.7
Vector sum length [µm] = 26.8, 24.8, 8.2, 19.3
15
10
5
0
−5
−10
−15
−20
−15
−10
−5
0
5
10

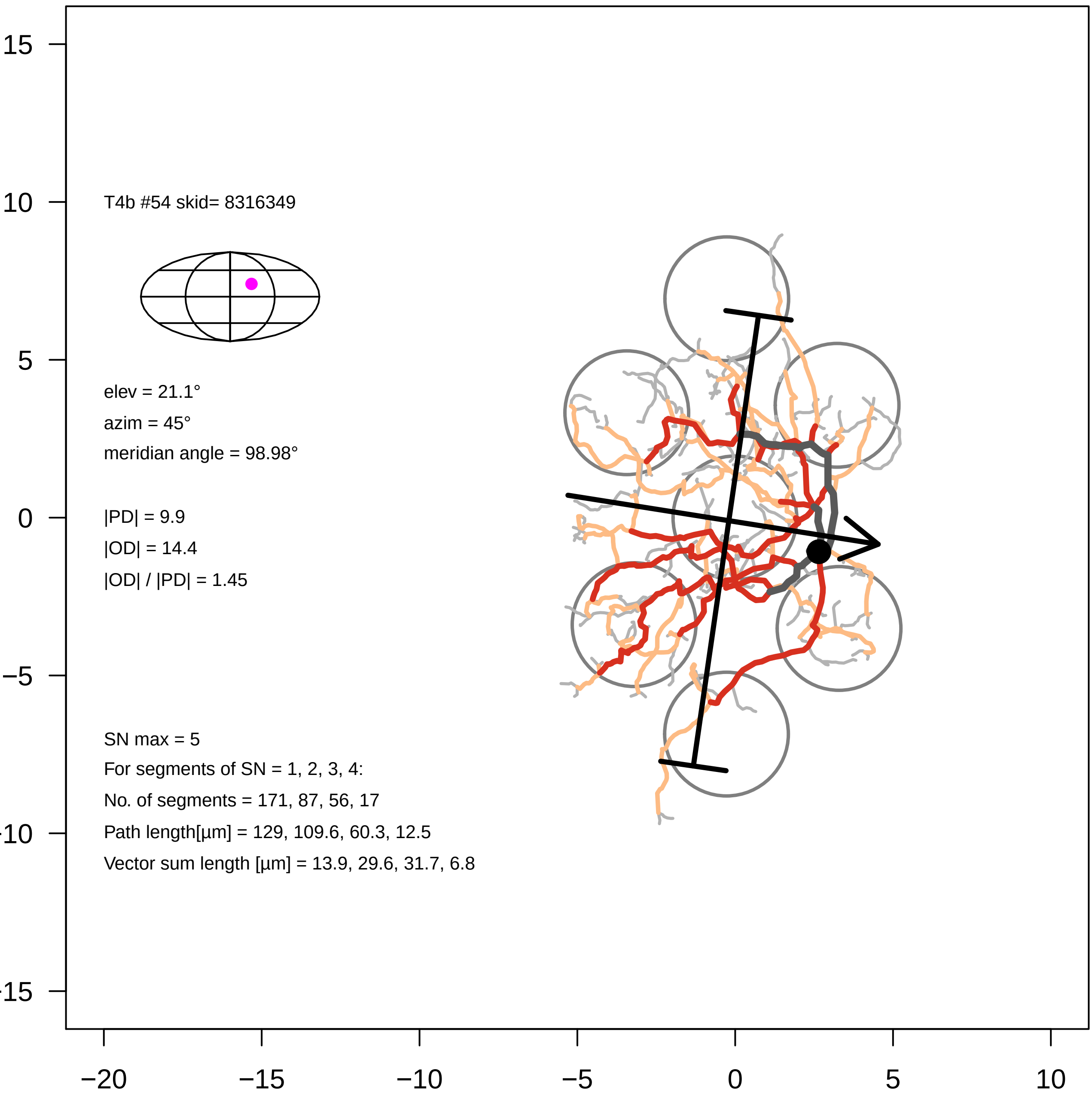
T4b #54 skid= 8316349
elev = 21.1°
azim = 45°
meridian angle = 98.98°
|PD| = 9.9
|OD| = 14.4
|OD| / |PD| = 1.45
SN max = 5
For segments of SN = 1, 2, 3, 4:
No. of segments = 171, 87, 56, 17
Path length[µm] = 129, 109.6, 60.3, 12.5
Vector sum length [µm] = 13.9, 29.6, 31.7, 6.8
15
10
5
0
−5
−10
−15
−20
−15
−10
−5
0
5
10

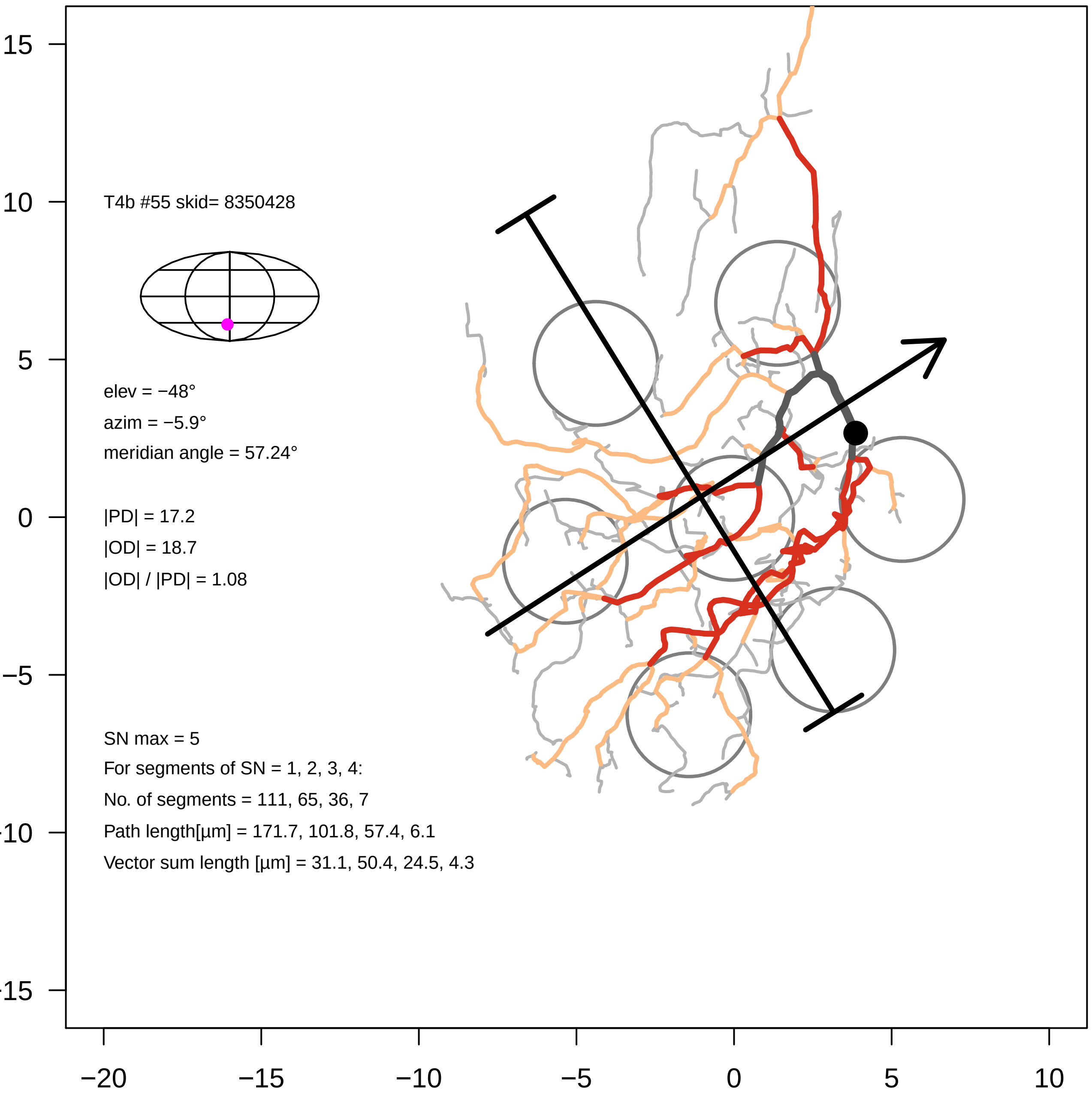
T4b #55 skid= 8350428
elev = −48°
azim = −5.9°
meridian angle = 57.24°
|PD| = 17.2
|OD| = 18.7
|OD| / |PD| = 1.08
SN max = 5
For segments of SN = 1, 2, 3, 4:
No. of segments = 111, 65, 36, 7
Path length[µm] = 171.7, 101.8, 57.4, 6.1
Vector sum length [µm] = 31.1, 50.4, 24.5, 4.3
15
10
5
0
−5
−10
−15
−20
−15
−10
−5
0
5
10

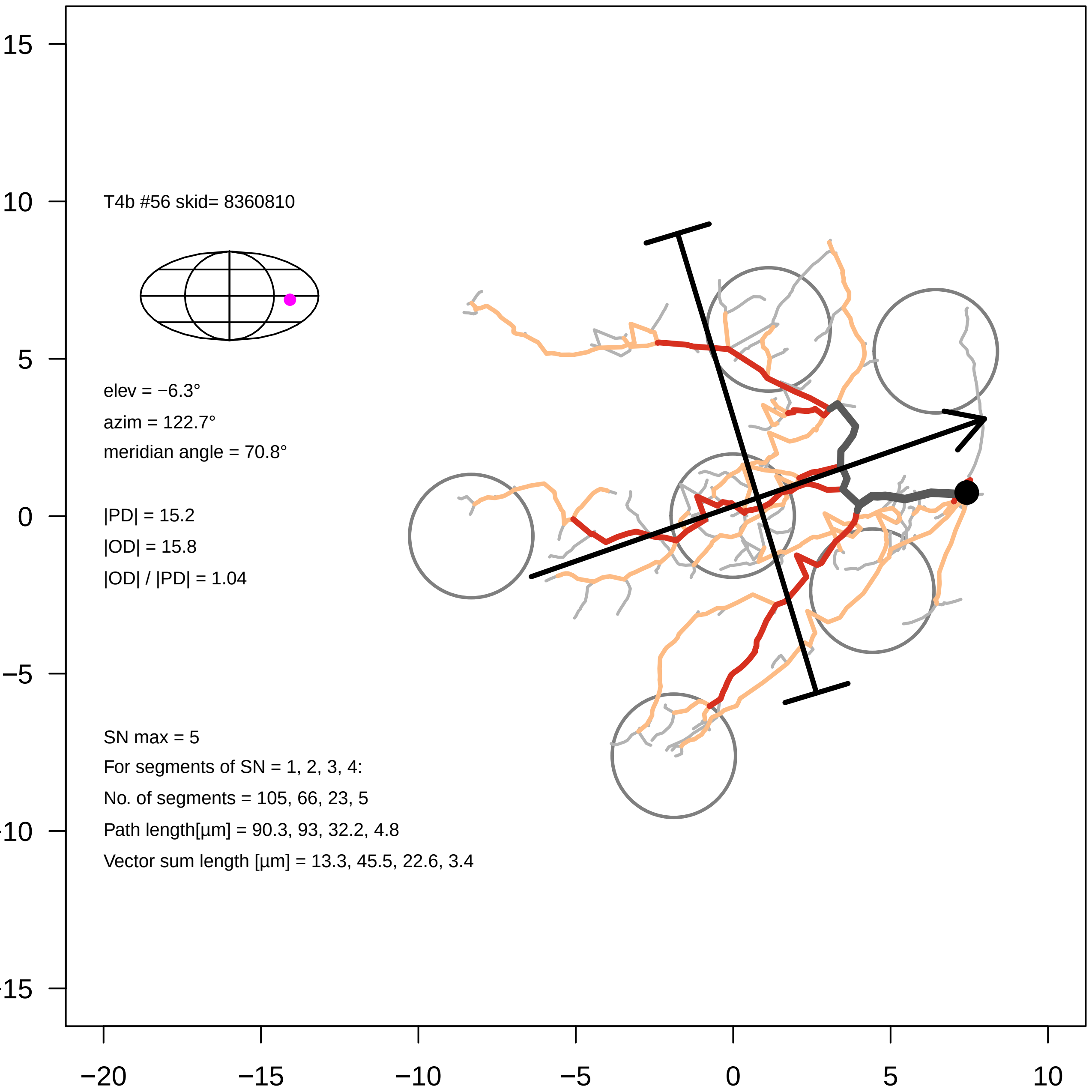

T4b #56 skid= 8360810
elev = −6.3°
azim = 122.7°
meridian angle = 70.8°
|PD| = 15.2
|OD| = 15.8
|OD| / |PD| = 1.04
SN max = 5
For segments of SN = 1, 2, 3, 4:
No. of segments = 105, 66, 23, 5
Path length[µm] = 90.3, 93, 32.2, 4.8
Vector sum length [µm] = 13.3, 45.5, 22.6, 3.4
15
10
5
0
−5
−10
−15
−20
−15
−10
−5
0
5
10

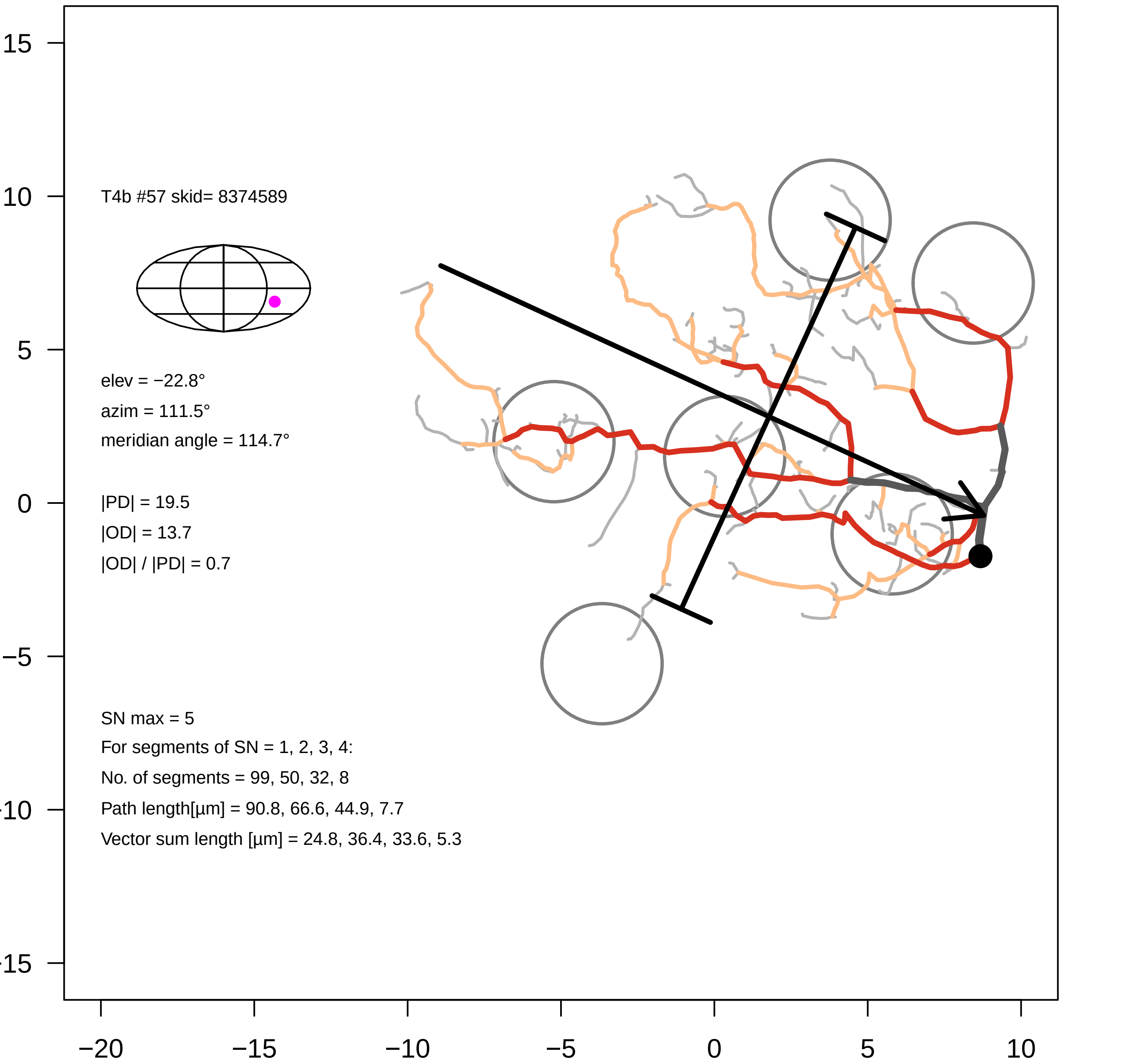
T4b #57 skid= 8374589
elev = −22.8°
azim = 111.5°
meridian angle = 114.7°
|PD| = 19.5
|OD| = 13.7
|OD| / |PD| = 0.7
SN max = 5
For segments of SN = 1, 2, 3, 4:
No. of segments = 99, 50, 32, 8
Path length[µm] = 90.8, 66.6, 44.9, 7.7
Vector sum length [µm] = 24.8, 36.4, 33.6, 5.3
15
10
5
0
−5
−10
−15
−20
−15
−10
−5
0
5
10

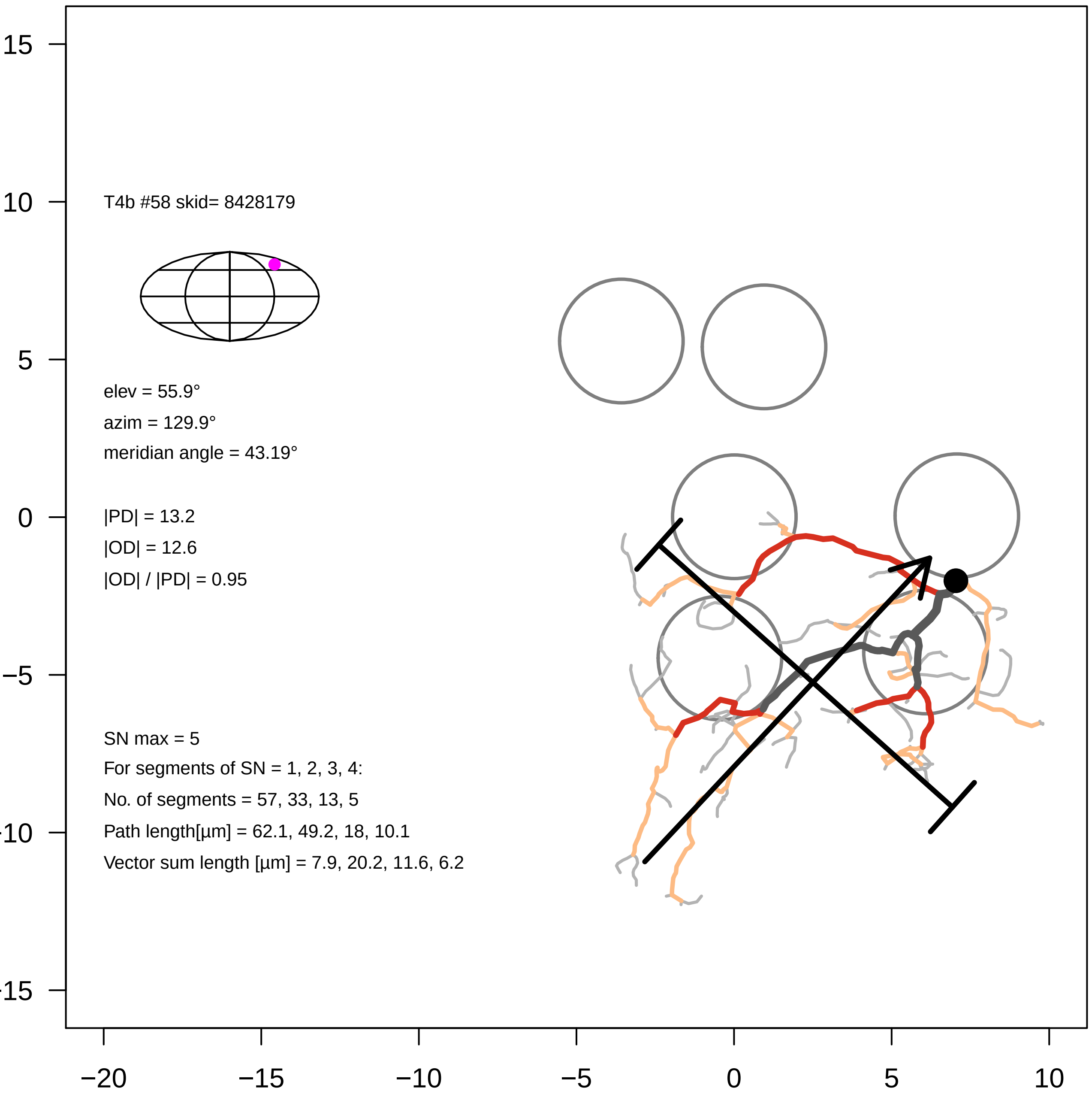

T4b #58 skid= 8428179
elev = 55.9°
azim = 129.9°
meridian angle = 43.19°
|PD| = 13.2
|OD| = 12.6
|OD| / |PD| = 0.95
SN max = 5
For segments of SN = 1, 2, 3, 4:
No. of segments = 57, 33, 13, 5
Path length[µm] = 62.1, 49.2, 18, 10.1
Vector sum length [µm] = 7.9, 20.2, 11.6, 6.2
15
10
5
0
−5
−10
−15
−20
−15
−10
−5
0
5
10

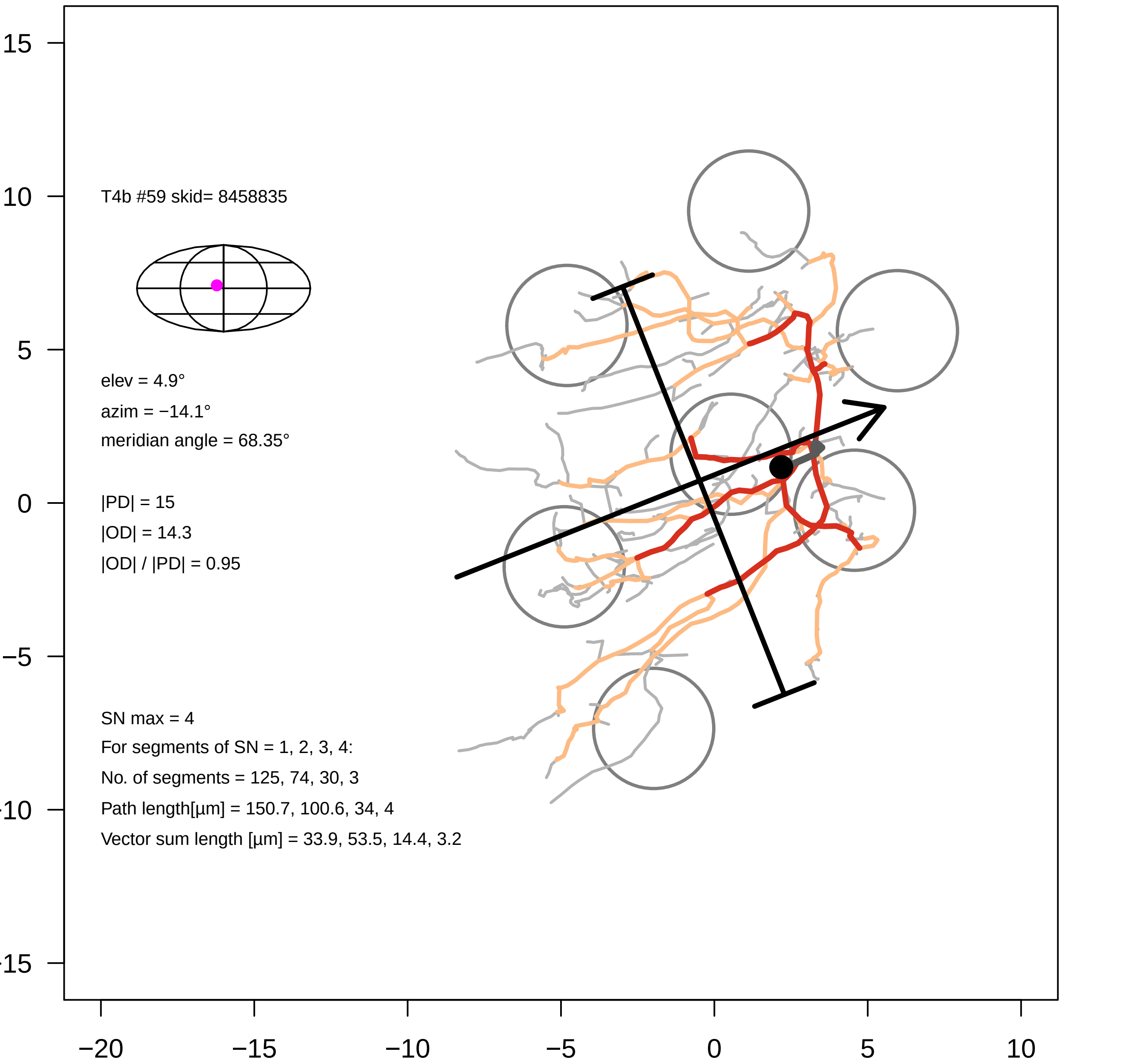
T4b #59 skid= 8458835
elev = 4.9°
azim = −14.1°
meridian angle = 68.35°
|PD| = 15
|OD| = 14.3
|OD| / |PD| = 0.95
SN max = 4
For segments of SN = 1, 2, 3, 4:
No. of segments = 125, 74, 30, 3
Path length[µm] = 150.7, 100.6, 34, 4
Vector sum length [µm] = 33.9, 53.5, 14.4, 3.2
15
10
5
0
−5
−10
−15
−20
−15
−10
−5
0
5
10

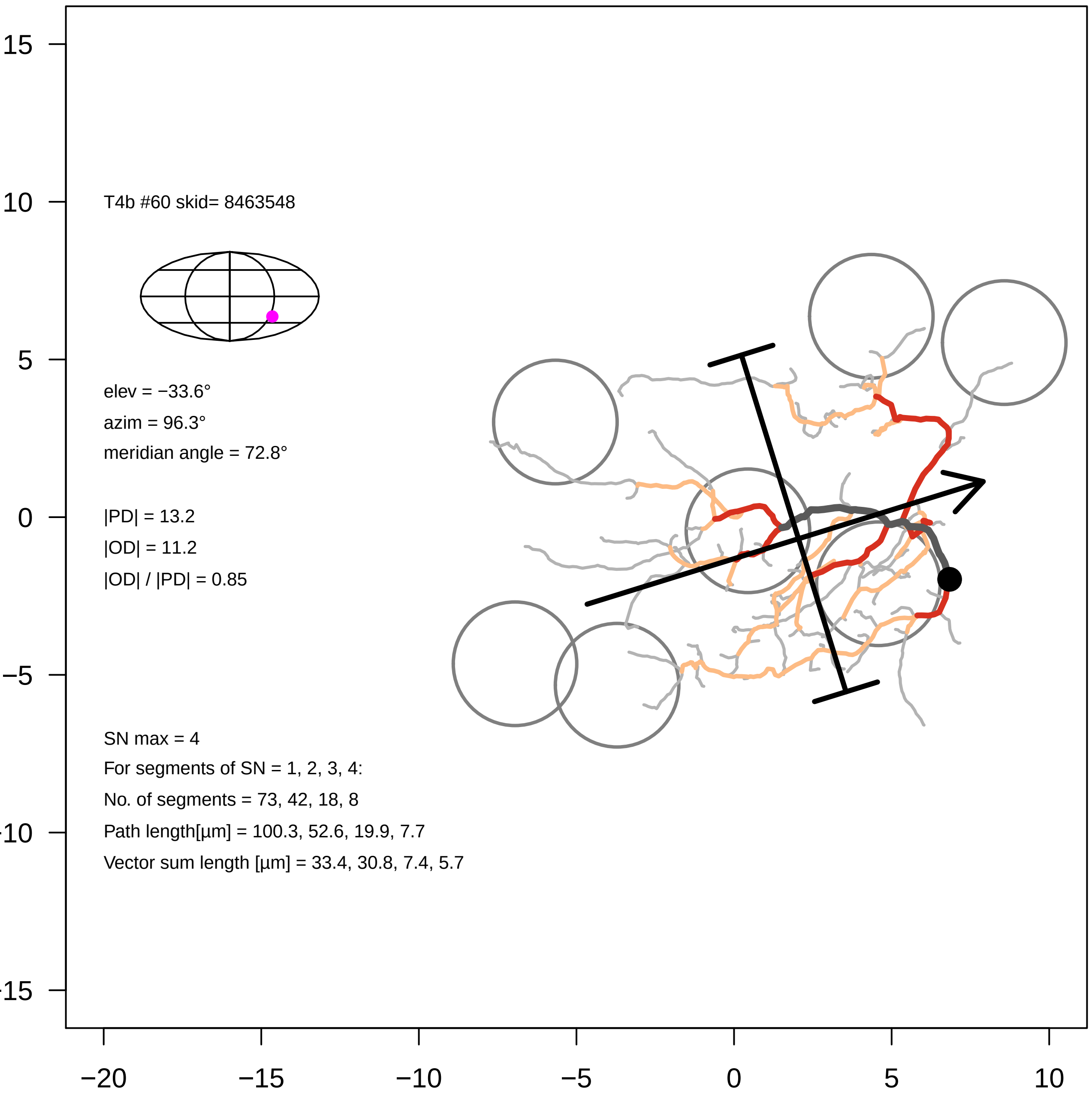
T4b #60 skid= 8463548
elev = −33.6°
azim = 96.3°
meridian angle = 72.8°
|PD| = 13.2
|OD| = 11.2
|OD| / |PD| = 0.85
SN max = 4
For segments of SN = 1, 2, 3, 4:
No. of segments = 73, 42, 18, 8
Path length[µm] = 100.3, 52.6, 19.9, 7.7
Vector sum length [µm] = 33.4, 30.8, 7.4, 5.7
15
10
5
0
−5
-10
-15
−20
−15
−10
−5
0
5
10

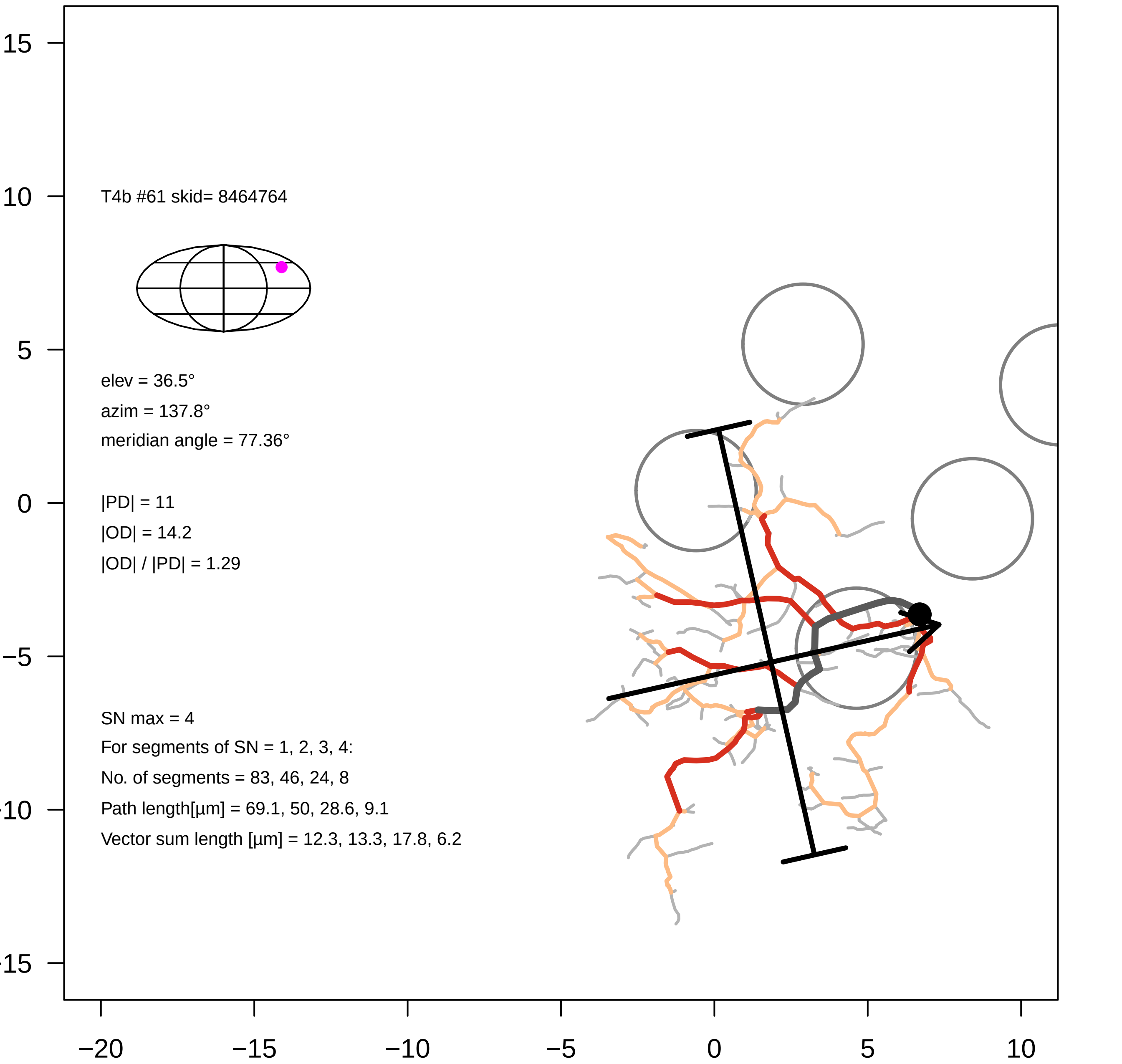

T4b #61 skid= 8464764
elev = 36.5°
azim = 137.8°
meridian angle = 77.36°
|PD| = 11
|OD| = 14.2
|OD| / |PD| = 1.29
SN max = 4
For segments of SN = 1, 2, 3, 4:
No. of segments = 83, 46, 24, 8
Path length[µm] = 69.1, 50, 28.6, 9.1
Vector sum length [µm] = 12.3, 13.3, 17.8, 6.2
15
10
5
0
−5
−10
−15
−20
−15
−10
−5
0
5
10

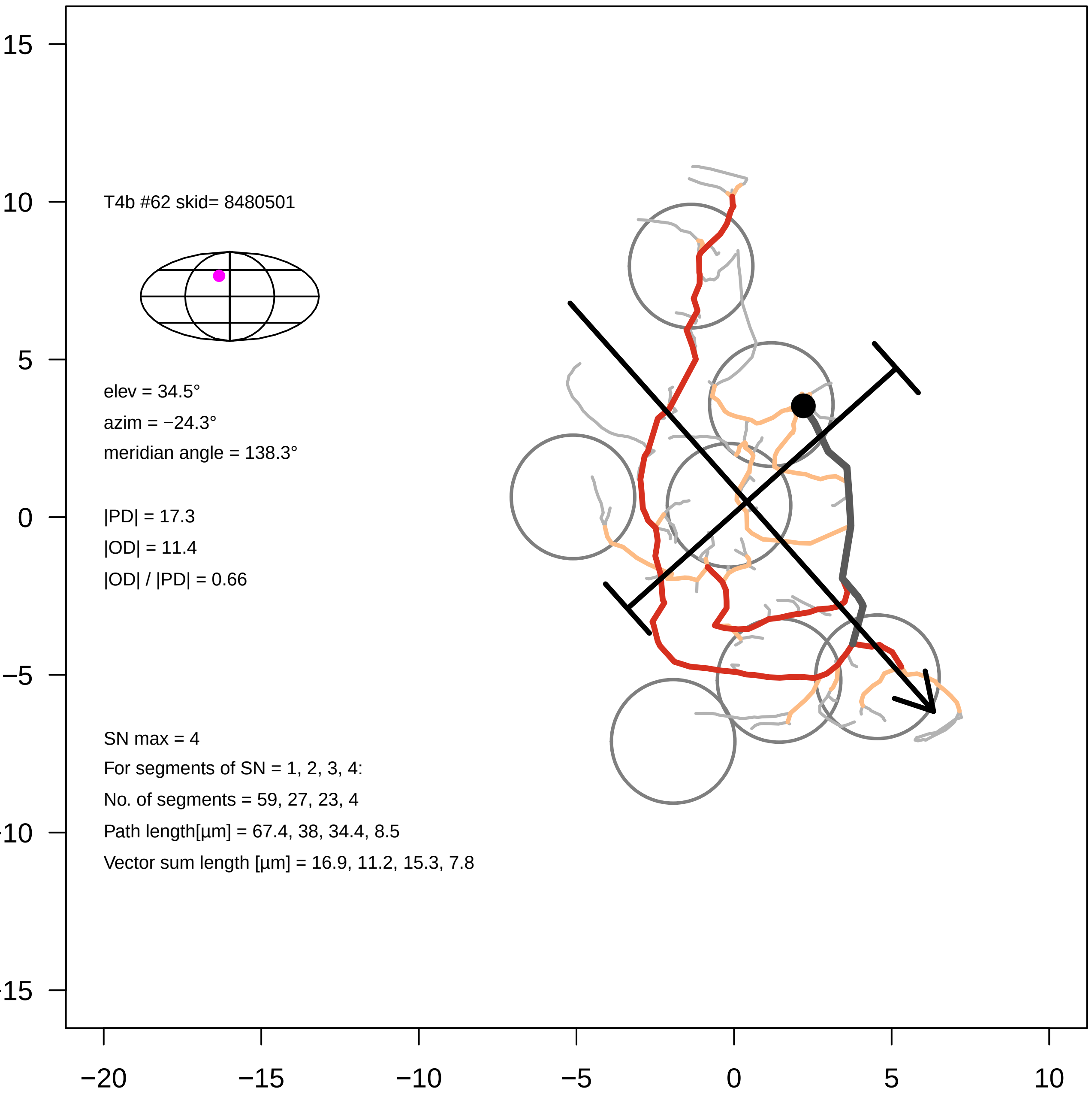

T4b #62 skid= 8480501
elev = 34.5°
azim = −24.3°
meridian angle = 138.3°
|PD| = 17.3
|OD| = 11.4
|OD| / |PD| = 0.66
SN max = 4
For segments of SN = 1, 2, 3, 4:
No. of segments = 59, 27, 23, 4
Path length[µm] = 67.4, 38, 34.4, 8.5
Vector sum length [µm] = 16.9, 11.2, 15.3, 7.8
15
10
5
0
−5
−10
−15
−20
−15
−10
−5
0
5
10

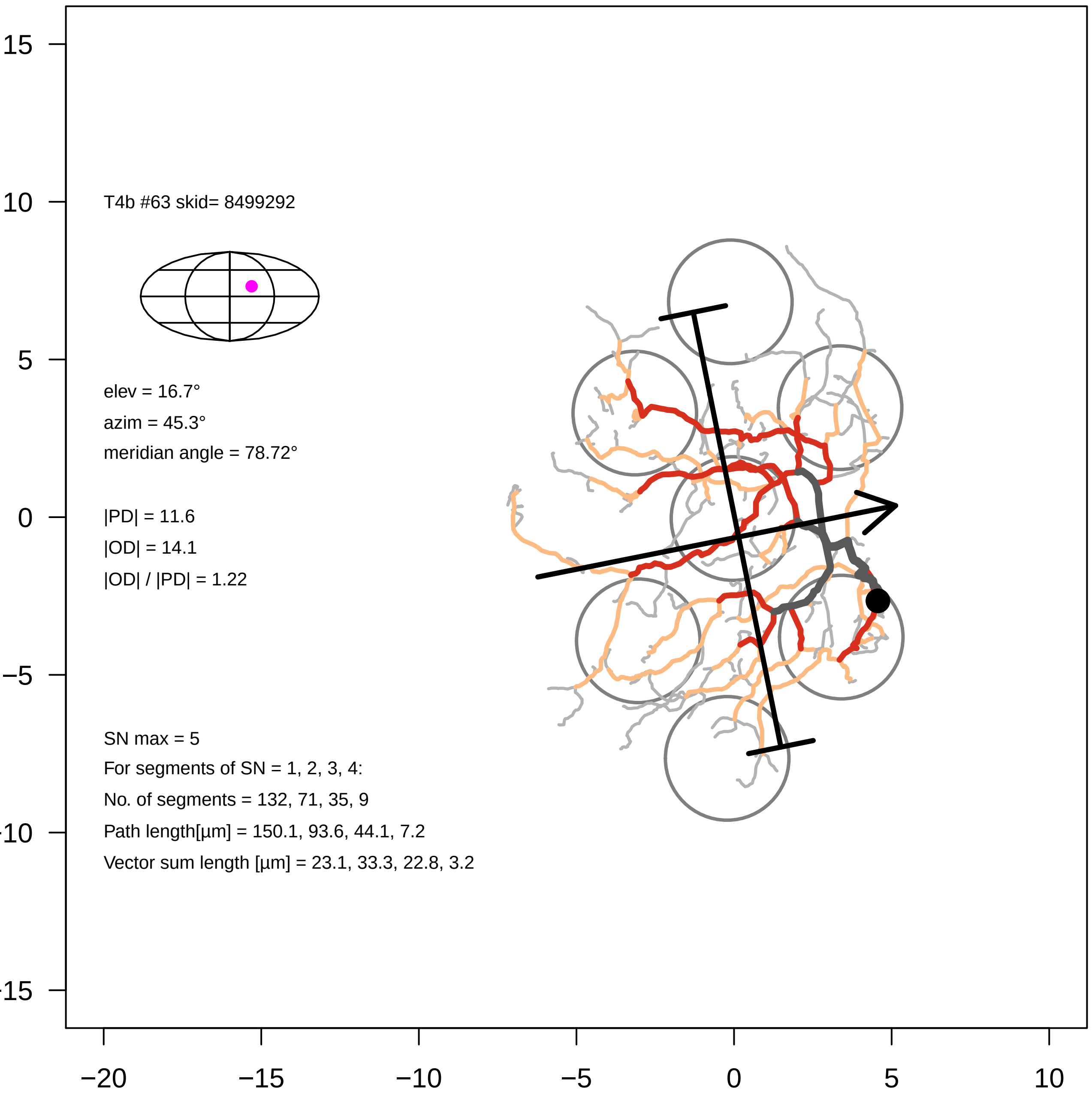

T4b #63 skid= 8499292
elev = 16.7°
azim = 45.3°
meridian angle = 78.72°
|PD| = 11.6
|OD| = 14.1
|OD| / |PD| = 1.22
SN max = 5
For segments of SN = 1, 2, 3, 4:
No. of segments = 132, 71, 35, 9
Path length[µm] = 150.1, 93.6, 44.1, 7.2
Vector sum length [µm] = 23.1, 33.3, 22.8, 3.2

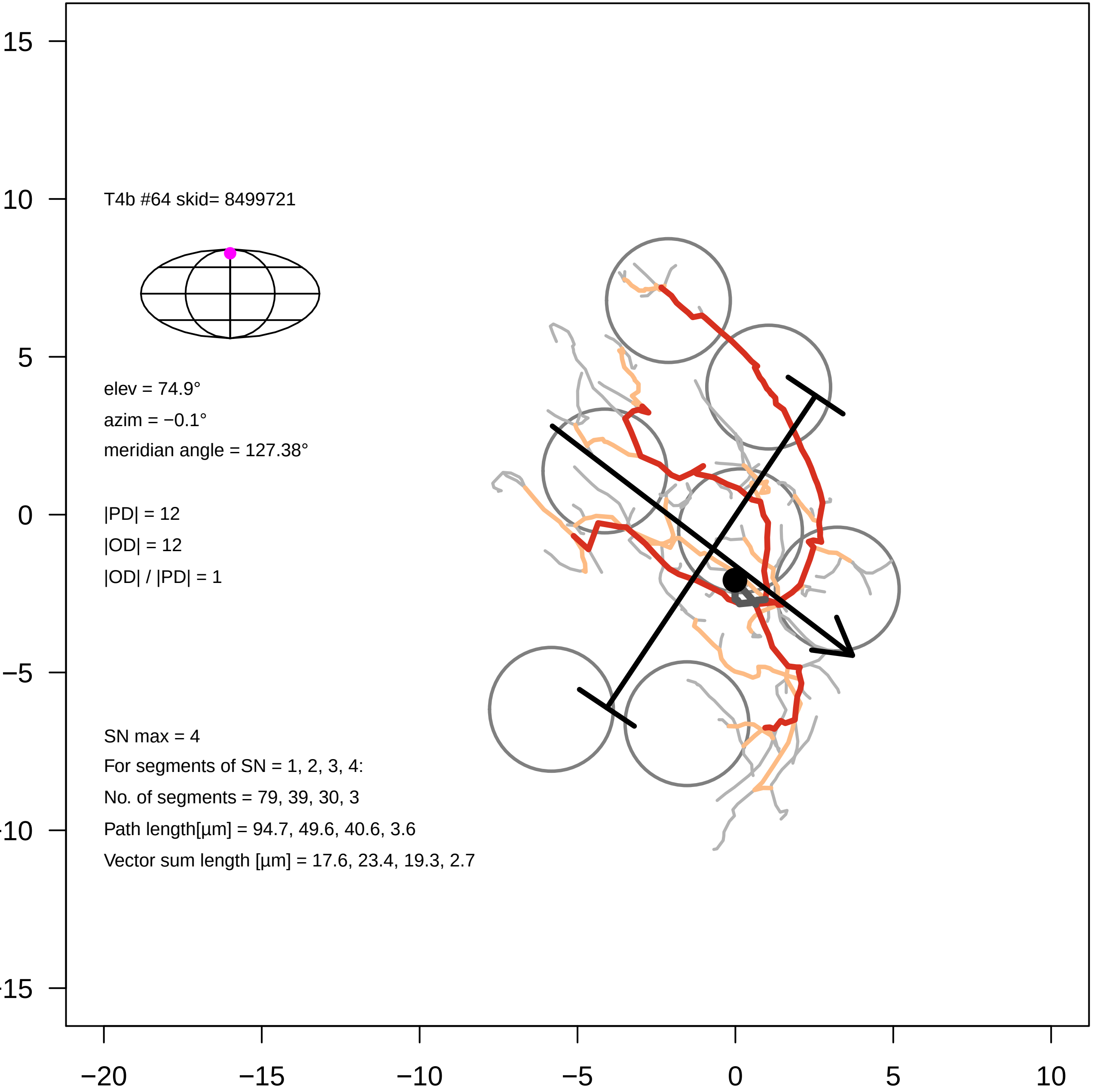
T4b #64 skid= 8499721
elev = 74.9°
azim = −0.1°
meridian angle = 127.38°
|PD| = 12
|OD| = 12
|OD| / |PD| = 1
SN max = 4
For segments of SN = 1, 2, 3, 4:
No. of segments = 79, 39, 30, 3
Path length[µm] = 94.7, 49.6, 40.6, 3.6
Vector sum length [µm] = 17.6, 23.4, 19.3, 2.7
15
10
5
0
−5
−20
−15
−10
−5
0
5
10

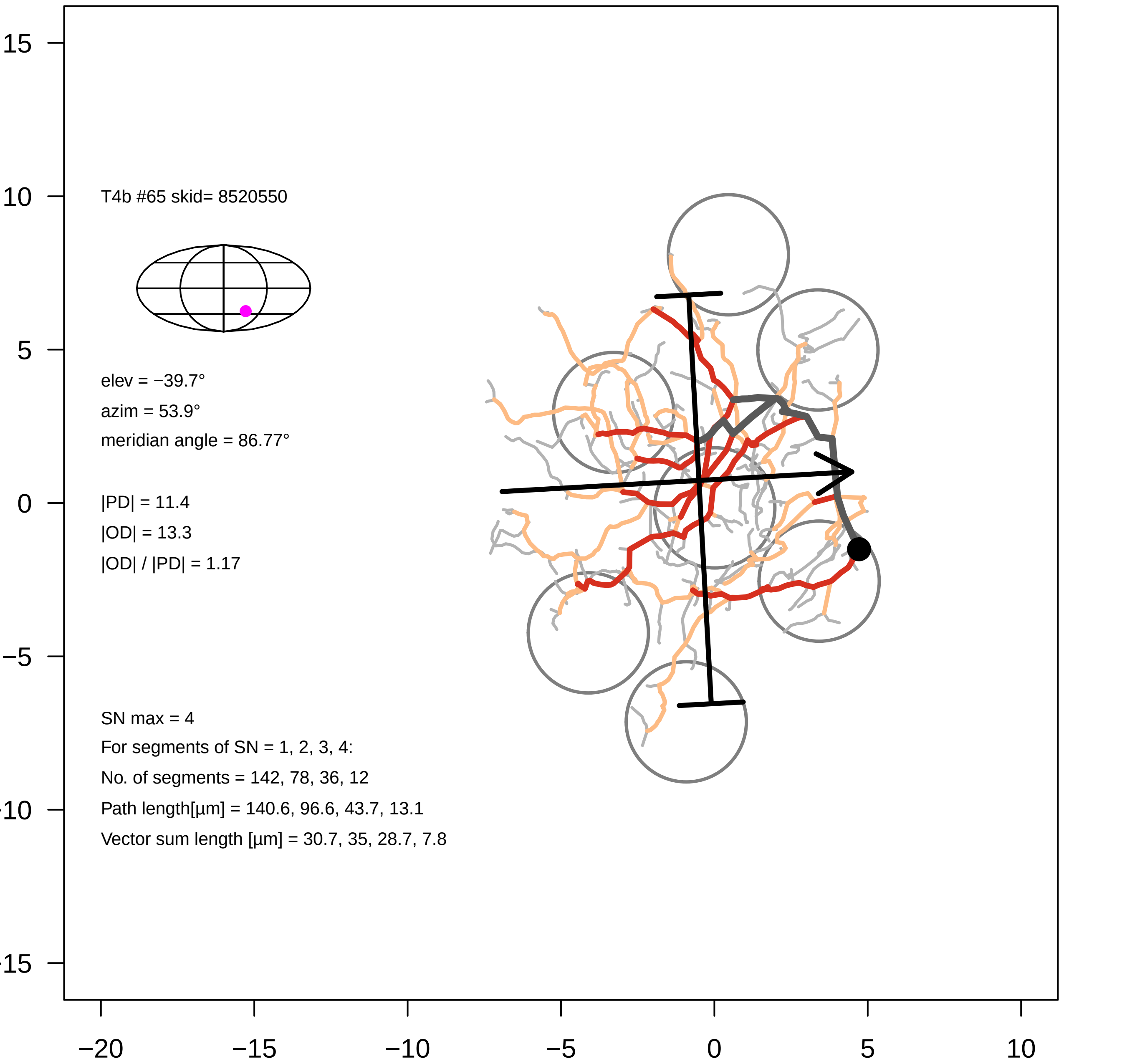
T4b #65 skid= 8520550
elev = −39.7°
azim = 53.9°
meridian angle = 86.77°
|PD| = 11.4
|OD| = 13.3
|OD| / |PD| = 1.17
SN max = 4
For segments of SN = 1, 2, 3, 4:
No. of segments = 142, 78, 36, 12
Path length[µm] = 140.6, 96.6, 43.7, 13.1
Vector sum length [µm] = 30.7, 35, 28.7, 7.8
15
10
5
0
−5
−20
−15
−10
−5
0
5
10

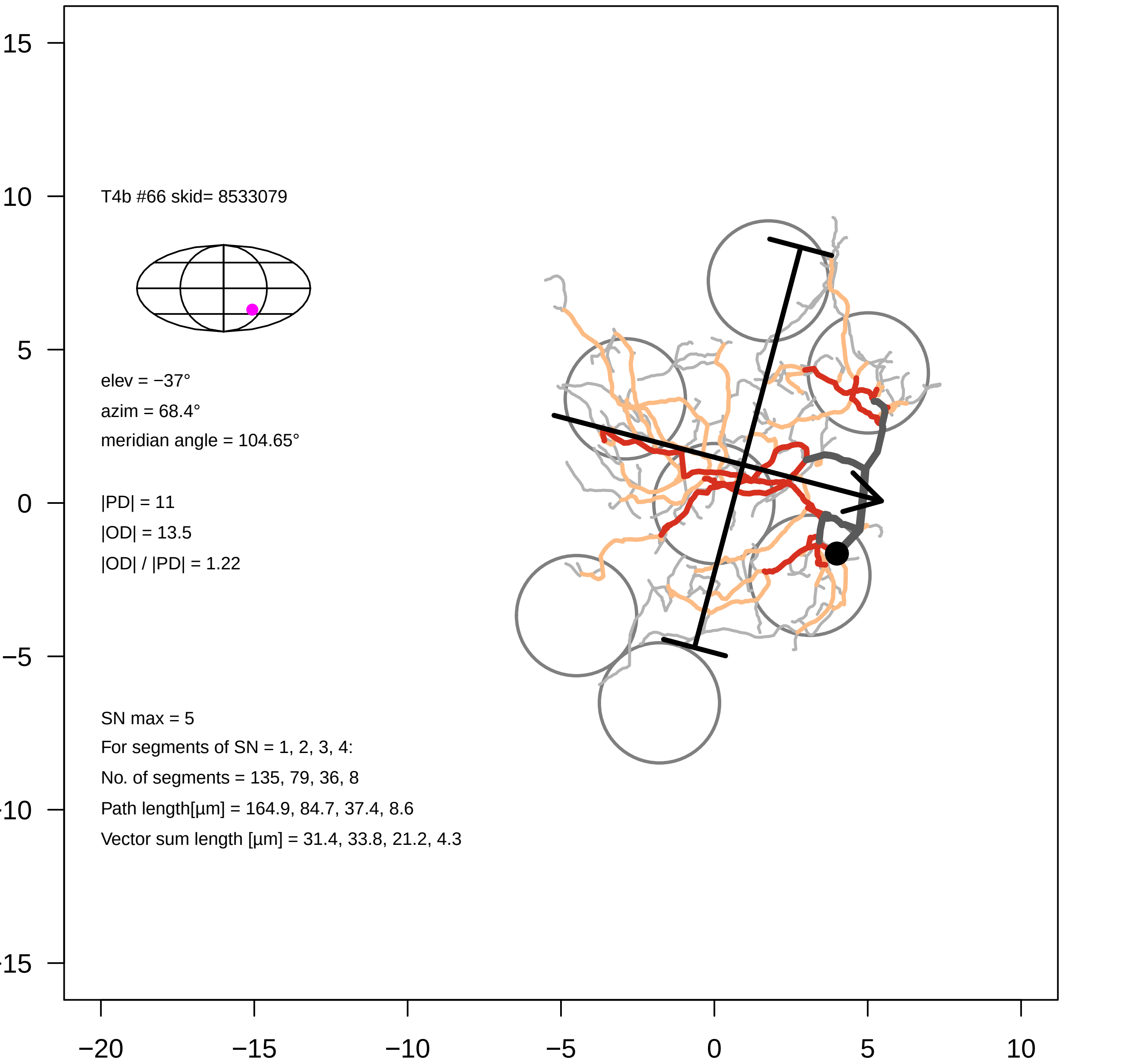

15
10
5
0
−5
−10
−15
−20
−15
−10
−5
0
5
10
T4b #66 skid= 8533079
elev = −37°
azim = 68.4°
meridian angle = 104.65°
|PD| = 11
|OD| = 13.5
|OD| / |PD| = 1.22
SN max = 5
For segments of SN = 1, 2, 3, 4:
No. of segments = 135, 79, 36, 8
Path length[µm] = 164.9, 84.7, 37.4, 8.6
Vector sum length [µm] = 31.4, 33.8, 21.2, 4.3

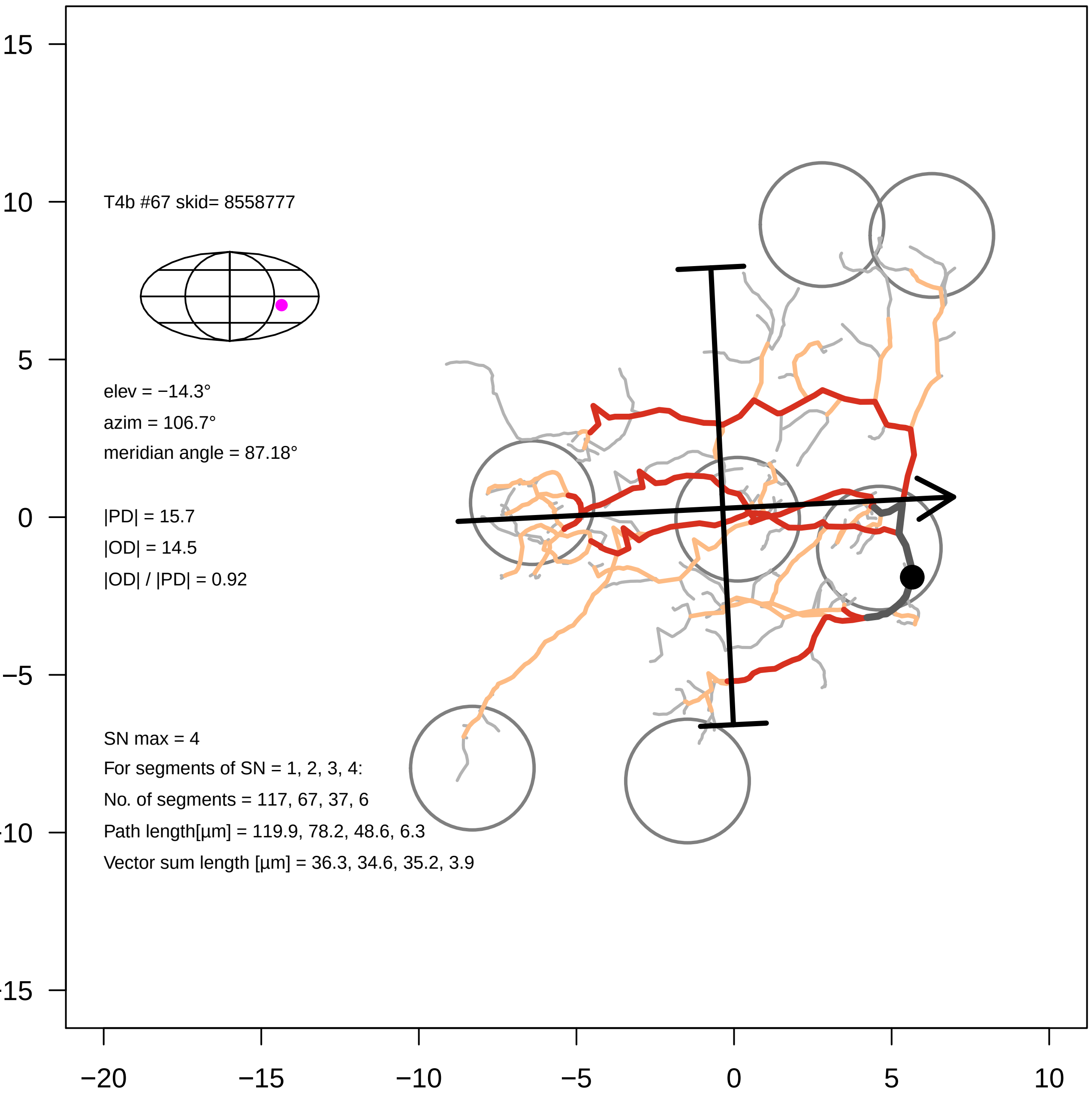

T4b #67 skid= 8558777
elev = −14.3°
azim = 106.7°
meridian angle = 87.18°
|PD| = 15.7
|OD| = 14.5
|OD| / |PD| = 0.92
SN max = 4
For segments of SN = 1, 2, 3, 4:
No. of segments = 117, 67, 37, 6
Path length[μm] = 119.9, 78.2, 48.6, 6.3
Vector sum length [μm] = 36.3, 34.6, 35.2, 3.9
15
10
5
0
−5
-10
-15
−20
−15
−10
−5
0
5
10

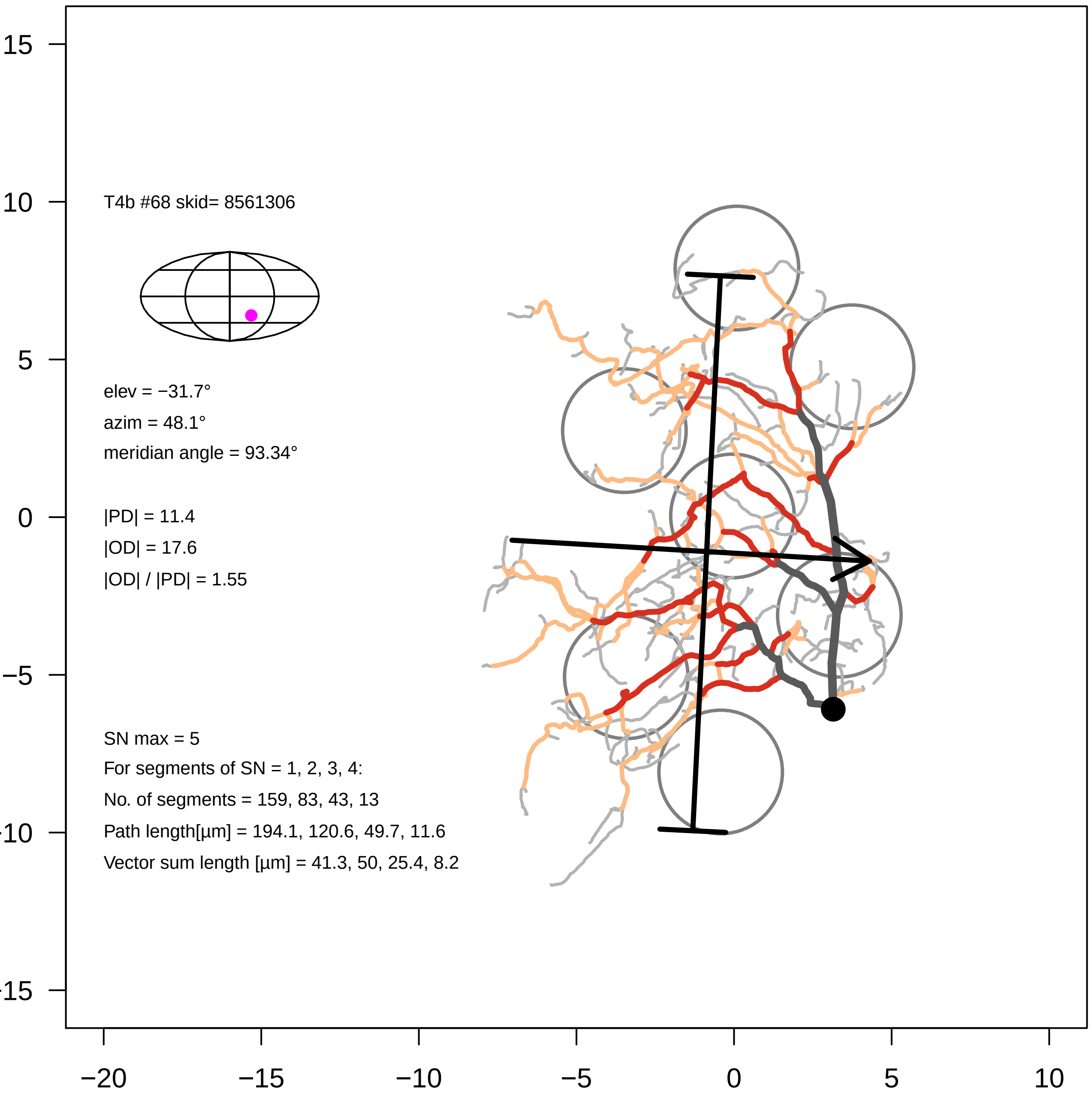

T4b #68 skid= 8561306
elev = −31.7°
azim = 48.1°
meridian angle = 93.34°
|PD| = 11.4
|OD| = 17.6
|OD| / |PD| = 1.55
SN max = 5
For segments of SN = 1, 2, 3, 4:
No. of segments = 159, 83, 43, 13
Path length[µm] = 194.1, 120.6, 49.7, 11.6
Vector sum length [µm] = 41.3, 50, 25.4, 8.2
15
10
5
0
−5
−10
−15
−20
−15
−10
−5
0
5
10

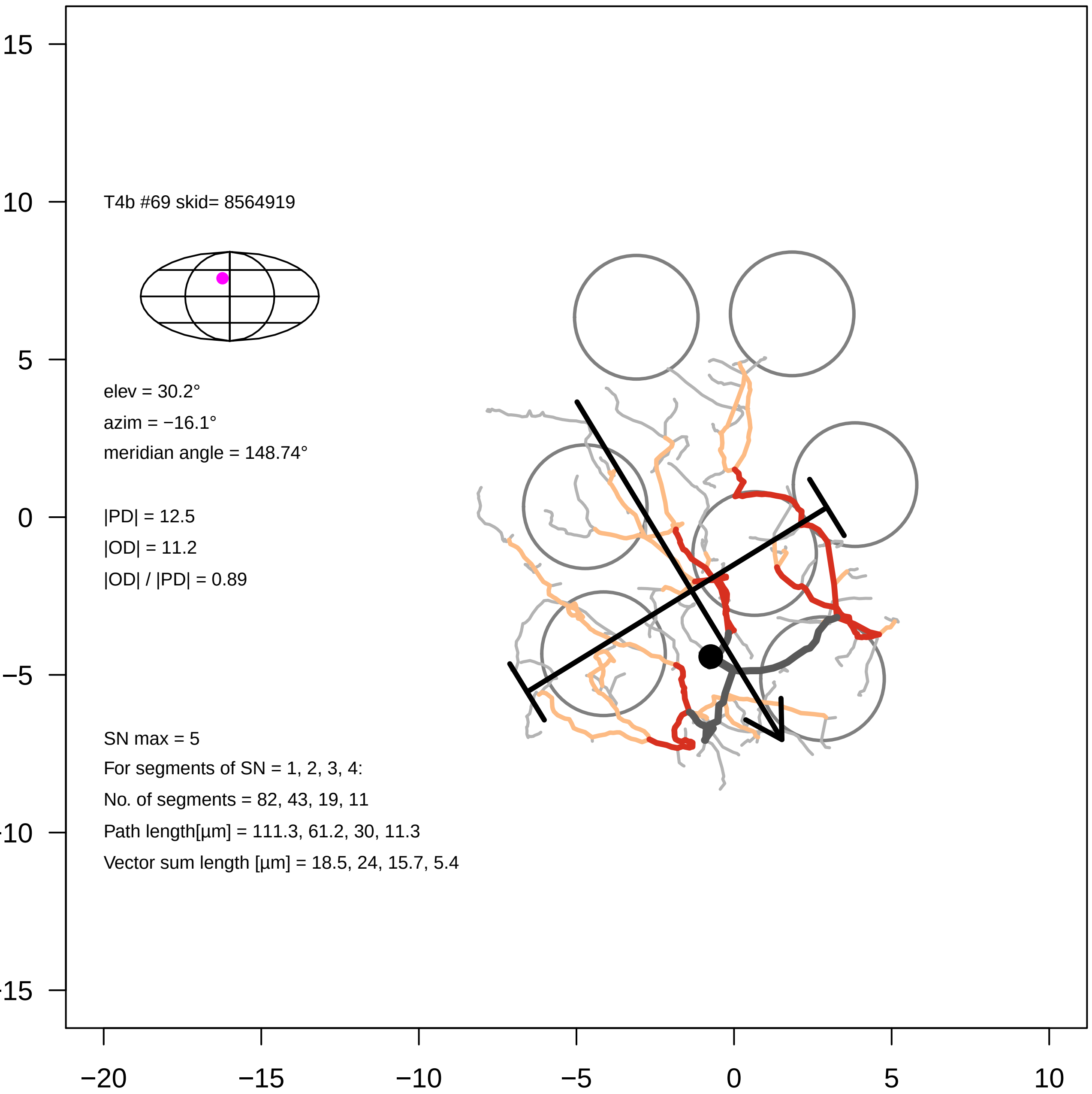
T4b #69 skid= 8564919
elev = 30.2°
azim = −16.1°
meridian angle = 148.74°
|PD| = 12.5
|OD| = 11.2
|OD| / |PD| = 0.89
SN max = 5
For segments of SN = 1, 2, 3, 4:
No. of segments = 82, 43, 19, 11
Path length[µm] = 111.3, 61.2, 30, 11.3
Vector sum length [µm] = 18.5, 24, 15.7, 5.4
15
10
5
0
−5
−10
−15
−20
−15
−10
−5
0
5
10

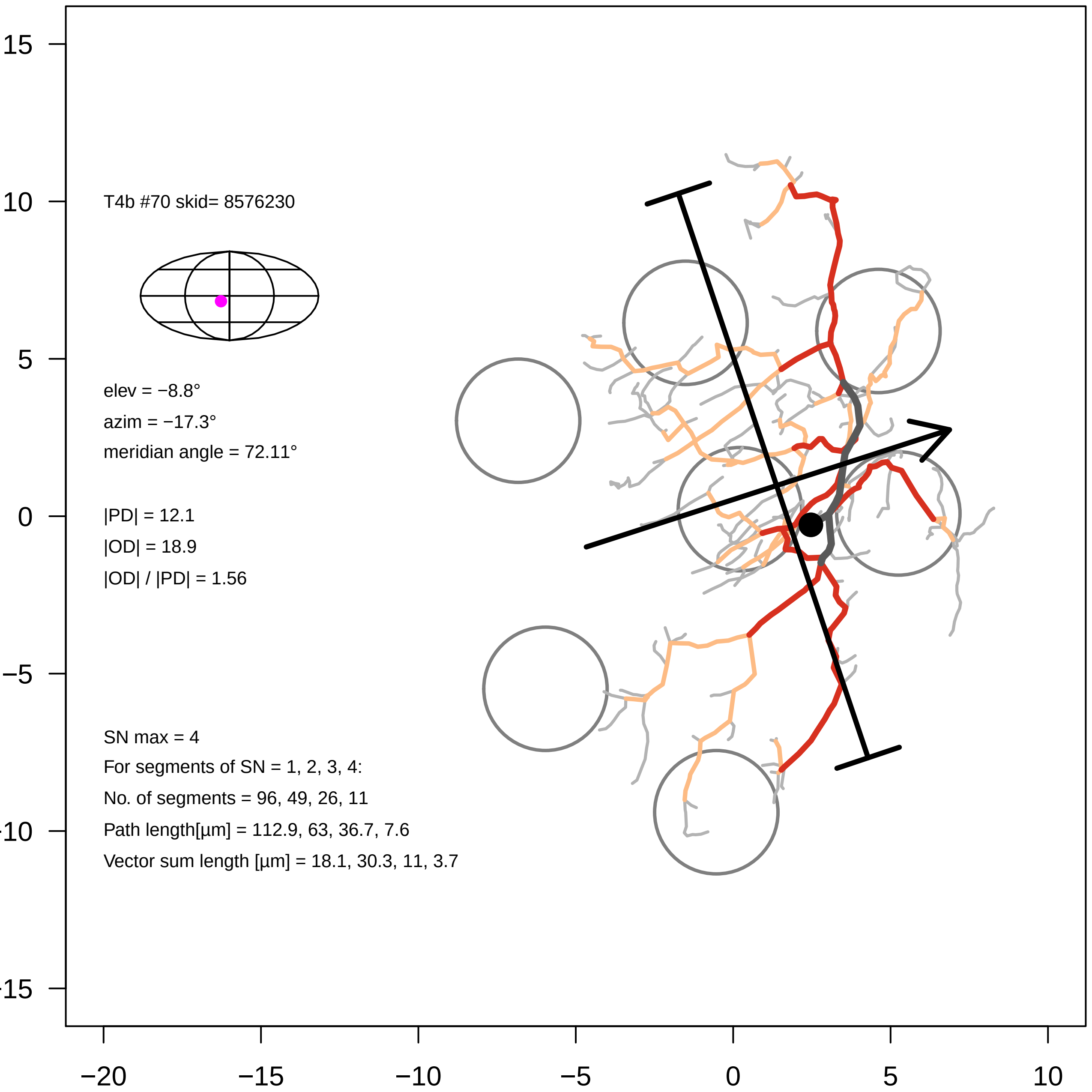

T4b #70 skid= 8576230
elev = −8.8°
azim = −17.3°
meridian angle = 72.11°
|PD| = 12.1
|OD| = 18.9
|OD| / |PD| = 1.56
SN max = 4
For segments of SN = 1, 2, 3, 4:
No. of segments = 96, 49, 26, 11
Path length[µm] = 112.9, 63, 36.7, 7.6
Vector sum length [µm] = 18.1, 30.3, 11, 3.7
15
10
5
0
−5
−10
−15
−20
−15
−10
−5
0
5
10

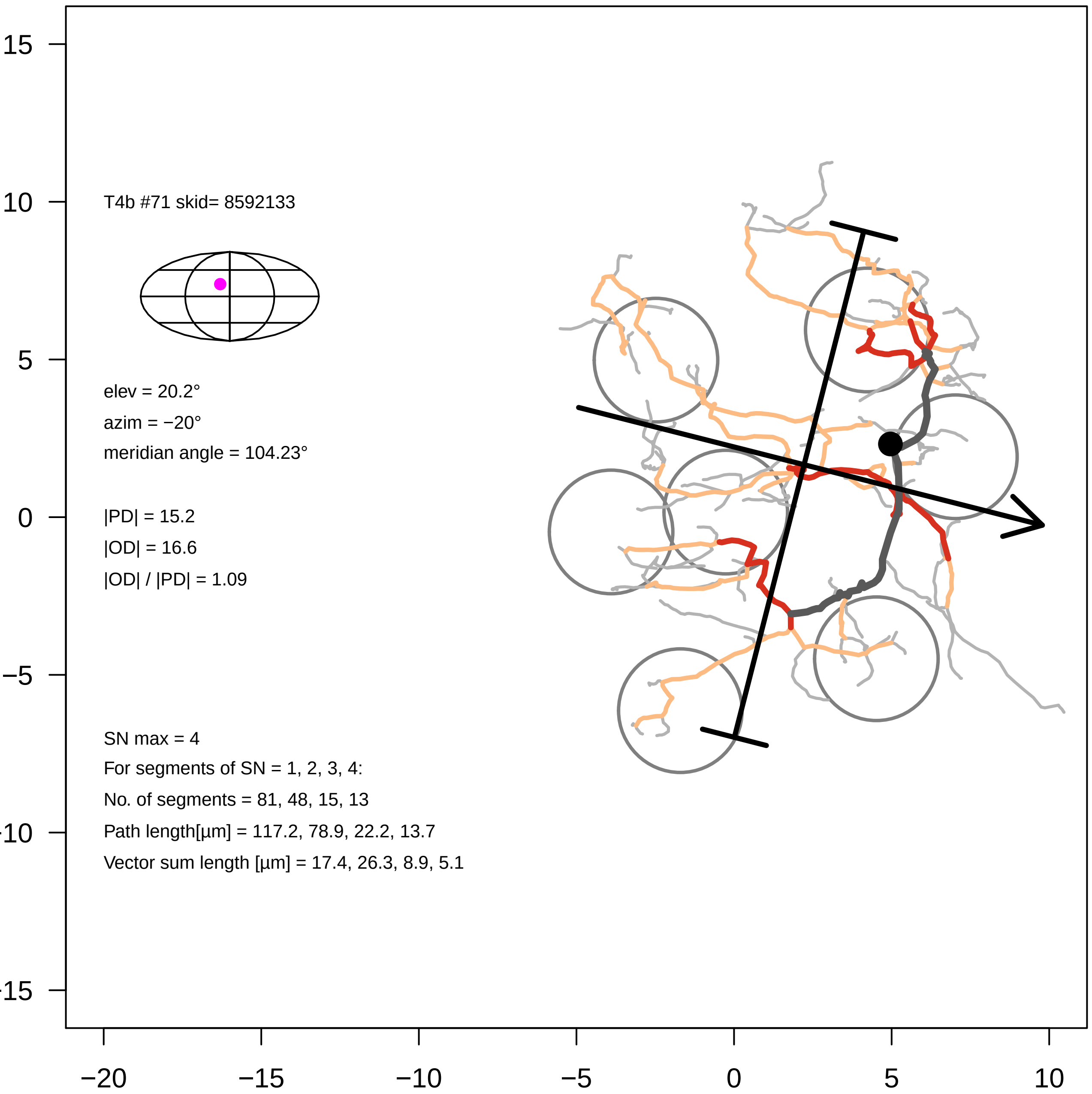
T4b #71 skid= 8592133
elev = 20.2°
azim = −20°
meridian angle = 104.23°
|PD| = 15.2
|OD| = 16.6
|OD| / |PD| = 1.09
SN max = 4
For segments of SN = 1, 2, 3, 4:
No. of segments = 81, 48, 15, 13
Path length[µm] = 117.2, 78.9, 22.2, 13.7
Vector sum length [µm] = 17.4, 26.3, 8.9, 5.1
15
10
5
0
−5
−10
−15
−20
−15
−10
−5
0
5
10

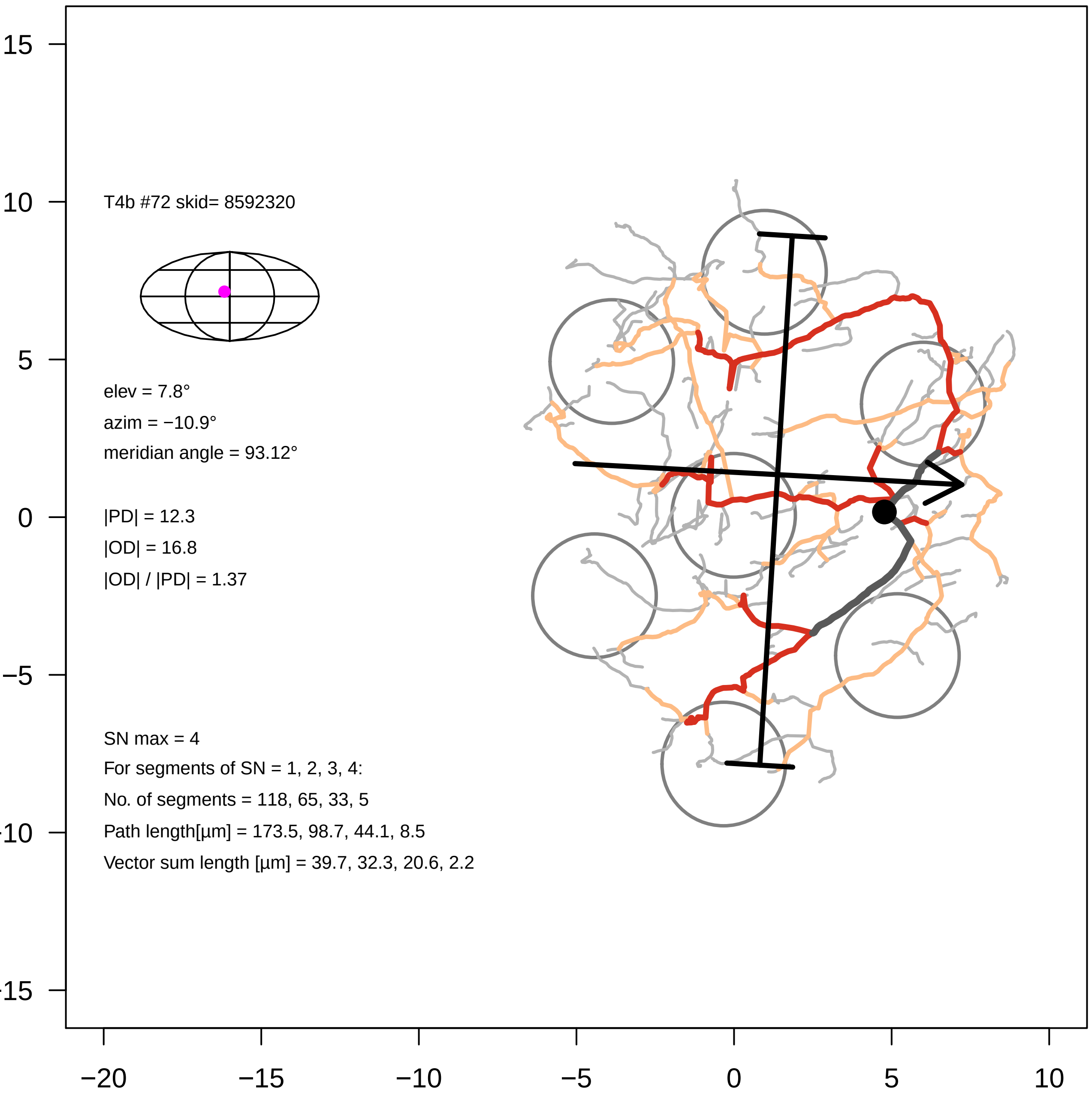
T4b #72 skid= 8592320
elev = 7.8°
azim = −10.9°
meridian angle = 93.12°
|PD| = 12.3
|OD| = 16.8
|OD| / |PD| = 1.37
SN max = 4
For segments of SN = 1, 2, 3, 4:
No. of segments = 118, 65, 33, 5
Path length[µm] = 173.5, 98.7, 44.1, 8.5
Vector sum length [µm] = 39.7, 32.3, 20.6, 2.2
15
10
5
0
−5
−20
−15
−10
−5
0
5
10

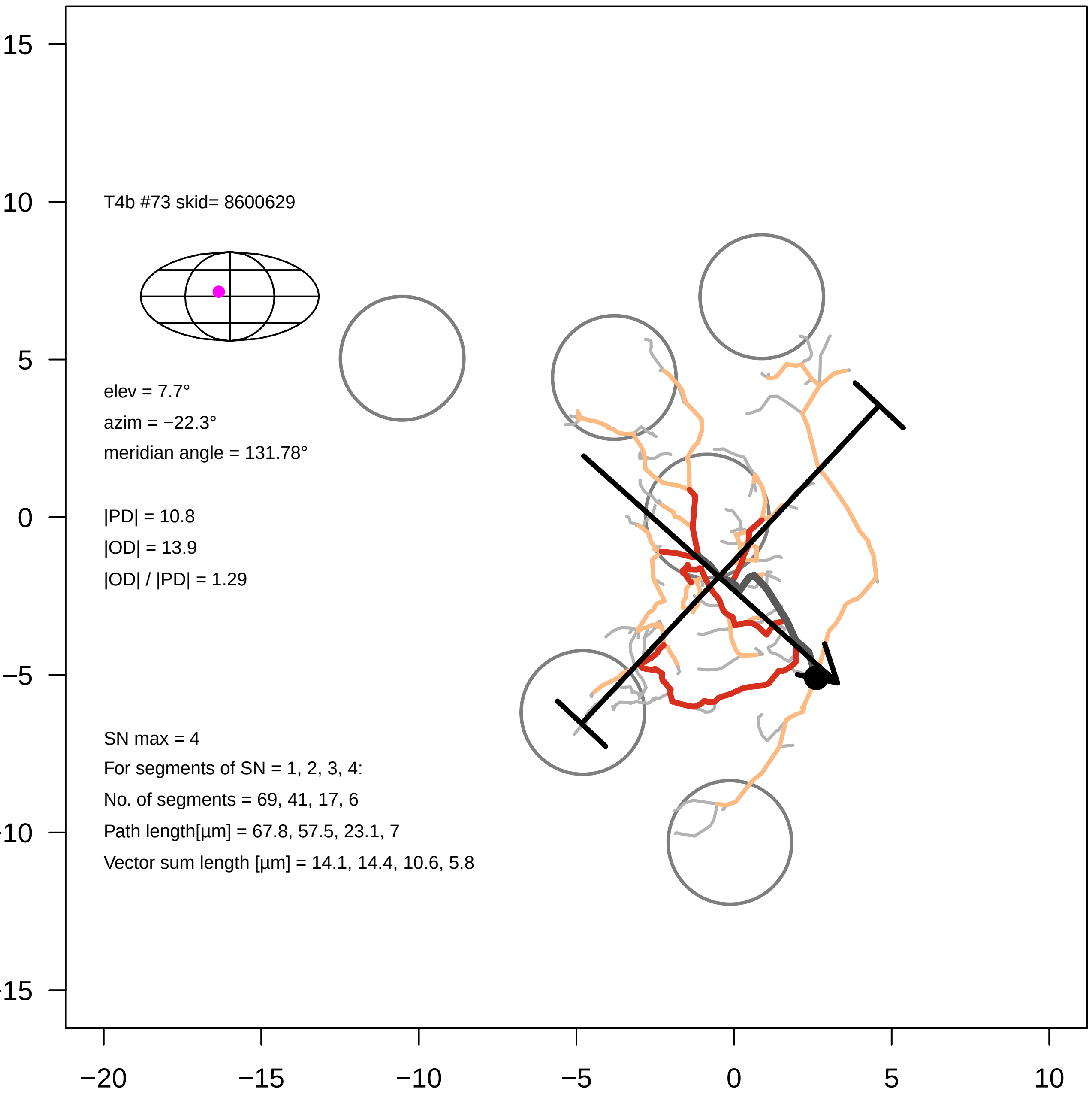

T4b #73 skid= 8600629
elev = 7.7°
azim = −22.3°
meridian angle = 131.78°
|PD| = 10.8
|OD| = 13.9
|OD| / |PD| = 1.29
SN max = 4
For segments of SN = 1, 2, 3, 4:
No. of segments = 69, 41, 17, 6
Path length[µm] = 67.8, 57.5, 23.1, 7
Vector sum length [µm] = 14.1, 14.4, 10.6, 5.8
15
10
5
0
−5
−10
−15
−20
−15
−10
−5
0
5
10

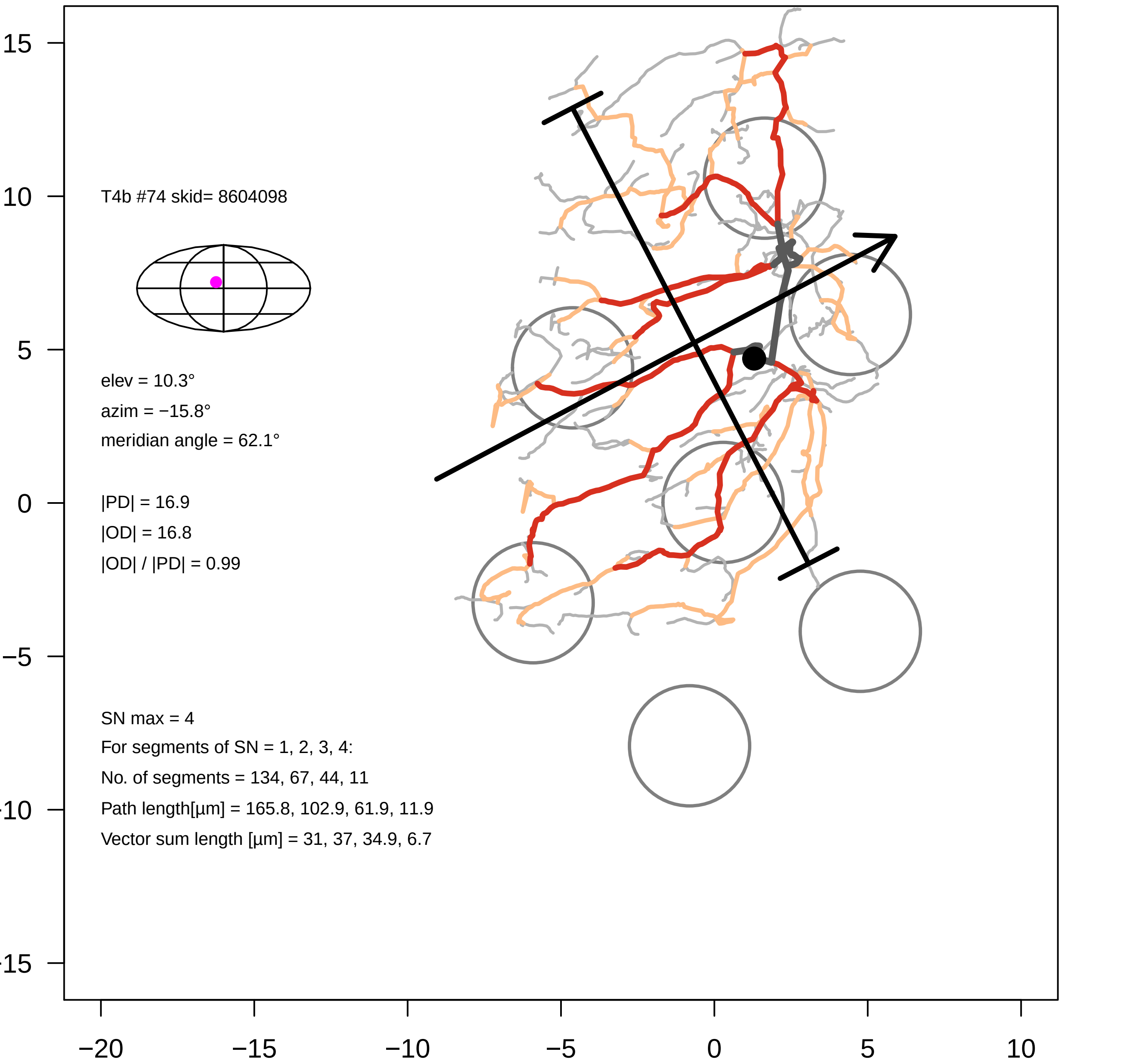

T4b #74 skid= 8604098
elev = 10.3°
azim = −15.8°
meridian angle = 62.1°
|PD| = 16.9
|OD| = 16.8
|OD| / |PD| = 0.99
SN max = 4
For segments of SN = 1, 2, 3, 4:
No. of segments = 134, 67, 44, 11
Path length[µm] = 165.8, 102.9, 61.9, 11.9
Vector sum length [µm] = 31, 37, 34.9, 6.7
15
10
5
0
−5
−10
−15
−20
−15
−10
−5
0
5
10

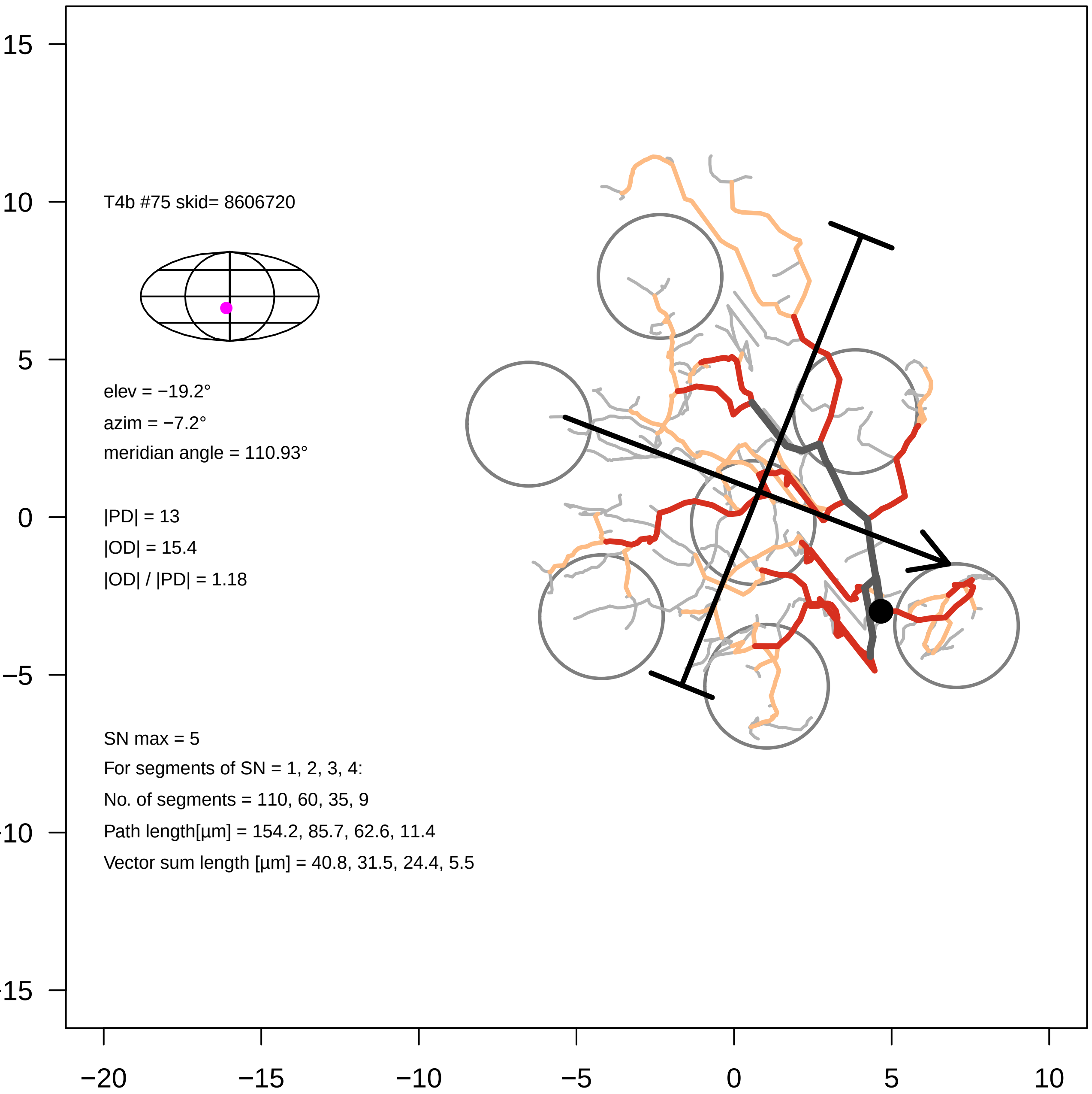

15
10
5
0
−5
−10
−15
−20
−15
−10
−5
0
5
10
T4b #75 skid= 8606720
elev = −19.2°
azim = −7.2°
meridian angle = 110.93°
|PD| = 13
|OD| = 15.4
|OD| / |PD| = 1.18
SN max = 5
For segments of SN = 1, 2, 3, 4:
No. of segments = 110, 60, 35, 9
Path length[µm] = 154.2, 85.7, 62.6, 11.4
Vector sum length [µm] = 40.8, 31.5, 24.4, 5.5

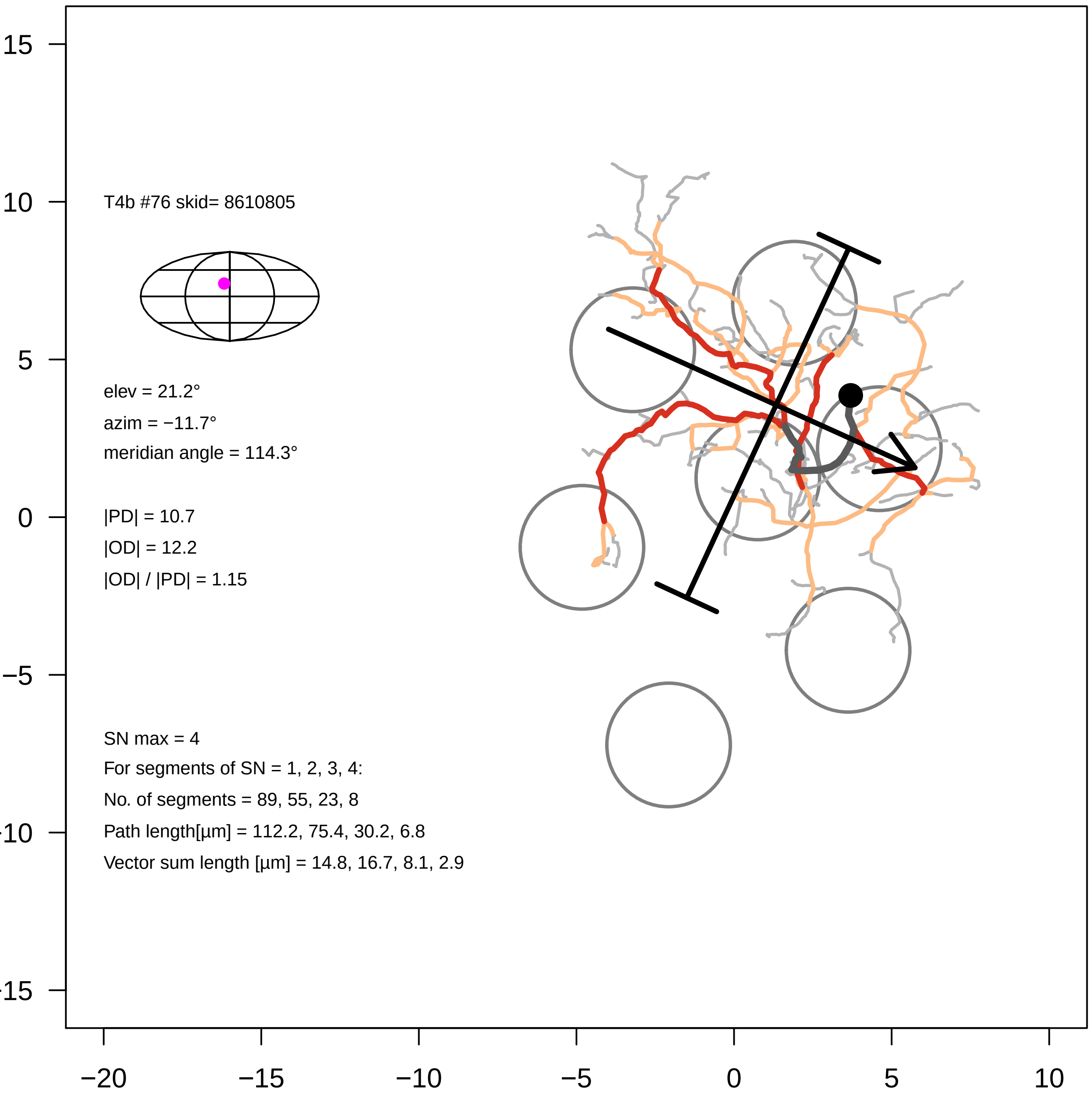

15
10
5
0
−5
−10
−15
−20
−15
−10
−5
0
5
10
T4b #76 skid= 8610805
elev = 21.2°
azim = −11.7°
meridian angle = 114.3°
|PD| = 10.7
|OD| = 12.2
|OD| / |PD| = 1.15
SN max = 4
For segments of SN = 1, 2, 3, 4:
No. of segments = 89, 55, 23, 8
Path length[µm] = 112.2, 75.4, 30.2, 6.8
Vector sum length [µm] = 14.8, 16.7, 8.1, 2.9

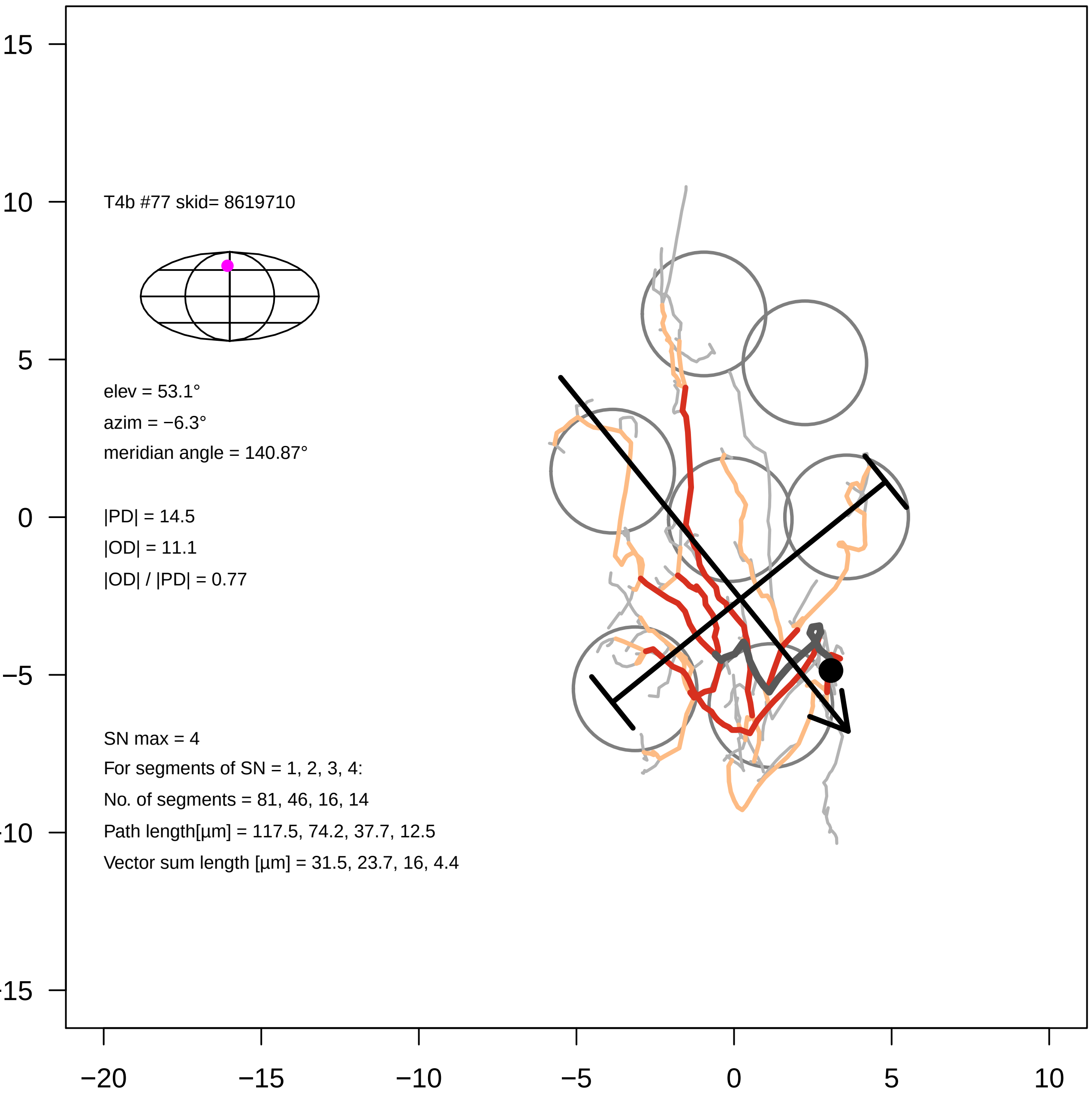

T4b #77 skid= 8619710
elev = 53.1°
azim = −6.3°
meridian angle = 140.87°
|PD| = 14.5
|OD| = 11.1
|OD| / |PD| = 0.77
SN max = 4
For segments of SN = 1, 2, 3, 4:
No. of segments = 81, 46, 16, 14
Path length[µm] = 117.5, 74.2, 37.7, 12.5
Vector sum length [µm] = 31.5, 23.7, 16, 4.4
15
10
5
0
−5
−10
−15
−20
−15
−10
−5
0
5
10

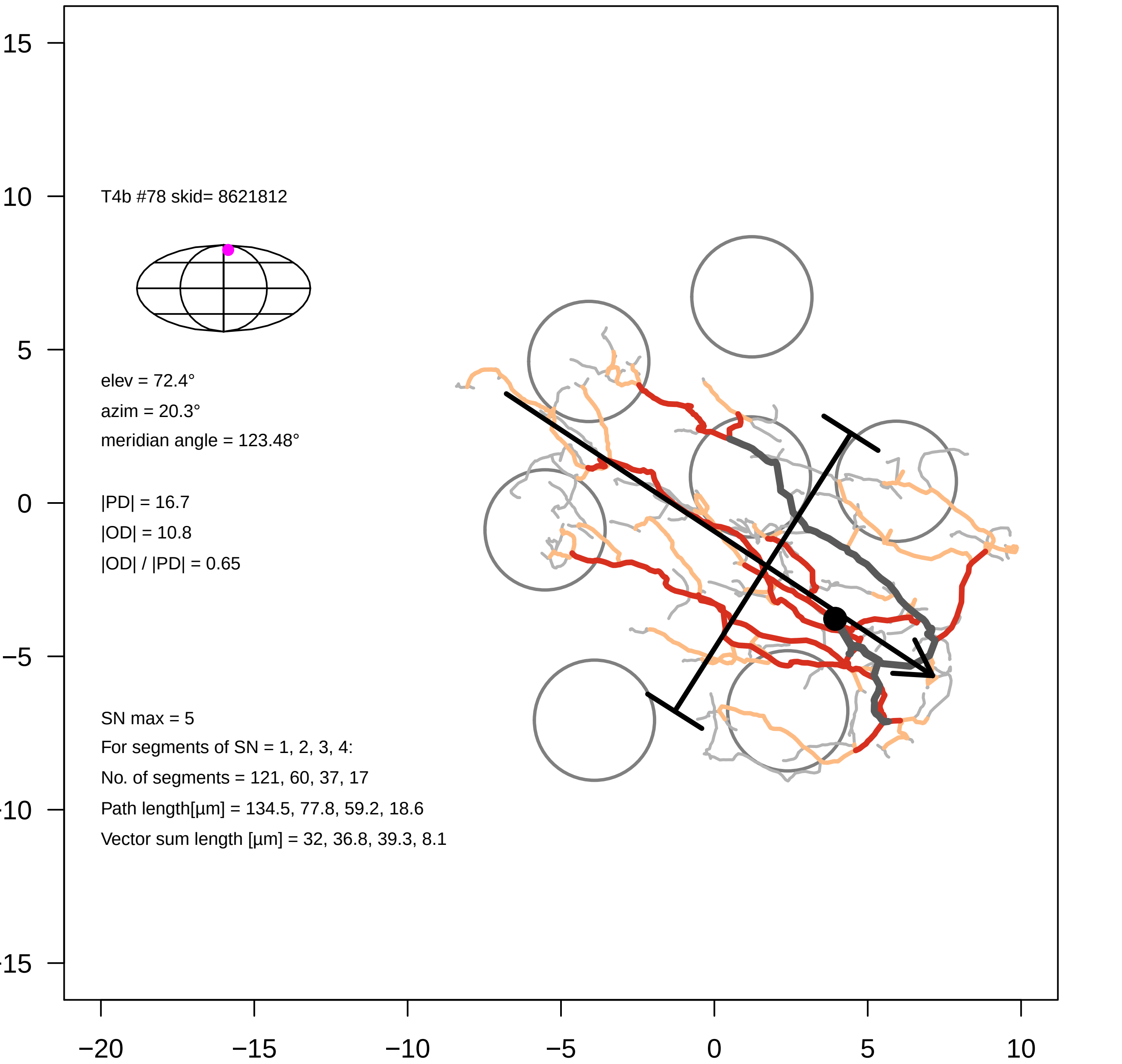

15
10
5
0
−5
−10
−15
−20
−15
−10
−5
0
5
10
T4b #78 skid= 8621812
elev = 72.4°
azim = 20.3°
meridian angle = 123.48°
|PD| = 16.7
|OD| = 10.8
|OD| / |PD| = 0.65
SN max = 5
For segments of SN = 1, 2, 3, 4:
No. of segments = 121, 60, 37, 17
Path length[µm] = 134.5, 77.8, 59.2, 18.6
Vector sum length [µm] = 32, 36.8, 39.3, 8.1

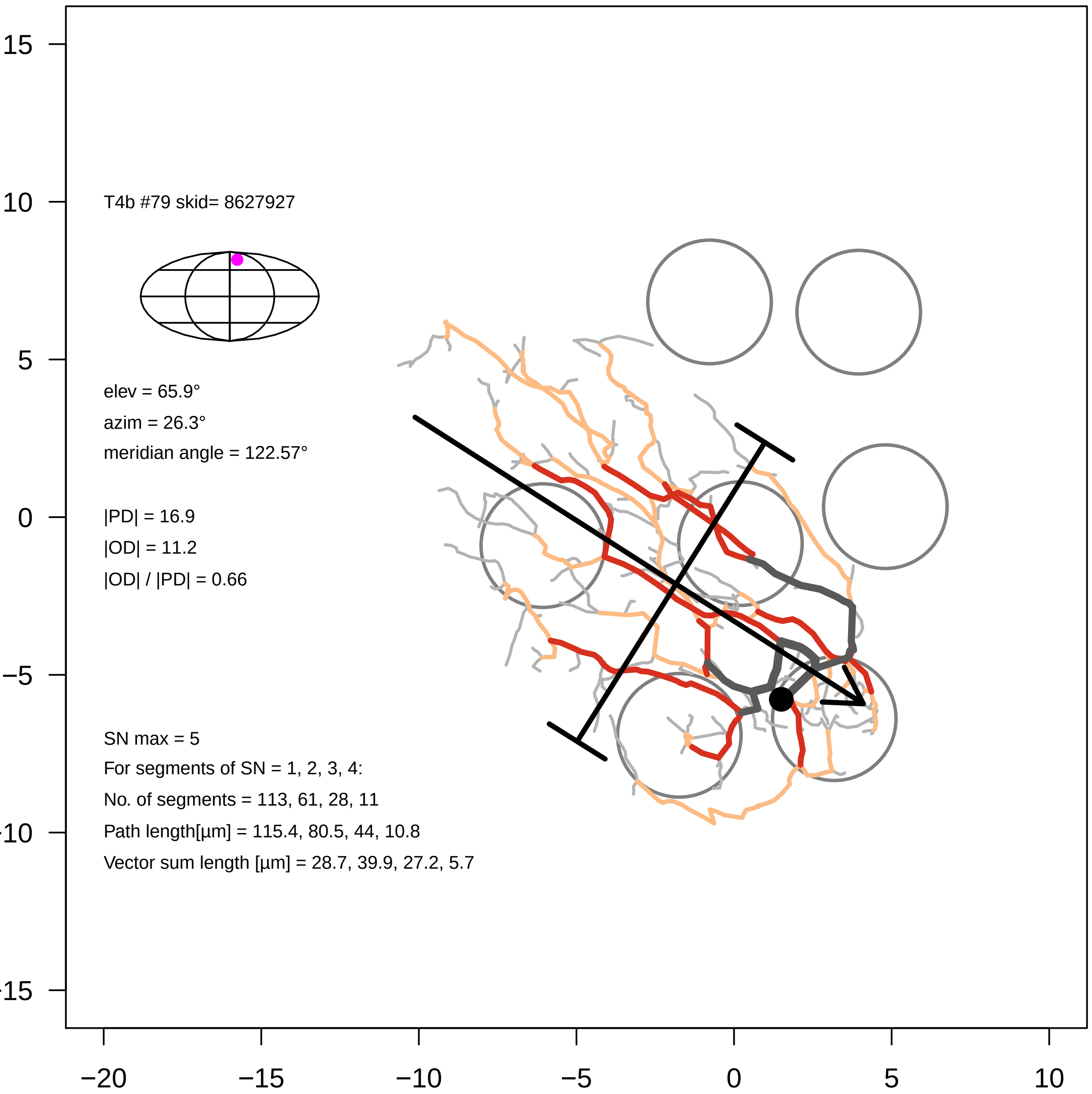

T4b #79 skid= 8627927
elev = 65.9°
azim = 26.3°
meridian angle = 122.57°
|PD| = 16.9
|OD| = 11.2
|OD| / |PD| = 0.66
SN max = 5
For segments of SN = 1, 2, 3, 4:
No. of segments = 113, 61, 28, 11
Path length[µm] = 115.4, 80.5, 44, 10.8
Vector sum length [µm] = 28.7, 39.9, 27.2, 5.7
15
10
5
0
−5
−10
−15
−20
−15
−10
−5
0
5
10

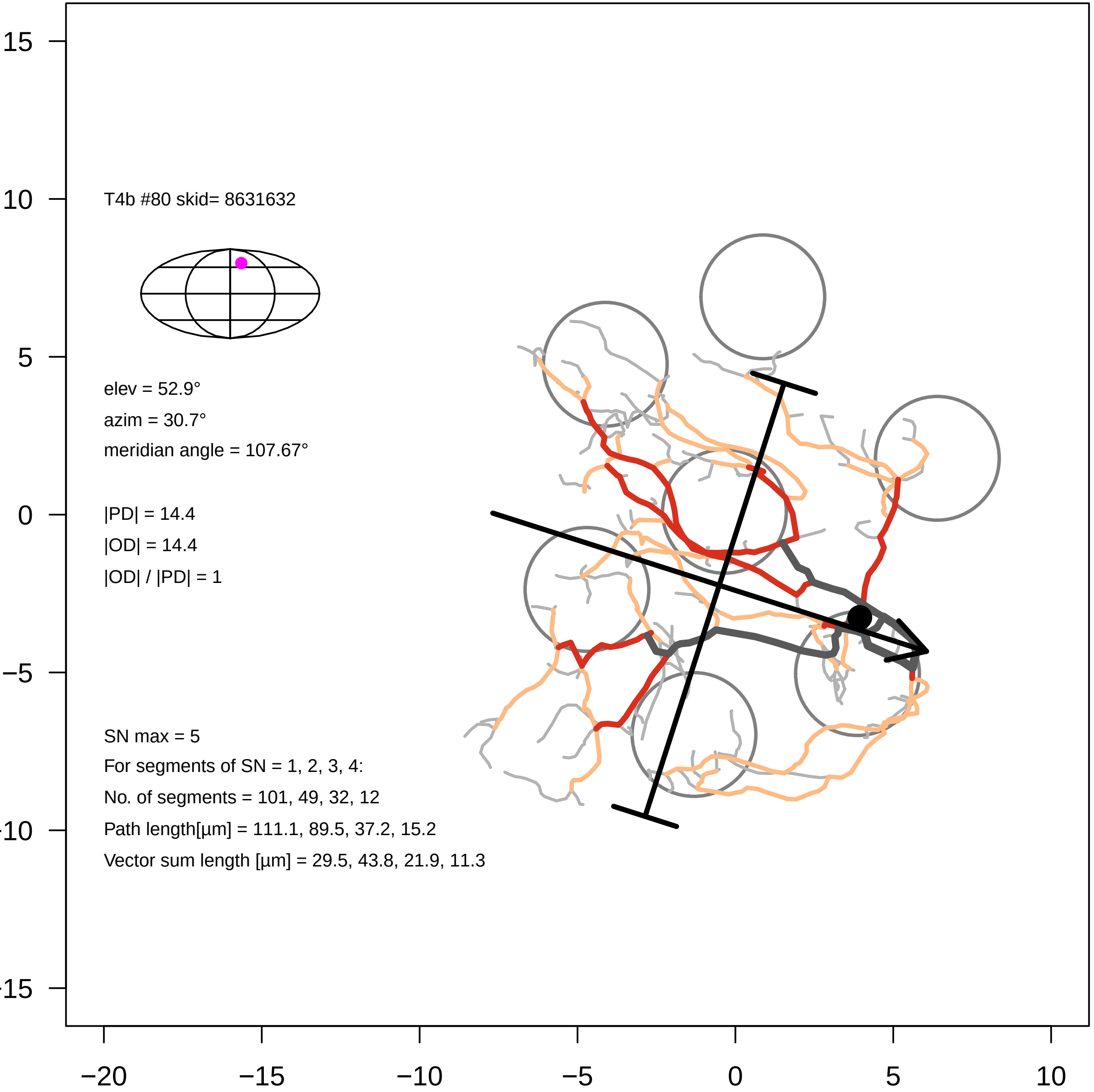
T4b #80 skid= 8631632
elev = 52.9°
azim = 30.7°
meridian angle = 107.67°
|PD| = 14.4
|OD| = 14.4
|OD| / |PD| = 1
SN max = 5
For segments of SN = 1, 2, 3, 4:
No. of segments = 101, 49, 32, 12
Path length[µm] = 111.1, 89.5, 37.2, 15.2
Vector sum length [µm] = 29.5, 43.8, 21.9, 11.3
15
10
5
0
−5
−10
−15
−20
−15
−10
−5
0
5
10

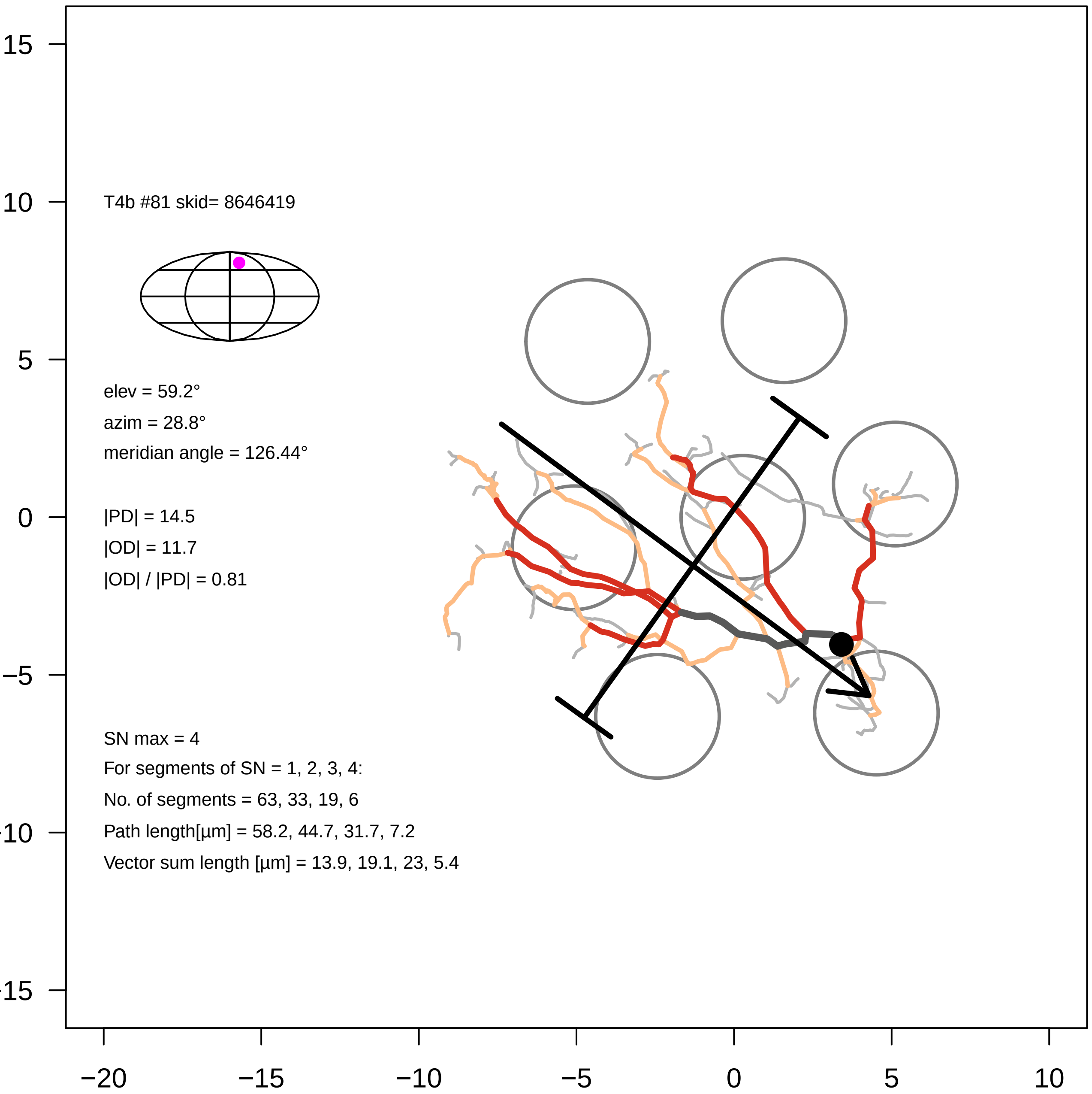

T4b #81 skid= 8646419
elev = 59.2°
azim = 28.8°
meridian angle = 126.44°
|PD| = 14.5
|OD| = 11.7
|OD| / |PD| = 0.81
SN max = 4
For segments of SN = 1, 2, 3, 4:
No. of segments = 63, 33, 19, 6
Path length[µm] = 58.2, 44.7, 31.7, 7.2
Vector sum length [µm] = 13.9, 19.1, 23, 5.4
15
10
5
0
−5
−10
−15
−20
−15
−10
−5
0
5
10

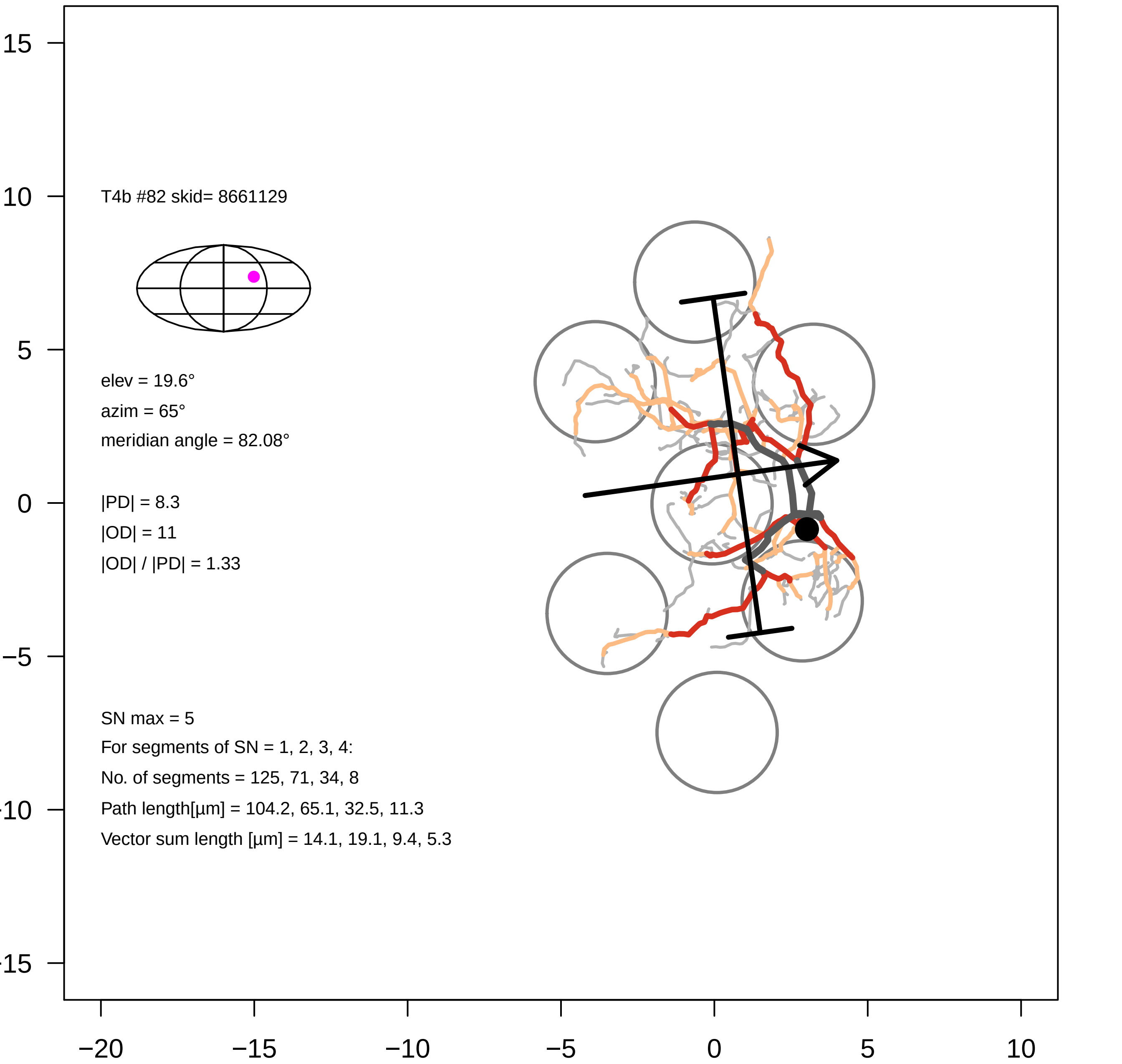

T4b #82 skid= 8661129
elev = 19.6°
azim = 65°
meridian angle = 82.08°
|PD| = 8.3
|OD| = 11
|OD| / |PD| = 1.33
SN max = 5
For segments of SN = 1, 2, 3, 4:
No. of segments = 125, 71, 34, 8
Path length[µm] = 104.2, 65.1, 32.5, 11.3
Vector sum length [µm] = 14.1, 19.1, 9.4, 5.3
15
10
5
0
−5
−10
−15
−20
−15
−10
−5
0
5
10

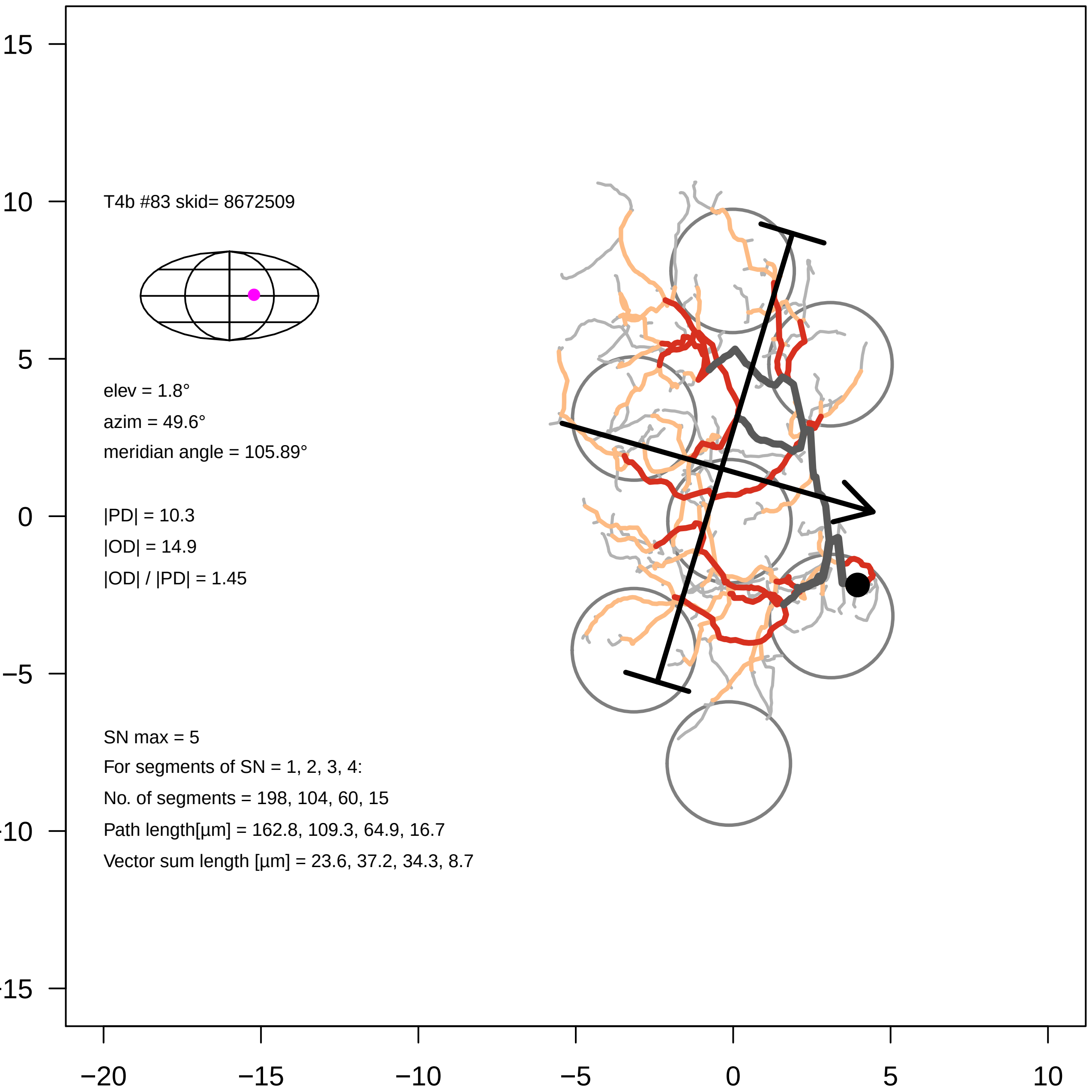
T4b #83 skid= 8672509
elev = 1.8°
azim = 49.6°
meridian angle = 105.89°
|PD| = 10.3
|OD| = 14.9
|OD| / |PD| = 1.45
SN max = 5
For segments of SN = 1, 2, 3, 4:
No. of segments = 198, 104, 60, 15
Path length[µm] = 162.8, 109.3, 64.9, 16.7
Vector sum length [µm] = 23.6, 37.2, 34.3, 8.7
15
10
5
0
−5
−10
−15
−20
−15
−10
−5
0
5
10

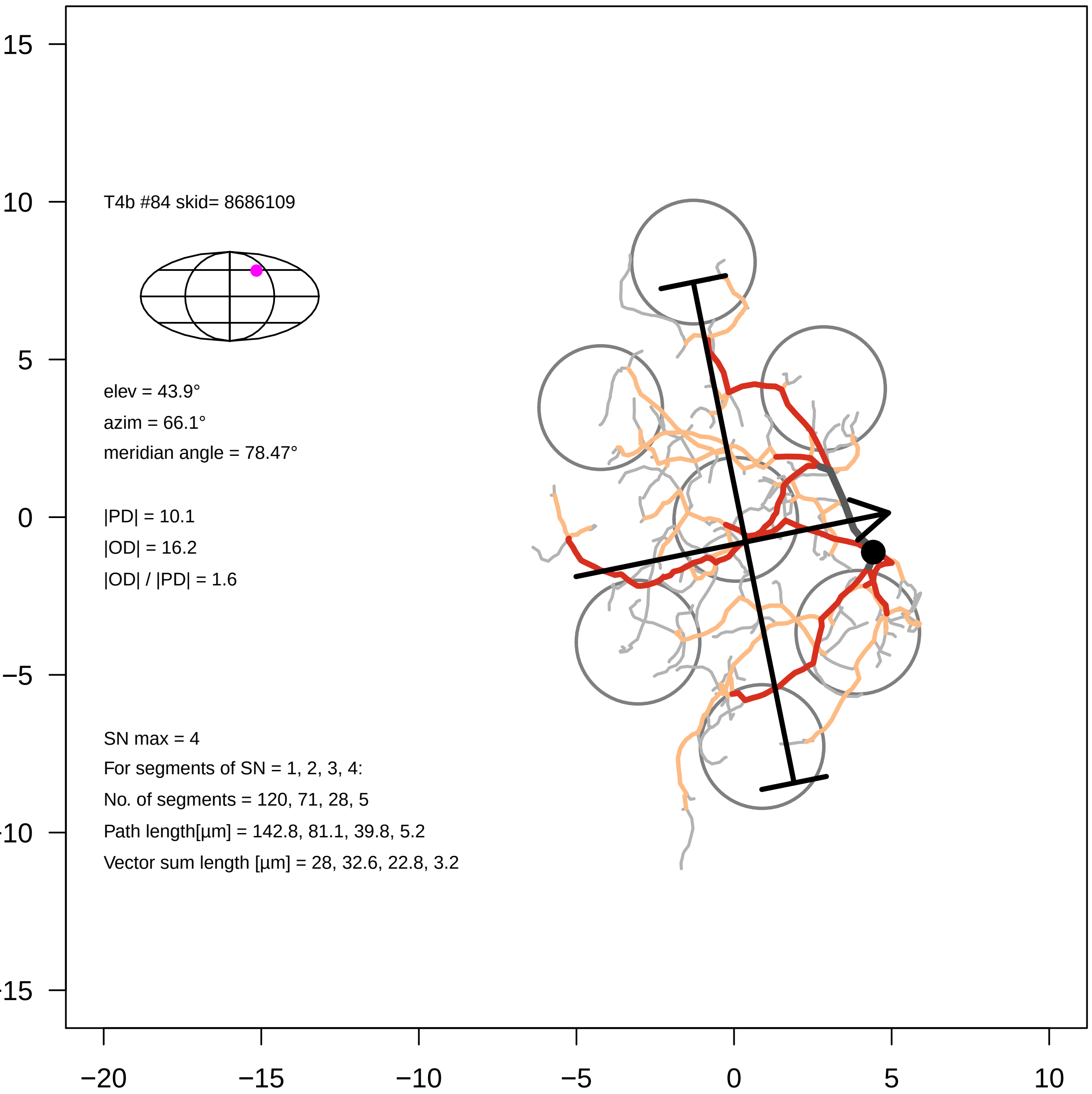
T4b #84 skid= 8686109
elev = 43.9°
azim = 66.1°
meridian angle = 78.47°
|PD| = 10.1
|OD| = 16.2
|OD| / |PD| = 1.6
SN max = 4
For segments of SN = 1, 2, 3, 4:
No. of segments = 120, 71, 28, 5
Path length[µm] = 142.8, 81.1, 39.8, 5.2
Vector sum length [µm] = 28, 32.6, 22.8, 3.2
15
10
5
0
−5
−10
−15
−20
−15
−10
−5
0
5
10

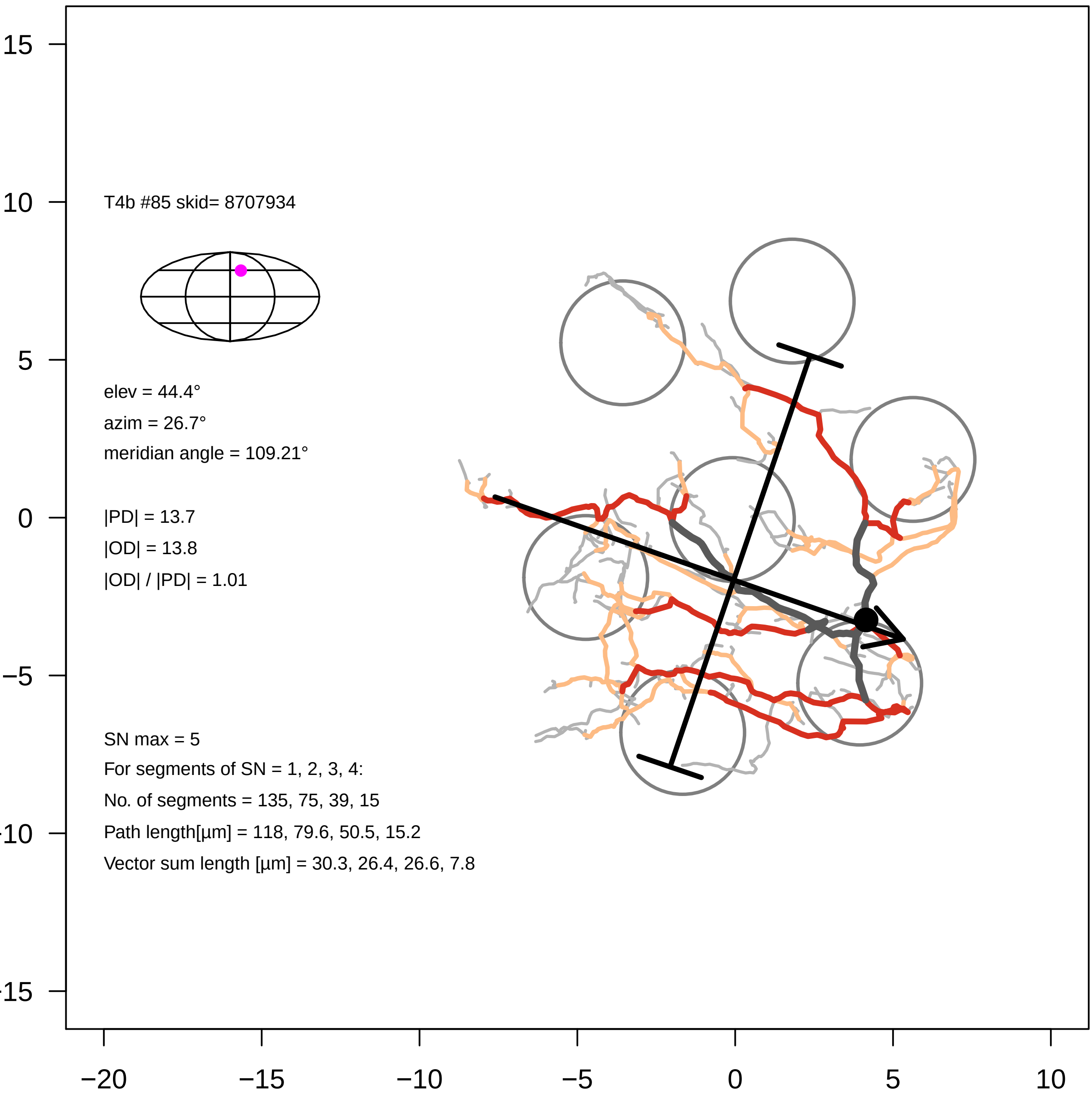
T4b #85 skid= 8707934
elev = 44.4°
azim = 26.7°
meridian angle = 109.21°
|PD| = 13.7
|OD| = 13.8
|OD| / |PD| = 1.01
SN max = 5
For segments of SN = 1, 2, 3, 4:
No. of segments = 135, 75, 39, 15
Path length[µm] = 118, 79.6, 50.5, 15.2
Vector sum length [µm] = 30.3, 26.4, 26.6, 7.8
15
10
5
0
−5
−10
−15
−20
−15
−10
−5
0
5
10

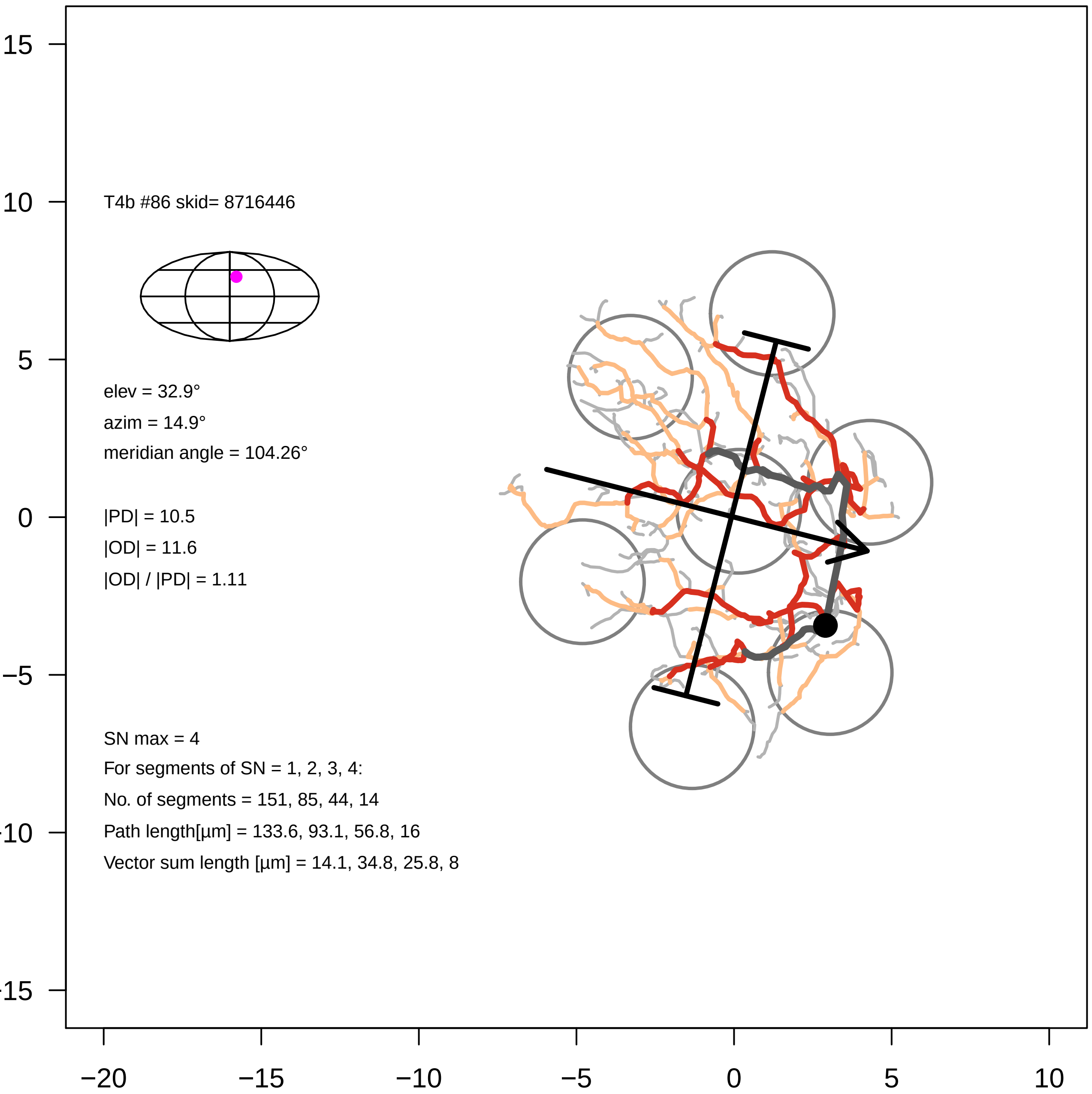

15
10
5
0
−5
−10
−15
−20
−15
−10
−5
0
5
10
T4b #86 skid= 8716446
elev = 32.9°
azim = 14.9°
meridian angle = 104.26°
|PD| = 10.5
|OD| = 11.6
|OD| / |PD| = 1.11
SN max = 4
For segments of SN = 1, 2, 3, 4:
No. of segments = 151, 85, 44, 14
Path length[µm] = 133.6, 93.1, 56.8, 16
Vector sum length [µm] = 14.1, 34.8, 25.8, 8

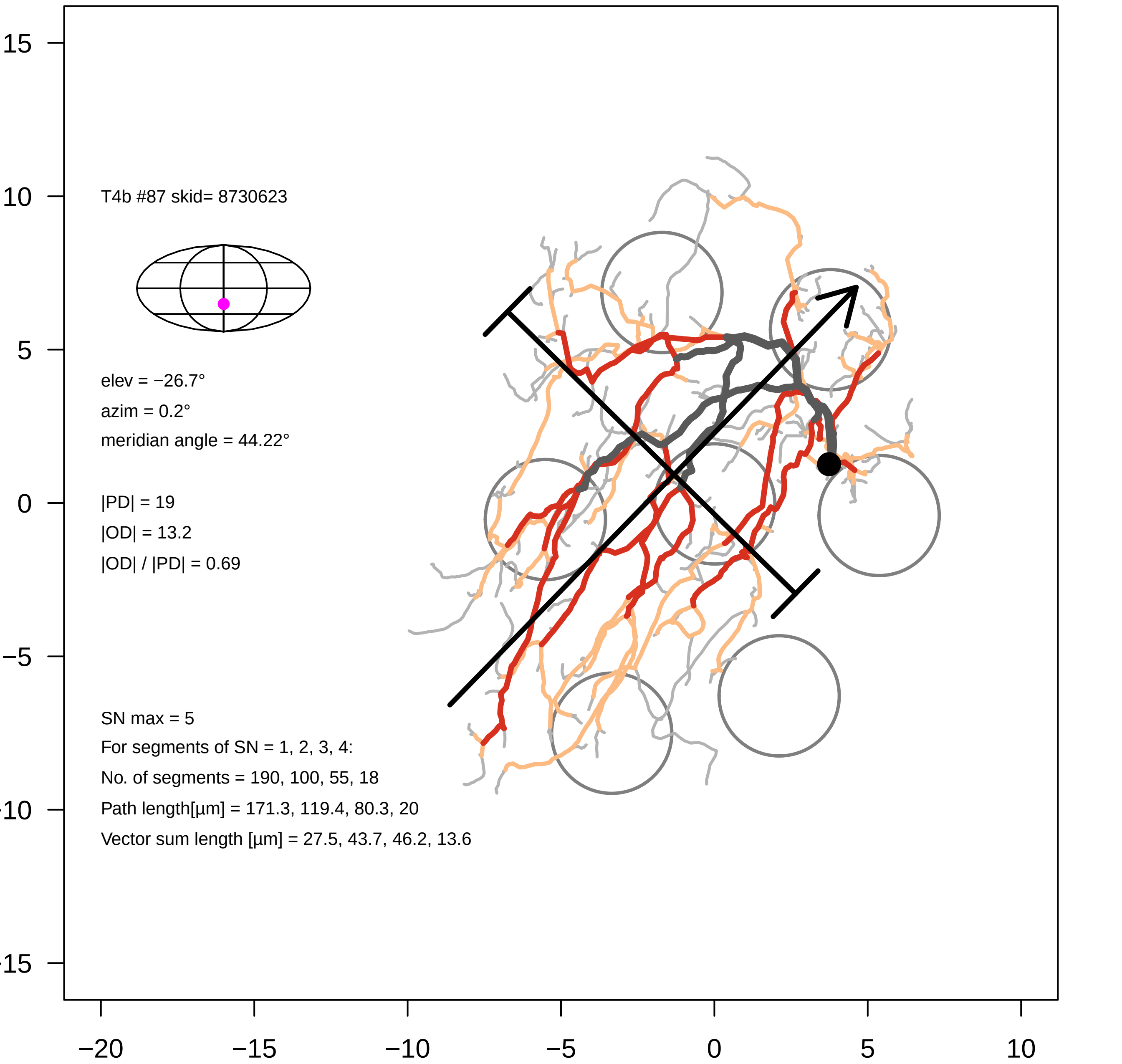

T4b #87 skid= 8730623
elev = −26.7°
azim = 0.2°
meridian angle = 44.22°
|PD| = 19
|OD| = 13.2
|OD| / |PD| = 0.69
SN max = 5
For segments of SN = 1, 2, 3, 4:
No. of segments = 190, 100, 55, 18
Path length[µm] = 171.3, 119.4, 80.3, 20
Vector sum length [µm] = 27.5, 43.7, 46.2, 13.6
15
10
5
0
−5
−10
−15
−20
−15
−10
−5
0
5
10

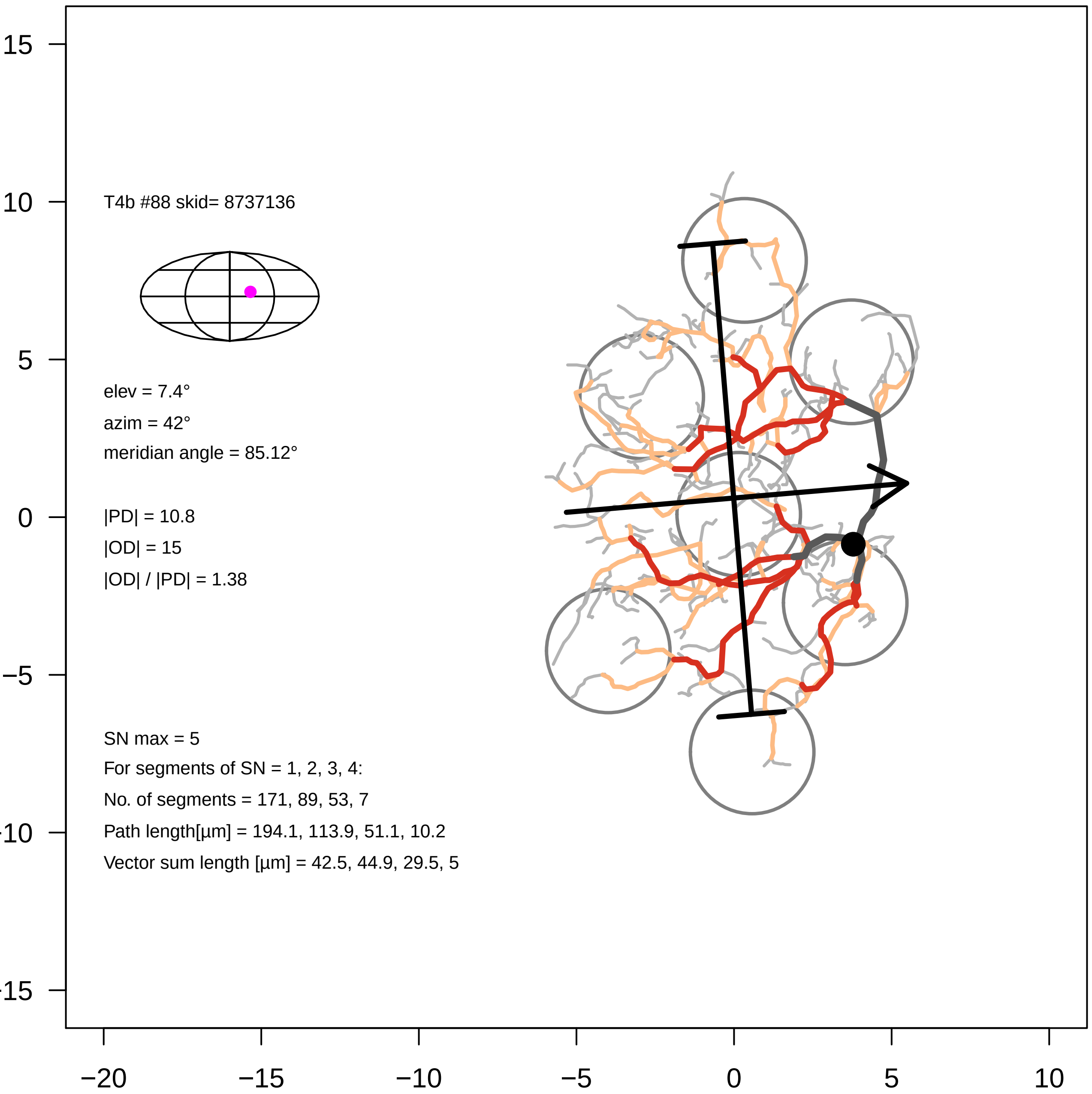
T4b #88 skid= 8737136
elev = 7.4°
azim = 42°
meridian angle = 85.12°
|PD| = 10.8
|OD| = 15
|OD| / |PD| = 1.38
SN max = 5
For segments of SN = 1, 2, 3, 4:
No. of segments = 171, 89, 53, 7
Path length[µm] = 194.1, 113.9, 51.1, 10.2
Vector sum length [µm] = 42.5, 44.9, 29.5, 5
15
10
5
0
−5
−10
−15
−20
−15
−10
−5
0
5
10

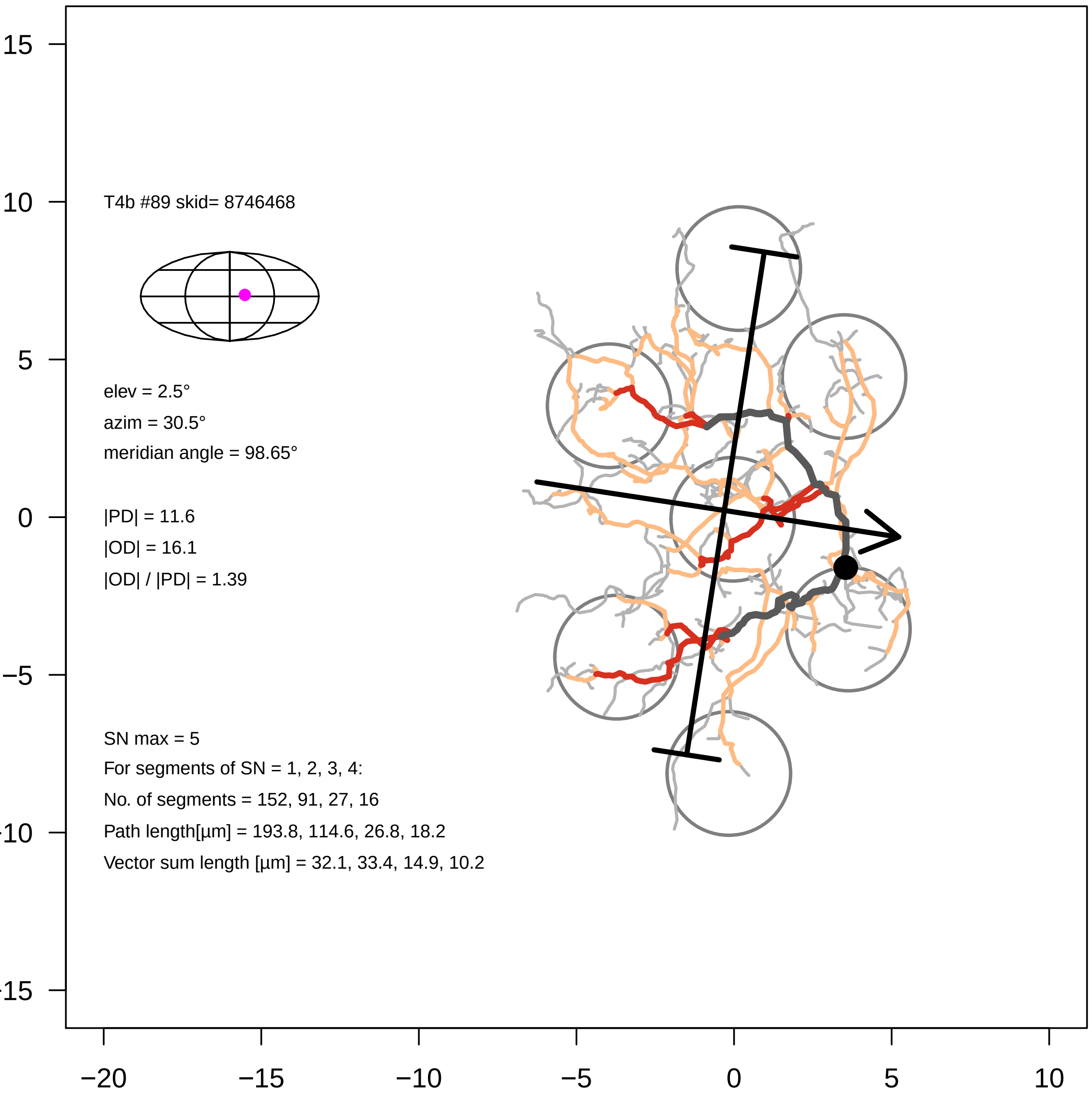

T4b #89 skid= 8746468
elev = 2.5°
azim = 30.5°
meridian angle = 98.65°
|PD| = 11.6
|OD| = 16.1
|OD| / |PD| = 1.39
SN max = 5
For segments of SN = 1, 2, 3, 4:
No. of segments = 152, 91, 27, 16
Path length[µm] = 193.8, 114.6, 26.8, 18.2
Vector sum length [µm] = 32.1, 33.4, 14.9, 10.2
15
10
5
0
−5
−10
−15
−20
−15
−10
−5
0
5
10

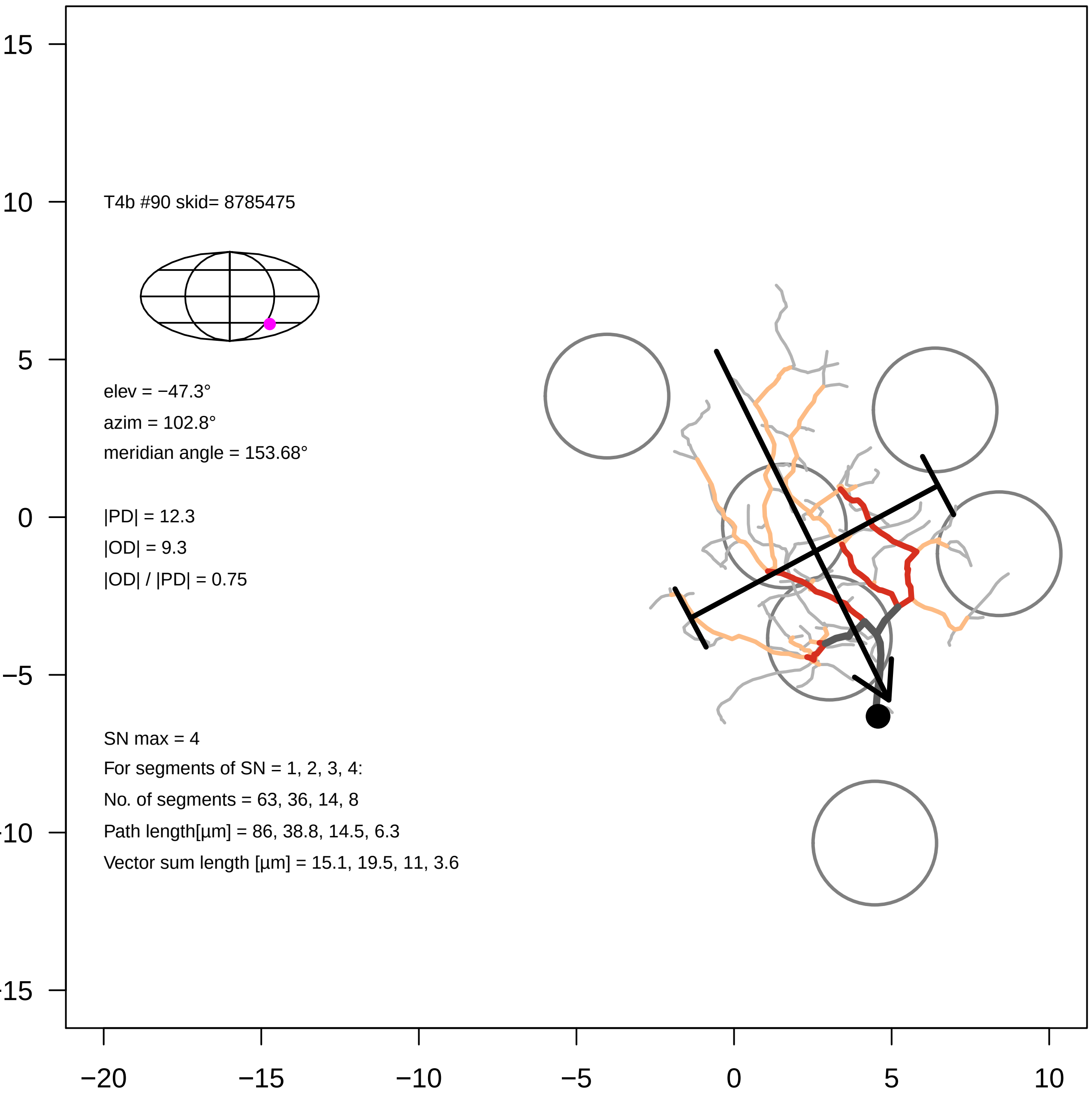

T4b #90 skid= 8785475
elev = −47.3°
azim = 102.8°
meridian angle = 153.68°
|PD| = 12.3
|OD| = 9.3
|OD| / |PD| = 0.75
SN max = 4
For segments of SN = 1, 2, 3, 4:
No. of segments = 63, 36, 14, 8
Path length[µm] = 86, 38.8, 14.5, 6.3
Vector sum length [µm] = 15.1, 19.5, 11, 3.6
15
10
5
0
−5
−10
−15
−20
−15
−10
−5
0
5
10

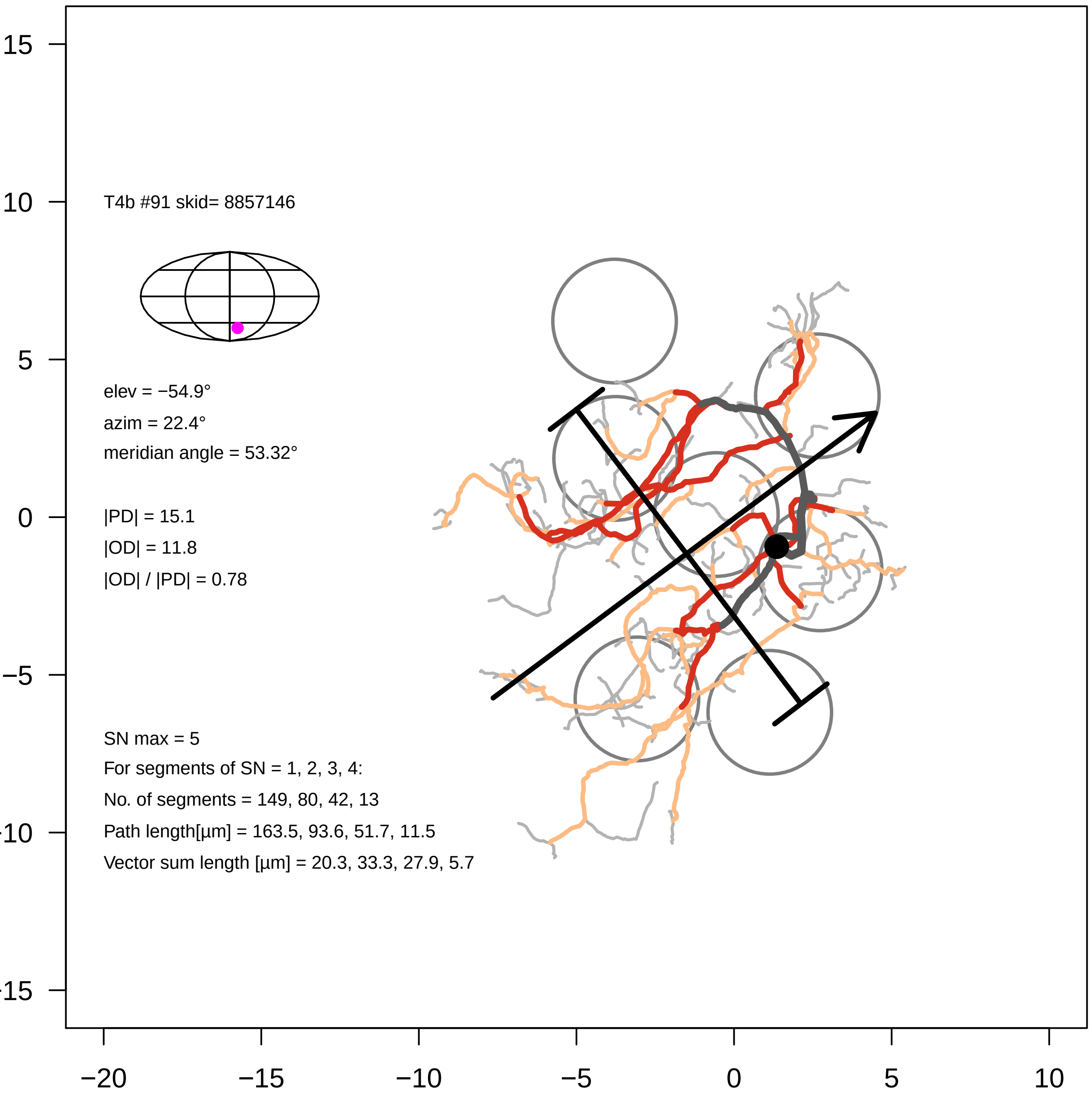
T4b #91 skid= 8857146
elev = −54.9°
azim = 22.4°
meridian angle = 53.32°
|PD| = 15.1
|OD| = 11.8
|OD| / |PD| = 0.78
SN max = 5
For segments of SN = 1, 2, 3, 4:
No. of segments = 149, 80, 42, 13
Path length[µm] = 163.5, 93.6, 51.7, 11.5
Vector sum length [µm] = 20.3, 33.3, 27.9, 5.7
15
10
5
0
−5
−10
−15
−20
−15
−10
−5
0
5
10

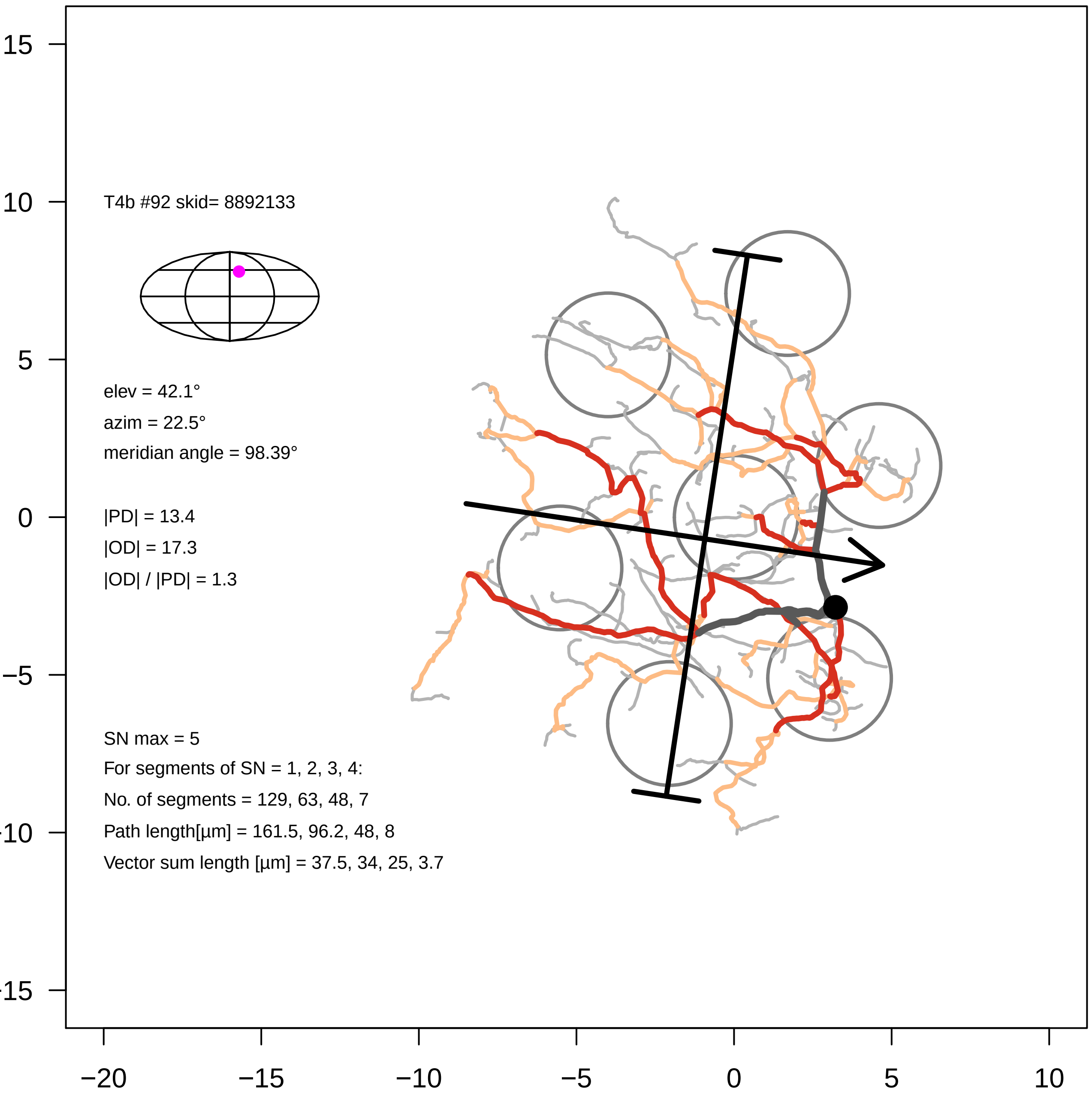

T4b #92 skid= 8892133
elev = 42.1°
azim = 22.5°
meridian angle = 98.39°
|PD| = 13.4
|OD| = 17.3
|OD| / |PD| = 1.3
SN max = 5
For segments of SN = 1, 2, 3, 4:
No. of segments = 129, 63, 48, 7
Path length[µm] = 161.5, 96.2, 48, 8
Vector sum length [µm] = 37.5, 34, 25, 3.7
15
10
5
0
−5
−10
−15
−20
−15
−10
−5
0
5
10

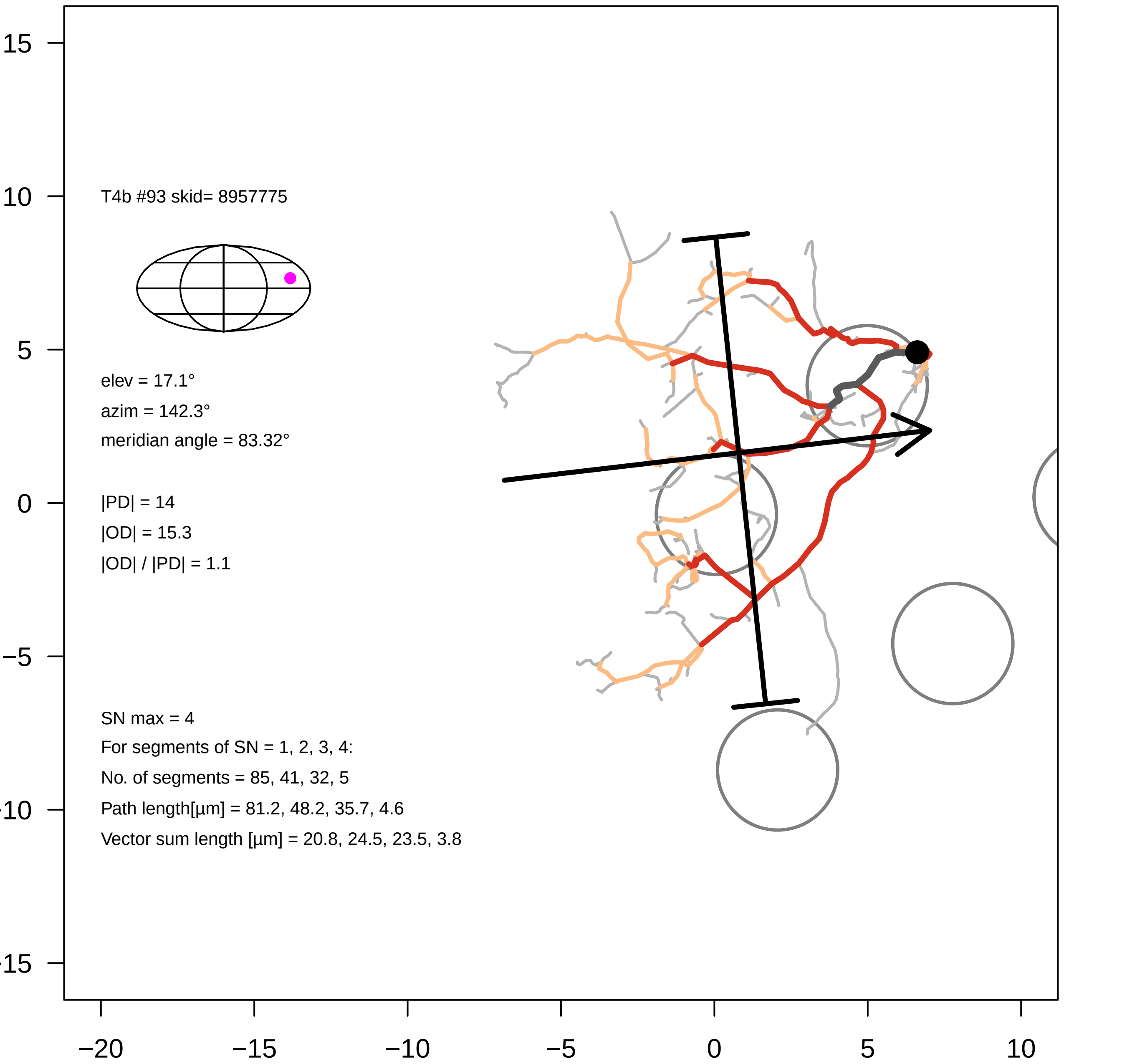
T4b #93 skid= 8957775
elev = 17.1°
azim = 142.3°
meridian angle = 83.32°
|PD| = 14
|OD| = 15.3
|OD| / |PD| = 1.1
SN max = 4
For segments of SN = 1, 2, 3, 4:
No. of segments = 85, 41, 32, 5
Path length[µm] = 81.2, 48.2, 35.7, 4.6
Vector sum length [µm] = 20.8, 24.5, 23.5, 3.8
15
10
5
0
−5
−10
−15
−20
−15
−10
−5
0
5
10

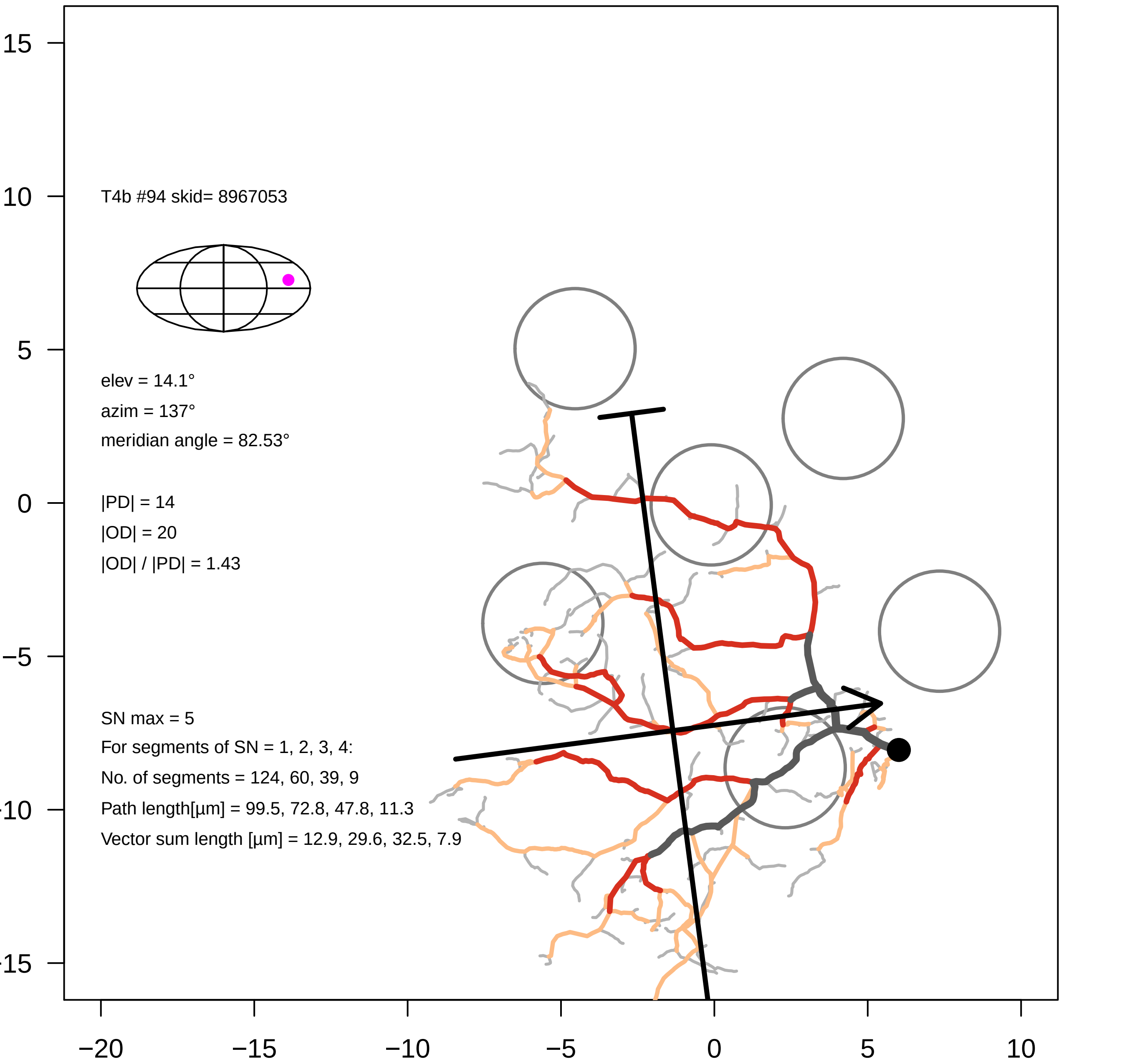

T4b #94 skid= 8967053
elev = 14.1°
azim = 137°
meridian angle = 82.53°
|PD| = 14
|OD| = 20
|OD| / |PD| = 1.43
SN max = 5
For segments of SN = 1, 2, 3, 4:
No. of segments = 124, 60, 39, 9
Path length[µm] = 99.5, 72.8, 47.8, 11.3
Vector sum length [µm] = 12.9, 29.6, 32.5, 7.9
15
10
5
0
−5
−10
−15
−20
−15
−10
−5
0
5
10

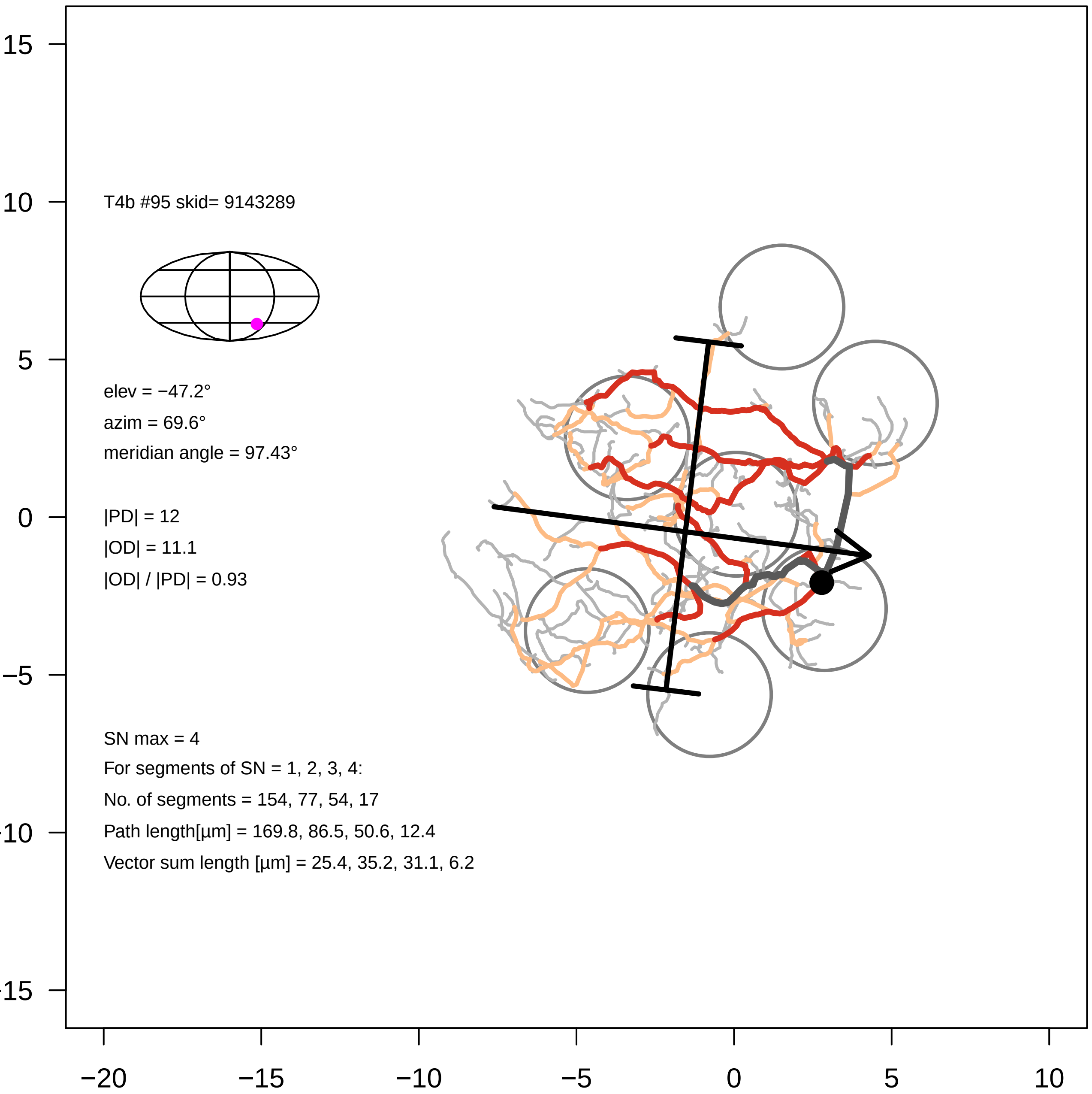

15
10
5
0
−5
−10
−15
−20
−15
−10
−5
0
5
10
T4b #95 skid= 9143289
elev = −47.2°
azim = 69.6°
meridian angle = 97.43°
|PD| = 12
|OD| = 11.1
|OD| / |PD| = 0.93
SN max = 4
For segments of SN = 1, 2, 3, 4:
No. of segments = 154, 77, 54, 17
Path length[µm] = 169.8, 86.5, 50.6, 12.4
Vector sum length [µm] = 25.4, 35.2, 31.1, 6.2

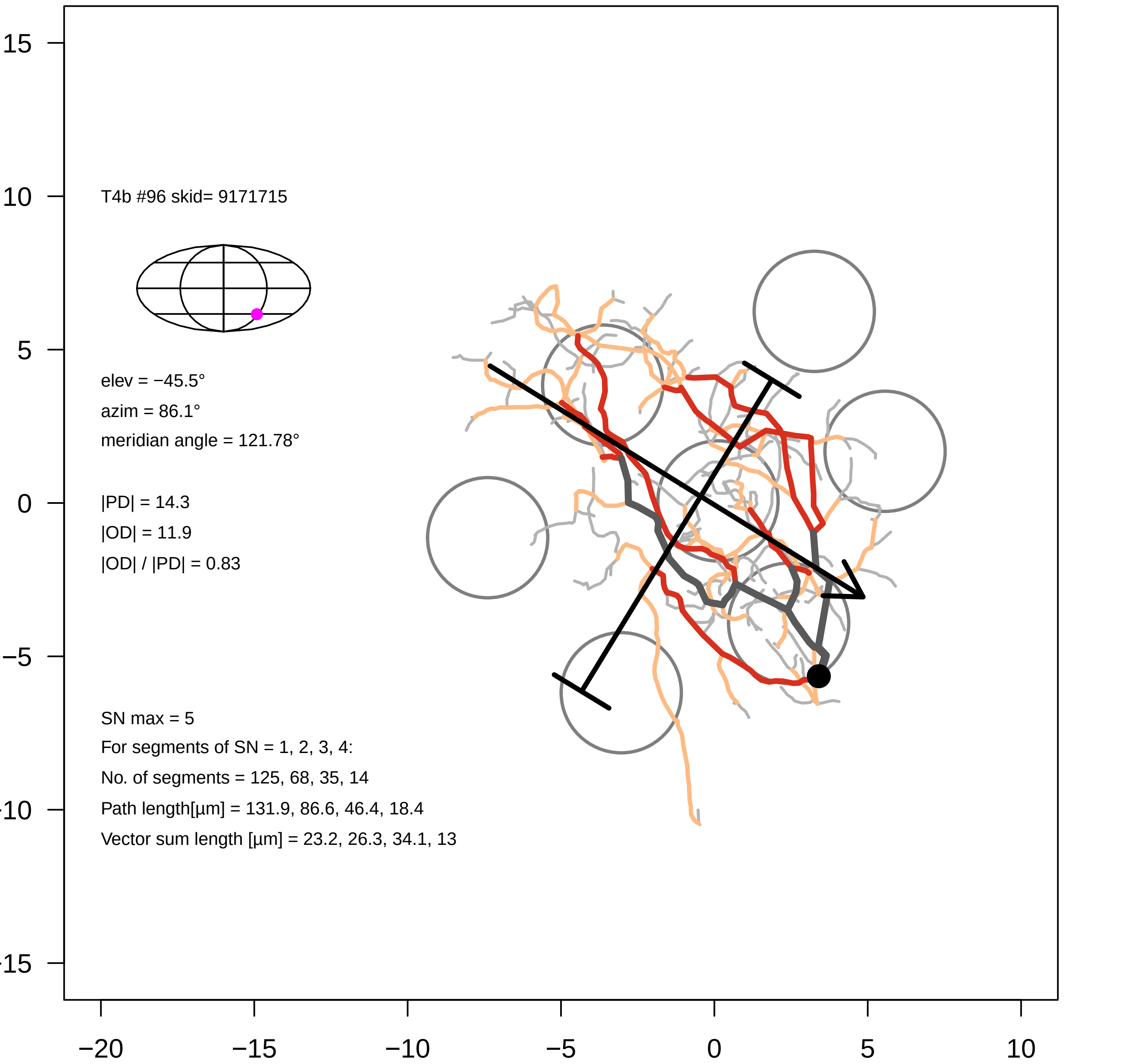

T4b #96 skid= 9171715
elev = −45.5°
azim = 86.1°
meridian angle = 121.78°
|PD| = 14.3
|OD| = 11.9
|OD| / |PD| = 0.83
SN max = 5
For segments of SN = 1, 2, 3, 4:
No. of segments = 125, 68, 35, 14
Path length[µm] = 131.9, 86.6, 46.4, 18.4
Vector sum length [µm] = 23.2, 26.3, 34.1, 13
15
10
5
0
−5
−10
−15
−20
−15
−10
−5
0
5
10

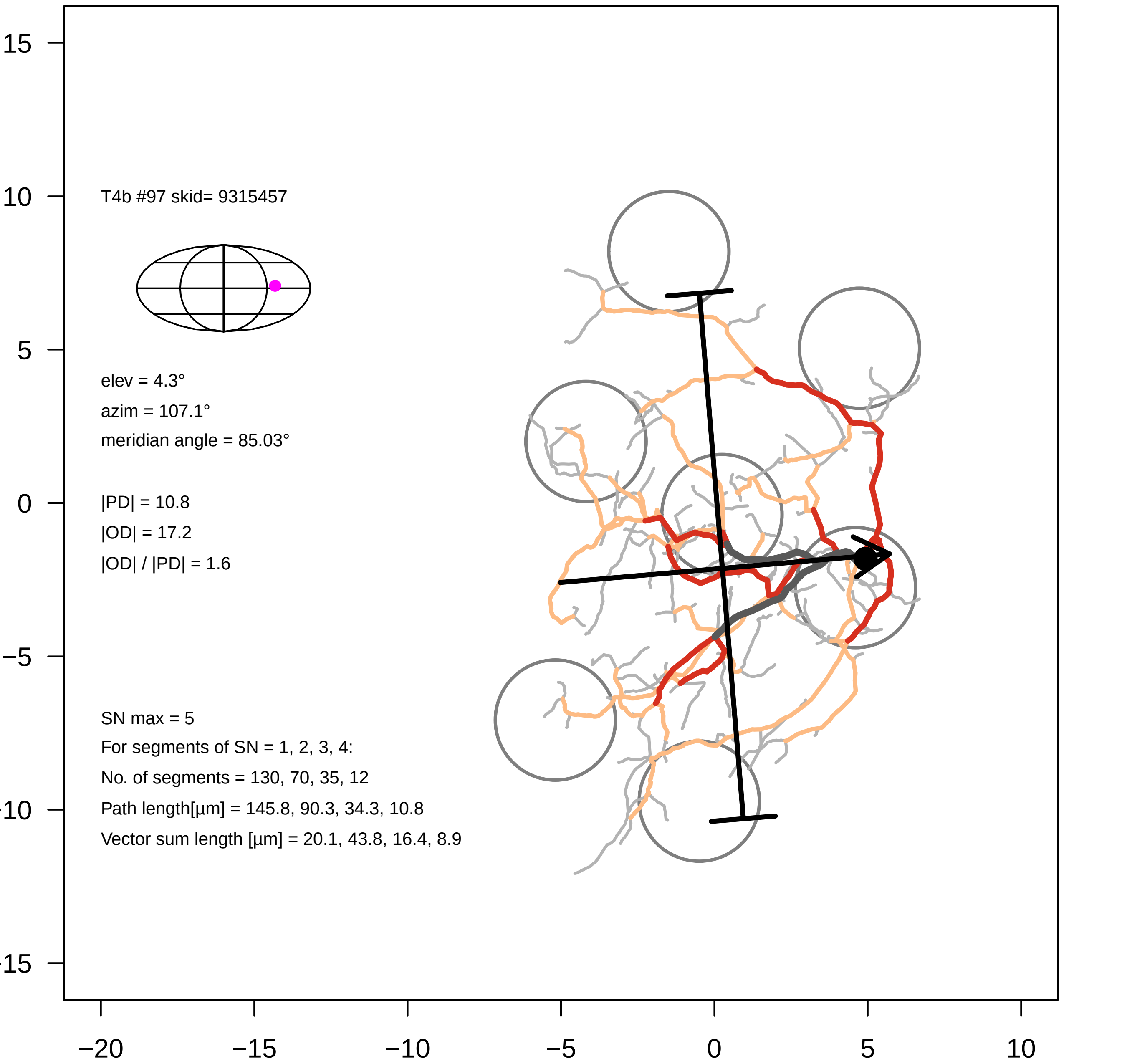

T4b #97 skid= 9315457
elev = 4.3°
azim = 107.1°
meridian angle = 85.03°
|PD| = 10.8
|OD| = 17.2
|OD| / |PD| = 1.6
SN max = 5
For segments of SN = 1, 2, 3, 4:
No. of segments = 130, 70, 35, 12
Path length[µm] = 145.8, 90.3, 34.3, 10.8
Vector sum length [µm] = 20.1, 43.8, 16.4, 8.9
15
10
5
0
−5
−10
−15
−20
−15
−10
−5
0
5
10

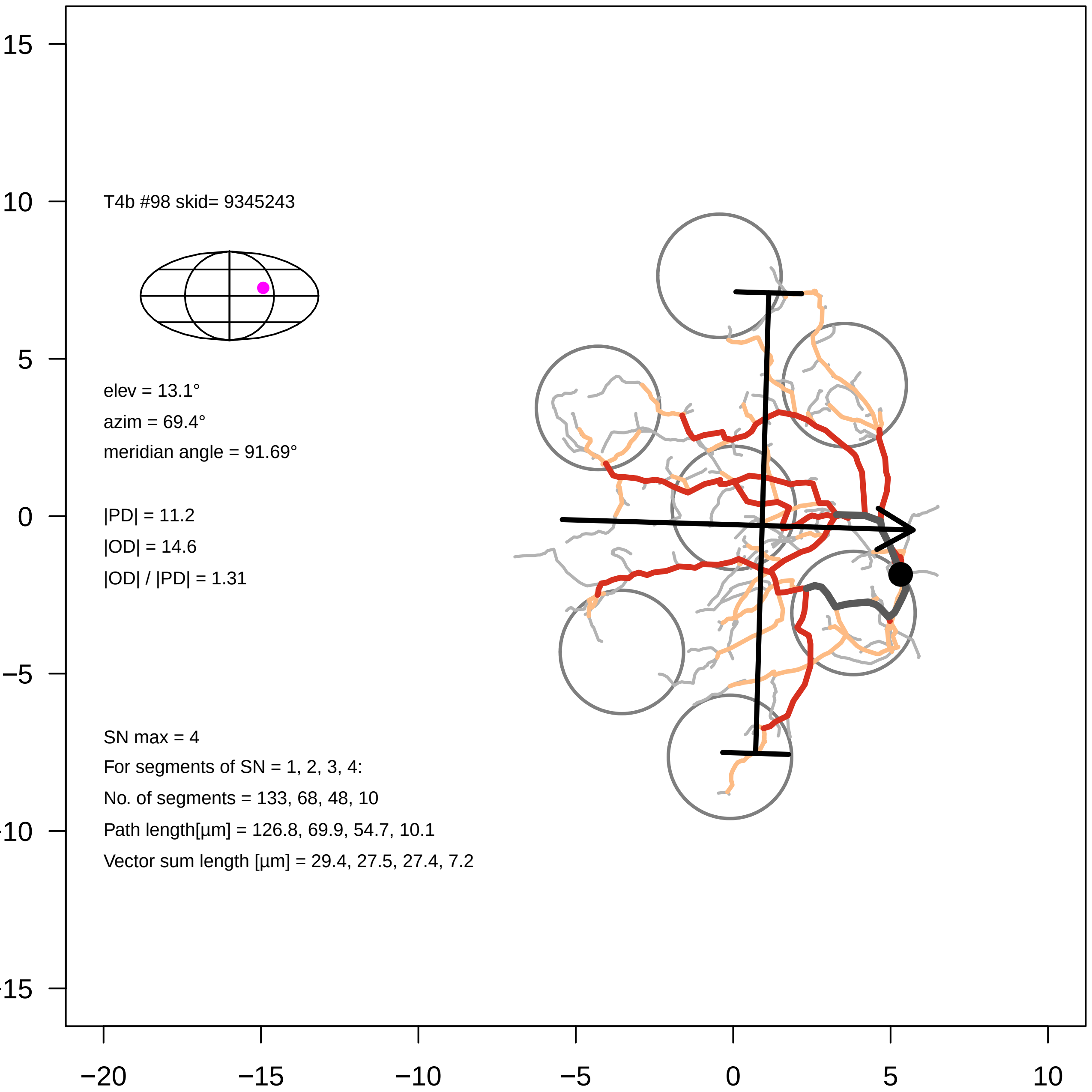

T4b #98 skid= 9345243
elev = 13.1°
azim = 69.4°
meridian angle = 91.69°
|PD| = 11.2
|OD| = 14.6
|OD| / |PD| = 1.31
SN max = 4
For segments of SN = 1, 2, 3, 4:
No. of segments = 133, 68, 48, 10
Path length[µm] = 126.8, 69.9, 54.7, 10.1
Vector sum length [µm] = 29.4, 27.5, 27.4, 7.2
15
10
5
0
−5
−10
−15
−20
−15
−10
−5
0
5
10

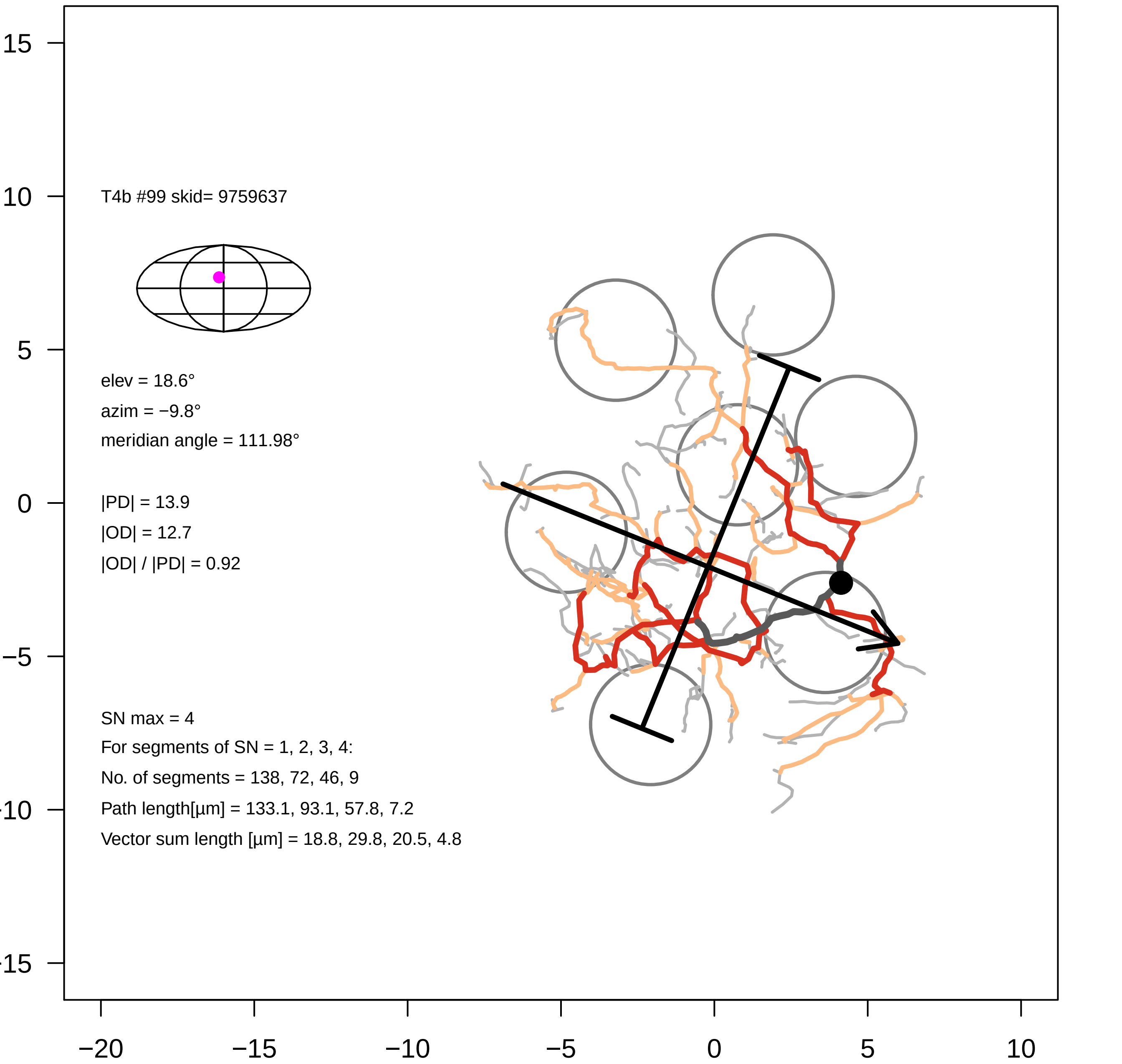
T4b #99 skid= 9759637
elev = 18.6°
azim = −9.8°
meridian angle = 111.98°
|PD| = 13.9
|OD| = 12.7
|OD| / |PD| = 0.92
SN max = 4
For segments of SN = 1, 2, 3, 4:
No. of segments = 138, 72, 46, 9
Path length[µm] = 133.1, 93.1, 57.8, 7.2
Vector sum length [µm] = 18.8, 29.8, 20.5, 4.8

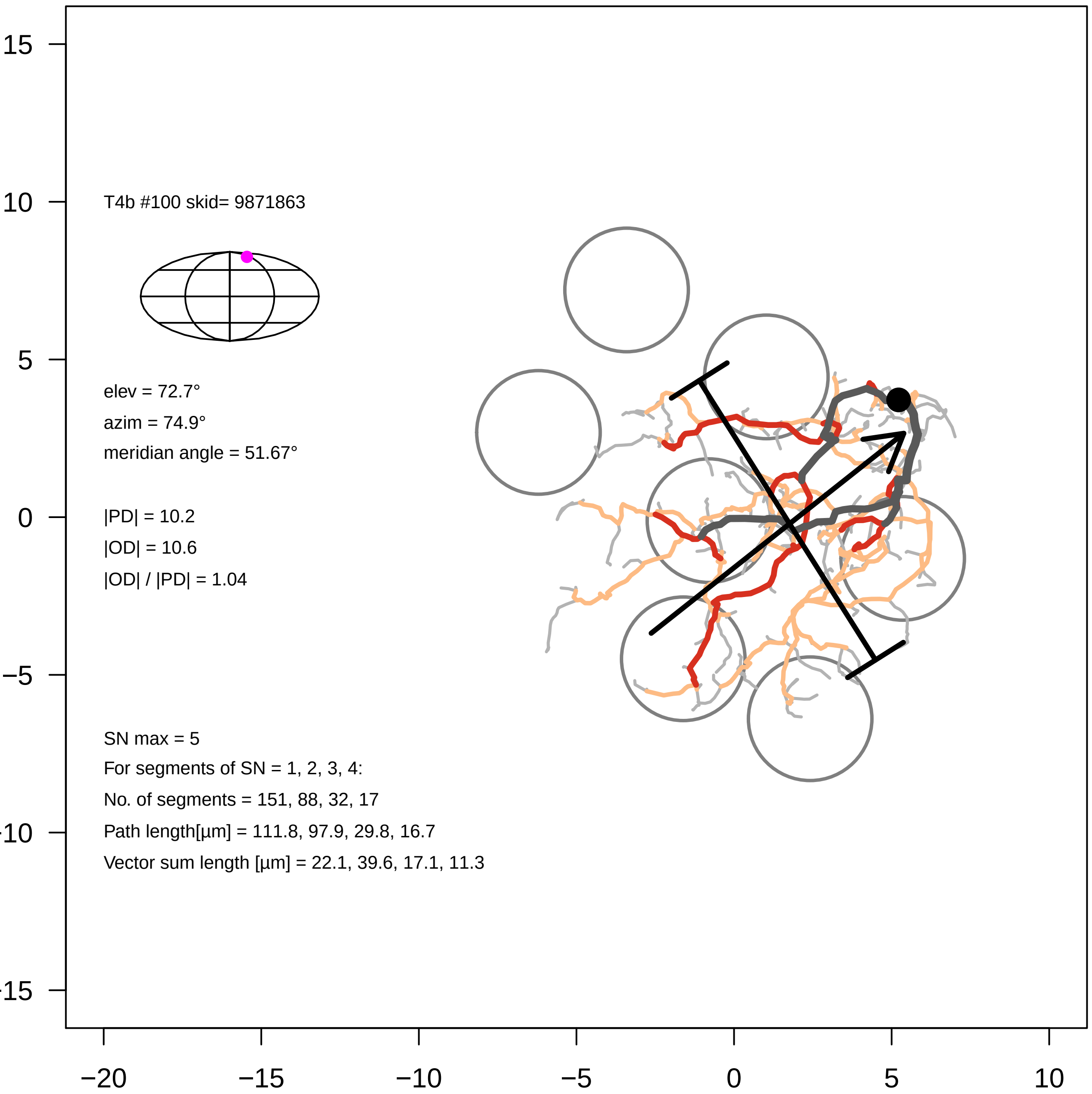

T4b #100 skid= 9871863
elev = 72.7°
azim = 74.9°
meridian angle = 51.67°
|PD| = 10.2
|OD| = 10.6
|OD| / |PD| = 1.04
SN max = 5
For segments of SN = 1, 2, 3, 4:
No. of segments = 151, 88, 32, 17
Path length[µm] = 111.8, 97.9, 29.8, 16.7
Vector sum length [µm] = 22.1, 39.6, 17.1, 11.3
15
10
5
0
−5
−10
−15
−20
−15
−10
−5
0
5
10

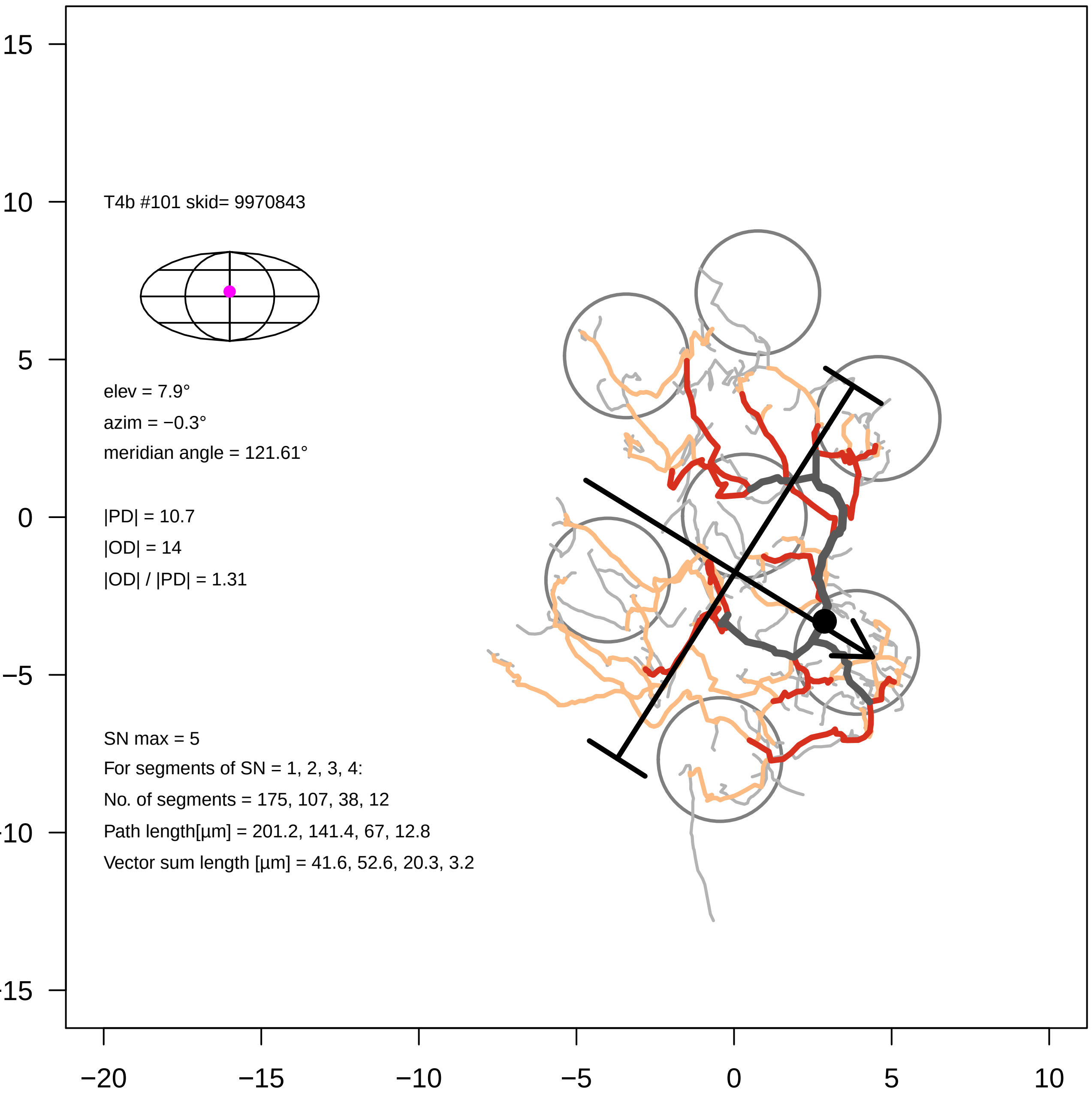

T4b #101 skid= 9970843
elev = 7.9°
azim = −0.3°
meridian angle = 121.61°
|PD| = 10.7
|OD| = 14
|OD| / |PD| = 1.31
SN max = 5
For segments of SN = 1, 2, 3, 4:
No. of segments = 175, 107, 38, 12
Path length[µm] = 201.2, 141.4, 67, 12.8
Vector sum length [µm] = 41.6, 52.6, 20.3, 3.2
15
10
5
0
−5
−20
−15
−10
−5
0
5
10

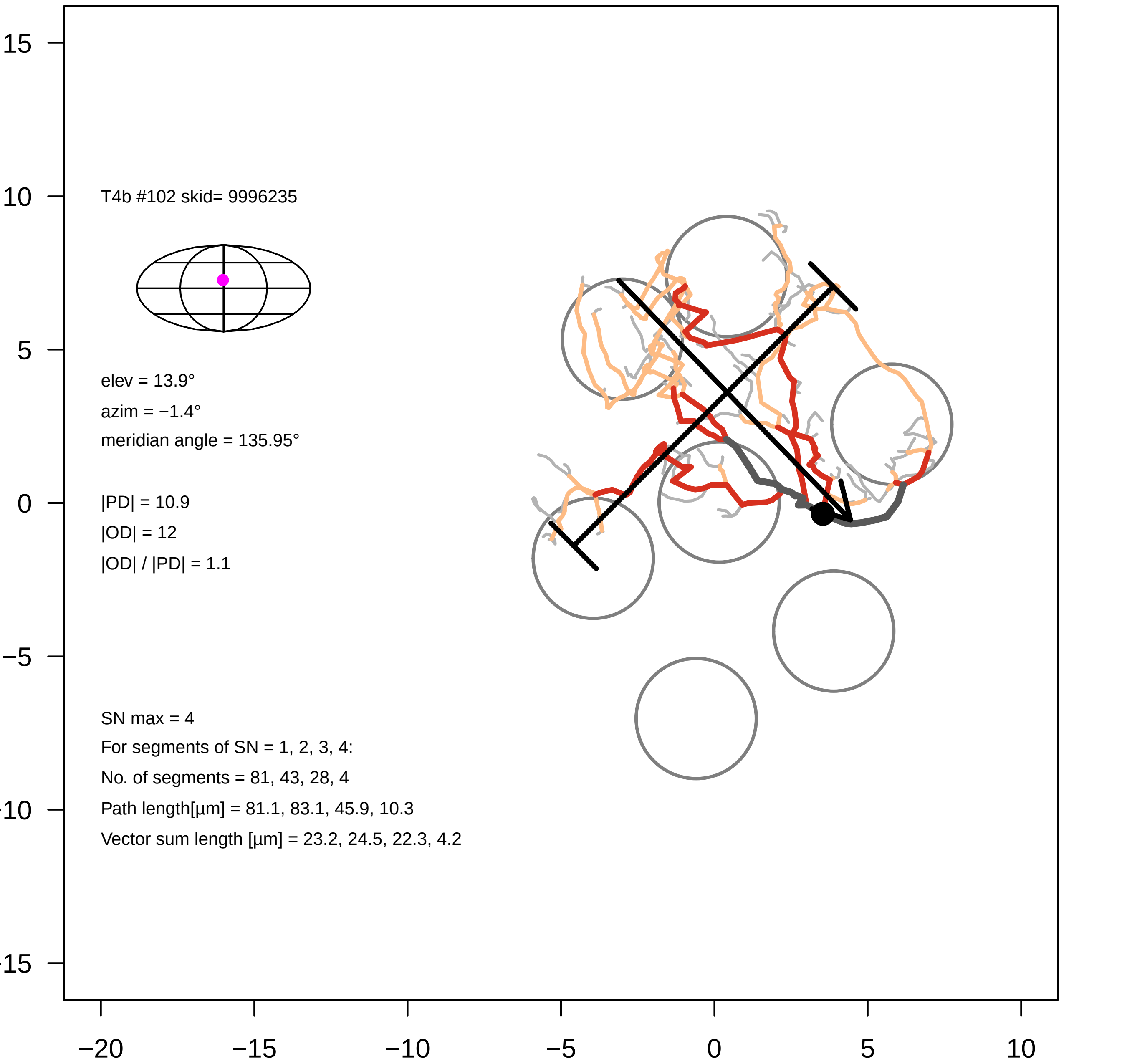
T4b #102 skid= 9996235
elev = 13.9°
azim = −1.4°
meridian angle = 135.95°
|PD| = 10.9
|OD| = 12
|OD| / |PD| = 1.1
SN max = 4
For segments of SN = 1, 2, 3, 4:
No. of segments = 81, 43, 28, 4
Path length[µm] = 81.1, 83.1, 45.9, 10.3
Vector sum length [µm] = 23.2, 24.5, 22.3, 4.2
15
10
5
0
−5
−20
−15
−10
−5
0
5
10

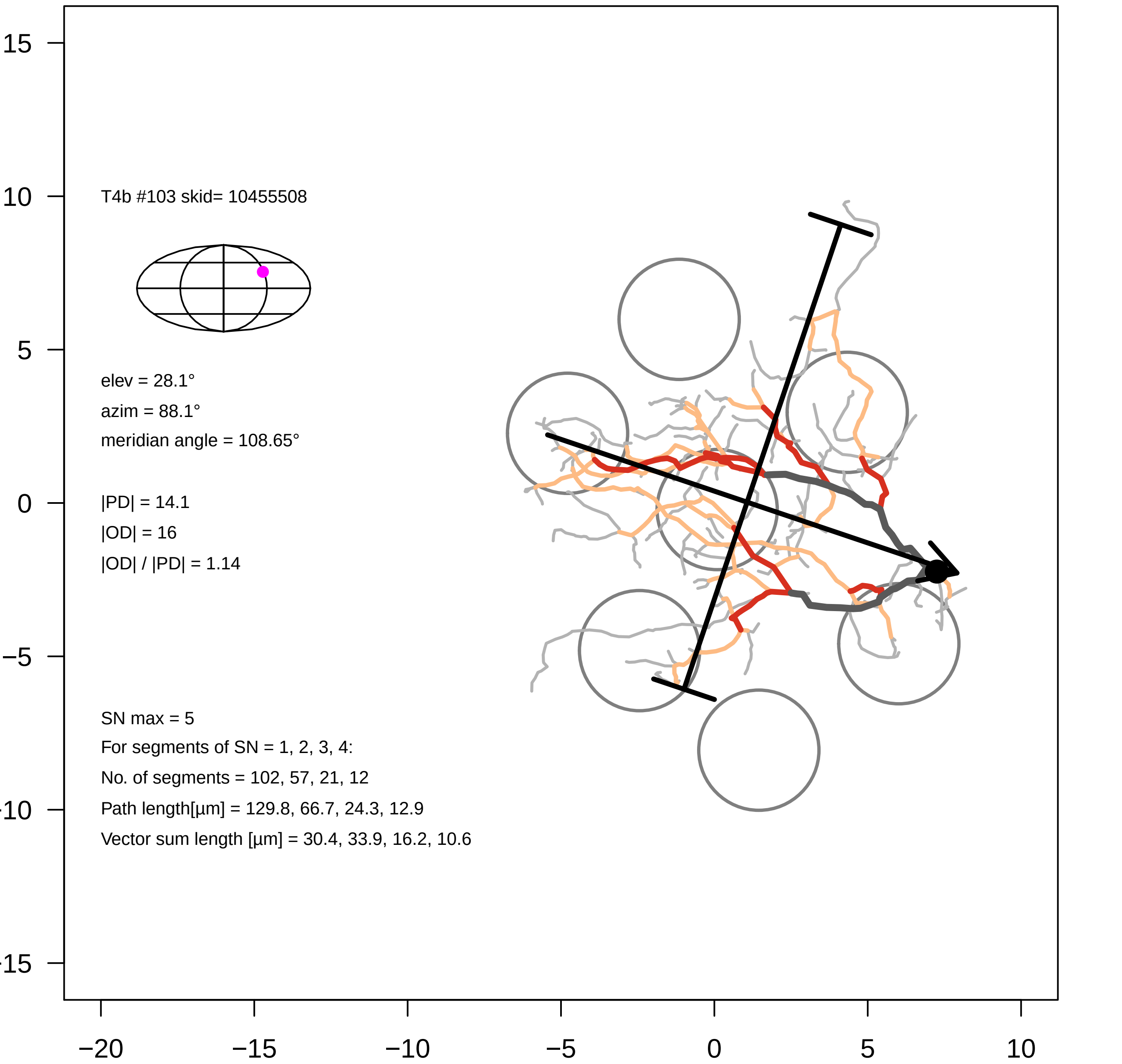

T4b #103 skid= 10455508
elev = 28.1°
azim = 88.1°
meridian angle = 108.65°
|PD| = 14.1
|OD| = 16
|OD| / |PD| = 1.14
SN max = 5
For segments of SN = 1, 2, 3, 4:
No. of segments = 102, 57, 21, 12
Path length[µm] = 129.8, 66.7, 24.3, 12.9
Vector sum length [µm] = 30.4, 33.9, 16.2, 10.6
15
10
5
0
−5
−10
−15
−20
−15
−10
−5
0
5
10

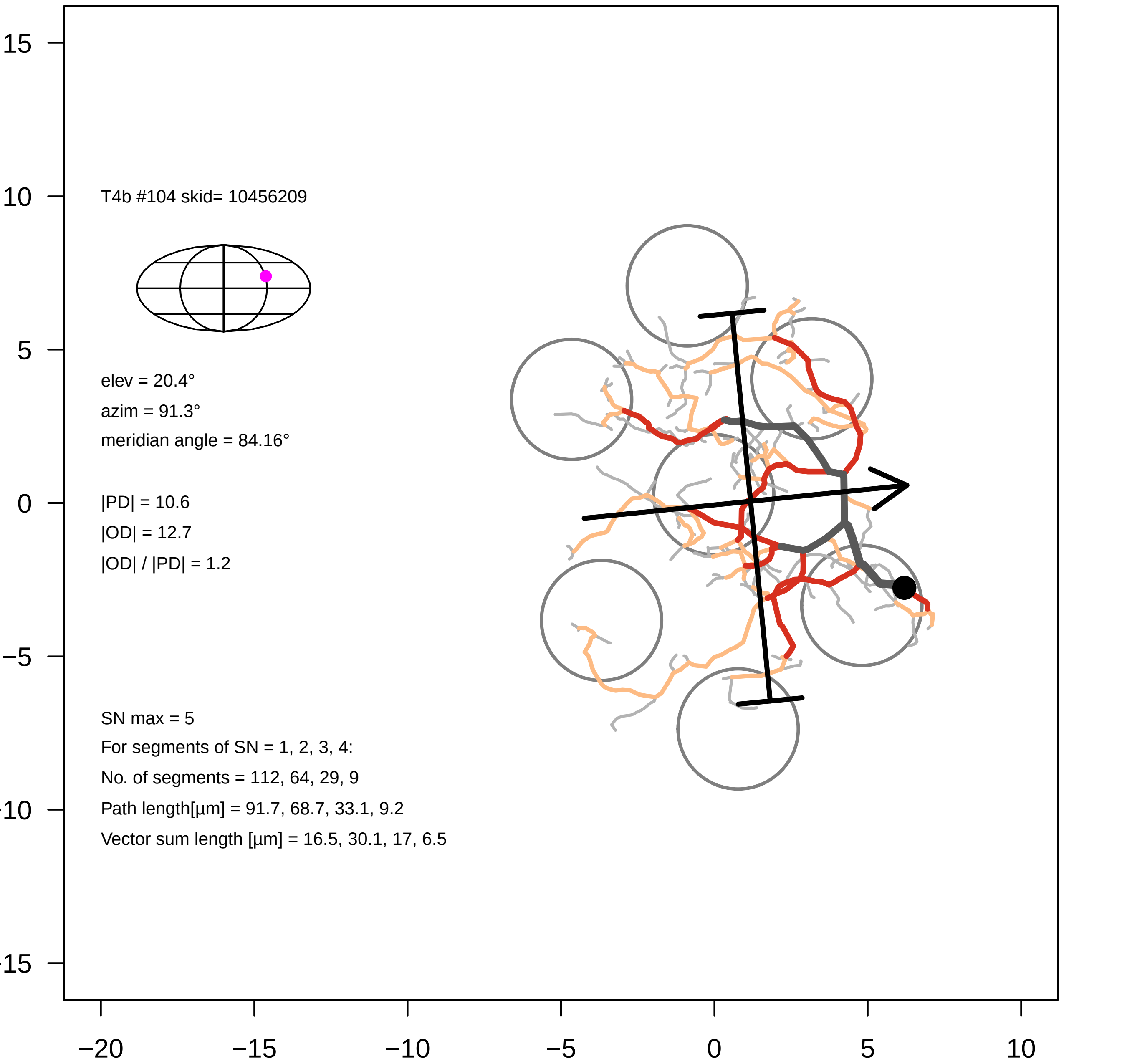
T4b #104 skid= 10456209
elev = 20.4°
azim = 91.3°
meridian angle = 84.16°
|PD| = 10.6
|OD| = 12.7
|OD| / |PD| = 1.2
SN max = 5
For segments of SN = 1, 2, 3, 4:
No. of segments = 112, 64, 29, 9
Path length[µm] = 91.7, 68.7, 33.1, 9.2
Vector sum length [µm] = 16.5, 30.1, 17, 6.5
15
10
5
0
−5
−10
−15
−20
−15
−10
−5
0
5
10

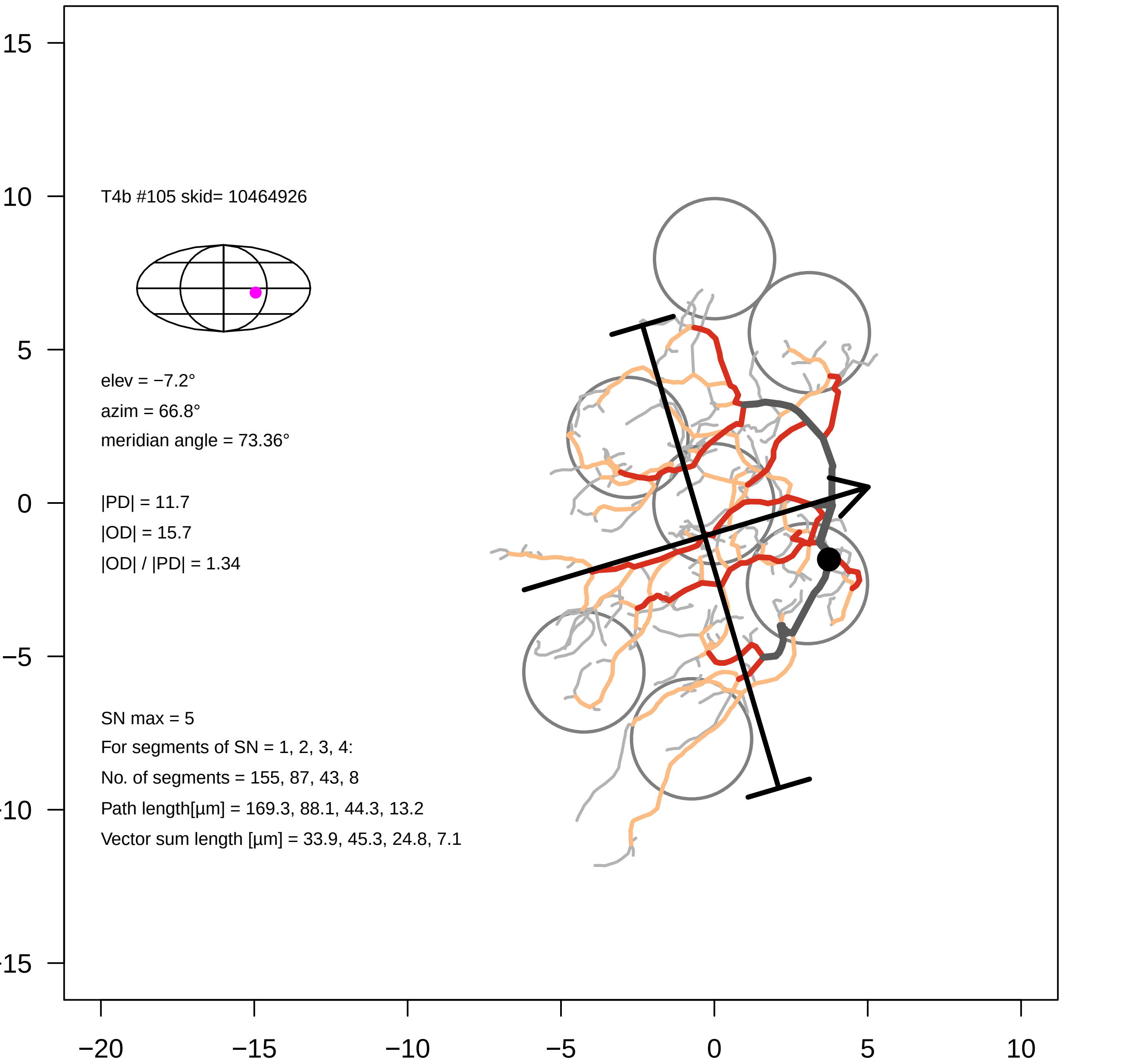

T4b #105 skid= 10464926
elev = −7.2°
azim = 66.8°
meridian angle = 73.36°
|PD| = 11.7
|OD| = 15.7
|OD| / |PD| = 1.34
SN max = 5
For segments of SN = 1, 2, 3, 4:
No. of segments = 155, 87, 43, 8
Path length[µm] = 169.3, 88.1, 44.3, 13.2
Vector sum length [µm] = 33.9, 45.3, 24.8, 7.1
15
10
5
0
−5
−10
−15
−20
−15
−10
−5
0
5
10

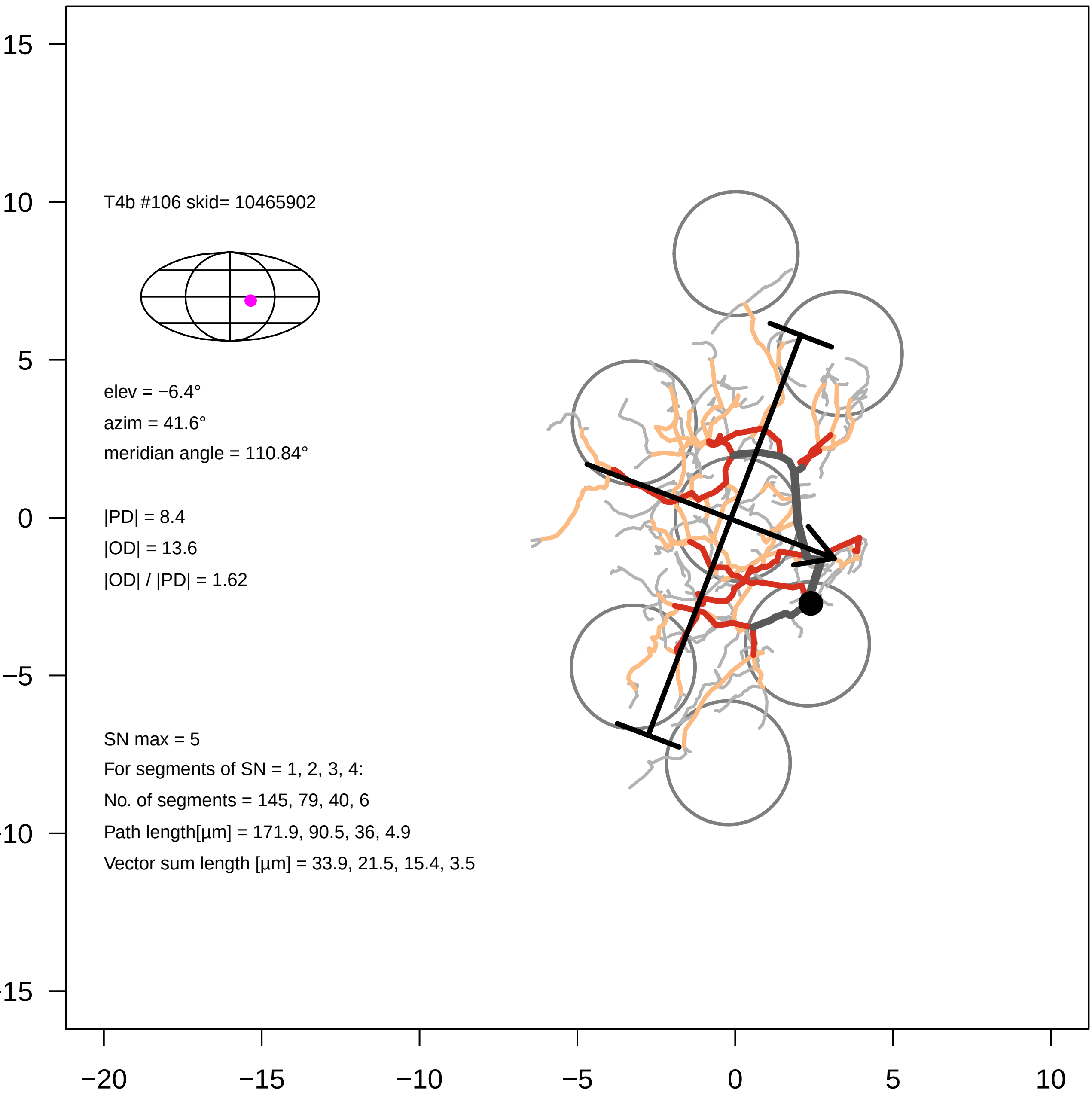
T4b #106 skid= 10465902
elev = −6.4°
azim = 41.6°
meridian angle = 110.84°
|PD| = 8.4
|OD| = 13.6
|OD| / |PD| = 1.62
SN max = 5
For segments of SN = 1, 2, 3, 4:
No. of segments = 145, 79, 40, 6
Path length[µm] = 171.9, 90.5, 36, 4.9
Vector sum length [µm] = 33.9, 21.5, 15.4, 3.5
15
10
5
0
−5
−10
−15
−20
−15
−10
−5
0
5
10

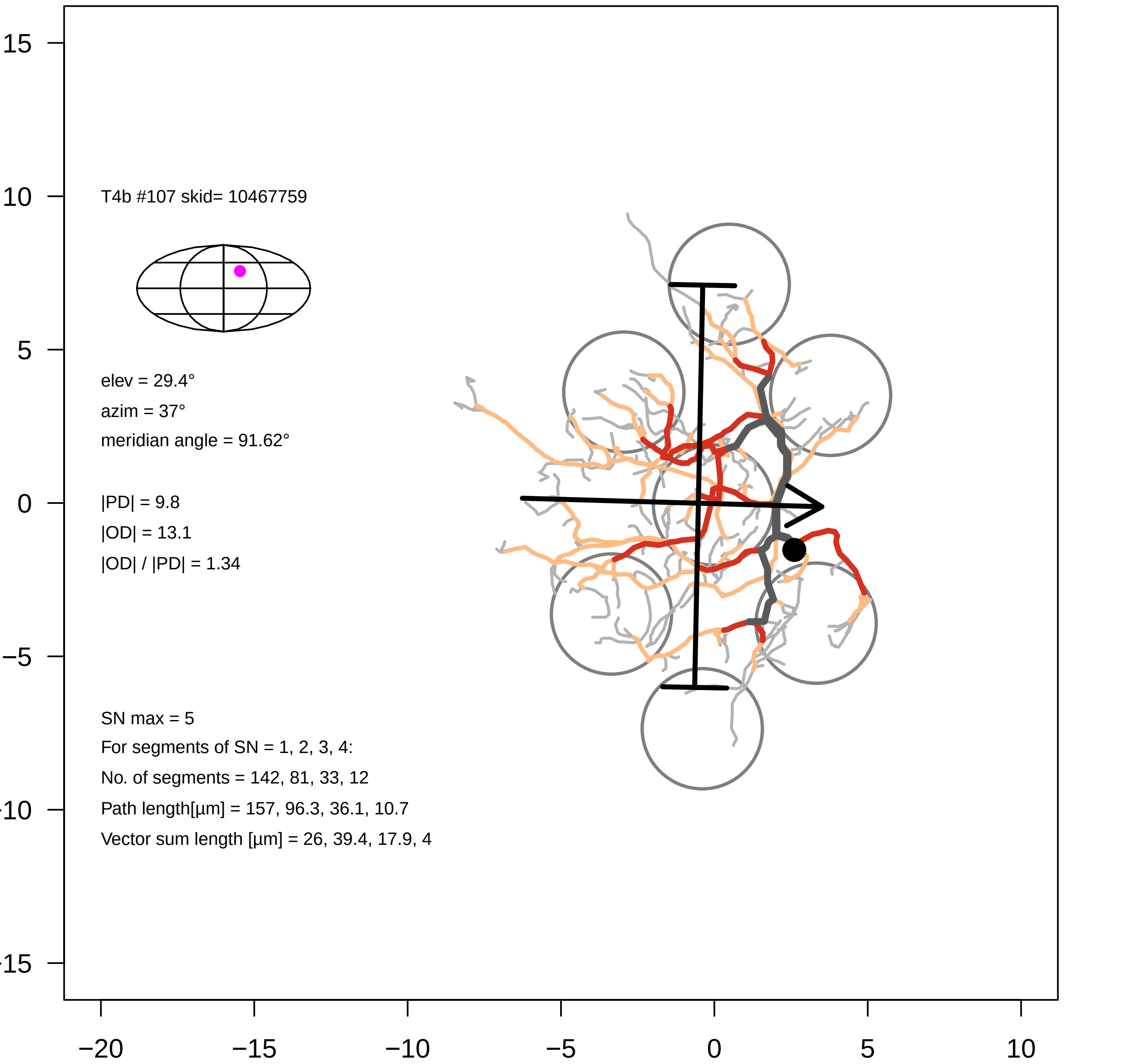

T4b #107 skid= 10467759
elev = 29.4°
azim = 37°
meridian angle = 91.62°
|PD| = 9.8
|OD| = 13.1
|OD| / |PD| = 1.34
SN max = 5
For segments of SN = 1, 2, 3, 4:
No. of segments = 142, 81, 33, 12
Path length[µm] = 157, 96.3, 36.1, 10.7
Vector sum length [µm] = 26, 39.4, 17.9, 4
15
10
5
0
−5
-10
-15
−20
−15
−10
−5
0
5
10

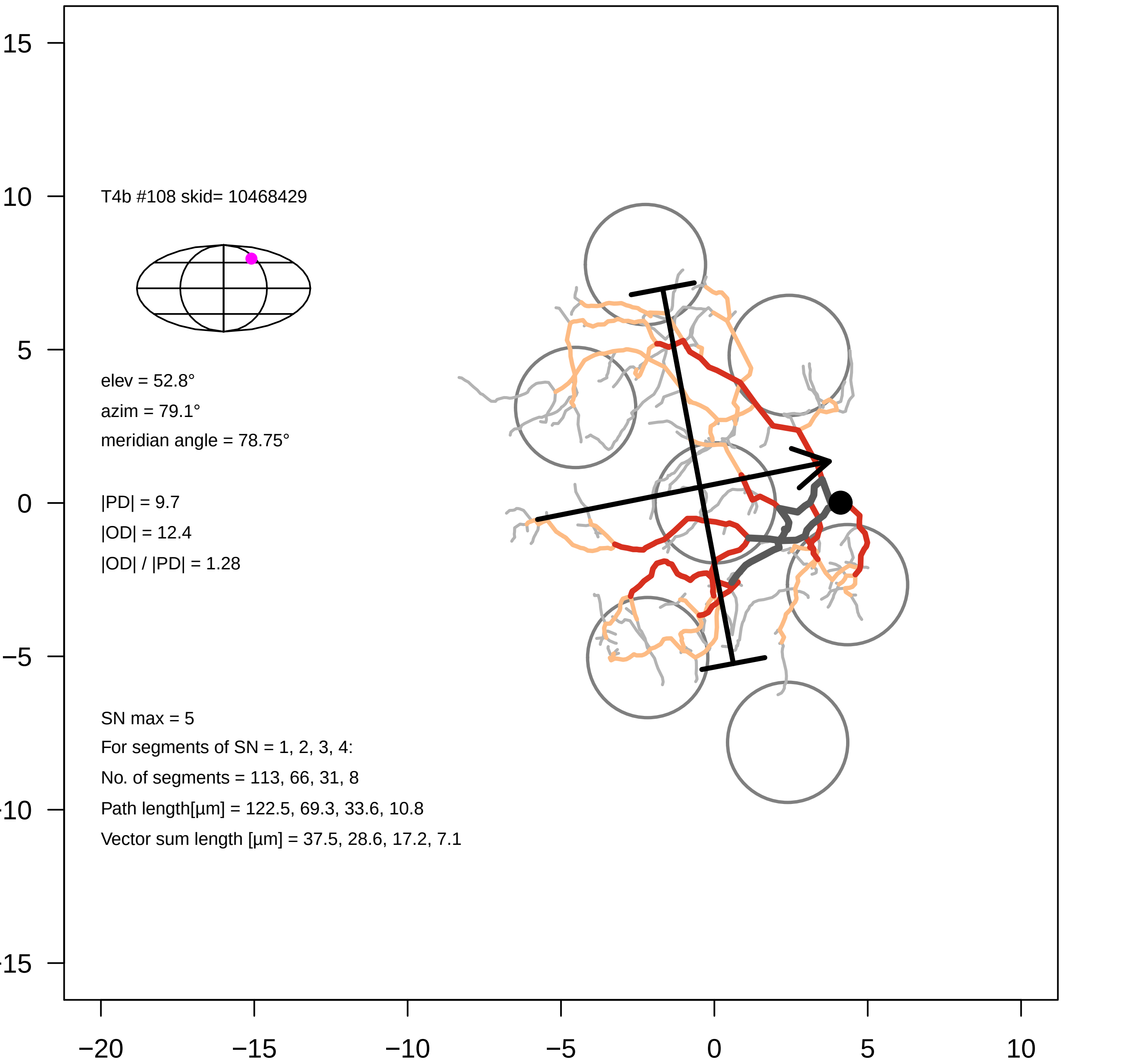

T4b #108 skid= 10468429
elev = 52.8°
azim = 79.1°
meridian angle = 78.75°
|PD| = 9.7
|OD| = 12.4
|OD| / |PD| = 1.28
SN max = 5
For segments of SN = 1, 2, 3, 4:
No. of segments = 113, 66, 31, 8
Path length[µm] = 122.5, 69.3, 33.6, 10.8
Vector sum length [µm] = 37.5, 28.6, 17.2, 7.1
15
10
5
0
−5
−10
−15
−20
−15
−10
−5
0
5
10

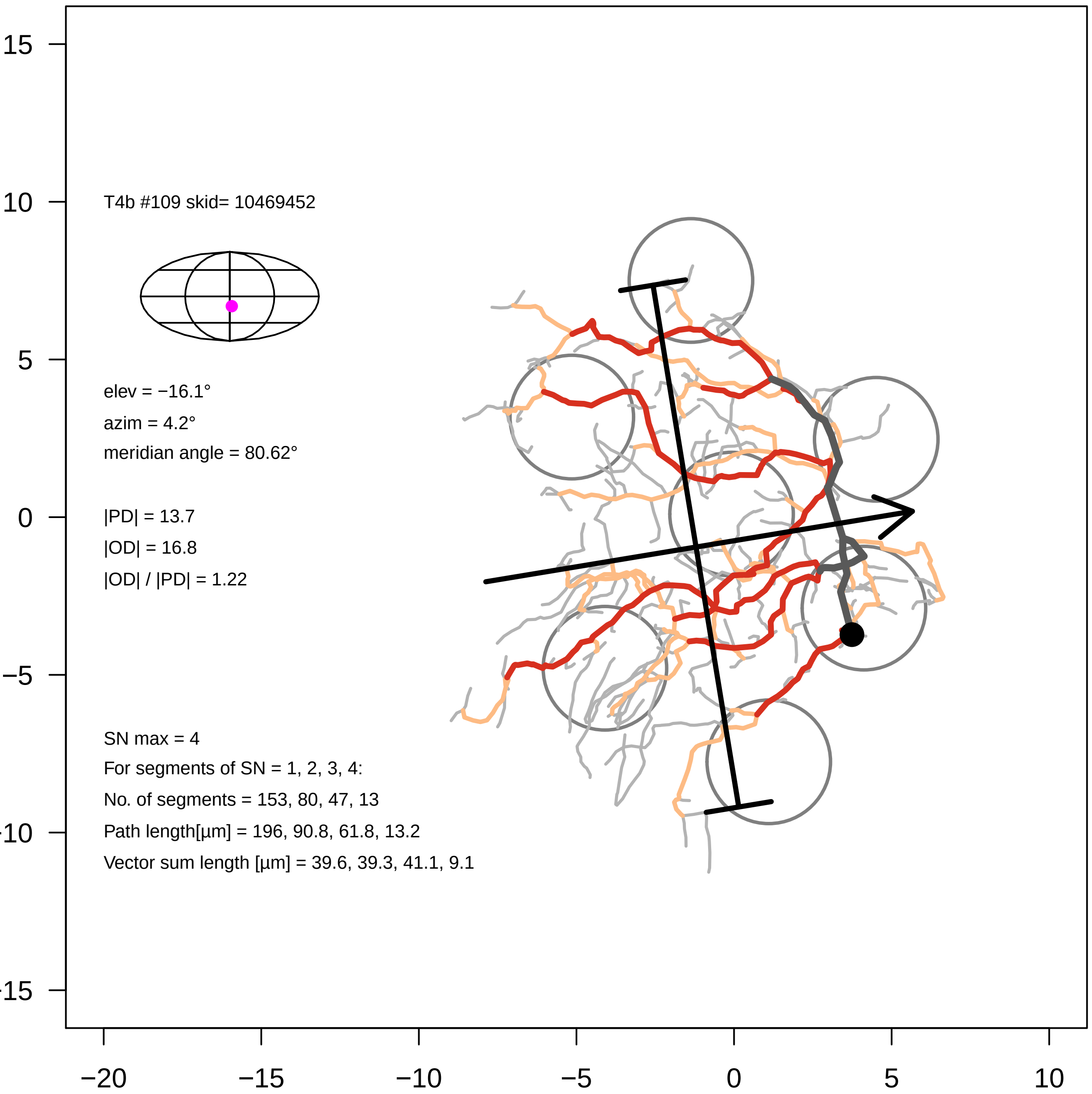
T4b #109 skid= 10469452
elev = −16.1°
azim = 4.2°
meridian angle = 80.62°
|PD| = 13.7
|OD| = 16.8
|OD| / |PD| = 1.22
SN max = 4
For segments of SN = 1, 2, 3, 4:
No. of segments = 153, 80, 47, 13
Path length[µm] = 196, 90.8, 61.8, 13.2
Vector sum length [µm] = 39.6, 39.3, 41.1, 9.1
15
10
5
0
−5
−10
−15
−20
−15
−10
−5
0
5
10

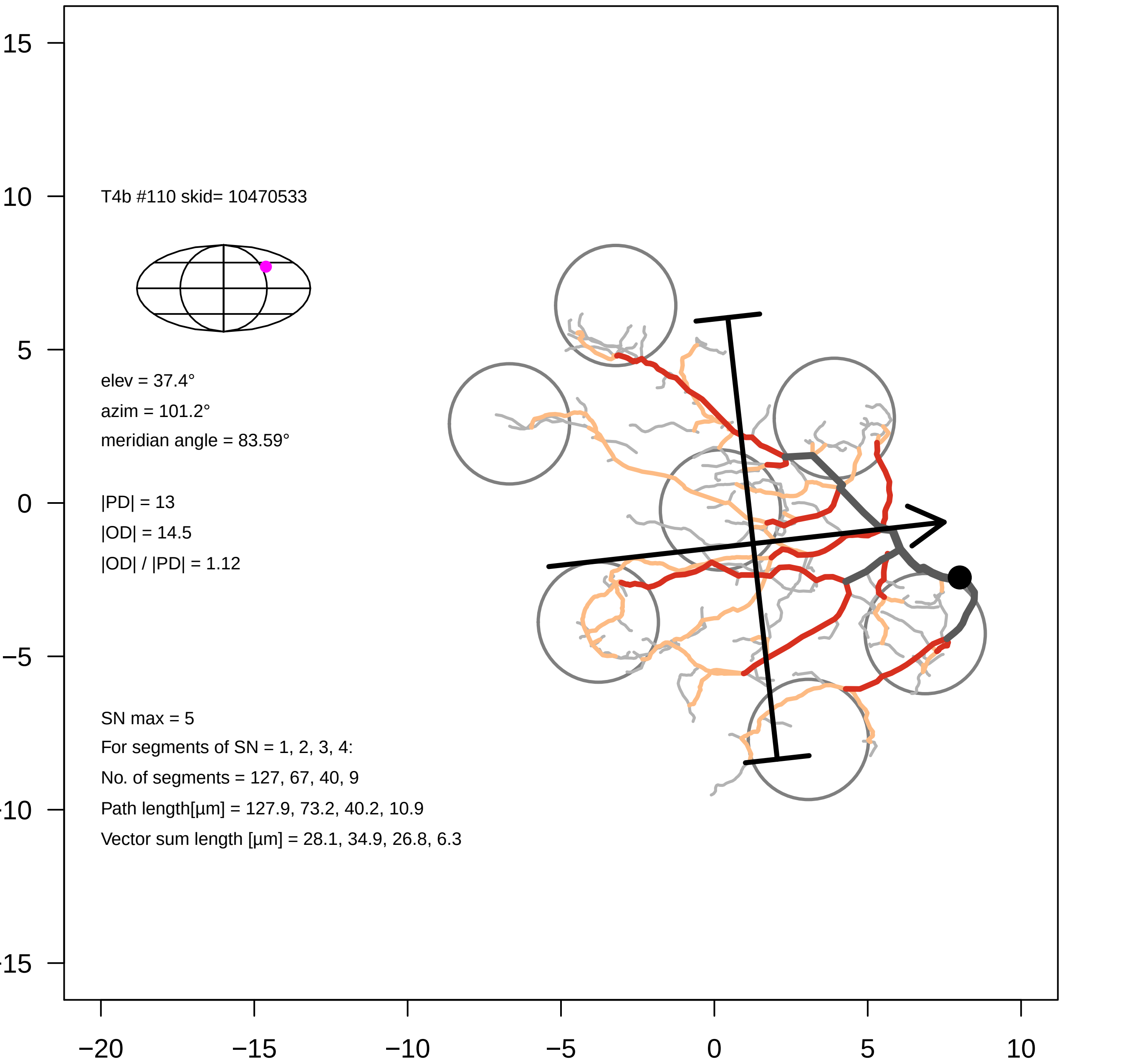

T4b #110 skid= 10470533
elev = 37.4°
azim = 101.2°
meridian angle = 83.59°
|PD| = 13
|OD| = 14.5
|OD| / |PD| = 1.12
SN max = 5
For segments of SN = 1, 2, 3, 4:
No. of segments = 127, 67, 40, 9
Path length[µm] = 127.9, 73.2, 40.2, 10.9
Vector sum length [µm] = 28.1, 34.9, 26.8, 6.3
15
10
5
0
−5
−10
−15
−20
−15
−10
−5
0
5
10

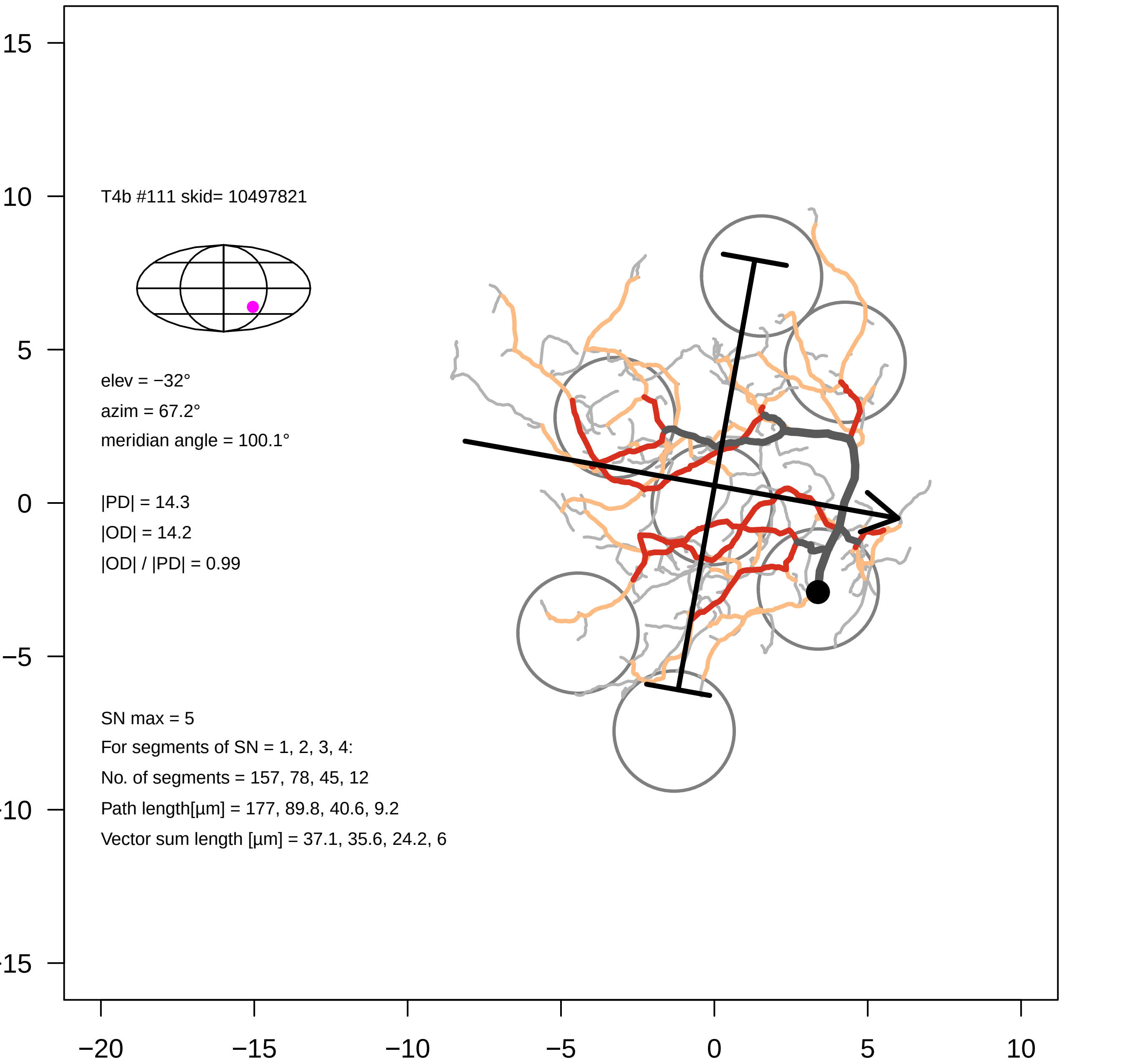

T4b #111 skid= 10497821
elev = −32°
azim = 67.2°
meridian angle = 100.1°
|PD| = 14.3
|OD| = 14.2
|OD| / |PD| = 0.99
SN max = 5
For segments of SN = 1, 2, 3, 4:
No. of segments = 157, 78, 45, 12
Path length[µm] = 177, 89.8, 40.6, 9.2
Vector sum length [µm] = 37.1, 35.6, 24.2, 6
15
10
5
0
−5
−10
−15
−20
−15
−10
−5
0
5
10

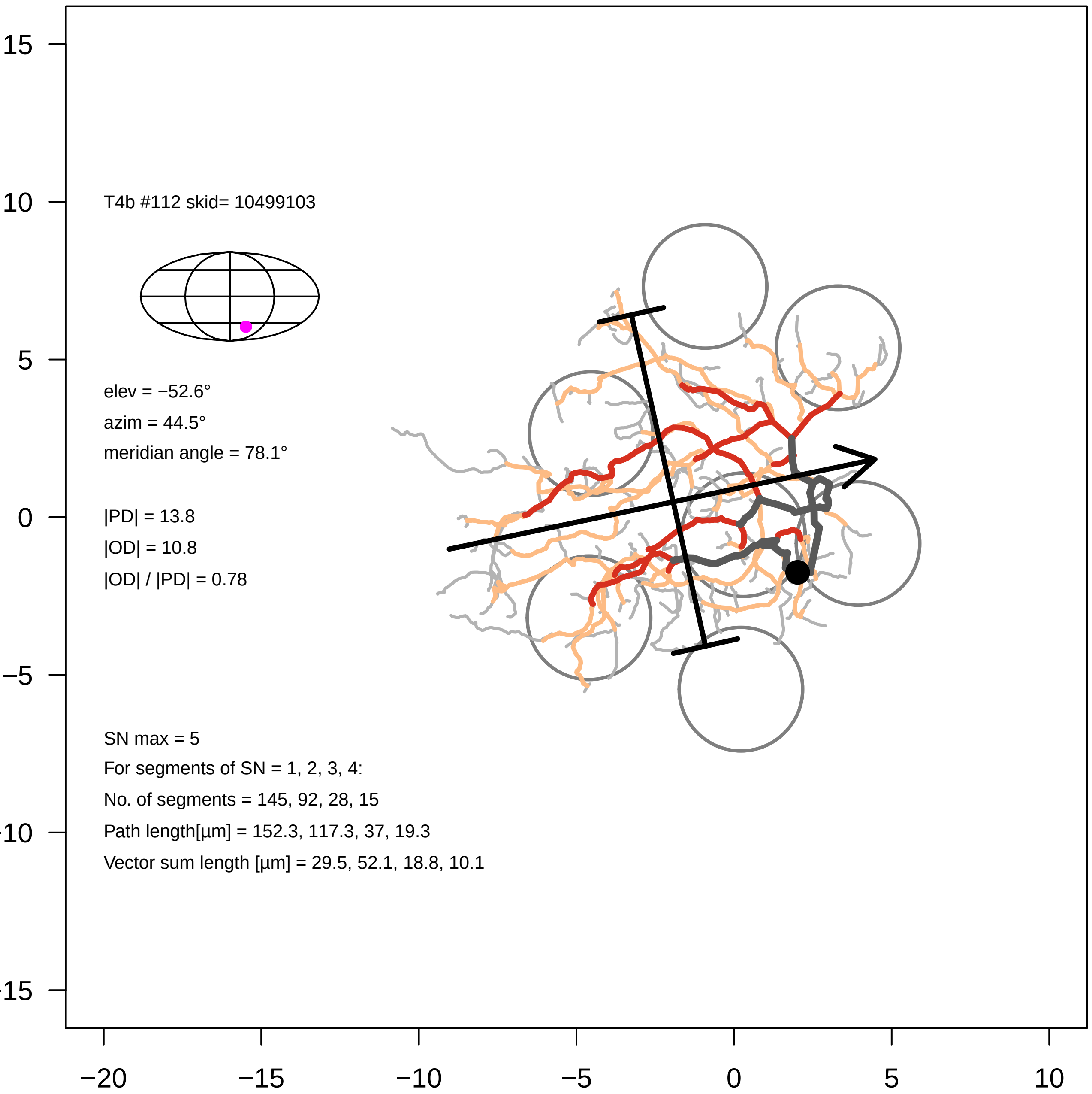
T4b #112 skid= 10499103
elev = −52.6°
azim = 44.5°
meridian angle = 78.1°
|PD| = 13.8
|OD| = 10.8
|OD| / |PD| = 0.78
SN max = 5
For segments of SN = 1, 2, 3, 4:
No. of segments = 145, 92, 28, 15
Path length[µm] = 152.3, 117.3, 37, 19.3
Vector sum length [µm] = 29.5, 52.1, 18.8, 10.1
15
10
5
0
−5
−20
−15
−10
−5
0
5
10

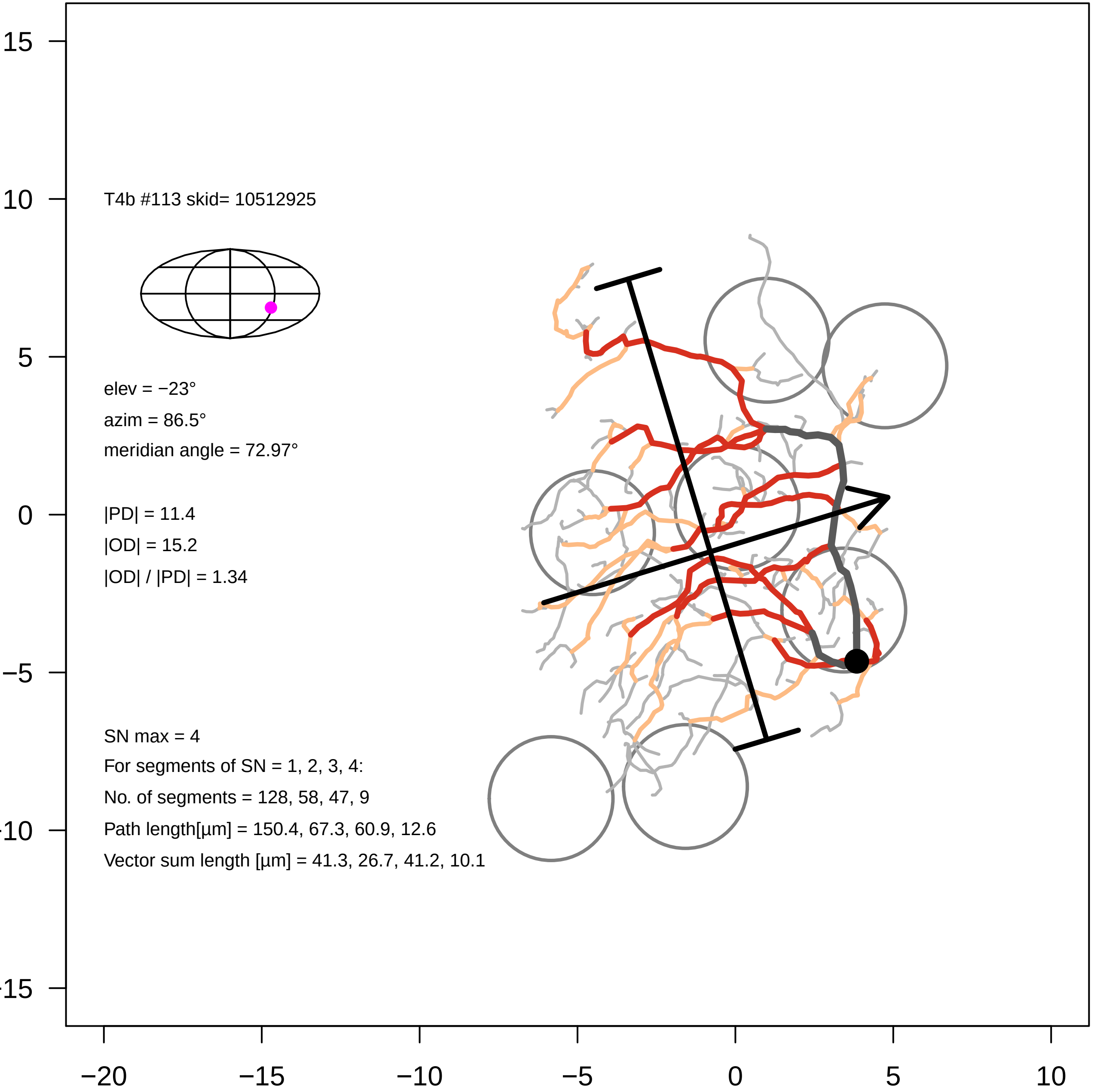
T4b #113 skid= 10512925
elev = −23°
azim = 86.5°
meridian angle = 72.97°
|PD| = 11.4
|OD| = 15.2
|OD| / |PD| = 1.34
SN max = 4
For segments of SN = 1, 2, 3, 4:
No. of segments = 128, 58, 47, 9
Path length[µm] = 150.4, 67.3, 60.9, 12.6
Vector sum length [µm] = 41.3, 26.7, 41.2, 10.1
15
10
5
0
−5
−10
−15
−20
−15
−10
−5
0
5
10

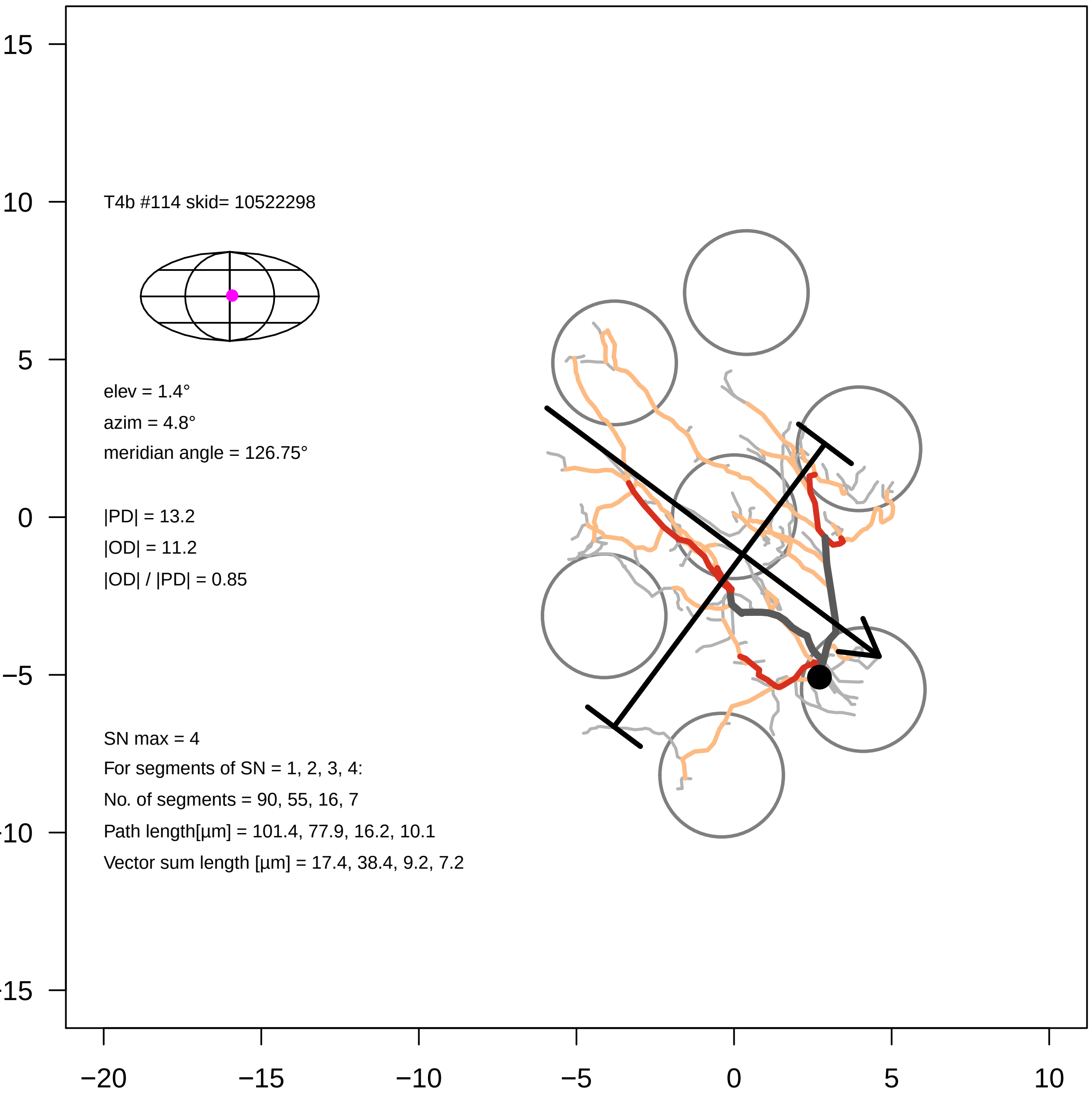
T4b #114 skid= 10522298
elev = 1.4°
azim = 4.8°
meridian angle = 126.75°
|PD| = 13.2
|OD| = 11.2
|OD| / |PD| = 0.85
SN max = 4
For segments of SN = 1, 2, 3, 4:
No. of segments = 90, 55, 16, 7
Path length[µm] = 101.4, 77.9, 16.2, 10.1
Vector sum length [µm] = 17.4, 38.4, 9.2, 7.2
15
10
5
0
−5
−10
−15
−20
−15
−10
−5
0
5
10

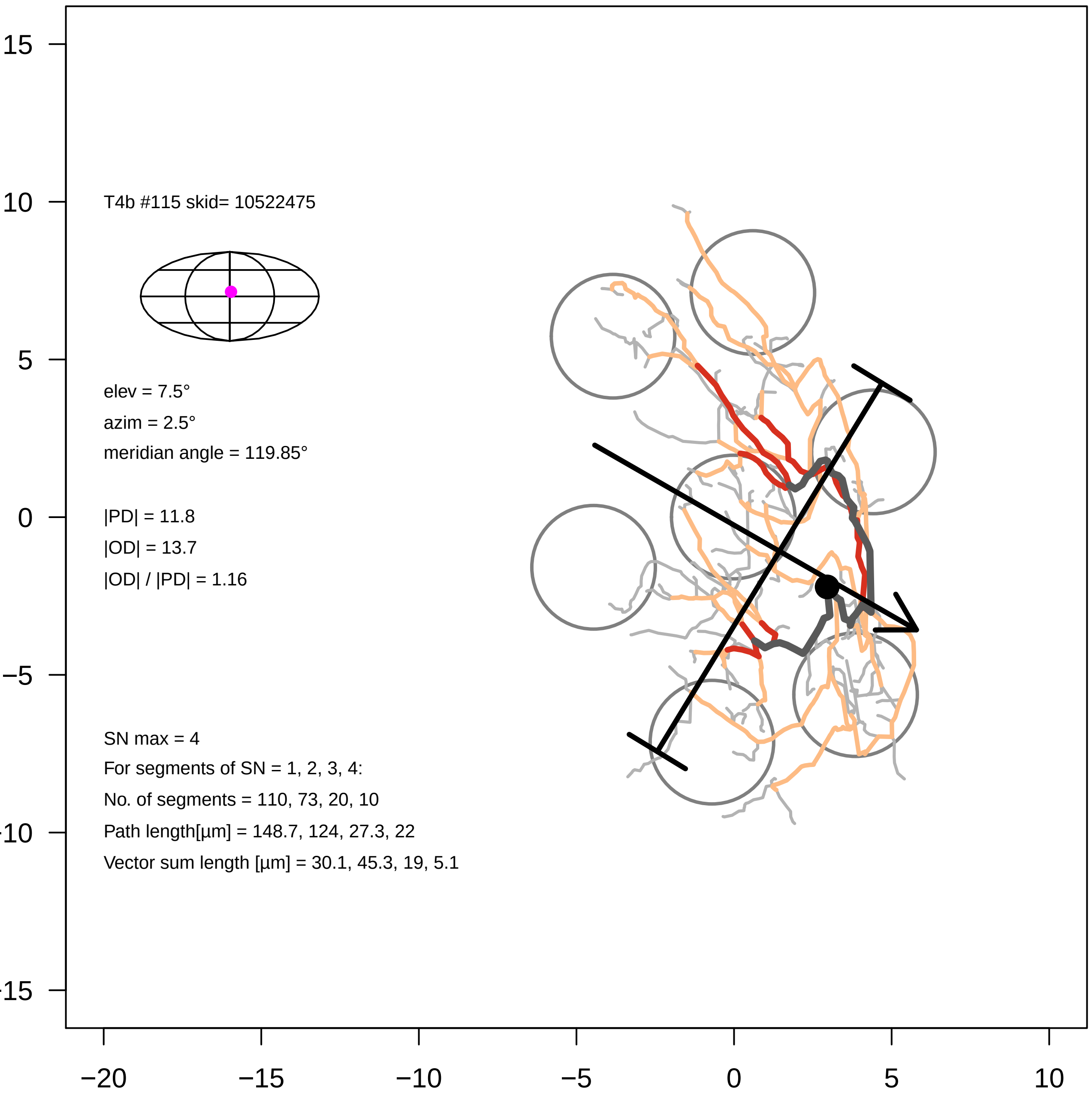

T4b #115 skid= 10522475
elev = 7.5°
azim = 2.5°
meridian angle = 119.85°
|PD| = 11.8
|OD| = 13.7
|OD| / |PD| = 1.16
SN max = 4
For segments of SN = 1, 2, 3, 4:
No. of segments = 110, 73, 20, 10
Path length[µm] = 148.7, 124, 27.3, 22
Vector sum length [µm] = 30.1, 45.3, 19, 5.1
15
10
5
0
−5
−10
−15
−20
−15
−10
−5
0
5
10

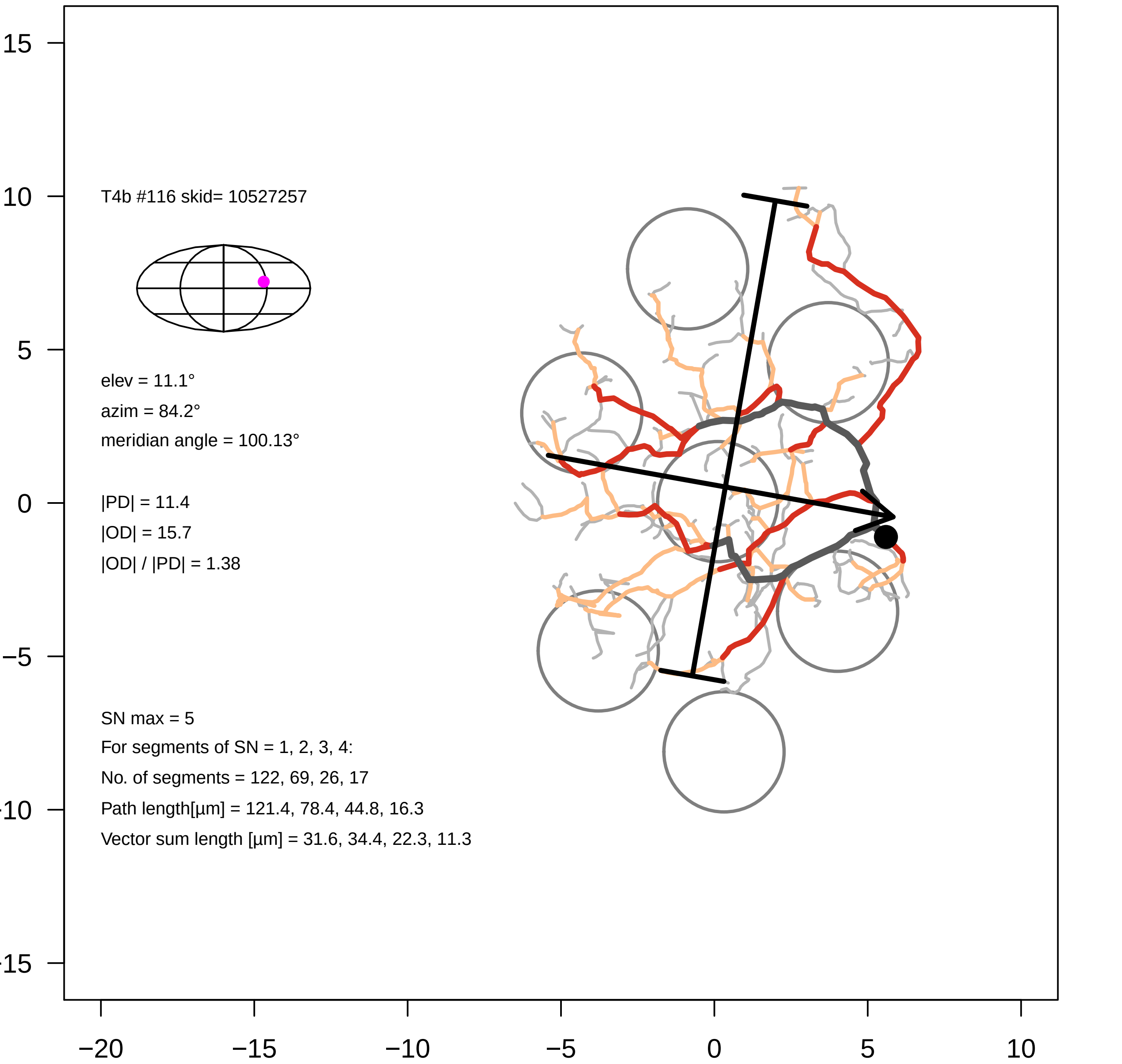

T4b #116 skid= 10527257
elev = 11.1°
azim = 84.2°
meridian angle = 100.13°
|PD| = 11.4
|OD| = 15.7
|OD| / |PD| = 1.38
SN max = 5
For segments of SN = 1, 2, 3, 4:
No. of segments = 122, 69, 26, 17
Path length[µm] = 121.4, 78.4, 44.8, 16.3
Vector sum length [µm] = 31.6, 34.4, 22.3, 11.3
−20 −15 −10 −5 0 5 10
15 10 5 0 −5 −10 −15

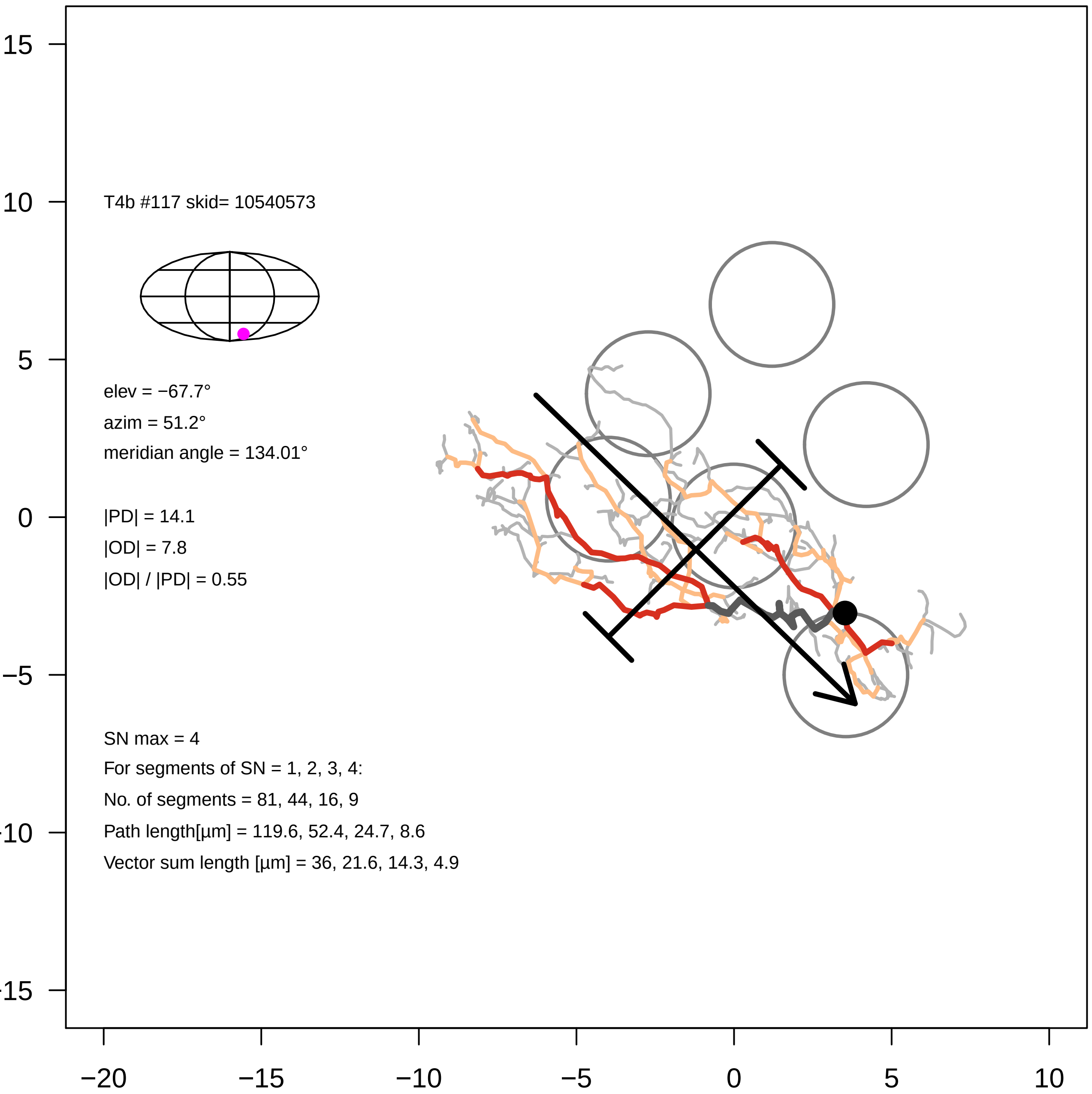

T4b #117 skid= 10540573
elev = −67.7°
azim = 51.2°
meridian angle = 134.01°
|PD| = 14.1
|OD| = 7.8
|OD| / |PD| = 0.55
SN max = 4
For segments of SN = 1, 2, 3, 4:
No. of segments = 81, 44, 16, 9
Path length[µm] = 119.6, 52.4, 24.7, 8.6
Vector sum length [µm] = 36, 21.6, 14.3, 4.9
15
10
5
0
−5
−10
−15
−20
−15
−10
−5
0
5
10

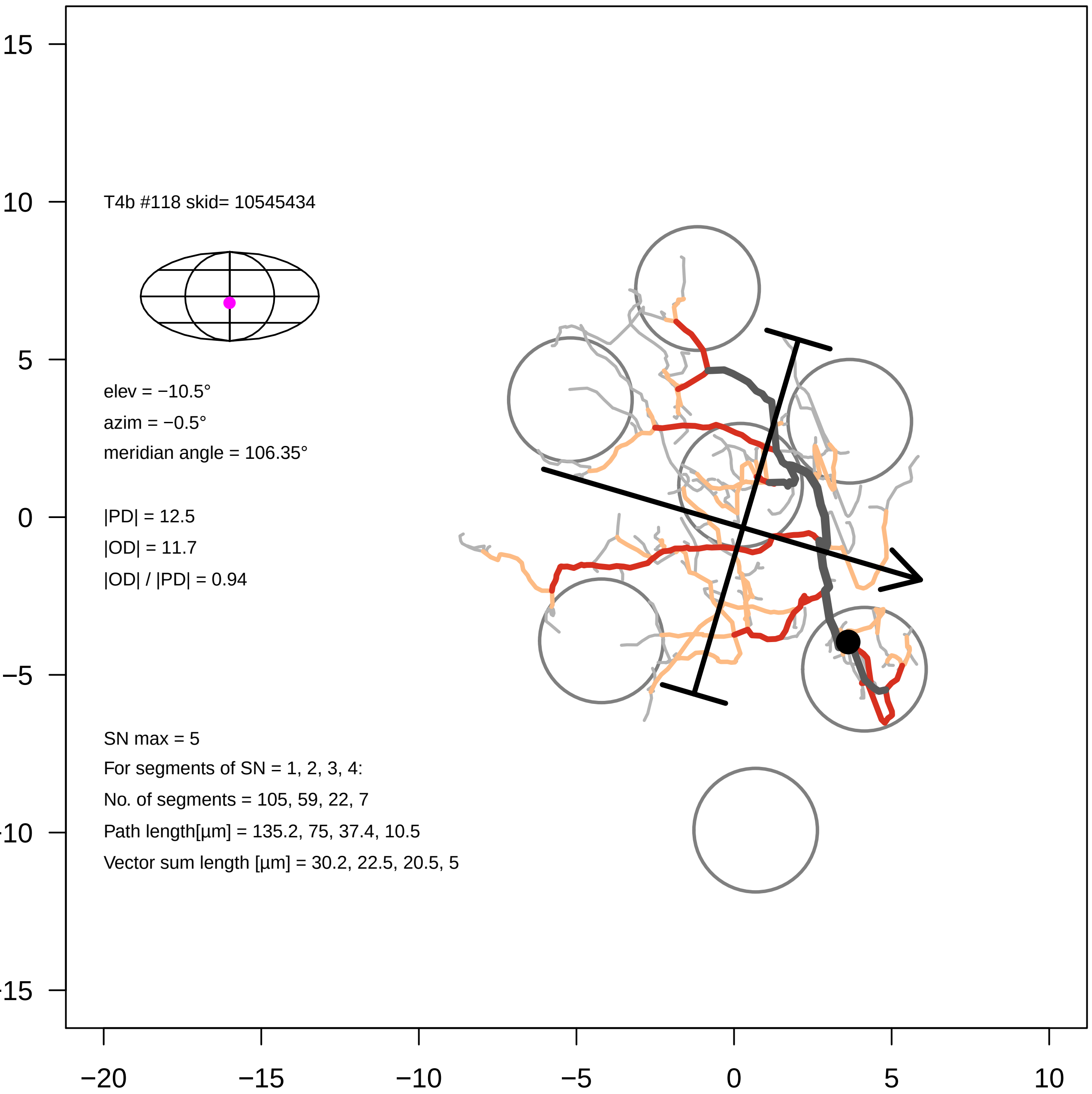
T4b #118 skid= 10545434
elev = −10.5°
azim = −0.5°
meridian angle = 106.35°
|PD| = 12.5
|OD| = 11.7
|OD| / |PD| = 0.94
SN max = 5
For segments of SN = 1, 2, 3, 4:
No. of segments = 105, 59, 22, 7
Path length[µm] = 135.2, 75, 37.4, 10.5
Vector sum length [µm] = 30.2, 22.5, 20.5, 5
15
10
5
0
−5
−10
−15
−20
−15
−10
−5
0
5
10

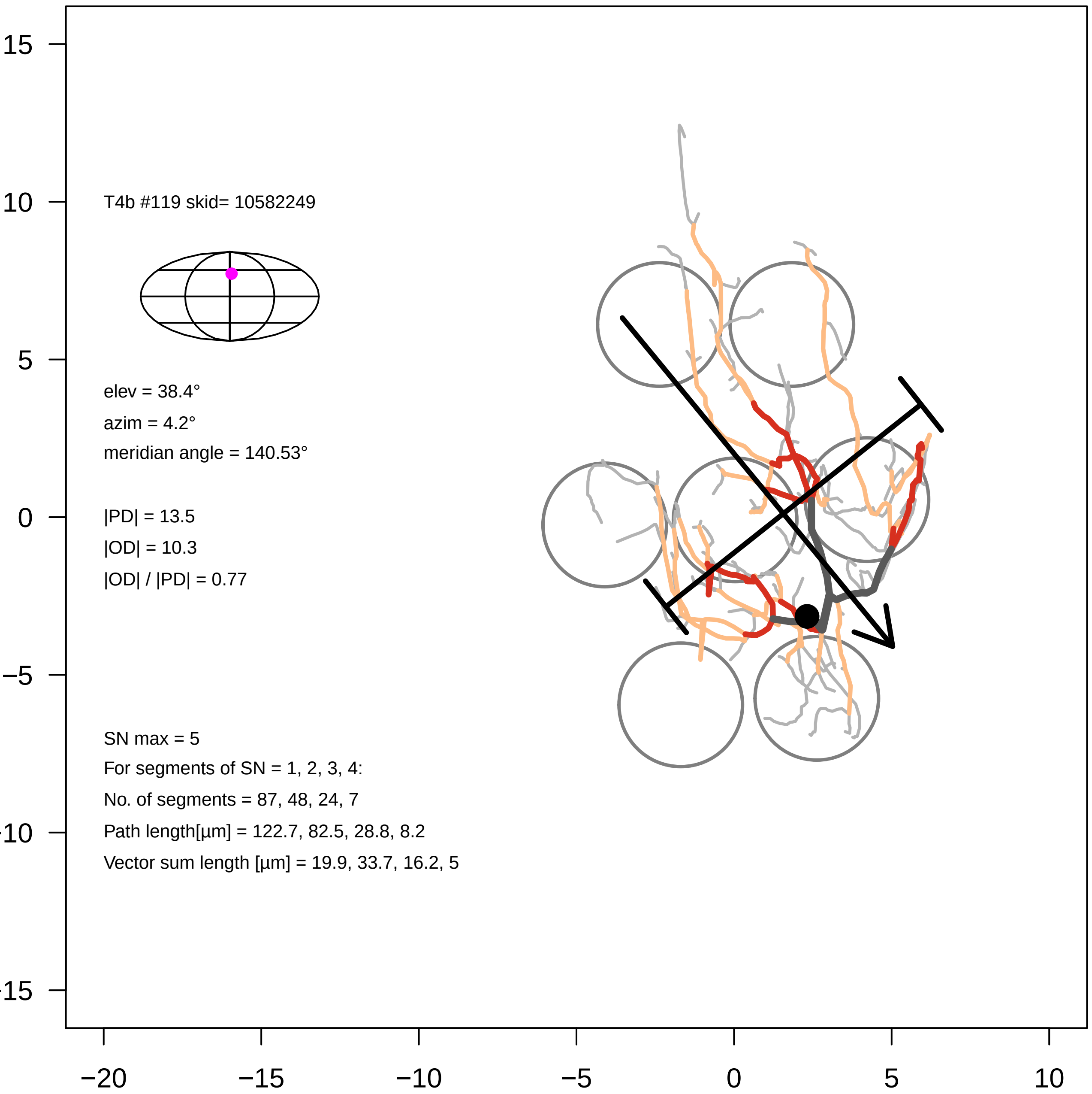
T4b #119 skid= 10582249
elev = 38.4°
azim = 4.2°
meridian angle = 140.53°
|PD| = 13.5
|OD| = 10.3
|OD| / |PD| = 0.77
SN max = 5
For segments of SN = 1, 2, 3, 4:
No. of segments = 87, 48, 24, 7
Path length[µm] = 122.7, 82.5, 28.8, 8.2
Vector sum length [µm] = 19.9, 33.7, 16.2, 5
15
10
5
0
−5
−10
−15
−20
−15
−10
−5
0
5
10

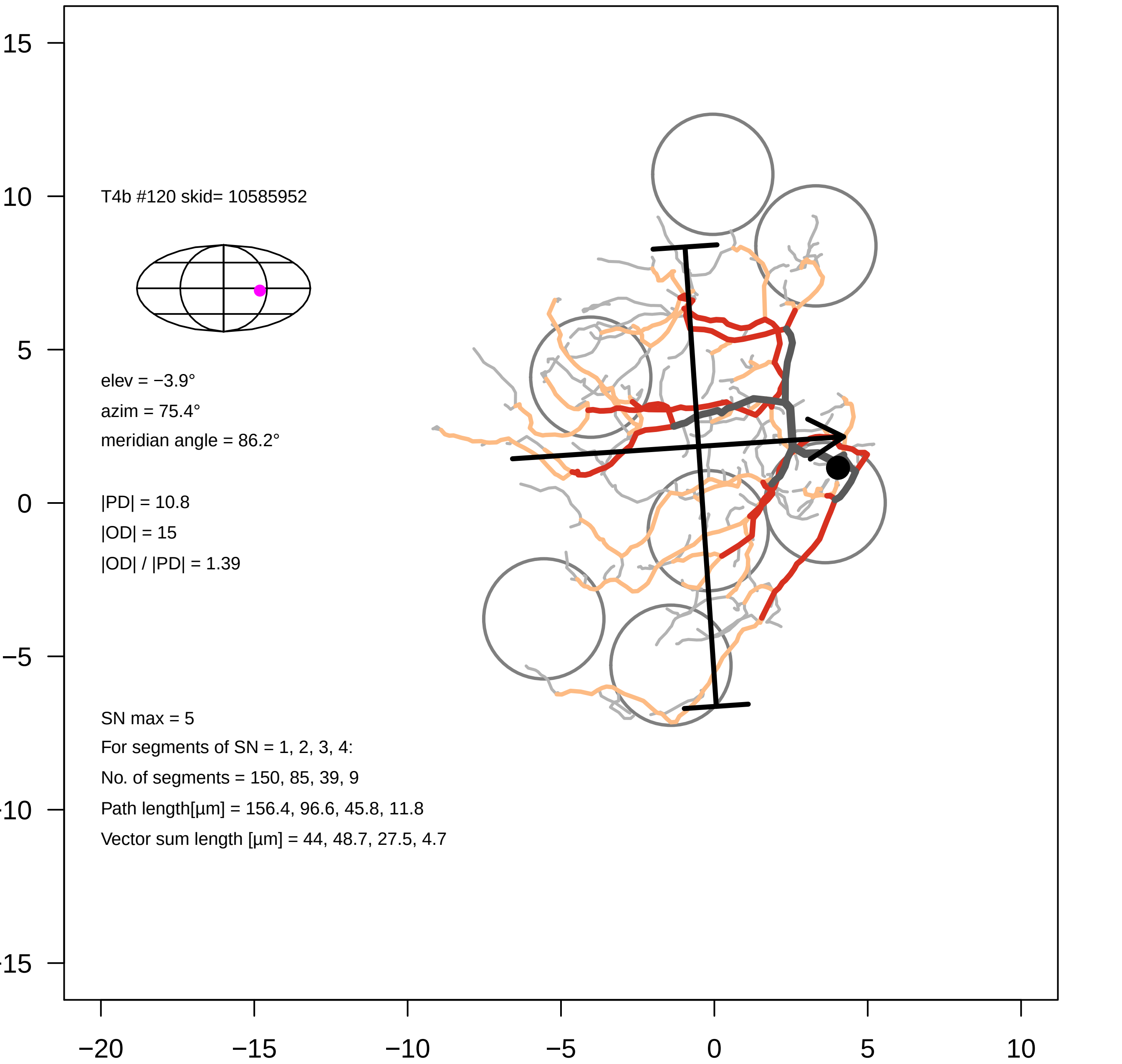
T4b #120 skid= 10585952
elev = −3.9°
azim = 75.4°
meridian angle = 86.2°
|PD| = 10.8
|OD| = 15
|OD| / |PD| = 1.39
SN max = 5
For segments of SN = 1, 2, 3, 4:
No. of segments = 150, 85, 39, 9
Path length[µm] = 156.4, 96.6, 45.8, 11.8
Vector sum length [µm] = 44, 48.7, 27.5, 4.7
15
10
5
0
−5
−10
−15
−20
−15
−10
−5
0
5
10

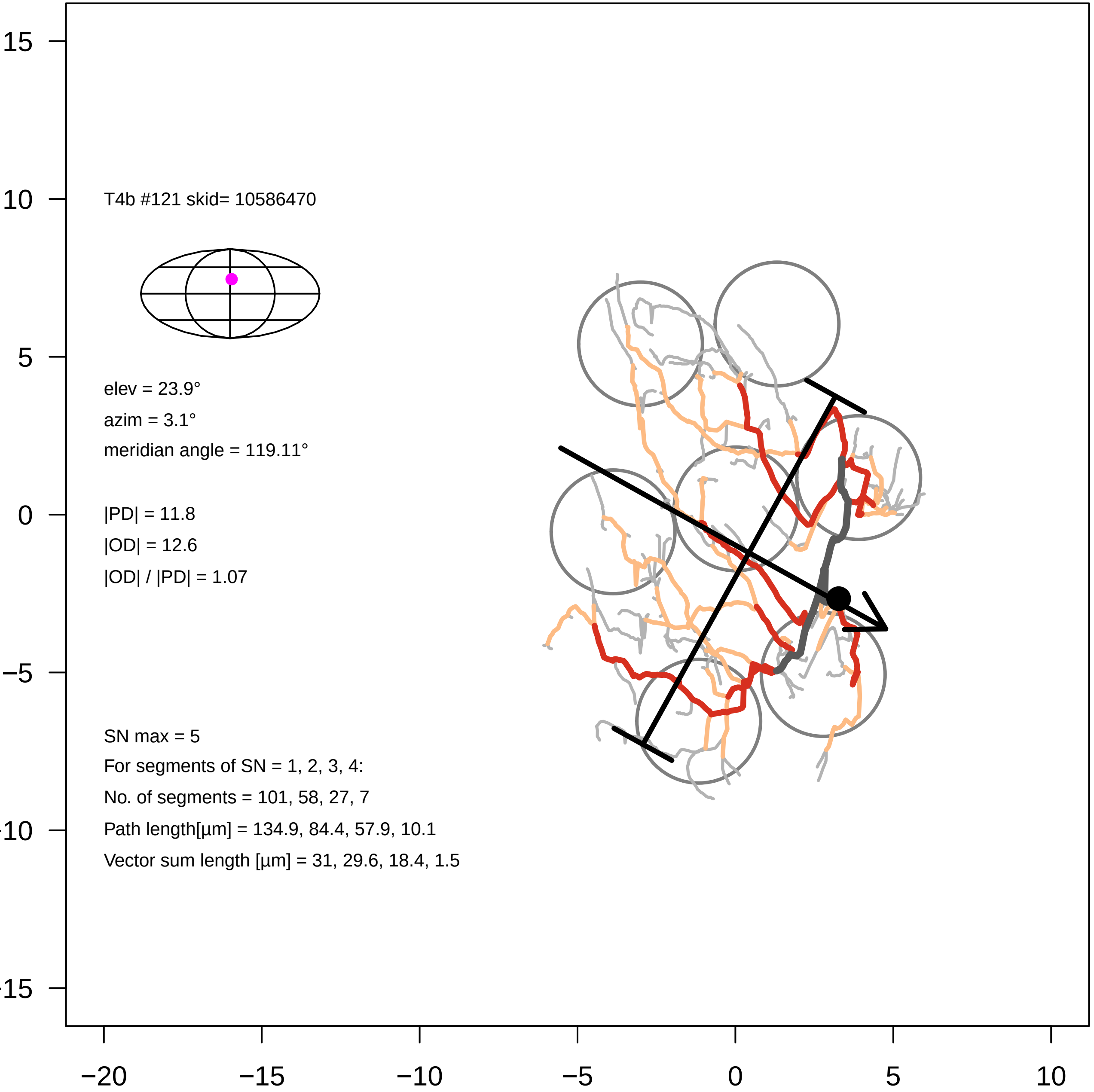

T4b #121 skid= 10586470
elev = 23.9°
azim = 3.1°
meridian angle = 119.11°
|PD| = 11.8
|OD| = 12.6
|OD| / |PD| = 1.07
SN max = 5
For segments of SN = 1, 2, 3, 4:
No. of segments = 101, 58, 27, 7
Path length[µm] = 134.9, 84.4, 57.9, 10.1
Vector sum length [µm] = 31, 29.6, 18.4, 1.5
15
10
5
0
−5
−10
−15
−20
−15
−10
−5
0
5
10

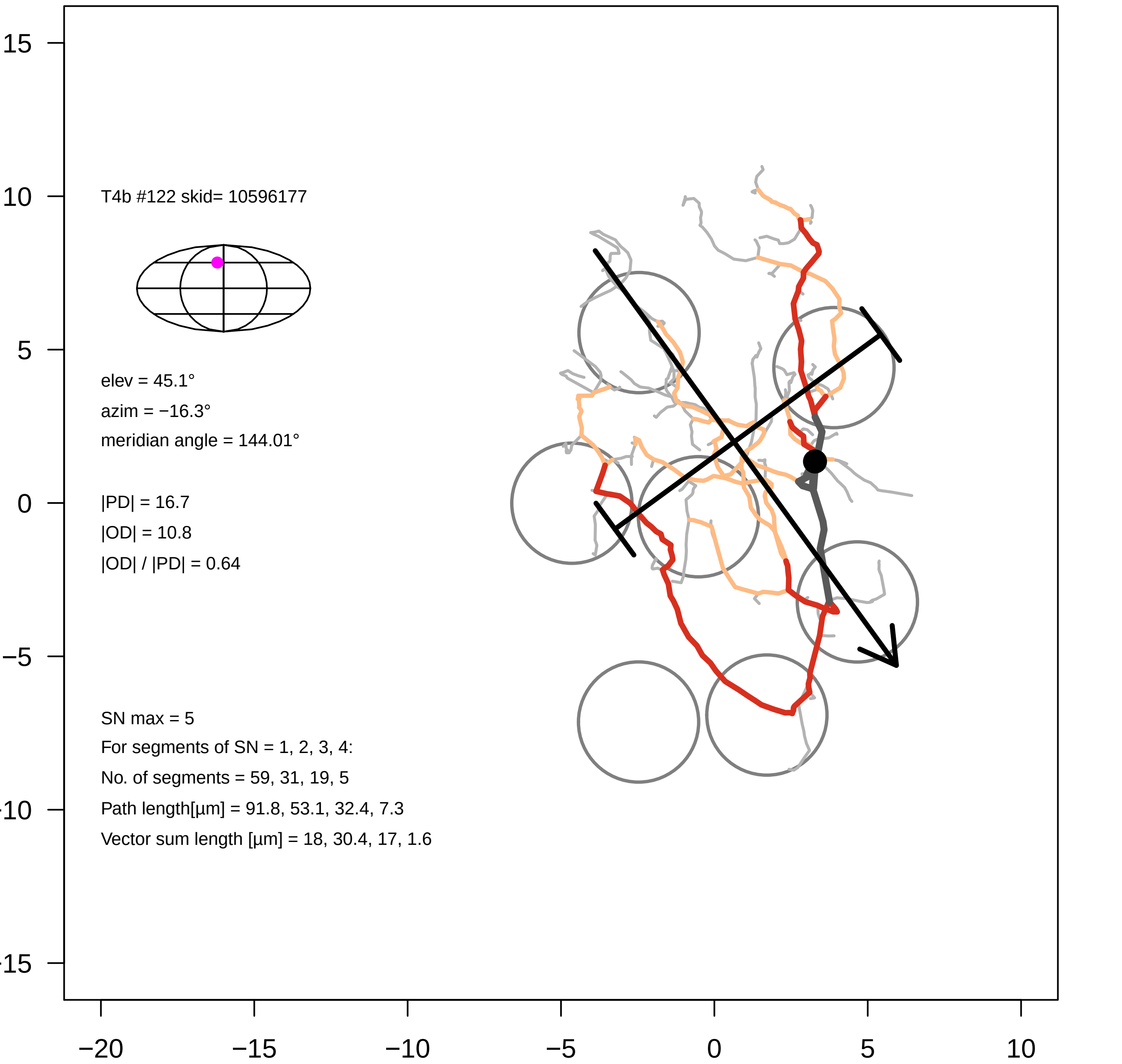

T4b #122 skid= 10596177
elev = 45.1°
azim = −16.3°
meridian angle = 144.01°
|PD| = 16.7
|OD| = 10.8
|OD| / |PD| = 0.64
SN max = 5
For segments of SN = 1, 2, 3, 4:
No. of segments = 59, 31, 19, 5
Path length[µm] = 91.8, 53.1, 32.4, 7.3
Vector sum length [µm] = 18, 30.4, 17, 1.6
15
10
5
0
−5
-10
-15
−20
−15
−10
−5
0
5
10

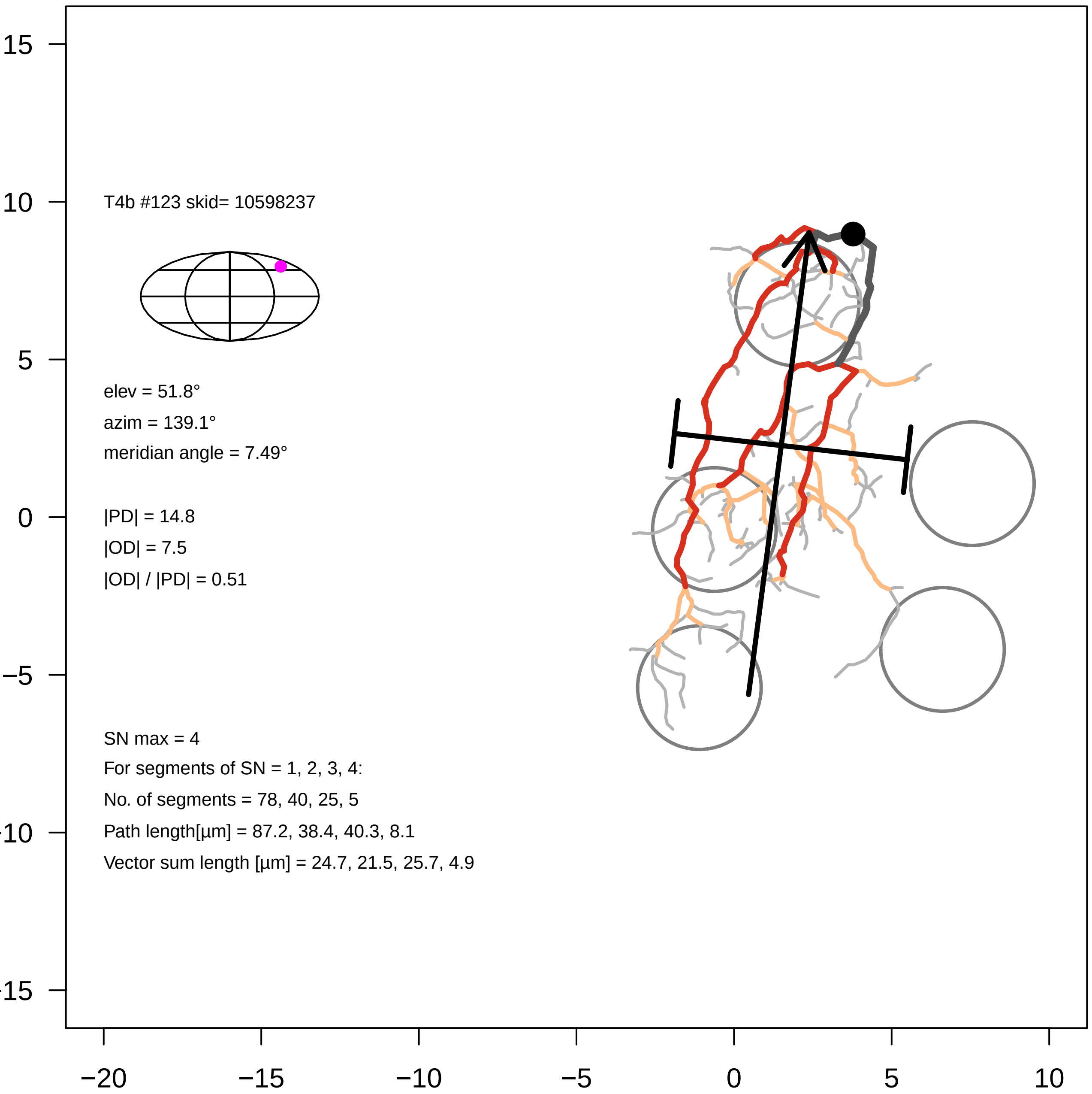
T4b #123 skid= 10598237
elev = 51.8°
azim = 139.1°
meridian angle = 7.49°
|PD| = 14.8
|OD| = 7.5
|OD| / |PD| = 0.51
SN max = 4
For segments of SN = 1, 2, 3, 4:
No. of segments = 78, 40, 25, 5
Path length[µm] = 87.2, 38.4, 40.3, 8.1
Vector sum length [µm] = 24.7, 21.5, 25.7, 4.9
15
10
5
0
−5
-10
-15
−20
−15
−10
−5
0
5
10

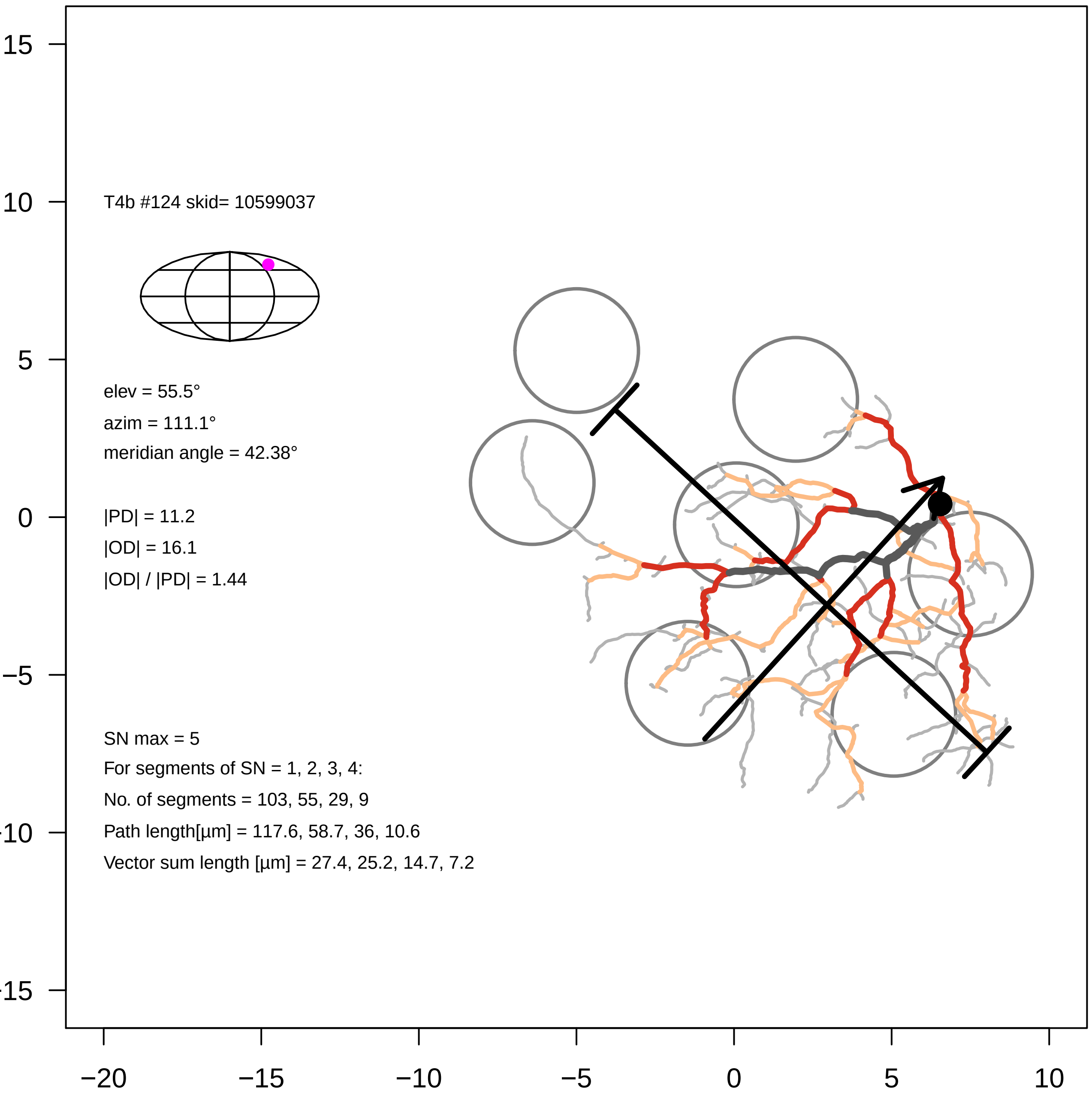
T4b #124 skid= 10599037
elev = 55.5°
azim = 111.1°
meridian angle = 42.38°
|PD| = 11.2
|OD| = 16.1
|OD| / |PD| = 1.44
SN max = 5
For segments of SN = 1, 2, 3, 4:
No. of segments = 103, 55, 29, 9
Path length[µm] = 117.6, 58.7, 36, 10.6
Vector sum length [µm] = 27.4, 25.2, 14.7, 7.2
15
10
5
0
−5
−10
−15
−20
−15
−10
−5
0
5
10

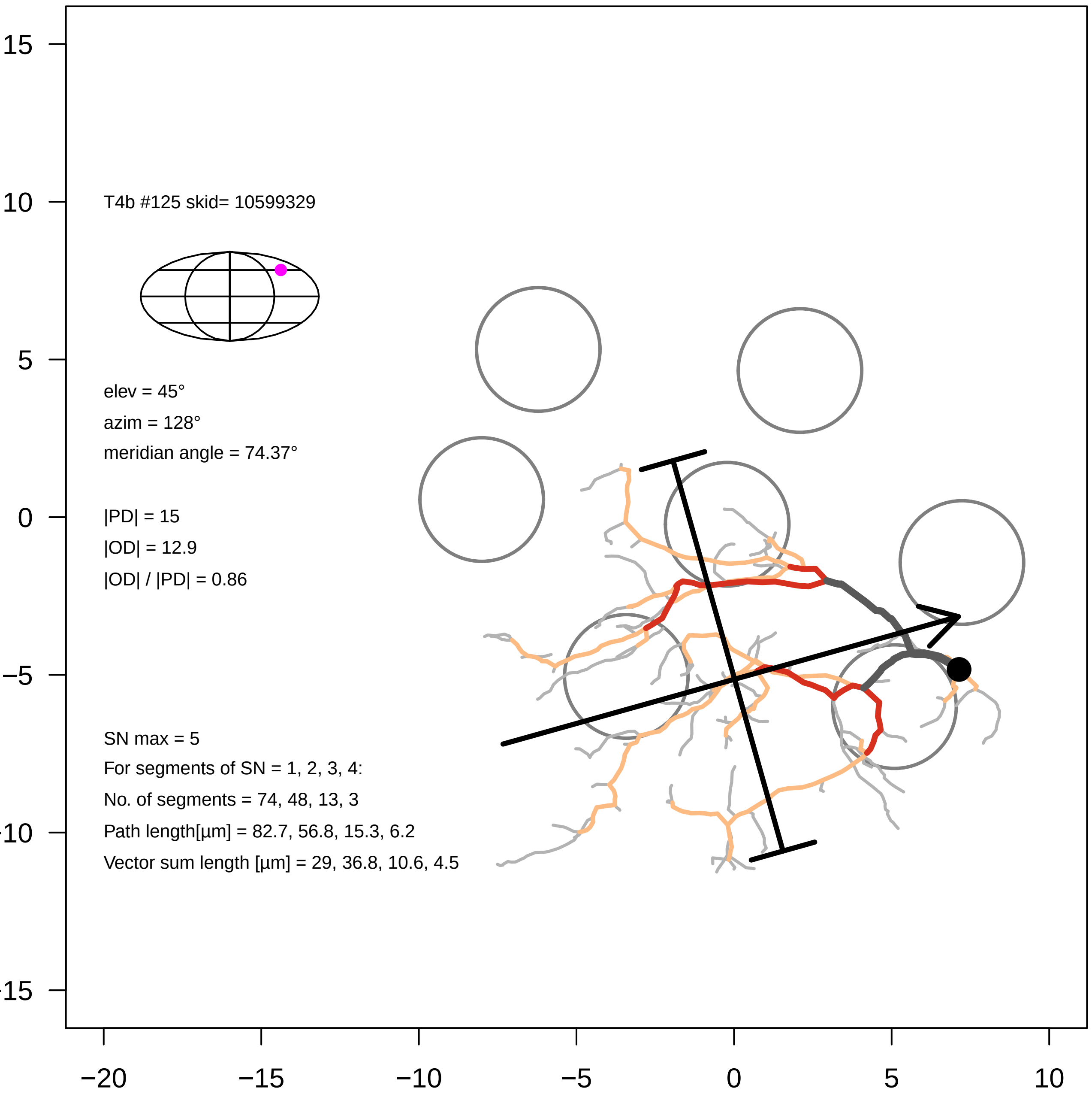
T4b #125 skid= 10599329
elev = 45°
azim = 128°
meridian angle = 74.37°
|PD| = 15
|OD| = 12.9
|OD| / |PD| = 0.86
SN max = 5
For segments of SN = 1, 2, 3, 4:
No. of segments = 74, 48, 13, 3
Path length[µm] = 82.7, 56.8, 15.3, 6.2
Vector sum length [µm] = 29, 36.8, 10.6, 4.5
15
10
5
0
−5
−10
−15
−20
−15
−10
−5
0
5
10

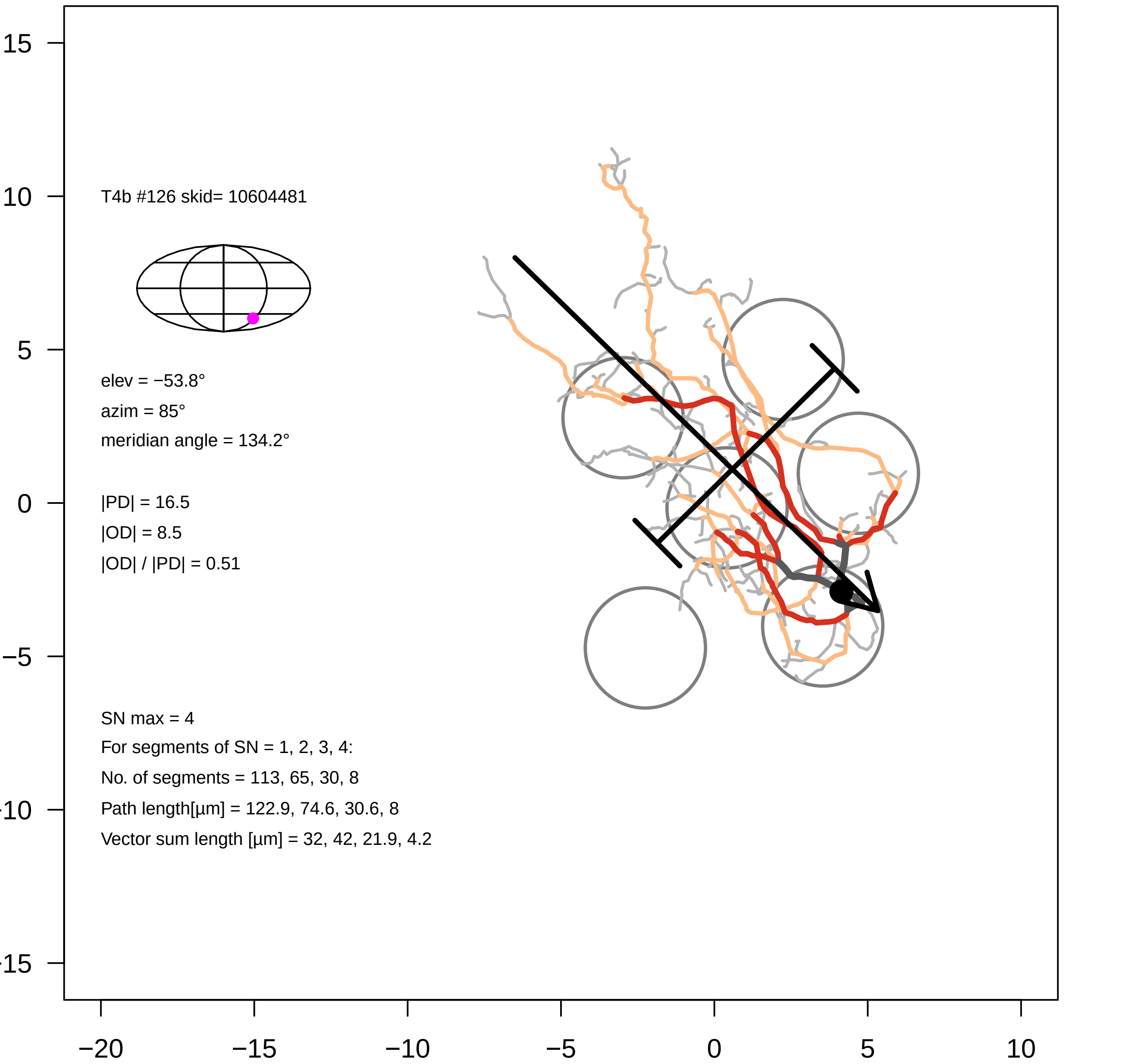

T4b #126 skid= 10604481
elev = −53.8°
azim = 85°
meridian angle = 134.2°
|PD| = 16.5
|OD| = 8.5
|OD| / |PD| = 0.51
SN max = 4
For segments of SN = 1, 2, 3, 4:
No. of segments = 113, 65, 30, 8
Path length[µm] = 122.9, 74.6, 30.6, 8
Vector sum length [µm] = 32, 42, 21.9, 4.2
15
10
5
0
−5
-10
-15
−20
−15
−10
−5
0
5
10

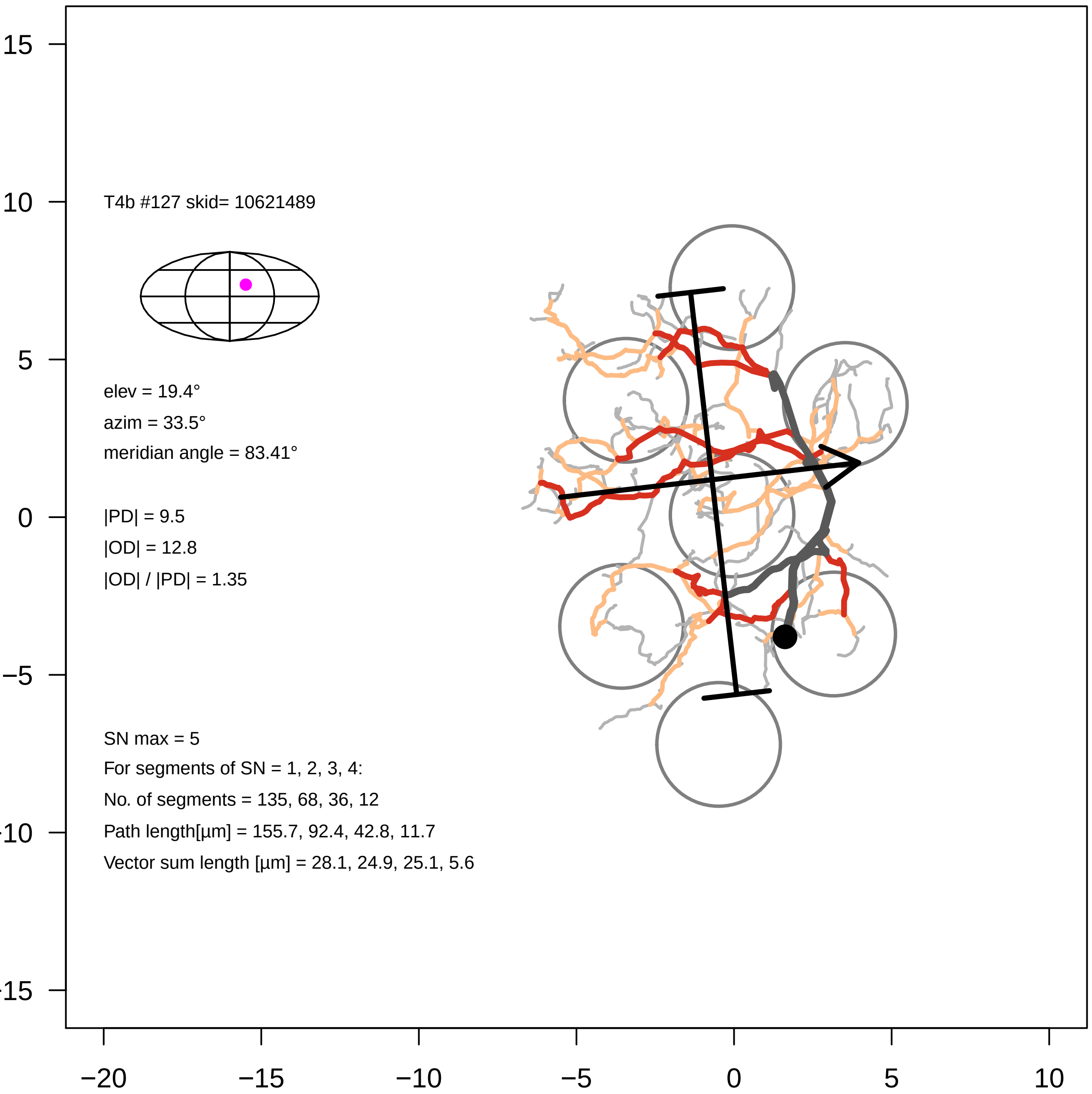

T4b #127 skid= 10621489
elev = 19.4°
azim = 33.5°
meridian angle = 83.41°
|PD| = 9.5
|OD| = 12.8
|OD| / |PD| = 1.35
SN max = 5
For segments of SN = 1, 2, 3, 4:
No. of segments = 135, 68, 36, 12
Path length[µm] = 155.7, 92.4, 42.8, 11.7
Vector sum length [µm] = 28.1, 24.9, 25.1, 5.6
15
10
5
0
−5
−10
−15
−20
−15
−10
−5
0
5
10

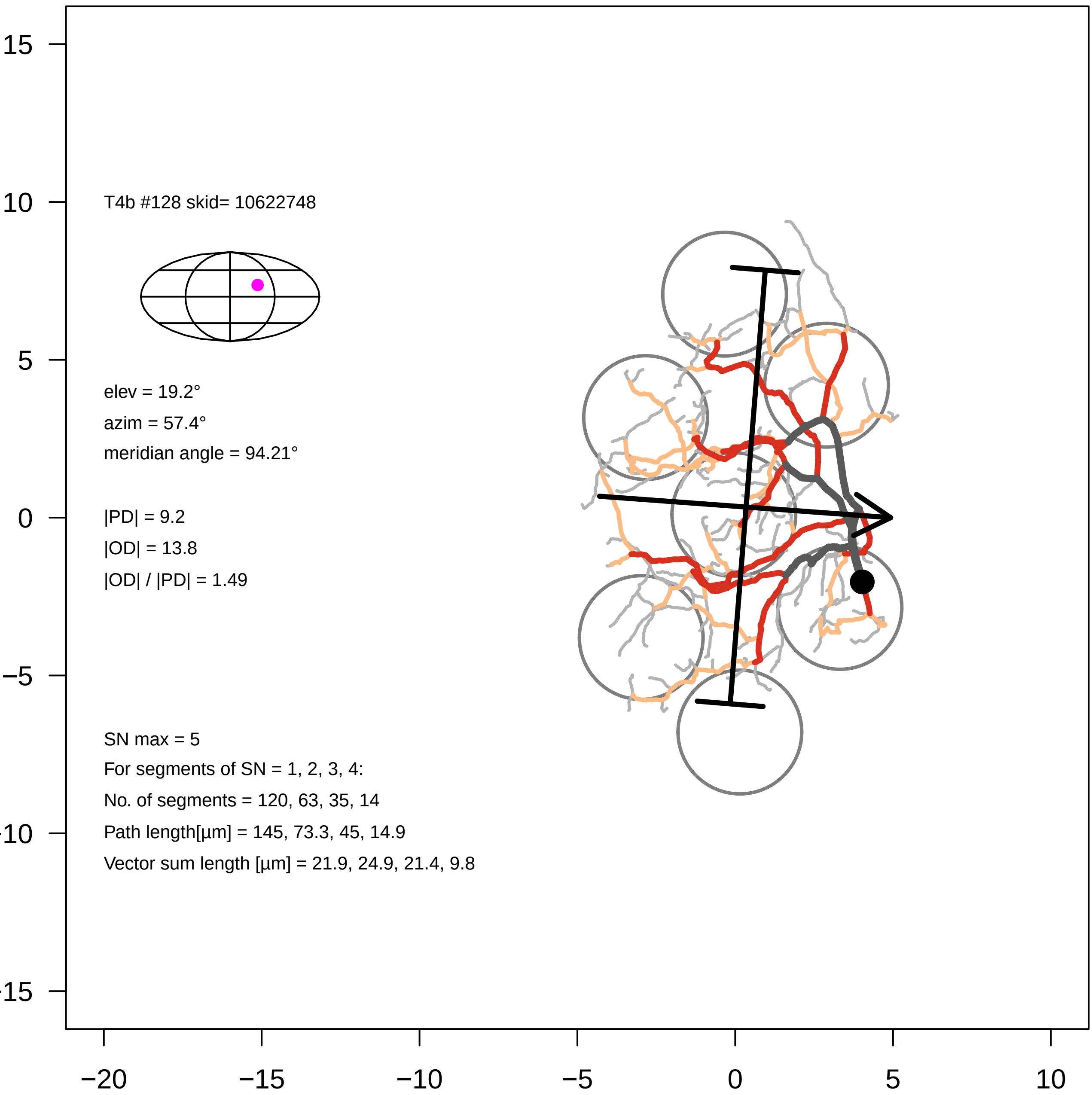
T4b #128 skid= 10622748
elev = 19.2°
azim = 57.4°
meridian angle = 94.21°
|PD| = 9.2
|OD| = 13.8
|OD| / |PD| = 1.49
SN max = 5
For segments of SN = 1, 2, 3, 4:
No. of segments = 120, 63, 35, 14
Path length[µm] = 145, 73.3, 45, 14.9
Vector sum length [µm] = 21.9, 24.9, 21.4, 9.8
15
10
5
0
−5
−10
−15
−20
−15
−10
−5
0
5
10

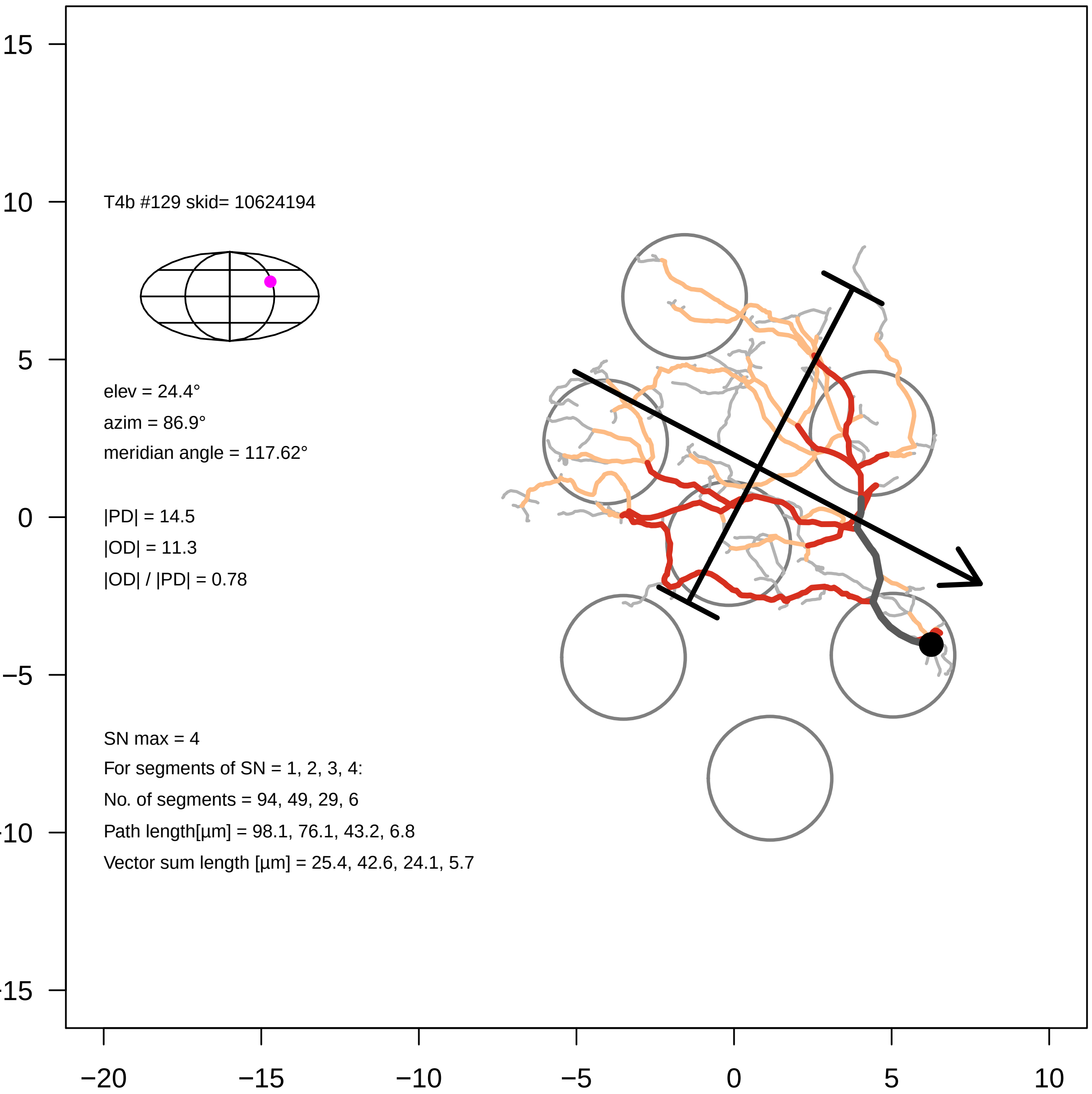
T4b #129 skid= 10624194
elev = 24.4°
azim = 86.9°
meridian angle = 117.62°
|PD| = 14.5
|OD| = 11.3
|OD| / |PD| = 0.78
SN max = 4
For segments of SN = 1, 2, 3, 4:
No. of segments = 94, 49, 29, 6
Path length[µm] = 98.1, 76.1, 43.2, 6.8
Vector sum length [µm] = 25.4, 42.6, 24.1, 5.7
15
10
5
0
−5
−10
−15
−20
−15
−10
−5
0
5
10

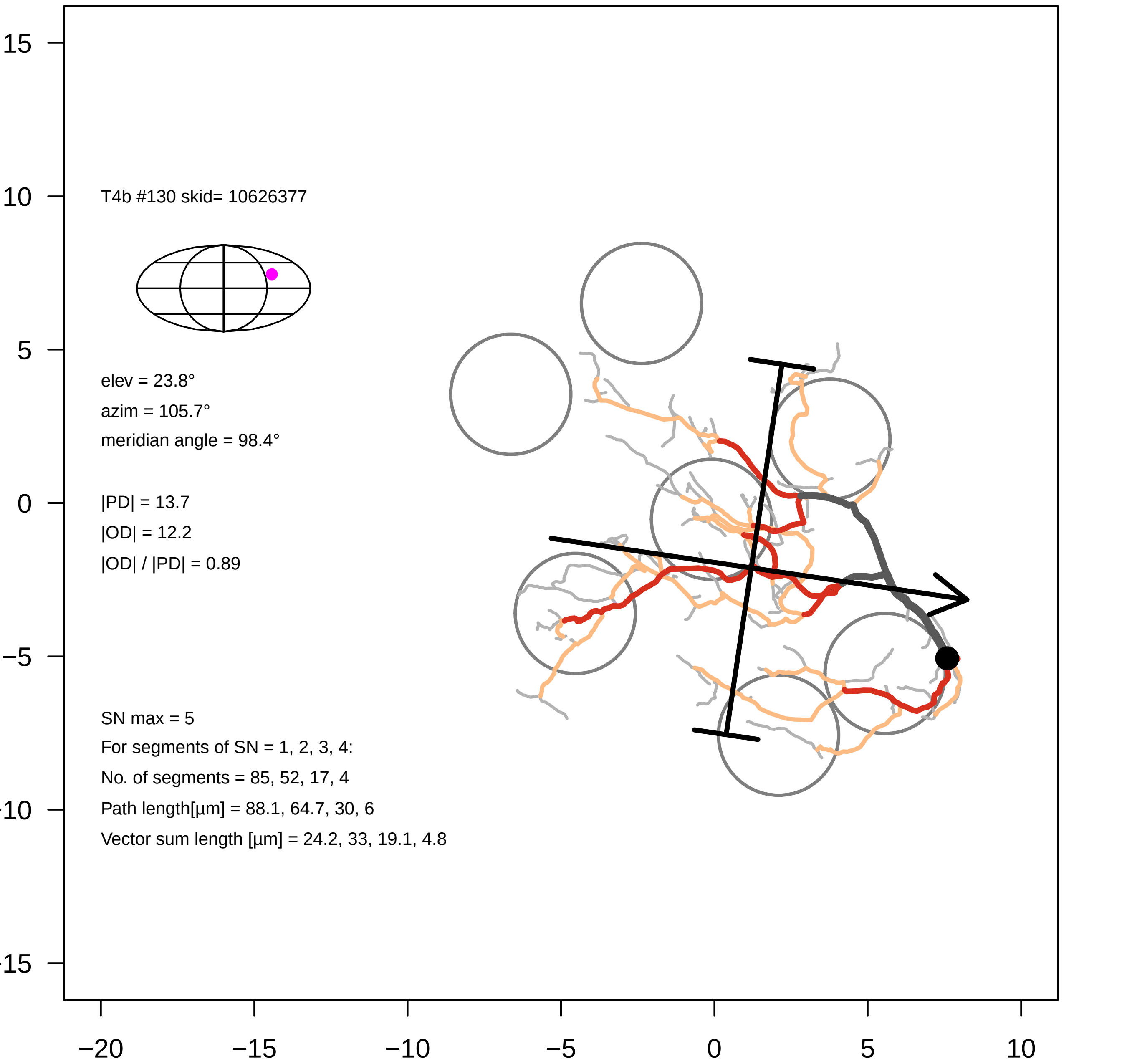
T4b #130 skid= 10626377
elev = 23.8°
azim = 105.7°
meridian angle = 98.4°
|PD| = 13.7
|OD| = 12.2
|OD| / |PD| = 0.89
SN max = 5
For segments of SN = 1, 2, 3, 4:
No. of segments = 85, 52, 17, 4
Path length[µm] = 88.1, 64.7, 30, 6
Vector sum length [µm] = 24.2, 33, 19.1, 4.8
15
10
5
0
−5
−10
−15
−20
−15
−10
−5
0
5
10

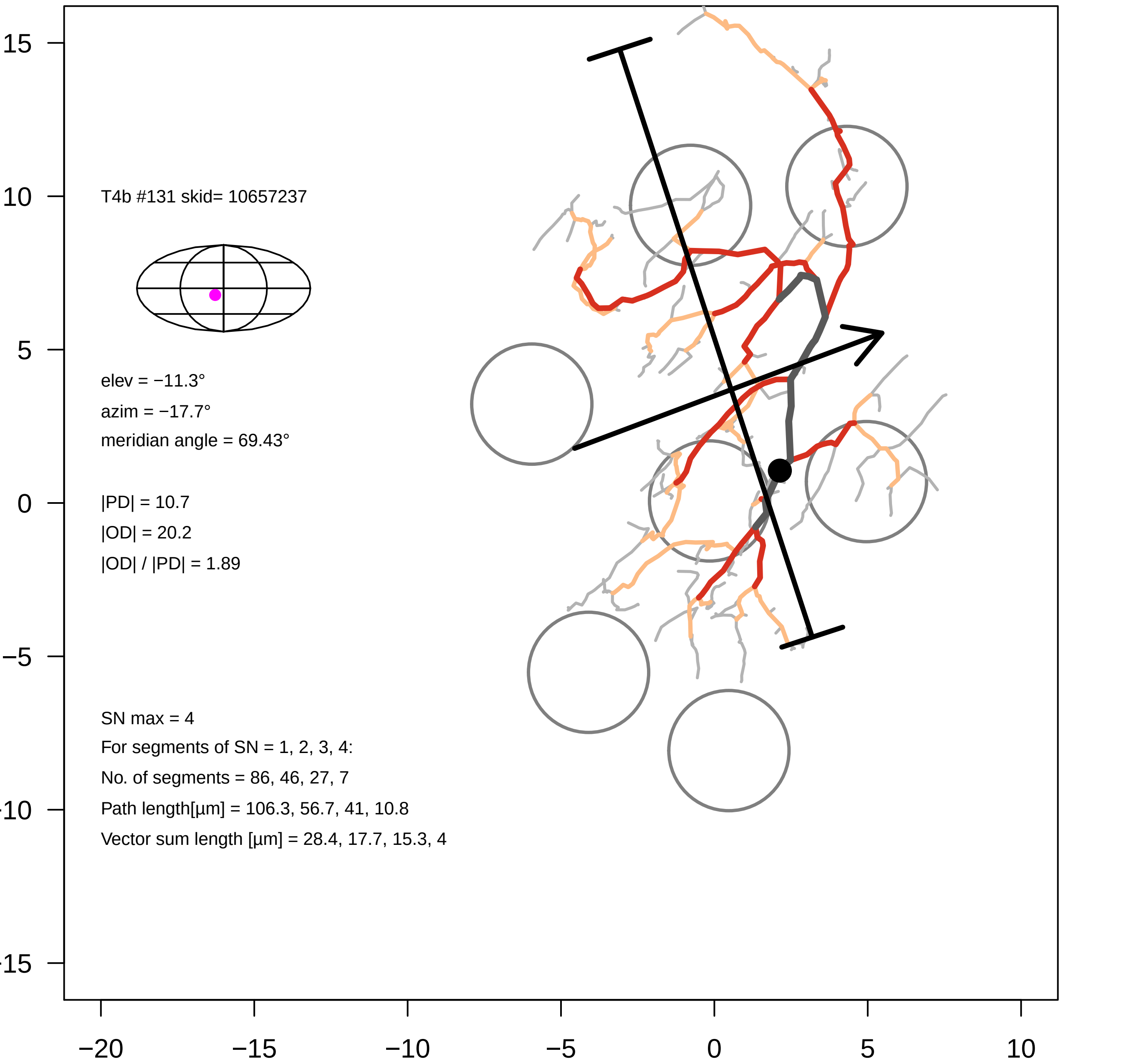

T4b #131 skid= 10657237
elev = −11.3°
azim = −17.7°
meridian angle = 69.43°
|PD| = 10.7
|OD| = 20.2
|OD| / |PD| = 1.89
SN max = 4
For segments of SN = 1, 2, 3, 4:
No. of segments = 86, 46, 27, 7
Path length[µm] = 106.3, 56.7, 41, 10.8
Vector sum length [µm] = 28.4, 17.7, 15.3, 4
15
10
5
0
−5
−10
−15
−20
−15
−10
−5
0
5
10

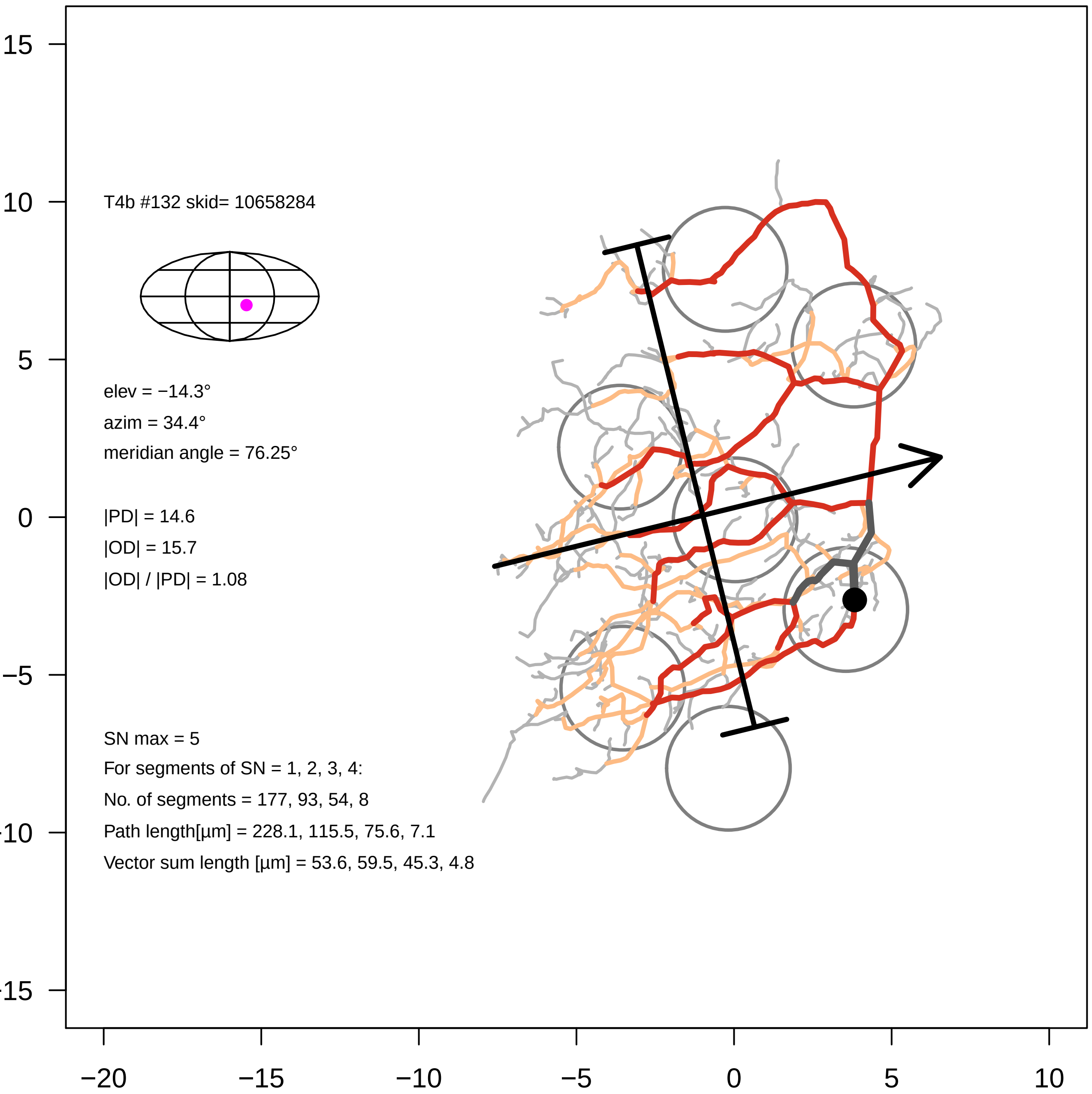
T4b #132 skid= 10658284
elev = −14.3°
azim = 34.4°
meridian angle = 76.25°
|PD| = 14.6
|OD| = 15.7
|OD| / |PD| = 1.08
SN max = 5
For segments of SN = 1, 2, 3, 4:
No. of segments = 177, 93, 54, 8
Path length[µm] = 228.1, 115.5, 75.6, 7.1
Vector sum length [µm] = 53.6, 59.5, 45.3, 4.8
15
10
5
0
−5
−10
−15
−20
−15
−10
−5
0
5
10

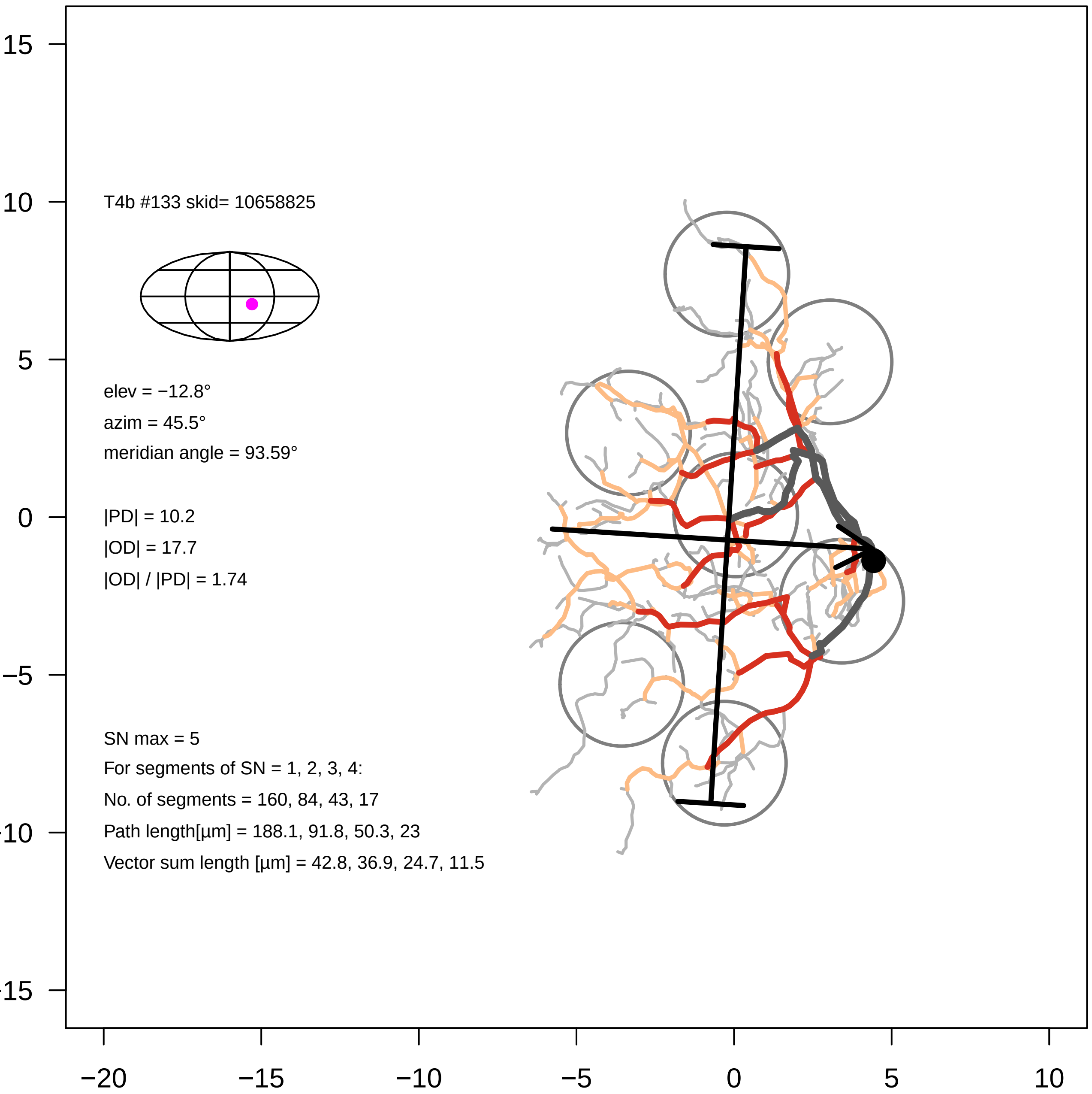
T4b #133 skid= 10658825
elev = −12.8°
azim = 45.5°
meridian angle = 93.59°
|PD| = 10.2
|OD| = 17.7
|OD| / |PD| = 1.74
SN max = 5
For segments of SN = 1, 2, 3, 4:
No. of segments = 160, 84, 43, 17
Path length[µm] = 188.1, 91.8, 50.3, 23
Vector sum length [µm] = 42.8, 36.9, 24.7, 11.5
15
10
5
0
−5
−20
−15
−10
−5
0
5
10

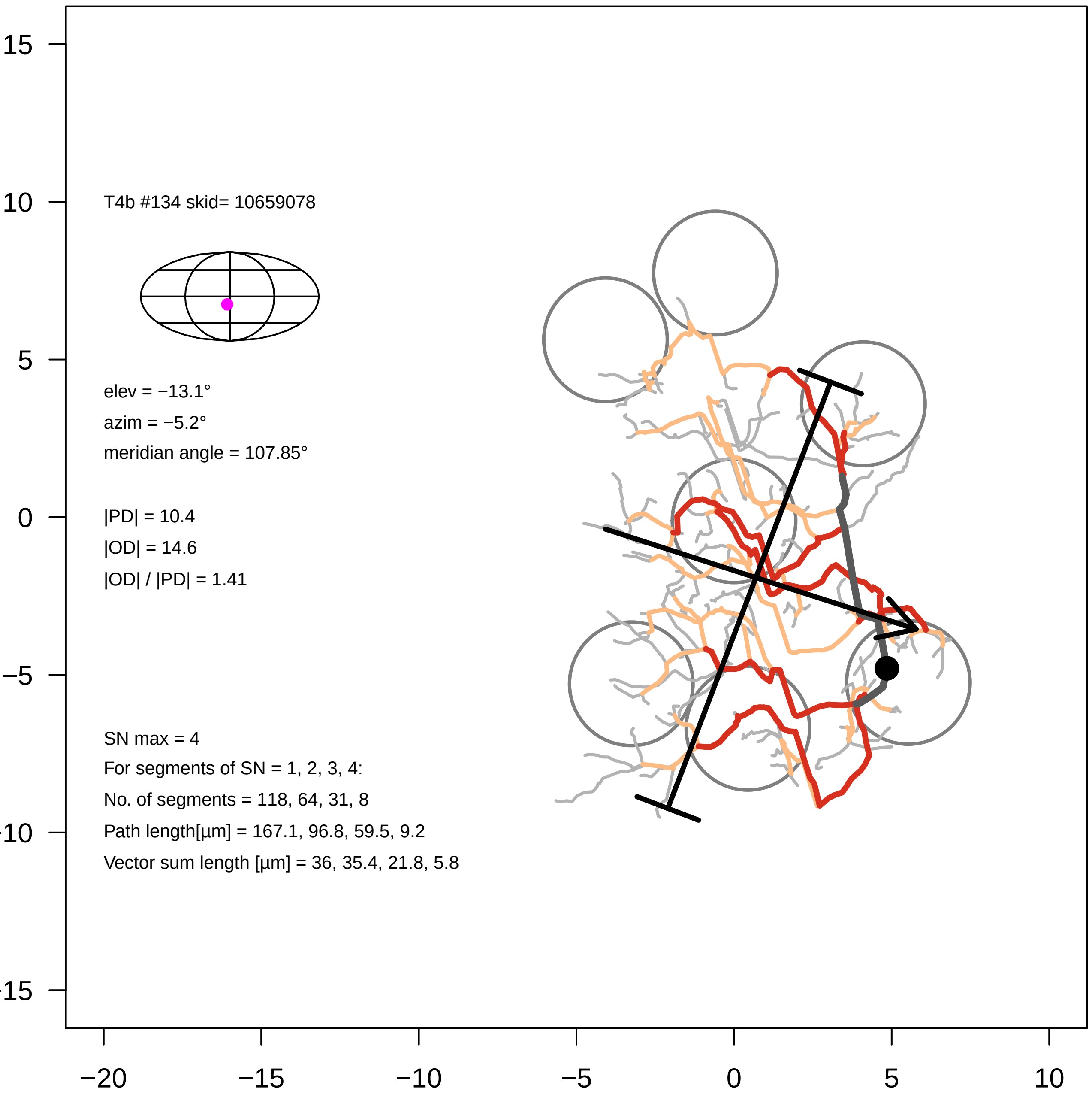
T4b #134 skid= 10659078
elev = −13.1°
azim = −5.2°
meridian angle = 107.85°
|PD| = 10.4
|OD| = 14.6
|OD| / |PD| = 1.41
SN max = 4
For segments of SN = 1, 2, 3, 4:
No. of segments = 118, 64, 31, 8
Path length[µm] = 167.1, 96.8, 59.5, 9.2
Vector sum length [µm] = 36, 35.4, 21.8, 5.8
15
10
5
0
−5
-10
-15
−20
−15
−10
−5
0
5
10

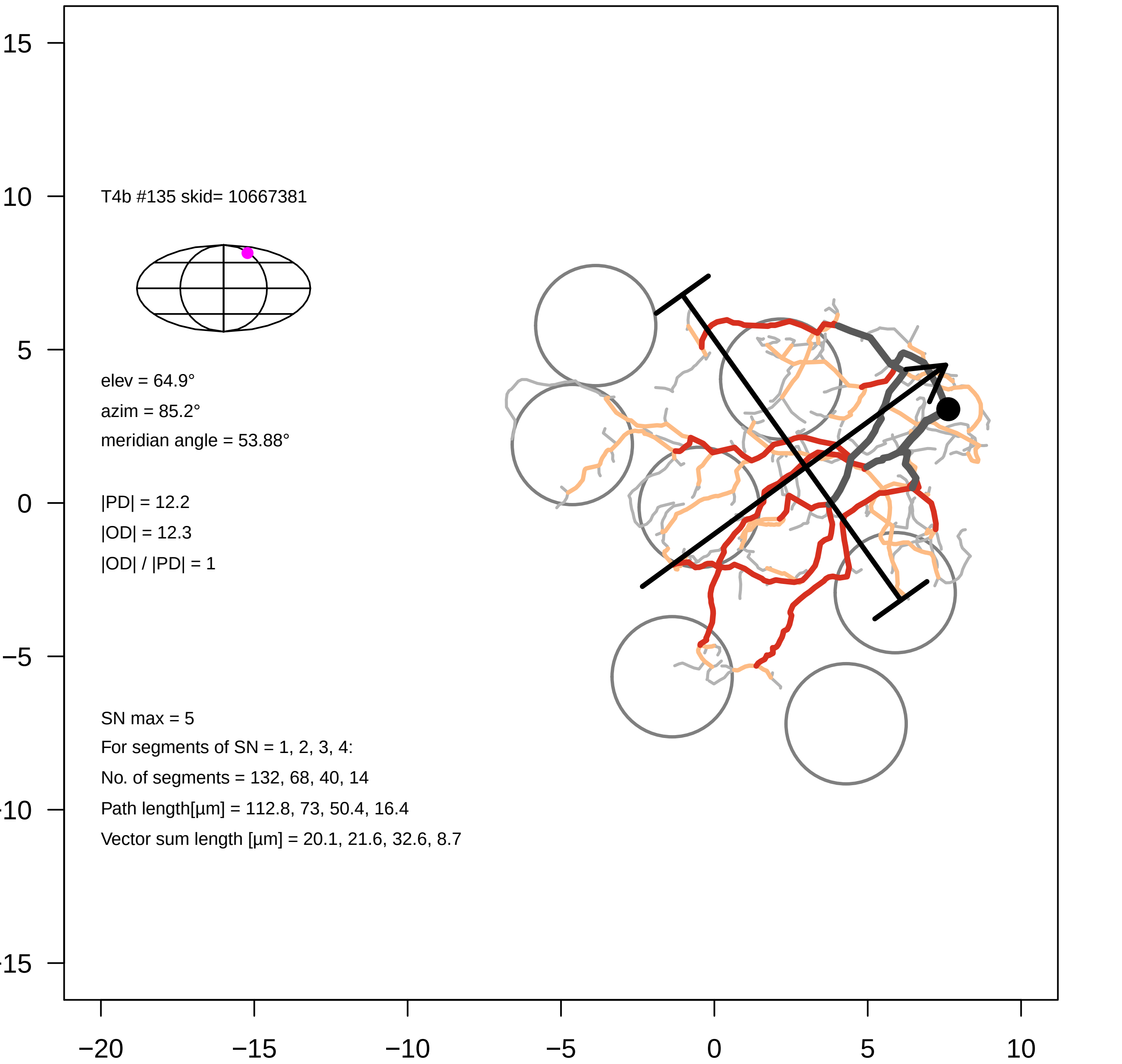
T4b #135 skid= 10667381
elev = 64.9°
azim = 85.2°
meridian angle = 53.88°
|PD| = 12.2
|OD| = 12.3
|OD| / |PD| = 1
SN max = 5
For segments of SN = 1, 2, 3, 4:
No. of segments = 132, 68, 40, 14
Path length[µm] = 112.8, 73, 50.4, 16.4
Vector sum length [µm] = 20.1, 21.6, 32.6, 8.7
15
10
5
0
−5
−10
−15
−20
−15
−10
−5
0
5
10

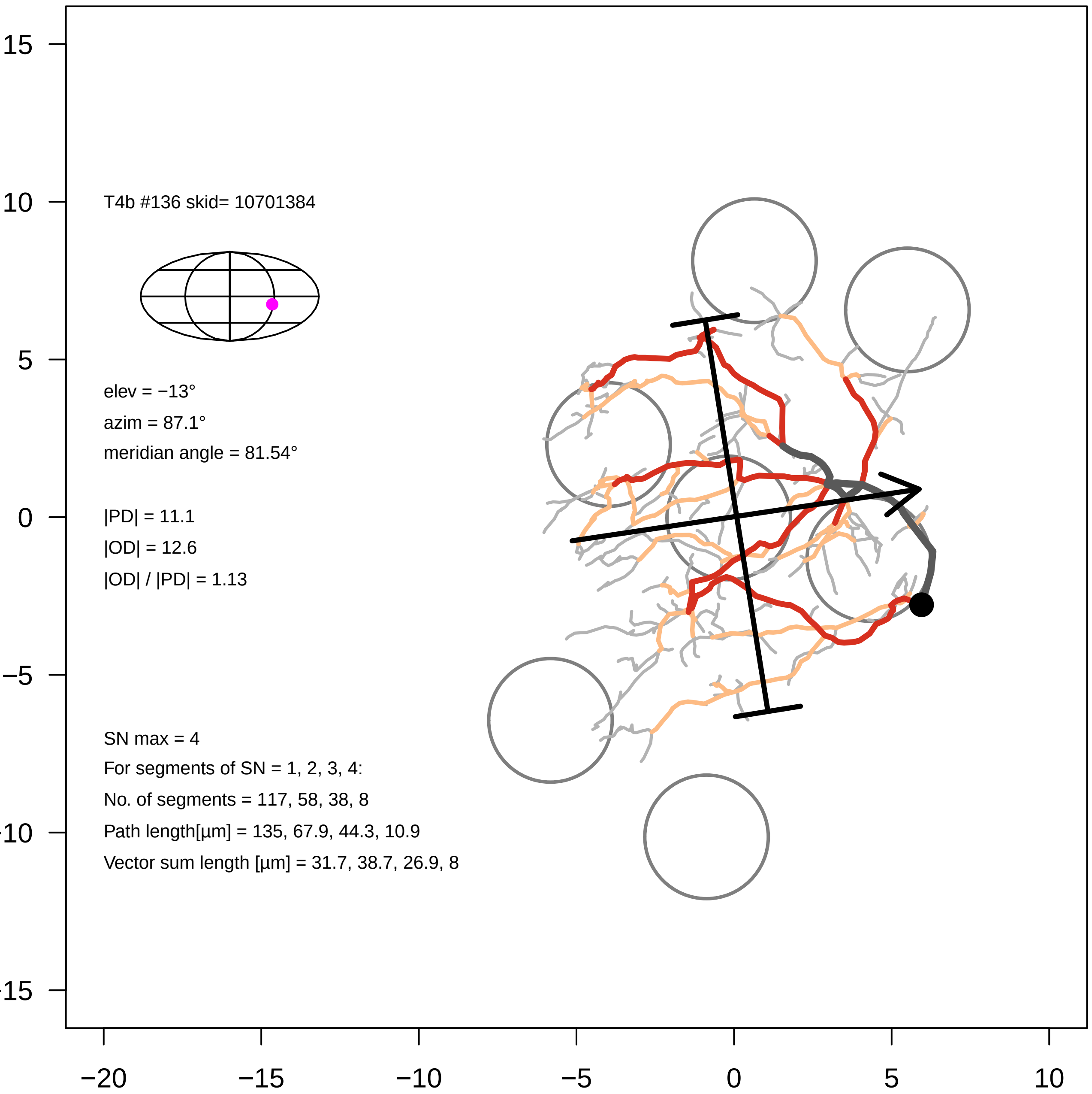
T4b #136 skid= 10701384
elev = −13°
azim = 87.1°
meridian angle = 81.54°
|PD| = 11.1
|OD| = 12.6
|OD| / |PD| = 1.13
SN max = 4
For segments of SN = 1, 2, 3, 4:
No. of segments = 117, 58, 38, 8
Path length[µm] = 135, 67.9, 44.3, 10.9
Vector sum length [µm] = 31.7, 38.7, 26.9, 8
15
10
5
0
−5
−10
−15
−20
−15
−10
−5
0
5
10

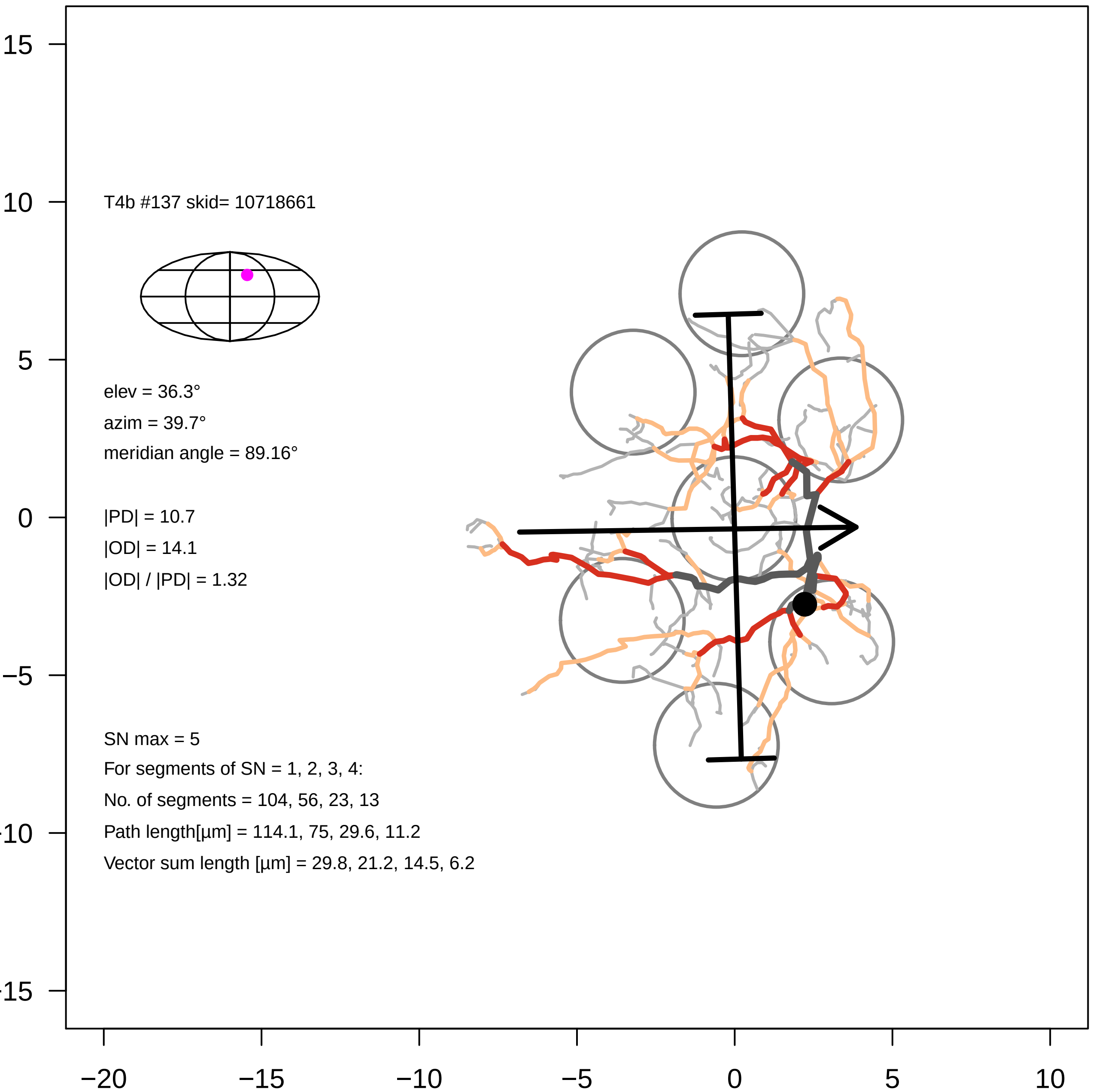
T4b #137 skid= 10718661
elev = 36.3°
azim = 39.7°
meridian angle = 89.16°
|PD| = 10.7
|OD| = 14.1
|OD| / |PD| = 1.32
SN max = 5
For segments of SN = 1, 2, 3, 4:
No. of segments = 104, 56, 23, 13
Path length[µm] = 114.1, 75, 29.6, 11.2
Vector sum length [µm] = 29.8, 21.2, 14.5, 6.2
15
10
5
0
−5
−10
−15
−20
−15
−10
−5
0
5
10

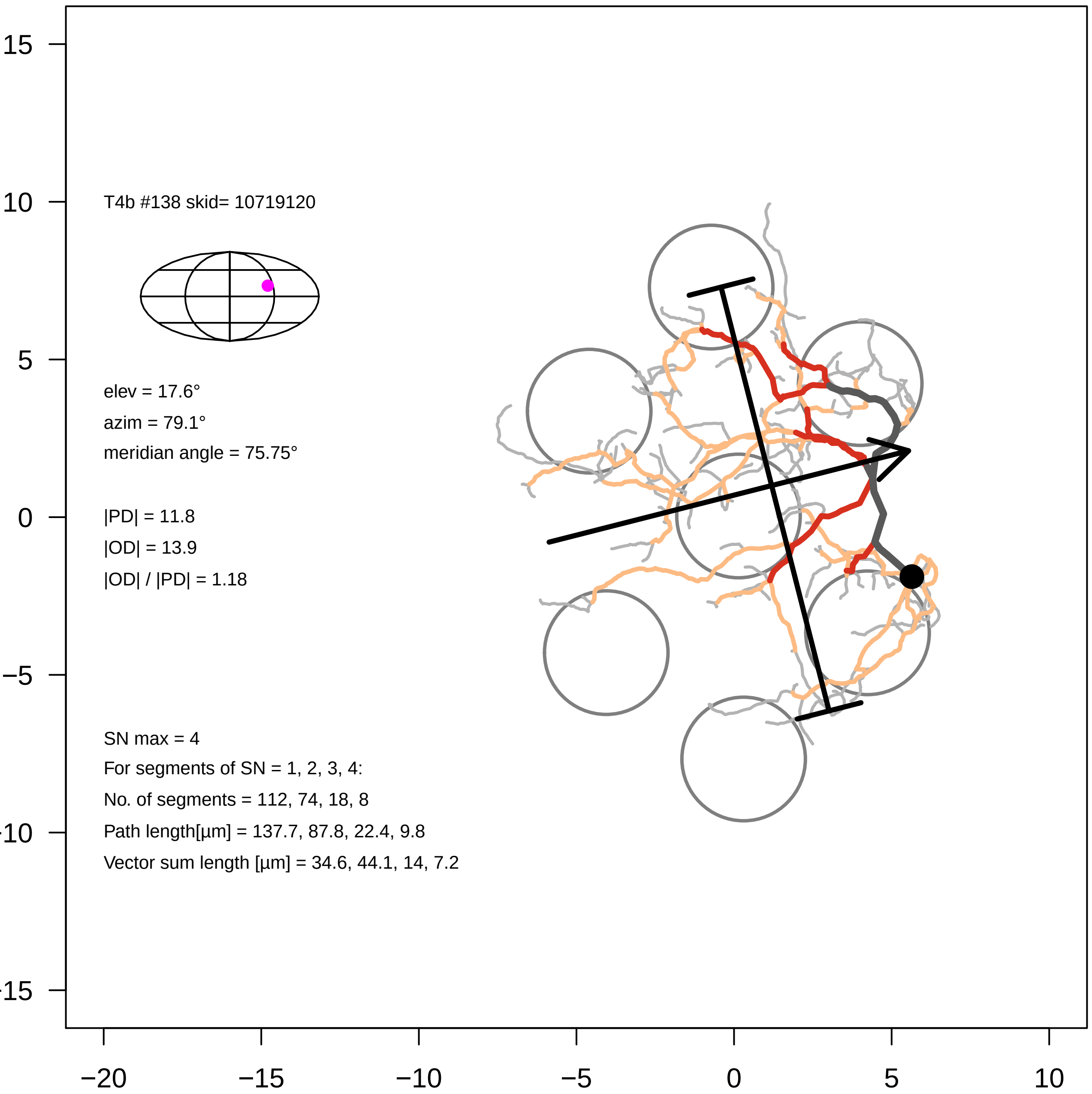
T4b #138 skid= 10719120
elev = 17.6°
azim = 79.1°
meridian angle = 75.75°
|PD| = 11.8
|OD| = 13.9
|OD| / |PD| = 1.18
SN max = 4
For segments of SN = 1, 2, 3, 4:
No. of segments = 112, 74, 18, 8
Path length[µm] = 137.7, 87.8, 22.4, 9.8
Vector sum length [µm] = 34.6, 44.1, 14, 7.2
15
10
5
0
−5
−10
−15
−20
−15
−10
−5
0
5
10

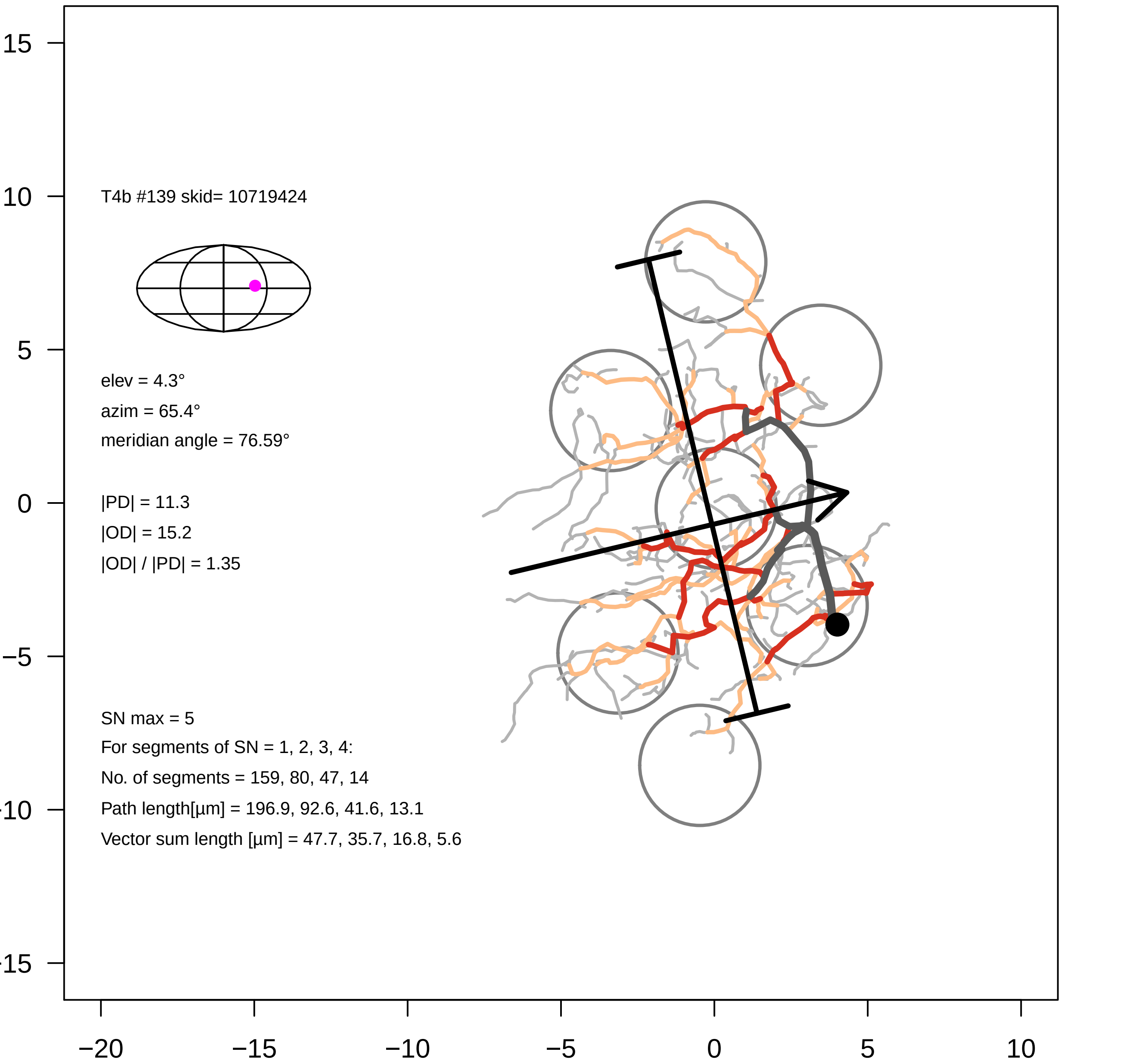

T4b #139 skid= 10719424
elev = 4.3°
azim = 65.4°
meridian angle = 76.59°
|PD| = 11.3
|OD| = 15.2
|OD| / |PD| = 1.35
SN max = 5
For segments of SN = 1, 2, 3, 4:
No. of segments = 159, 80, 47, 14
Path length[µm] = 196.9, 92.6, 41.6, 13.1
Vector sum length [µm] = 47.7, 35.7, 16.8, 5.6
15
10
5
0
−5
−10
−15
−20
−15
−10
−5
0
5
10

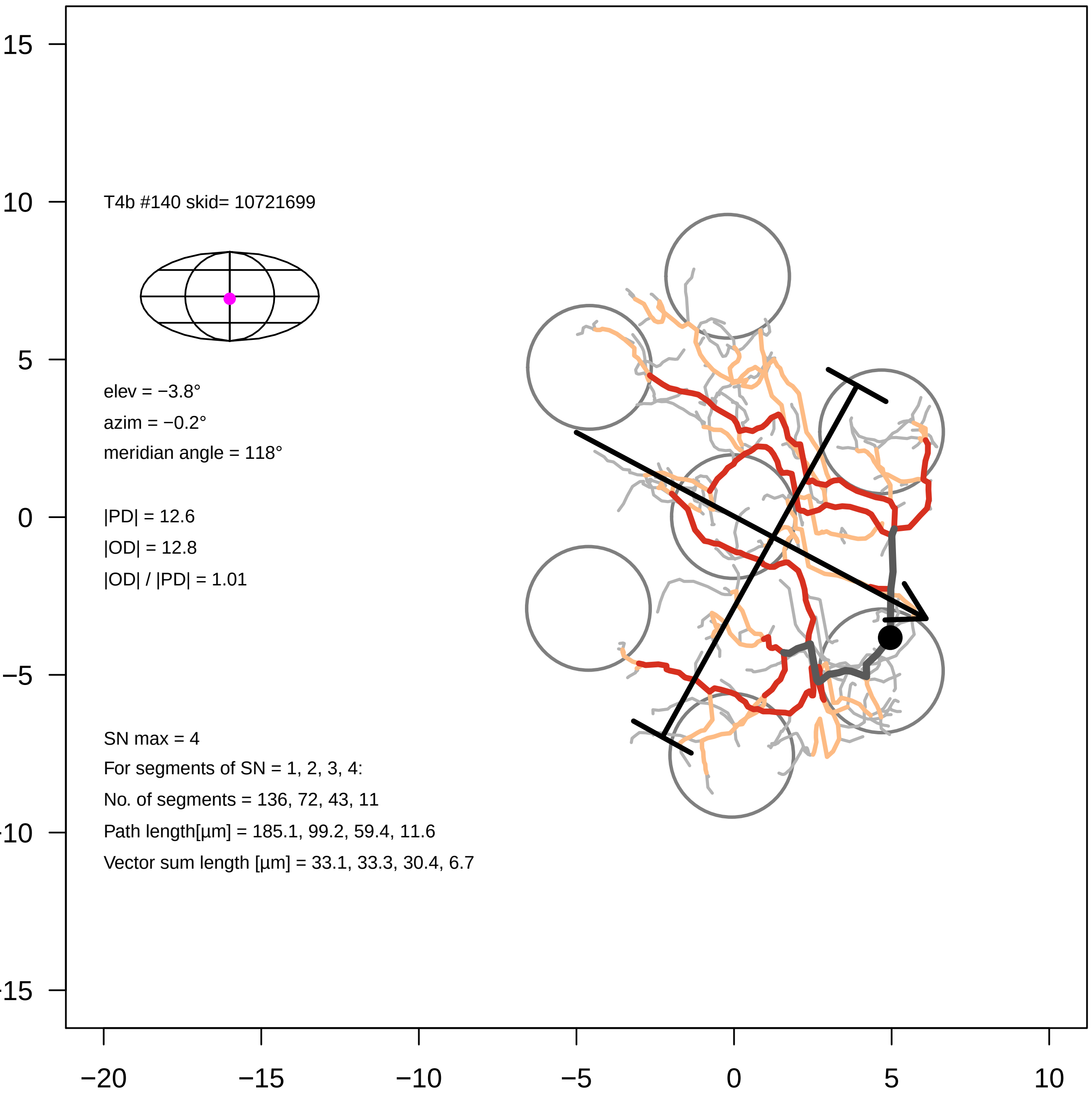
T4b #140 skid= 10721699
elev = −3.8°
azim = −0.2°
meridian angle = 118°
|PD| = 12.6
|OD| = 12.8
|OD| / |PD| = 1.01
SN max = 4
For segments of SN = 1, 2, 3, 4:
No. of segments = 136, 72, 43, 11
Path length[µm] = 185.1, 99.2, 59.4, 11.6
Vector sum length [µm] = 33.1, 33.3, 30.4, 6.7
15
10
5
0
−5
−10
−15
−20
−15
−10
−5
0
5
10

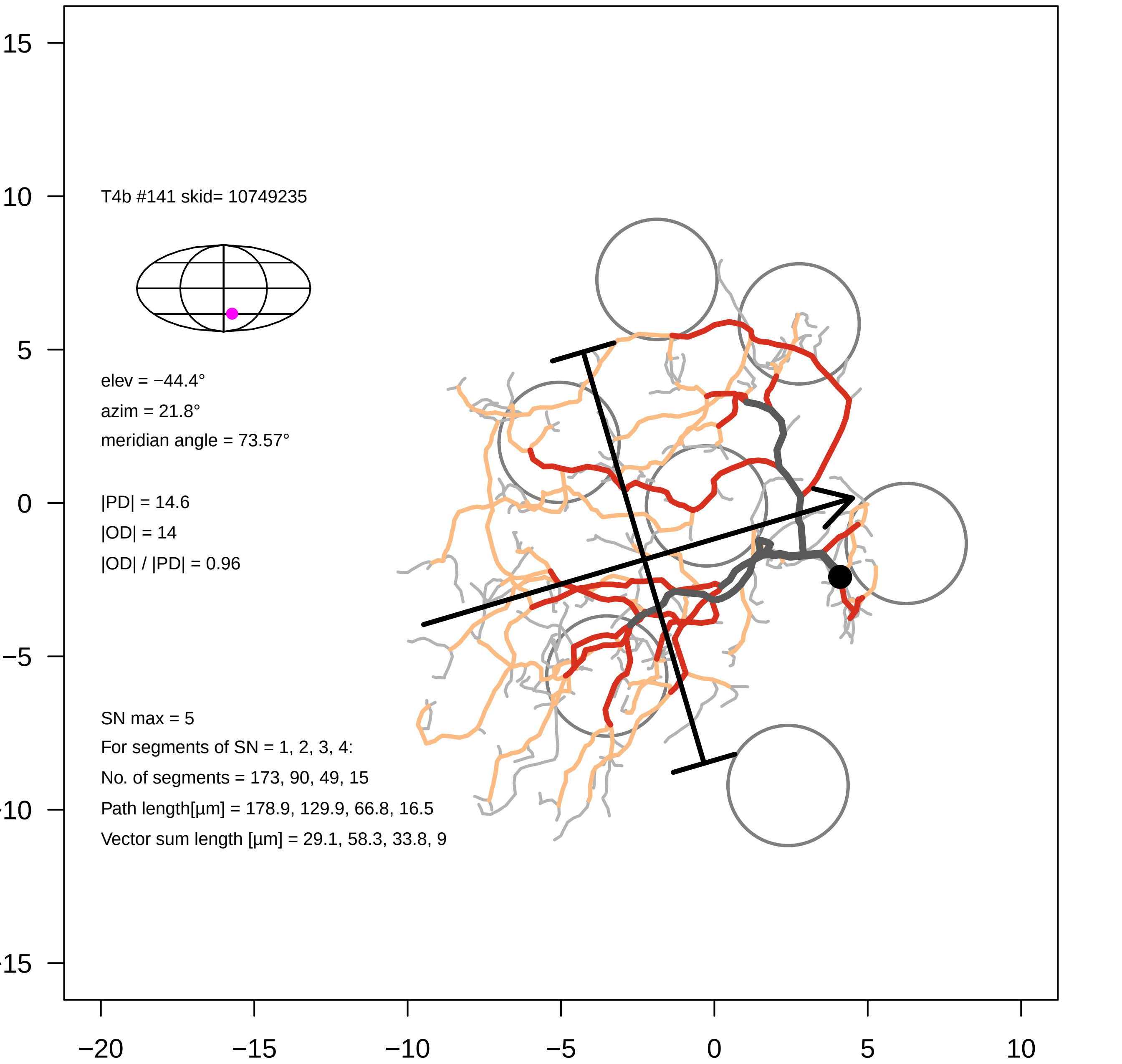
T4b #141 skid= 10749235
elev = −44.4°
azim = 21.8°
meridian angle = 73.57°
|PD| = 14.6
|OD| = 14
|OD| / |PD| = 0.96
SN max = 5
For segments of SN = 1, 2, 3, 4:
No. of segments = 173, 90, 49, 15
Path length[µm] = 178.9, 129.9, 66.8, 16.5
Vector sum length [µm] = 29.1, 58.3, 33.8, 9
15
10
5
0
−5
−10
−15
−20
−15
−10
−5
0
5
10

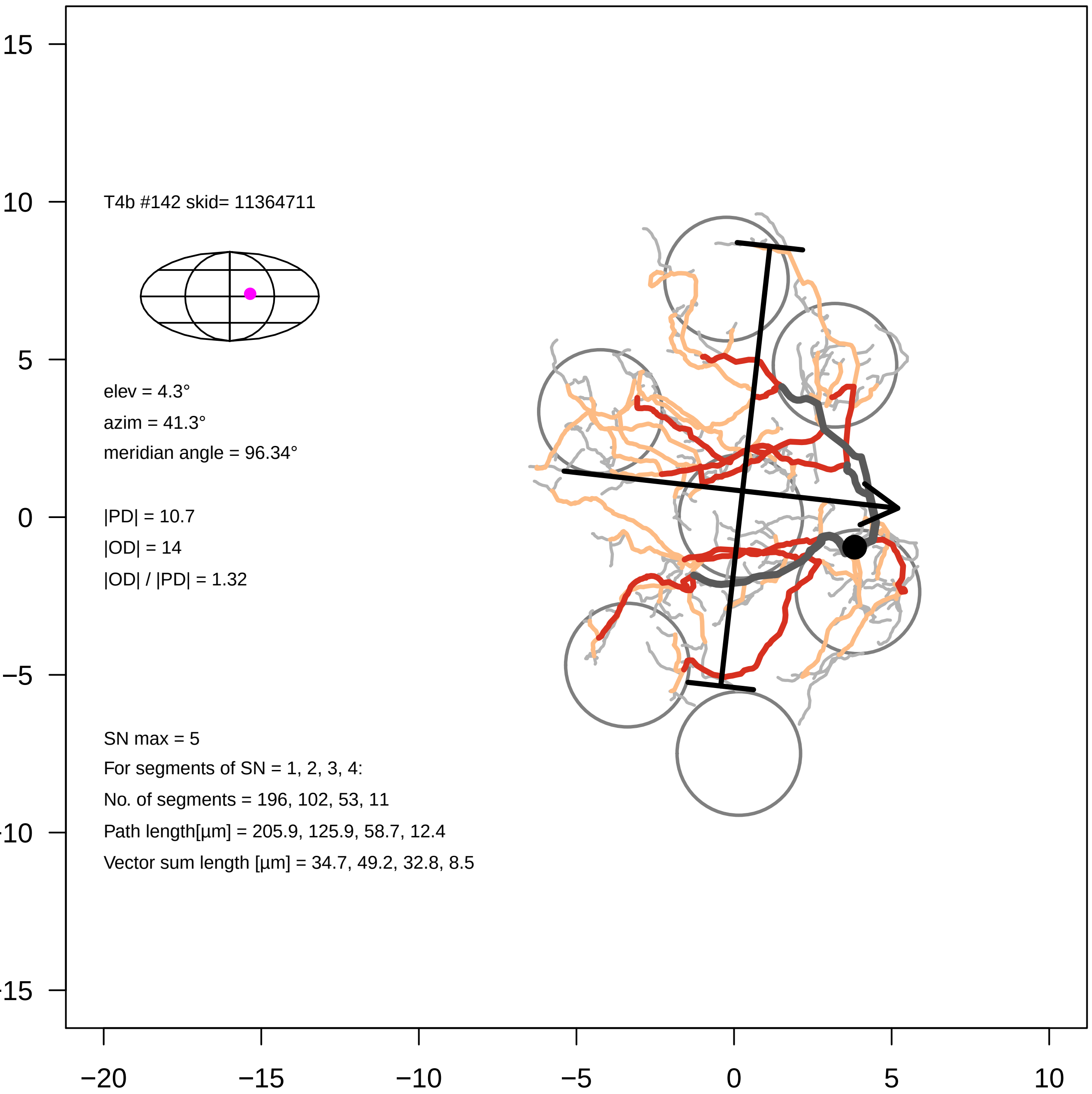

T4b #142 skid= 11364711
elev = 4.3°
azim = 41.3°
meridian angle = 96.34°
|PD| = 10.7
|OD| = 14
|OD| / |PD| = 1.32
SN max = 5
For segments of SN = 1, 2, 3, 4:
No. of segments = 196, 102, 53, 11
Path length[µm] = 205.9, 125.9, 58.7, 12.4
Vector sum length [µm] = 34.7, 49.2, 32.8, 8.5
15
10
5
0
-5
-10
-15
−20
−15
−10
−5
0
5
10

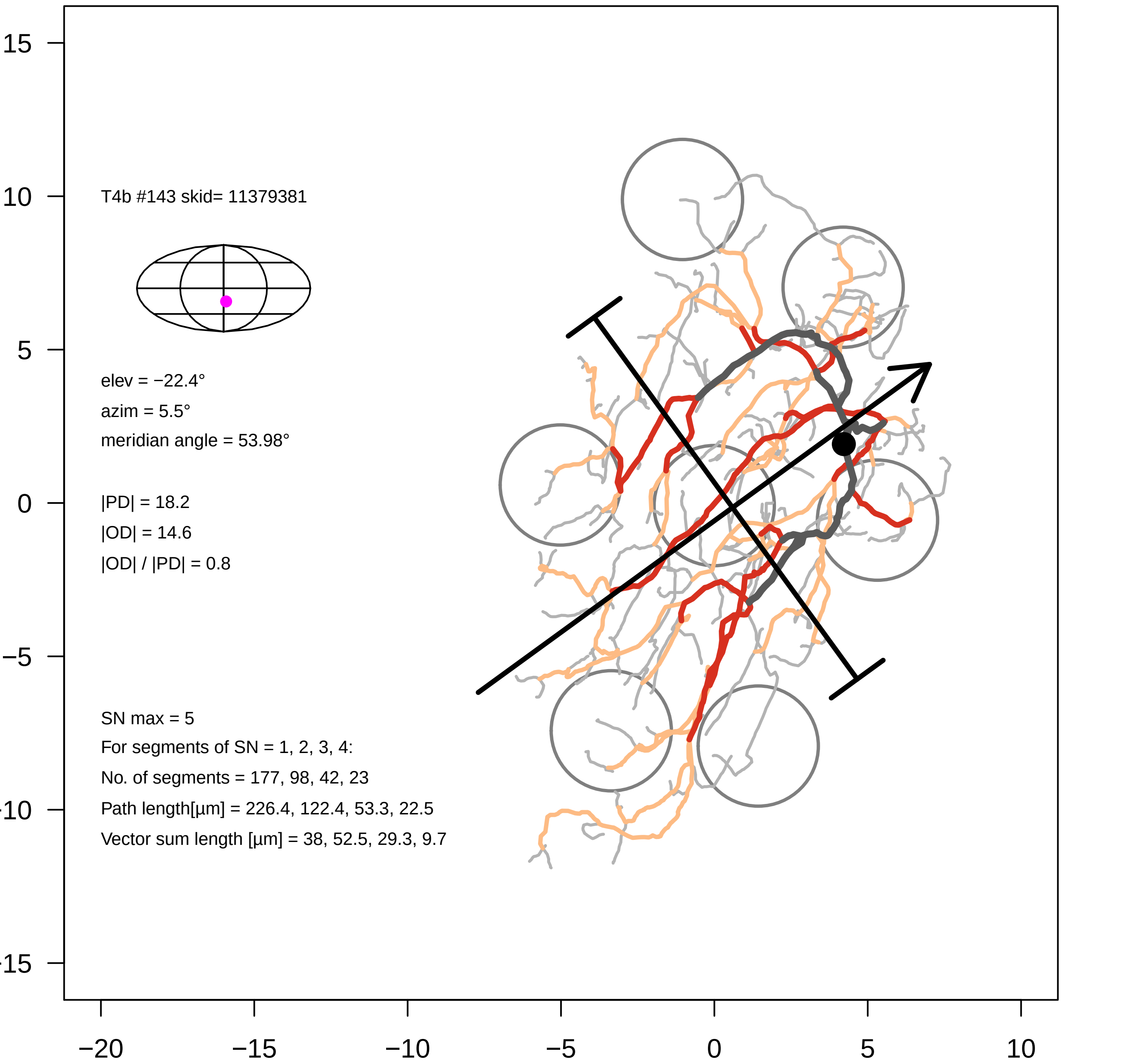

T4b #143 skid= 11379381
elev = −22.4°
azim = 5.5°
meridian angle = 53.98°
|PD| = 18.2
|OD| = 14.6
|OD| / |PD| = 0.8
SN max = 5
For segments of SN = 1, 2, 3, 4:
No. of segments = 177, 98, 42, 23
Path length[µm] = 226.4, 122.4, 53.3, 22.5
Vector sum length [µm] = 38, 52.5, 29.3, 9.7
15
10
5
0
−5
−20
−15
−10
−5
0
5
10

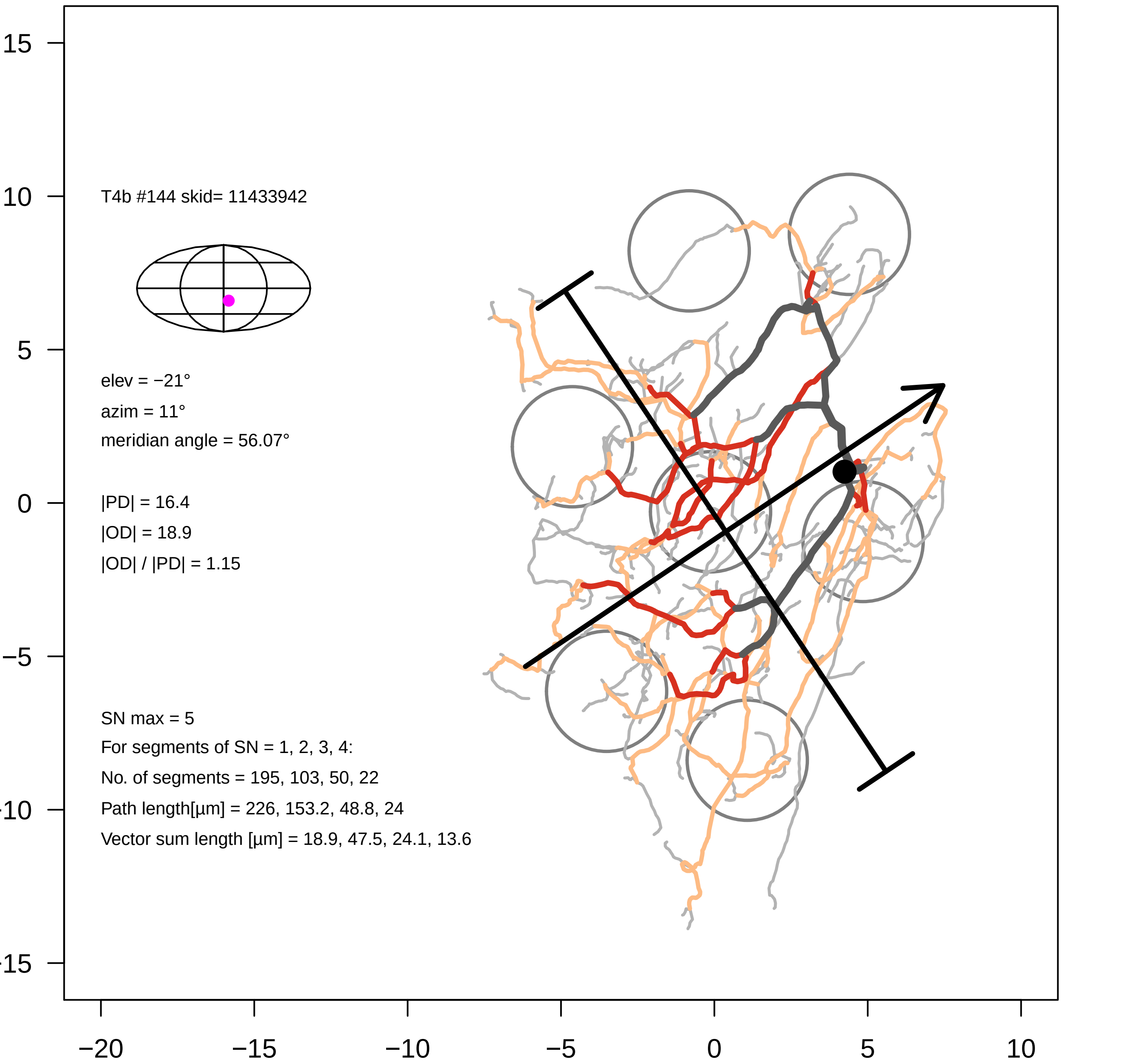

T4b #144 skid= 11433942
elev = −21°
azim = 11°
meridian angle = 56.07°
|PD| = 16.4
|OD| = 18.9
|OD| / |PD| = 1.15
SN max = 5
For segments of SN = 1, 2, 3, 4:
No. of segments = 195, 103, 50, 22
Path length[µm] = 226, 153.2, 48.8, 24
Vector sum length [µm] = 18.9, 47.5, 24.1, 13.6
15
10
5
0
−5
−10
−15
−20
−15
−10
−5
0
5
10

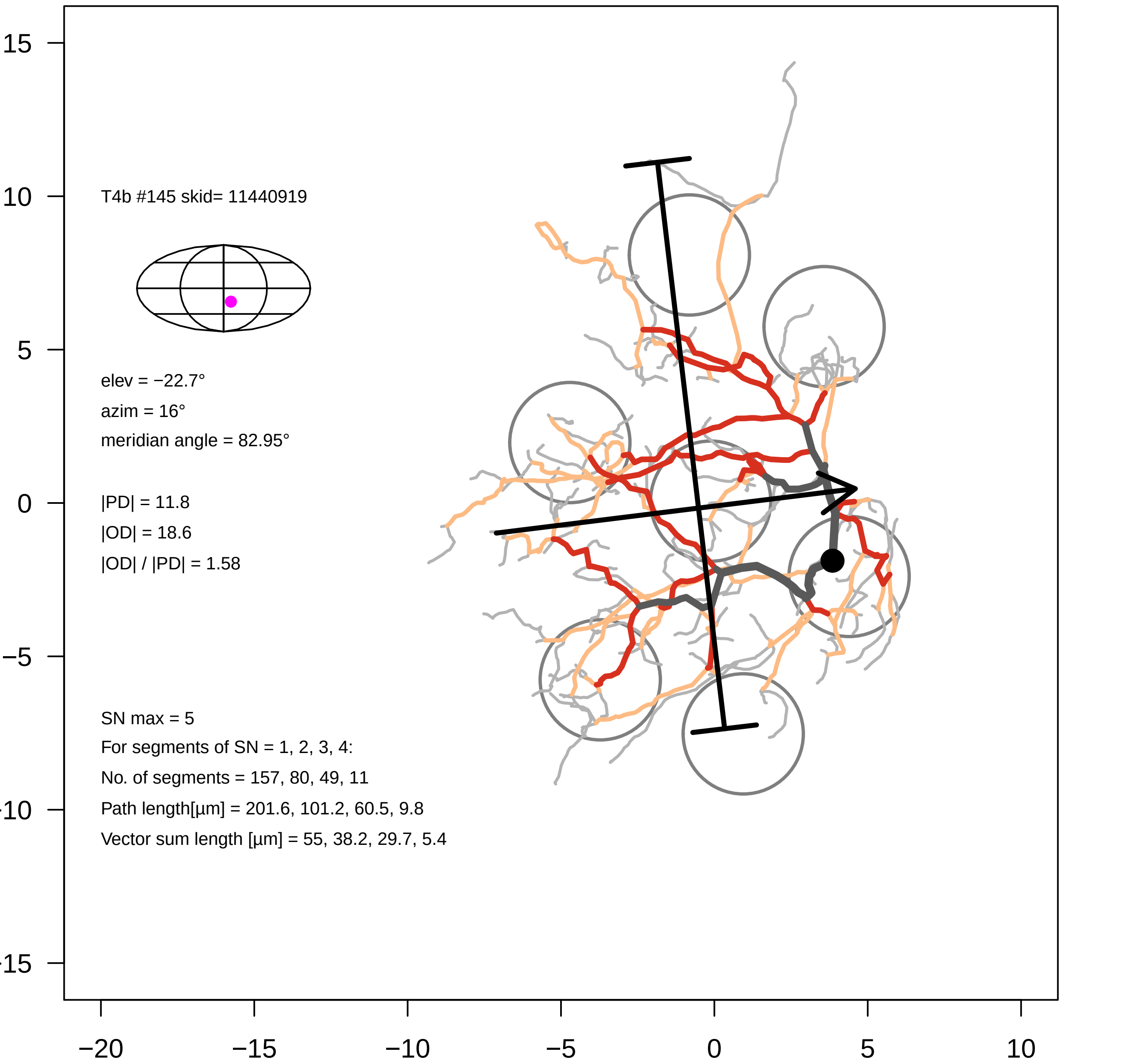

T4b #145 skid= 11440919
elev = −22.7°
azim = 16°
meridian angle = 82.95°
|PD| = 11.8
|OD| = 18.6
|OD| / |PD| = 1.58
SN max = 5
For segments of SN = 1, 2, 3, 4:
No. of segments = 157, 80, 49, 11
Path length[µm] = 201.6, 101.2, 60.5, 9.8
Vector sum length [µm] = 55, 38.2, 29.7, 5.4

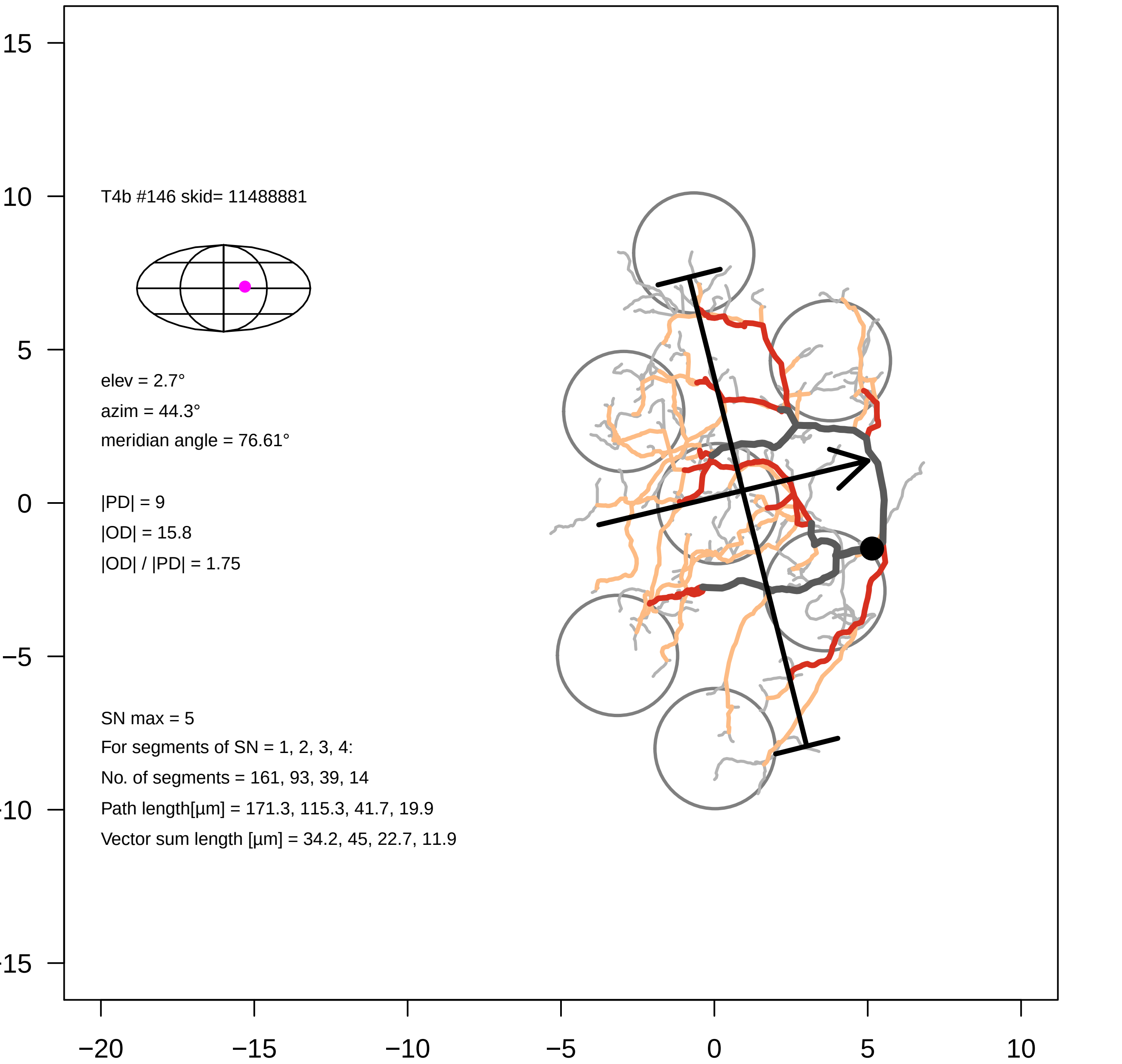

T4b #146 skid= 11488881
elev = 2.7°
azim = 44.3°
meridian angle = 76.61°
|PD| = 9
|OD| = 15.8
|OD| / |PD| = 1.75
SN max = 5
For segments of SN = 1, 2, 3, 4:
No. of segments = 161, 93, 39, 14
Path length[µm] = 171.3, 115.3, 41.7, 19.9
Vector sum length [µm] = 34.2, 45, 22.7, 11.9
15
10
5
0
−5
−10
−15
−20
−15
−10
−5
0
5
10

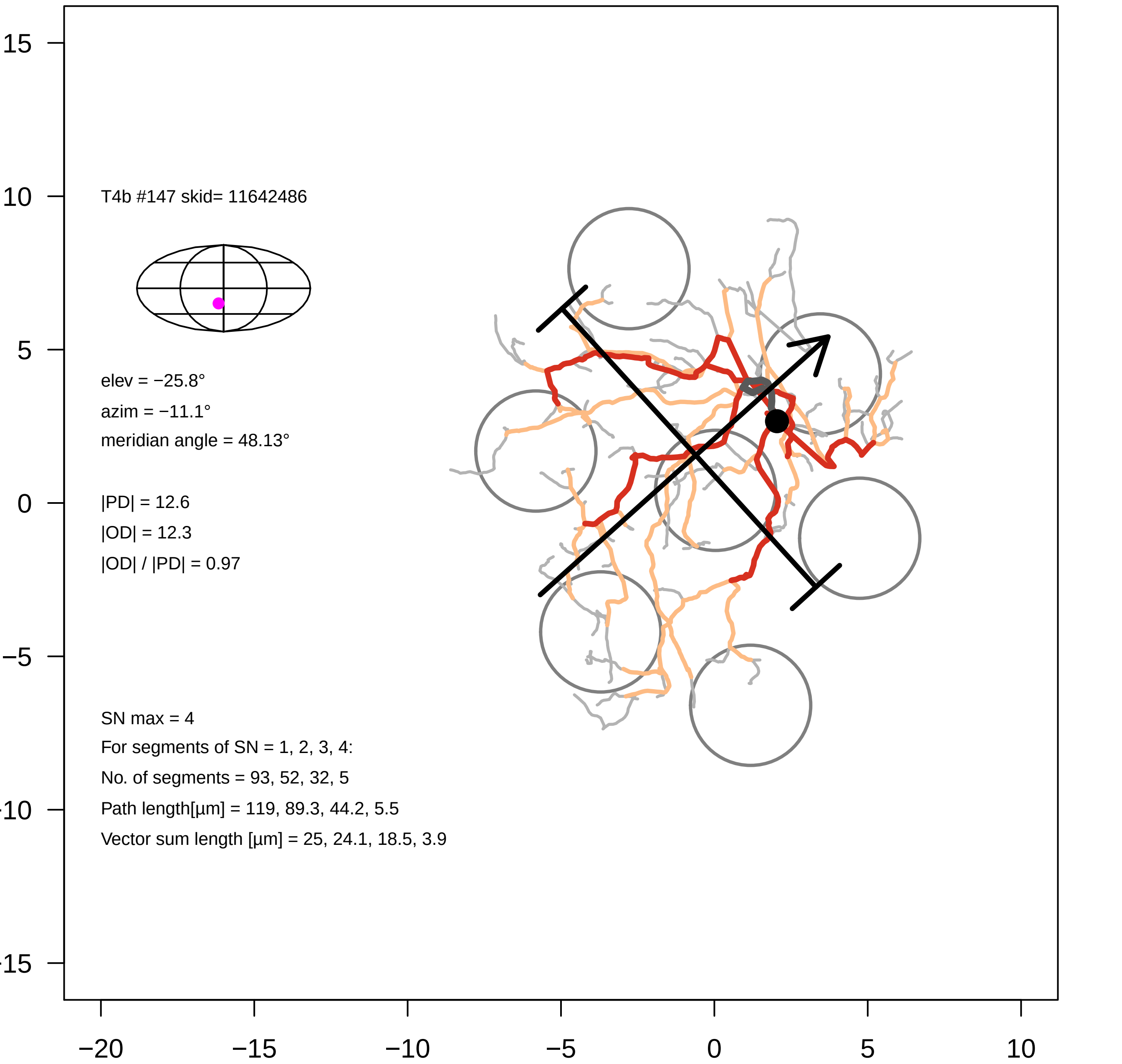
T4b #147 skid= 11642486
elev = −25.8°
azim = −11.1°
meridian angle = 48.13°
|PD| = 12.6
|OD| = 12.3
|OD| / |PD| = 0.97
SN max = 4
For segments of SN = 1, 2, 3, 4:
No. of segments = 93, 52, 32, 5
Path length[µm] = 119, 89.3, 44.2, 5.5
Vector sum length [µm] = 25, 24.1, 18.5, 3.9
15
10
5
0
−5
−10
−15
−20
−15
−10
−5
0
5
10

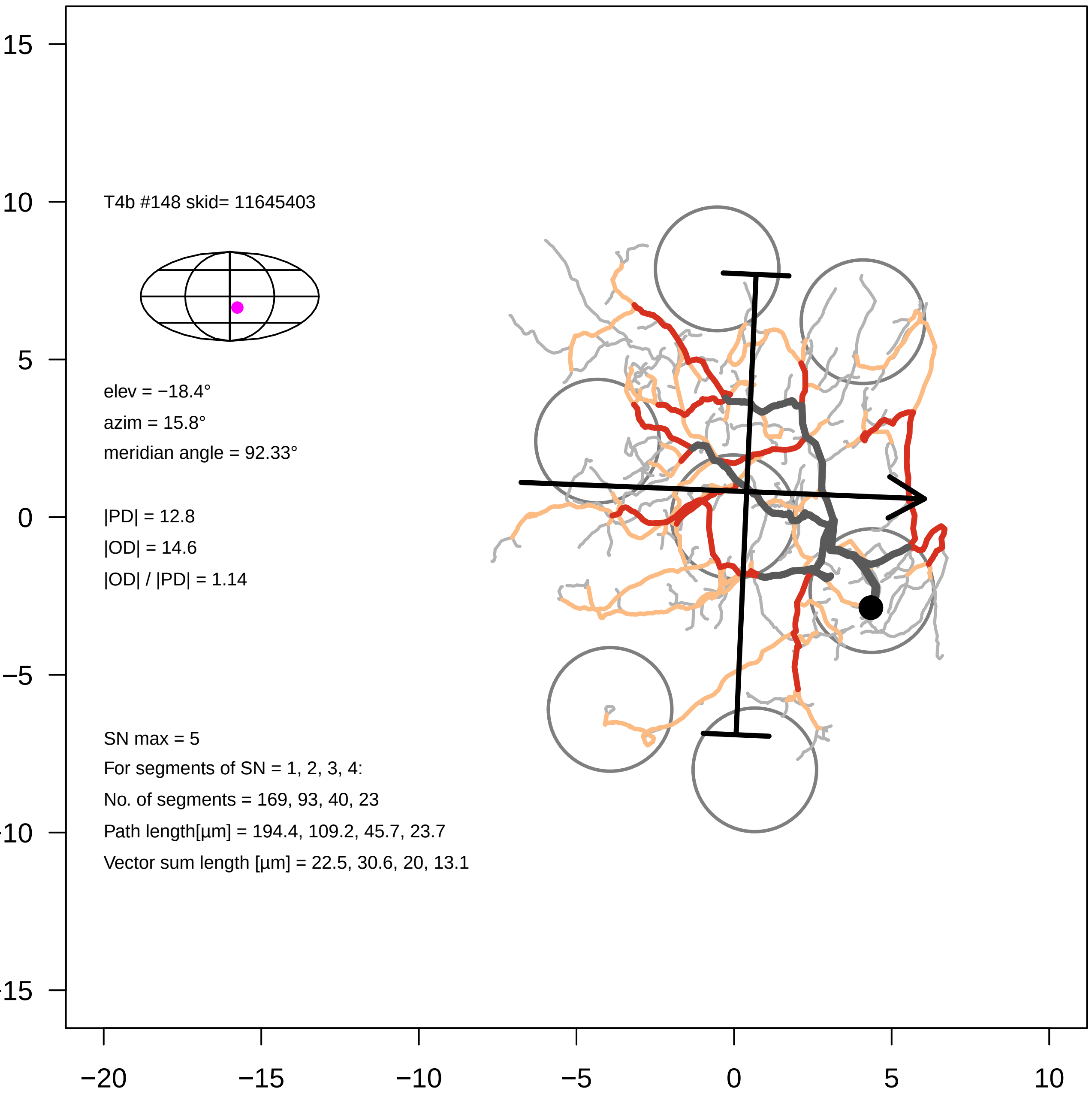
T4b #148 skid= 11645403
elev = −18.4°
azim = 15.8°
meridian angle = 92.33°
|PD| = 12.8
|OD| = 14.6
|OD| / |PD| = 1.14
SN max = 5
For segments of SN = 1, 2, 3, 4:
No. of segments = 169, 93, 40, 23
Path length[µm] = 194.4, 109.2, 45.7, 23.7
Vector sum length [µm] = 22.5, 30.6, 20, 13.1
15
10
5
0
−5
−10
−15
−20
−15
−10
−5
0
5
10

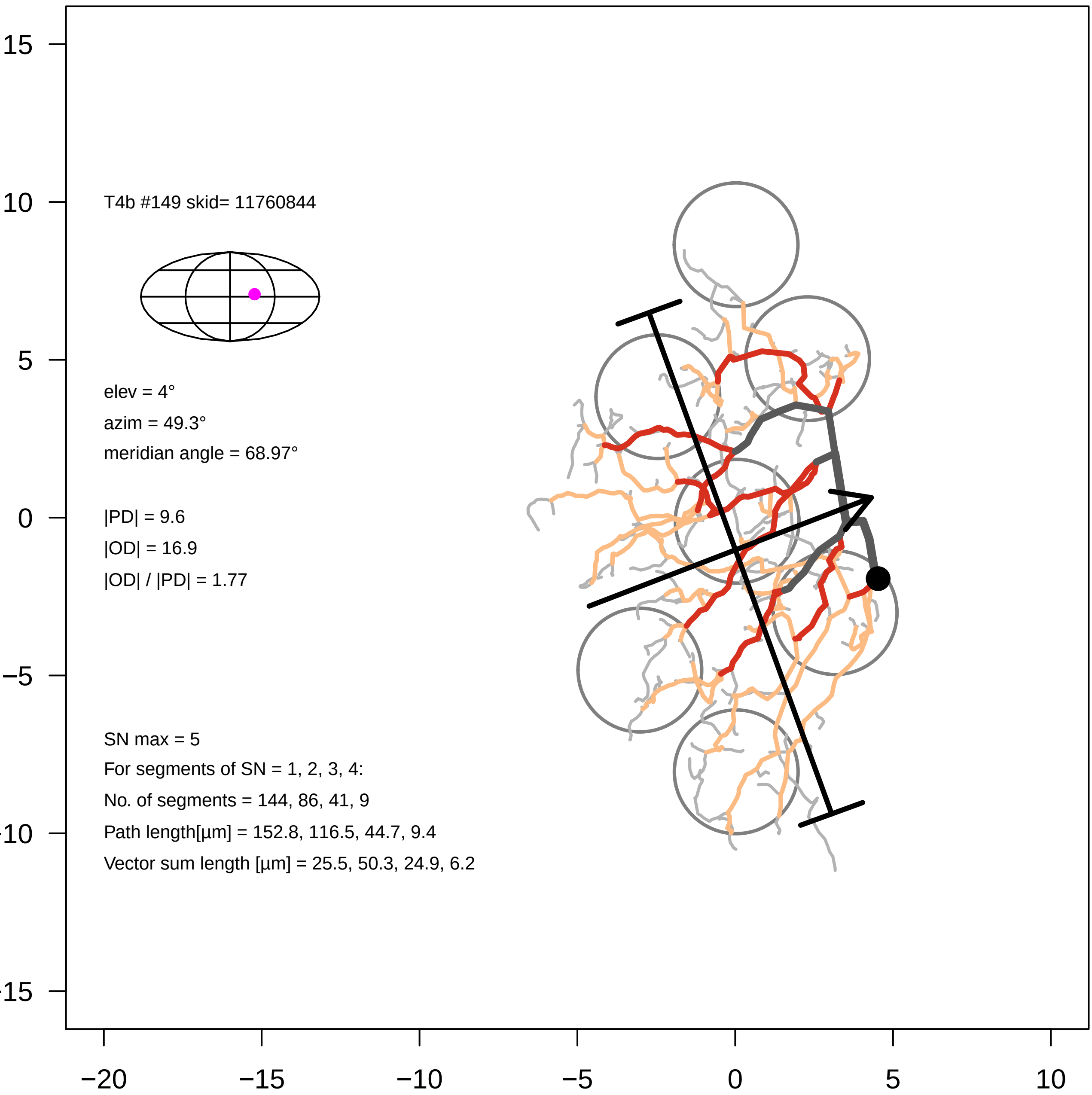

T4b #149 skid= 11760844
elev = 4°
azim = 49.3°
meridian angle = 68.97°
|PD| = 9.6
|OD| = 16.9
|OD| / |PD| = 1.77
SN max = 5
For segments of SN = 1, 2, 3, 4:
No. of segments = 144, 86, 41, 9
Path length[µm] = 152.8, 116.5, 44.7, 9.4
Vector sum length [µm] = 25.5, 50.3, 24.9, 6.2
15
10
5
0
−5
−10
−15
−20
−15
−10
−5
0
5
10

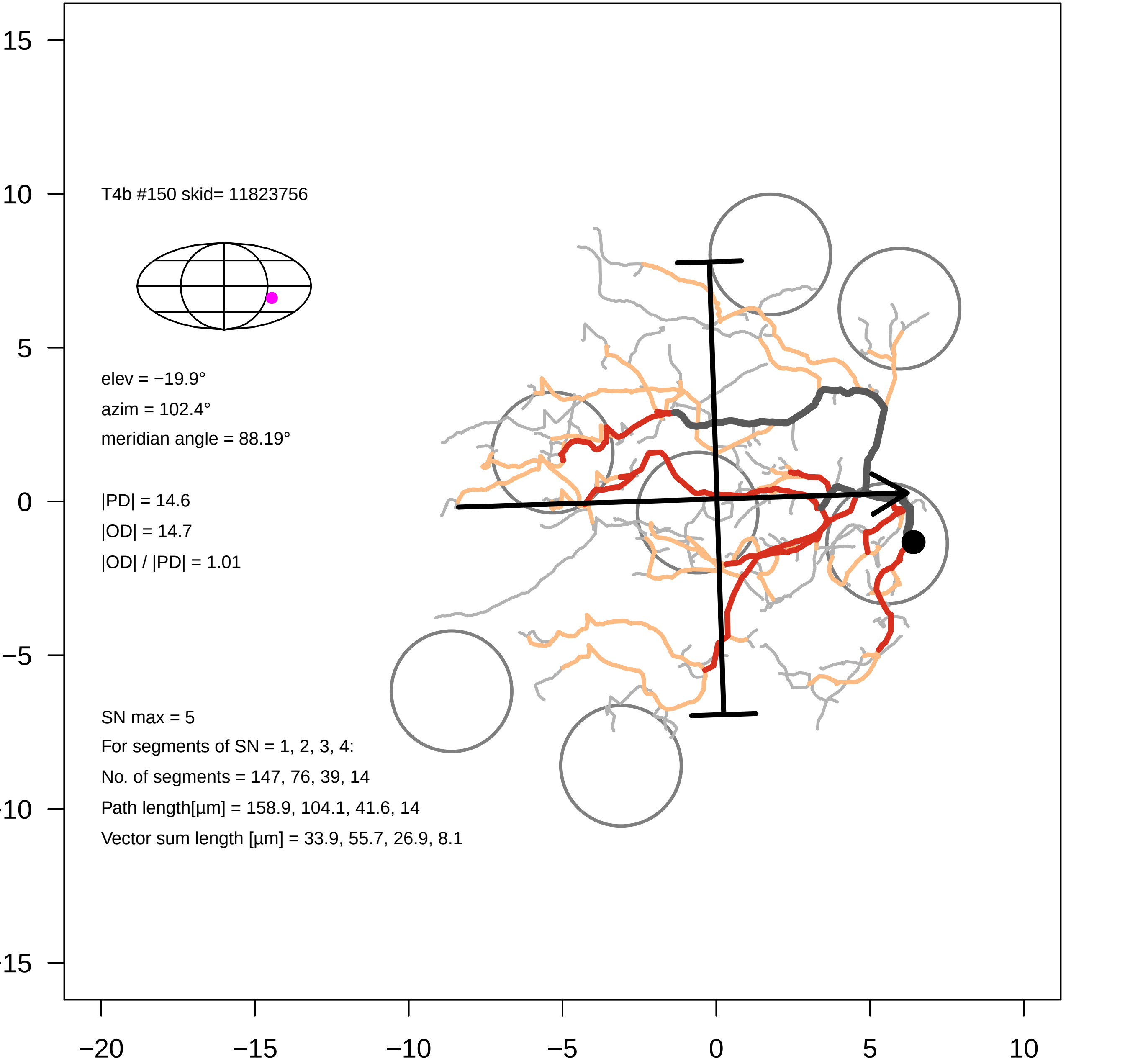
T4b #150 skid= 11823756
elev = −19.9°
azim = 102.4°
meridian angle = 88.19°
|PD| = 14.6
|OD| = 14.7
|OD| / |PD| = 1.01
SN max = 5
For segments of SN = 1, 2, 3, 4:
No. of segments = 147, 76, 39, 14
Path length[µm] = 158.9, 104.1, 41.6, 14
Vector sum length [µm] = 33.9, 55.7, 26.9, 8.1
15
10
5
0
−5
−10
−15
−20
−15
−10
−5
0
5
10

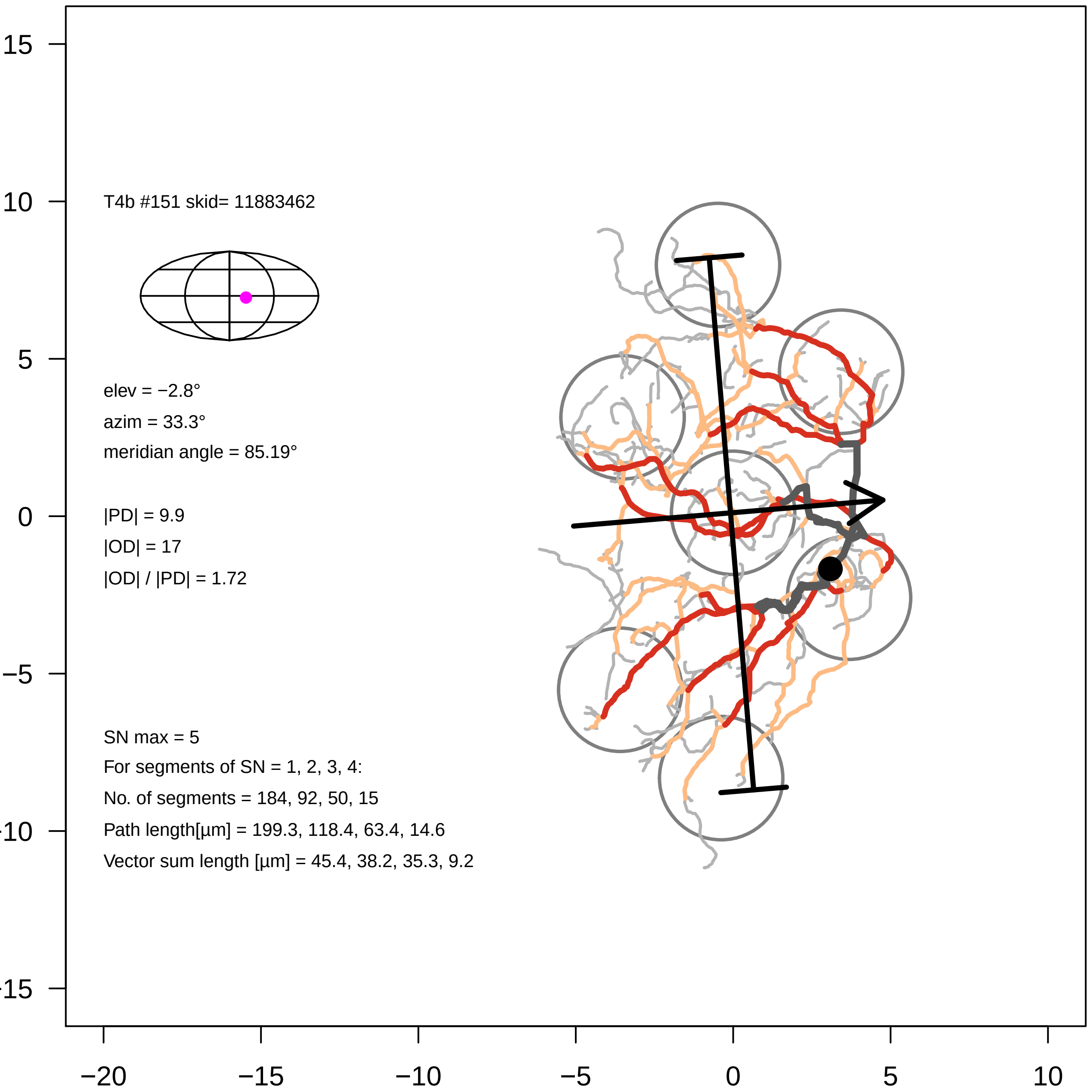
T4b #151 skid= 11883462
elev = −2.8°
azim = 33.3°
meridian angle = 85.19°
|PD| = 9.9
|OD| = 17
|OD| / |PD| = 1.72
SN max = 5
For segments of SN = 1, 2, 3, 4:
No. of segments = 184, 92, 50, 15
Path length[µm] = 199.3, 118.4, 63.4, 14.6
Vector sum length [µm] = 45.4, 38.2, 35.3, 9.2
15
10
5
0
−5
−10
−15
−20
−15
−10
−5
0
5
10

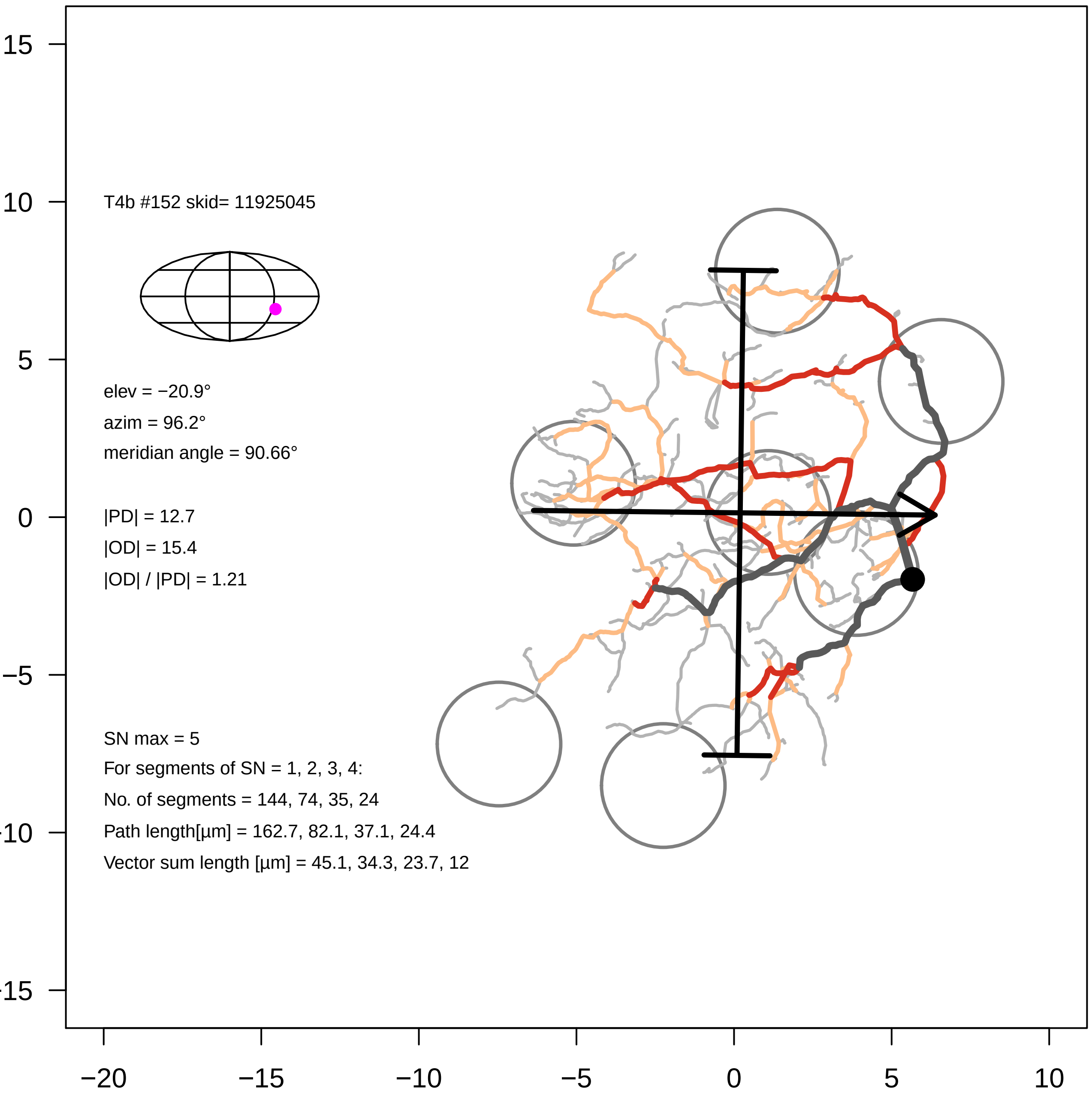
T4b #152 skid= 11925045
elev = −20.9°
azim = 96.2°
meridian angle = 90.66°
|PD| = 12.7
|OD| = 15.4
|OD| / |PD| = 1.21
SN max = 5
For segments of SN = 1, 2, 3, 4:
No. of segments = 144, 74, 35, 24
Path length[µm] = 162.7, 82.1, 37.1, 24.4
Vector sum length [µm] = 45.1, 34.3, 23.7, 12
−20
−15
−10
−5
0
5
10
15
10
5
0
−5

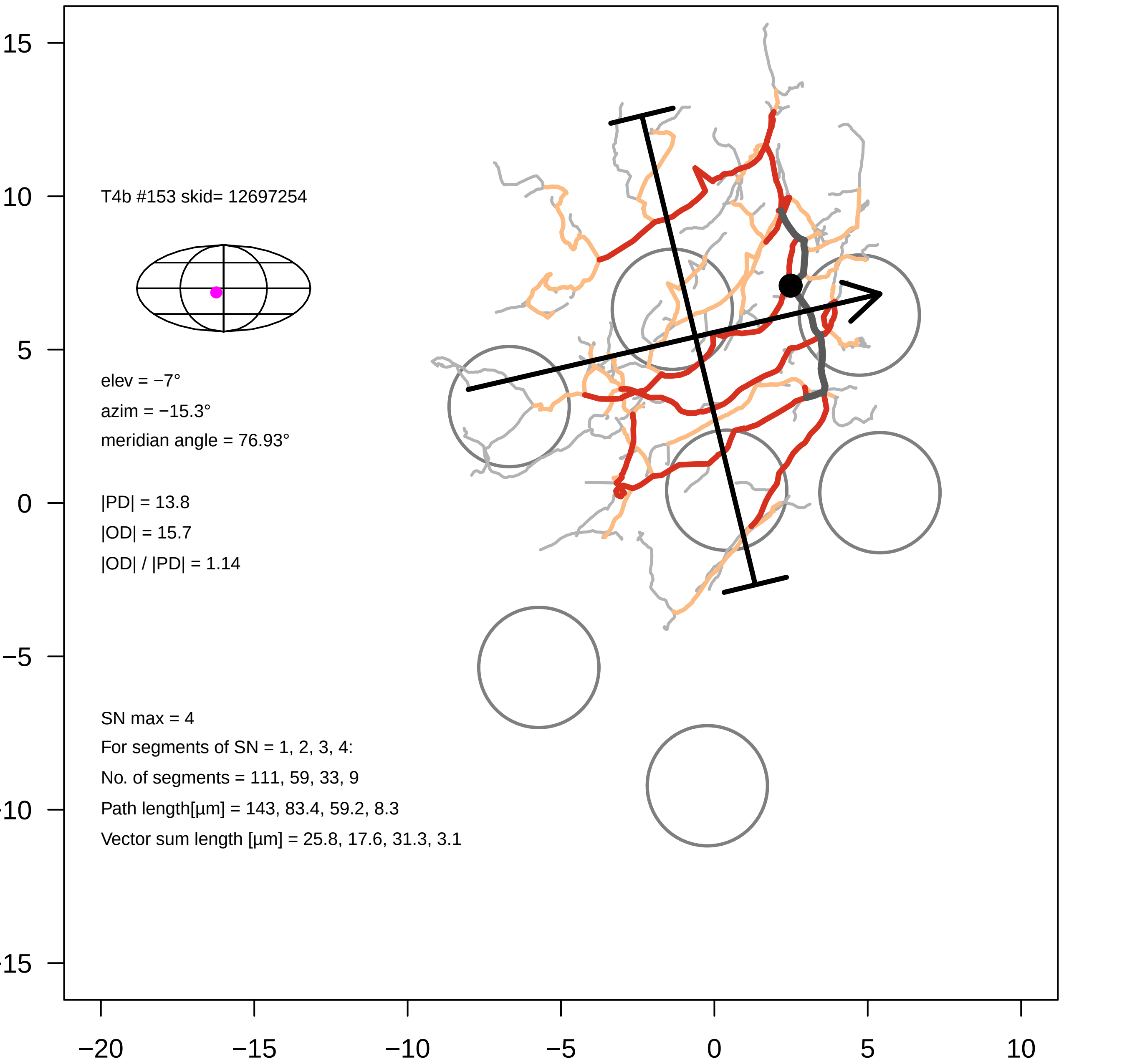

T4b #153 skid= 12697254
elev = −7°
azim = −15.3°
meridian angle = 76.93°
|PD| = 13.8
|OD| = 15.7
|OD| / |PD| = 1.14
SN max = 4
For segments of SN = 1, 2, 3, 4:
No. of segments = 111, 59, 33, 9
Path length[µm] = 143, 83.4, 59.2, 8.3
Vector sum length [µm] = 25.8, 17.6, 31.3, 3.1
15
10
5
0
−5
−10
−15
−20
−15
−10
−5
0
5
10

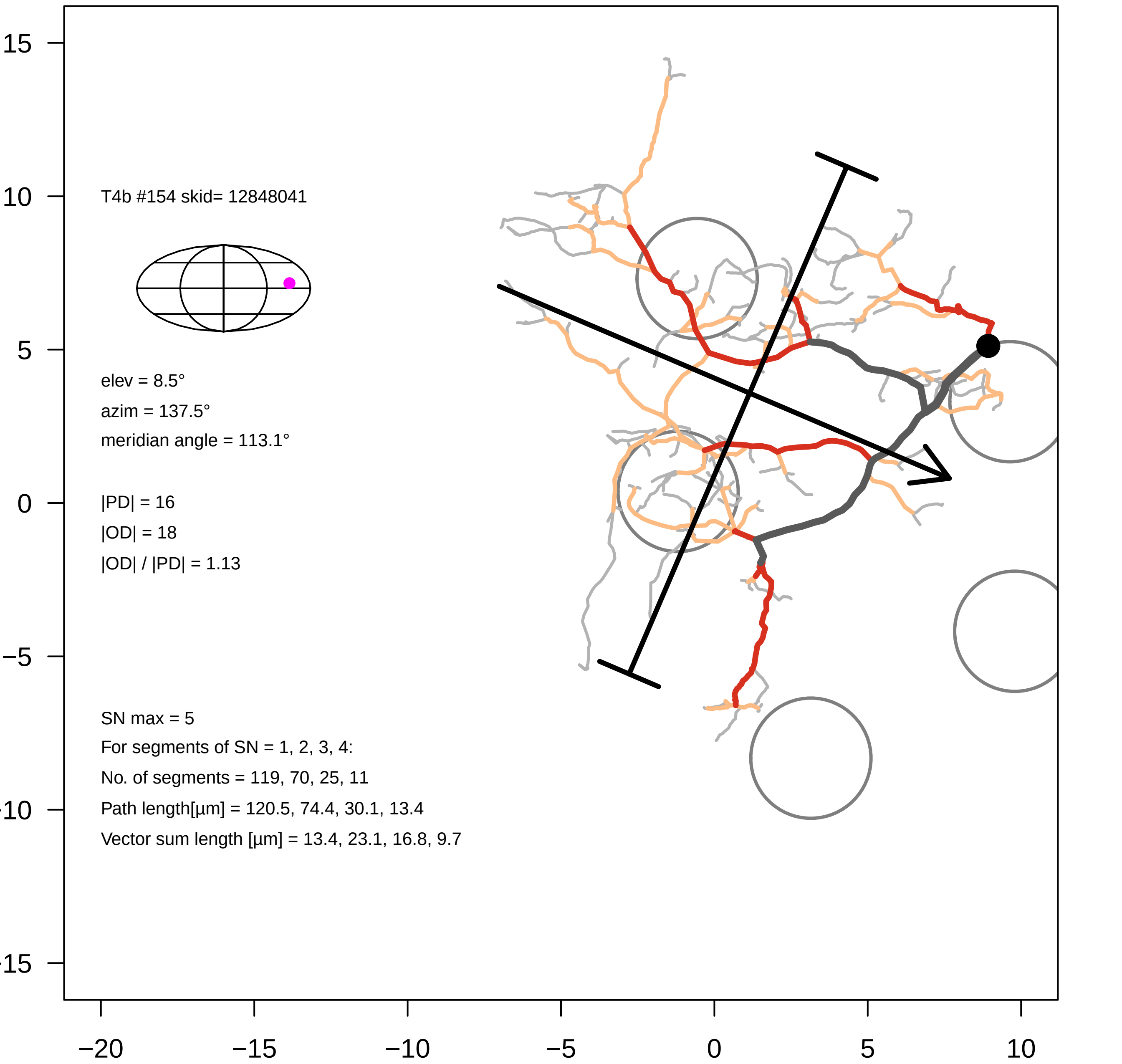

15
10
5
0
−5
−10
−15
−20
−15
−10
−5
0
5
10
T4b #154 skid= 12848041
elev = 8.5°
azim = 137.5°
meridian angle = 113.1°
|PD| = 16
|OD| = 18
|OD| / |PD| = 1.13
SN max = 5
For segments of SN = 1, 2, 3, 4:
No. of segments = 119, 70, 25, 11
Path length[µm] = 120.5, 74.4, 30.1, 13.4
Vector sum length [µm] = 13.4, 23.1, 16.8, 9.7

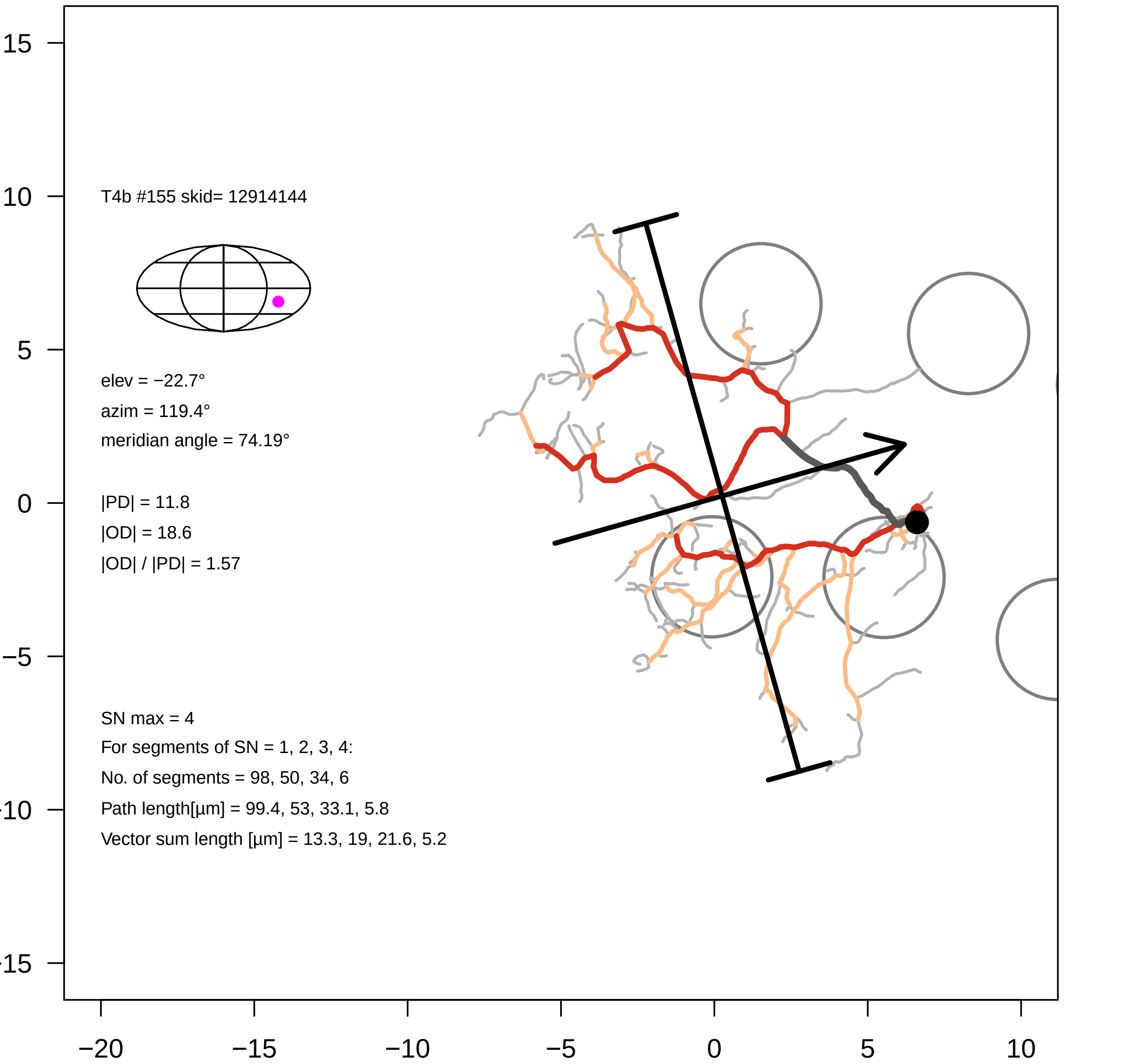
T4b #155 skid= 12914144
elev = −22.7°
azim = 119.4°
meridian angle = 74.19°
|PD| = 11.8
|OD| = 18.6
|OD| / |PD| = 1.57
SN max = 4
For segments of SN = 1, 2, 3, 4:
No. of segments = 98, 50, 34, 6
Path length[µm] = 99.4, 53, 33.1, 5.8
Vector sum length [µm] = 13.3, 19, 21.6, 5.2
15
10
5
0
−5
−10
−15
−20
−15
−10
−5
0
5
10

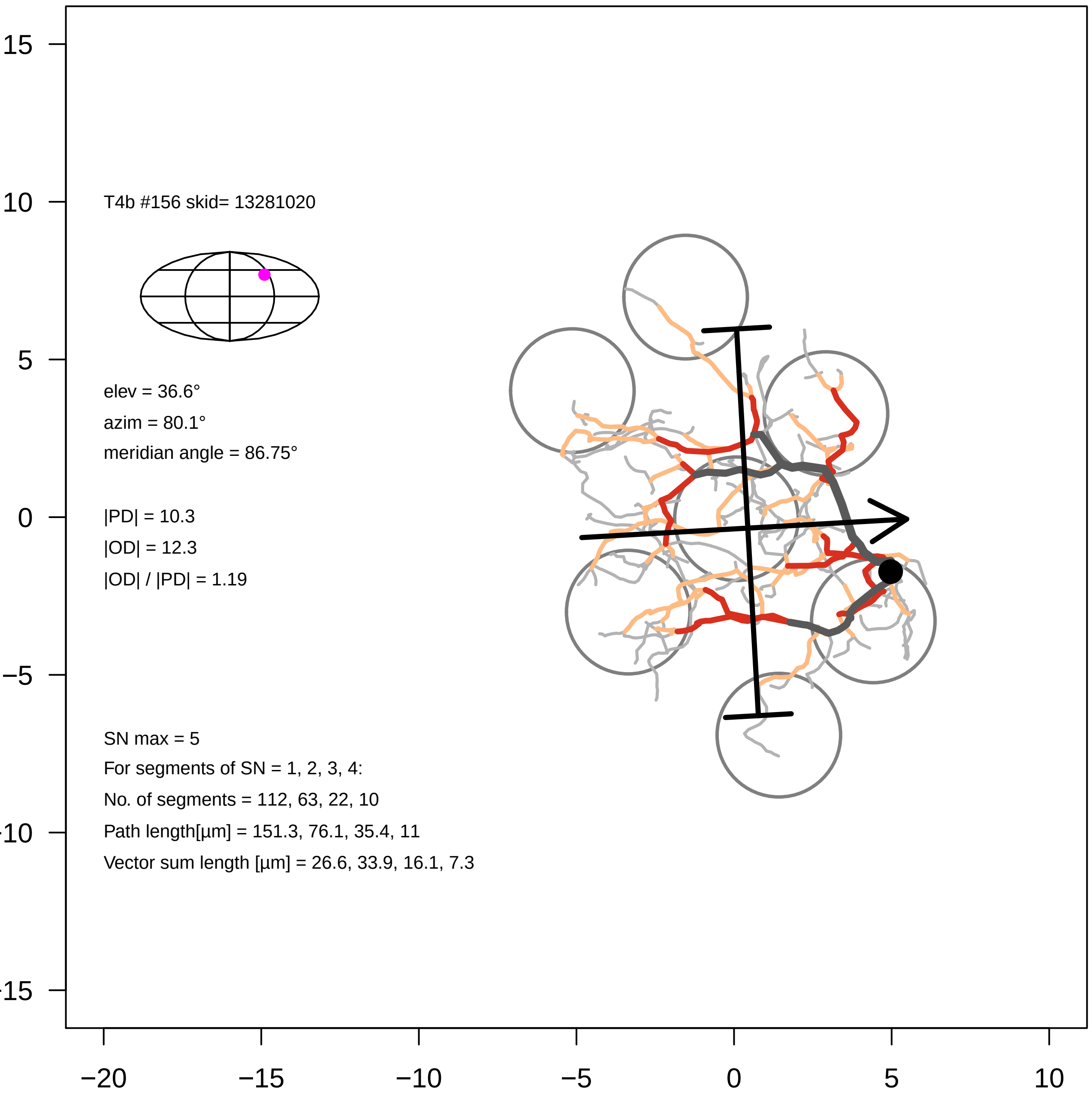

T4b #156 skid= 13281020
elev = 36.6°
azim = 80.1°
meridian angle = 86.75°
|PD| = 10.3
|OD| = 12.3
|OD| / |PD| = 1.19
SN max = 5
For segments of SN = 1, 2, 3, 4:
No. of segments = 112, 63, 22, 10
Path length[µm] = 151.3, 76.1, 35.4, 11
Vector sum length [µm] = 26.6, 33.9, 16.1, 7.3
15
10
5
0
−5
−10
−15
−20
−15
−10
−5
0
5
10

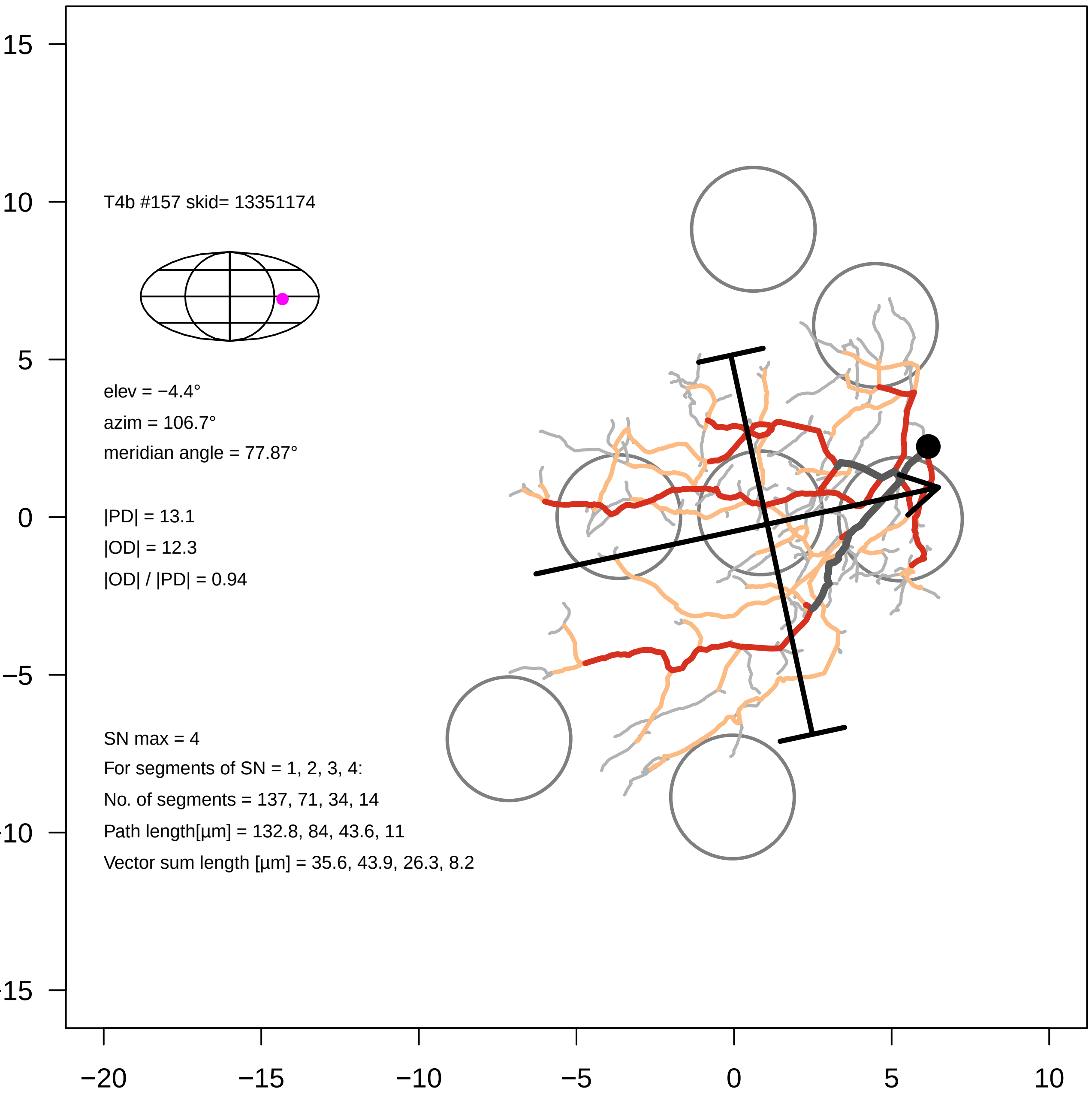
T4b #157 skid= 13351174
elev = −4.4°
azim = 106.7°
meridian angle = 77.87°
|PD| = 13.1
|OD| = 12.3
|OD| / |PD| = 0.94
SN max = 4
For segments of SN = 1, 2, 3, 4:
No. of segments = 137, 71, 34, 14
Path length[µm] = 132.8, 84, 43.6, 11
Vector sum length [µm] = 35.6, 43.9, 26.3, 8.2
15
10
5
0
−5
−10
−15
−20
−15
−10
−5
0
5
10

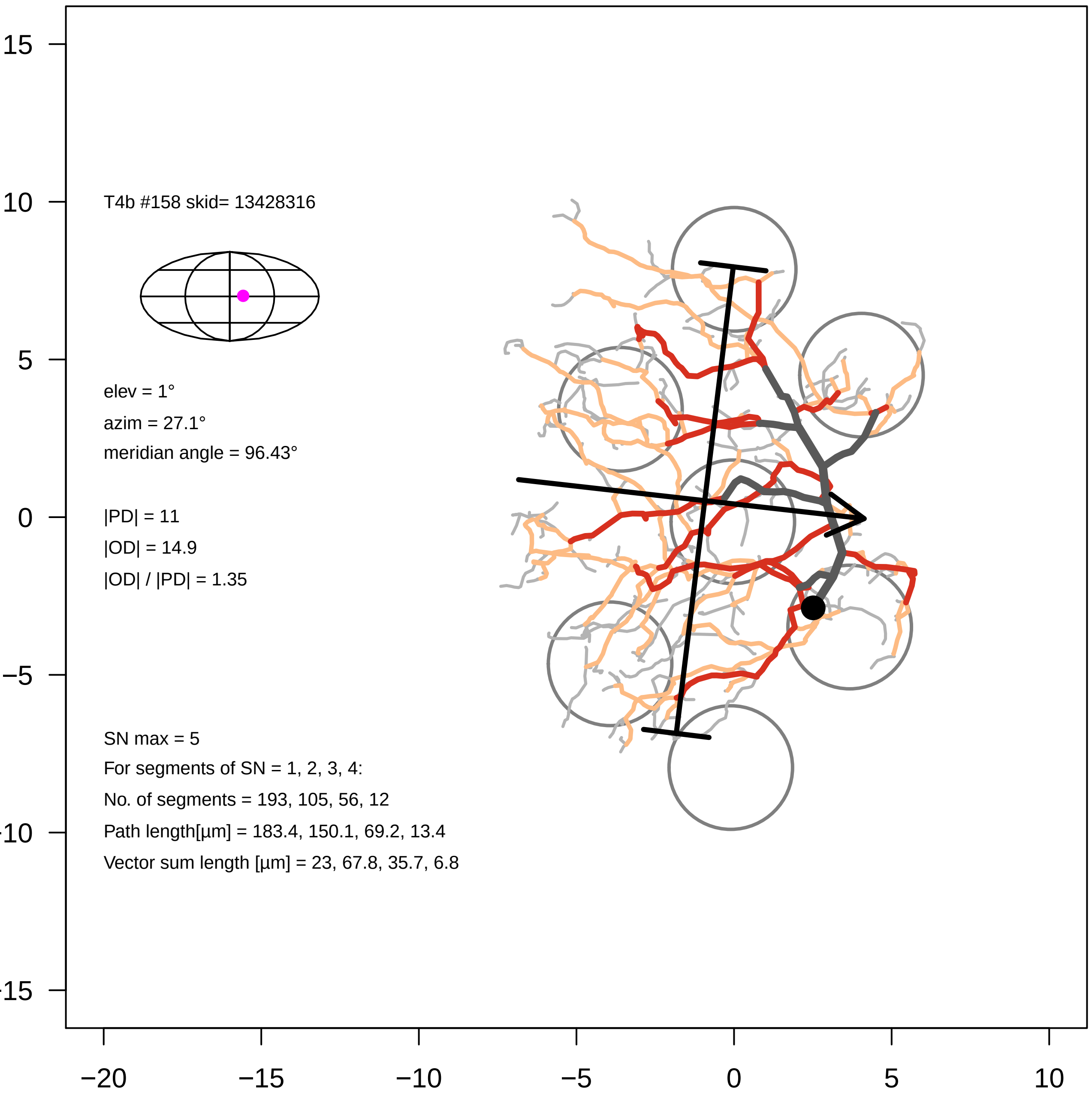

T4b #158 skid= 13428316
elev = 1°
azim = 27.1°
meridian angle = 96.43°
|PD| = 11
|OD| = 14.9
|OD| / |PD| = 1.35
SN max = 5
For segments of SN = 1, 2, 3, 4:
No. of segments = 193, 105, 56, 12
Path length[µm] = 183.4, 150.1, 69.2, 13.4
Vector sum length [µm] = 23, 67.8, 35.7, 6.8
15
10
5
0
−5
−10
−15
−20
−15
−10
−5
0
5
10

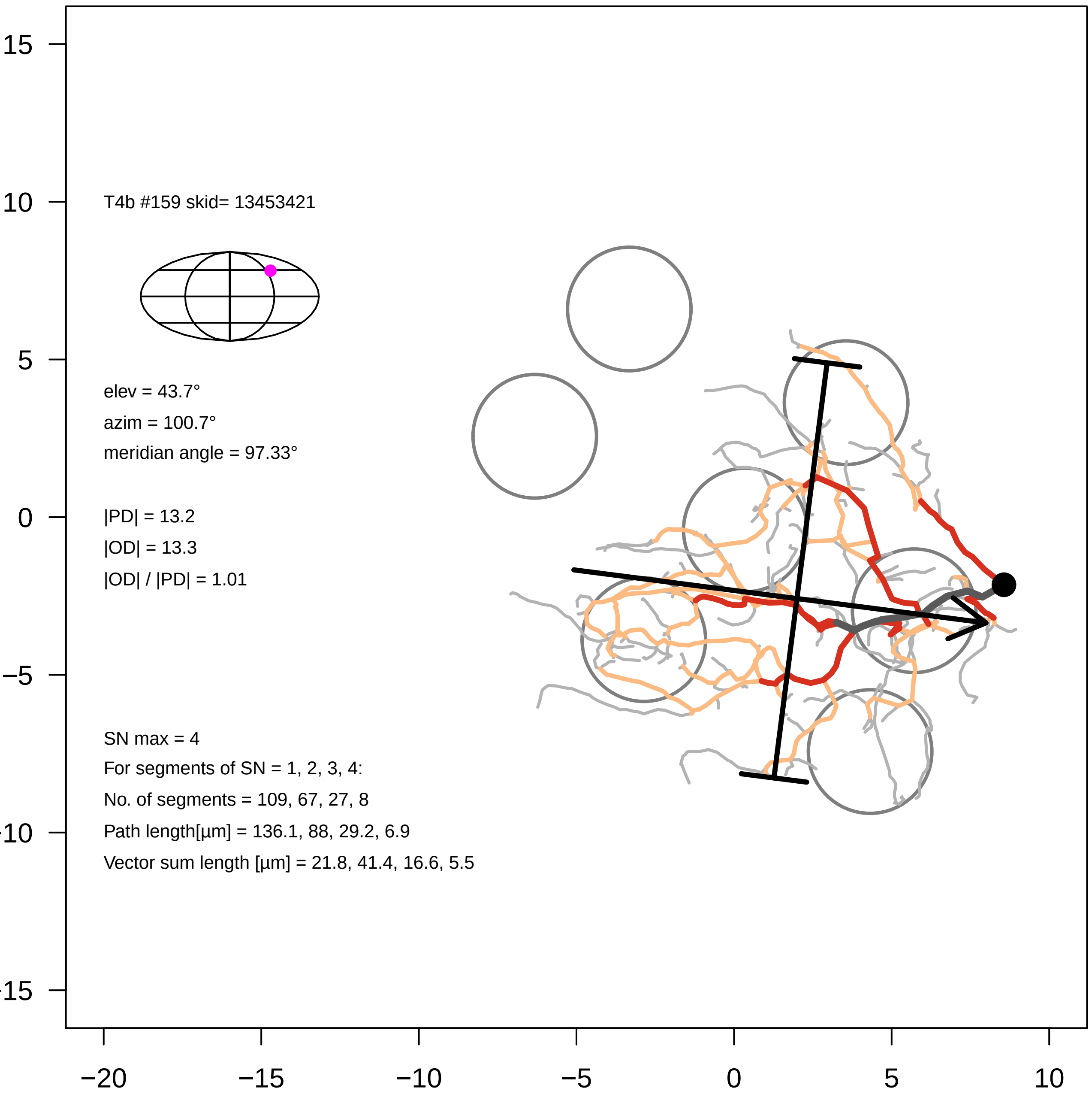
T4b #159 skid= 13453421
elev = 43.7°
azim = 100.7°
meridian angle = 97.33°
|PD| = 13.2
|OD| = 13.3
|OD| / |PD| = 1.01
SN max = 4
For segments of SN = 1, 2, 3, 4:
No. of segments = 109, 67, 27, 8
Path length[µm] = 136.1, 88, 29.2, 6.9
Vector sum length [µm] = 21.8, 41.4, 16.6, 5.5
15
10
5
0
−5
−10
−15
−20
−15
−10
−5
0
5
10

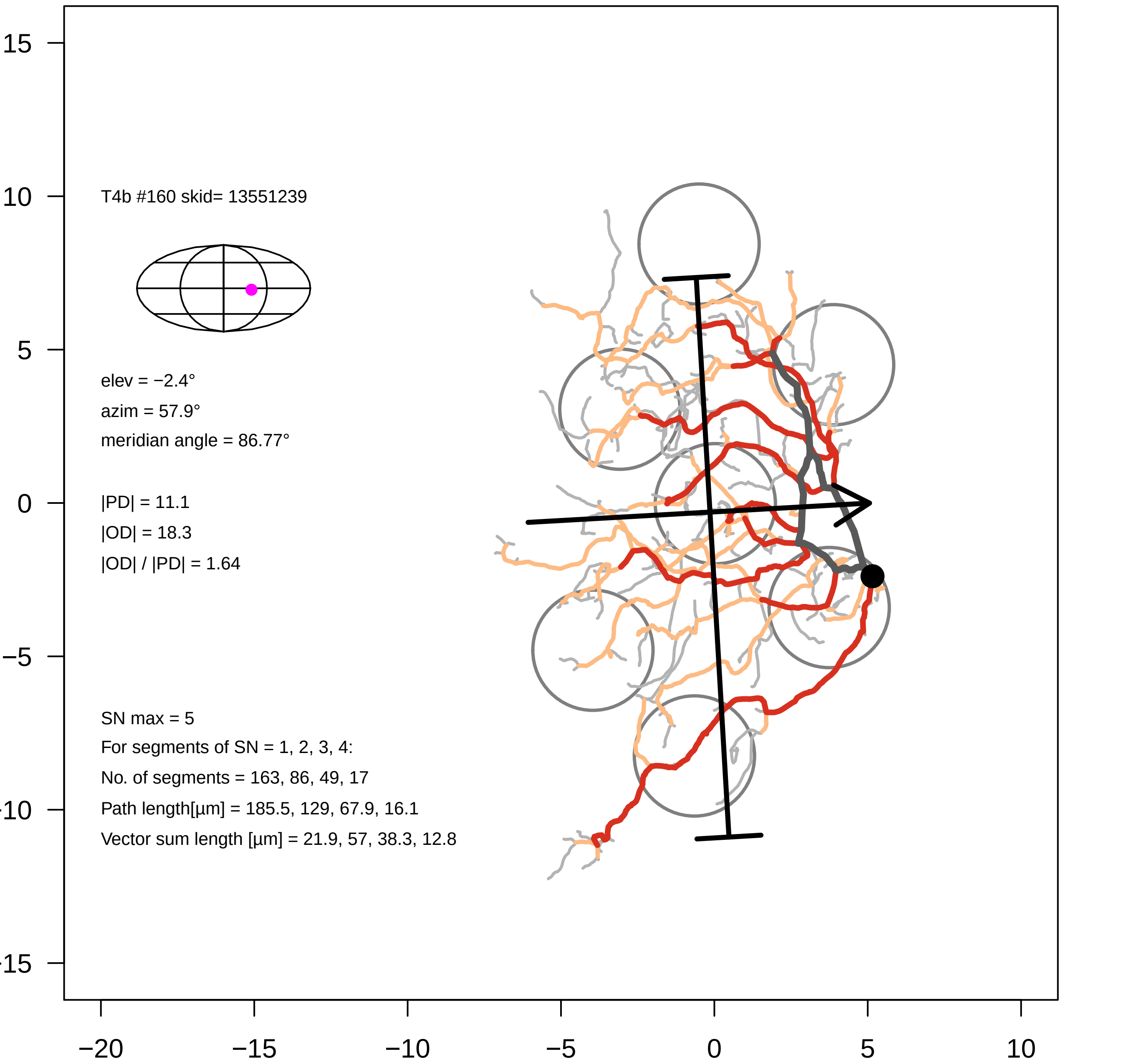

T4b #160 skid= 13551239
elev = −2.4°
azim = 57.9°
meridian angle = 86.77°
|PD| = 11.1
|OD| = 18.3
|OD| / |PD| = 1.64
SN max = 5
For segments of SN = 1, 2, 3, 4:
No. of segments = 163, 86, 49, 17
Path length[µm] = 185.5, 129, 67.9, 16.1
Vector sum length [µm] = 21.9, 57, 38.3, 12.8
15
10
5
0
−5
−10
−15
−20
−15
−10
−5
0
5
10

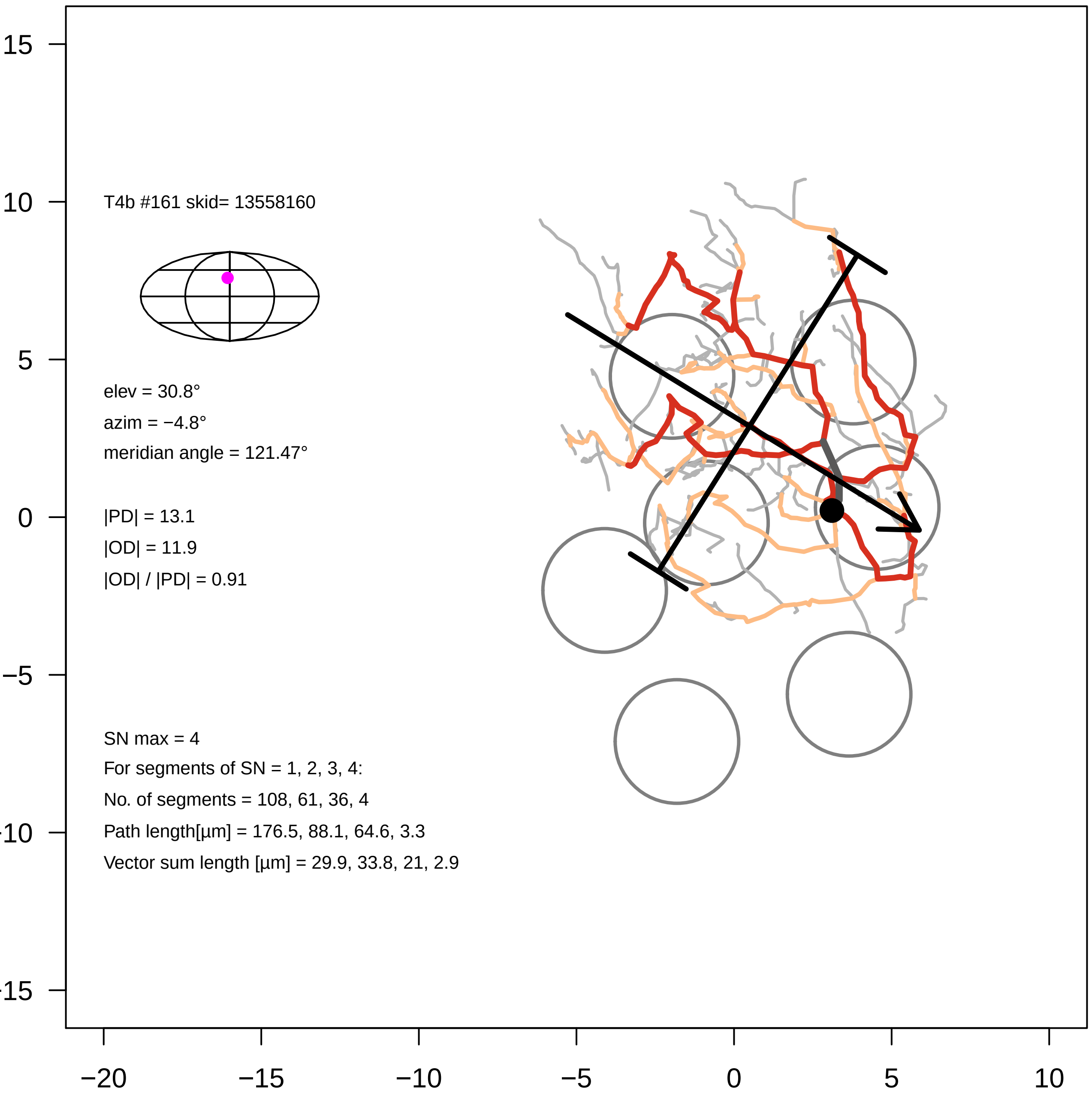

T4b #161 skid= 13558160
elev = 30.8°
azim = −4.8°
meridian angle = 121.47°
|PD| = 13.1
|OD| = 11.9
|OD| / |PD| = 0.91
SN max = 4
For segments of SN = 1, 2, 3, 4:
No. of segments = 108, 61, 36, 4
Path length[µm] = 176.5, 88.1, 64.6, 3.3
Vector sum length [µm] = 29.9, 33.8, 21, 2.9
15
10
5
0
−5
−10
−15
−20
−15
−10
−5
0
5
10

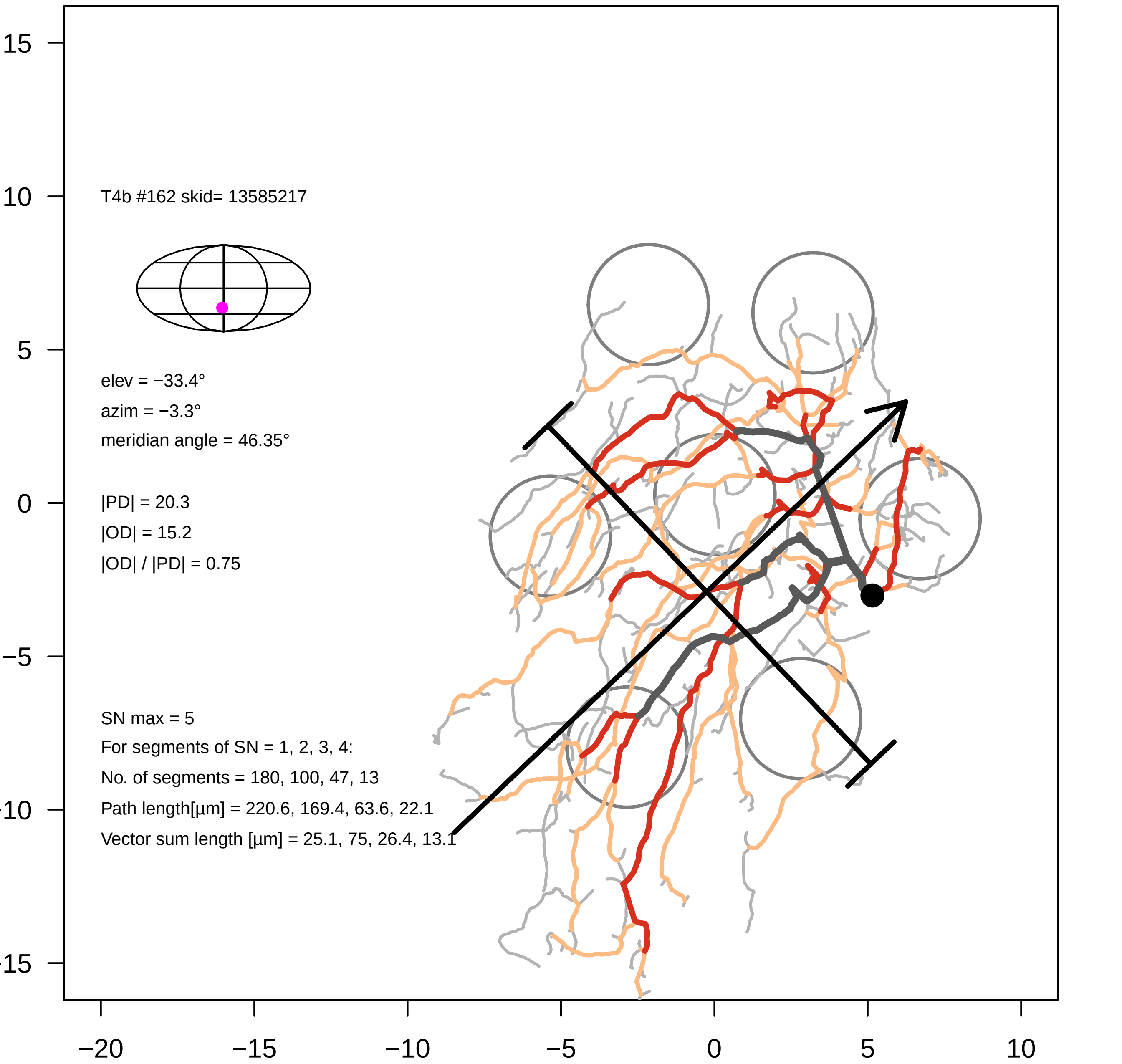

T4b #162 skid= 13585217
elev = −33.4°
azim = −3.3°
meridian angle = 46.35°
|PD| = 20.3
|OD| = 15.2
|OD| / |PD| = 0.75
SN max = 5
For segments of SN = 1, 2, 3, 4:
No. of segments = 180, 100, 47, 13
Path length[µm] = 220.6, 169.4, 63.6, 22.1
Vector sum length [µm] = 25.1, 75, 26.4, 13.1
15
10
5
0
−5
−10
−15
−20
−15
−10
−5
0
5
10

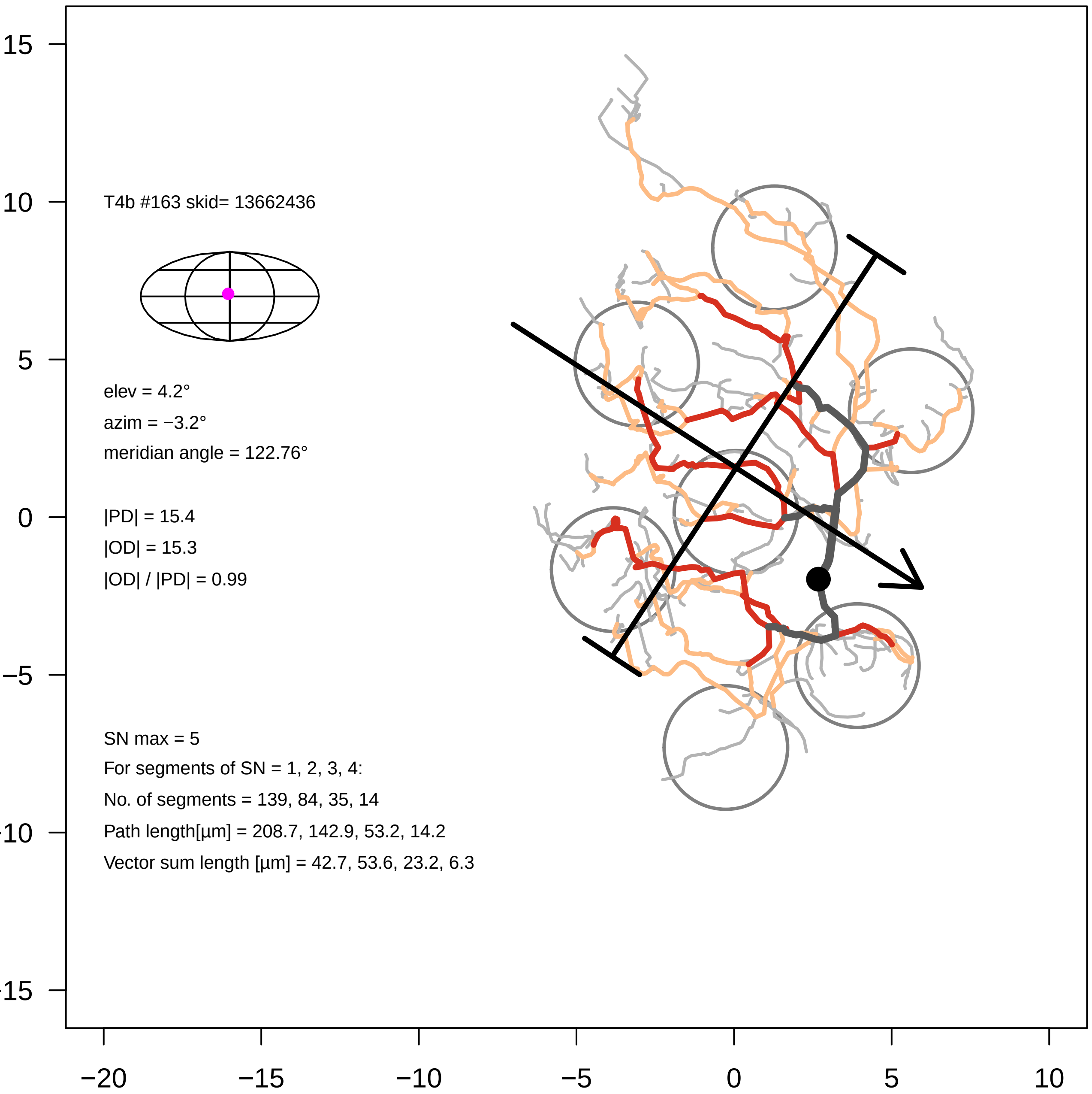

T4b #163 skid= 13662436
elev = 4.2°
azim = −3.2°
meridian angle = 122.76°
|PD| = 15.4
|OD| = 15.3
|OD| / |PD| = 0.99
SN max = 5
For segments of SN = 1, 2, 3, 4:
No. of segments = 139, 84, 35, 14
Path length[µm] = 208.7, 142.9, 53.2, 14.2
Vector sum length [µm] = 42.7, 53.6, 23.2, 6.3
15
10
5
0
−5
−10
−15
−20
−15
−10
−5
0
5
10

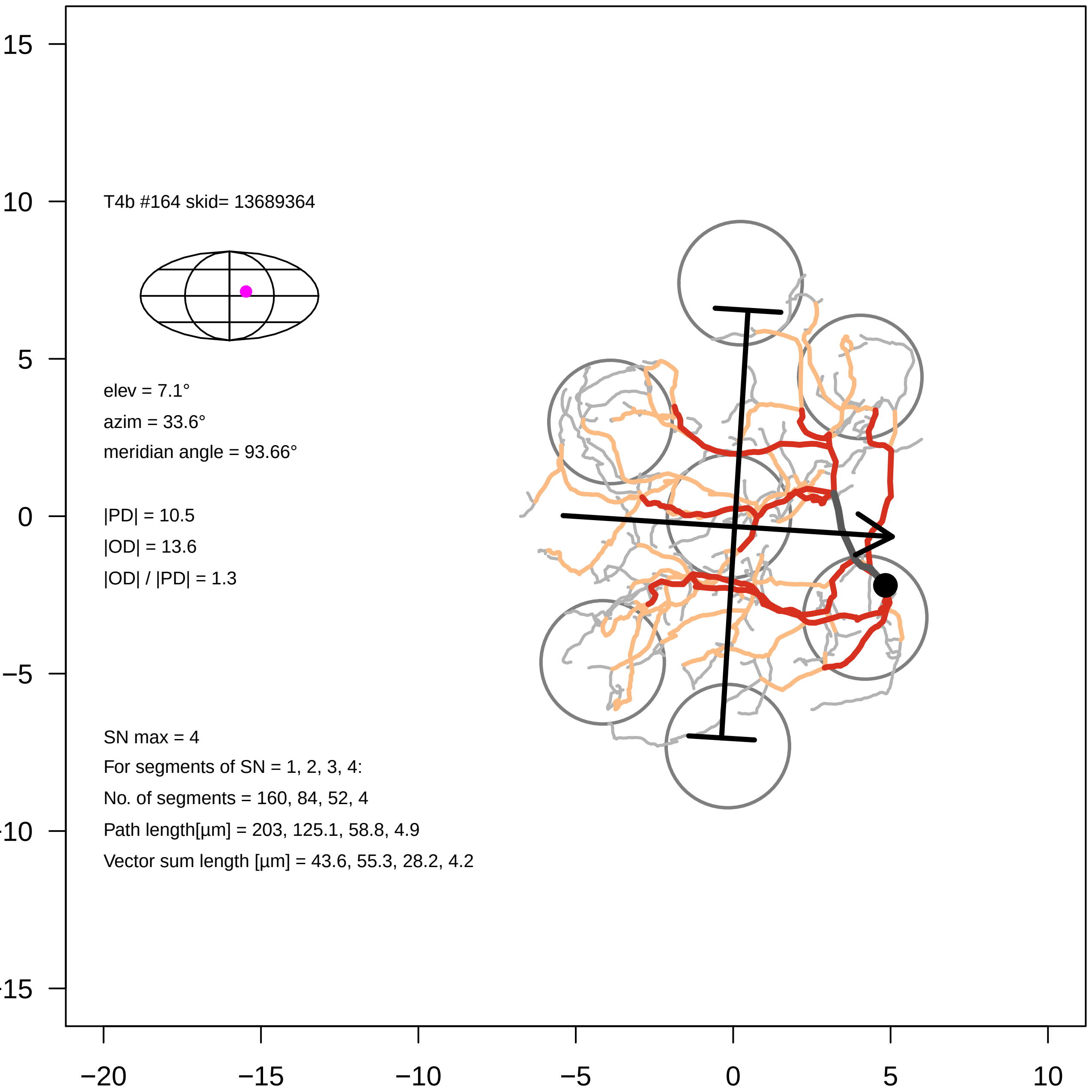
T4b #164 skid= 13689364
elev = 7.1°
azim = 33.6°
meridian angle = 93.66°
|PD| = 10.5
|OD| = 13.6
|OD| / |PD| = 1.3
SN max = 4
For segments of SN = 1, 2, 3, 4:
No. of segments = 160, 84, 52, 4
Path length[µm] = 203, 125.1, 58.8, 4.9
Vector sum length [µm] = 43.6, 55.3, 28.2, 4.2
15
10
5
0
−5
−10
−15
−20
−15
−10
−5
0
5
10

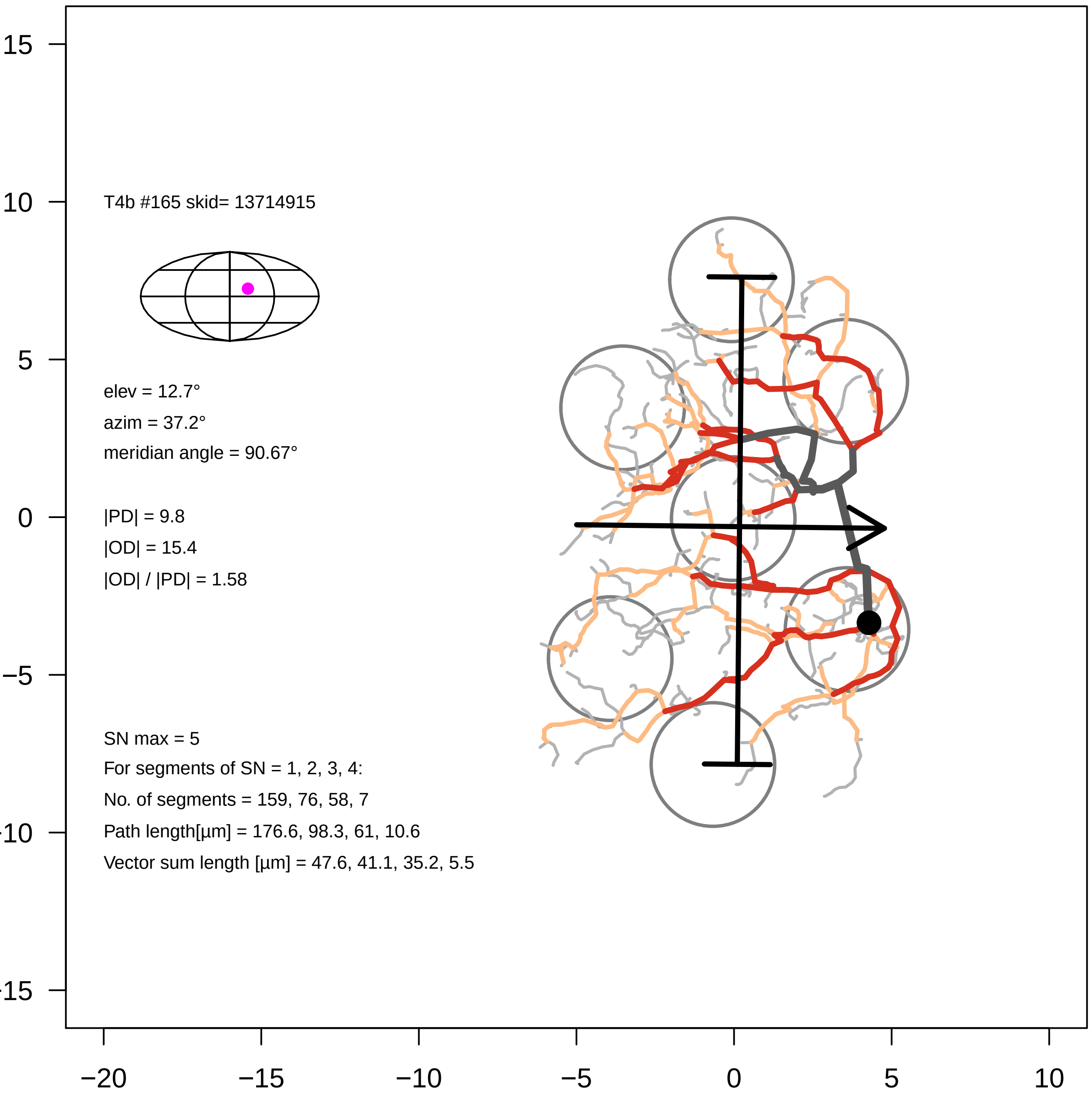

T4b #165 skid= 13714915
elev = 12.7°
azim = 37.2°
meridian angle = 90.67°
|PD| = 9.8
|OD| = 15.4
|OD| / |PD| = 1.58
SN max = 5
For segments of SN = 1, 2, 3, 4:
No. of segments = 159, 76, 58, 7
Path length[µm] = 176.6, 98.3, 61, 10.6
Vector sum length [µm] = 47.6, 41.1, 35.2, 5.5
15
10
5
0
−5
−10
−15
−20
−15
−10
−5
0
5
10

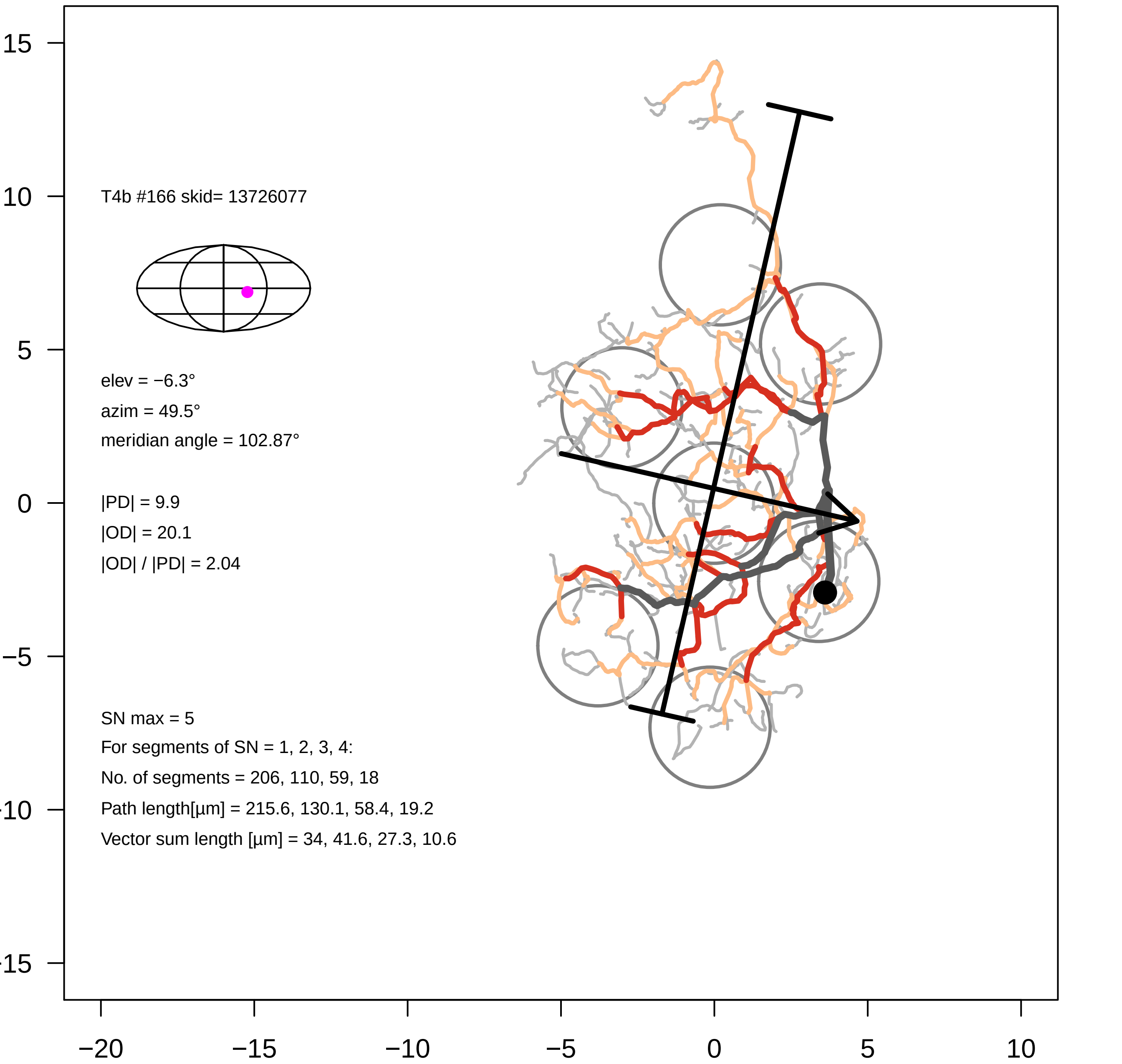
T4b #166 skid= 13726077
elev = −6.3°
azim = 49.5°
meridian angle = 102.87°
|PD| = 9.9
|OD| = 20.1
|OD| / |PD| = 2.04
SN max = 5
For segments of SN = 1, 2, 3, 4:
No. of segments = 206, 110, 59, 18
Path length[µm] = 215.6, 130.1, 58.4, 19.2
Vector sum length [µm] = 34, 41.6, 27.3, 10.6
15
10
5
0
−5
−10
−15
−20
−15
−10
−5
0
5
10

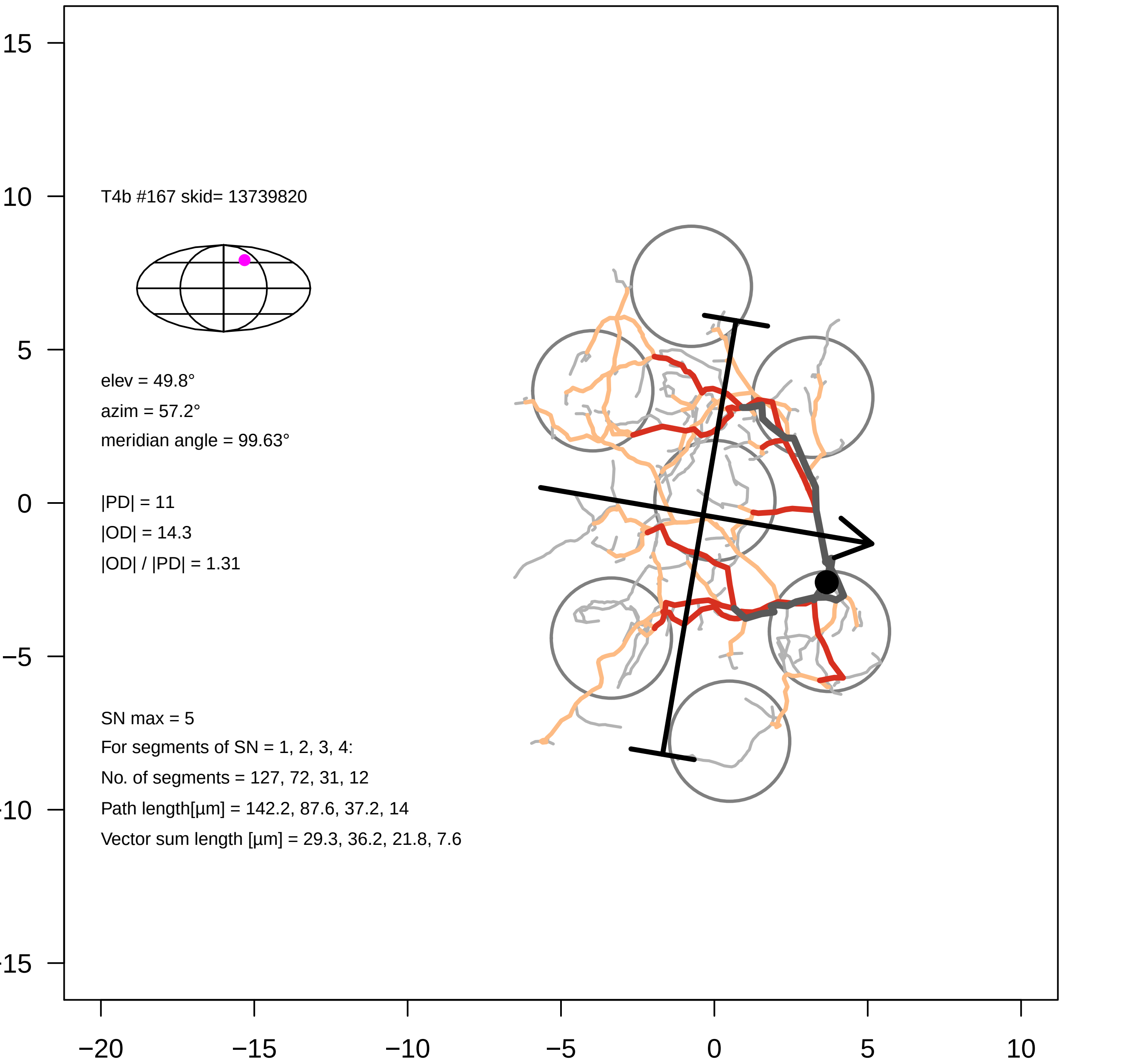

T4b #167 skid= 13739820
elev = 49.8°
azim = 57.2°
meridian angle = 99.63°
|PD| = 11
|OD| = 14.3
|OD| / |PD| = 1.31
SN max = 5
For segments of SN = 1, 2, 3, 4:
No. of segments = 127, 72, 31, 12
Path length[µm] = 142.2, 87.6, 37.2, 14
Vector sum length [µm] = 29.3, 36.2, 21.8, 7.6
15
10
5
0
−5
−10
−15
−20
−15
−10
−5
0
5
10

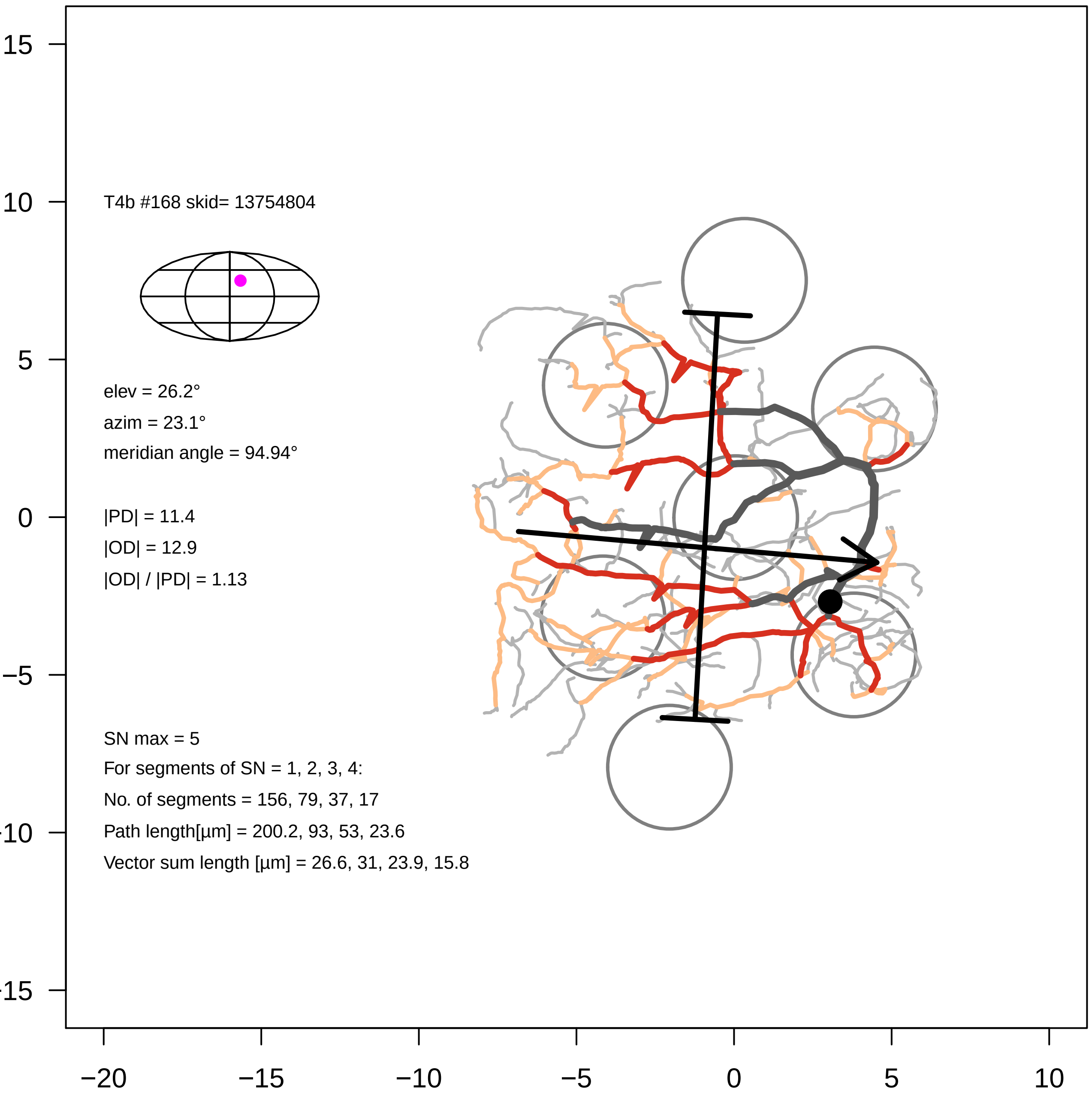

T4b #168 skid= 13754804
elev = 26.2°
azim = 23.1°
meridian angle = 94.94°
|PD| = 11.4
|OD| = 12.9
|OD| / |PD| = 1.13
SN max = 5
For segments of SN = 1, 2, 3, 4:
No. of segments = 156, 79, 37, 17
Path length[µm] = 200.2, 93, 53, 23.6
Vector sum length [µm] = 26.6, 31, 23.9, 15.8
15
10
5
0
−5
−10
−15
−20
−15
−10
−5
0
5
10

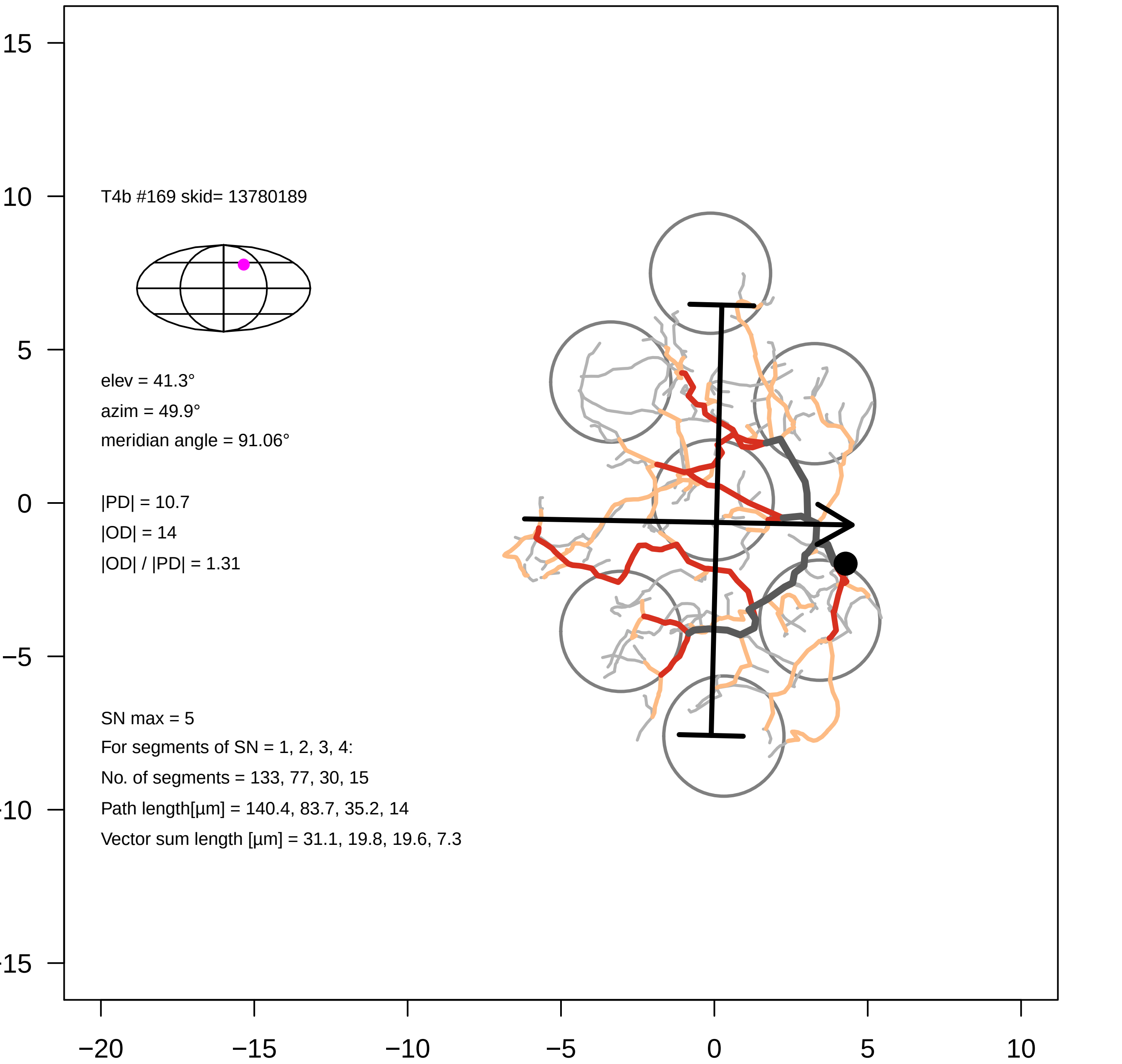

T4b #169 skid= 13780189
elev = 41.3°
azim = 49.9°
meridian angle = 91.06°
|PD| = 10.7
|OD| = 14
|OD| / |PD| = 1.31
SN max = 5
For segments of SN = 1, 2, 3, 4:
No. of segments = 133, 77, 30, 15
Path length[µm] = 140.4, 83.7, 35.2, 14
Vector sum length [µm] = 31.1, 19.8, 19.6, 7.3
15
10
5
0
−5
−10
−15
−20
−15
−10
−5
0
5
10

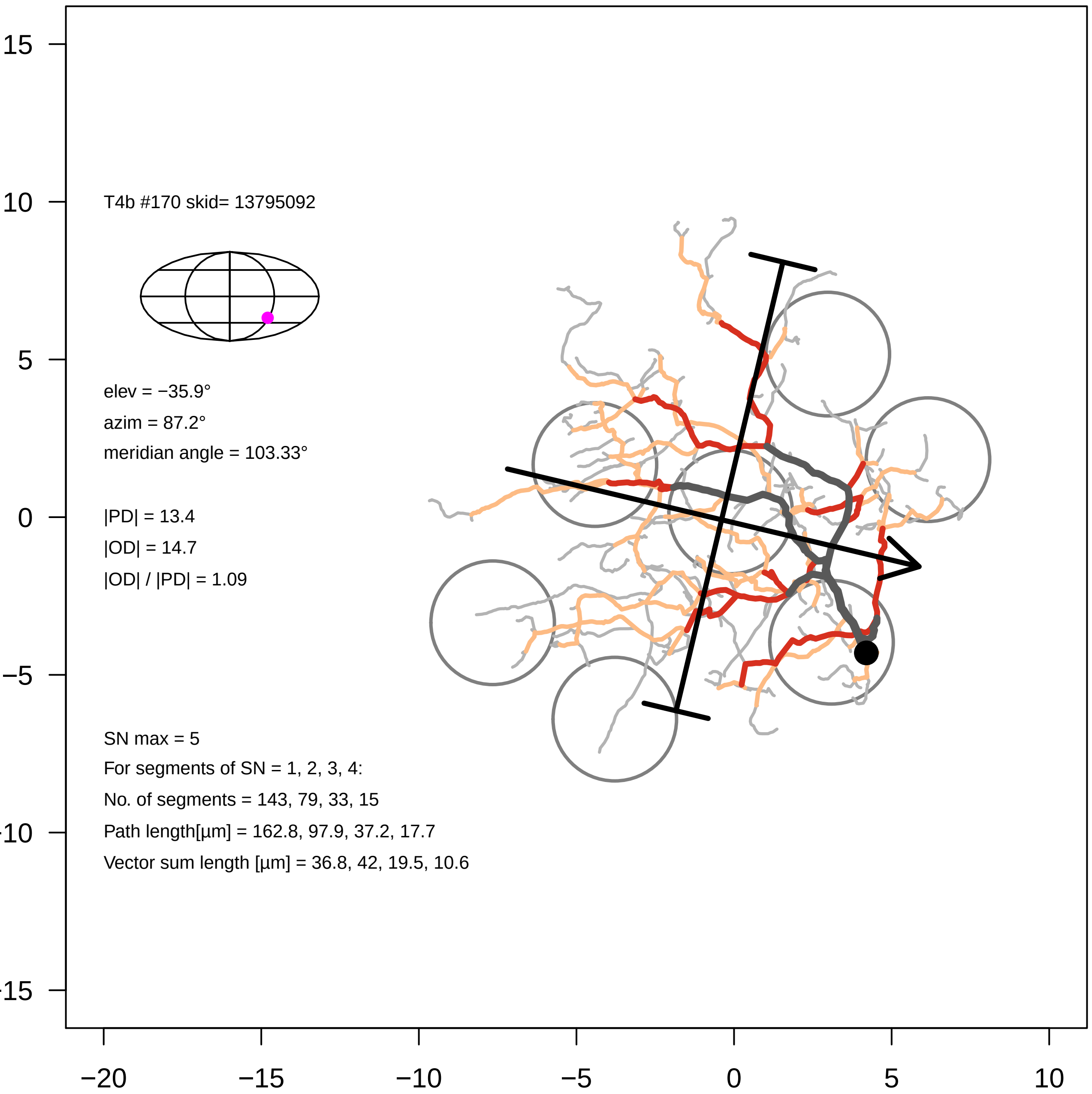
T4b #170 skid= 13795092
elev = −35.9°
azim = 87.2°
meridian angle = 103.33°
|PD| = 13.4
|OD| = 14.7
|OD| / |PD| = 1.09
SN max = 5
For segments of SN = 1, 2, 3, 4:
No. of segments = 143, 79, 33, 15
Path length[µm] = 162.8, 97.9, 37.2, 17.7
Vector sum length [µm] = 36.8, 42, 19.5, 10.6
15
10
5
0
−5
-10
-15
−20
−15
−10
−5
0
5
10

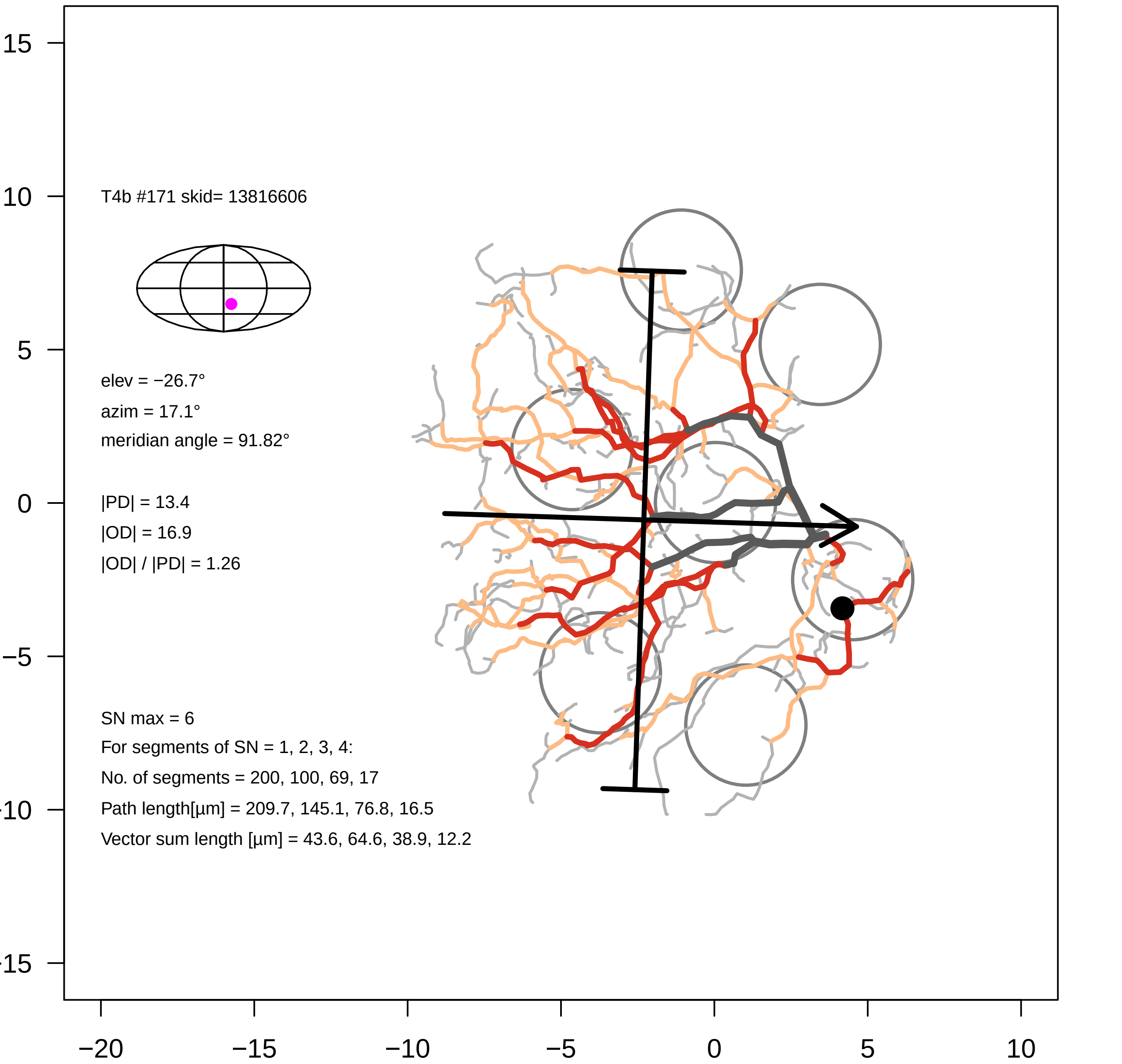
T4b #171 skid= 13816606
elev = −26.7°
azim = 17.1°
meridian angle = 91.82°
|PD| = 13.4
|OD| = 16.9
|OD| / |PD| = 1.26
SN max = 6
For segments of SN = 1, 2, 3, 4:
No. of segments = 200, 100, 69, 17
Path length[µm] = 209.7, 145.1, 76.8, 16.5
Vector sum length [µm] = 43.6, 64.6, 38.9, 12.2
15
10
5
0
−5
−10
−15
−20
−15
−10
−5
0
5
10

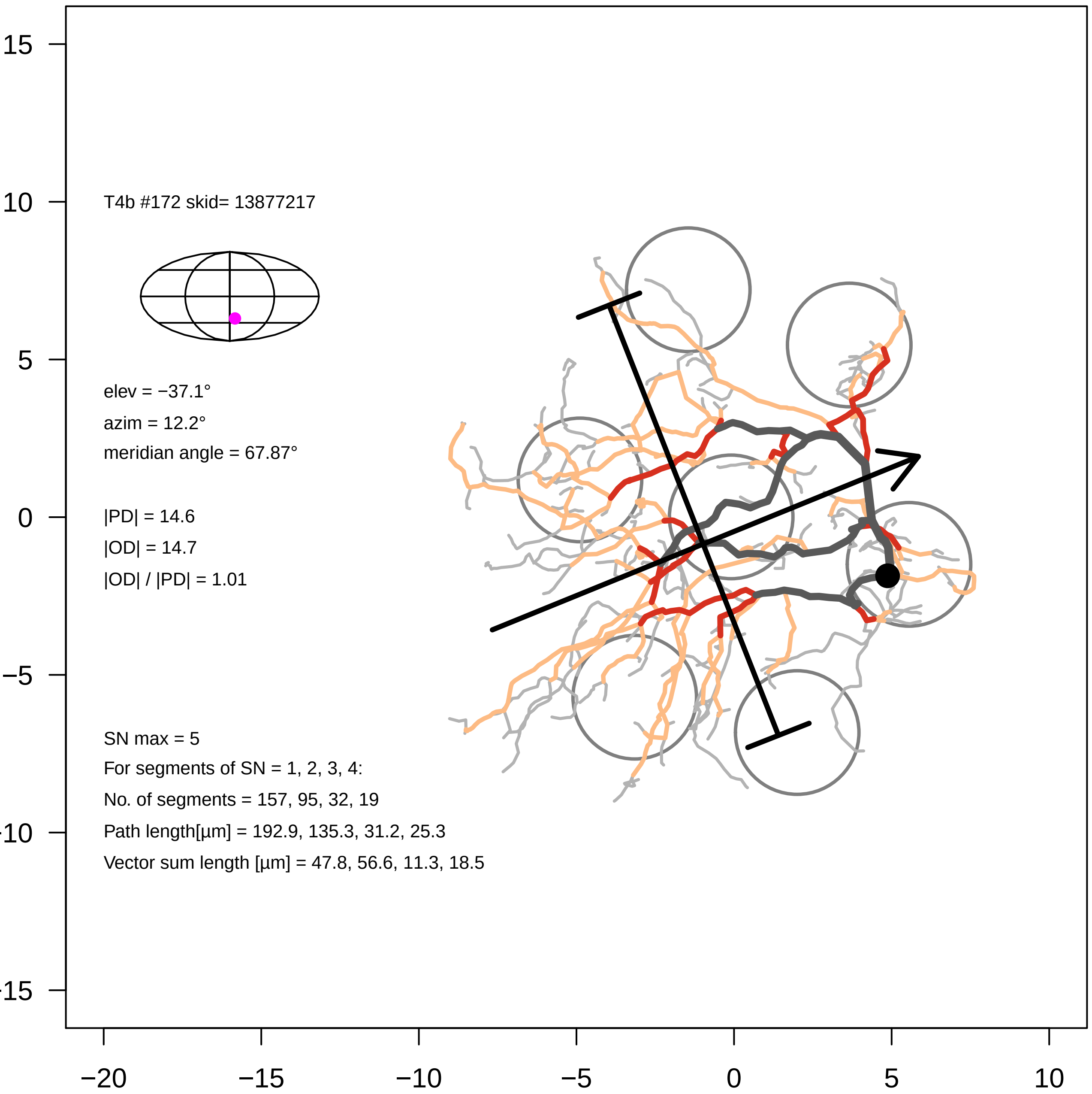

T4b #172 skid= 13877217
elev = −37.1°
azim = 12.2°
meridian angle = 67.87°
|PD| = 14.6
|OD| = 14.7
|OD| / |PD| = 1.01
SN max = 5
For segments of SN = 1, 2, 3, 4:
No. of segments = 157, 95, 32, 19
Path length[µm] = 192.9, 135.3, 31.2, 25.3
Vector sum length [µm] = 47.8, 56.6, 11.3, 18.5
15
10
5
0
−5
−10
−15
−20
−15
−10
−5
0
5
10

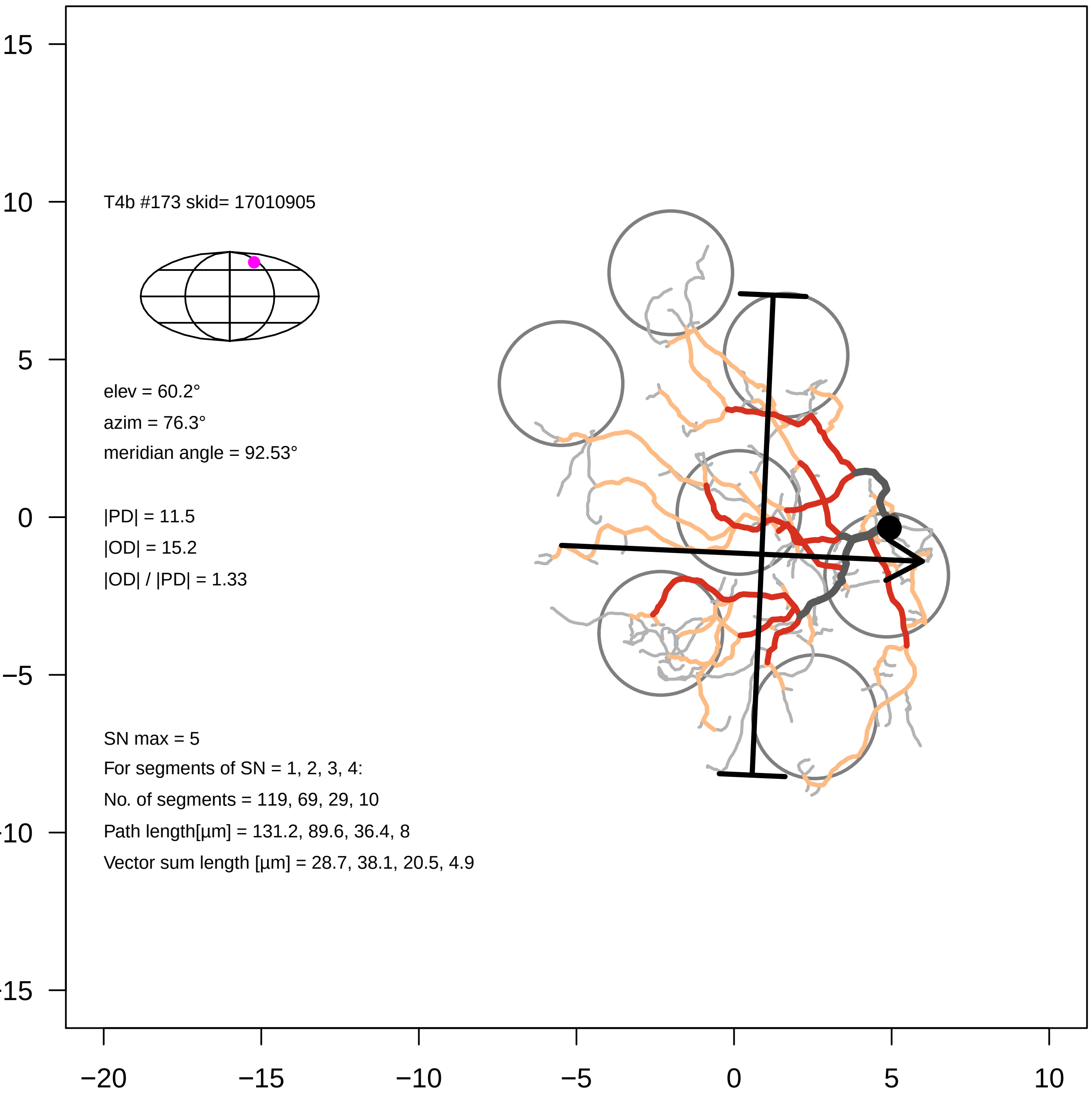
T4b #173 skid= 17010905
elev = 60.2°
azim = 76.3°
meridian angle = 92.53°
|PD| = 11.5
|OD| = 15.2
|OD| / |PD| = 1.33
SN max = 5
For segments of SN = 1, 2, 3, 4:
No. of segments = 119, 69, 29, 10
Path length[µm] = 131.2, 89.6, 36.4, 8
Vector sum length [µm] = 28.7, 38.1, 20.5, 4.9
15
10
5
0
−5
−10
−15
−20
−15
−10
−5
0
5
10

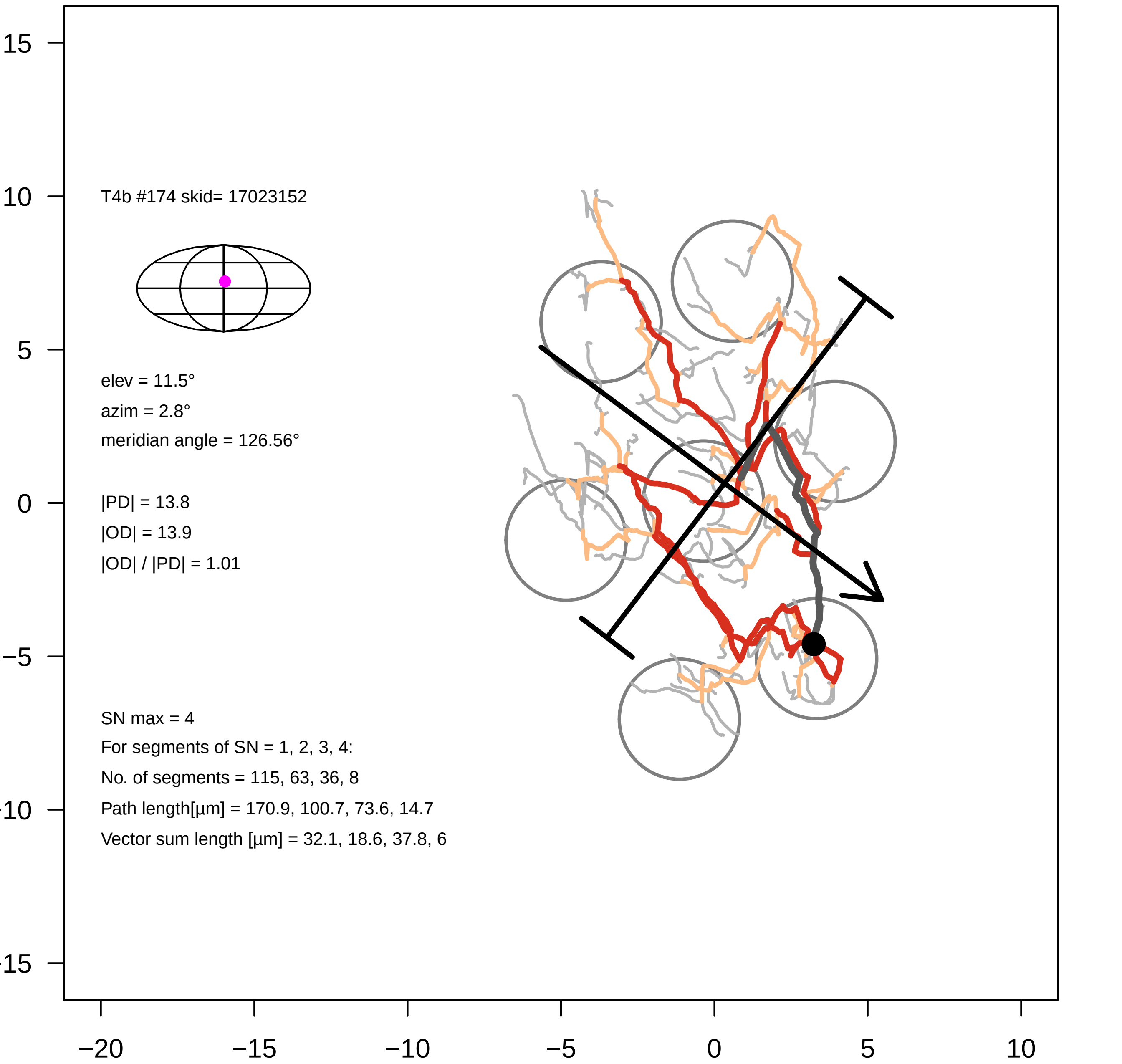

T4b #174 skid= 17023152
elev = 11.5°
azim = 2.8°
meridian angle = 126.56°
|PD| = 13.8
|OD| = 13.9
|OD| / |PD| = 1.01
SN max = 4
For segments of SN = 1, 2, 3, 4:
No. of segments = 115, 63, 36, 8
Path length[µm] = 170.9, 100.7, 73.6, 14.7
Vector sum length [µm] = 32.1, 18.6, 37.8, 6
15
10
5
0
−5
−10
−15
−20
−15
−10
−5
0
5
10

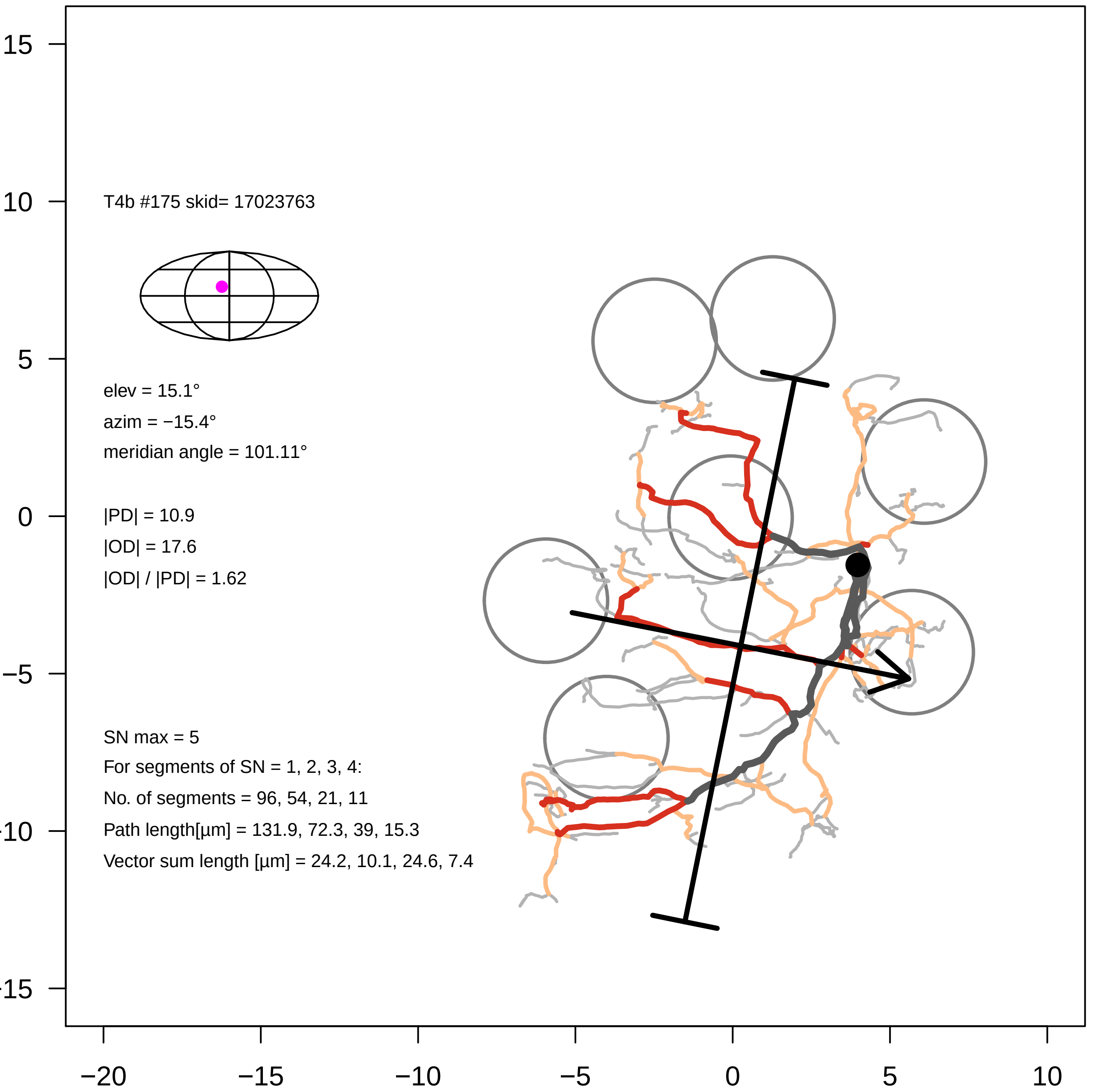

T4b #175 skid= 17023763
elev = 15.1°
azim = −15.4°
meridian angle = 101.11°
|PD| = 10.9
|OD| = 17.6
|OD| / |PD| = 1.62
SN max = 5
For segments of SN = 1, 2, 3, 4:
No. of segments = 96, 54, 21, 11
Path length[µm] = 131.9, 72.3, 39, 15.3
Vector sum length [µm] = 24.2, 10.1, 24.6, 7.4
15
10
5
0
−5
−10
−15
−20
−15
−10
−5
0
5
10

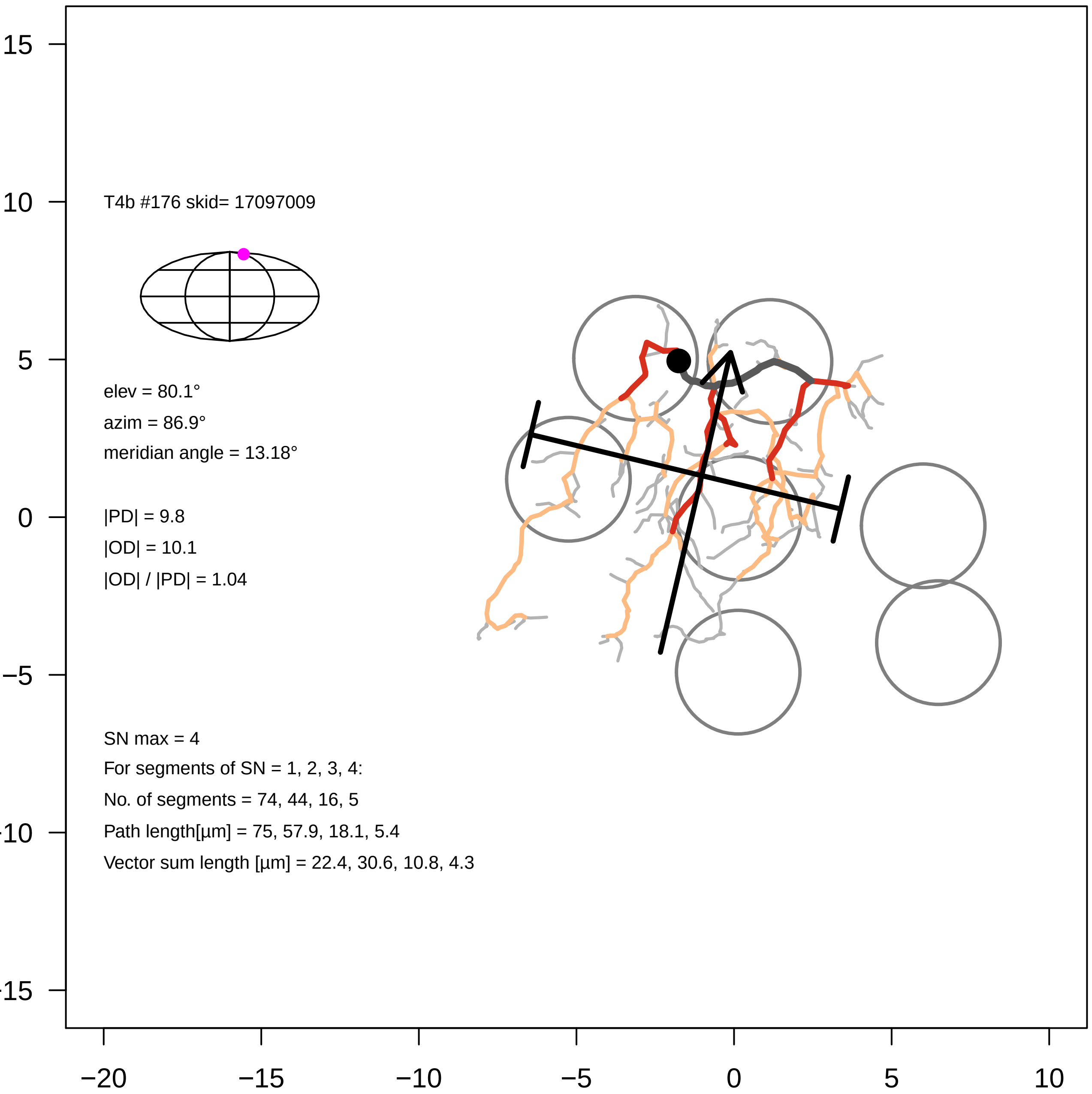

T4b #176 skid= 17097009
elev = 80.1°
azim = 86.9°
meridian angle = 13.18°
|PD| = 9.8
|OD| = 10.1
|OD| / |PD| = 1.04
SN max = 4
For segments of SN = 1, 2, 3, 4:
No. of segments = 74, 44, 16, 5
Path length[µm] = 75, 57.9, 18.1, 5.4
Vector sum length [µm] = 22.4, 30.6, 10.8, 4.3
15
10
5
0
−5
-10
-15
−20
−15
−10
−5
0
5
10
