## Supplementary Files, T4 galleries for "Eye structure shapes neuron function in *Drosophila* motion vision": Supplementary_Data_File3_T4c.pdf

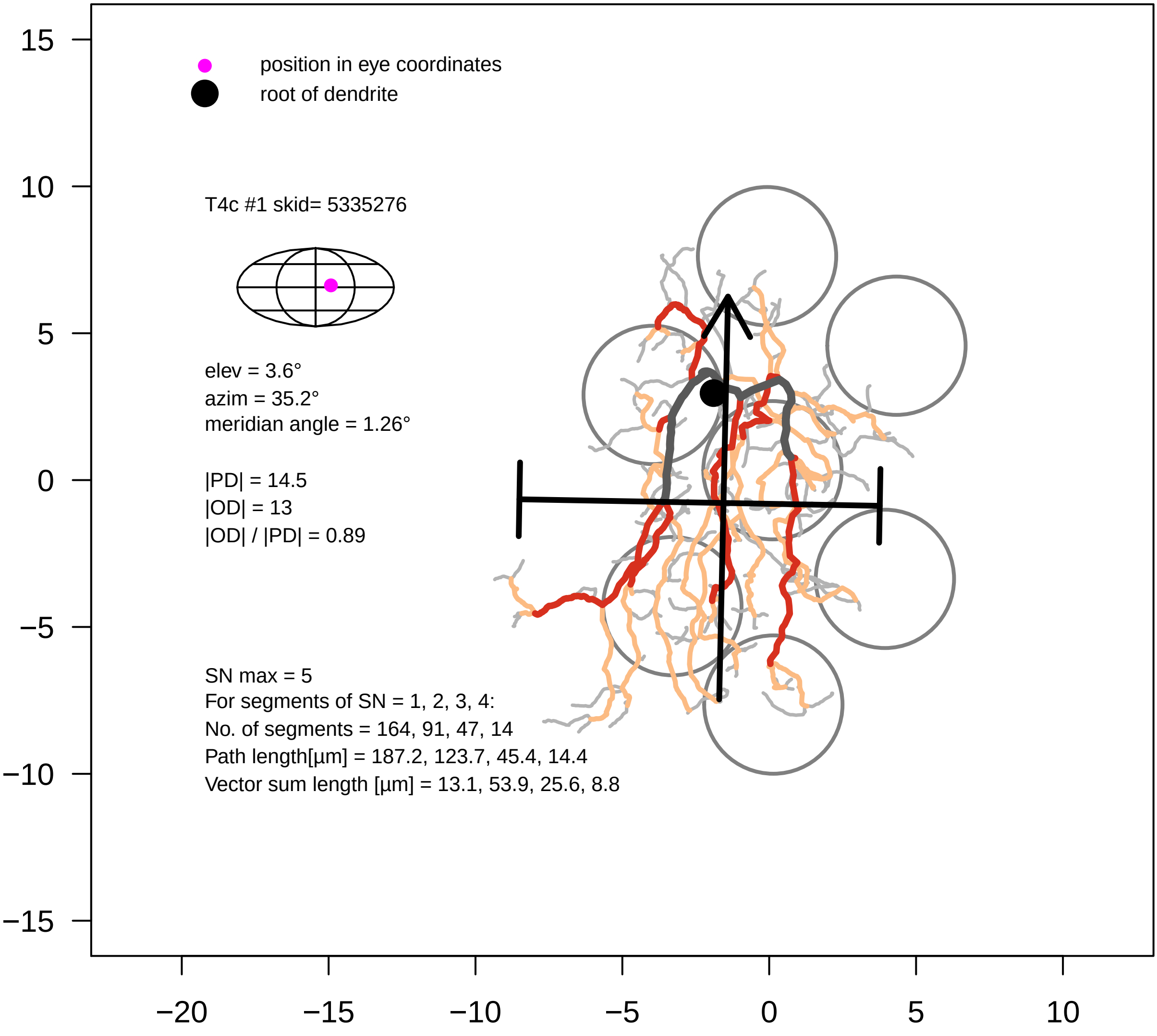
position in eye coordinates
root of dendrite
T4c #1 skid= 5335276
elev = 3.6°
azim = 35.2°
meridian angle = 1.26°
|PD| = 14.5
|OD| = 13
|OD| / |PD| = 0.89
SN max = 5
For segments of SN = 1, 2, 3, 4:
No. of segments = 164, 91, 47, 14
Path length[µm] = 187.2, 123.7, 45.4, 14.4
Vector sum length [µm] = 13.1, 53.9, 25.6, 8.8
15
10
5
0
−5
−10
−15
−20
−15
−10
−5
0
5
10

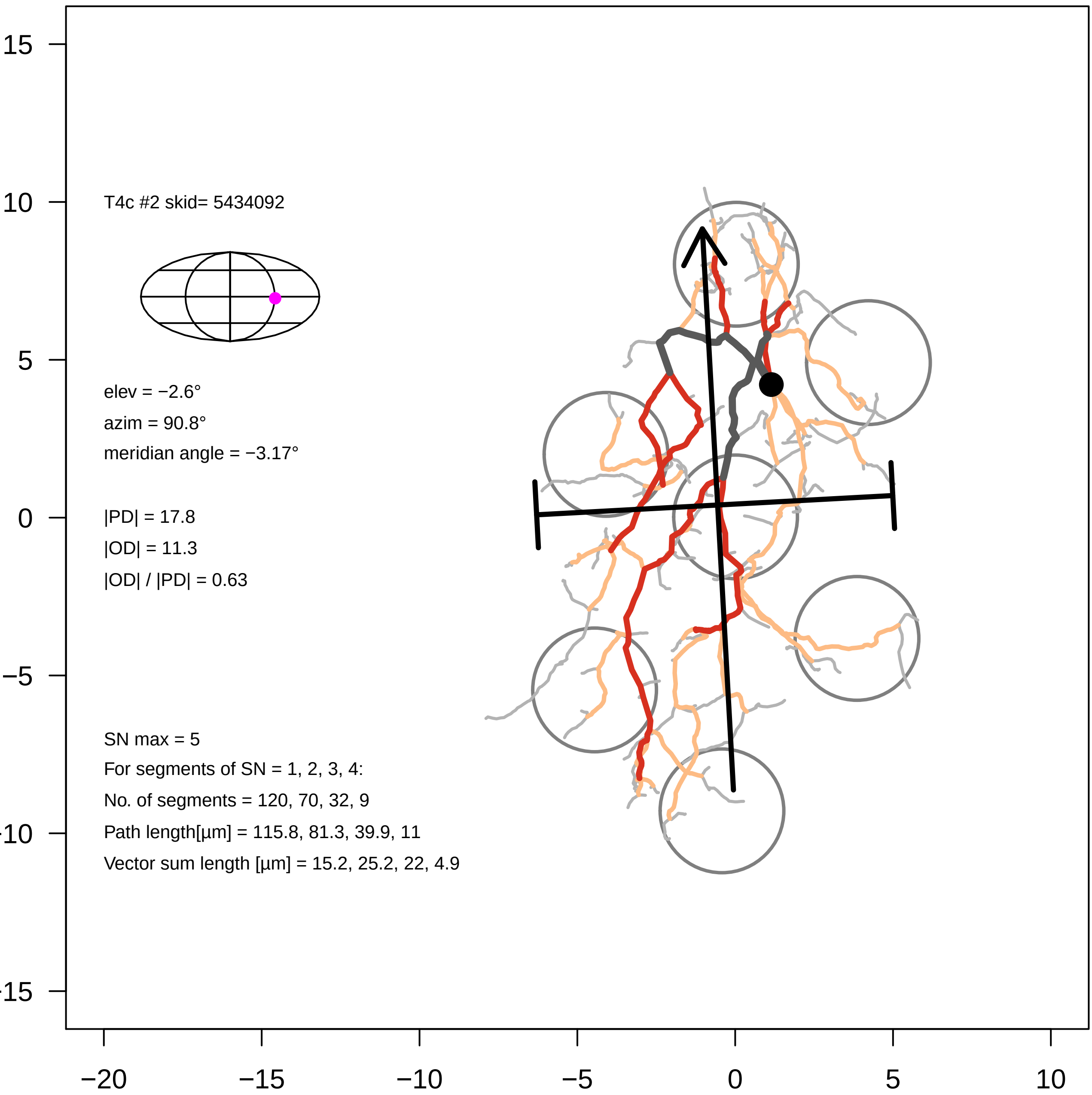

T4c #2 skid= 5434092
elev = −2.6°
azim = 90.8°
meridian angle = −3.17°
|PD| = 17.8
|OD| = 11.3
|OD| / |PD| = 0.63
SN max = 5
For segments of SN = 1, 2, 3, 4:
No. of segments = 120, 70, 32, 9
Path length[µm] = 115.8, 81.3, 39.9, 11
Vector sum length [µm] = 15.2, 25.2, 22, 4.9
15
10
5
0
−5
−10
−15
−20
−15
−10
−5
0
5
10

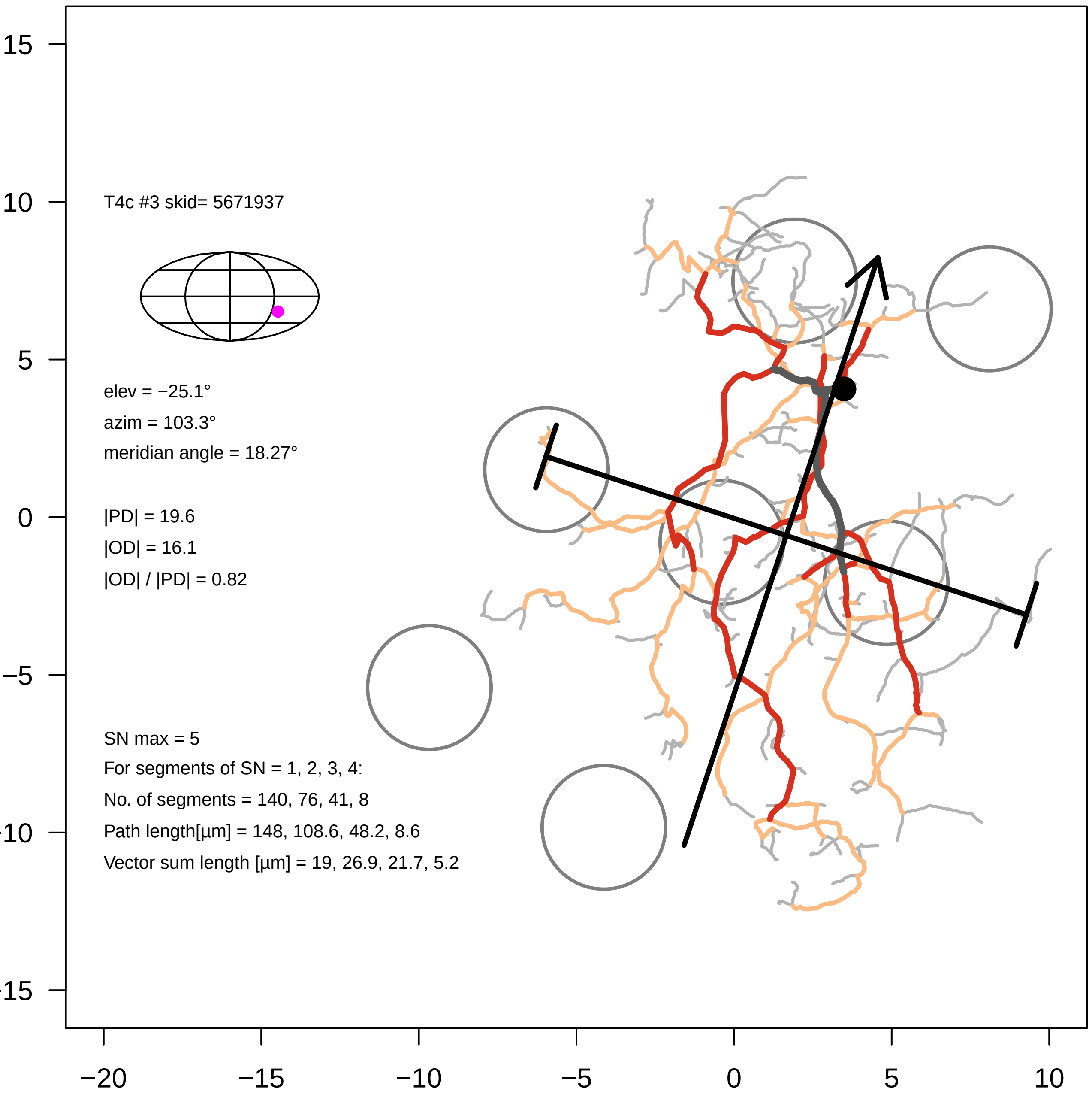
T4c #3 skid= 5671937
elev = −25.1°
azim = 103.3°
meridian angle = 18.27°
|PD| = 19.6
|OD| = 16.1
|OD| / |PD| = 0.82
SN max = 5
For segments of SN = 1, 2, 3, 4:
No. of segments = 140, 76, 41, 8
Path length[µm] = 148, 108.6, 48.2, 8.6
Vector sum length [µm] = 19, 26.9, 21.7, 5.2
15
10
5
0
−5
−10
−15
−20
−15
−10
−5
0
5
10

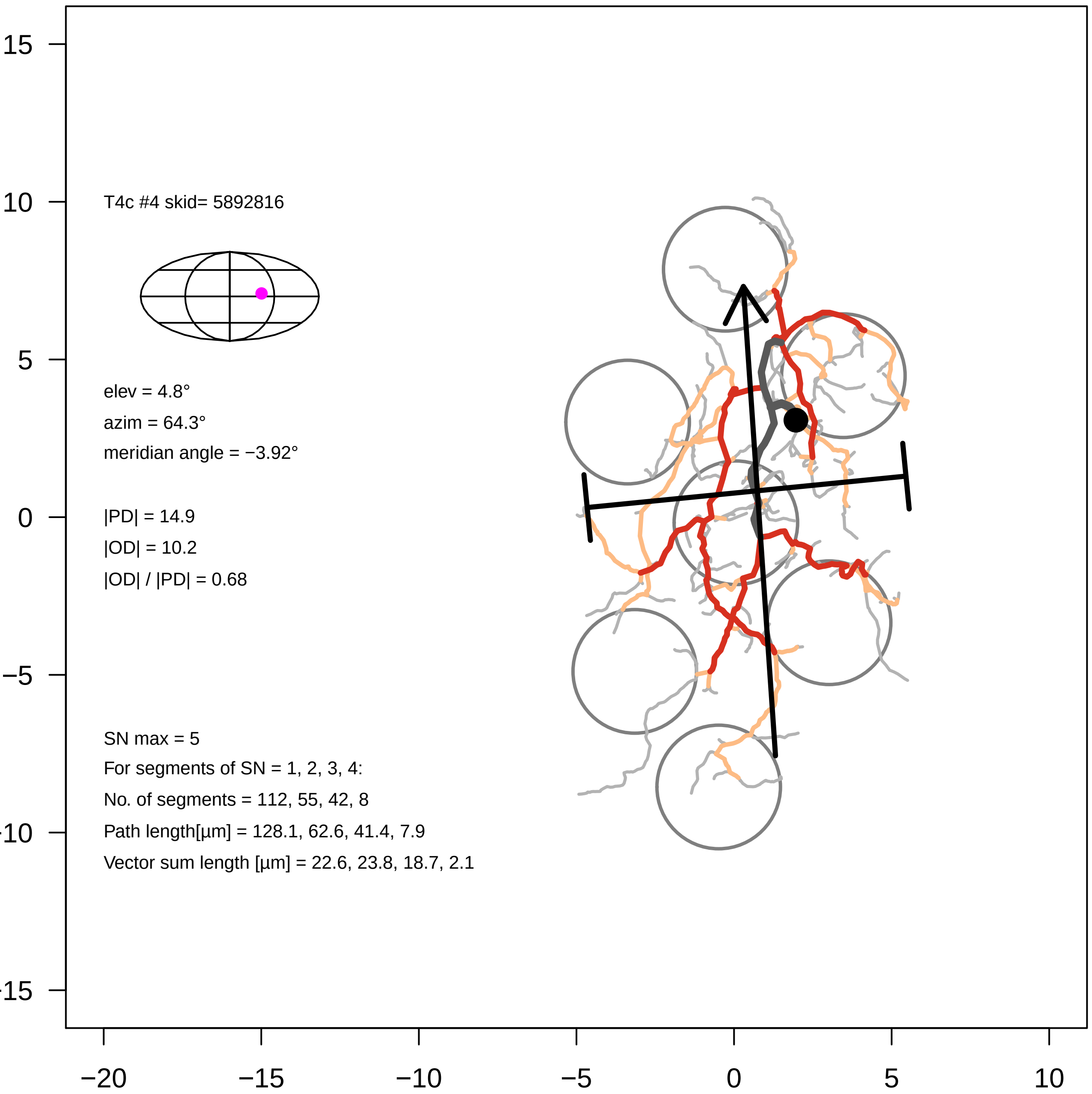

T4c #4 skid= 5892816
elev = 4.8°
azim = 64.3°
meridian angle = −3.92°
|PD| = 14.9
|OD| = 10.2
|OD| / |PD| = 0.68
SN max = 5
For segments of SN = 1, 2, 3, 4:
No. of segments = 112, 55, 42, 8
Path length[µm] = 128.1, 62.6, 41.4, 7.9
Vector sum length [µm] = 22.6, 23.8, 18.7, 2.1
15
10
5
0
−5
−10
−15
−20
−15
−10
−5
0
5
10

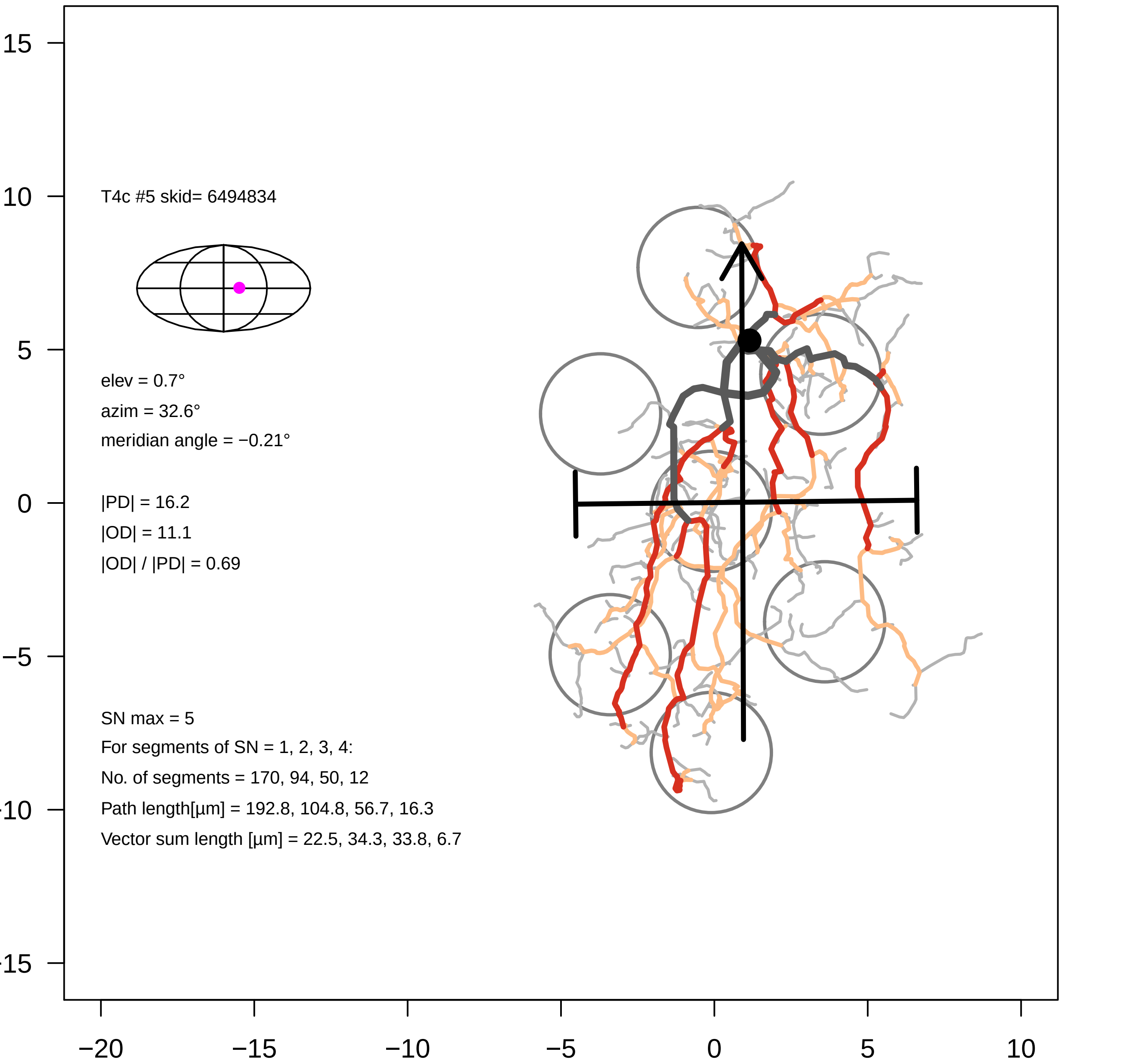

T4c #5 skid= 6494834
elev = 0.7°
azim = 32.6°
meridian angle = −0.21°
|PD| = 16.2
|OD| = 11.1
|OD| / |PD| = 0.69
SN max = 5
For segments of SN = 1, 2, 3, 4:
No. of segments = 170, 94, 50, 12
Path length[µm] = 192.8, 104.8, 56.7, 16.3
Vector sum length [µm] = 22.5, 34.3, 33.8, 6.7
15
10
5
0
−5
−10
−15
−20
−15
−10
−5
0
5
10

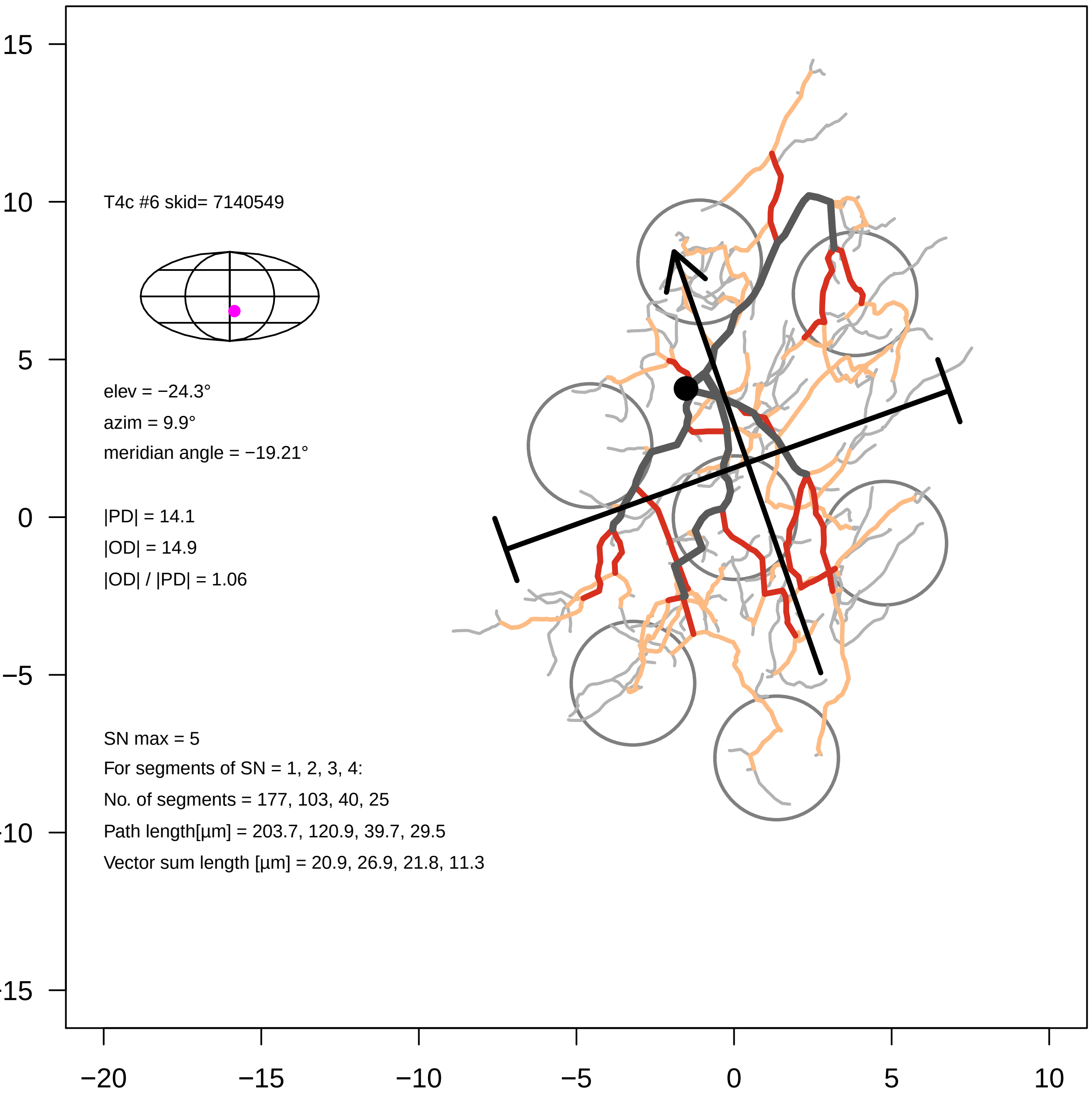

T4c #6 skid= 7140549
elev = −24.3°
azim = 9.9°
meridian angle = −19.21°
|PD| = 14.1
|OD| = 14.9
|OD| / |PD| = 1.06
SN max = 5
For segments of SN = 1, 2, 3, 4:
No. of segments = 177, 103, 40, 25
Path length[µm] = 203.7, 120.9, 39.7, 29.5
Vector sum length [µm] = 20.9, 26.9, 21.8, 11.3
15
10
5
0
−5
-10
-15
−20
−15
−10
−5
0
5
10

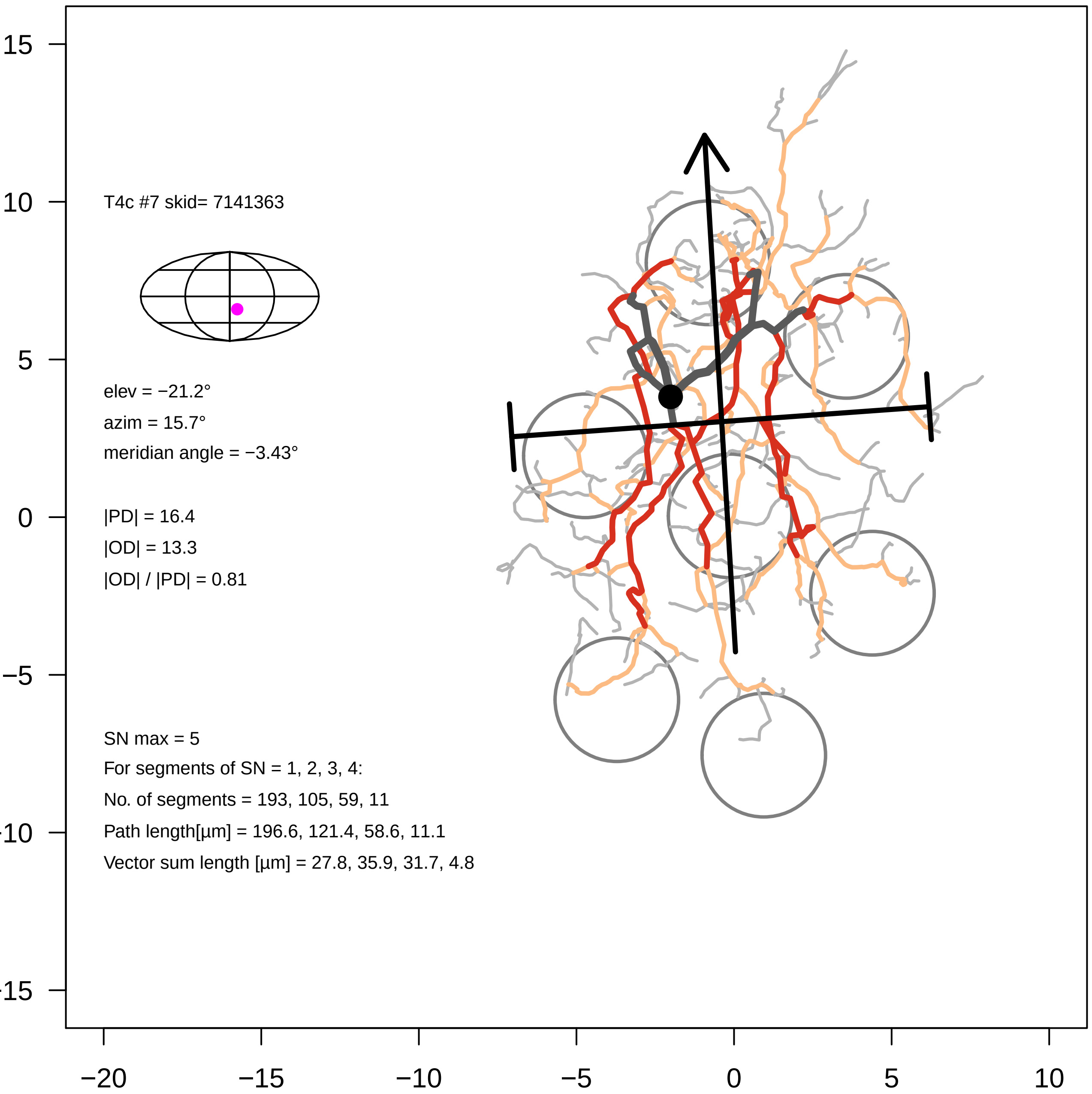

15
10
5
0
−5
−10
−15
−20
−15
−10
−5
0
5
10
T4c #7 skid= 7141363
elev = −21.2°
azim = 15.7°
meridian angle = −3.43°
|PD| = 16.4
|OD| = 13.3
|OD| / |PD| = 0.81
SN max = 5
For segments of SN = 1, 2, 3, 4:
No. of segments = 193, 105, 59, 11
Path length[µm] = 196.6, 121.4, 58.6, 11.1
Vector sum length [µm] = 27.8, 35.9, 31.7, 4.8

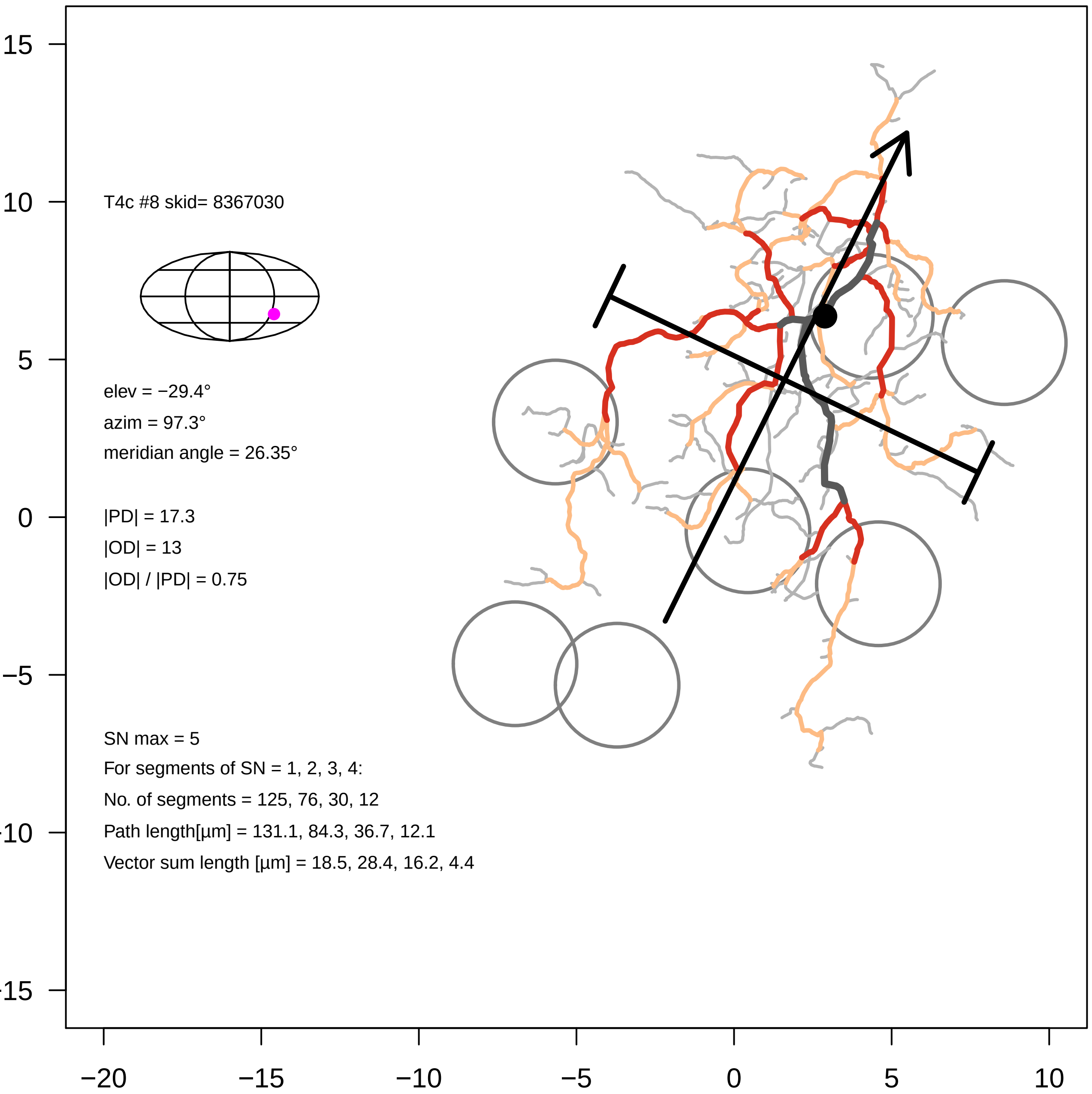
T4c #8 skid= 8367030
elev = −29.4°
azim = 97.3°
meridian angle = 26.35°
|PD| = 17.3
|OD| = 13
|OD| / |PD| = 0.75
SN max = 5
For segments of SN = 1, 2, 3, 4:
No. of segments = 125, 76, 30, 12
Path length[µm] = 131.1, 84.3, 36.7, 12.1
Vector sum length [µm] = 18.5, 28.4, 16.2, 4.4
15
10
5
0
−5
-10
-15
−20
−15
−10
−5
0
5
10

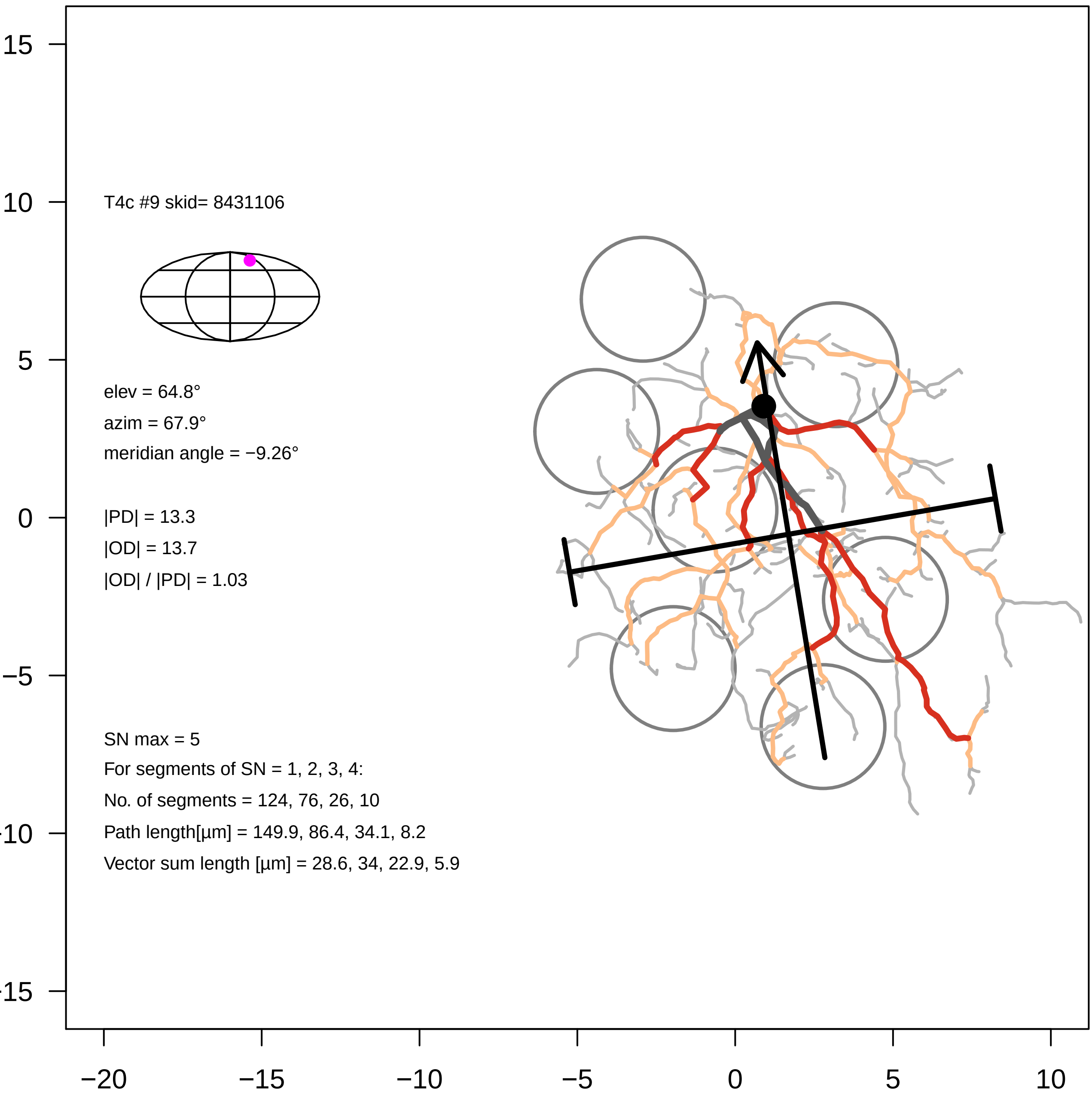

T4c #9 skid= 8431106
elev = 64.8°
azim = 67.9°
meridian angle = −9.26°
|PD| = 13.3
|OD| = 13.7
|OD| / |PD| = 1.03
SN max = 5
For segments of SN = 1, 2, 3, 4:
No. of segments = 124, 76, 26, 10
Path length[µm] = 149.9, 86.4, 34.1, 8.2
Vector sum length [µm] = 28.6, 34, 22.9, 5.9
15
10
5
0
−5
-10
-15
−20
−15
−10
−5
0
5
10

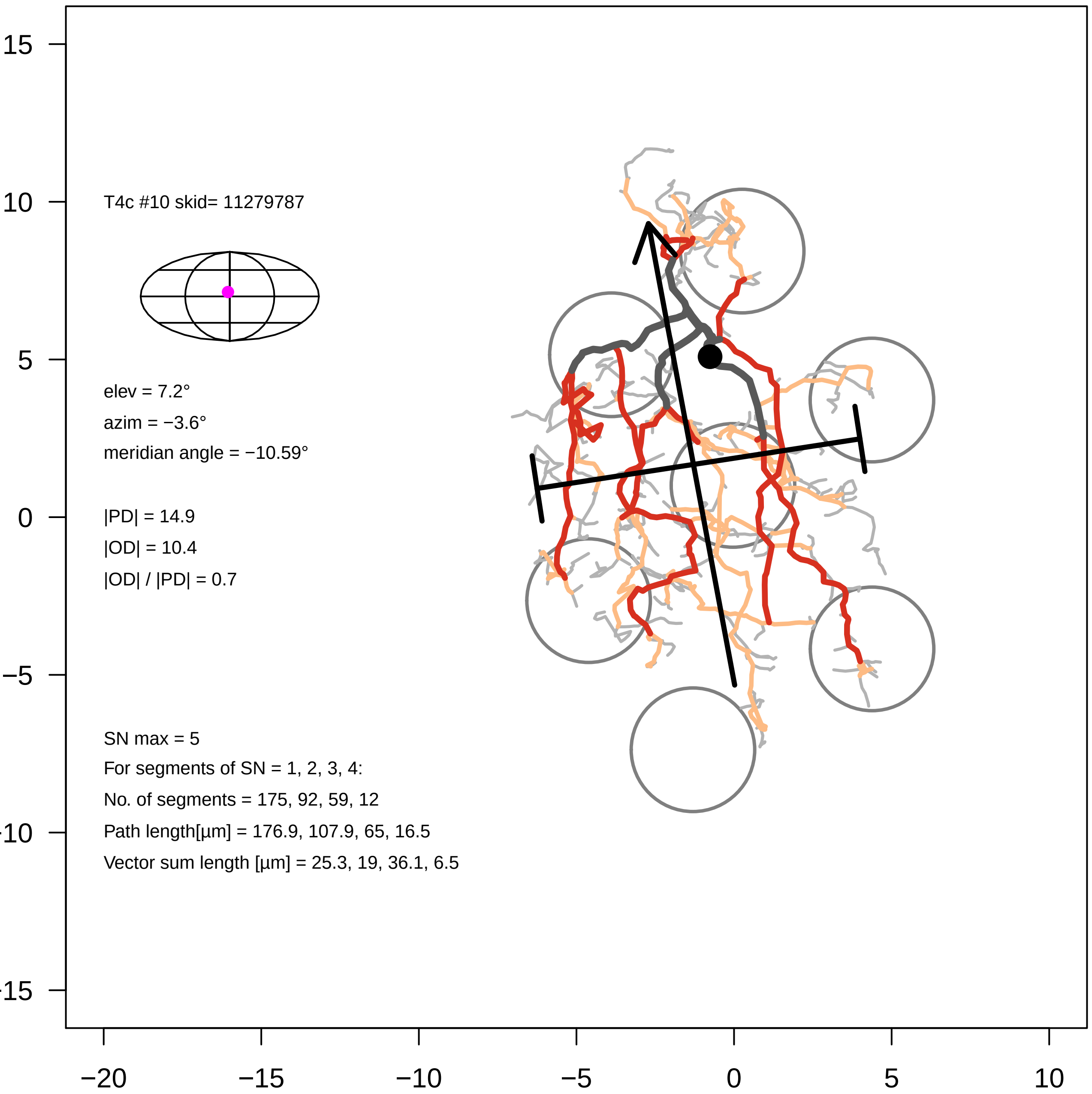

T4c #10 skid= 11279787
elev = 7.2°
azim = −3.6°
meridian angle = −10.59°
|PD| = 14.9
|OD| = 10.4
|OD| / |PD| = 0.7
SN max = 5
For segments of SN = 1, 2, 3, 4:
No. of segments = 175, 92, 59, 12
Path length[µm] = 176.9, 107.9, 65, 16.5
Vector sum length [µm] = 25.3, 19, 36.1, 6.5
15
10
5
0
−5
−10
−15
−20
−15
−10
−5
0
5
10

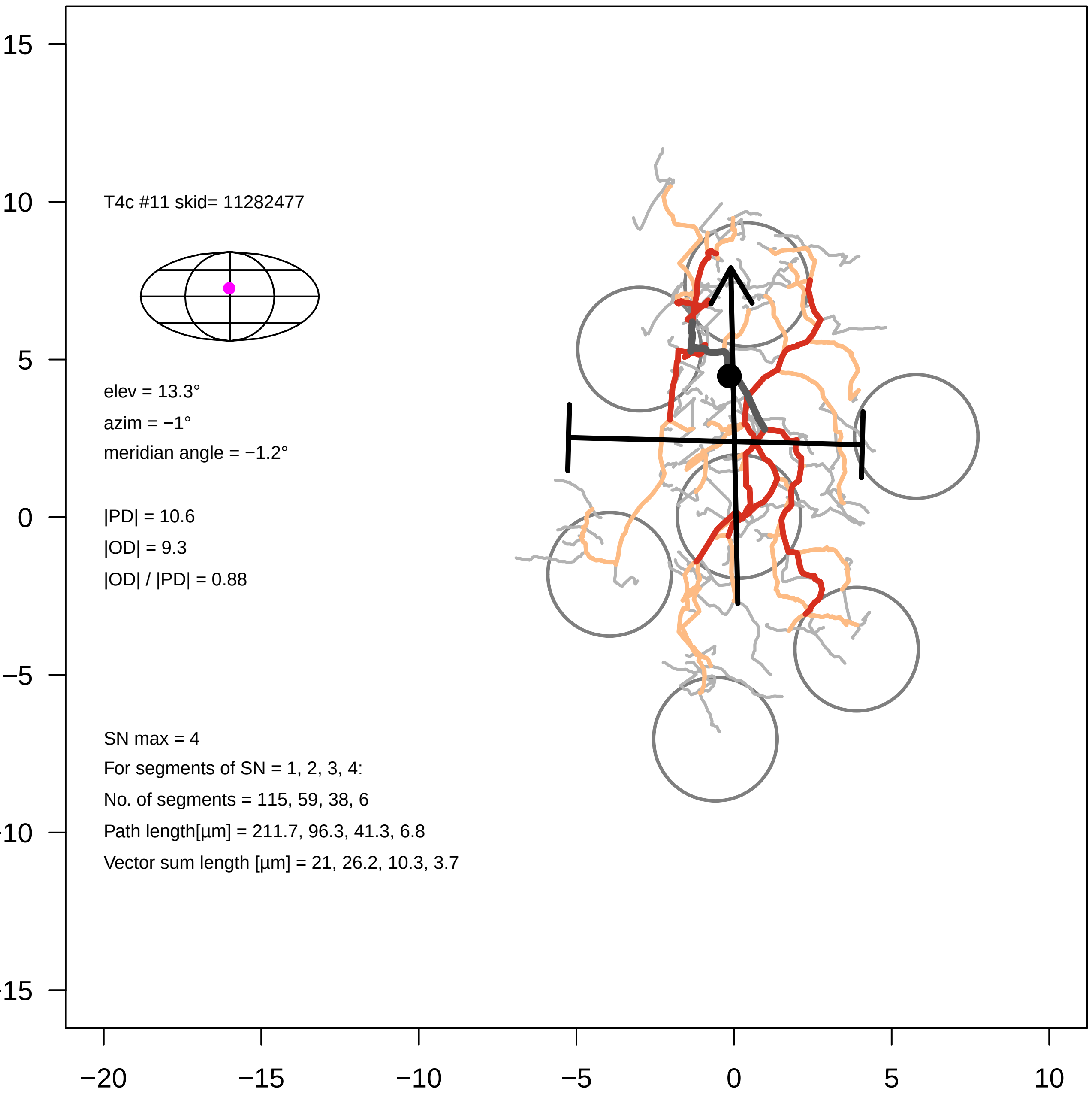

T4c #11 skid= 11282477
elev = 13.3°
azim = −1°
meridian angle = −1.2°
|PD| = 10.6
|OD| = 9.3
|OD| / |PD| = 0.88
SN max = 4
For segments of SN = 1, 2, 3, 4:
No. of segments = 115, 59, 38, 6
Path length[µm] = 211.7, 96.3, 41.3, 6.8
Vector sum length [µm] = 21, 26.2, 10.3, 3.7
15
10
5
0
−5
−10
−15
−20
−15
−10
−5
0
5
10

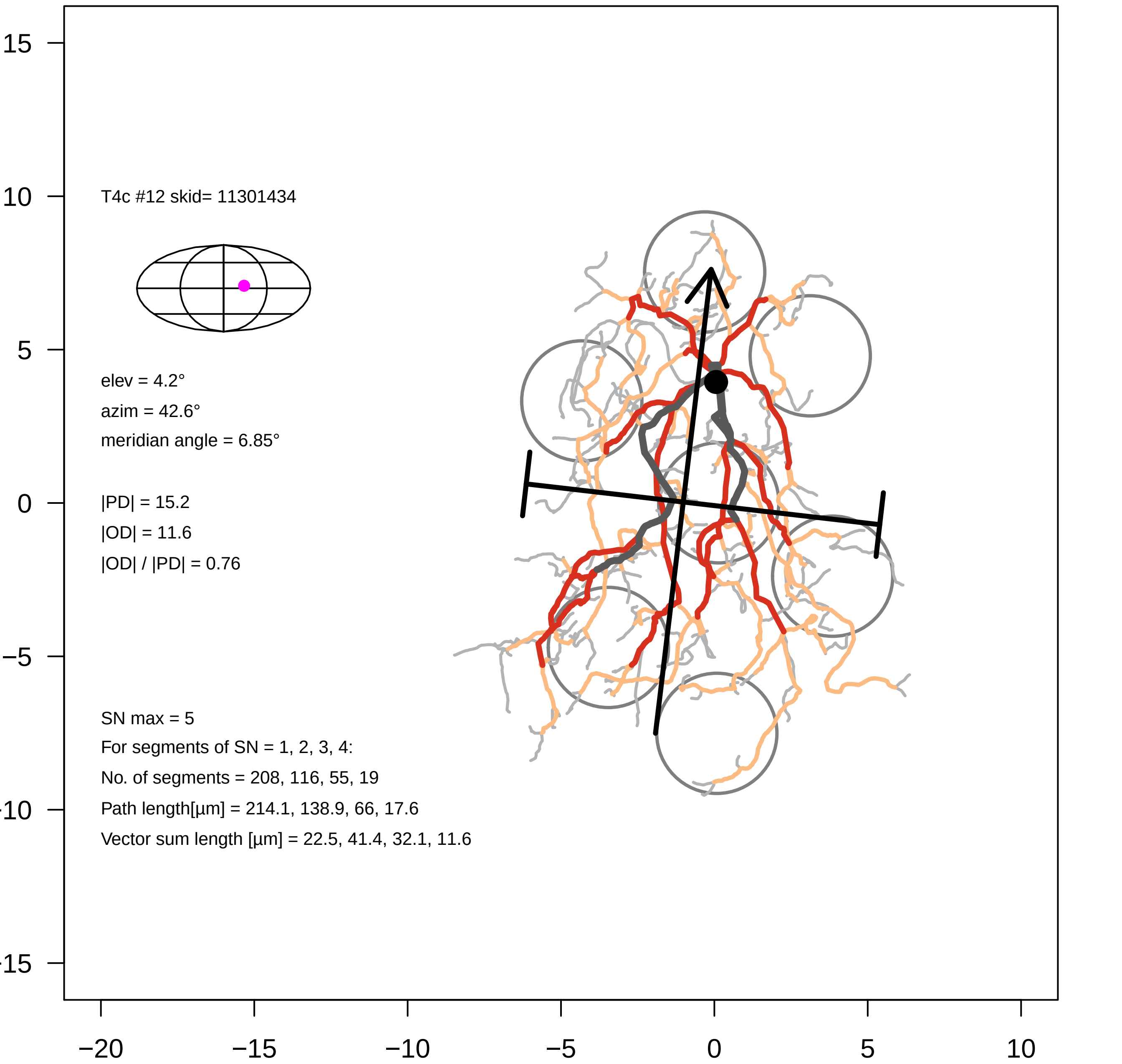

T4c #12 skid= 11301434
elev = 4.2°
azim = 42.6°
meridian angle = 6.85°
|PD| = 15.2
|OD| = 11.6
|OD| / |PD| = 0.76
SN max = 5
For segments of SN = 1, 2, 3, 4:
No. of segments = 208, 116, 55, 19
Path length[µm] = 214.1, 138.9, 66, 17.6
Vector sum length [µm] = 22.5, 41.4, 32.1, 11.6
15
10
5
0
−5
−10
−15
−20
−15
−10
−5
0
5
10

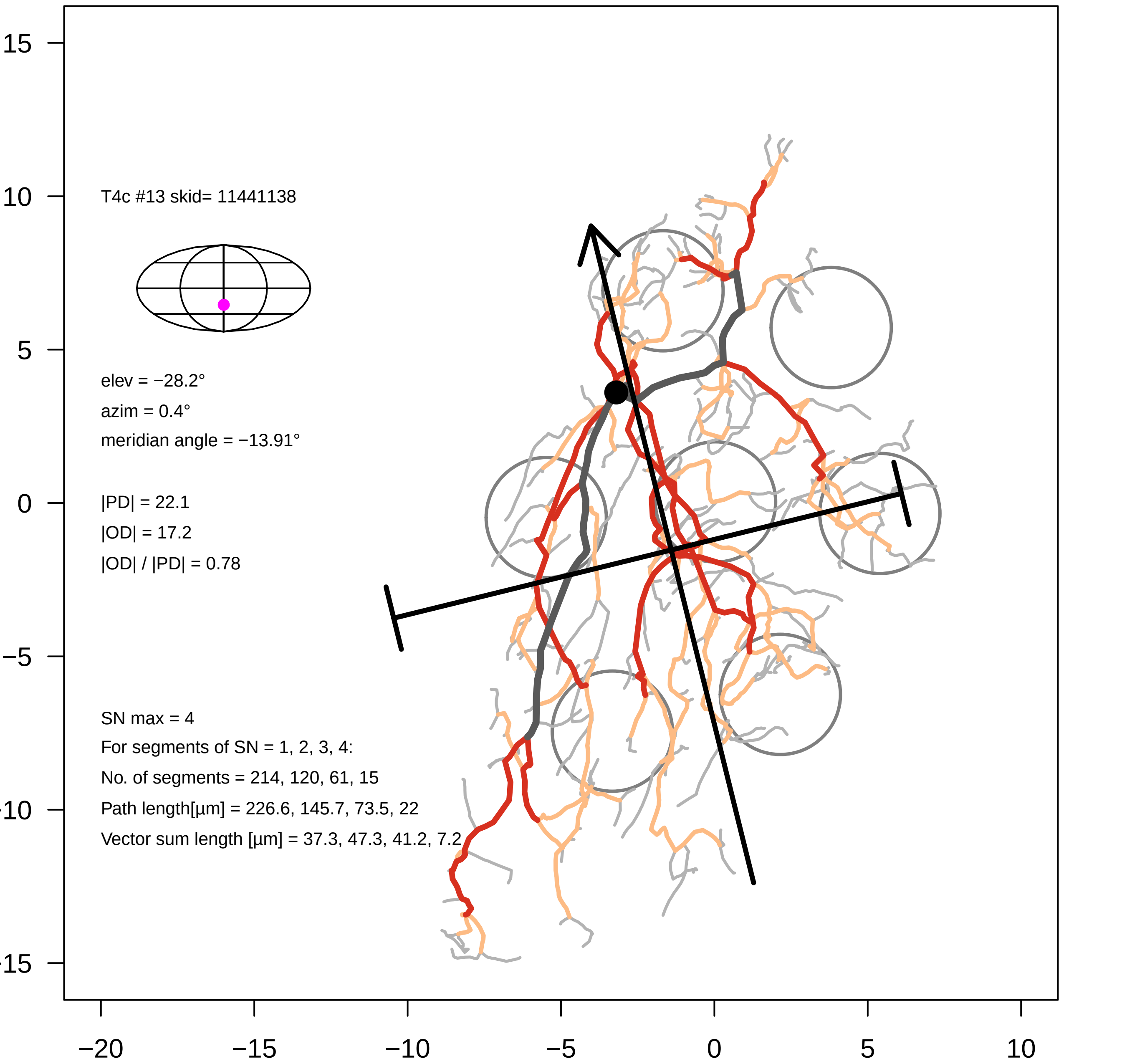

T4c #13 skid= 11441138
elev = −28.2°
azim = 0.4°
meridian angle = −13.91°
|PD| = 22.1
|OD| = 17.2
|OD| / |PD| = 0.78
SN max = 4
For segments of SN = 1, 2, 3, 4:
No. of segments = 214, 120, 61, 15
Path length[µm] = 226.6, 145.7, 73.5, 22
Vector sum length [µm] = 37.3, 47.3, 41.2, 7.2
15
10
5
0
−5
−10
−15
−20
−15
−10
−5
0
5
10

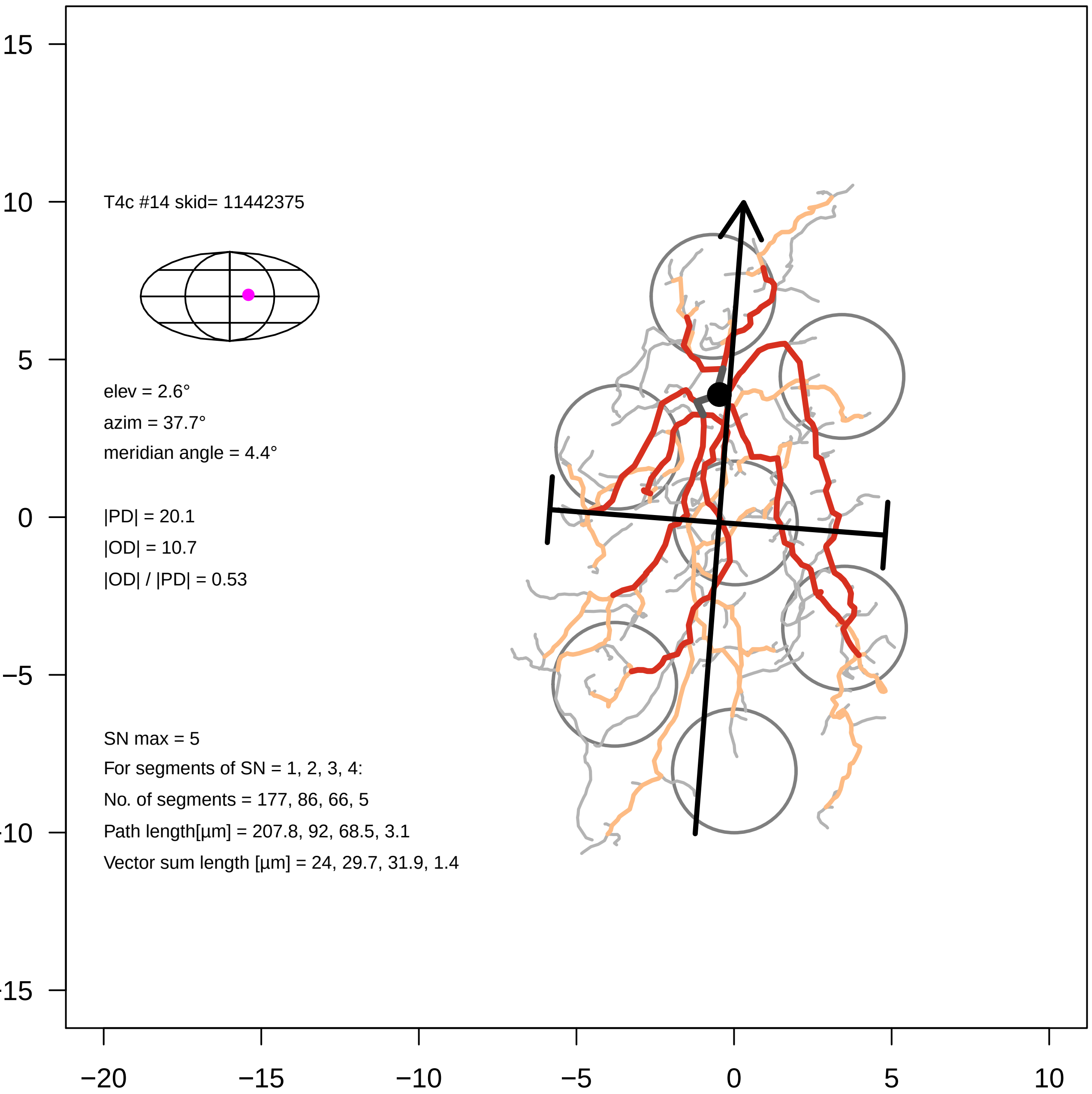

T4c #14 skid= 11442375
elev = 2.6°
azim = 37.7°
meridian angle = 4.4°
|PD| = 20.1
|OD| = 10.7
|OD| / |PD| = 0.53
SN max = 5
For segments of SN = 1, 2, 3, 4:
No. of segments = 177, 86, 66, 5
Path length[µm] = 207.8, 92, 68.5, 3.1
Vector sum length [µm] = 24, 29.7, 31.9, 1.4
15
10
5
0
−5
-10
-15
−20
−15
−10
−5
0
5
10

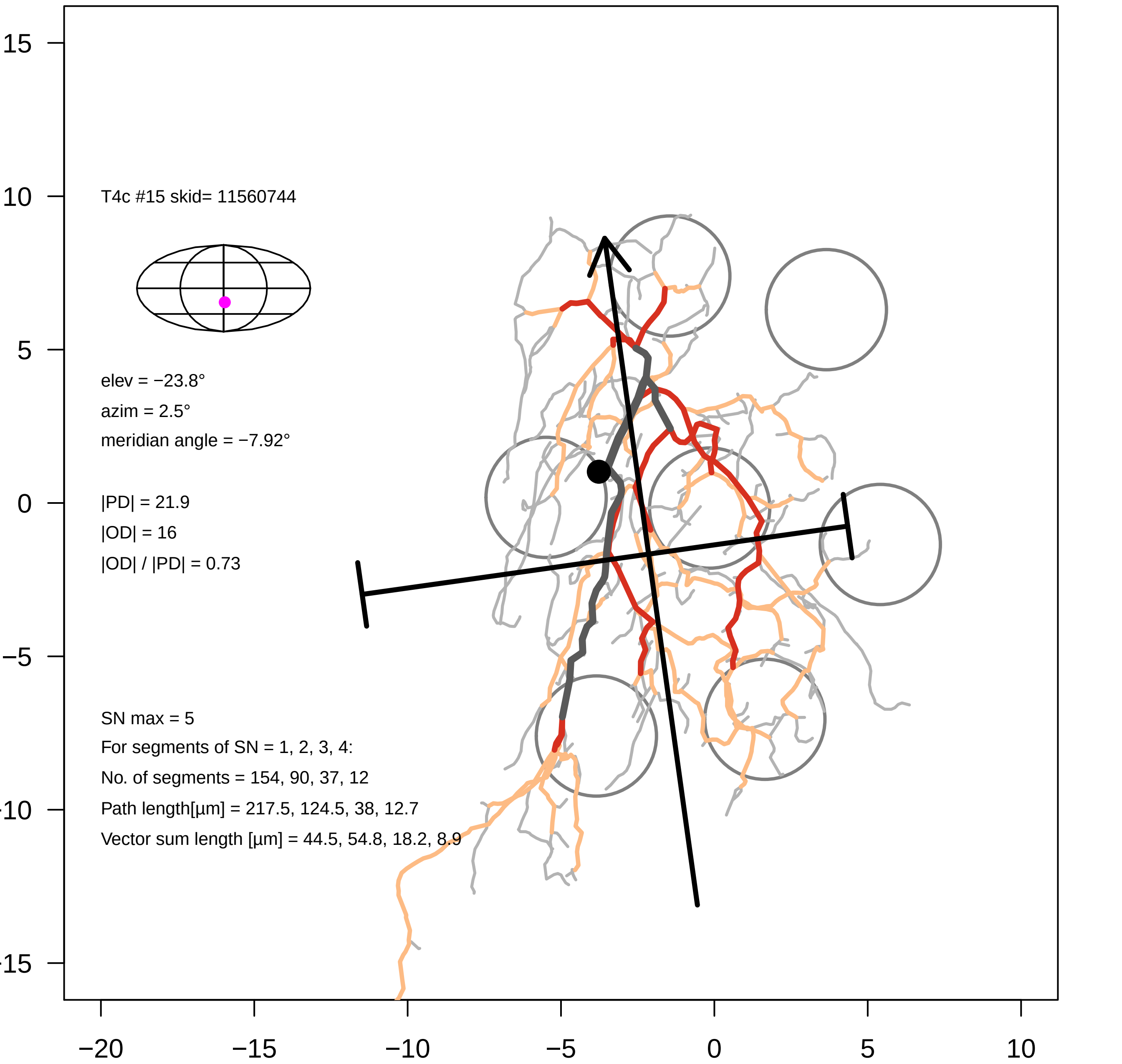

T4c #15 skid= 11560744
elev = −23.8°
azim = 2.5°
meridian angle = −7.92°
|PD| = 21.9
|OD| = 16
|OD| / |PD| = 0.73
SN max = 5
For segments of SN = 1, 2, 3, 4:
No. of segments = 154, 90, 37, 12
Path length[µm] = 217.5, 124.5, 38, 12.7
Vector sum length [µm] = 44.5, 54.8, 18.2, 8.9
15
10
5
0
−5
−10
−15
−20
−15
−10
−5
0
5
10

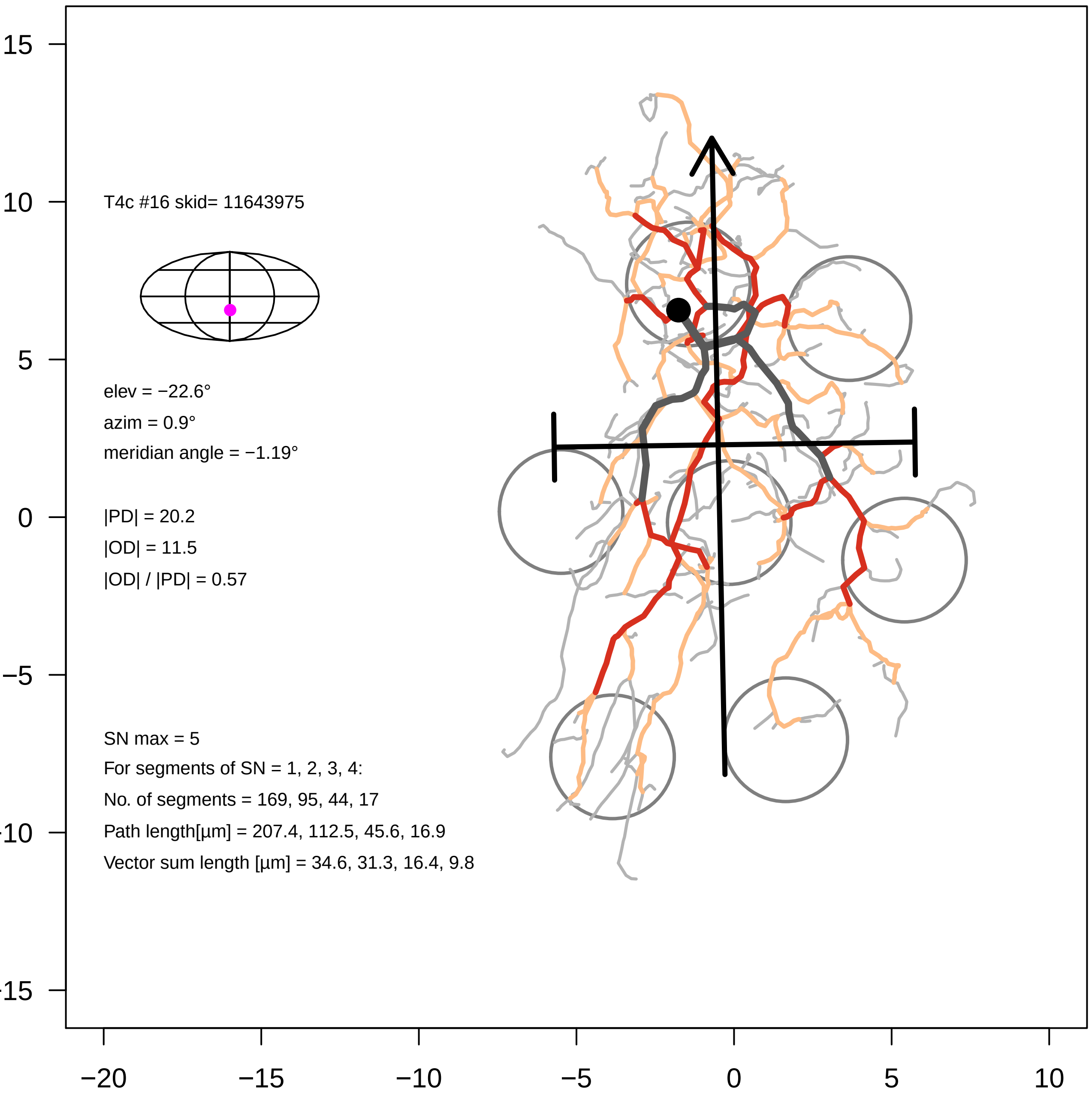

T4c #16 skid= 11643975
elev = −22.6°
azim = 0.9°
meridian angle = −1.19°
|PD| = 20.2
|OD| = 11.5
|OD| / |PD| = 0.57
SN max = 5
For segments of SN = 1, 2, 3, 4:
No. of segments = 169, 95, 44, 17
Path length[µm] = 207.4, 112.5, 45.6, 16.9
Vector sum length [µm] = 34.6, 31.3, 16.4, 9.8
15
10
5
0
−5
−10
−15
−20
−15
−10
−5
0
5
10

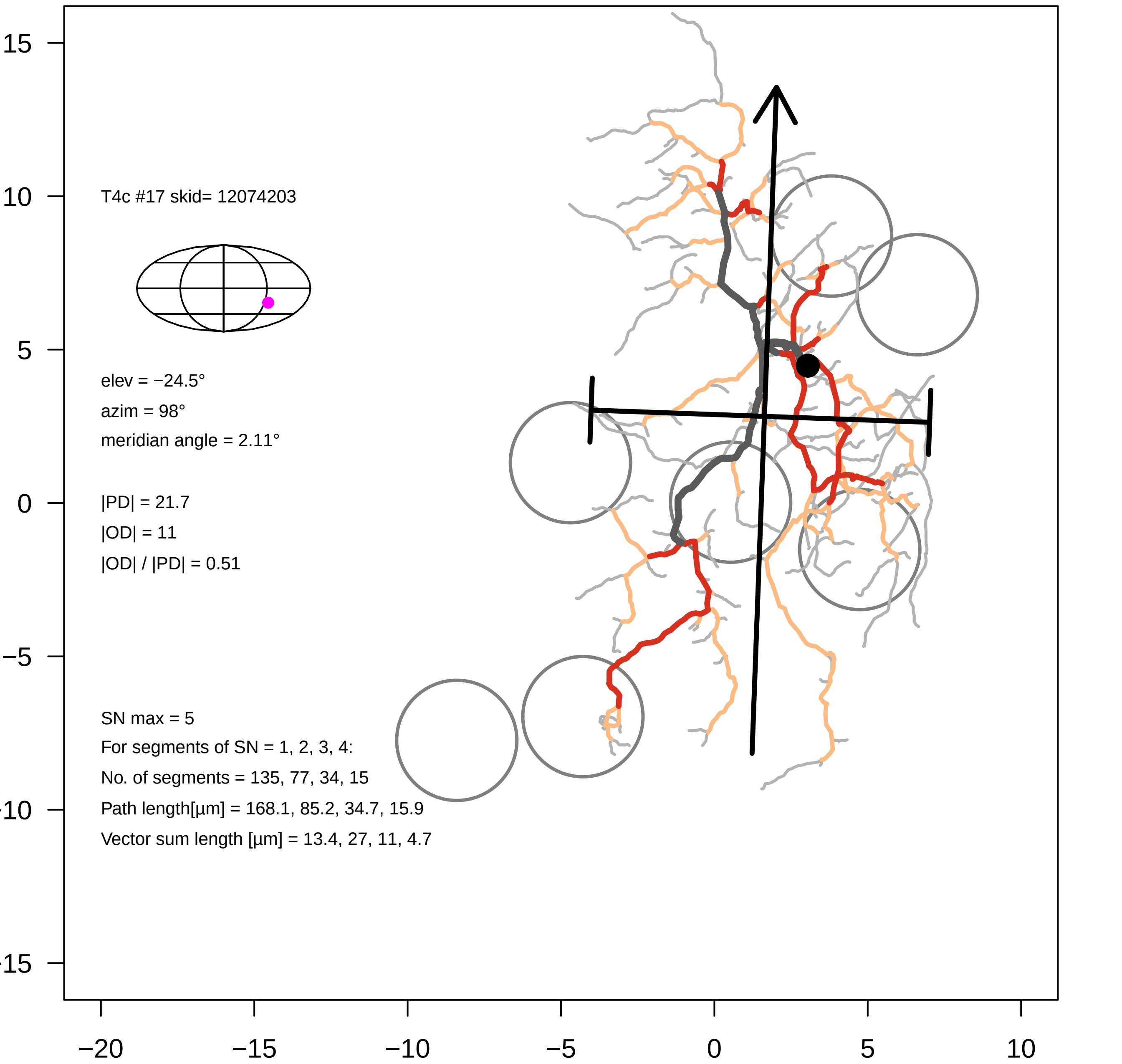

T4c #17 skid= 12074203
elev = −24.5°
azim = 98°
meridian angle = 2.11°
|PD| = 21.7
|OD| = 11
|OD| / |PD| = 0.51
SN max = 5
For segments of SN = 1, 2, 3, 4:
No. of segments = 135, 77, 34, 15
Path length[µm] = 168.1, 85.2, 34.7, 15.9
Vector sum length [µm] = 13.4, 27, 11, 4.7
15
10
5
0
−5
-10
-15
−20
−15
−10
−5
0
5
10

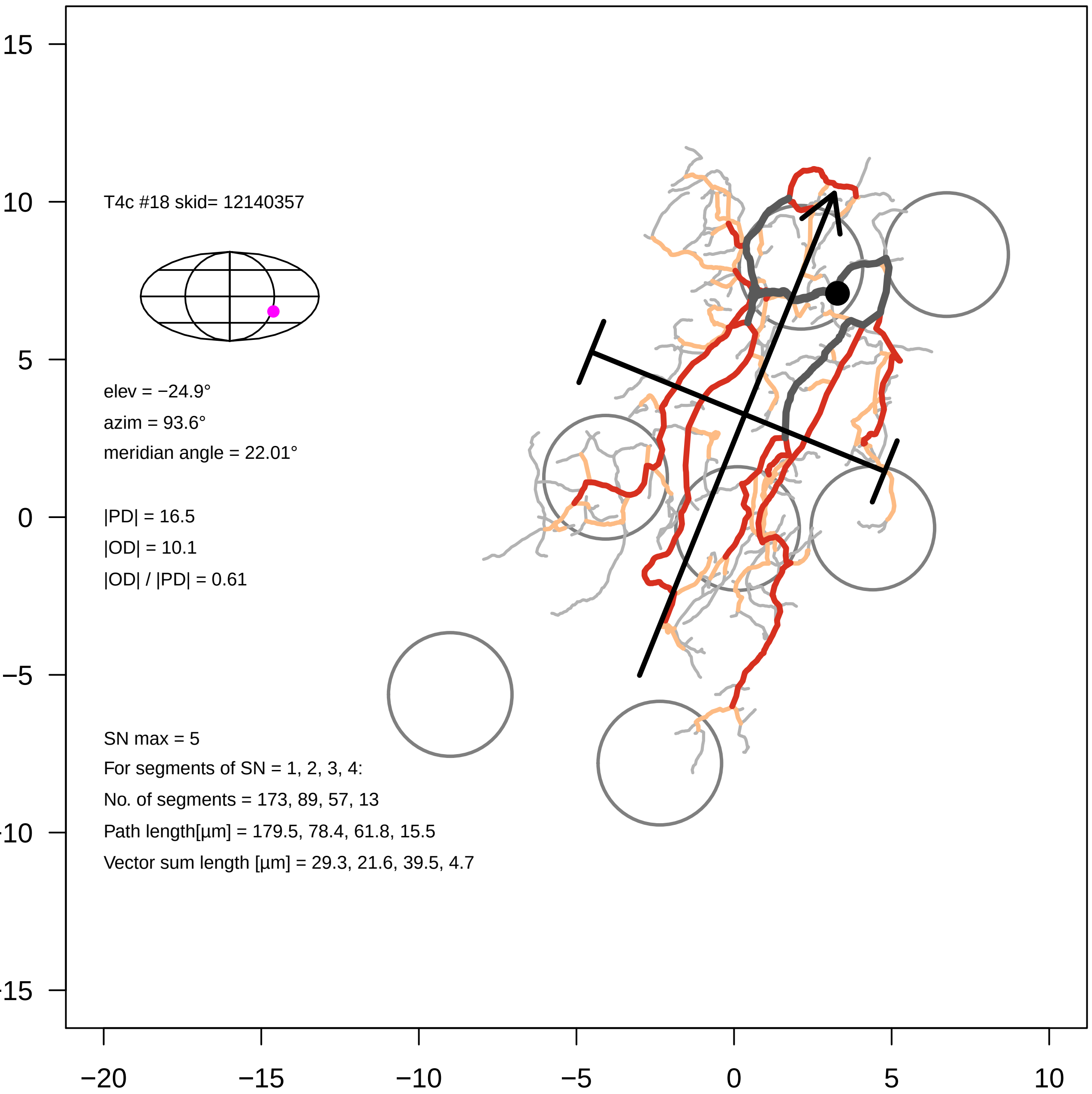
T4c #18 skid= 12140357
elev = −24.9°
azim = 93.6°
meridian angle = 22.01°
|PD| = 16.5
|OD| = 10.1
|OD| / |PD| = 0.61
SN max = 5
For segments of SN = 1, 2, 3, 4:
No. of segments = 173, 89, 57, 13
Path length[µm] = 179.5, 78.4, 61.8, 15.5
Vector sum length [µm] = 29.3, 21.6, 39.5, 4.7
15
10
5
0
−5
−10
−15
−20
−15
−10
−5
0
5
10

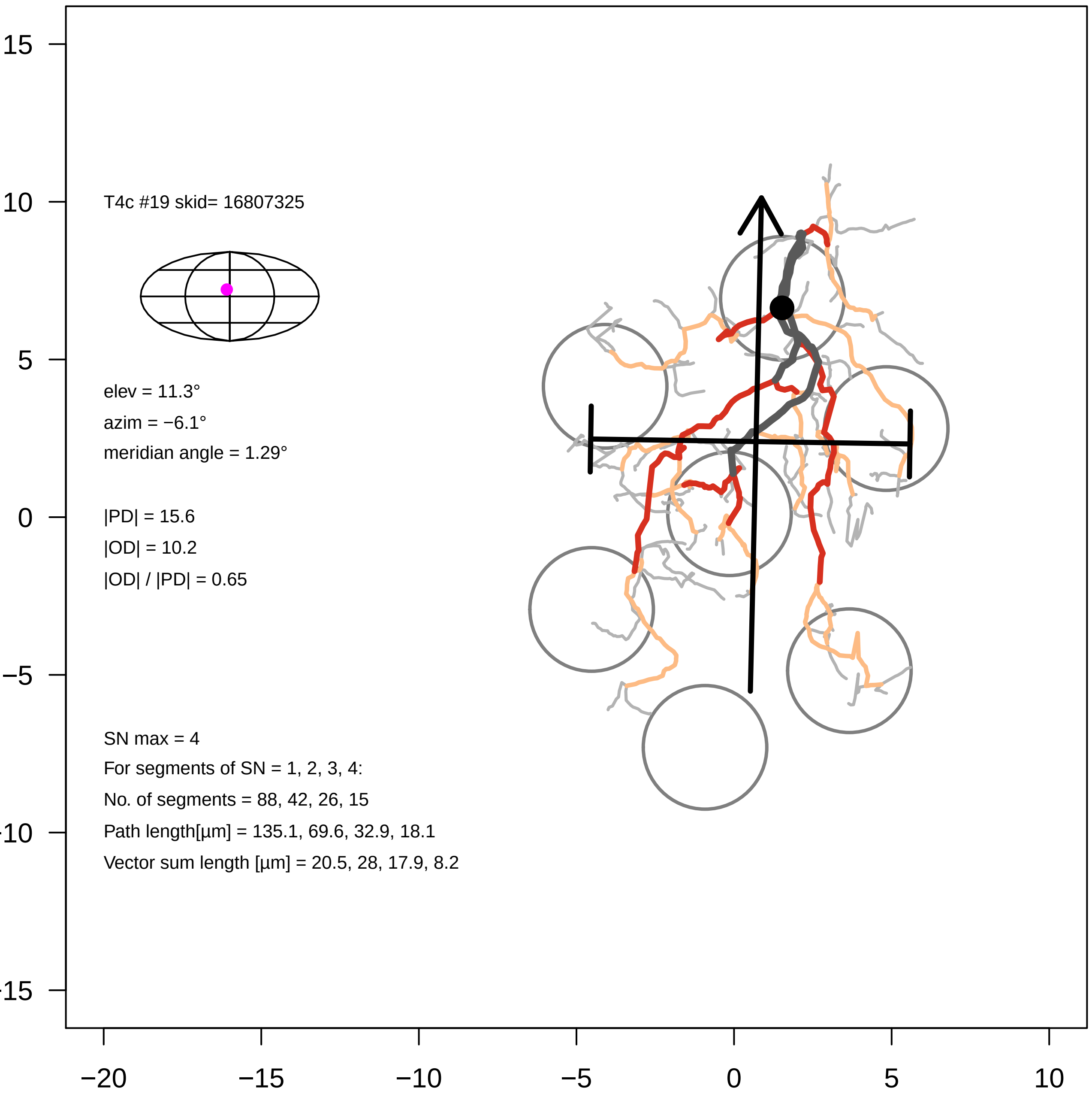

T4c #19 skid= 16807325
elev = 11.3°
azim = −6.1°
meridian angle = 1.29°
|PD| = 15.6
|OD| = 10.2
|OD| / |PD| = 0.65
SN max = 4
For segments of SN = 1, 2, 3, 4:
No. of segments = 88, 42, 26, 15
Path length[µm] = 135.1, 69.6, 32.9, 18.1
Vector sum length [µm] = 20.5, 28, 17.9, 8.2
15
10
5
0
−5
−10
−15
−20
−15
−10
−5
0
5
10

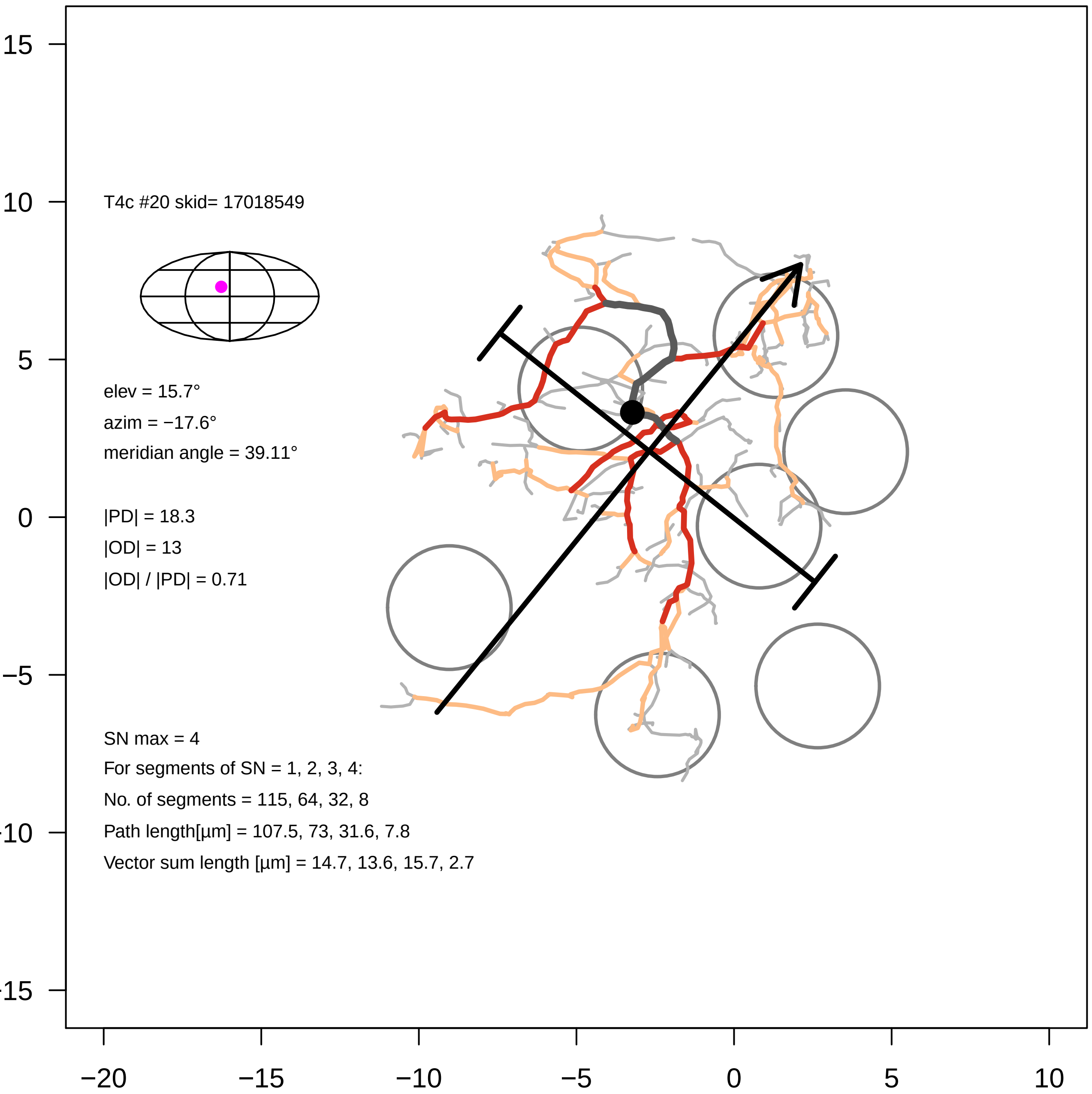

T4c #20 skid= 17018549
elev = 15.7°
azim = −17.6°
meridian angle = 39.11°
|PD| = 18.3
|OD| = 13
|OD| / |PD| = 0.71
SN max = 4
For segments of SN = 1, 2, 3, 4:
No. of segments = 115, 64, 32, 8
Path length[µm] = 107.5, 73, 31.6, 7.8
Vector sum length [µm] = 14.7, 13.6, 15.7, 2.7
15
10
5
0
−5
−10
−15
−20
−15
−10
−5
0
5
10

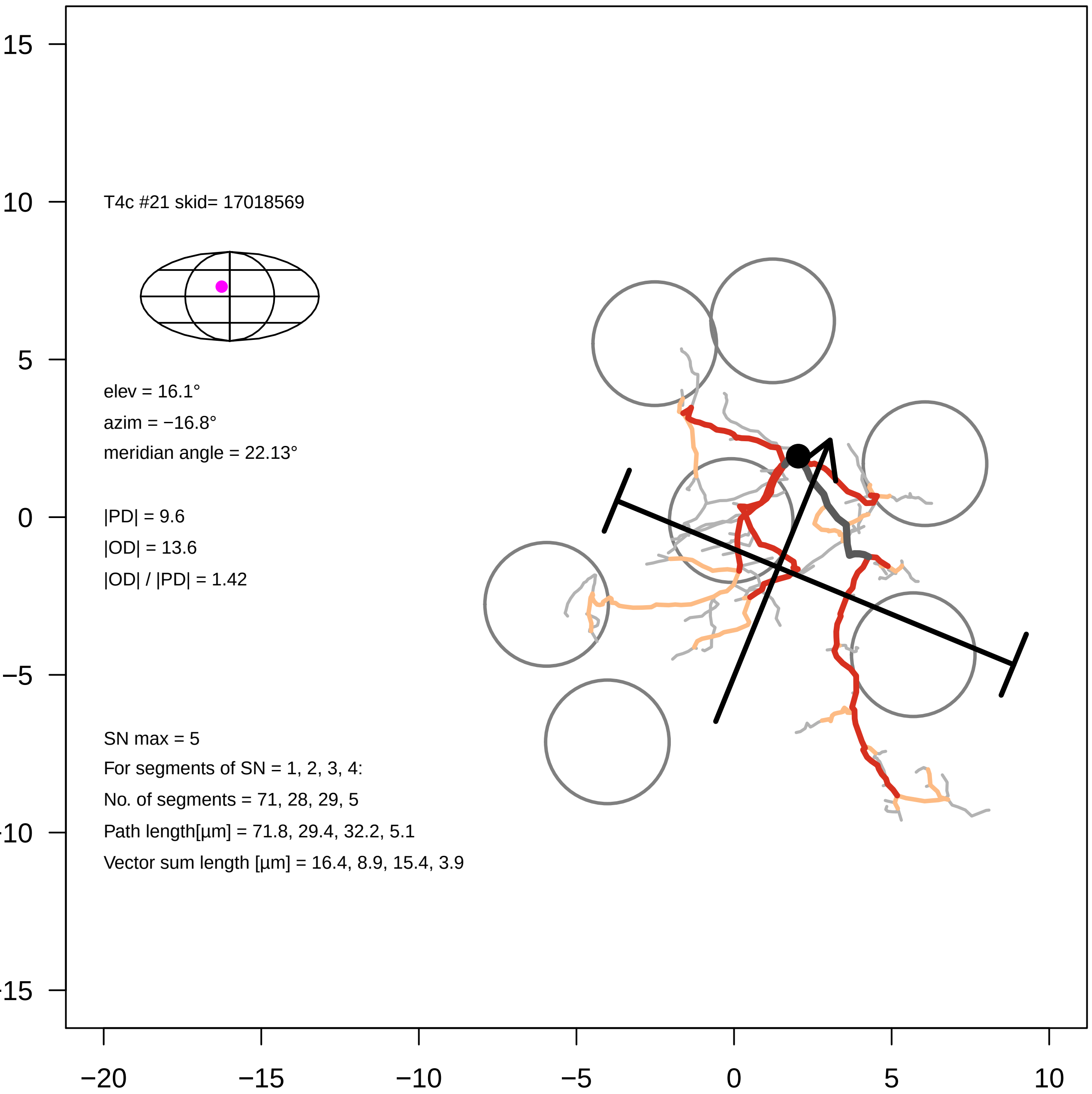

T4c #21 skid= 17018569
elev = 16.1°
azim = −16.8°
meridian angle = 22.13°
|PD| = 9.6
|OD| = 13.6
|OD| / |PD| = 1.42
SN max = 5
For segments of SN = 1, 2, 3, 4:
No. of segments = 71, 28, 29, 5
Path length[µm] = 71.8, 29.4, 32.2, 5.1
Vector sum length [µm] = 16.4, 8.9, 15.4, 3.9
15
10
5
0
−5
−10
−15
−20
−15
−10
−5
0
5
10

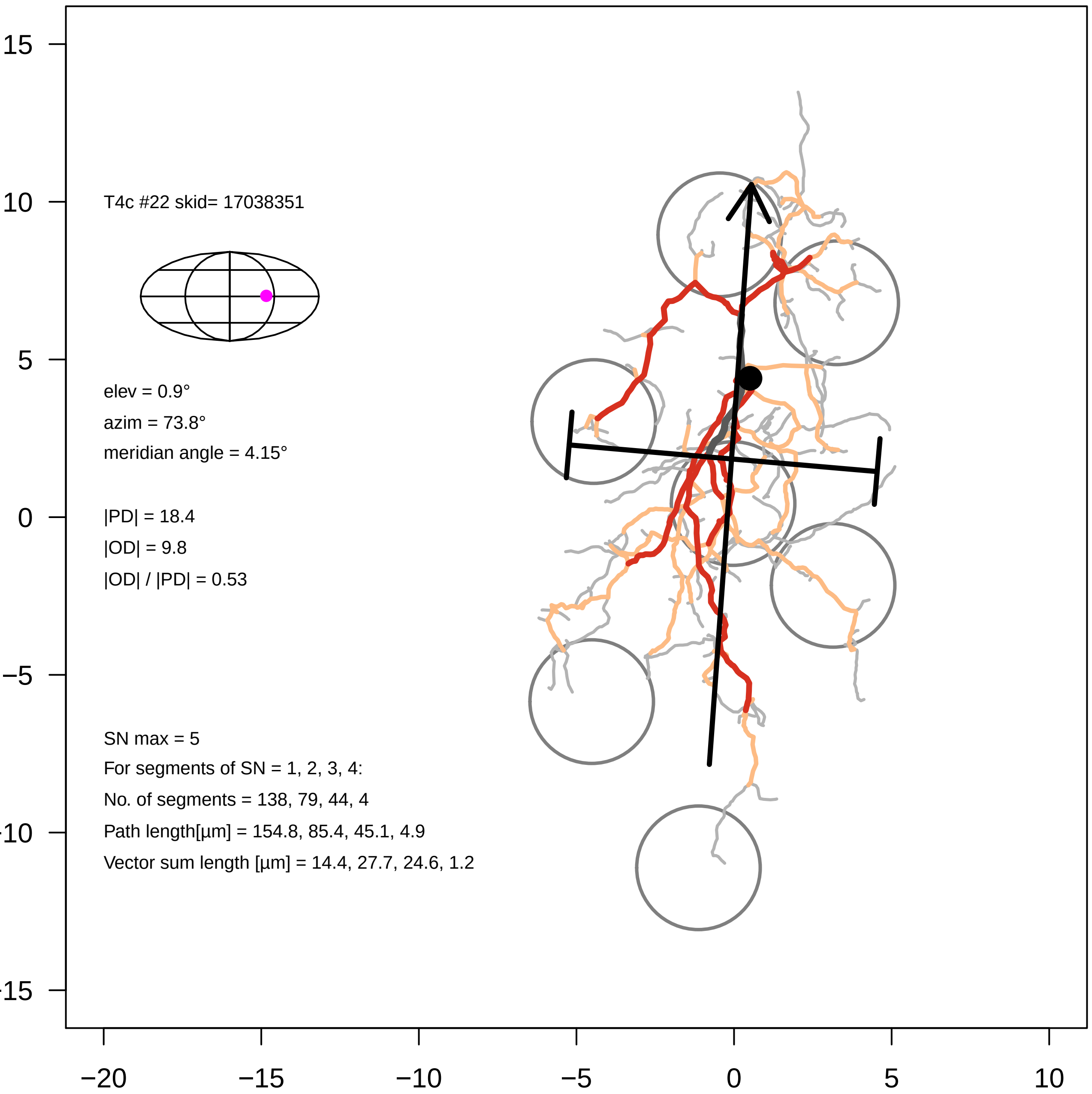
T4c #22 skid= 17038351
elev = 0.9°
azim = 73.8°
meridian angle = 4.15°
|PD| = 18.4
|OD| = 9.8
|OD| / |PD| = 0.53
SN max = 5
For segments of SN = 1, 2, 3, 4:
No. of segments = 138, 79, 44, 4
Path length[µm] = 154.8, 85.4, 45.1, 4.9
Vector sum length [µm] = 14.4, 27.7, 24.6, 1.2
15
10
5
0
−5
−10
−15
−20
−15
−10
−5
0
5
10
