## Supplementary Files, T4 galleries for "Eye structure shapes neuron function in *Drosophila* motion vision": Supplementary_Data_File4_T4d.pdf

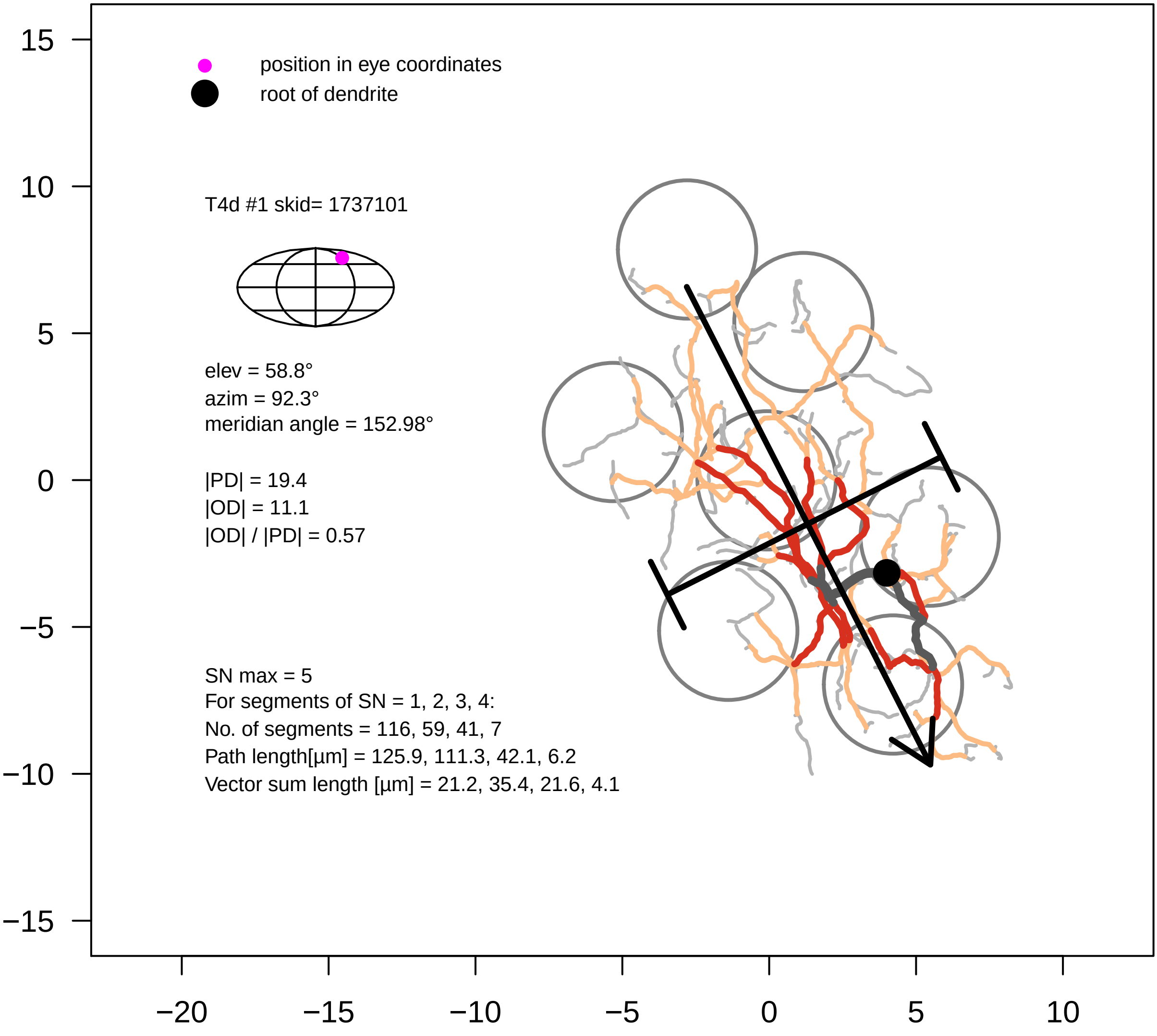

position in eye coordinates
root of dendrite
T4d #1 skid= 1737101
elev = 58.8°
azim = 92.3°
meridian angle = 152.98°
|PD| = 19.4
|OD| = 11.1
|OD| / |PD| = 0.57
SN max = 5
For segments of SN = 1, 2, 3, 4:
No. of segments = 116, 59, 41, 7
Path length[µm] = 125.9, 111.3, 42.1, 6.2
Vector sum length [µm] = 21.2, 35.4, 21.6, 4.1
15
10
5
0
−5
−10
−15
−20
−15
−10
−5
0
5
10

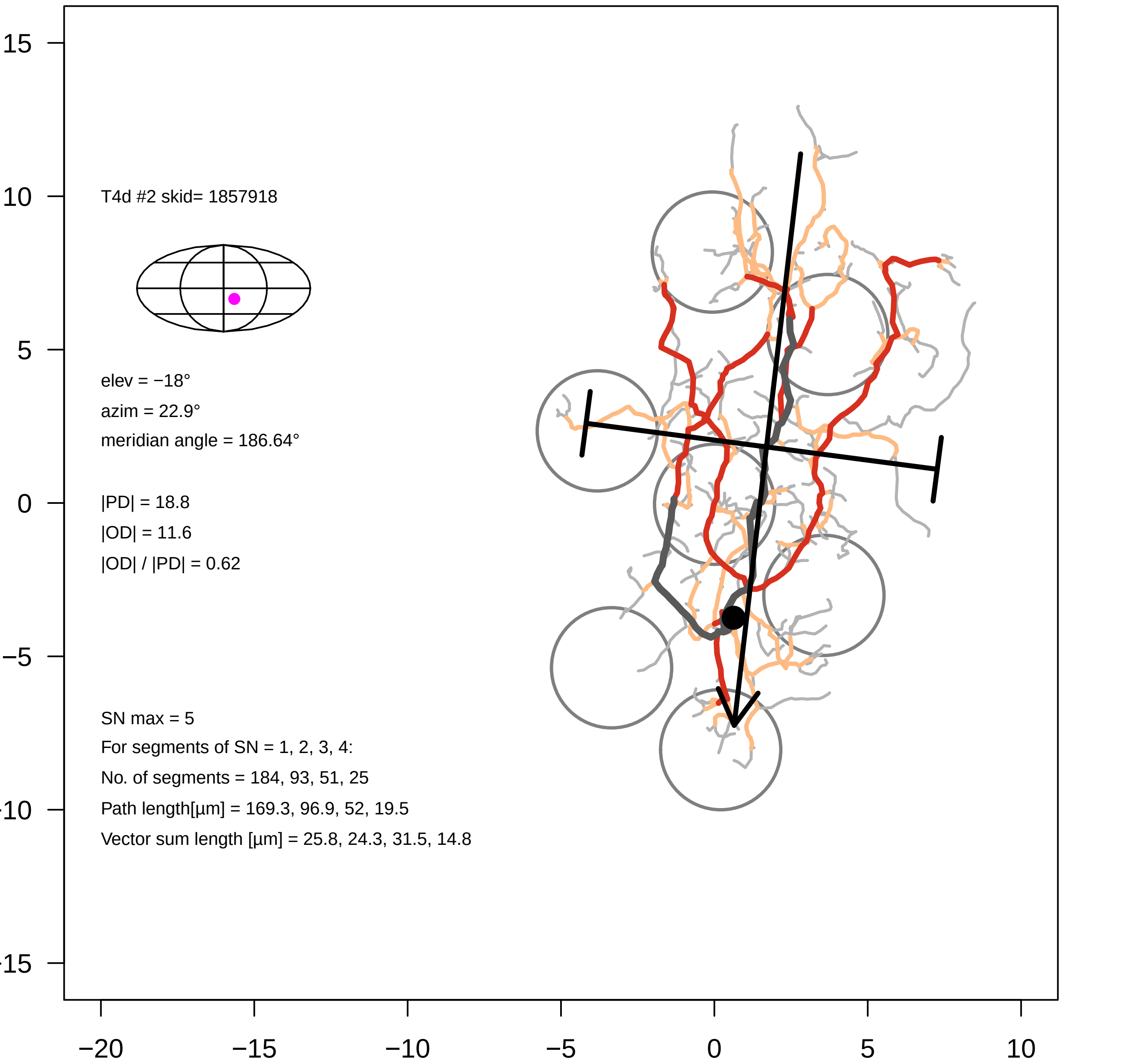
T4d #2 skid= 1857918
elev = −18°
azim = 22.9°
meridian angle = 186.64°
|PD| = 18.8
|OD| = 11.6
|OD| / |PD| = 0.62
SN max = 5
For segments of SN = 1, 2, 3, 4:
No. of segments = 184, 93, 51, 25
Path length[µm] = 169.3, 96.9, 52, 19.5
Vector sum length [µm] = 25.8, 24.3, 31.5, 14.8
15
10
5
0
−5
−10
−15
−20
−15
−10
−5
0
5
10

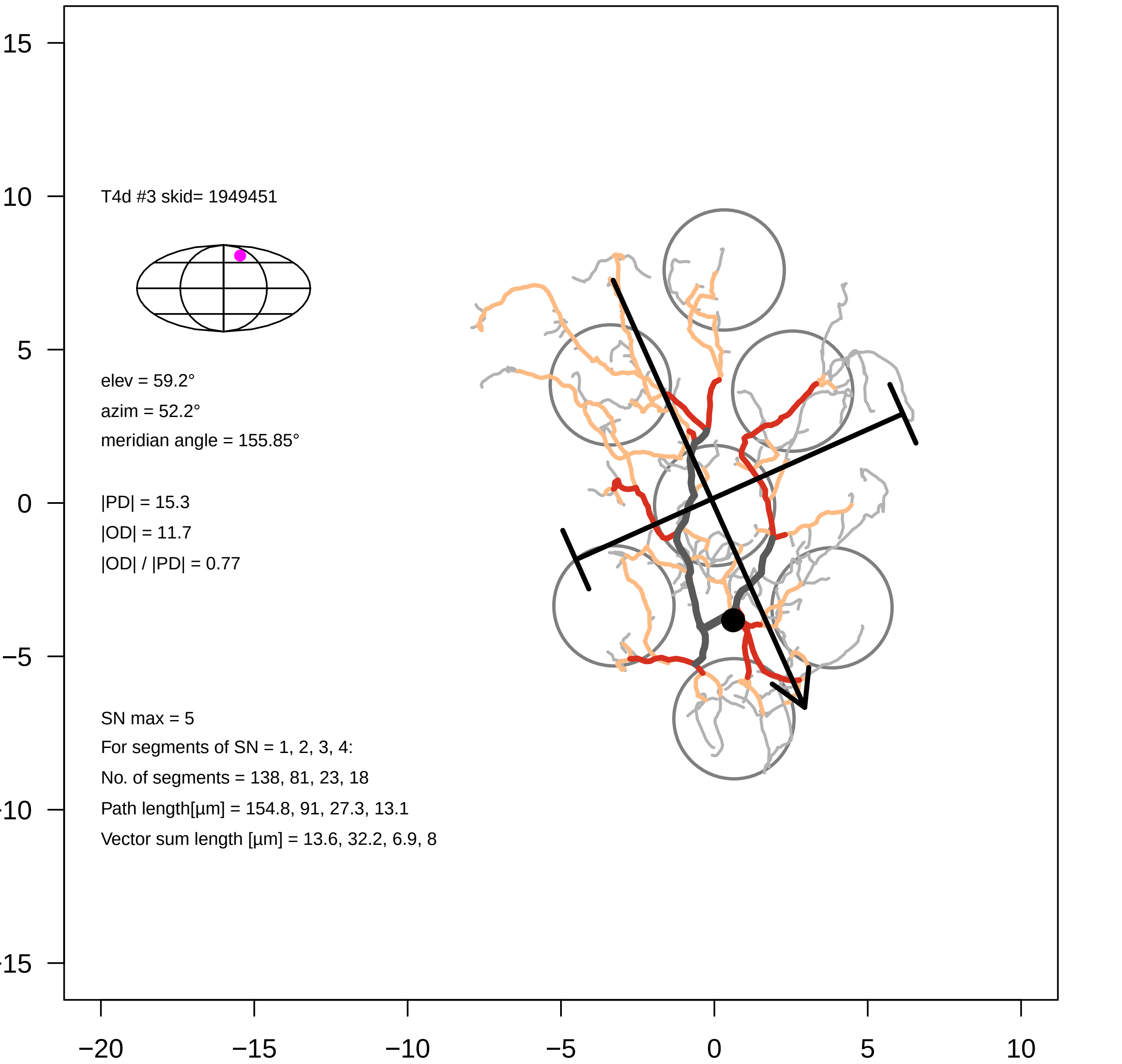

T4d #3 skid= 1949451
elev = 59.2°
azim = 52.2°
meridian angle = 155.85°
|PD| = 15.3
|OD| = 11.7
|OD| / |PD| = 0.77
SN max = 5
For segments of SN = 1, 2, 3, 4:
No. of segments = 138, 81, 23, 18
Path length[µm] = 154.8, 91, 27.3, 13.1
Vector sum length [µm] = 13.6, 32.2, 6.9, 8
15
10
5
0
−5
−20
−15
−10
−5
0
5
10

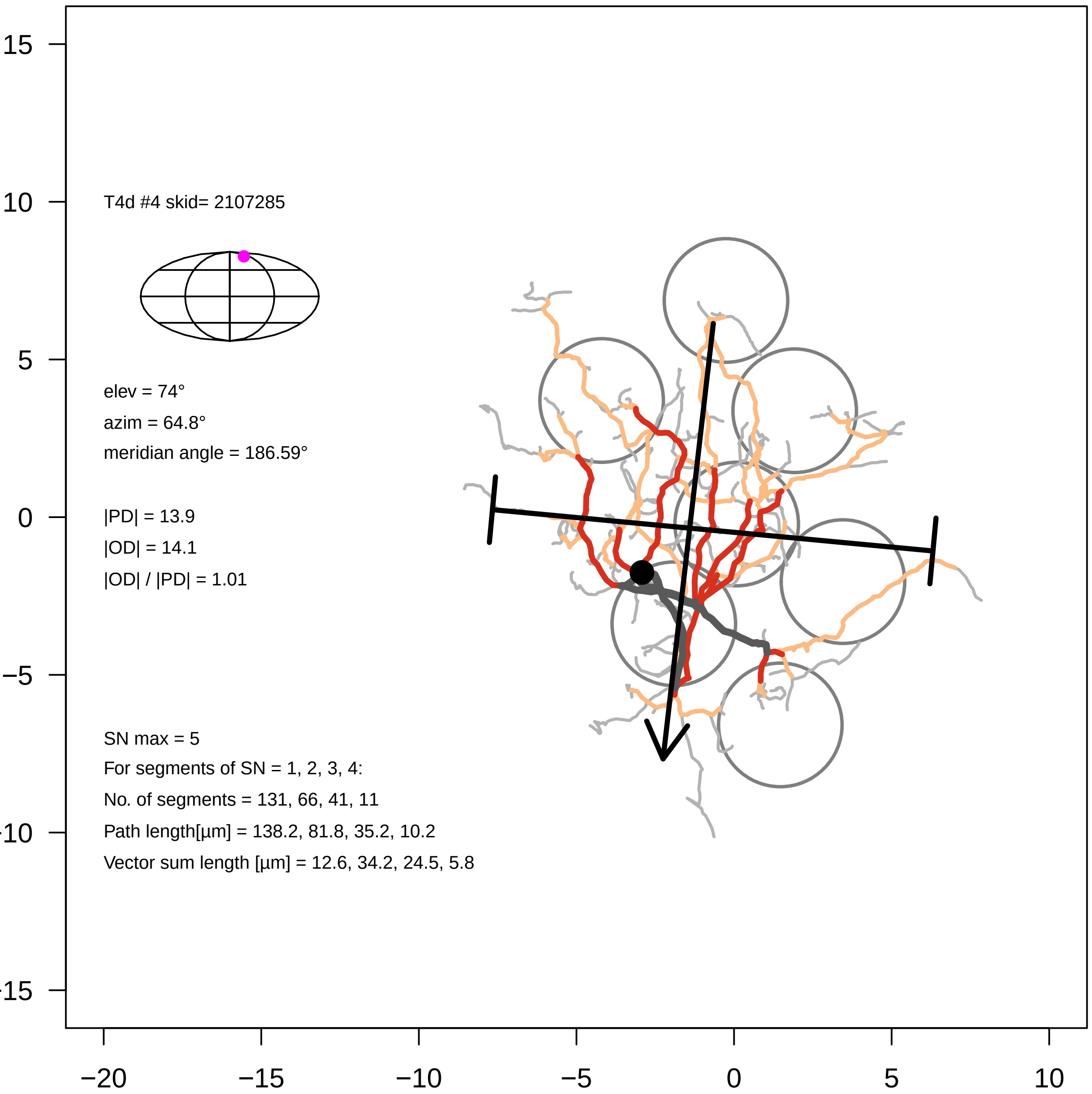

T4d #4 skid= 2107285
elev = 74°
azim = 64.8°
meridian angle = 186.59°
|PD| = 13.9
|OD| = 14.1
|OD| / |PD| = 1.01
SN max = 5
For segments of SN = 1, 2, 3, 4:
No. of segments = 131, 66, 41, 11
Path length[µm] = 138.2, 81.8, 35.2, 10.2
Vector sum length [µm] = 12.6, 34.2, 24.5, 5.8
15
10
5
0
−5
−10
−15
−20
−15
−10
−5
0
5
10

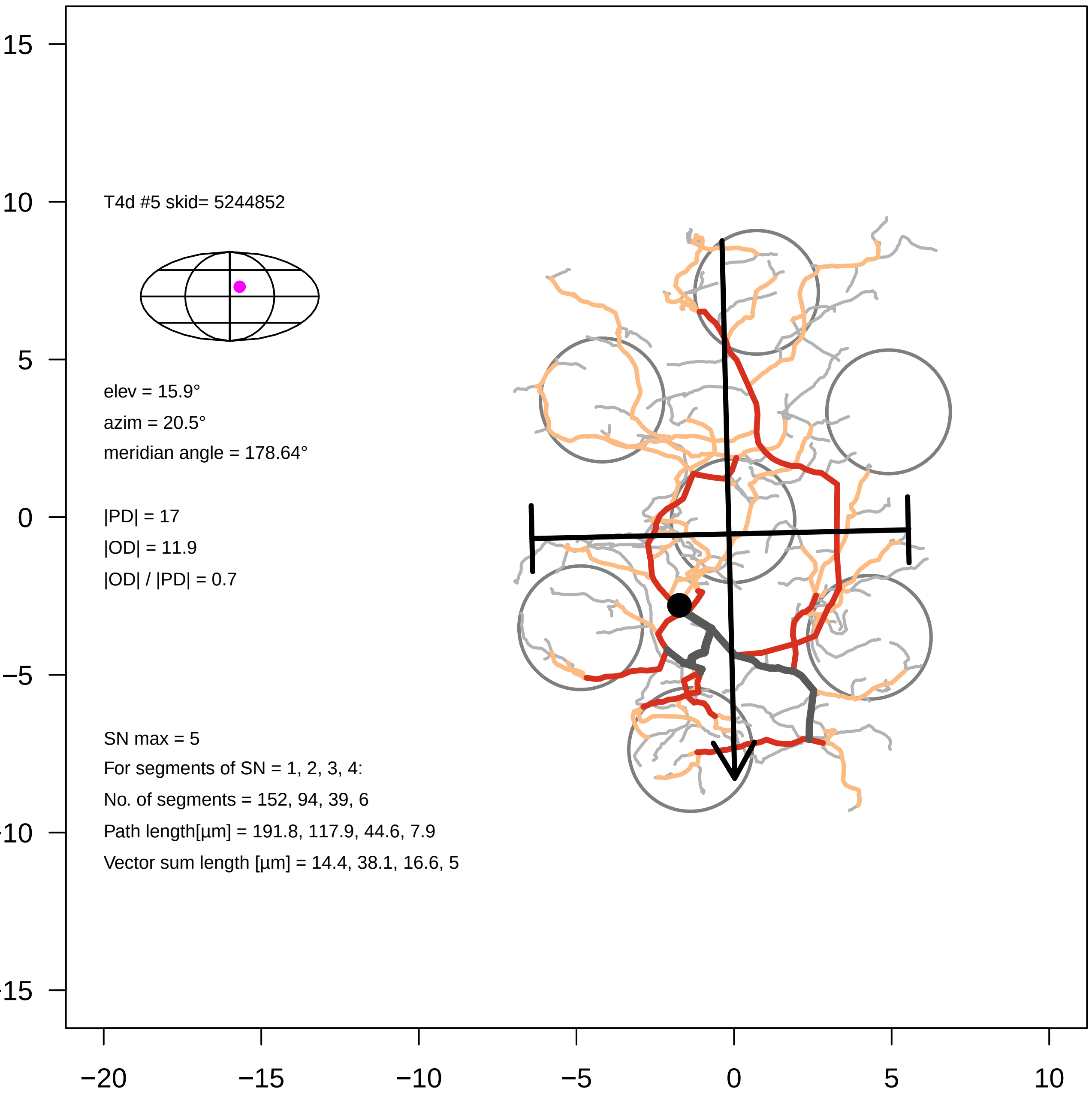
T4d #5 skid= 5244852
elev = 15.9°
azim = 20.5°
meridian angle = 178.64°
|PD| = 17
|OD| = 11.9
|OD| / |PD| = 0.7
SN max = 5
For segments of SN = 1, 2, 3, 4:
No. of segments = 152, 94, 39, 6
Path length[µm] = 191.8, 117.9, 44.6, 7.9
Vector sum length [µm] = 14.4, 38.1, 16.6, 5
15
10
5
0
−5
−20
−15
−10
−5
0
5
10

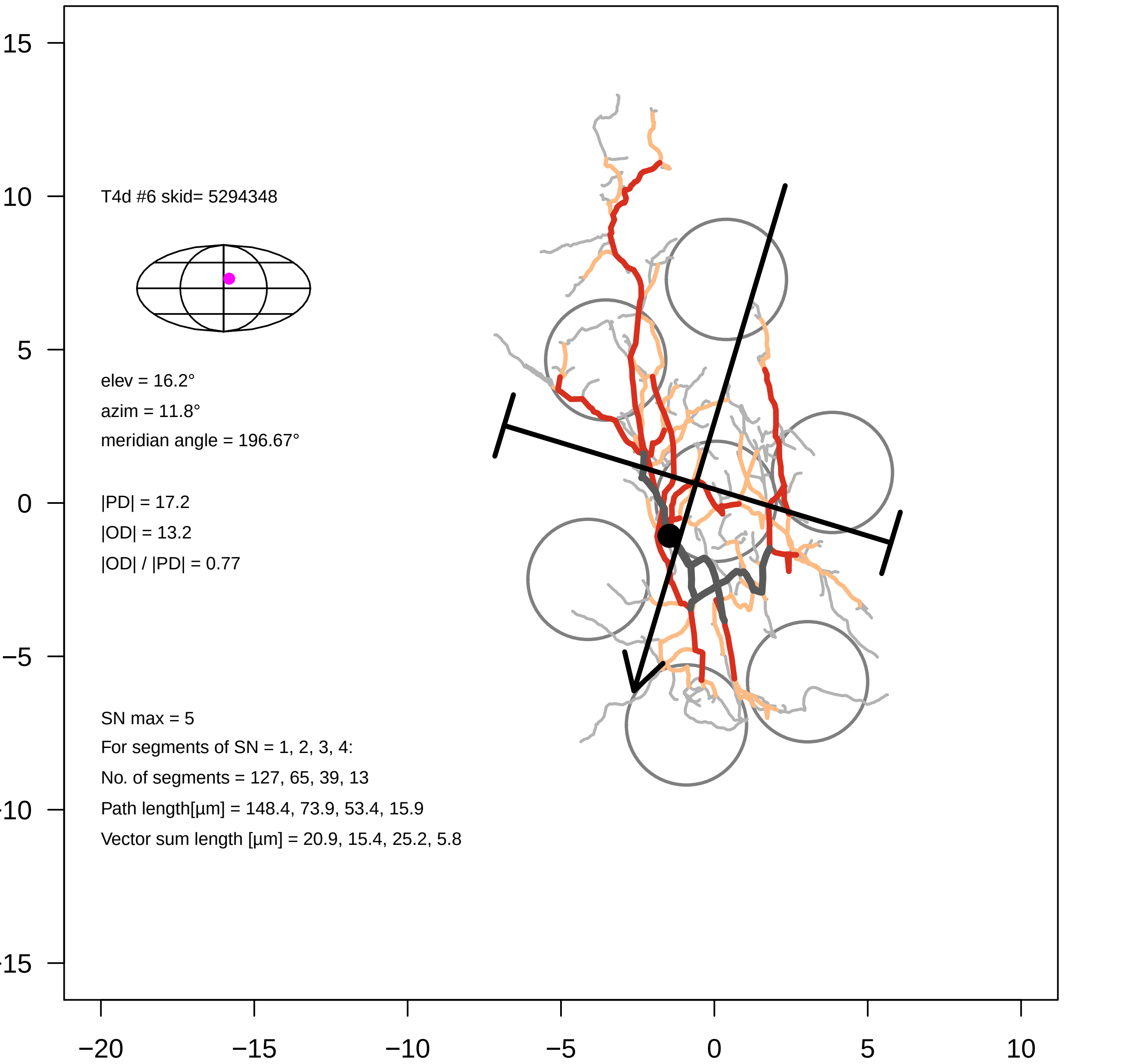
T4d #6 skid= 5294348
elev = 16.2°
azim = 11.8°
meridian angle = 196.67°
|PD| = 17.2
|OD| = 13.2
|OD| / |PD| = 0.77
SN max = 5
For segments of SN = 1, 2, 3, 4:
No. of segments = 127, 65, 39, 13
Path length[µm] = 148.4, 73.9, 53.4, 15.9
Vector sum length [µm] = 20.9, 15.4, 25.2, 5.8
15
10
5
0
−5
−10
−15
−20
−15
−10
−5
0
5
10

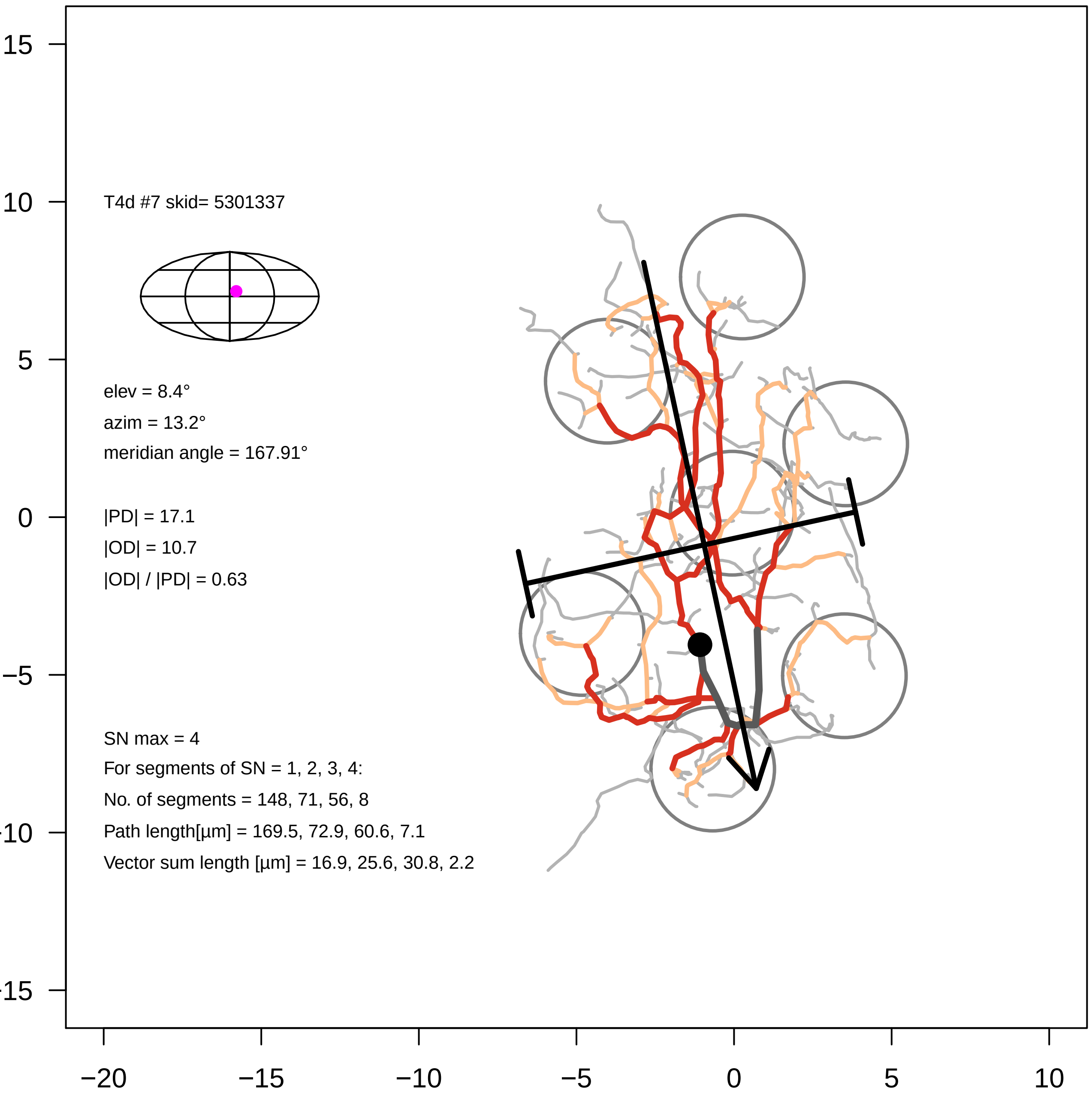

T4d #7 skid= 5301337
elev = 8.4°
azim = 13.2°
meridian angle = 167.91°
|PD| = 17.1
|OD| = 10.7
|OD| / |PD| = 0.63
SN max = 4
For segments of SN = 1, 2, 3, 4:
No. of segments = 148, 71, 56, 8
Path length[µm] = 169.5, 72.9, 60.6, 7.1
Vector sum length [µm] = 16.9, 25.6, 30.8, 2.2
15
10
5
0
−5
−20
−15
−10
−5
0
5
10

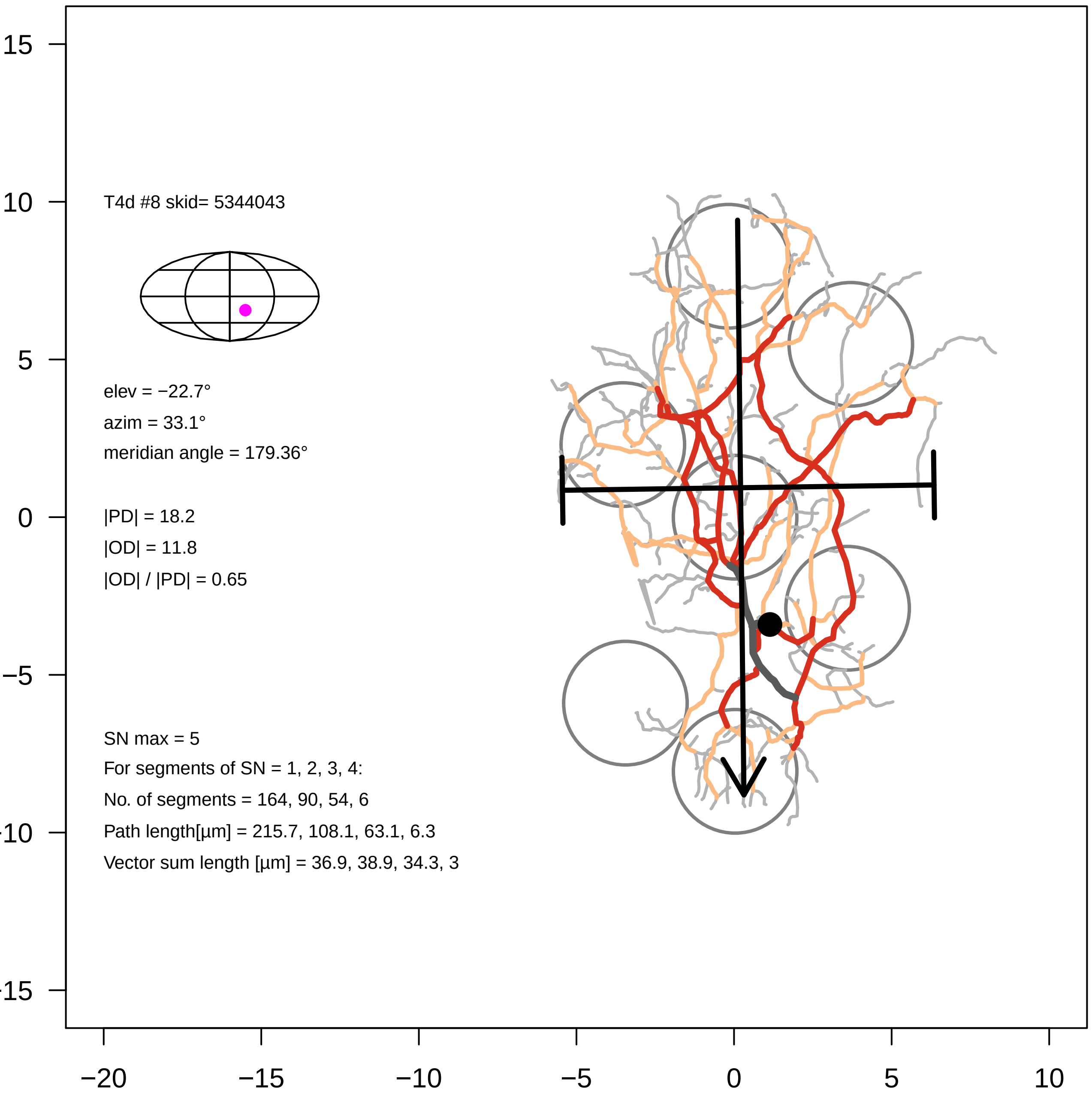
T4d #8 skid= 5344043
elev = −22.7°
azim = 33.1°
meridian angle = 179.36°
|PD| = 18.2
|OD| = 11.8
|OD| / |PD| = 0.65
SN max = 5
For segments of SN = 1, 2, 3, 4:
No. of segments = 164, 90, 54, 6
Path length[µm] = 215.7, 108.1, 63.1, 6.3
Vector sum length [µm] = 36.9, 38.9, 34.3, 3
15
10
5
0
−5
−20
−15
−10
−5
0
5
10

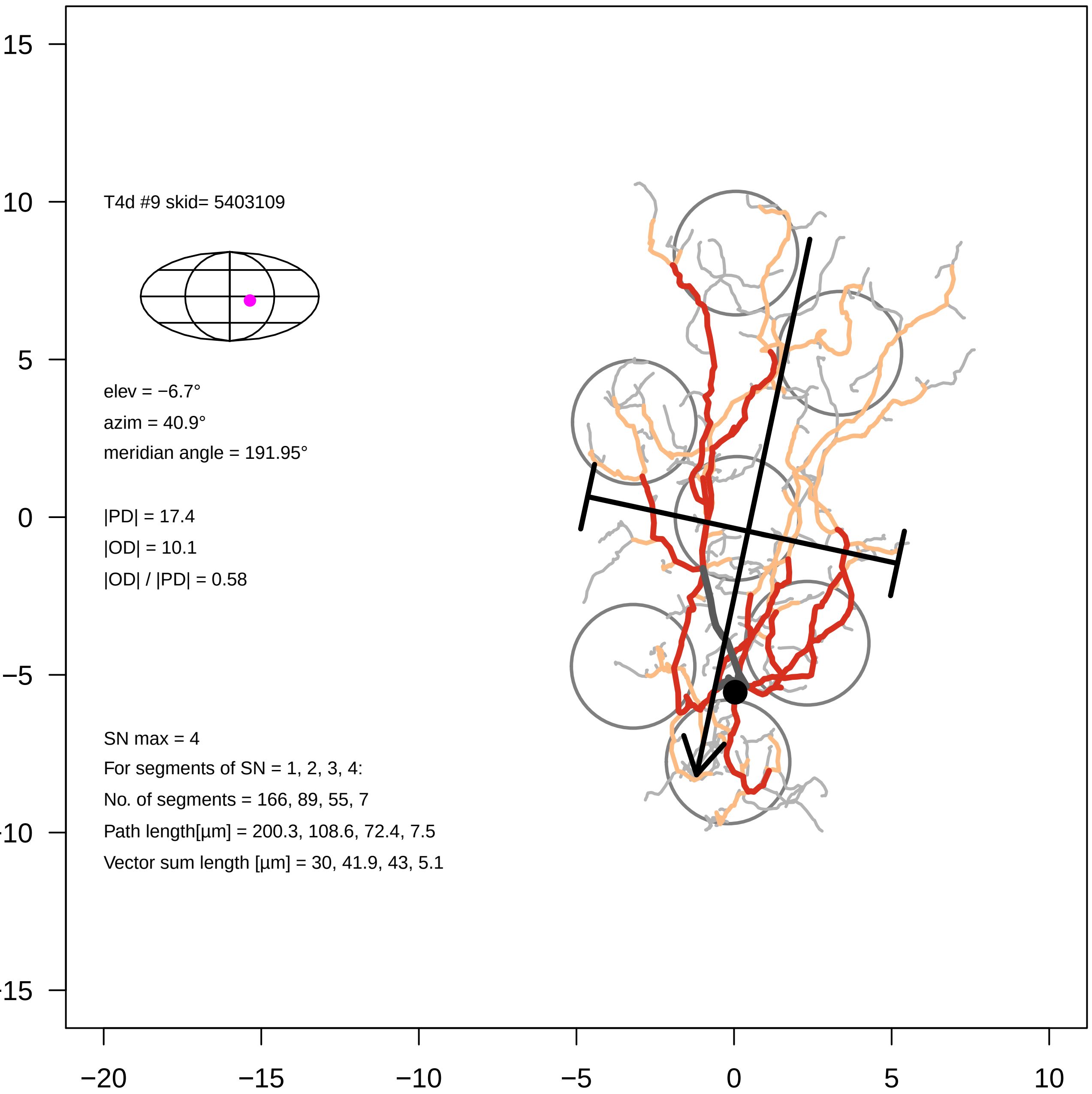

T4d #9 skid= 5403109
elev = −6.7°
azim = 40.9°
meridian angle = 191.95°
|PD| = 17.4
|OD| = 10.1
|OD| / |PD| = 0.58
SN max = 4
For segments of SN = 1, 2, 3, 4:
No. of segments = 166, 89, 55, 7
Path length[µm] = 200.3, 108.6, 72.4, 7.5
Vector sum length [µm] = 30, 41.9, 43, 5.1
15
10
5
0
−5
−10
−15
−20
−15
−10
−5
0
5
10

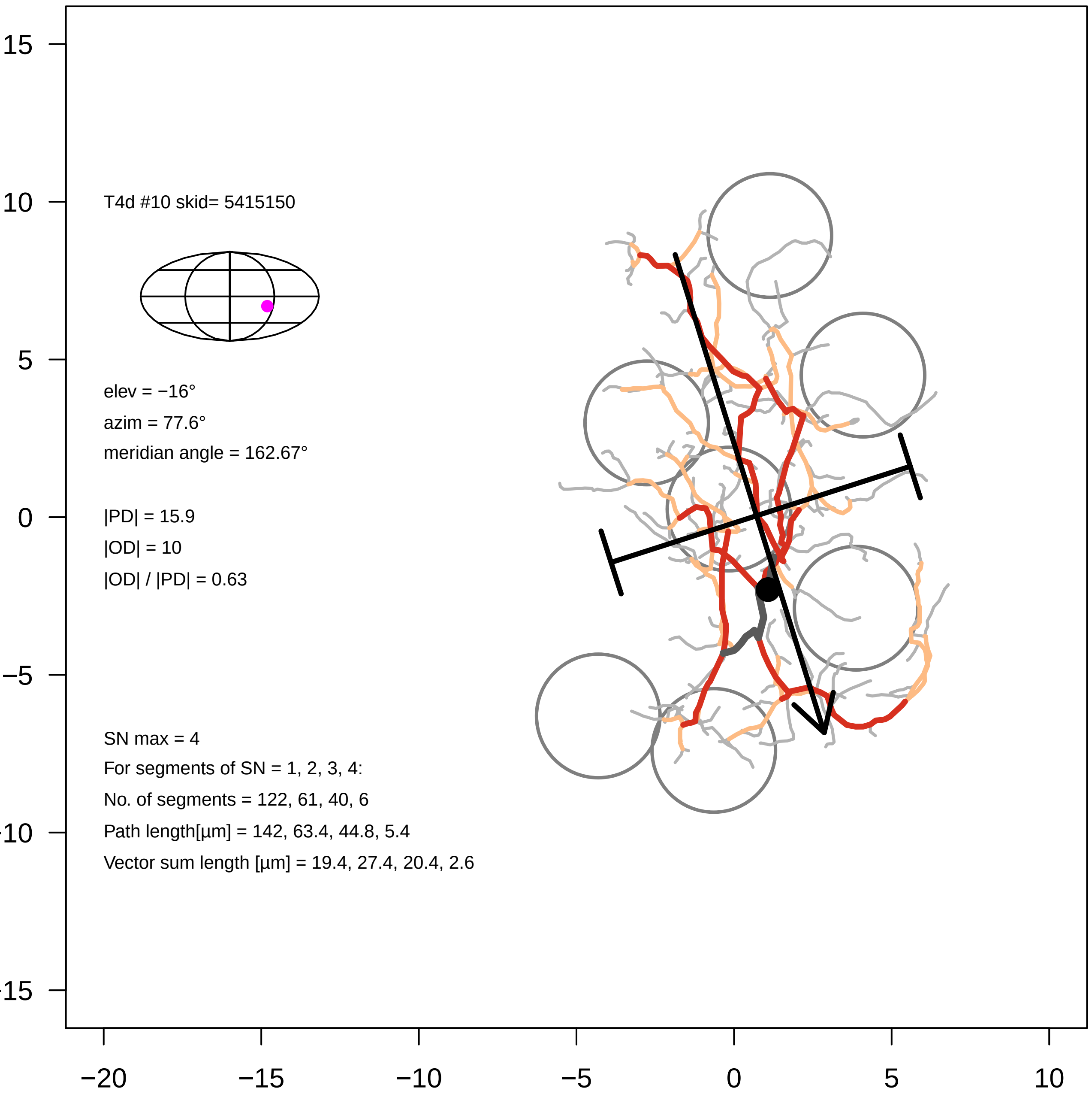

T4d #10 skid= 5415150
elev = −16°
azim = 77.6°
meridian angle = 162.67°
|PD| = 15.9
|OD| = 10
|OD| / |PD| = 0.63
SN max = 4
For segments of SN = 1, 2, 3, 4:
No. of segments = 122, 61, 40, 6
Path length[µm] = 142, 63.4, 44.8, 5.4
Vector sum length [µm] = 19.4, 27.4, 20.4, 2.6
15
10
5
0
−5
−10
−15
−20
−15
−10
−5
0
5
10

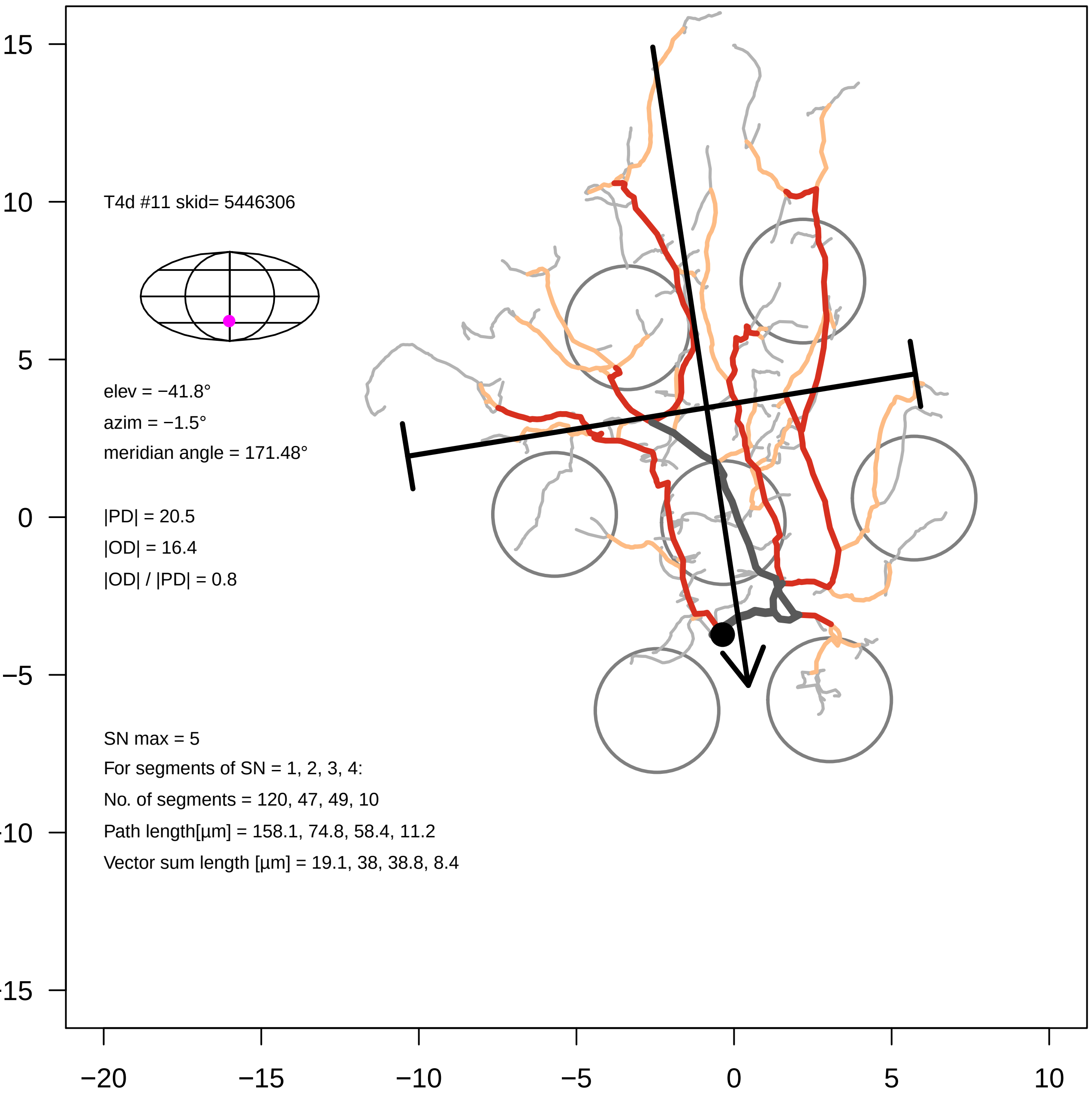

T4d #11 skid= 5446306
elev = −41.8°
azim = −1.5°
meridian angle = 171.48°
|PD| = 20.5
|OD| = 16.4
|OD| / |PD| = 0.8
SN max = 5
For segments of SN = 1, 2, 3, 4:
No. of segments = 120, 47, 49, 10
Path length[µm] = 158.1, 74.8, 58.4, 11.2
Vector sum length [µm] = 19.1, 38, 38.8, 8.4
15
10
5
0
−5
-10
-15
−20
−15
−10
−5
0
5
10

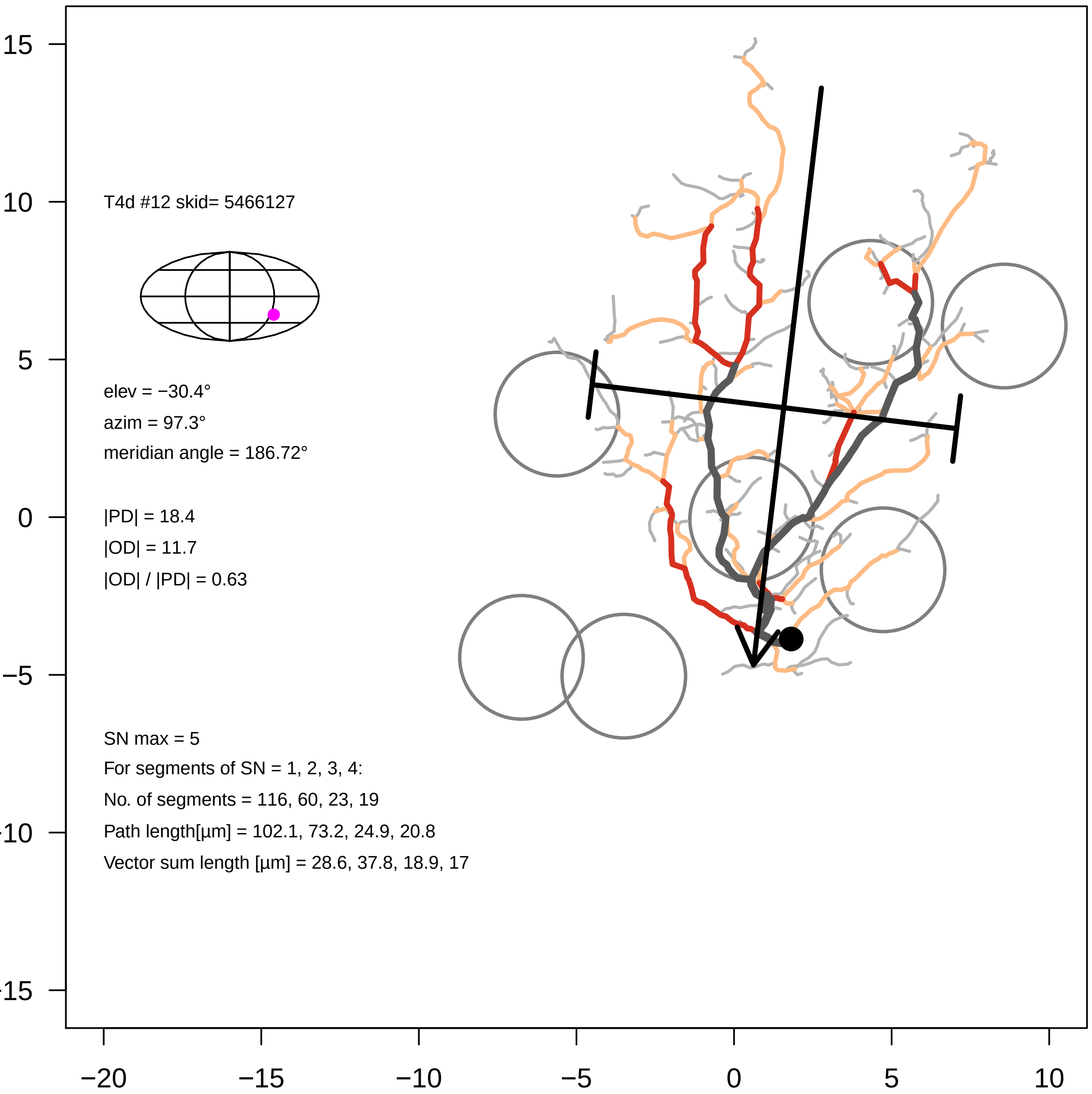
T4d #12 skid= 5466127
elev = −30.4°
azim = 97.3°
meridian angle = 186.72°
|PD| = 18.4
|OD| = 11.7
|OD| / |PD| = 0.63
SN max = 5
For segments of SN = 1, 2, 3, 4:
No. of segments = 116, 60, 23, 19
Path length[µm] = 102.1, 73.2, 24.9, 20.8
Vector sum length [µm] = 28.6, 37.8, 18.9, 17
15
10
5
0
−5
-10
-15
−20
−15
−10
−5
0
5
10

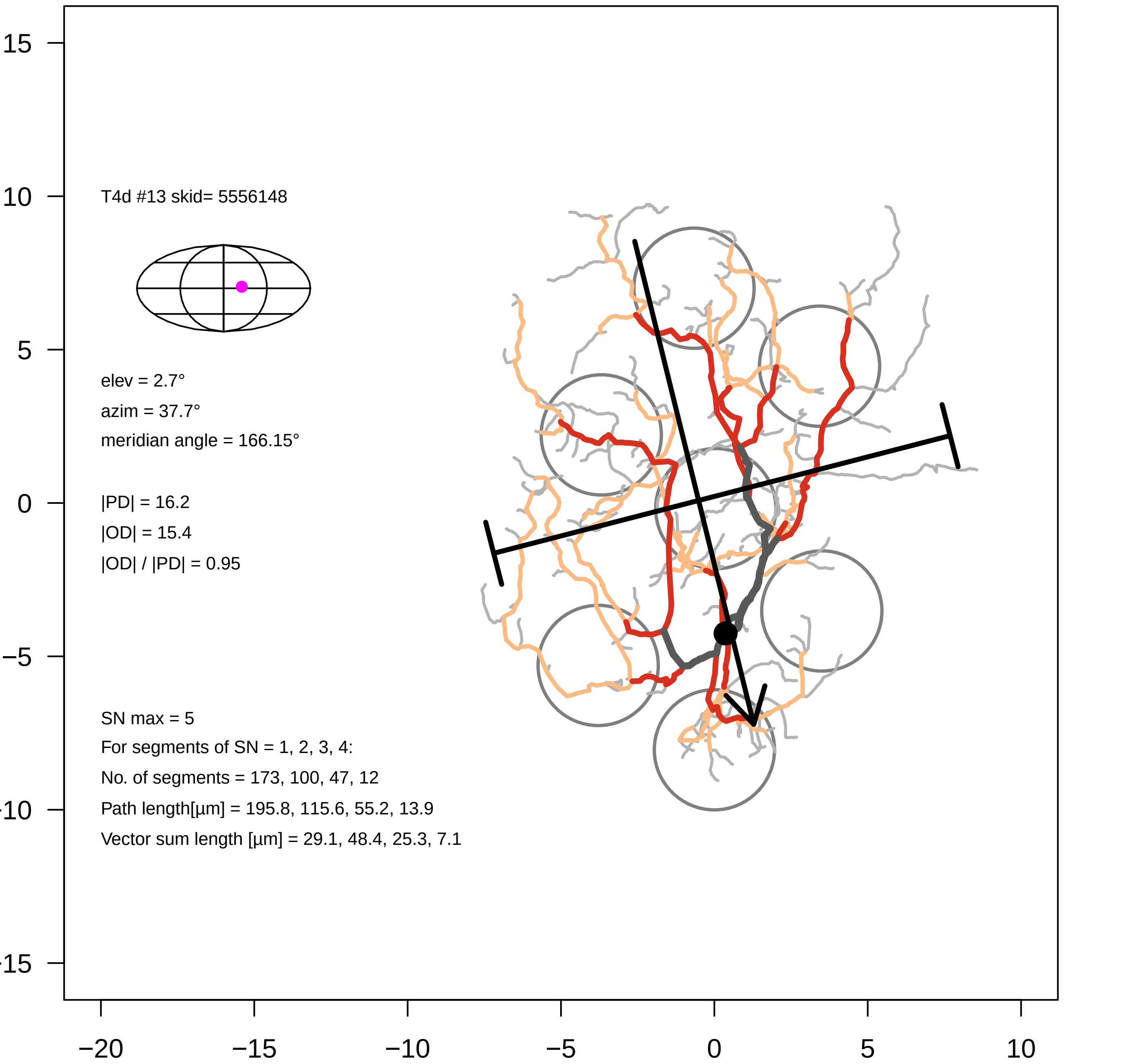

T4d #13 skid= 5556148
elev = 2.7°
azim = 37.7°
meridian angle = 166.15°
|PD| = 16.2
|OD| = 15.4
|OD| / |PD| = 0.95
SN max = 5
For segments of SN = 1, 2, 3, 4:
No. of segments = 173, 100, 47, 12
Path length[µm] = 195.8, 115.6, 55.2, 13.9
Vector sum length [µm] = 29.1, 48.4, 25.3, 7.1
15
10
5
0
−5
−10
−15
−20
−15
−10
−5
0
5
10

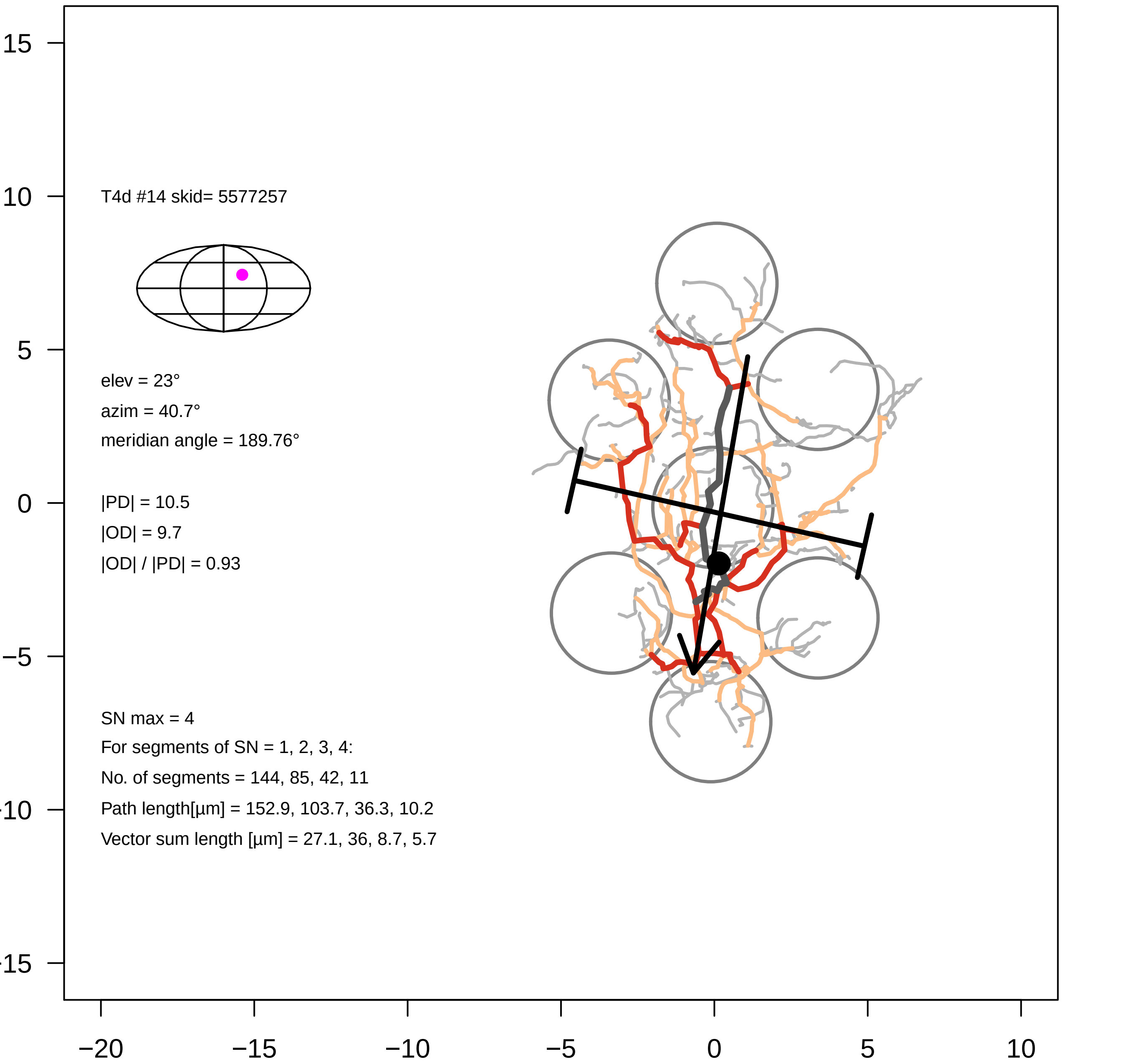

T4d #14 skid= 5577257
elev = 23°
azim = 40.7°
meridian angle = 189.76°
|PD| = 10.5
|OD| = 9.7
|OD| / |PD| = 0.93
SN max = 4
For segments of SN = 1, 2, 3, 4:
No. of segments = 144, 85, 42, 11
Path length[µm] = 152.9, 103.7, 36.3, 10.2
Vector sum length [µm] = 27.1, 36, 8.7, 5.7
15
10
5
0
−5
−10
−15
−20
−15
−10
−5
0
5
10

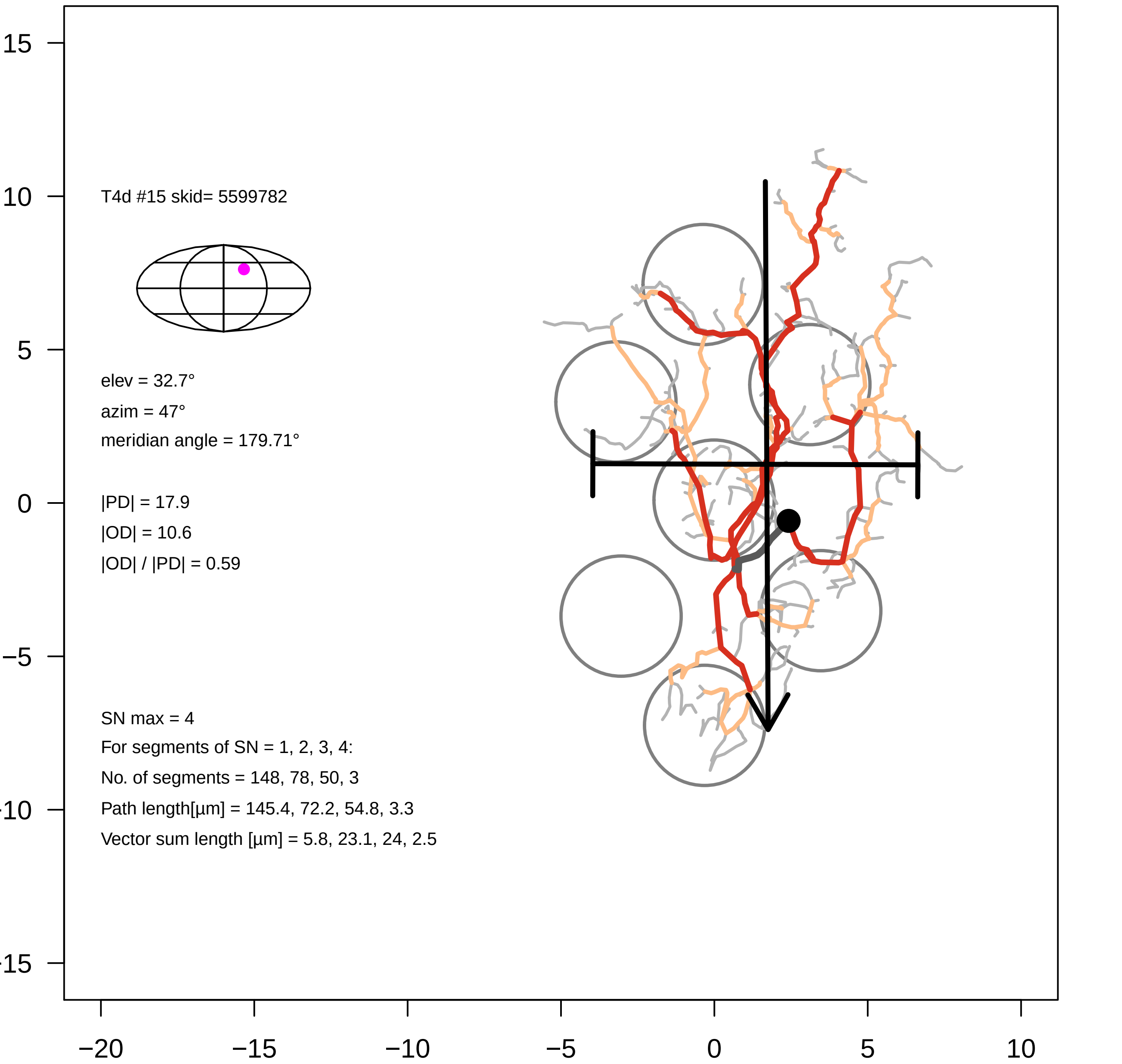
T4d #15 skid= 5599782
elev = 32.7°
azim = 47°
meridian angle = 179.71°
|PD| = 17.9
|OD| = 10.6
|OD| / |PD| = 0.59
SN max = 4
For segments of SN = 1, 2, 3, 4:
No. of segments = 148, 78, 50, 3
Path length[µm] = 145.4, 72.2, 54.8, 3.3
Vector sum length [µm] = 5.8, 23.1, 24, 2.5
15
10
5
0
−5
−10
−15
−20
−15
−10
−5
0
5
10

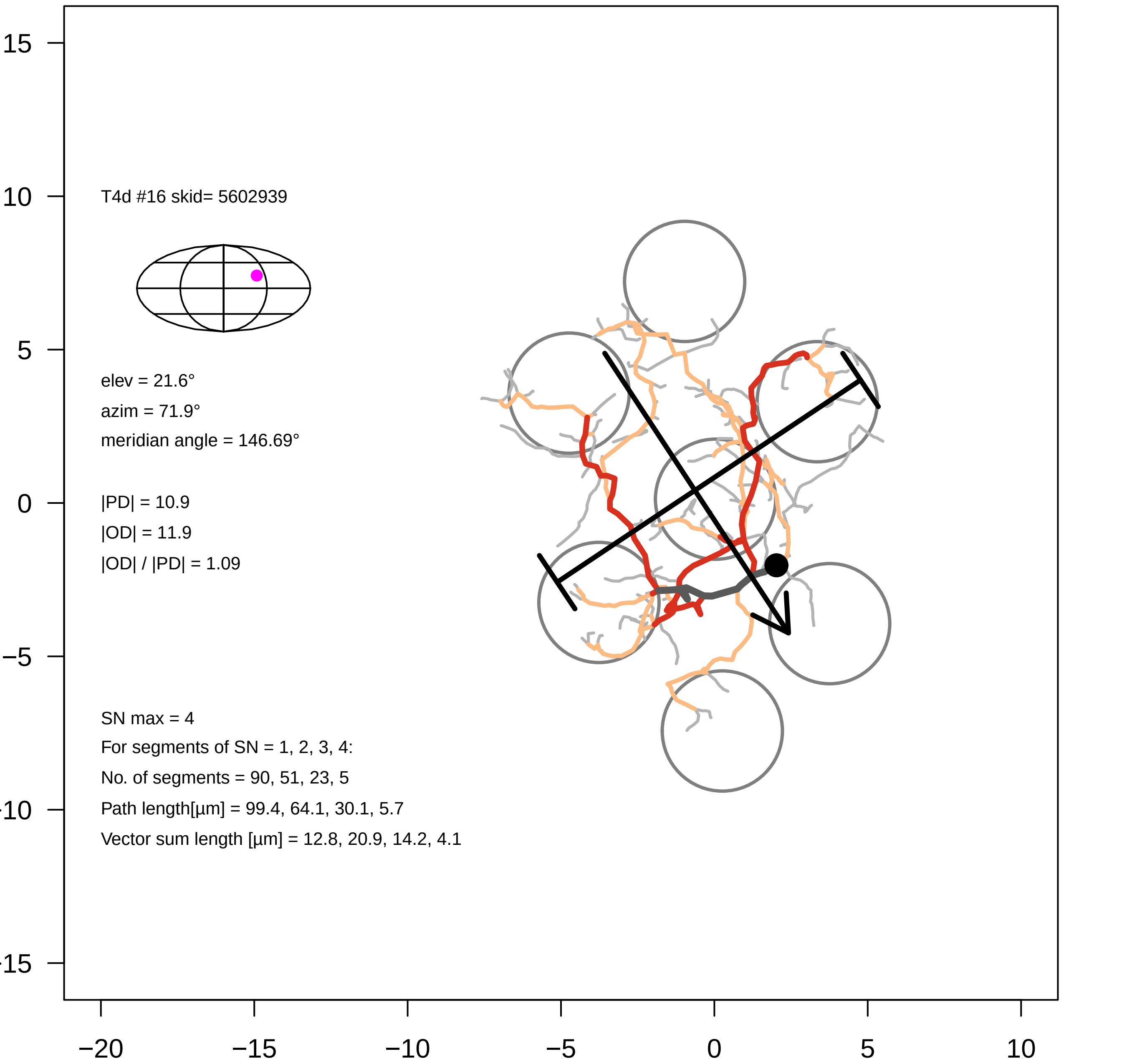

T4d #16 skid= 5602939
elev = 21.6°
azim = 71.9°
meridian angle = 146.69°
|PD| = 10.9
|OD| = 11.9
|OD| / |PD| = 1.09
SN max = 4
For segments of SN = 1, 2, 3, 4:
No. of segments = 90, 51, 23, 5
Path length[µm] = 99.4, 64.1, 30.1, 5.7
Vector sum length [µm] = 12.8, 20.9, 14.2, 4.1
15
10
5
0
−5
−10
−15
−20
−15
−10
−5
0
5
10

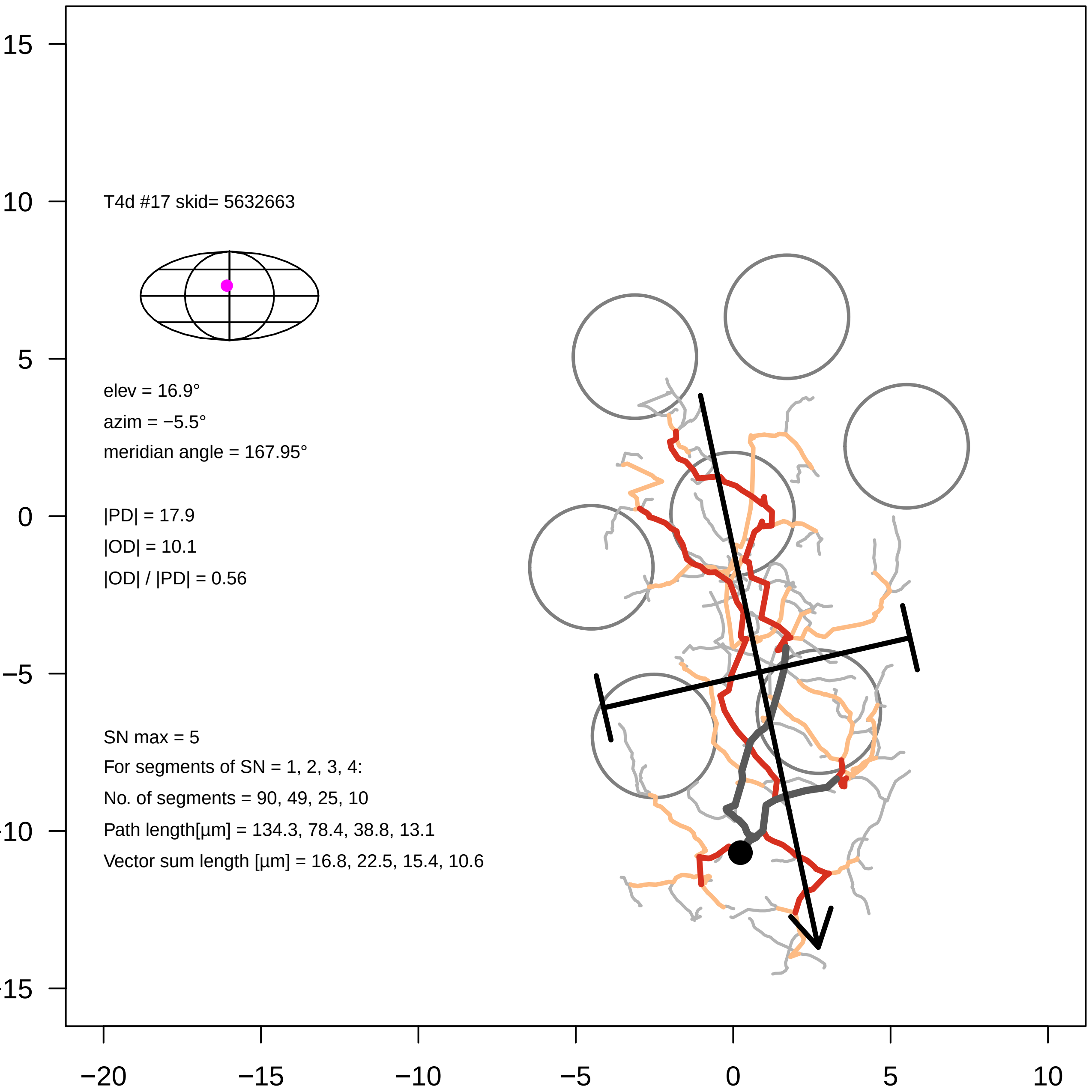
T4d #17 skid= 5632663
elev = 16.9°
azim = −5.5°
meridian angle = 167.95°
|PD| = 17.9
|OD| = 10.1
|OD| / |PD| = 0.56
SN max = 5
For segments of SN = 1, 2, 3, 4:
No. of segments = 90, 49, 25, 10
Path length[µm] = 134.3, 78.4, 38.8, 13.1
Vector sum length [µm] = 16.8, 22.5, 15.4, 10.6
15
10
5
0
−5
−10
−15
−20
−15
−10
−5
0
5
10

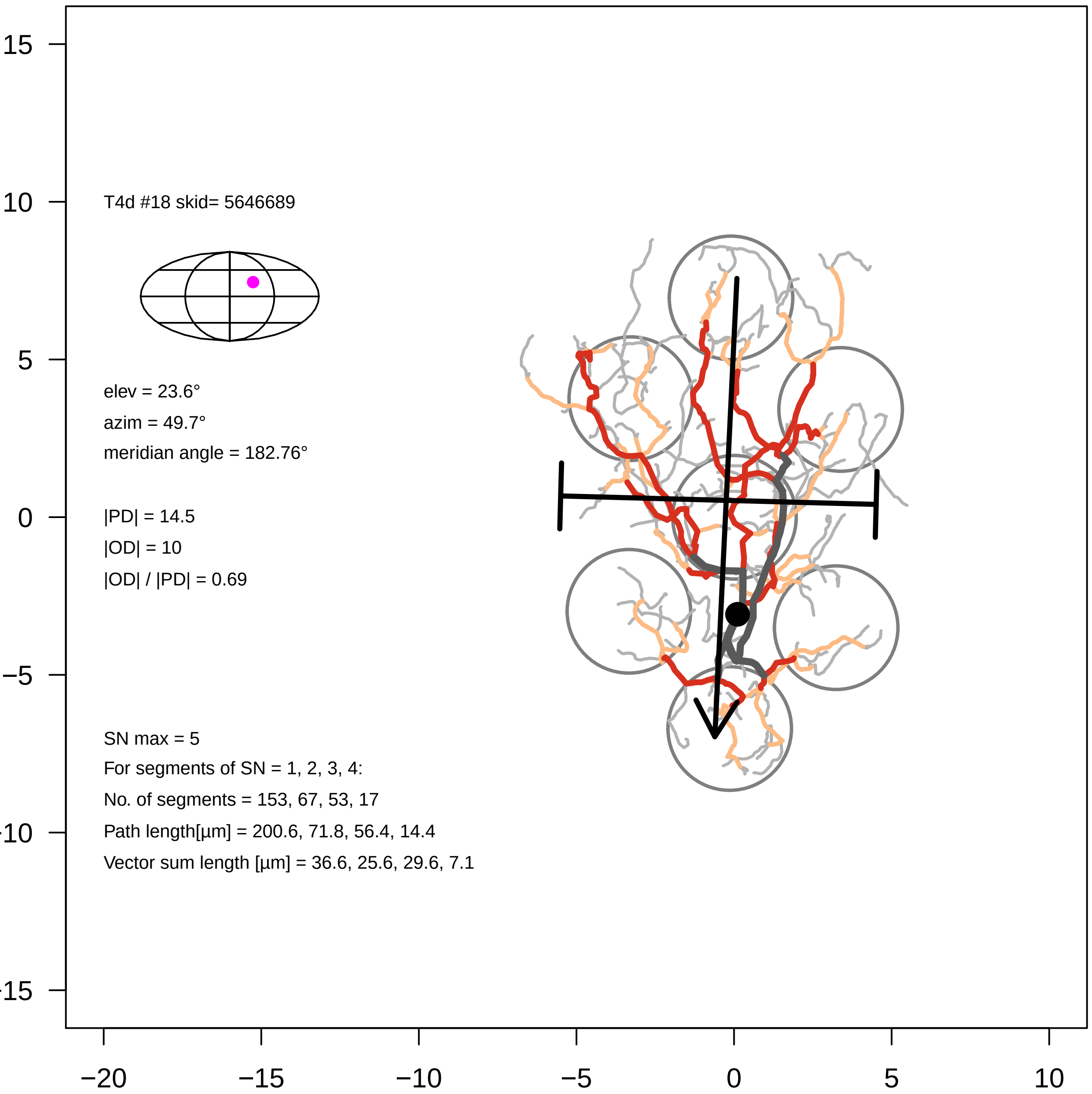

T4d #18 skid= 5646689
elev = 23.6°
azim = 49.7°
meridian angle = 182.76°
|PD| = 14.5
|OD| = 10
|OD| / |PD| = 0.69
SN max = 5
For segments of SN = 1, 2, 3, 4:
No. of segments = 153, 67, 53, 17
Path length[µm] = 200.6, 71.8, 56.4, 14.4
Vector sum length [µm] = 36.6, 25.6, 29.6, 7.1
15
10
5
0
-5
-10
-15
−20
−15
−10
−5
0
5
10

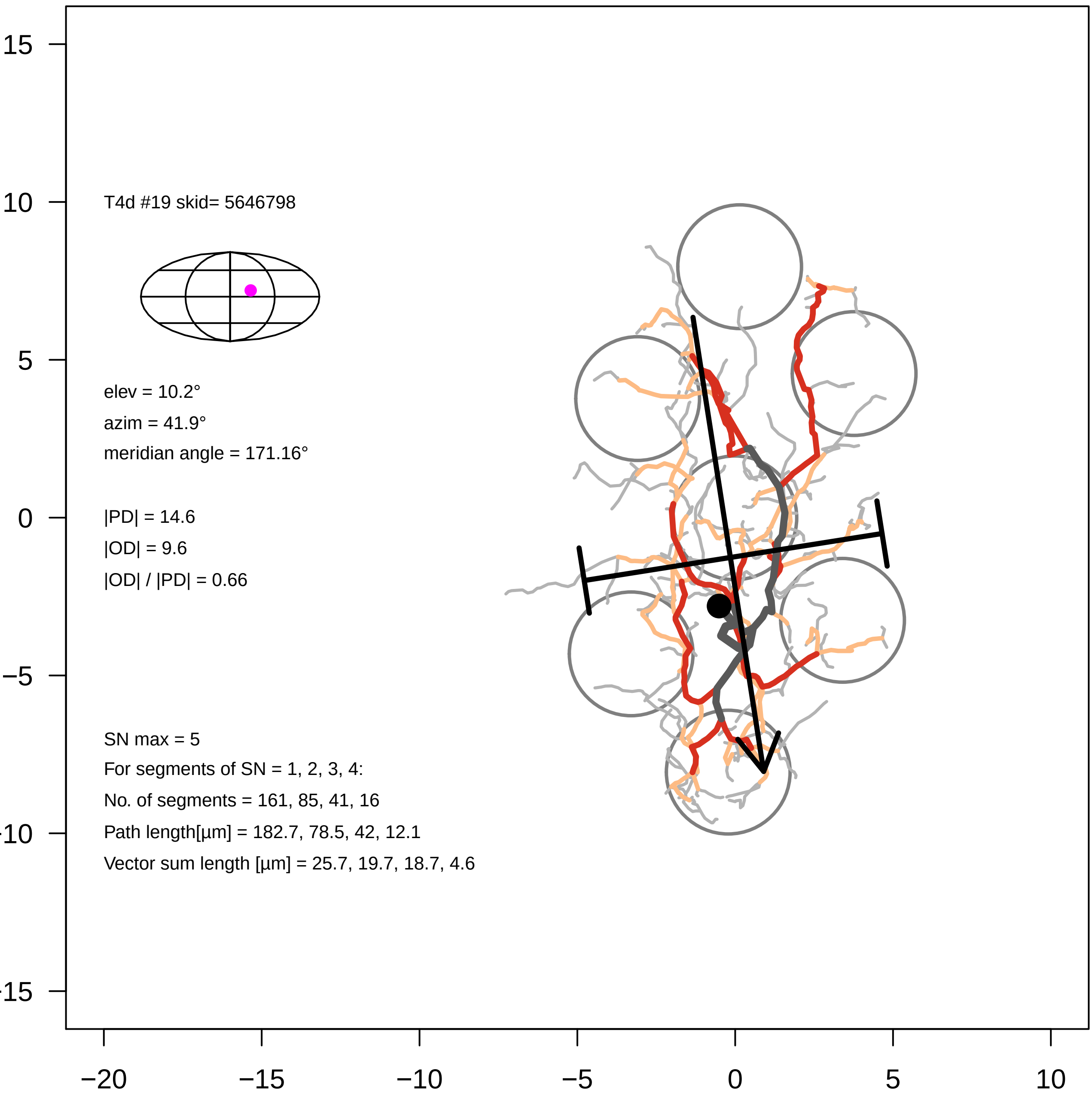

15
10
5
0
−5
−10
−15
−20
−15
−10
−5
0
5
10
T4d #19 skid= 5646798
elev = 10.2°
azim = 41.9°
meridian angle = 171.16°
|PD| = 14.6
|OD| = 9.6
|OD| / |PD| = 0.66
SN max = 5
For segments of SN = 1, 2, 3, 4:
No. of segments = 161, 85, 41, 16
Path length[µm] = 182.7, 78.5, 42, 12.1
Vector sum length [µm] = 25.7, 19.7, 18.7, 4.6

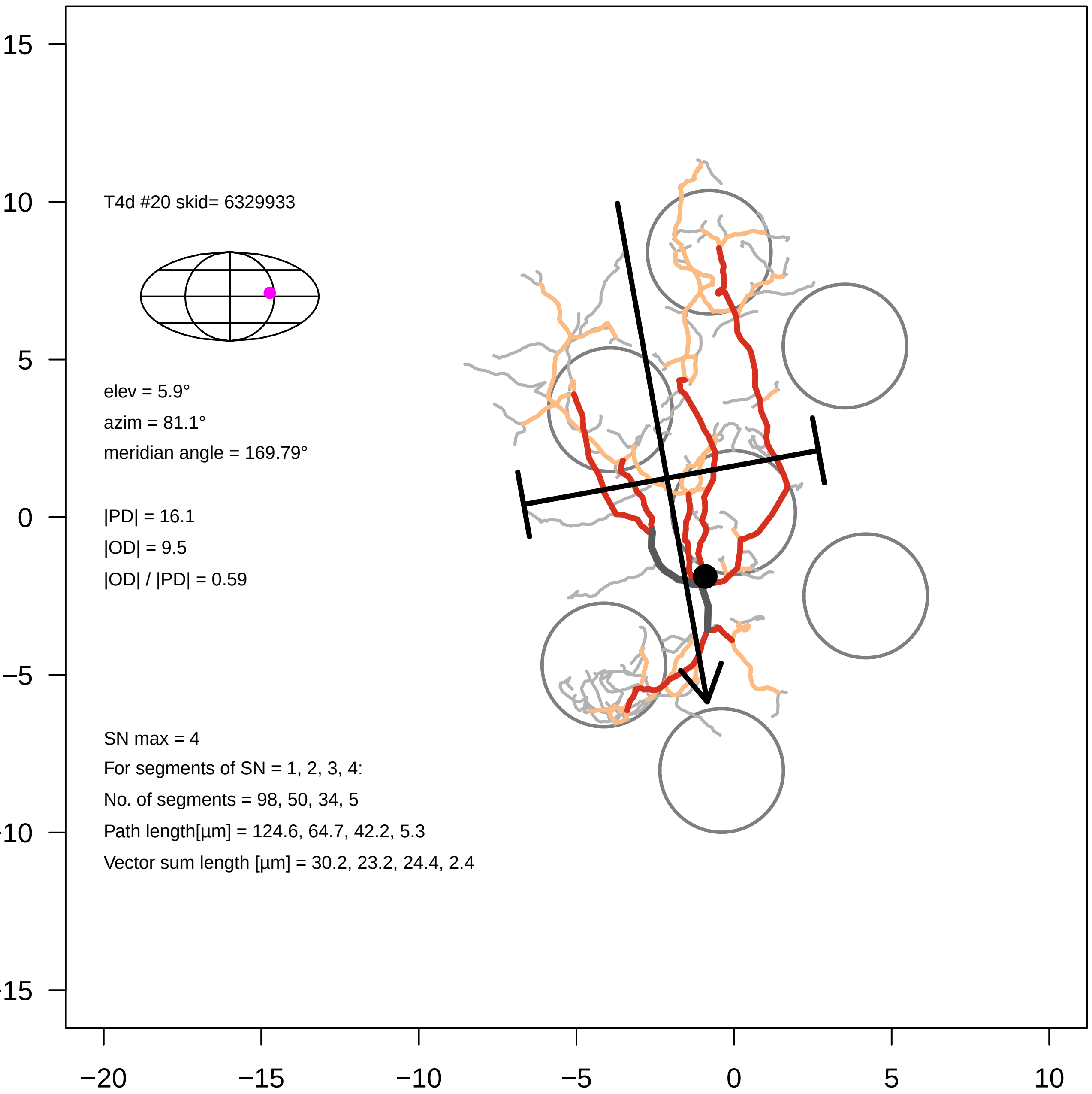

T4d #20 skid= 6329933
elev = 5.9°
azim = 81.1°
meridian angle = 169.79°
|PD| = 16.1
|OD| = 9.5
|OD| / |PD| = 0.59
SN max = 4
For segments of SN = 1, 2, 3, 4:
No. of segments = 98, 50, 34, 5
Path length[µm] = 124.6, 64.7, 42.2, 5.3
Vector sum length [µm] = 30.2, 23.2, 24.4, 2.4
15
10
5
0
−5
−10
−15
−20
−15
−10
−5
0
5
10

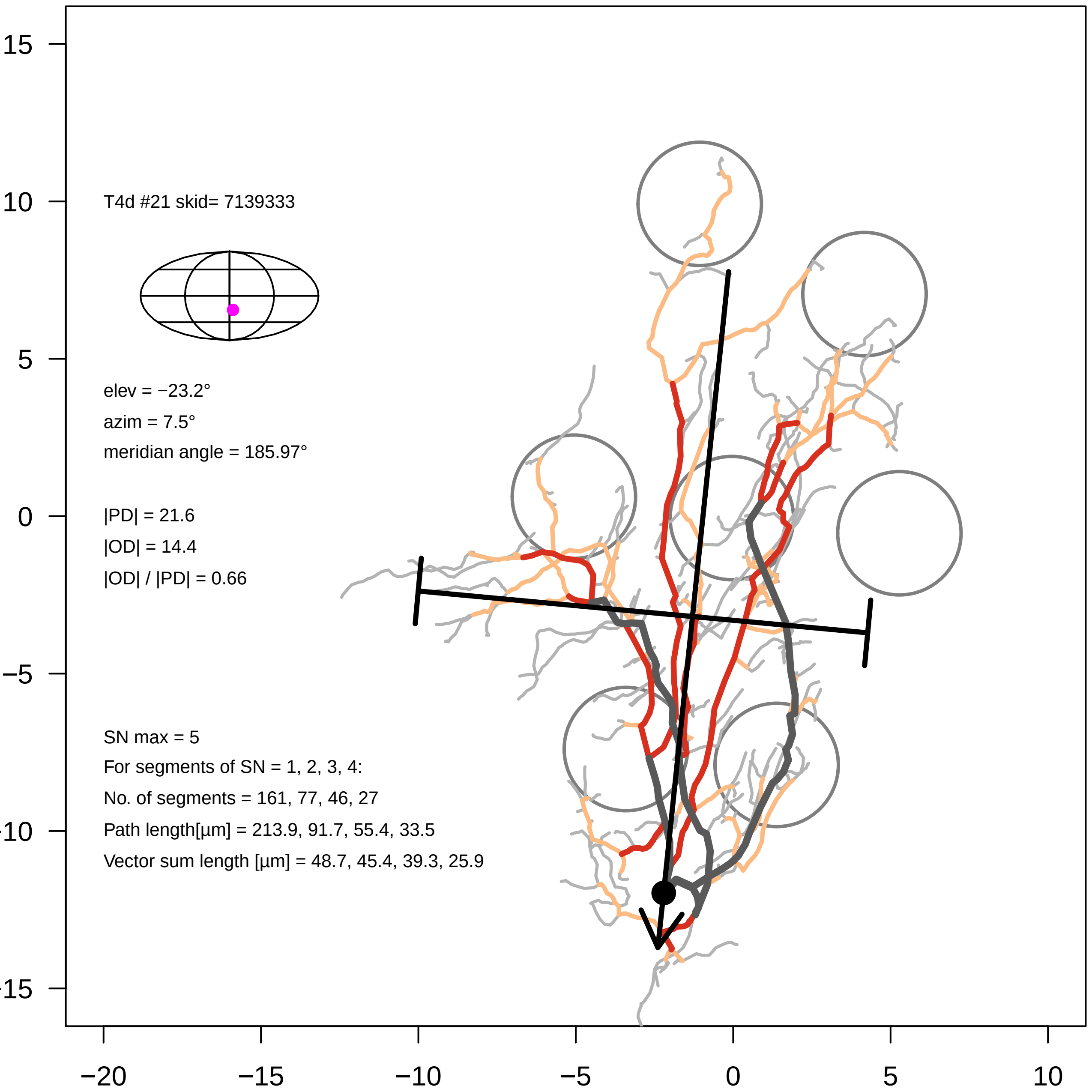

T4d #21 skid= 7139333
elev = −23.2°
azim = 7.5°
meridian angle = 185.97°
|PD| = 21.6
|OD| = 14.4
|OD| / |PD| = 0.66
SN max = 5
For segments of SN = 1, 2, 3, 4:
No. of segments = 161, 77, 46, 27
Path length[µm] = 213.9, 91.7, 55.4, 33.5
Vector sum length [µm] = 48.7, 45.4, 39.3, 25.9
15
10
5
0
−5
−10
−15
−20
−15
−10
−5
0
5
10

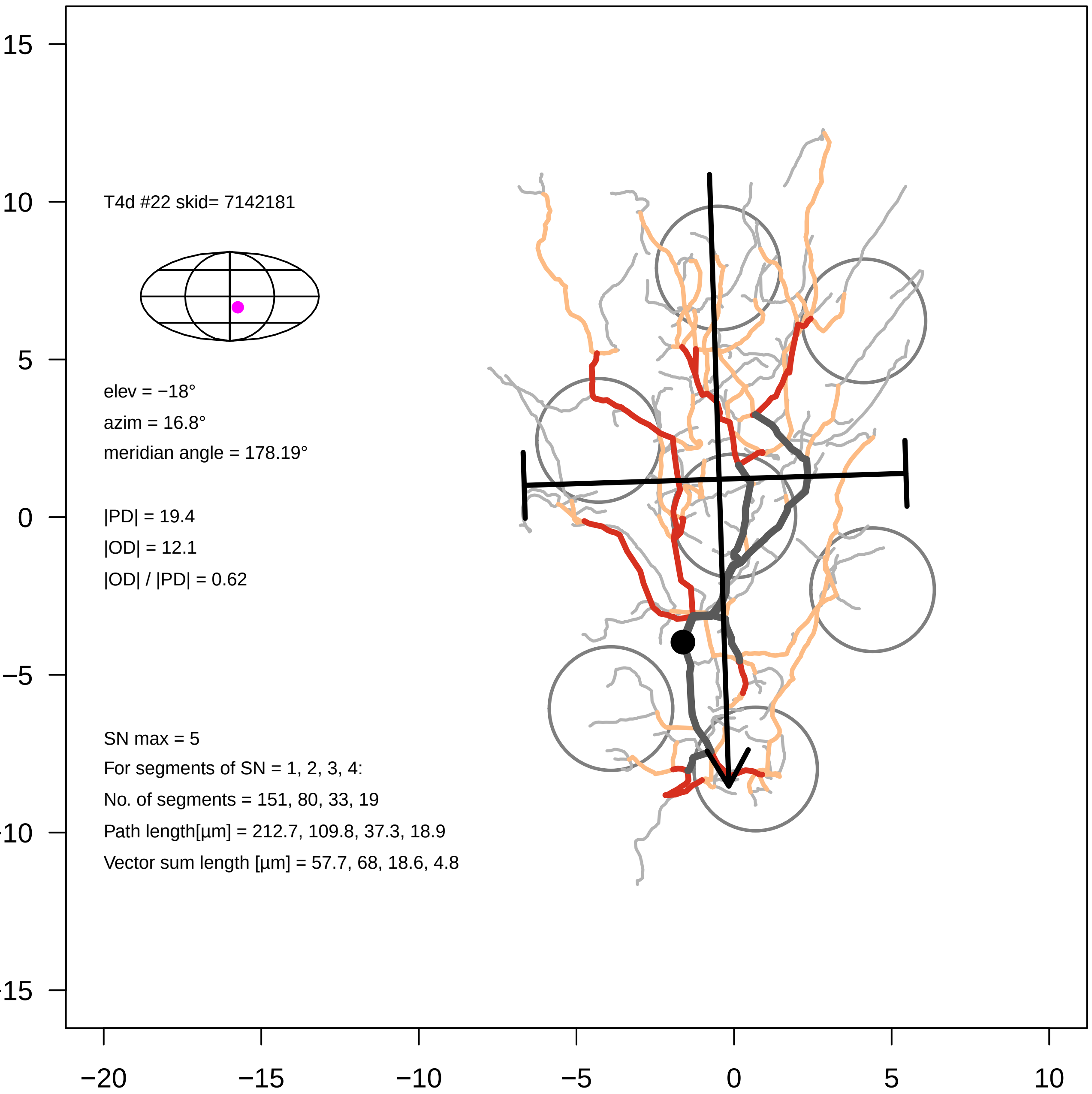

T4d #22 skid= 7142181
elev = −18°
azim = 16.8°
meridian angle = 178.19°
|PD| = 19.4
|OD| = 12.1
|OD| / |PD| = 0.62
SN max = 5
For segments of SN = 1, 2, 3, 4:
No. of segments = 151, 80, 33, 19
Path length[µm] = 212.7, 109.8, 37.3, 18.9
Vector sum length [µm] = 57.7, 68, 18.6, 4.8
15
10
5
0
−5
-10
-15
−20
−15
−10
−5
0
5
10

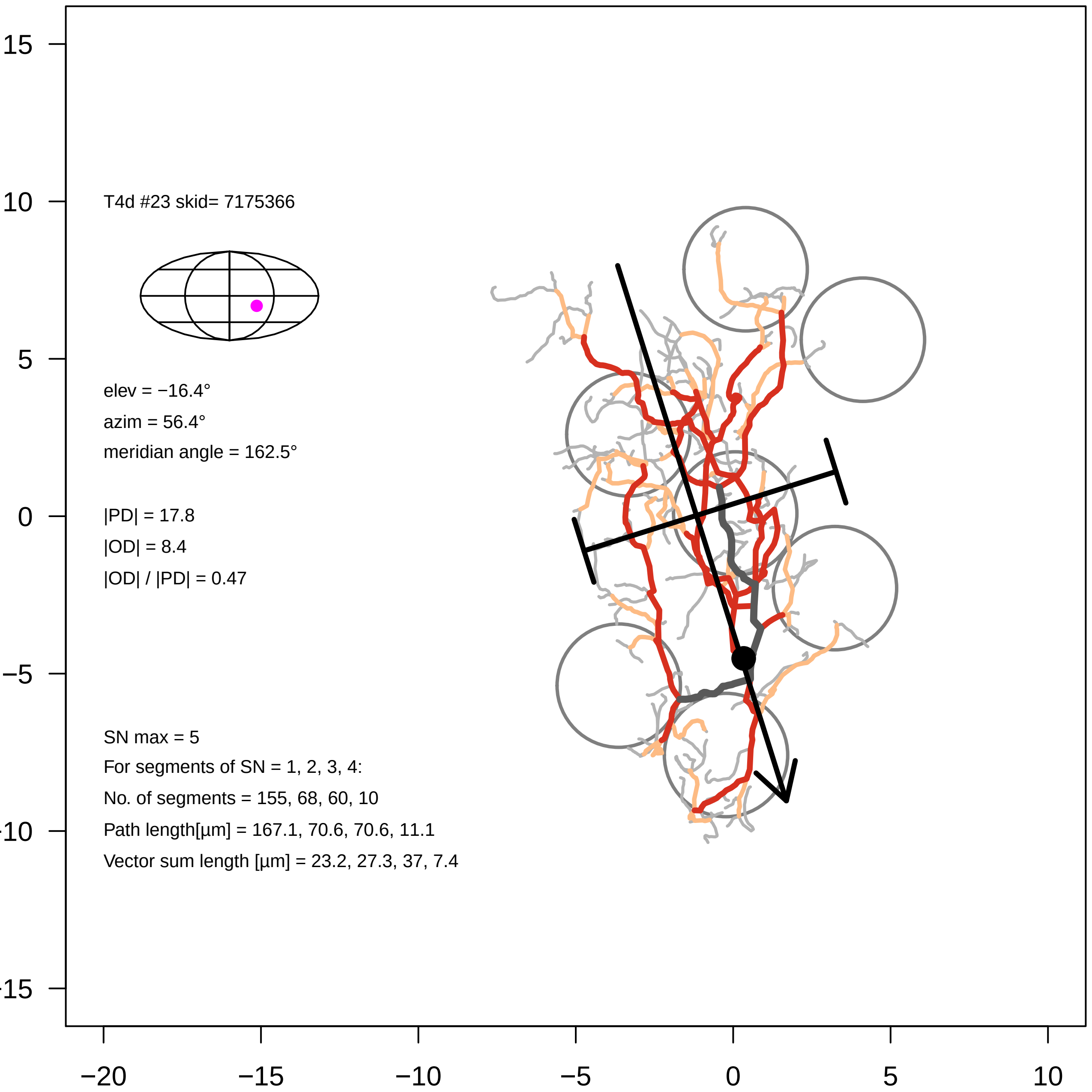

T4d #23 skid= 7175366
elev = −16.4°
azim = 56.4°
meridian angle = 162.5°
|PD| = 17.8
|OD| = 8.4
|OD| / |PD| = 0.47
SN max = 5
For segments of SN = 1, 2, 3, 4:
No. of segments = 155, 68, 60, 10
Path length[µm] = 167.1, 70.6, 70.6, 11.1
Vector sum length [µm] = 23.2, 27.3, 37, 7.4
15
10
5
0
−5
-10
-15
−20
−15
−10
−5
0
5
10

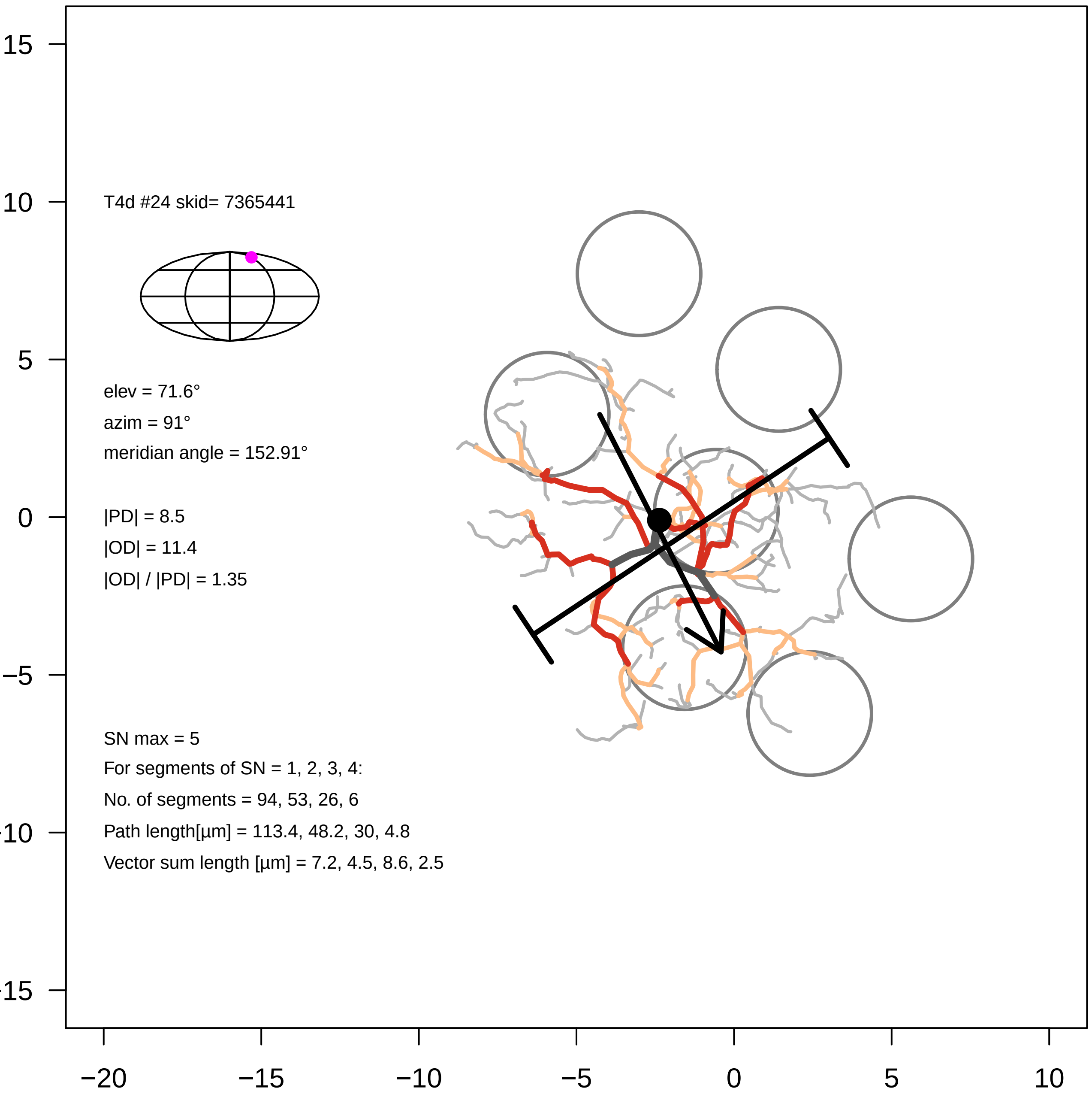

T4d #24 skid= 7365441
elev = 71.6°
azim = 91°
meridian angle = 152.91°
|PD| = 8.5
|OD| = 11.4
|OD| / |PD| = 1.35
SN max = 5
For segments of SN = 1, 2, 3, 4:
No. of segments = 94, 53, 26, 6
Path length[µm] = 113.4, 48.2, 30, 4.8
Vector sum length [µm] = 7.2, 4.5, 8.6, 2.5
15
10
5
0
−5
−10
−15
−20
−15
−10
−5
0
5
10

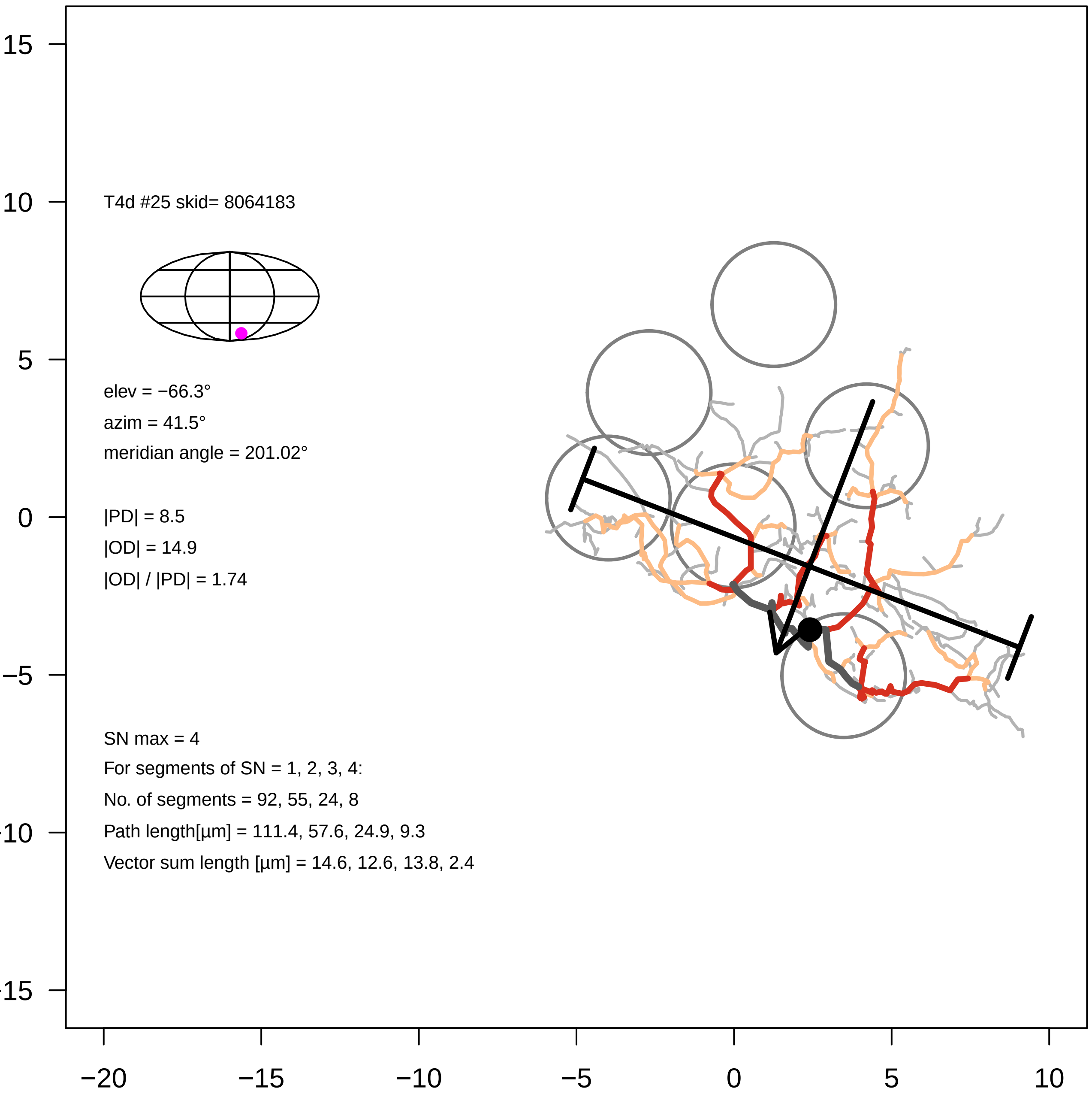

T4d #25 skid= 8064183
elev = −66.3°
azim = 41.5°
meridian angle = 201.02°
|PD| = 8.5
|OD| = 14.9
|OD| / |PD| = 1.74
SN max = 4
For segments of SN = 1, 2, 3, 4:
No. of segments = 92, 55, 24, 8
Path length[µm] = 111.4, 57.6, 24.9, 9.3
Vector sum length [µm] = 14.6, 12.6, 13.8, 2.4
15
10
5
0
−5
-10
-15
−20
−15
−10
−5
0
5
10

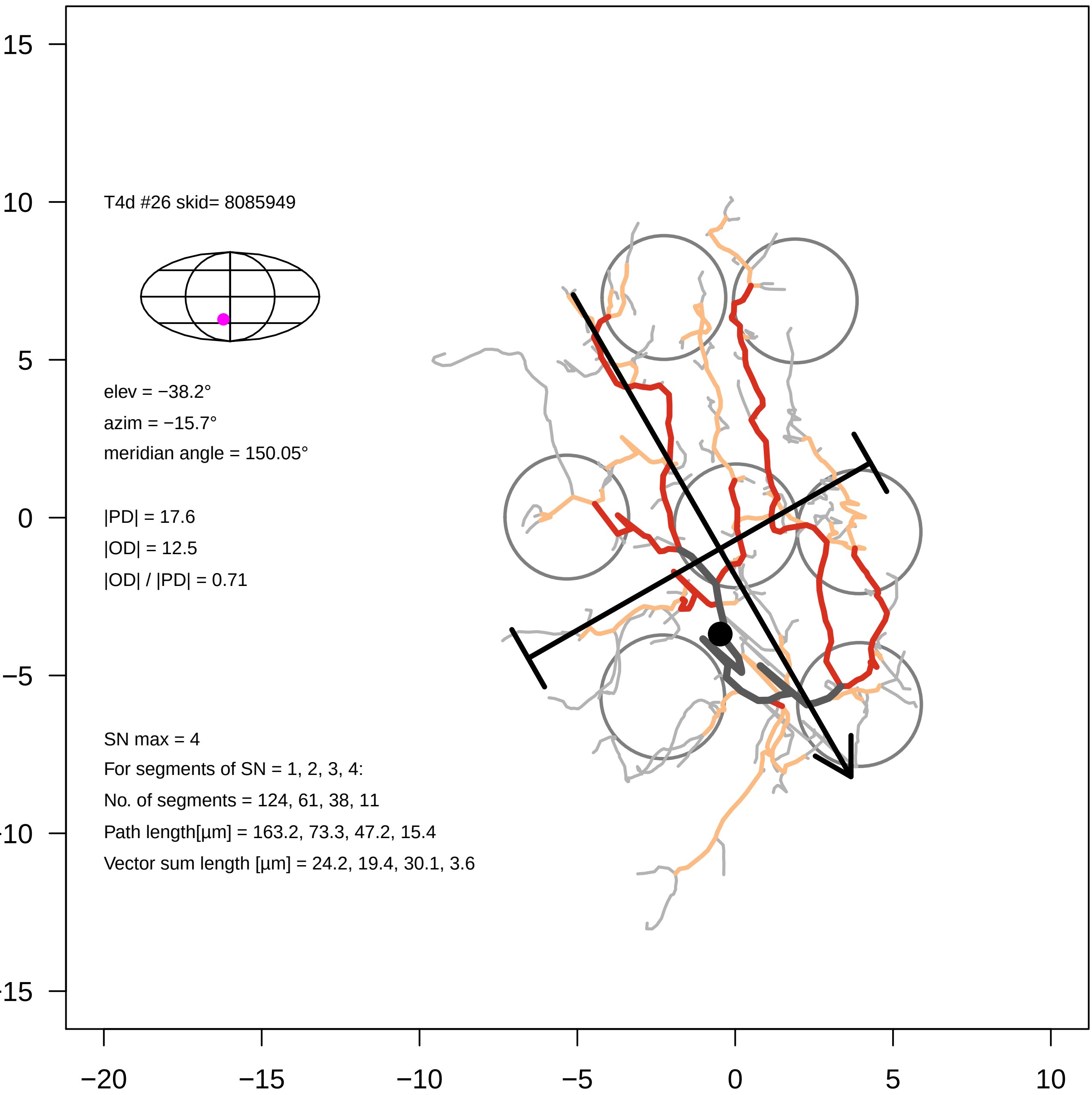
T4d #26 skid= 8085949
elev = −38.2°
azim = −15.7°
meridian angle = 150.05°
|PD| = 17.6
|OD| = 12.5
|OD| / |PD| = 0.71
SN max = 4
For segments of SN = 1, 2, 3, 4:
No. of segments = 124, 61, 38, 11
Path length[µm] = 163.2, 73.3, 47.2, 15.4
Vector sum length [µm] = 24.2, 19.4, 30.1, 3.6
15
10
5
0
−5
−10
−15
−20
−15
−10
−5
0
5
10

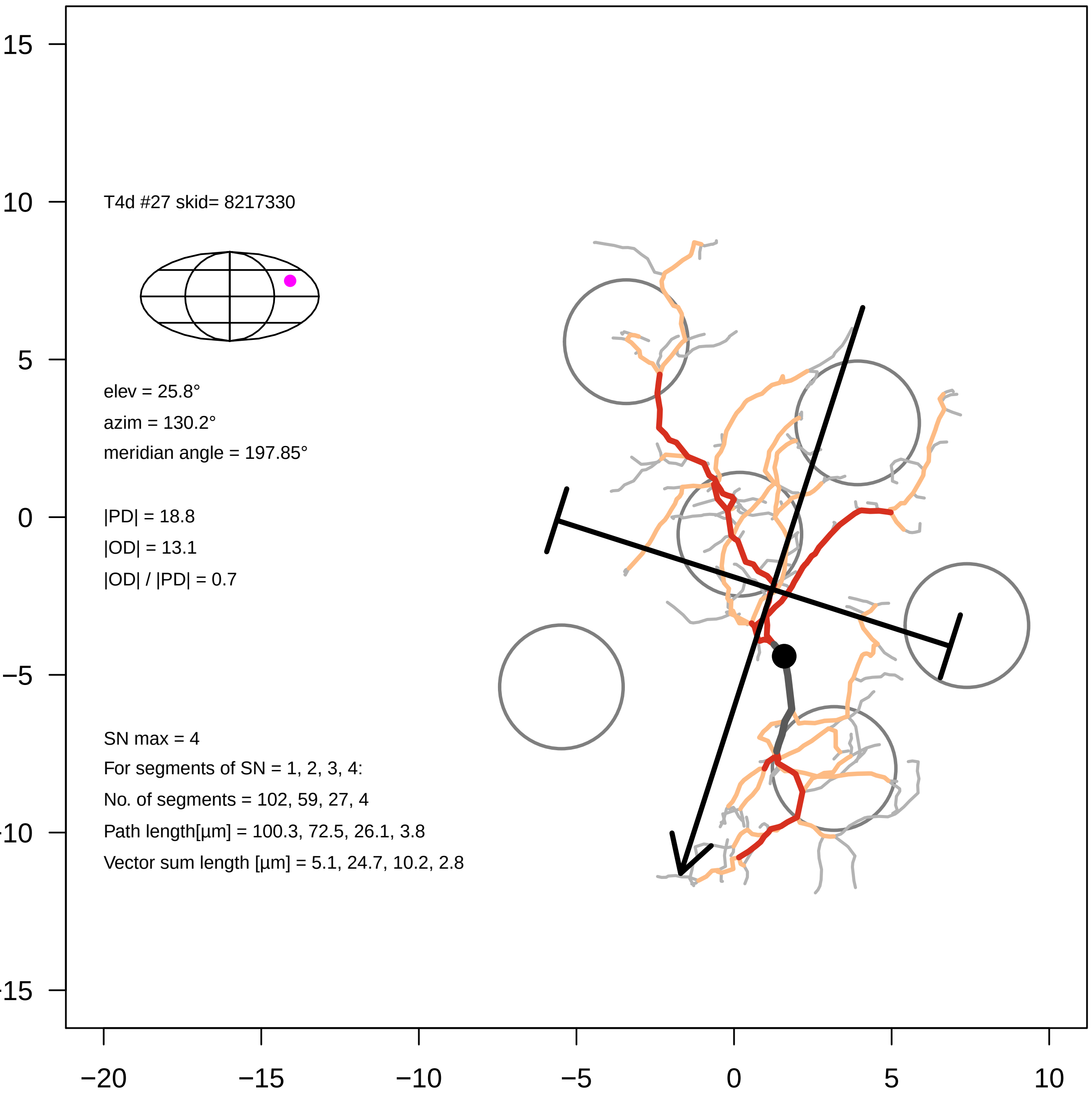

T4d #27 skid= 8217330
elev = 25.8°
azim = 130.2°
meridian angle = 197.85°
|PD| = 18.8
|OD| = 13.1
|OD| / |PD| = 0.7
SN max = 4
For segments of SN = 1, 2, 3, 4:
No. of segments = 102, 59, 27, 4
Path length[µm] = 100.3, 72.5, 26.1, 3.8
Vector sum length [µm] = 5.1, 24.7, 10.2, 2.8
15
10
5
0
−5
−10
−15
−20
−15
−10
−5
0
5
10

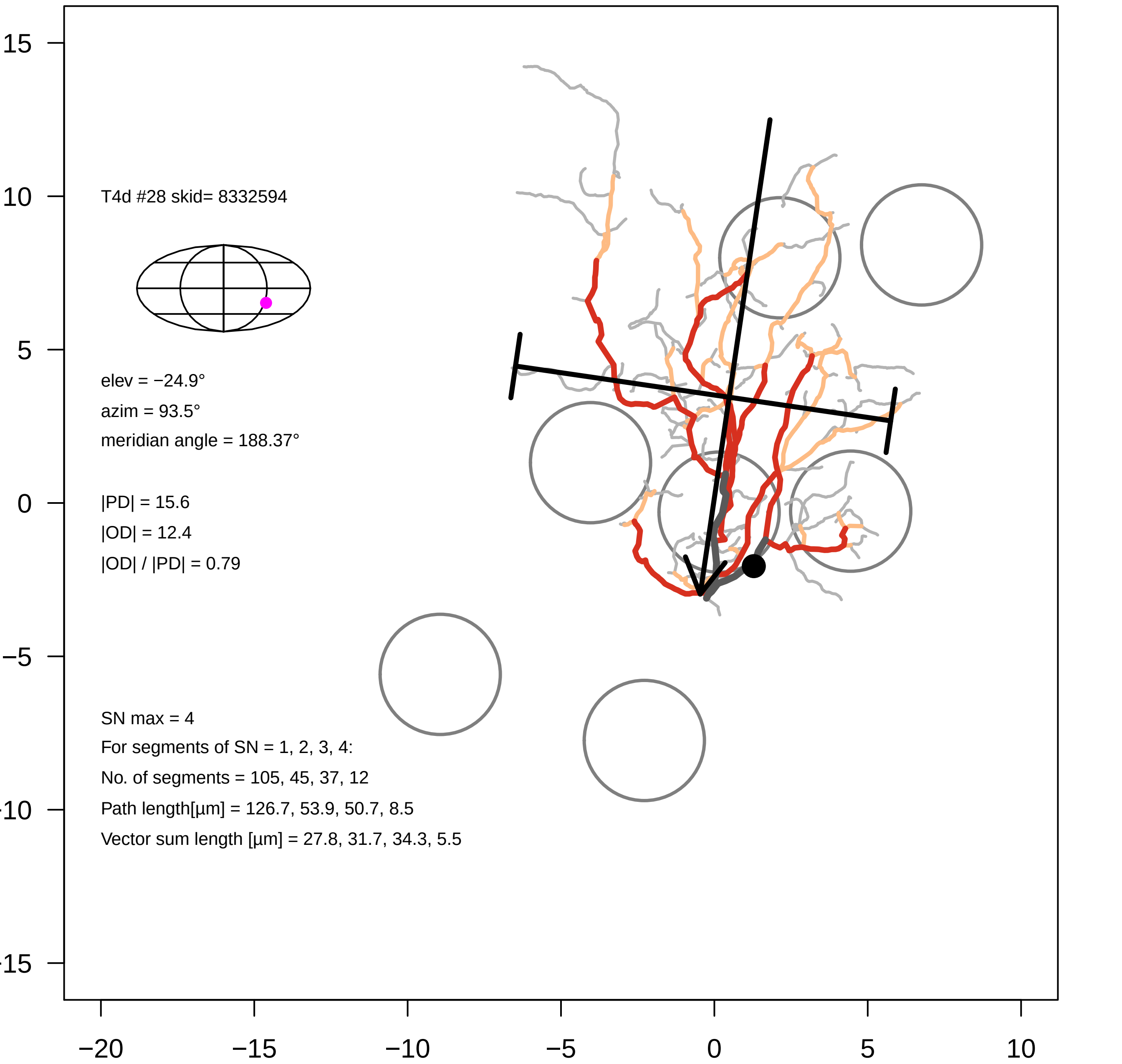
T4d #28 skid= 8332594
elev = −24.9°
azim = 93.5°
meridian angle = 188.37°
|PD| = 15.6
|OD| = 12.4
|OD| / |PD| = 0.79
SN max = 4
For segments of SN = 1, 2, 3, 4:
No. of segments = 105, 45, 37, 12
Path length[µm] = 126.7, 53.9, 50.7, 8.5
Vector sum length [µm] = 27.8, 31.7, 34.3, 5.5
15
10
5
0
−5
−20
−15
−10
−5
0
5
10

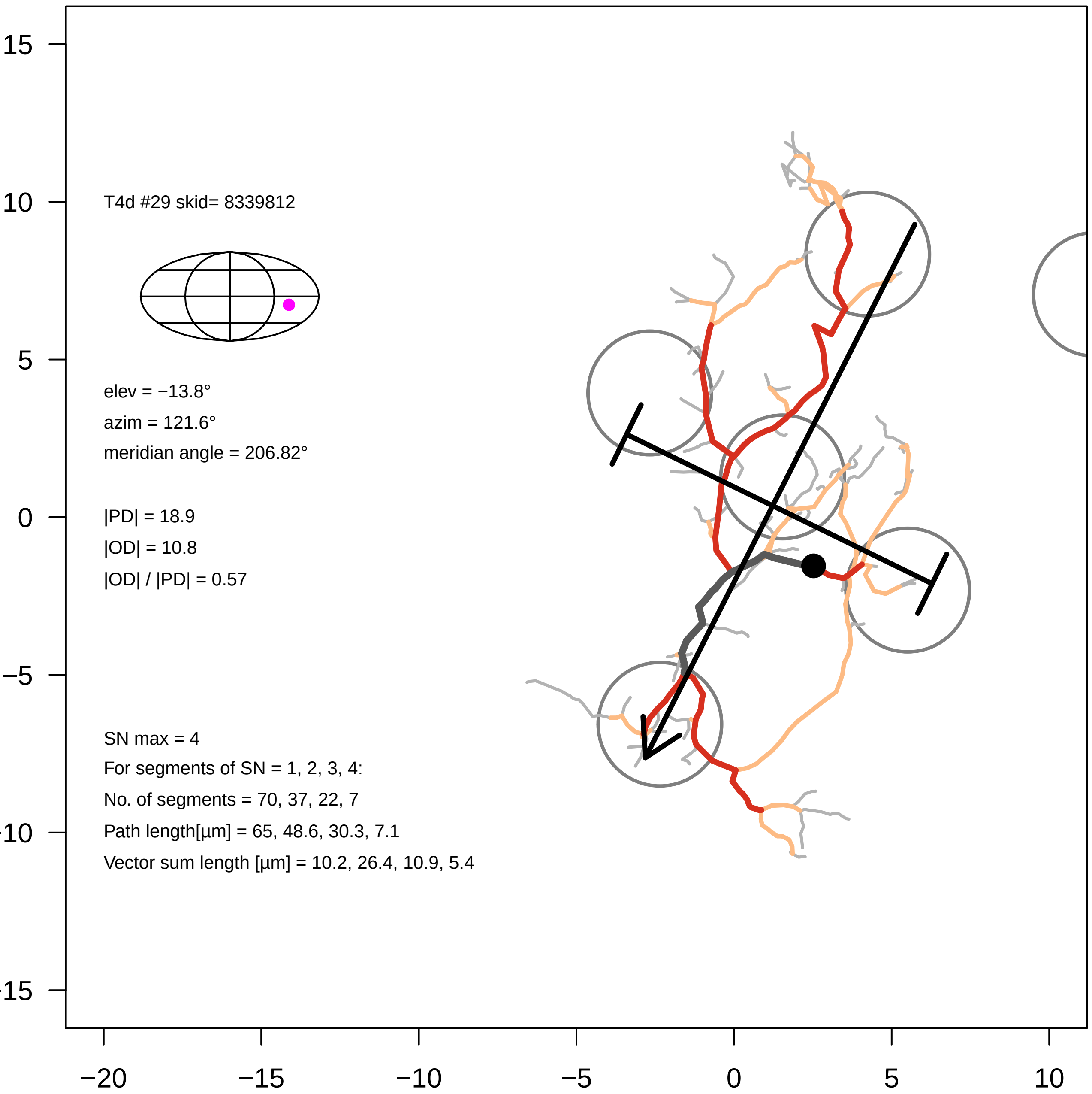

T4d #29 skid= 8339812
elev = −13.8°
azim = 121.6°
meridian angle = 206.82°
|PD| = 18.9
|OD| = 10.8
|OD| / |PD| = 0.57
SN max = 4
For segments of SN = 1, 2, 3, 4:
No. of segments = 70, 37, 22, 7
Path length[µm] = 65, 48.6, 30.3, 7.1
Vector sum length [µm] = 10.2, 26.4, 10.9, 5.4
15
10
5
0
−5
−10
−15
−20
−15
−10
−5
0
5
10

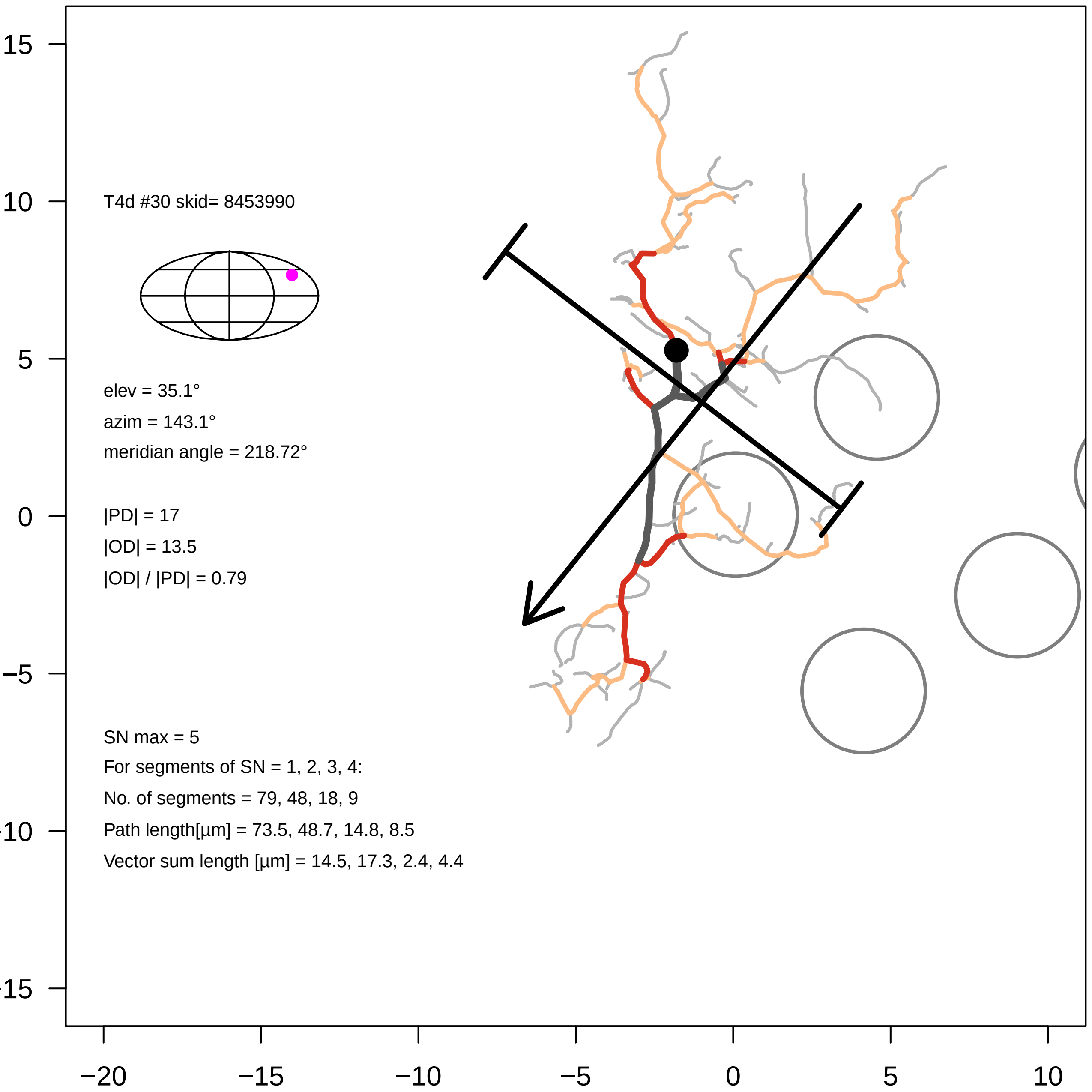
T4d #30 skid= 8453990
elev = 35.1°
azim = 143.1°
meridian angle = 218.72°
|PD| = 17
|OD| = 13.5
|OD| / |PD| = 0.79
SN max = 5
For segments of SN = 1, 2, 3, 4:
No. of segments = 79, 48, 18, 9
Path length[µm] = 73.5, 48.7, 14.8, 8.5
Vector sum length [µm] = 14.5, 17.3, 2.4, 4.4
15
10
5
0
−5
−10
−15
−20
−15
−10
−5
0
5
10

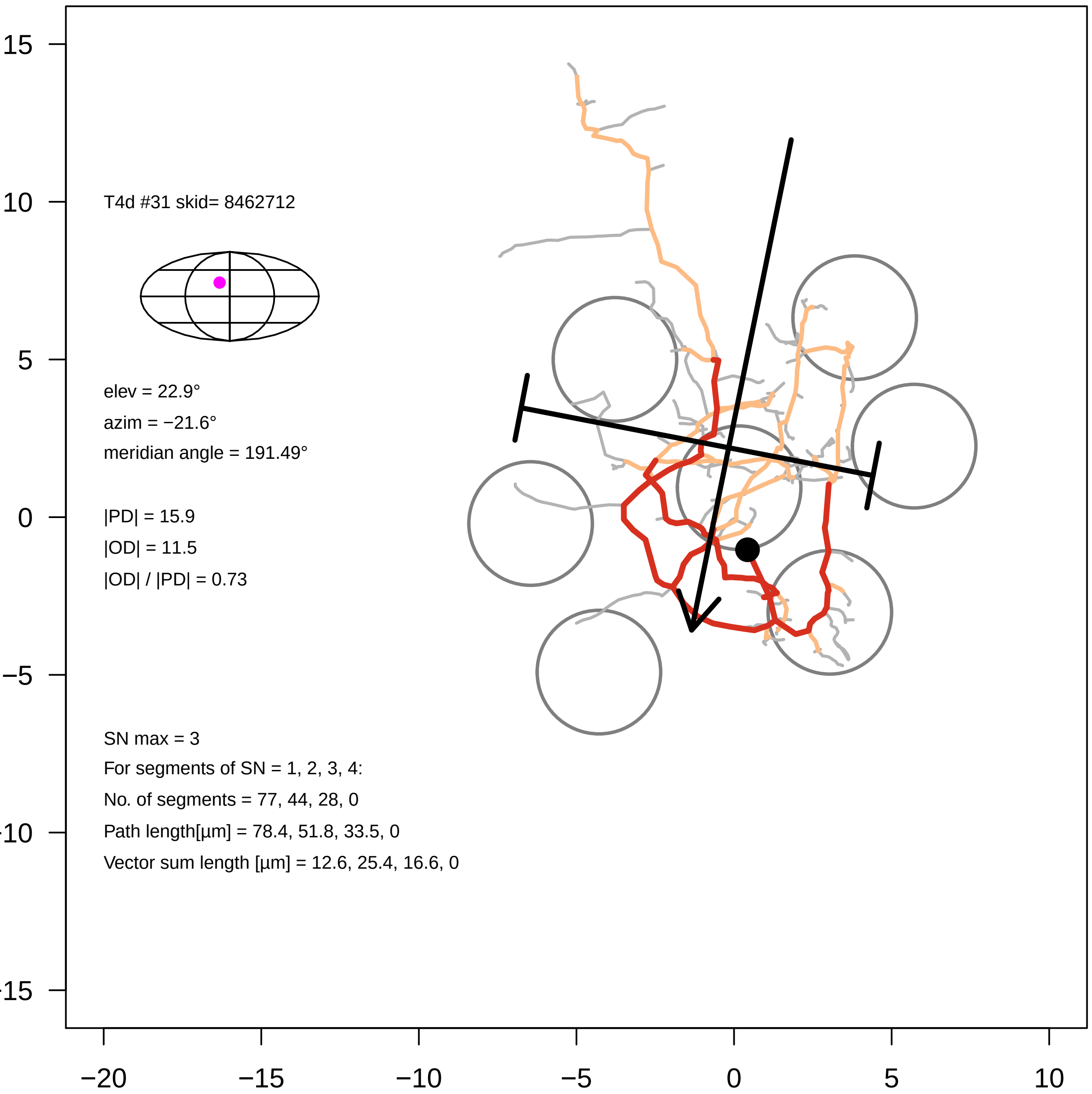

T4d #31 skid= 8462712
elev = 22.9°
azim = −21.6°
meridian angle = 191.49°
|PD| = 15.9
|OD| = 11.5
|OD| / |PD| = 0.73
SN max = 3
For segments of SN = 1, 2, 3, 4:
No. of segments = 77, 44, 28, 0
Path length[µm] = 78.4, 51.8, 33.5, 0
Vector sum length [µm] = 12.6, 25.4, 16.6, 0
15
10
5
0
−5
−20
−15
−10
−5
0
5
10

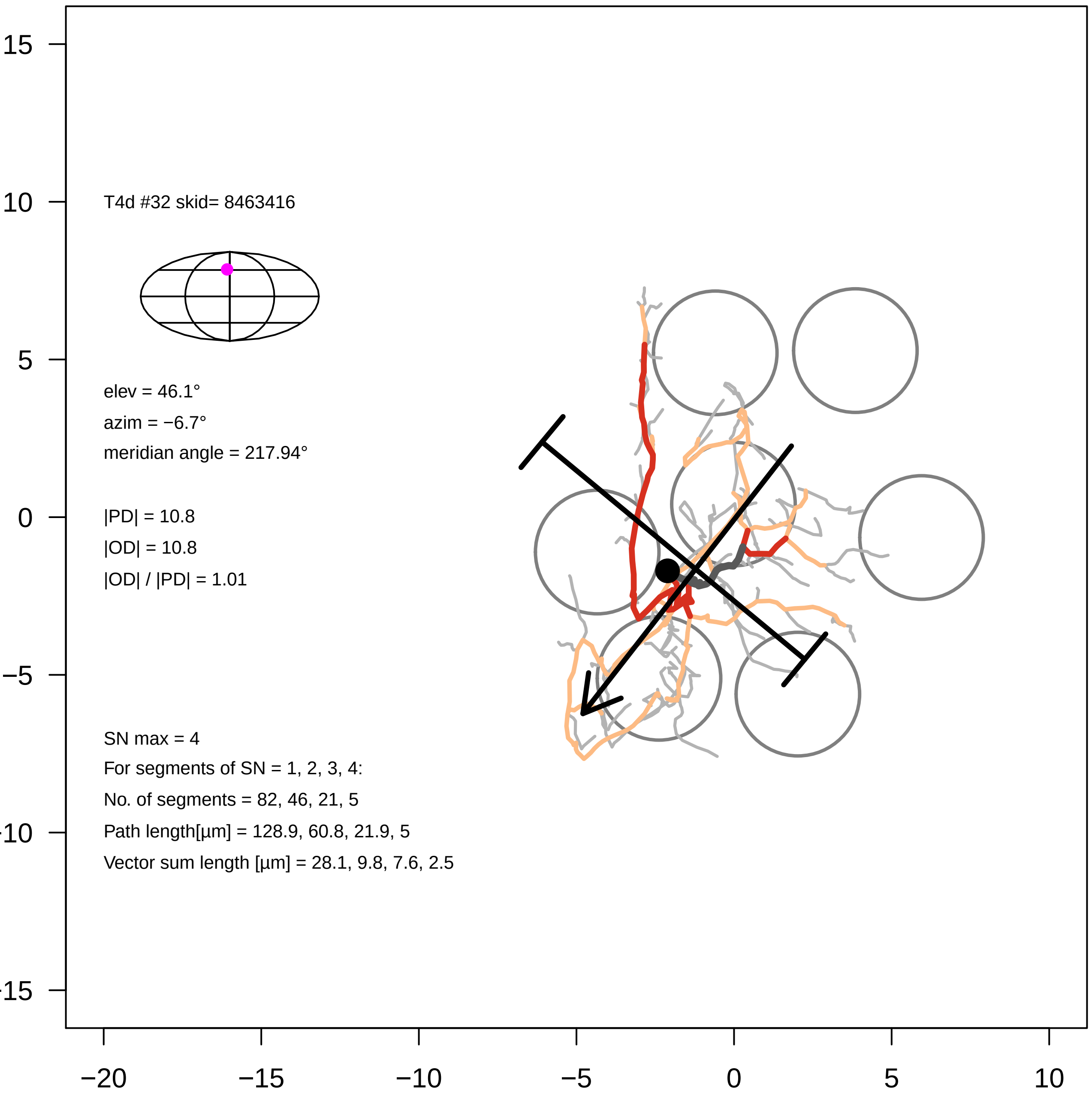

T4d #32 skid= 8463416
elev = 46.1°
azim = −6.7°
meridian angle = 217.94°
|PD| = 10.8
|OD| = 10.8
|OD| / |PD| = 1.01
SN max = 4
For segments of SN = 1, 2, 3, 4:
No. of segments = 82, 46, 21, 5
Path length[µm] = 128.9, 60.8, 21.9, 5
Vector sum length [µm] = 28.1, 9.8, 7.6, 2.5
15
10
5
0
−5
−10
−15
−20
−15
−10
−5
0
5
10

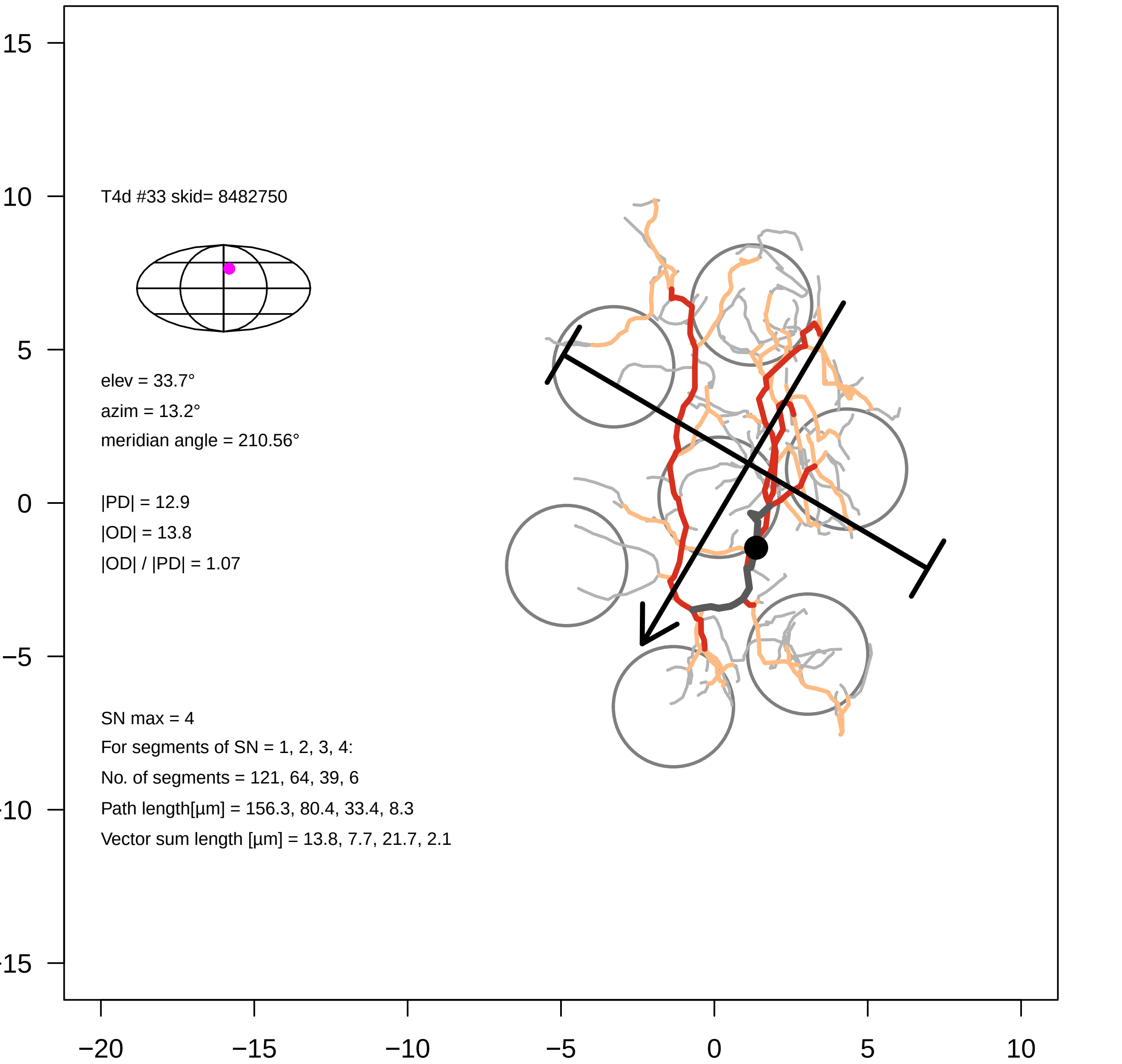
T4d #33 skid= 8482750
elev = 33.7°
azim = 13.2°
meridian angle = 210.56°
|PD| = 12.9
|OD| = 13.8
|OD| / |PD| = 1.07
SN max = 4
For segments of SN = 1, 2, 3, 4:
No. of segments = 121, 64, 39, 6
Path length[µm] = 156.3, 80.4, 33.4, 8.3
Vector sum length [µm] = 13.8, 7.7, 21.7, 2.1
15
10
5
0
−5
−10
−15
−20
−15
−10
−5
0
5
10

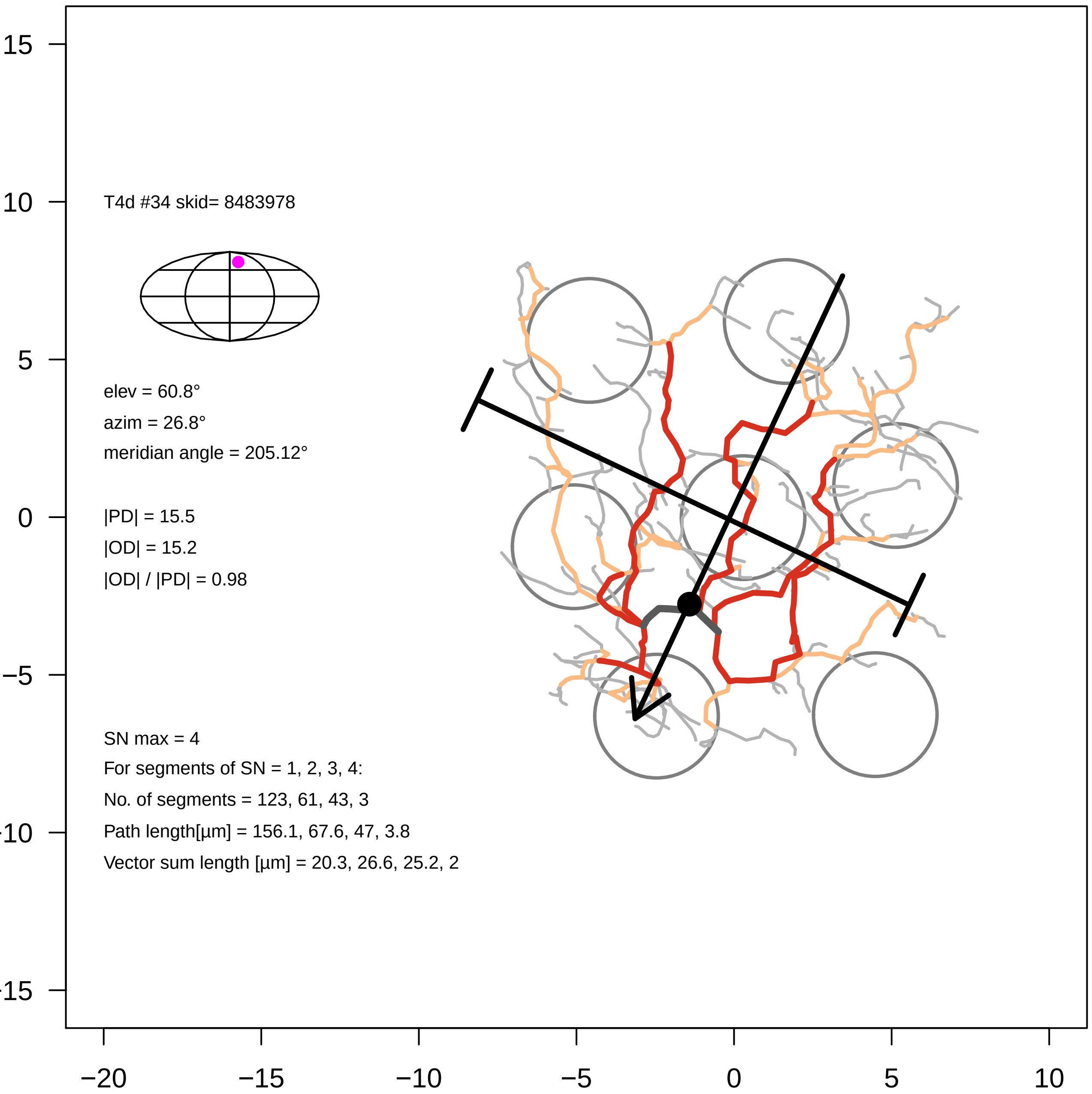

15
10
5
0
−5
−10
−15
−20
−15
−10
−5
0
5
10
T4d #34 skid= 8483978
elev = 60.8°
azim = 26.8°
meridian angle = 205.12°
|PD| = 15.5
|OD| = 15.2
|OD| / |PD| = 0.98
SN max = 4
For segments of SN = 1, 2, 3, 4:
No. of segments = 123, 61, 43, 3
Path length[µm] = 156.1, 67.6, 47, 3.8
Vector sum length [µm] = 20.3, 26.6, 25.2, 2

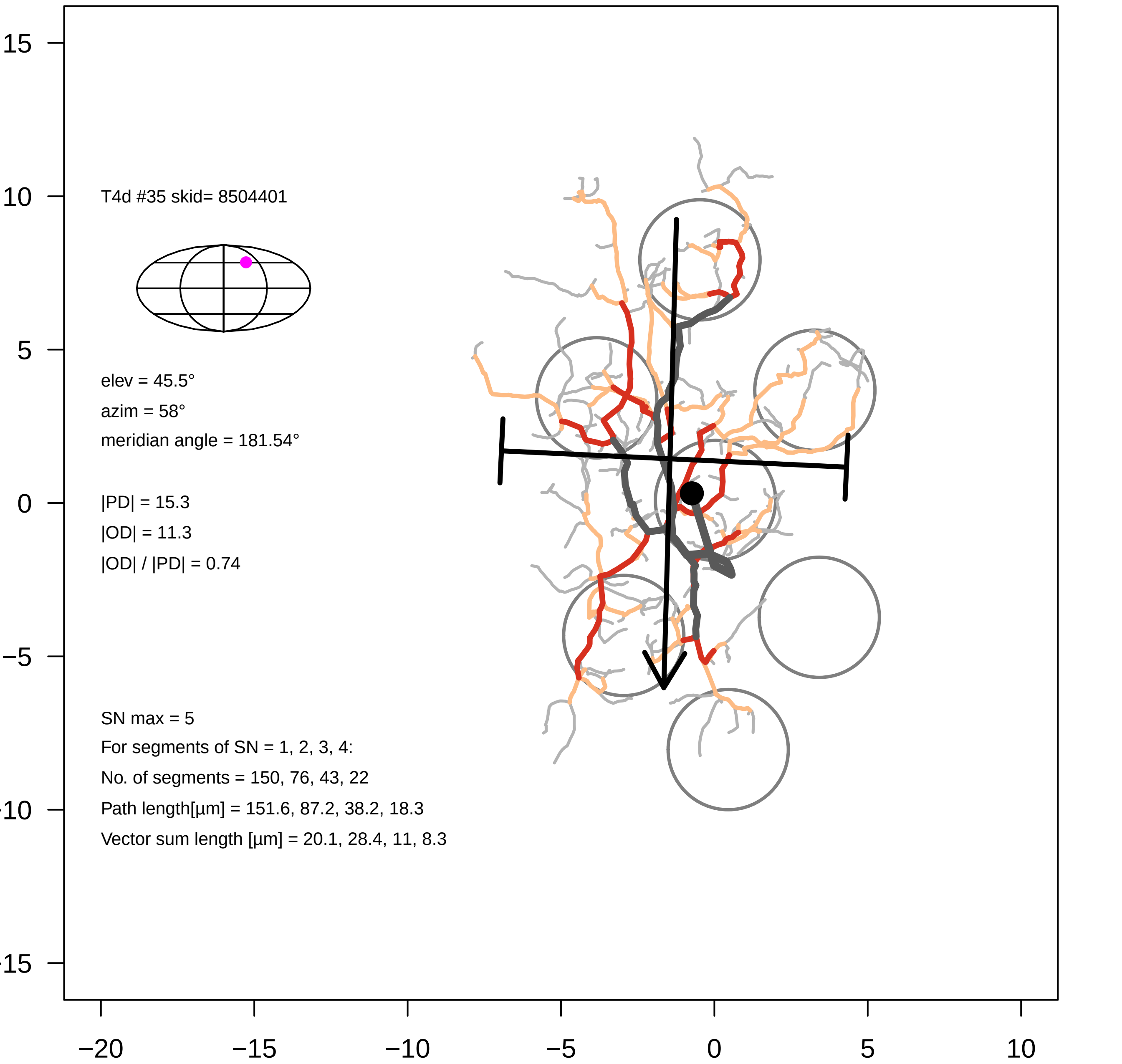

T4d #35 skid= 8504401
elev = 45.5°
azim = 58°
meridian angle = 181.54°
|PD| = 15.3
|OD| = 11.3
|OD| / |PD| = 0.74
SN max = 5
For segments of SN = 1, 2, 3, 4:
No. of segments = 150, 76, 43, 22
Path length[µm] = 151.6, 87.2, 38.2, 18.3
Vector sum length [µm] = 20.1, 28.4, 11, 8.3
15
10
5
0
−5
−10
−15
−20
−15
−10
−5
0
5
10

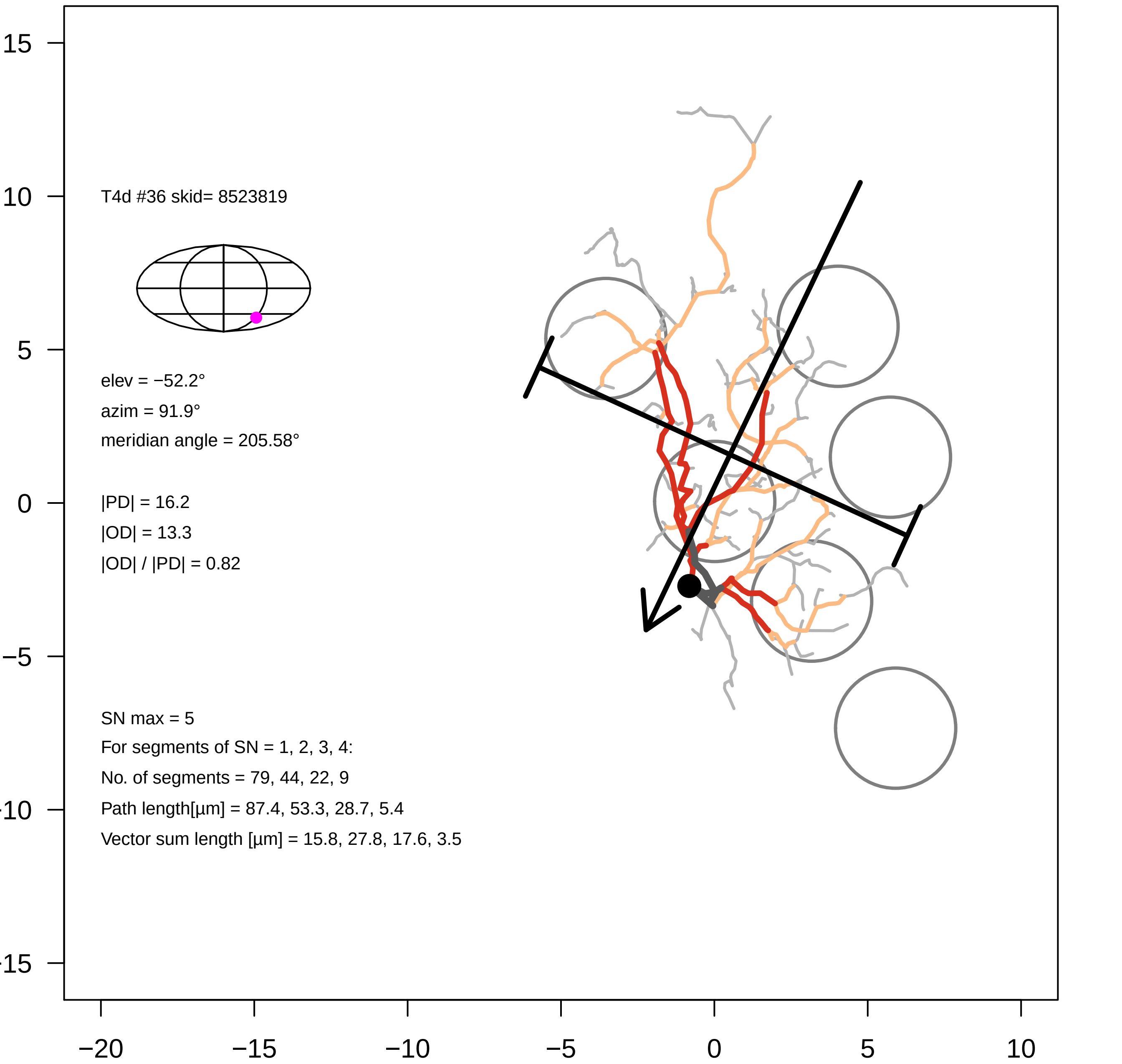
T4d #36 skid= 8523819
elev = −52.2°
azim = 91.9°
meridian angle = 205.58°
|PD| = 16.2
|OD| = 13.3
|OD| / |PD| = 0.82
SN max = 5
For segments of SN = 1, 2, 3, 4:
No. of segments = 79, 44, 22, 9
Path length[µm] = 87.4, 53.3, 28.7, 5.4
Vector sum length [µm] = 15.8, 27.8, 17.6, 3.5
15
10
5
0
−5
−10
−15
−20
−15
−10
−5
0
5
10

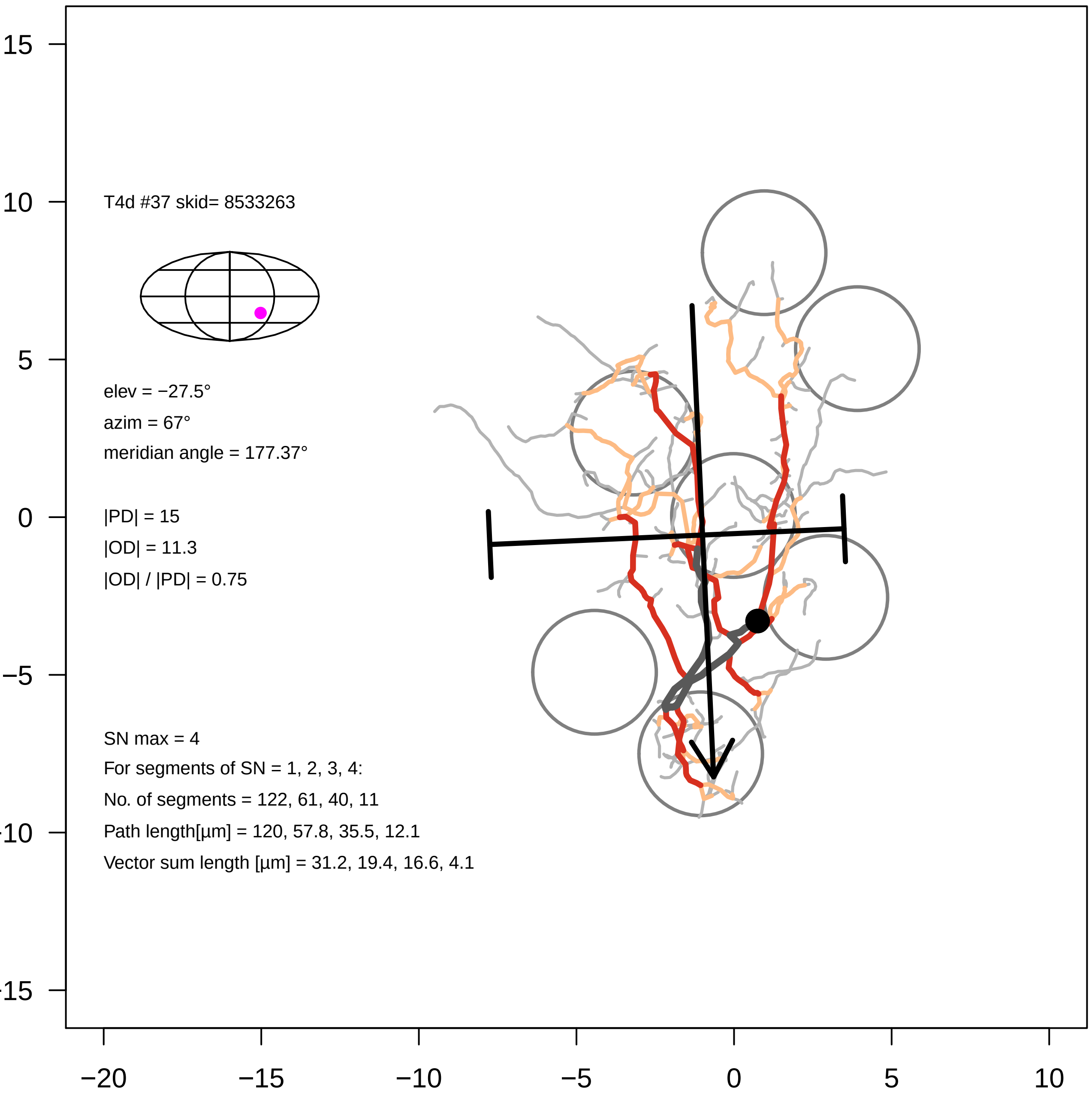

T4d #37 skid= 8533263
elev = −27.5°
azim = 67°
meridian angle = 177.37°
|PD| = 15
|OD| = 11.3
|OD| / |PD| = 0.75
SN max = 4
For segments of SN = 1, 2, 3, 4:
No. of segments = 122, 61, 40, 11
Path length[µm] = 120, 57.8, 35.5, 12.1
Vector sum length [µm] = 31.2, 19.4, 16.6, 4.1
15
10
5
0
−5
−10
−15
−20
−15
−10
−5
0
5
10

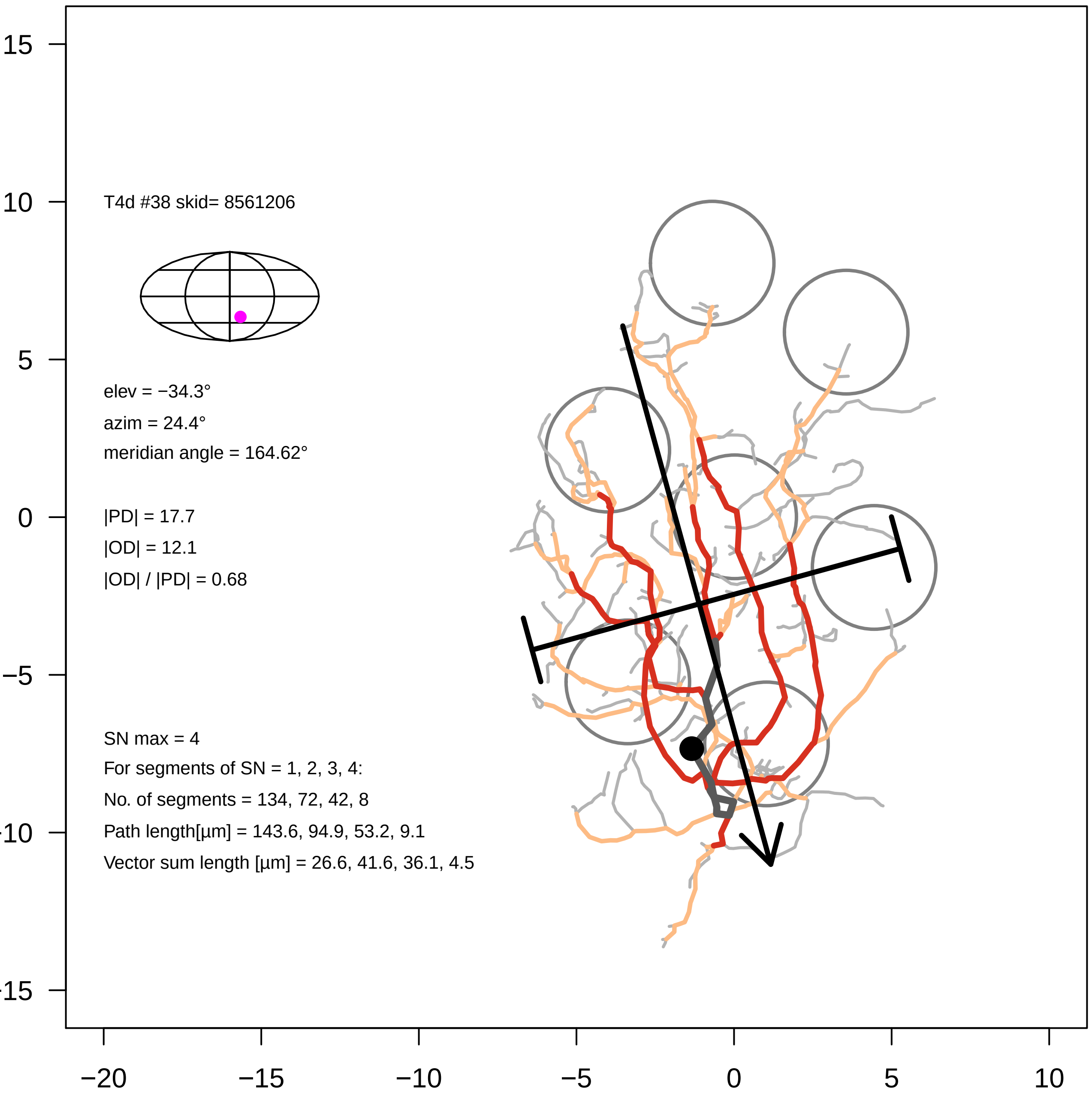

T4d #38 skid= 8561206
elev = −34.3°
azim = 24.4°
meridian angle = 164.62°
|PD| = 17.7
|OD| = 12.1
|OD| / |PD| = 0.68
SN max = 4
For segments of SN = 1, 2, 3, 4:
No. of segments = 134, 72, 42, 8
Path length[µm] = 143.6, 94.9, 53.2, 9.1
Vector sum length [µm] = 26.6, 41.6, 36.1, 4.5
15
10
5
0
−5
−10
−15
−20
−15
−10
−5
0
5
10

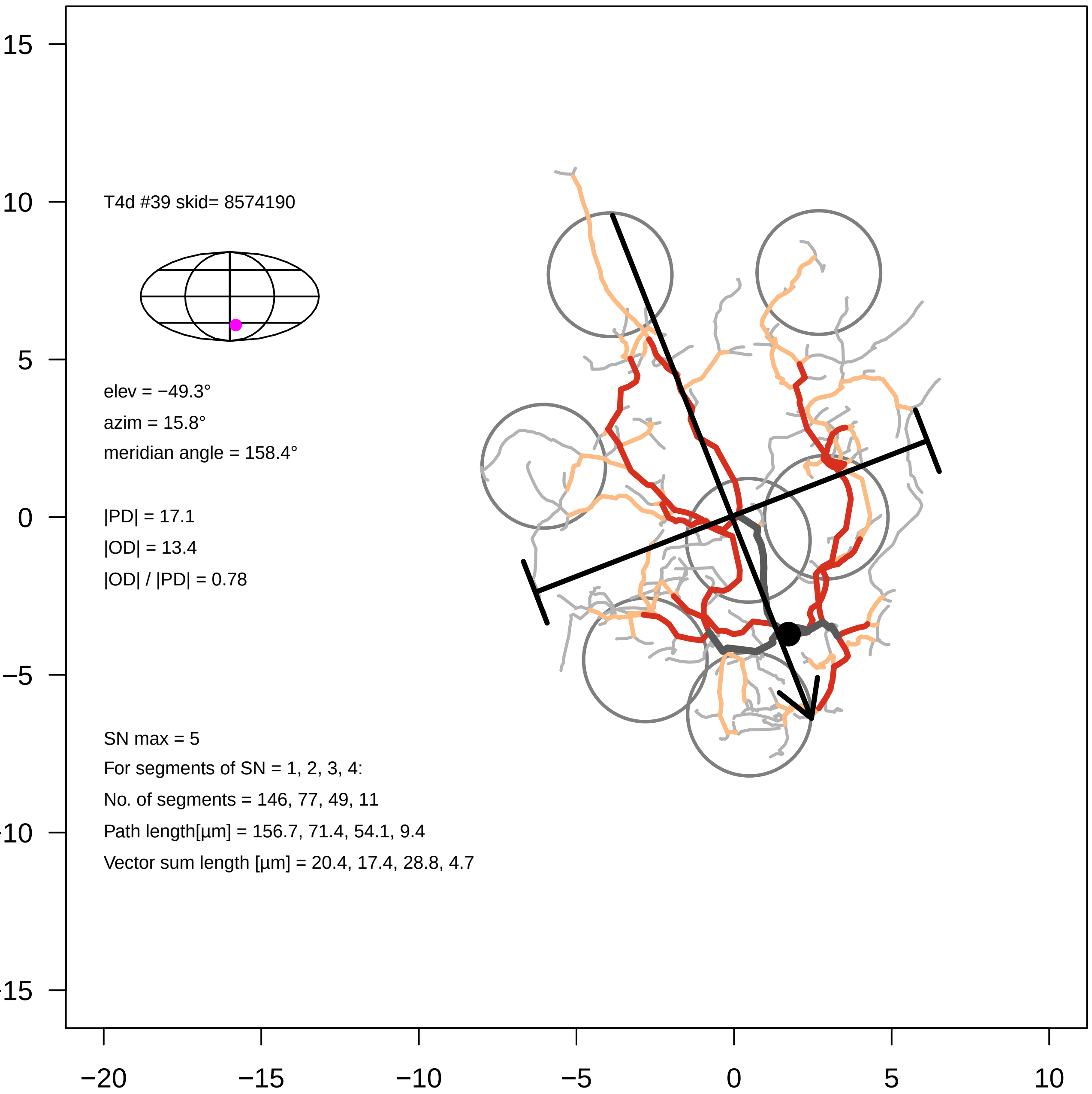
T4d #39 skid= 8574190
elev = −49.3°
azim = 15.8°
meridian angle = 158.4°
|PD| = 17.1
|OD| = 13.4
|OD| / |PD| = 0.78
SN max = 5
For segments of SN = 1, 2, 3, 4:
No. of segments = 146, 77, 49, 11
Path length[µm] = 156.7, 71.4, 54.1, 9.4
Vector sum length [µm] = 20.4, 17.4, 28.8, 4.7
15
10
5
0
−5
−10
−15
−20
−15
−10
−5
0
5
10

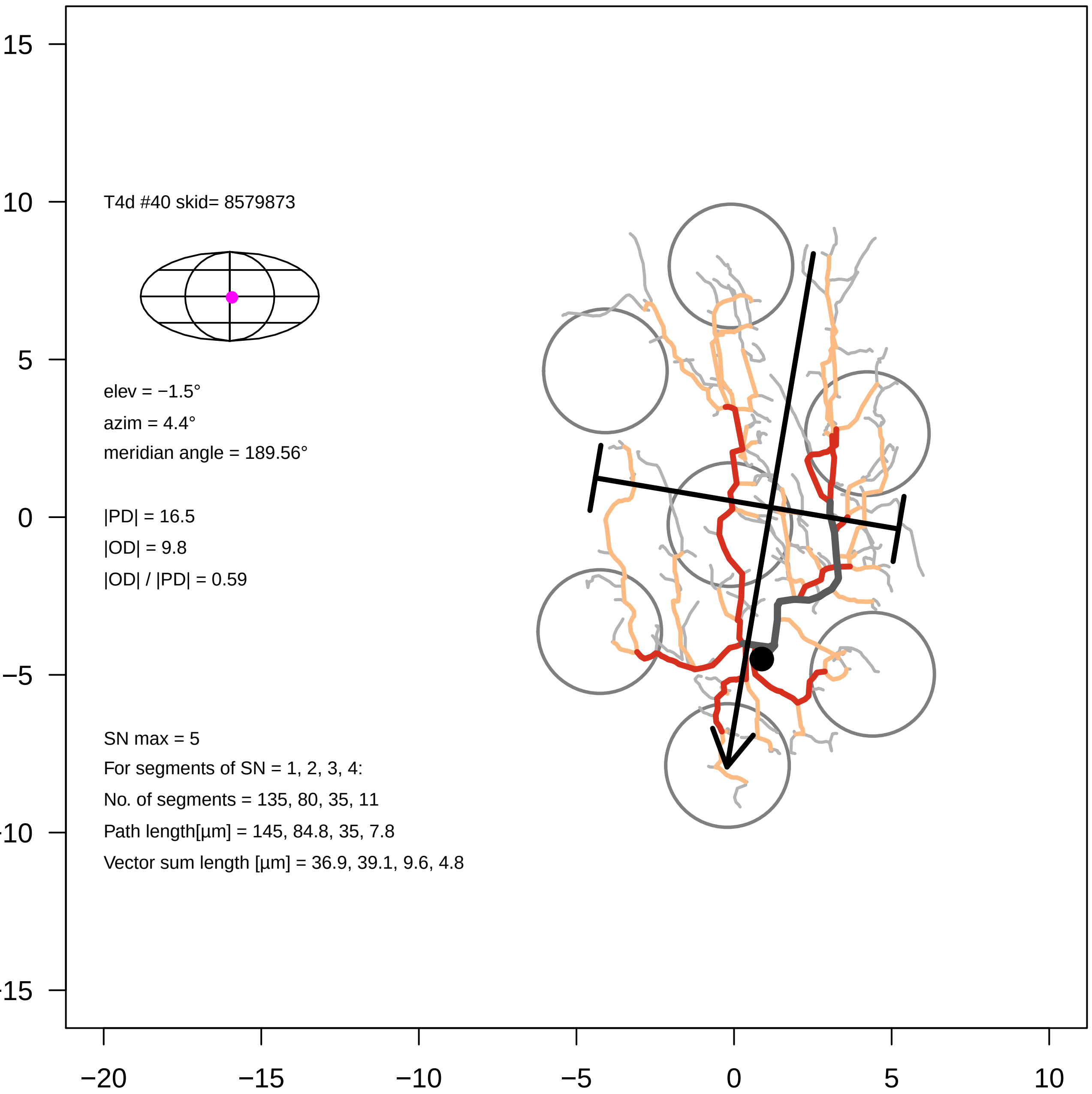
T4d #40 skid= 8579873
elev = −1.5°
azim = 4.4°
meridian angle = 189.56°
|PD| = 16.5
|OD| = 9.8
|OD| / |PD| = 0.59
SN max = 5
For segments of SN = 1, 2, 3, 4:
No. of segments = 135, 80, 35, 11
Path length[µm] = 145, 84.8, 35, 7.8
Vector sum length [µm] = 36.9, 39.1, 9.6, 4.8
15
10
5
0
−5
−10
−15
−20
−15
−10
−5
0
5
10

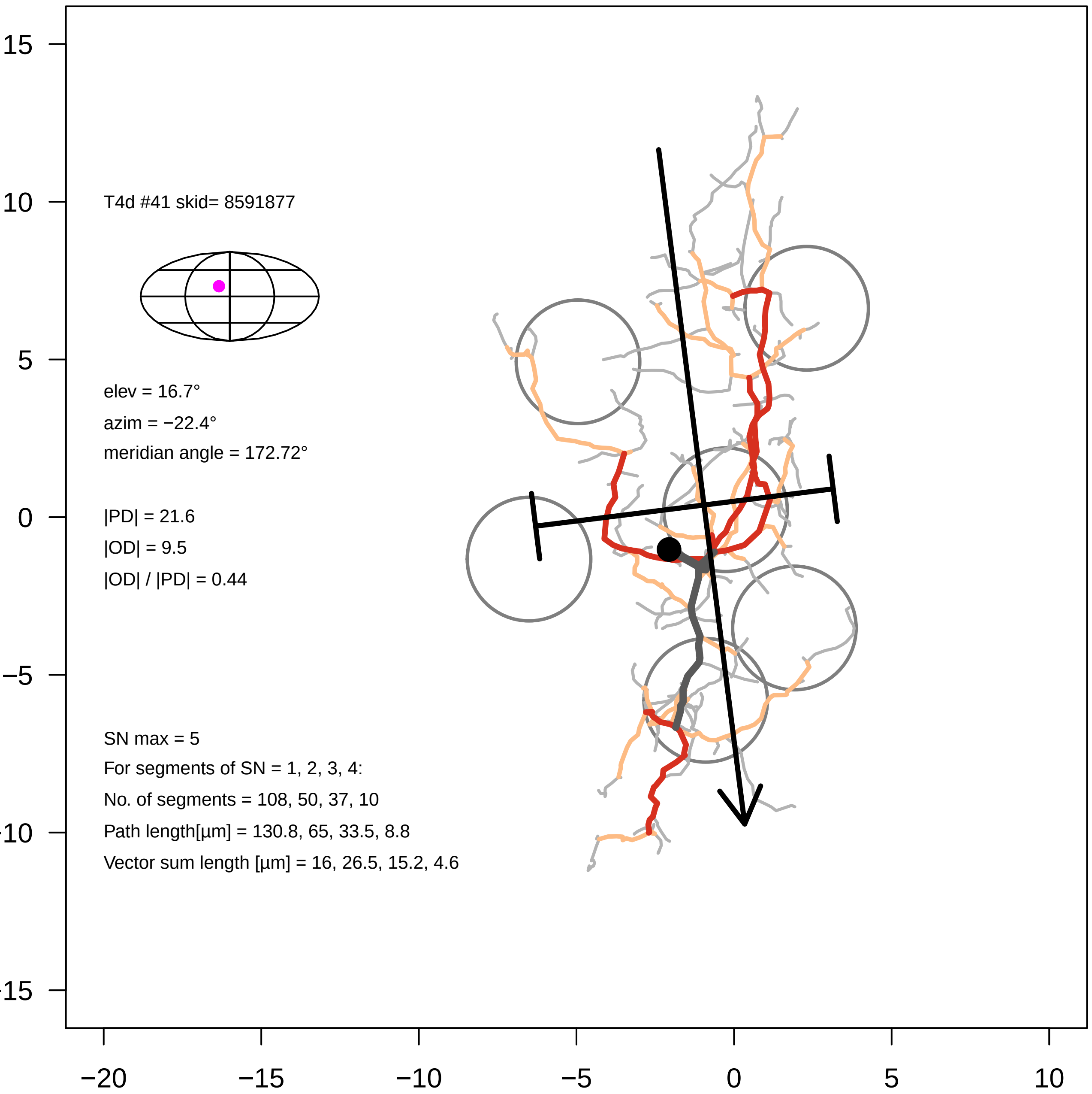

T4d #41 skid= 8591877
elev = 16.7°
azim = −22.4°
meridian angle = 172.72°
|PD| = 21.6
|OD| = 9.5
|OD| / |PD| = 0.44
SN max = 5
For segments of SN = 1, 2, 3, 4:
No. of segments = 108, 50, 37, 10
Path length[µm] = 130.8, 65, 33.5, 8.8
Vector sum length [µm] = 16, 26.5, 15.2, 4.6
15
10
5
0
−5
−10
−15
−20
−15
−10
−5
0
5
10

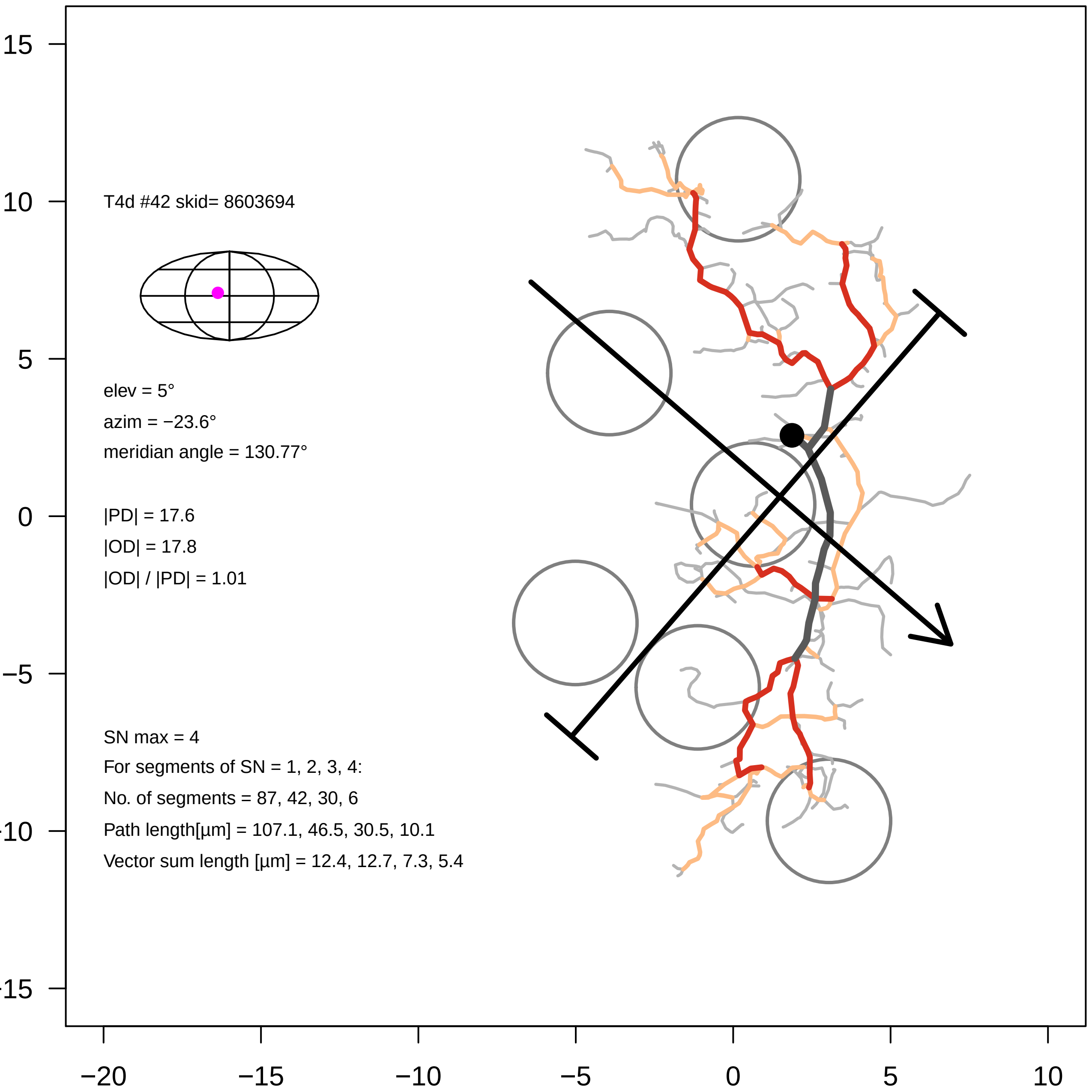
T4d #42 skid= 8603694
elev = 5°
azim = −23.6°
meridian angle = 130.77°
|PD| = 17.6
|OD| = 17.8
|OD| / |PD| = 1.01
SN max = 4
For segments of SN = 1, 2, 3, 4:
No. of segments = 87, 42, 30, 6
Path length[µm] = 107.1, 46.5, 30.5, 10.1
Vector sum length [µm] = 12.4, 12.7, 7.3, 5.4
15
10
5
0
−5
−10
−15
−20
−15
−10
−5
0
5
10

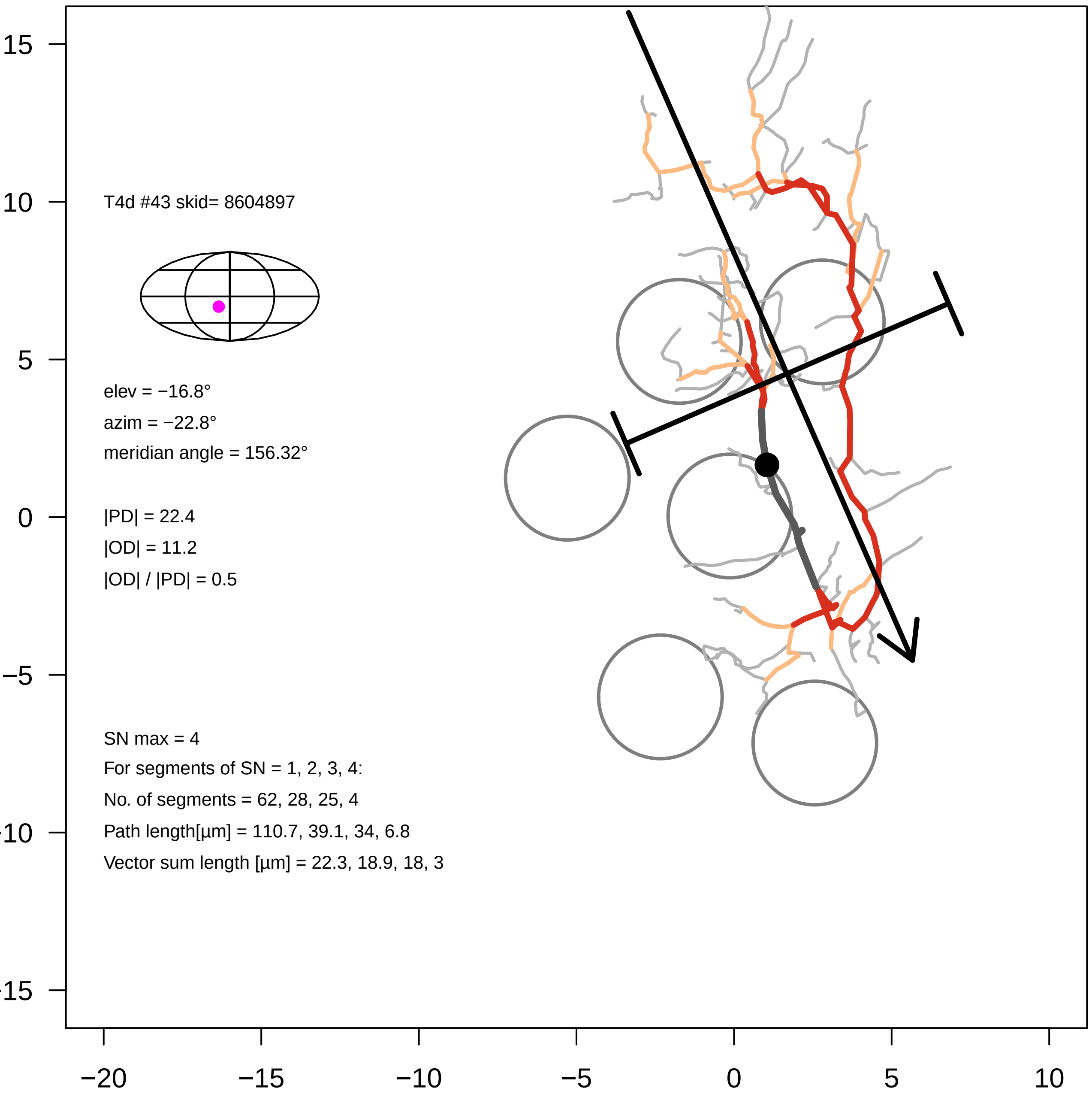

T4d #43 skid= 8604897
elev = −16.8°
azim = −22.8°
meridian angle = 156.32°
|PD| = 22.4
|OD| = 11.2
|OD| / |PD| = 0.5
SN max = 4
For segments of SN = 1, 2, 3, 4:
No. of segments = 62, 28, 25, 4
Path length[µm] = 110.7, 39.1, 34, 6.8
Vector sum length [µm] = 22.3, 18.9, 18, 3
15
10
5
0
−5
-10
-15
−20
−15
−10
−5
0
5
10

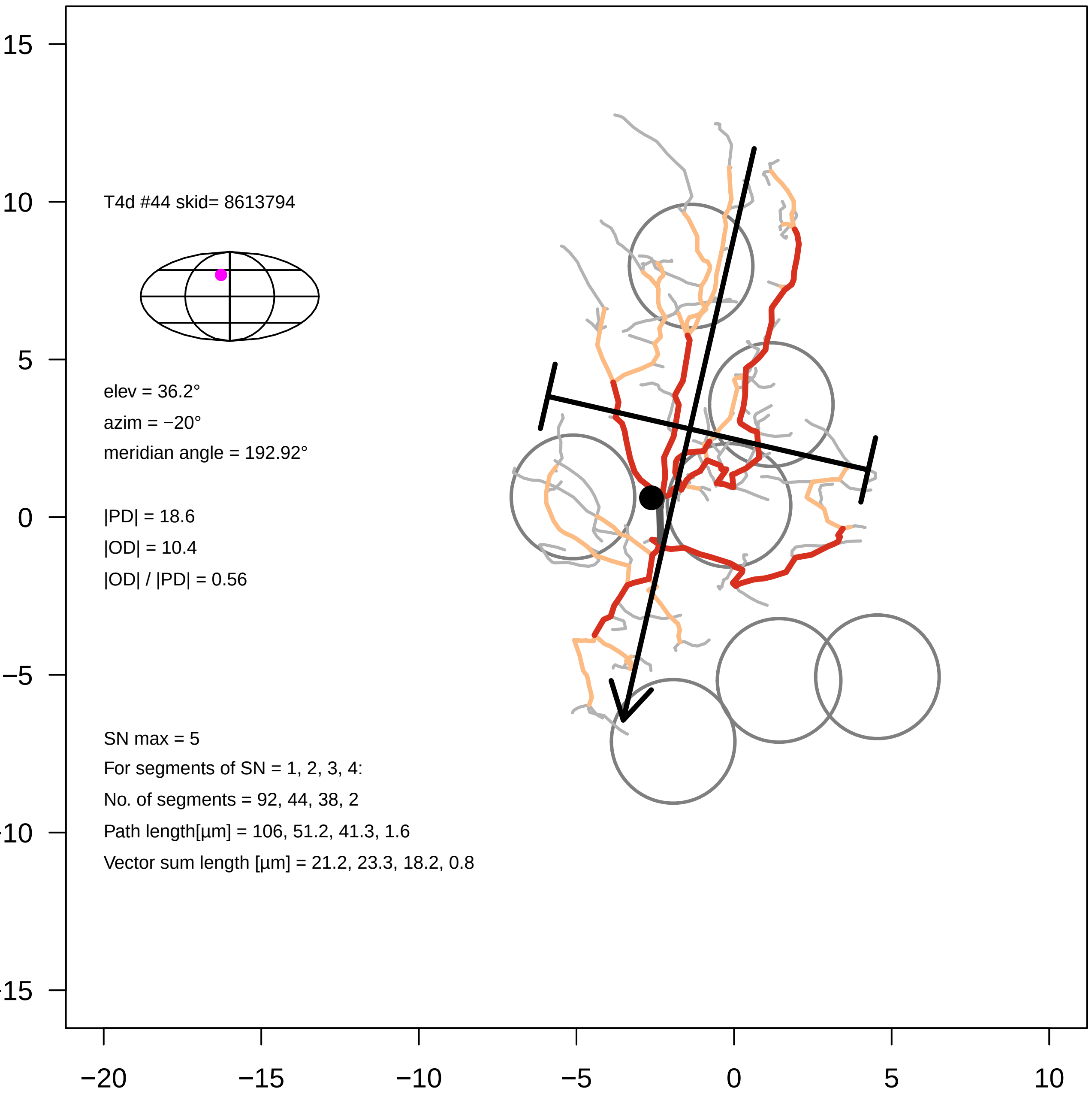

T4d #44 skid= 8613794
elev = 36.2°
azim = −20°
meridian angle = 192.92°
|PD| = 18.6
|OD| = 10.4
|OD| / |PD| = 0.56
SN max = 5
For segments of SN = 1, 2, 3, 4:
No. of segments = 92, 44, 38, 2
Path length[µm] = 106, 51.2, 41.3, 1.6
Vector sum length [µm] = 21.2, 23.3, 18.2, 0.8
15
10
5
0
−5
−10
−15
−20
−15
−10
−5
0
5
10

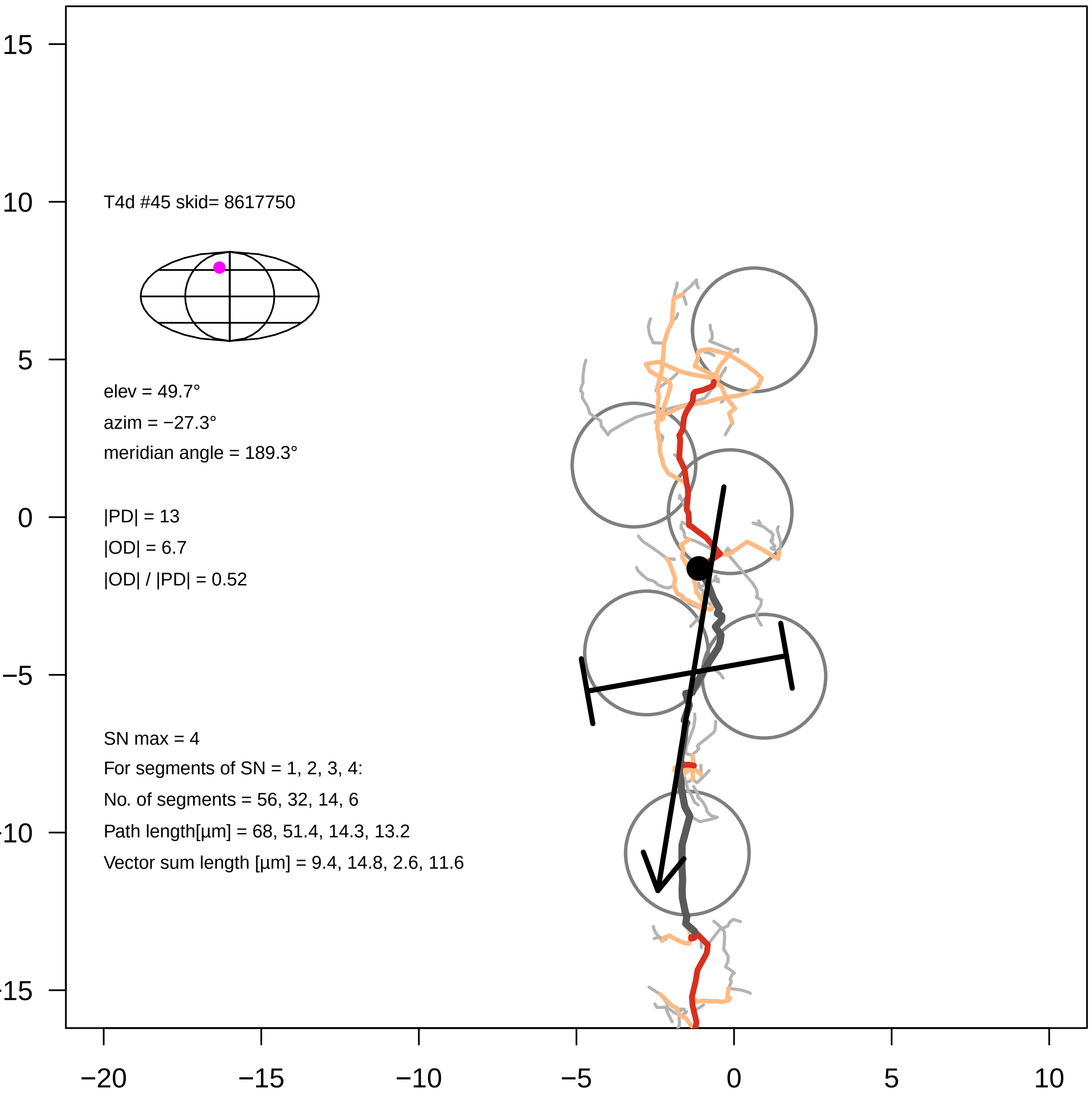

T4d #45 skid= 8617750
elev = 49.7°
azim = −27.3°
meridian angle = 189.3°
|PD| = 13
|OD| = 6.7
|OD| / |PD| = 0.52
SN max = 4
For segments of SN = 1, 2, 3, 4:
No. of segments = 56, 32, 14, 6
Path length[µm] = 68, 51.4, 14.3, 13.2
Vector sum length [µm] = 9.4, 14.8, 2.6, 11.6
15
10
5
0
−5
−10
−15
−20
−15
−10
−5
0
5
10

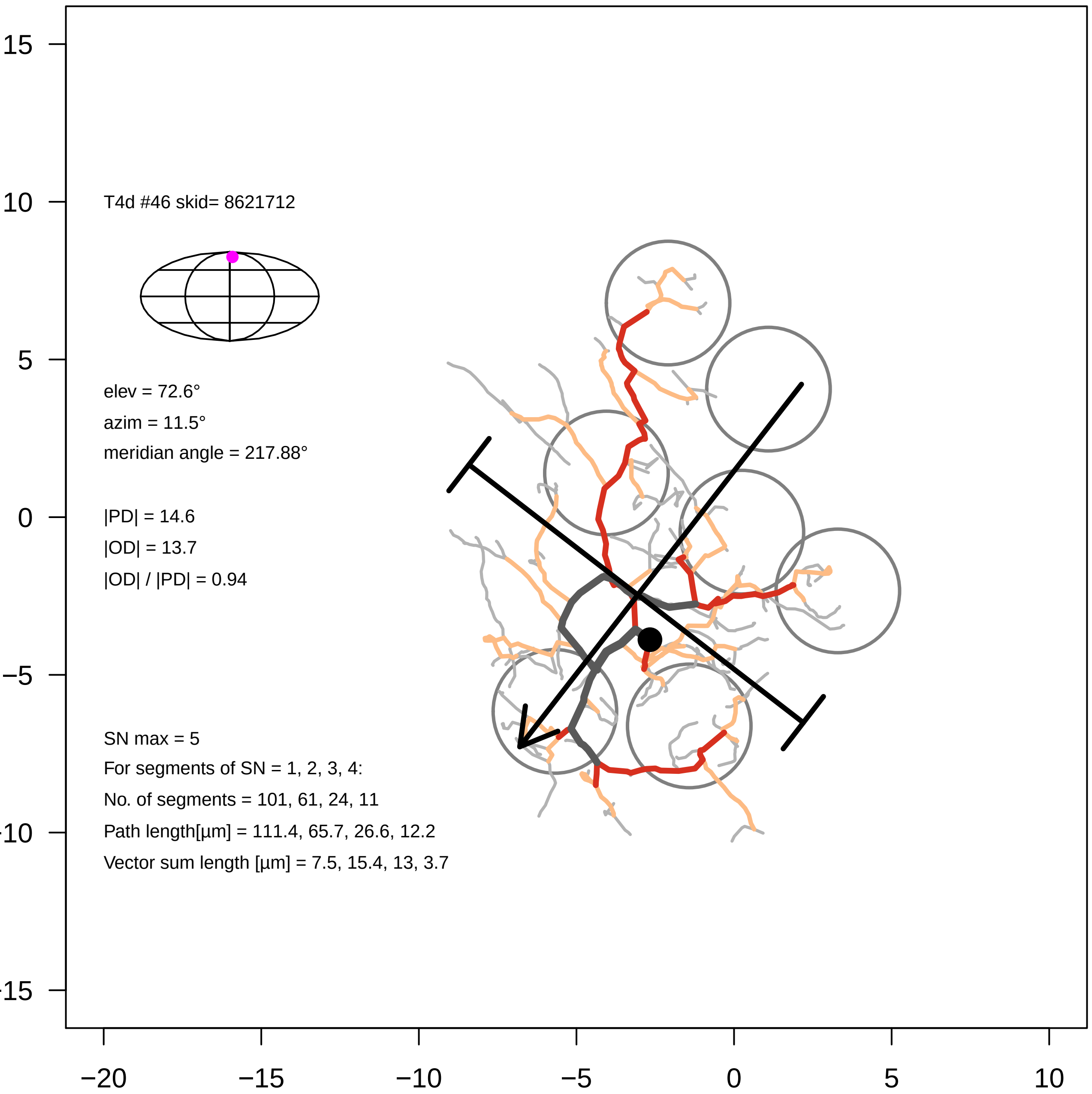

T4d #46 skid= 8621712
elev = 72.6°
azim = 11.5°
meridian angle = 217.88°
|PD| = 14.6
|OD| = 13.7
|OD| / |PD| = 0.94
SN max = 5
For segments of SN = 1, 2, 3, 4:
No. of segments = 101, 61, 24, 11
Path length[µm] = 111.4, 65.7, 26.6, 12.2
Vector sum length [µm] = 7.5, 15.4, 13, 3.7
15
10
5
0
−5
−10
−15
−20
−15
−10
−5
0
5
10

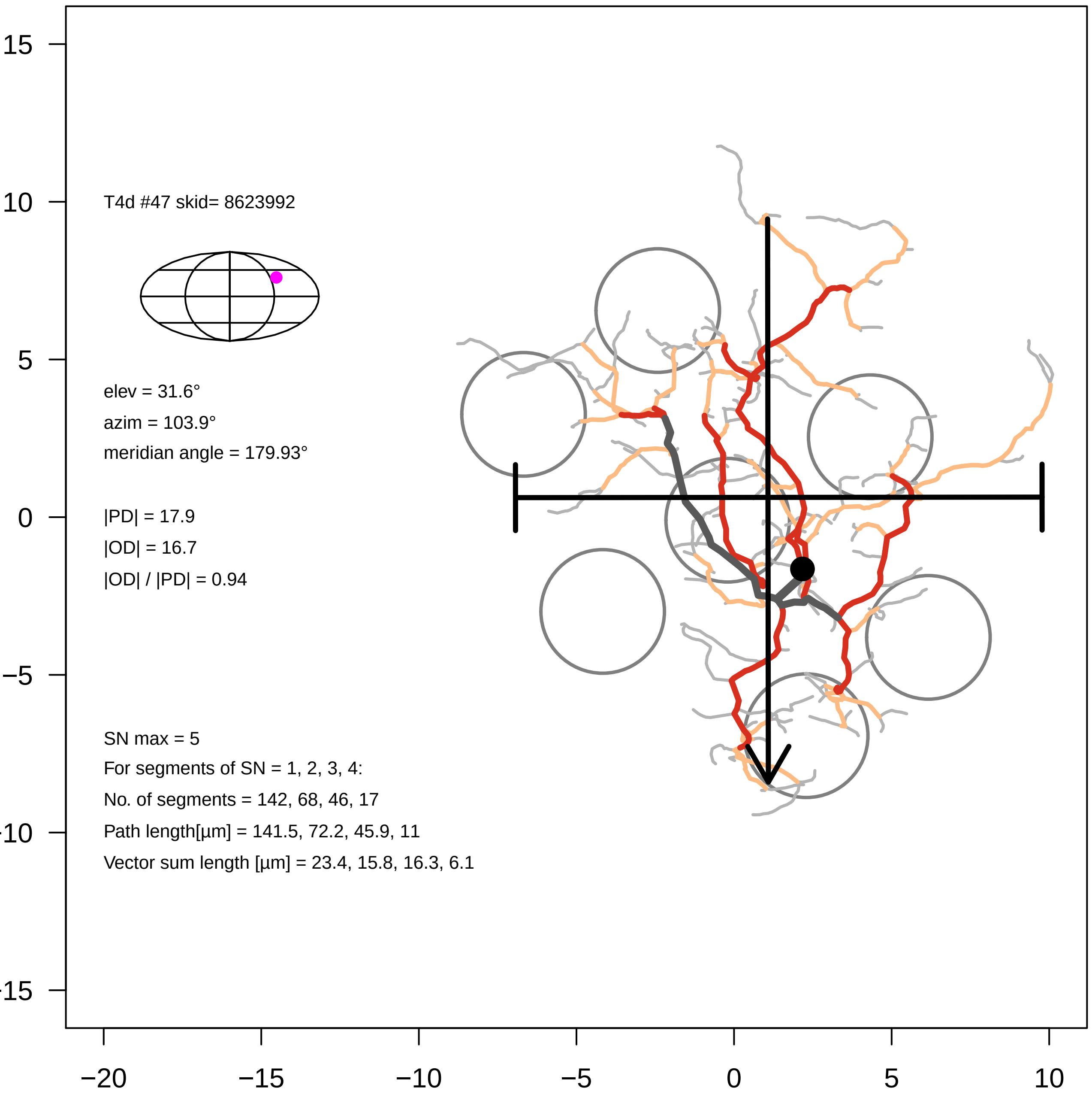
T4d #47 skid= 8623992
elev = 31.6°
azim = 103.9°
meridian angle = 179.93°
|PD| = 17.9
|OD| = 16.7
|OD| / |PD| = 0.94
SN max = 5
For segments of SN = 1, 2, 3, 4:
No. of segments = 142, 68, 46, 17
Path length[µm] = 141.5, 72.2, 45.9, 11
Vector sum length [µm] = 23.4, 15.8, 16.3, 6.1
15
10
5
0
−5
−10
−15
−20
−15
−10
−5
0
5
10

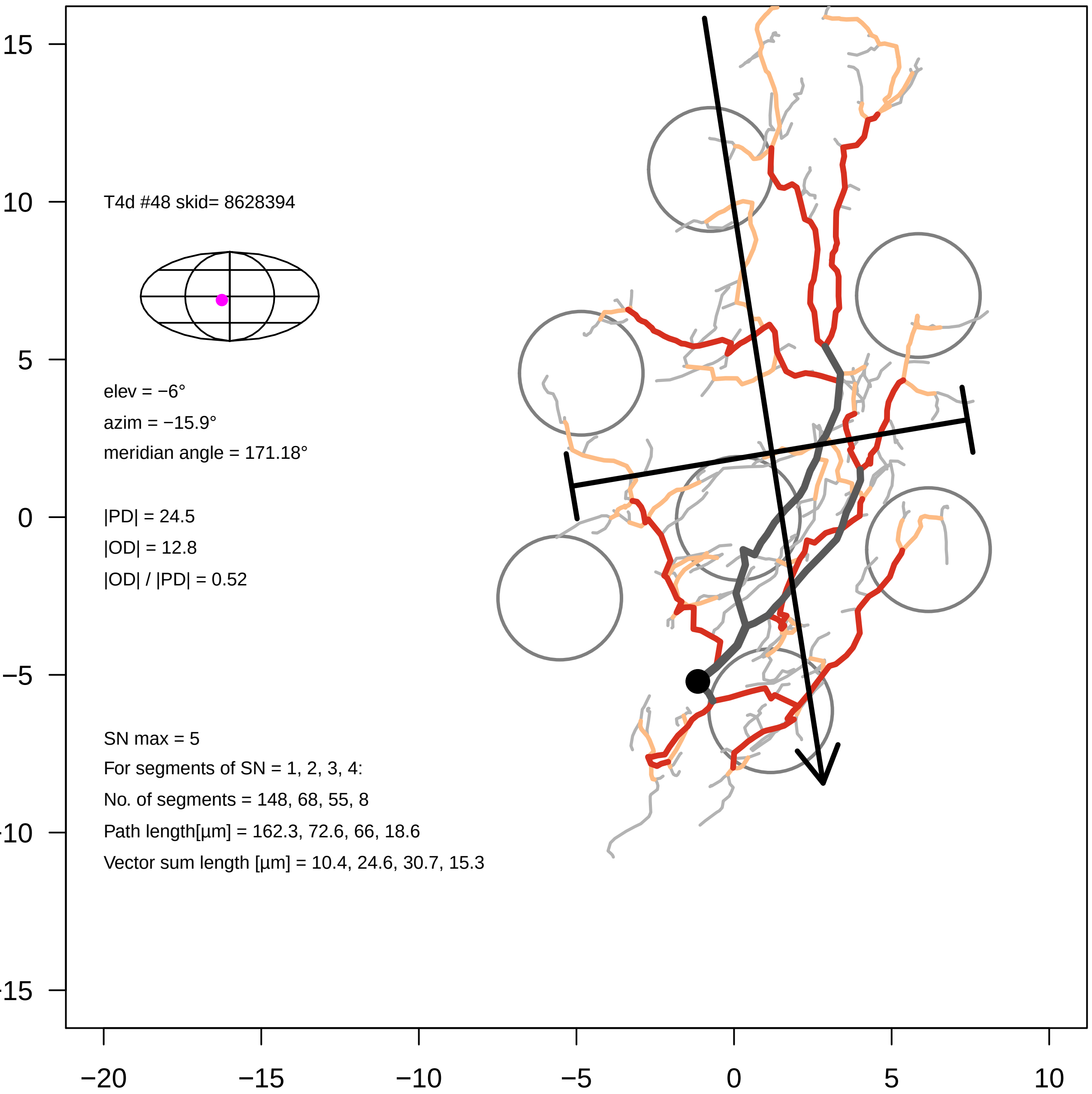

T4d #48 skid= 8628394
elev = −6°
azim = −15.9°
meridian angle = 171.18°
|PD| = 24.5
|OD| = 12.8
|OD| / |PD| = 0.52
SN max = 5
For segments of SN = 1, 2, 3, 4:
No. of segments = 148, 68, 55, 8
Path length[µm] = 162.3, 72.6, 66, 18.6
Vector sum length [µm] = 10.4, 24.6, 30.7, 15.3
15
10
5
0
−5
−10
−15
−20
−15
−10
−5
0
5
10

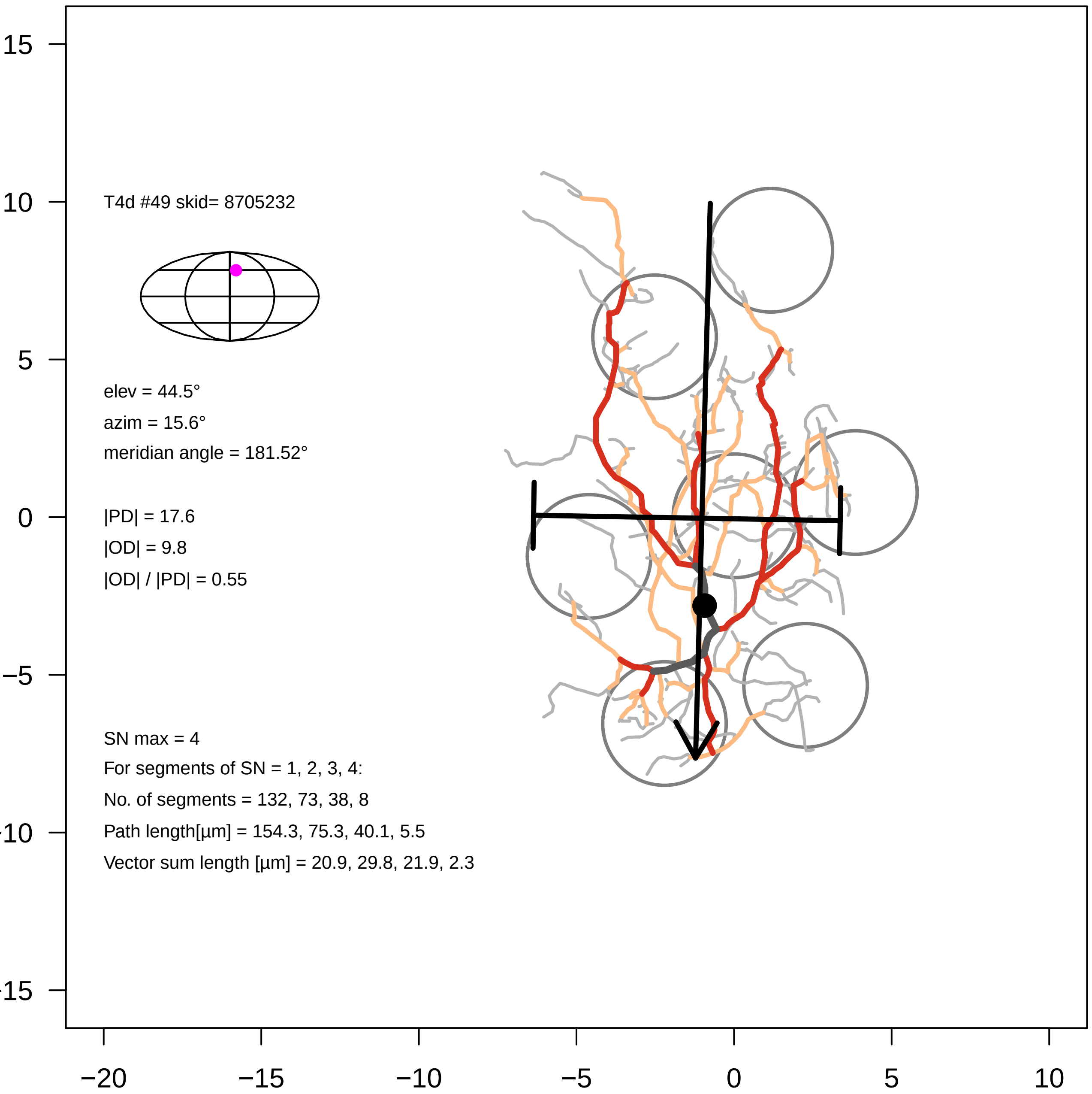
T4d #49 skid= 8705232
elev = 44.5°
azim = 15.6°
meridian angle = 181.52°
|PD| = 17.6
|OD| = 9.8
|OD| / |PD| = 0.55
SN max = 4
For segments of SN = 1, 2, 3, 4:
No. of segments = 132, 73, 38, 8
Path length[µm] = 154.3, 75.3, 40.1, 5.5
Vector sum length [µm] = 20.9, 29.8, 21.9, 2.3
15
10
5
0
−5
−10
−15
−20
−15
−10
−5
0
5
10

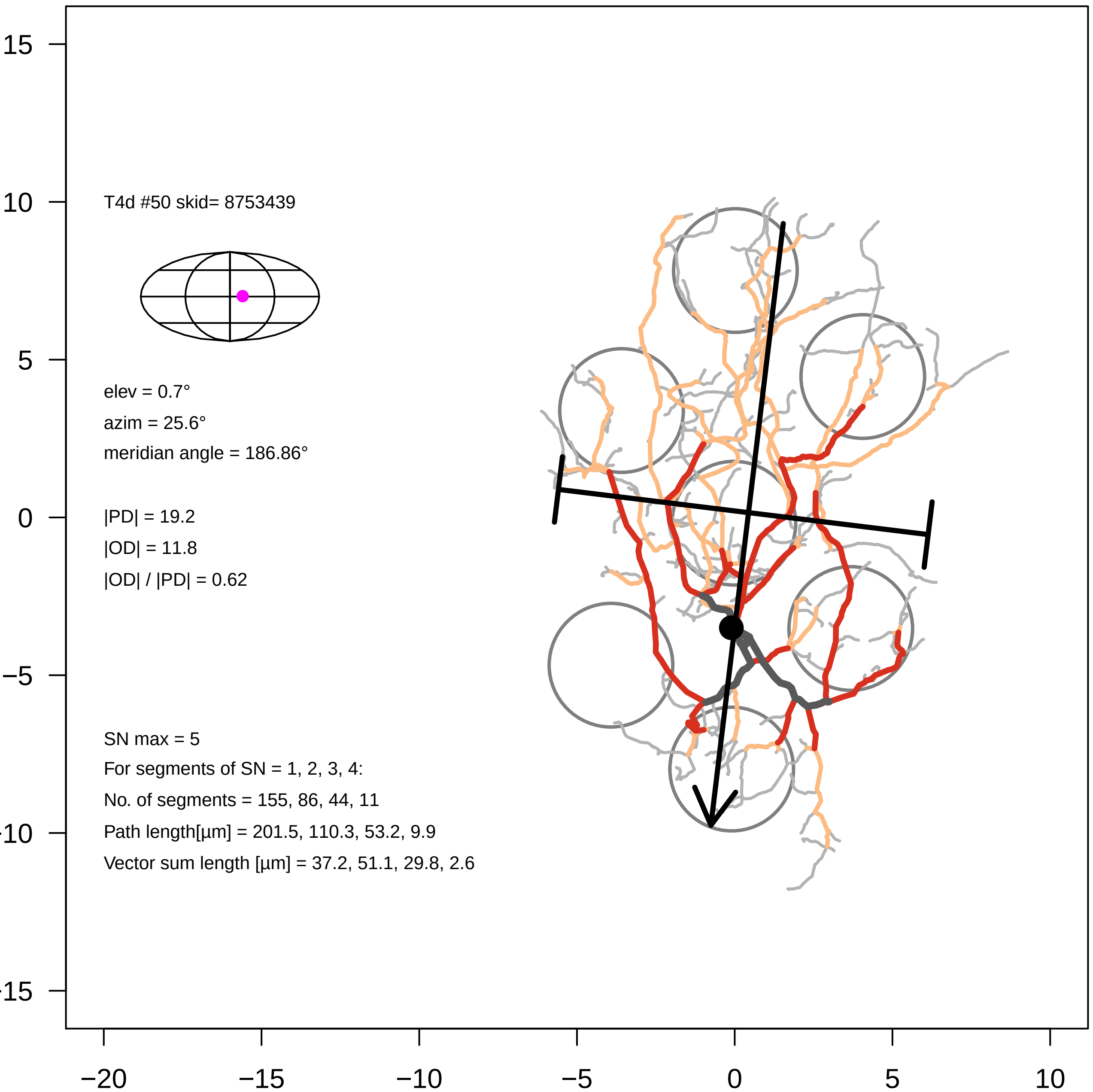

T4d #50 skid= 8753439
elev = 0.7°
azim = 25.6°
meridian angle = 186.86°
|PD| = 19.2
|OD| = 11.8
|OD| / |PD| = 0.62
SN max = 5
For segments of SN = 1, 2, 3, 4:
No. of segments = 155, 86, 44, 11
Path length[µm] = 201.5, 110.3, 53.2, 9.9
Vector sum length [µm] = 37.2, 51.1, 29.8, 2.6
15
10
5
0
−5
−10
−15
−20
−15
−10
−5
0
5
10

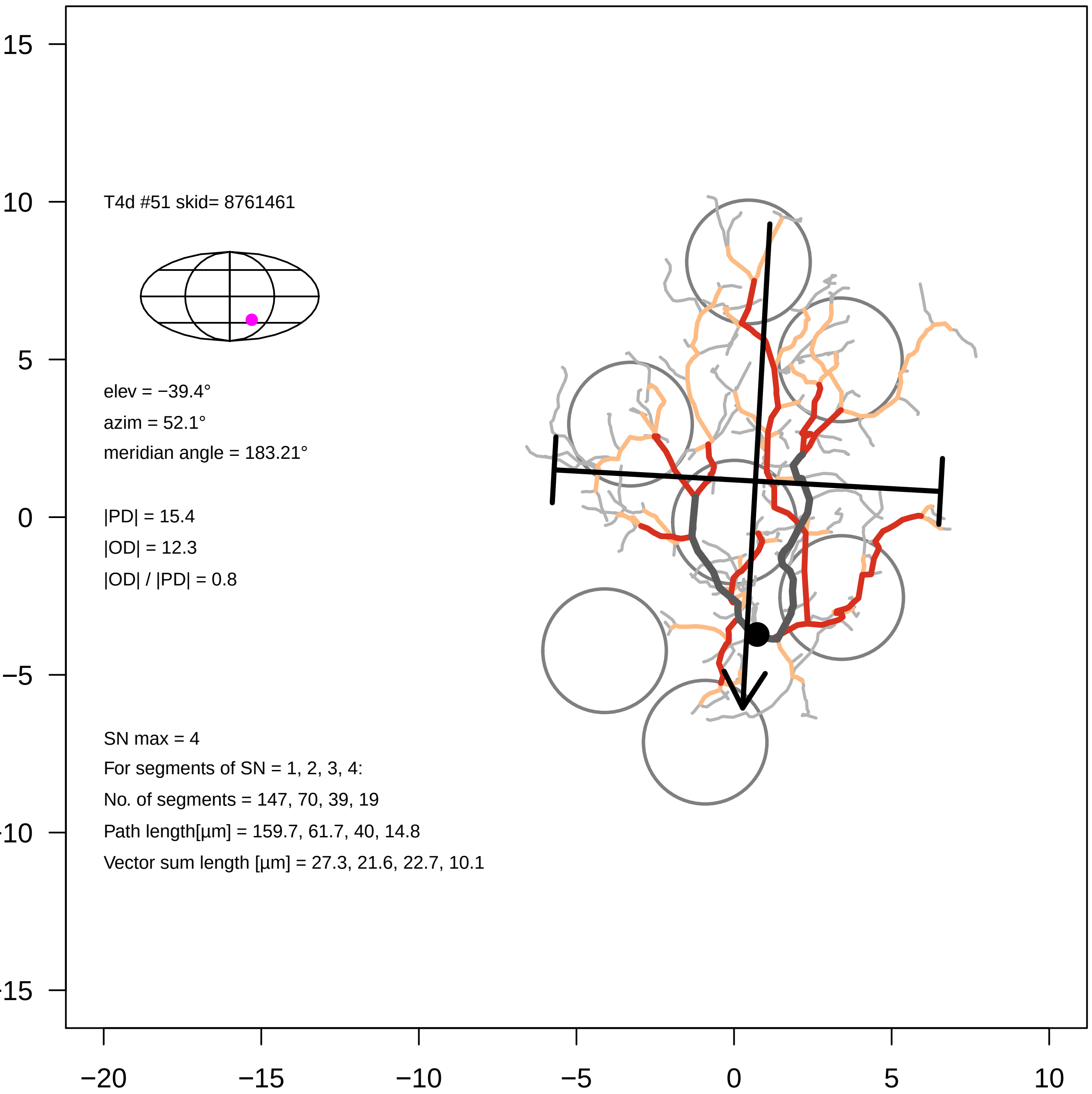

T4d #51 skid= 8761461
elev = −39.4°
azim = 52.1°
meridian angle = 183.21°
|PD| = 15.4
|OD| = 12.3
|OD| / |PD| = 0.8
SN max = 4
For segments of SN = 1, 2, 3, 4:
No. of segments = 147, 70, 39, 19
Path length[µm] = 159.7, 61.7, 40, 14.8
Vector sum length [µm] = 27.3, 21.6, 22.7, 10.1
15
10
5
0
−5
−20
−15
−10
−5
0
5
10

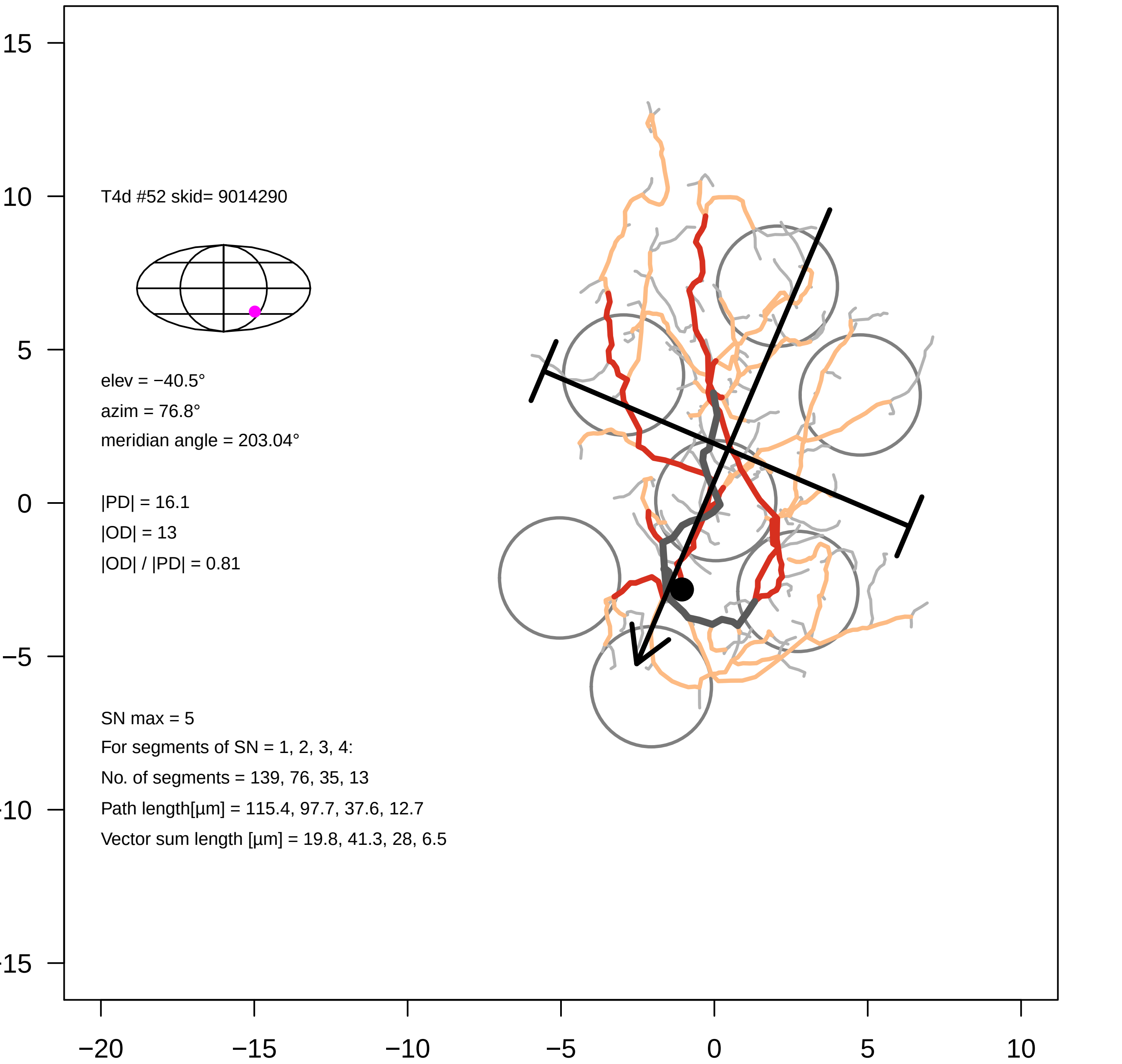
T4d #52 skid= 9014290
elev = −40.5°
azim = 76.8°
meridian angle = 203.04°
|PD| = 16.1
|OD| = 13
|OD| / |PD| = 0.81
SN max = 5
For segments of SN = 1, 2, 3, 4:
No. of segments = 139, 76, 35, 13
Path length[µm] = 115.4, 97.7, 37.6, 12.7
Vector sum length [µm] = 19.8, 41.3, 28, 6.5
15
10
5
0
−5
−10
−15
−20
−15
−10
−5
0
5
10

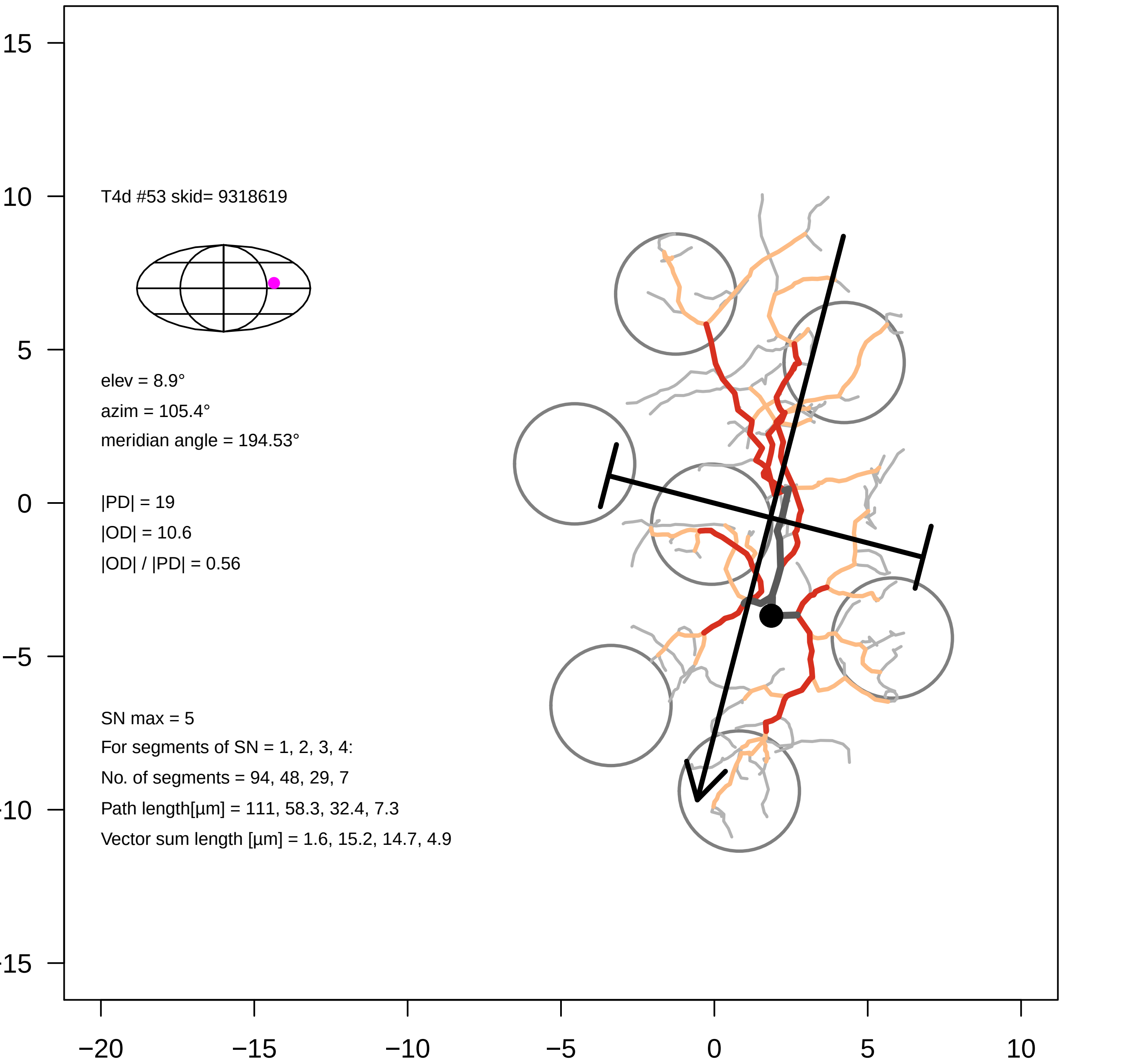

T4d #53 skid= 9318619
elev = 8.9°
azim = 105.4°
meridian angle = 194.53°
|PD| = 19
|OD| = 10.6
|OD| / |PD| = 0.56
SN max = 5
For segments of SN = 1, 2, 3, 4:
No. of segments = 94, 48, 29, 7
Path length[µm] = 111, 58.3, 32.4, 7.3
Vector sum length [µm] = 1.6, 15.2, 14.7, 4.9
15
10
5
0
−5
−10
−15
−20
−15
−10
−5
0
5
10

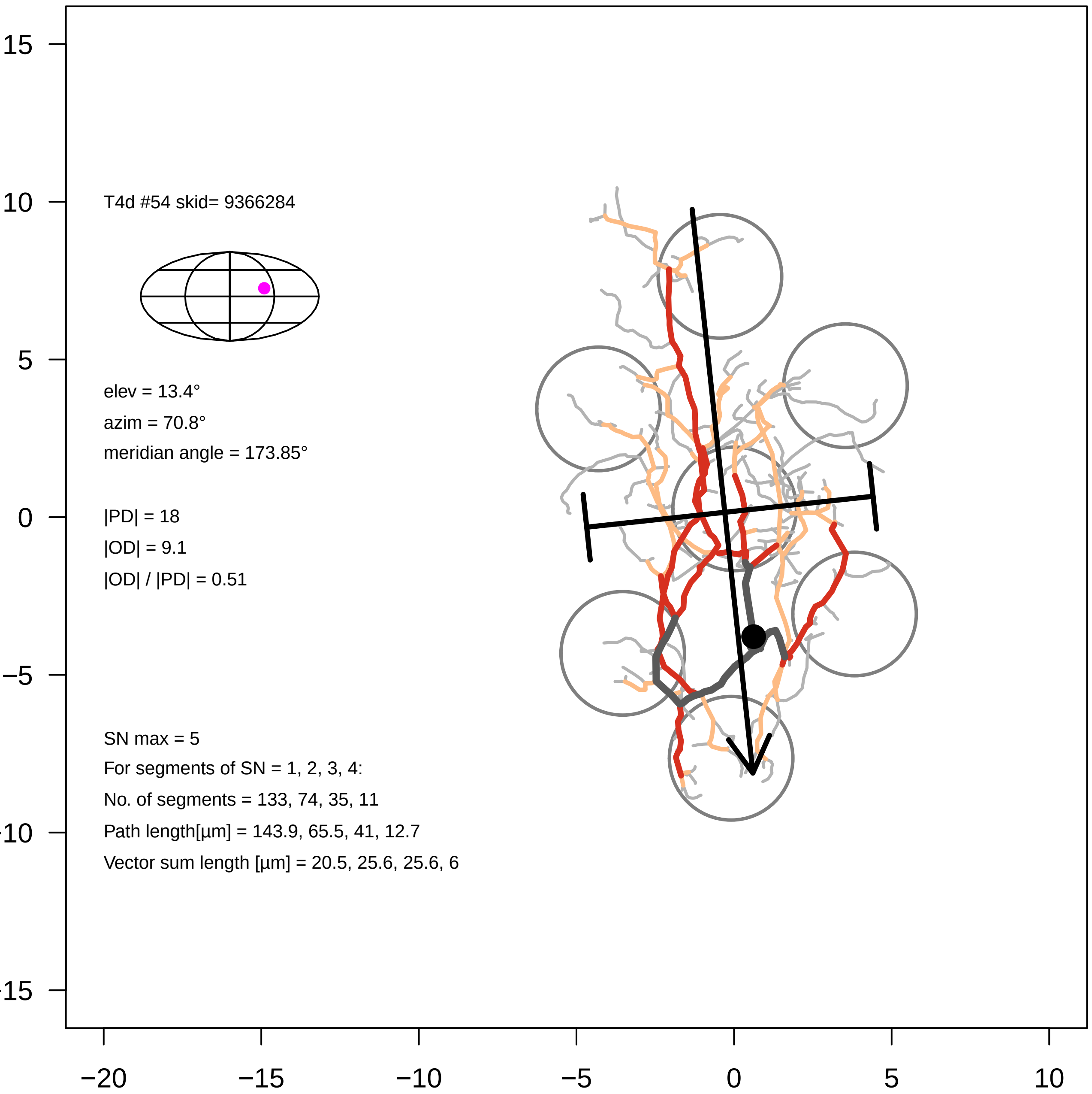

T4d #54 skid= 9366284
elev = 13.4°
azim = 70.8°
meridian angle = 173.85°
|PD| = 18
|OD| = 9.1
|OD| / |PD| = 0.51
SN max = 5
For segments of SN = 1, 2, 3, 4:
No. of segments = 133, 74, 35, 11
Path length[µm] = 143.9, 65.5, 41, 12.7
Vector sum length [µm] = 20.5, 25.6, 25.6, 6
15
10
5
0
−5
−10
−15
−20
−15
−10
−5
0
5
10

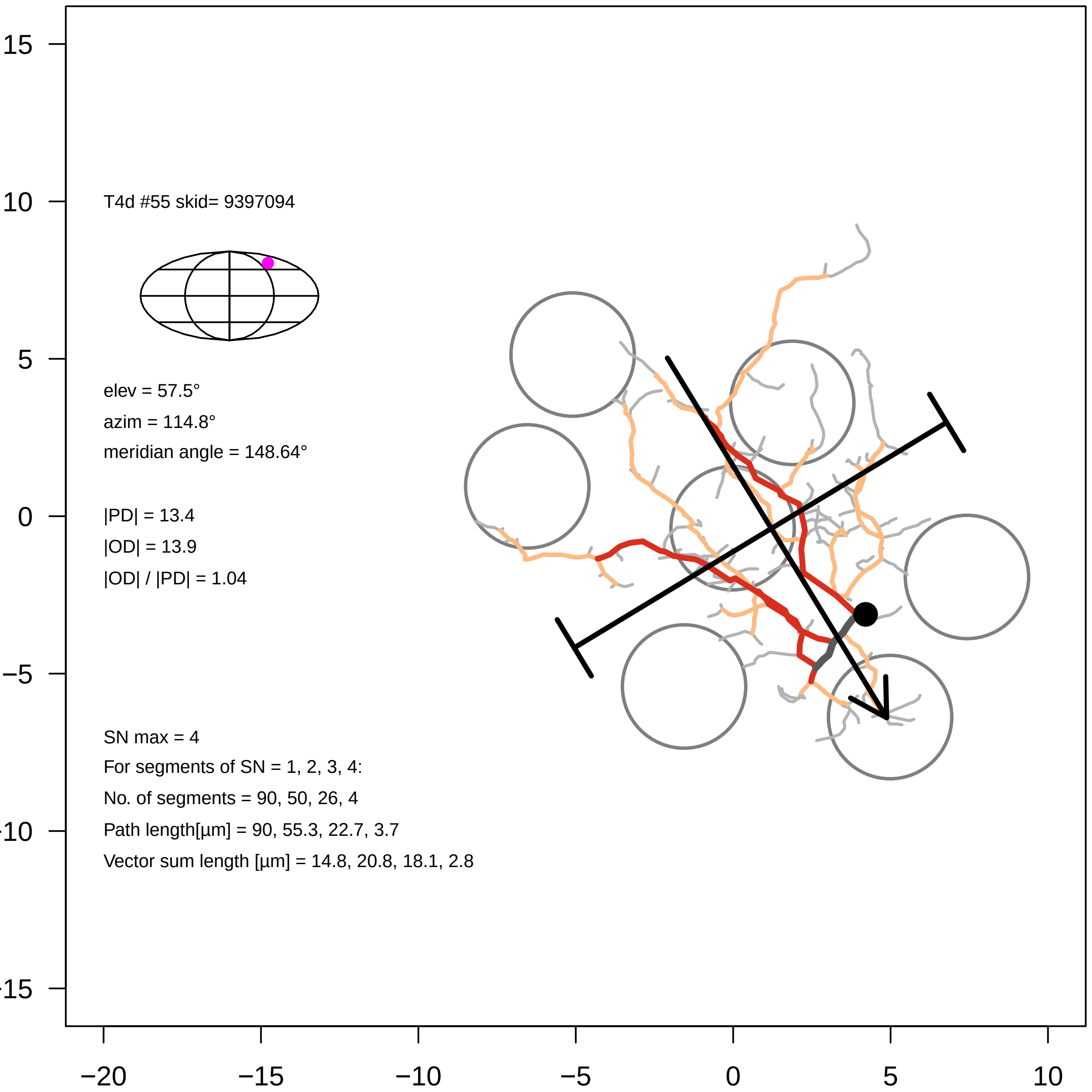
T4d #55 skid= 9397094
elev = 57.5°
azim = 114.8°
meridian angle = 148.64°
|PD| = 13.4
|OD| = 13.9
|OD| / |PD| = 1.04
SN max = 4
For segments of SN = 1, 2, 3, 4:
No. of segments = 90, 50, 26, 4
Path length[µm] = 90, 55.3, 22.7, 3.7
Vector sum length [µm] = 14.8, 20.8, 18.1, 2.8
15
10
5
0
−5
−10
−15
−20
−15
−10
−5
0
5
10

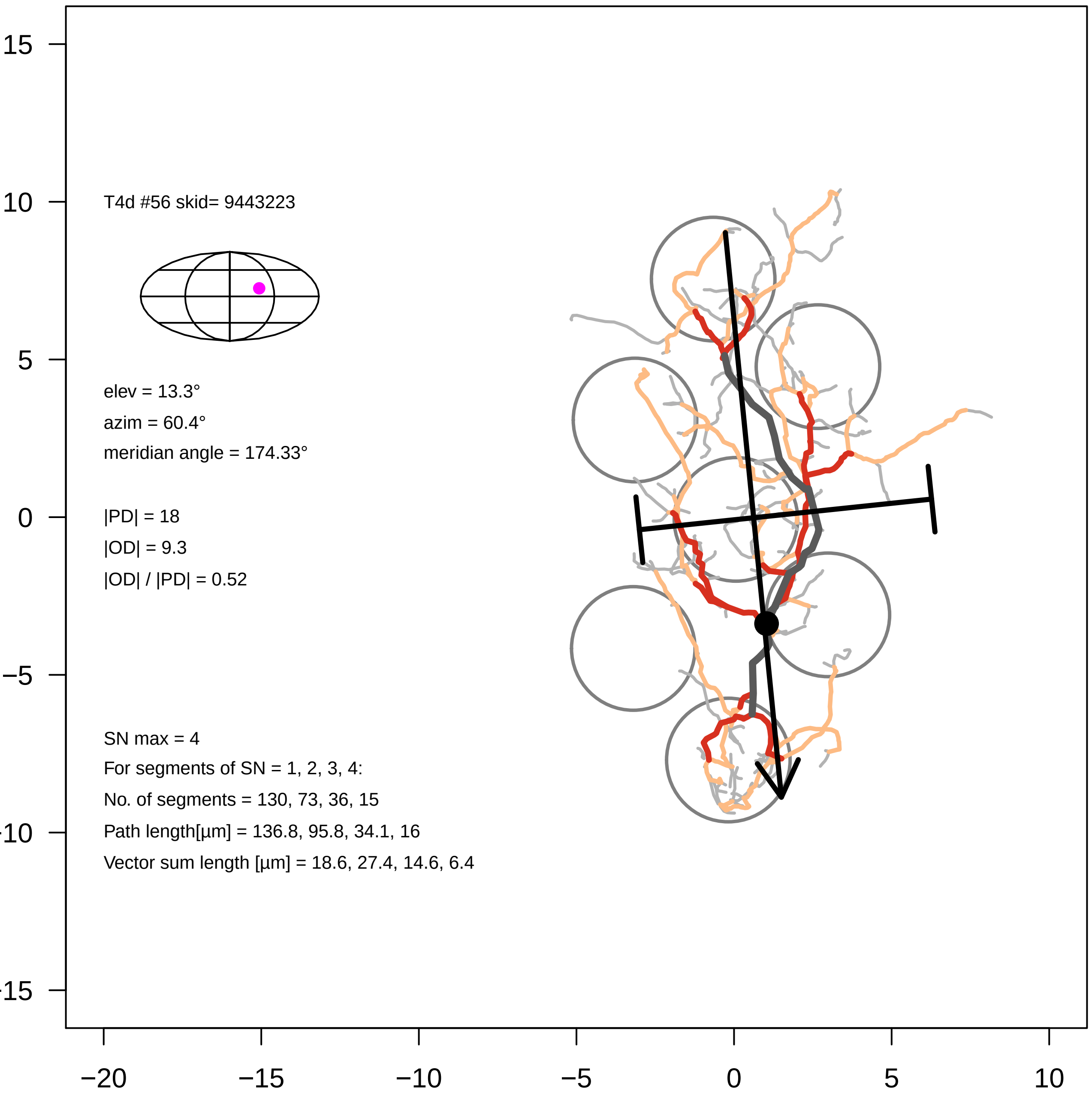

T4d #56 skid= 9443223
elev = 13.3°
azim = 60.4°
meridian angle = 174.33°
|PD| = 18
|OD| = 9.3
|OD| / |PD| = 0.52
SN max = 4
For segments of SN = 1, 2, 3, 4:
No. of segments = 130, 73, 36, 15
Path length[µm] = 136.8, 95.8, 34.1, 16
Vector sum length [µm] = 18.6, 27.4, 14.6, 6.4
15
10
5
0
−5
−10
−15
−20
−15
−10
−5
0
5
10

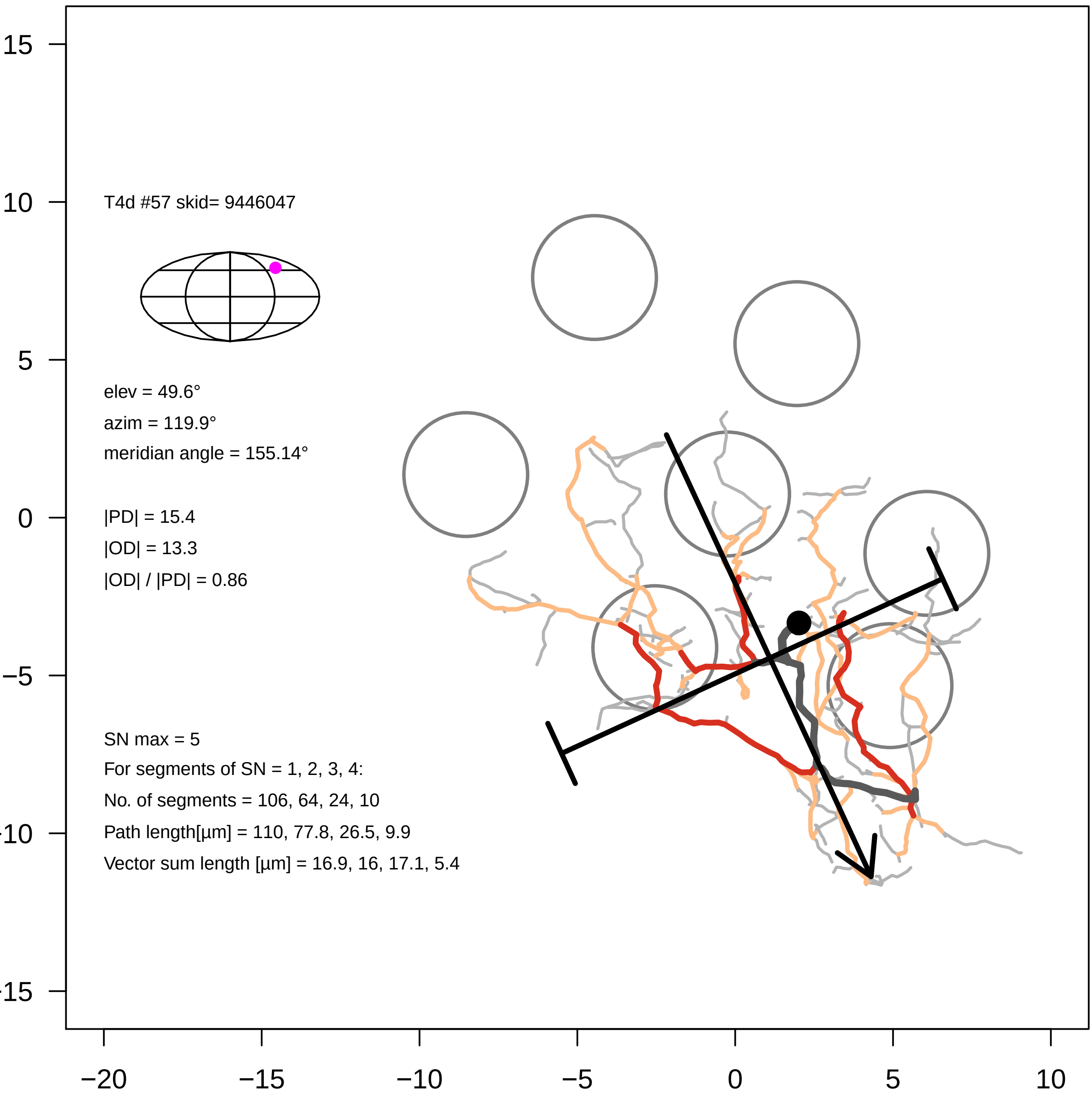

T4d #57 skid= 9446047
elev = 49.6°
azim = 119.9°
meridian angle = 155.14°
|PD| = 15.4
|OD| = 13.3
|OD| / |PD| = 0.86
SN max = 5
For segments of SN = 1, 2, 3, 4:
No. of segments = 106, 64, 24, 10
Path length[µm] = 110, 77.8, 26.5, 9.9
Vector sum length [µm] = 16.9, 16, 17.1, 5.4
15
10
5
0
−5
−10
−15
−20
−15
−10
−5
0
5
10

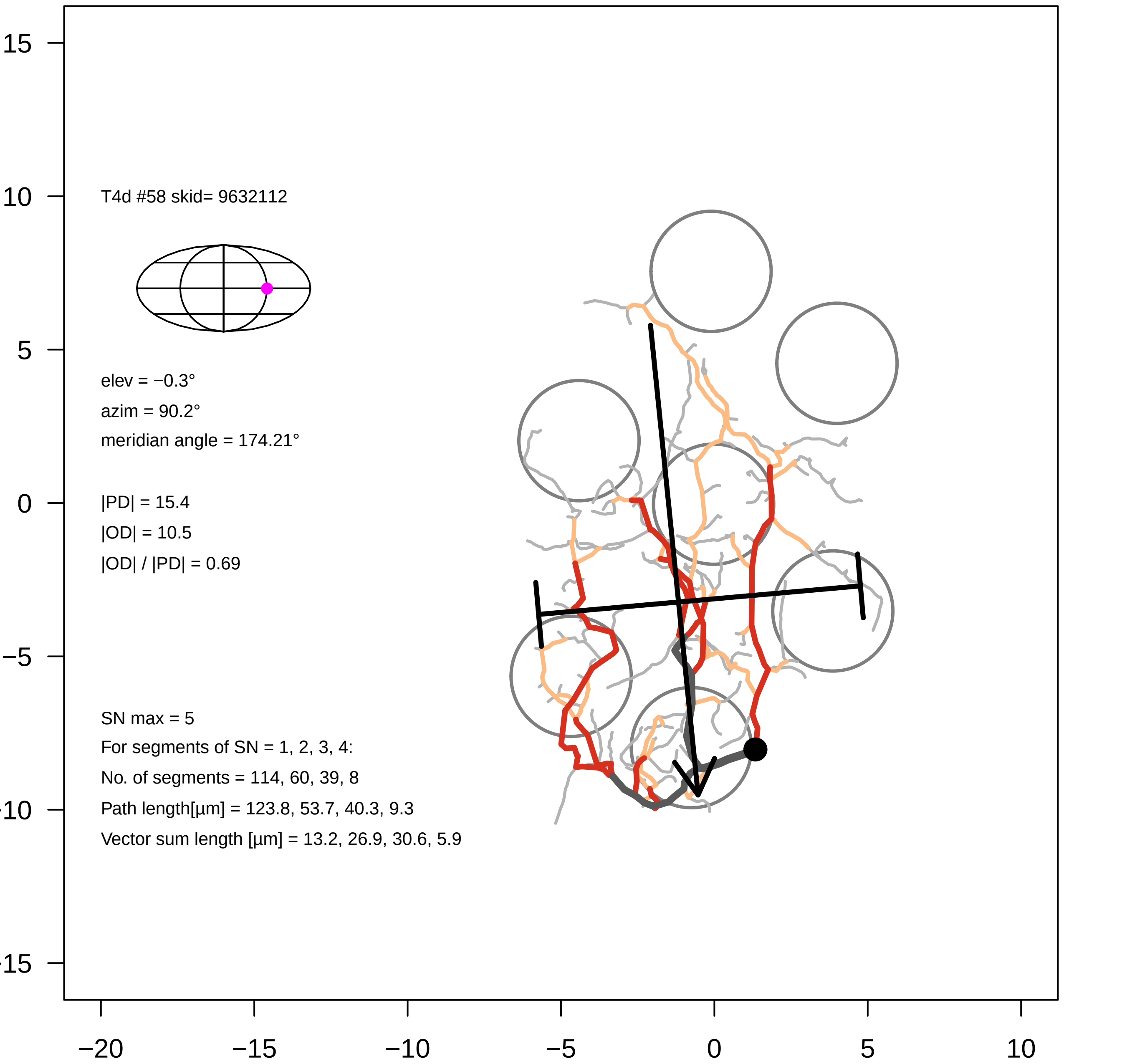
T4d #58 skid= 9632112
elev = −0.3°
azim = 90.2°
meridian angle = 174.21°
|PD| = 15.4
|OD| = 10.5
|OD| / |PD| = 0.69
SN max = 5
For segments of SN = 1, 2, 3, 4:
No. of segments = 114, 60, 39, 8
Path length[µm] = 123.8, 53.7, 40.3, 9.3
Vector sum length [µm] = 13.2, 26.9, 30.6, 5.9
15
10
5
0
−5
−10
−15
−20
−15
−10
−5
0
5
10

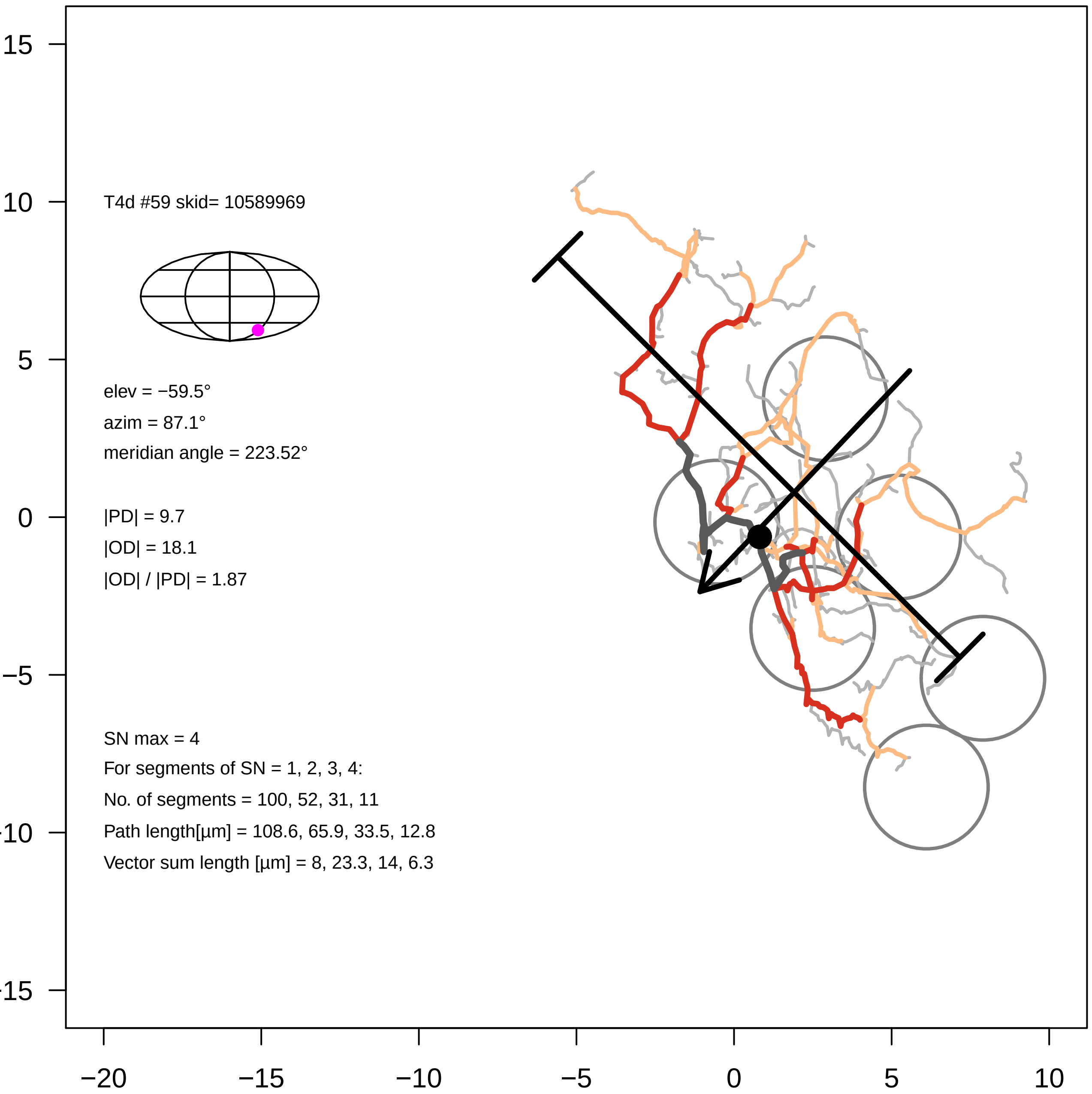

T4d #59 skid= 10589969
elev = −59.5°
azim = 87.1°
meridian angle = 223.52°
|PD| = 9.7
|OD| = 18.1
|OD| / |PD| = 1.87
SN max = 4
For segments of SN = 1, 2, 3, 4:
No. of segments = 100, 52, 31, 11
Path length[µm] = 108.6, 65.9, 33.5, 12.8
Vector sum length [µm] = 8, 23.3, 14, 6.3
15
10
5
0
−5
-10
-15
−20
−15
−10
−5
0
5
10

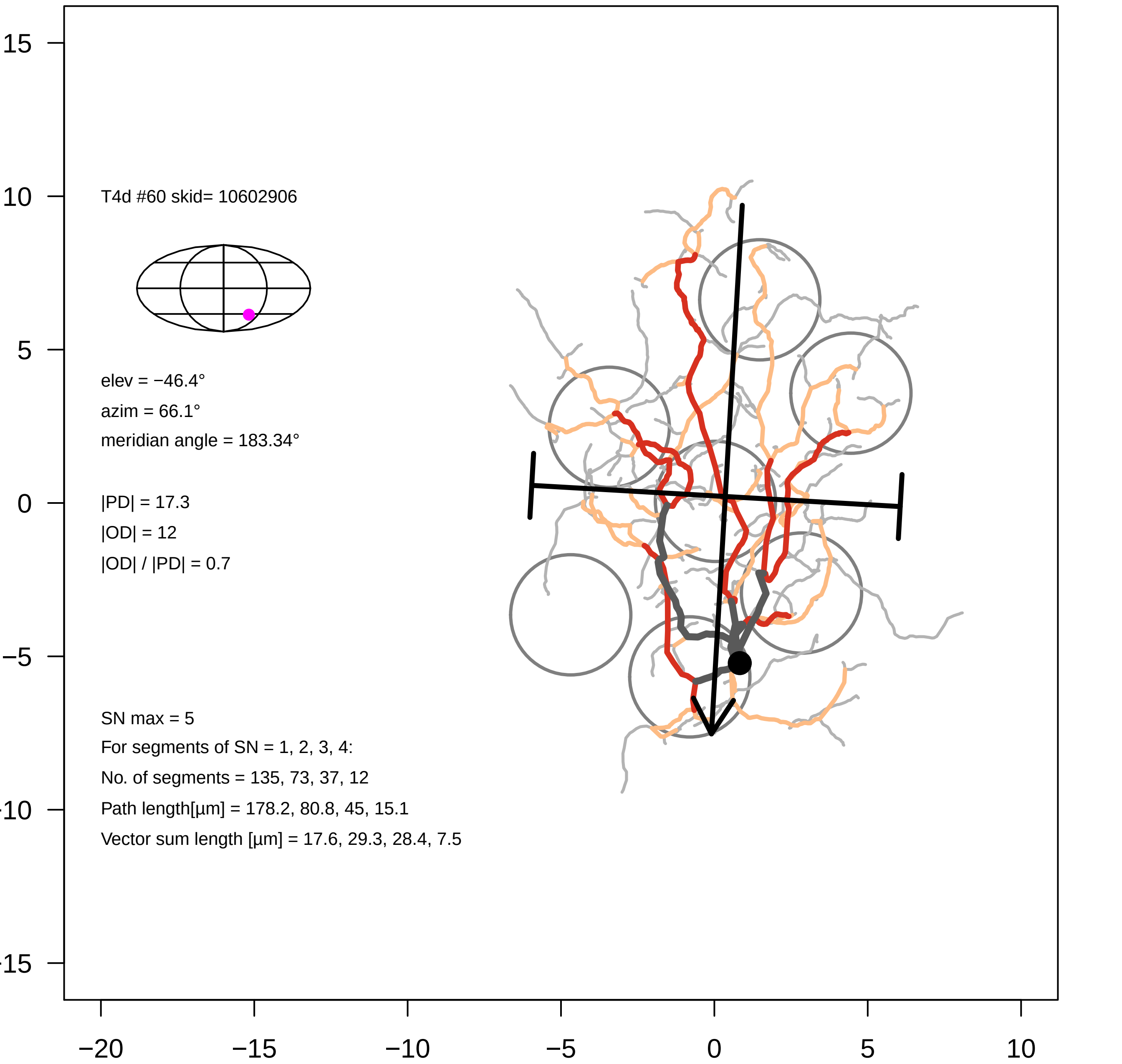

T4d #60 skid= 10602906
elev = −46.4°
azim = 66.1°
meridian angle = 183.34°
|PD| = 17.3
|OD| = 12
|OD| / |PD| = 0.7
SN max = 5
For segments of SN = 1, 2, 3, 4:
No. of segments = 135, 73, 37, 12
Path length[µm] = 178.2, 80.8, 45, 15.1
Vector sum length [µm] = 17.6, 29.3, 28.4, 7.5
15
10
5
0
−5
−10
−15
−20
−15
−10
−5
0
5
10

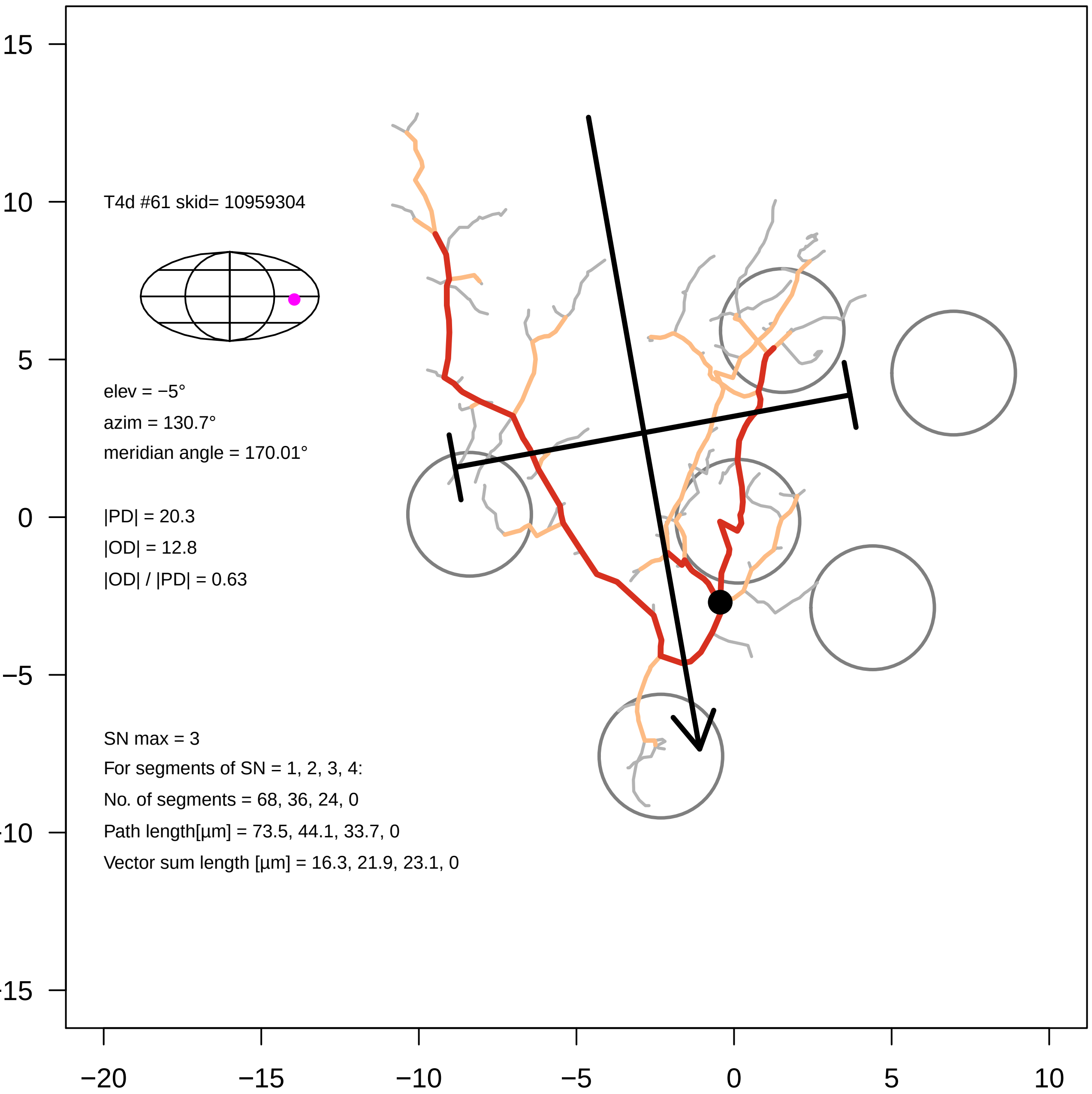

T4d #61 skid= 10959304
elev = −5°
azim = 130.7°
meridian angle = 170.01°
|PD| = 20.3
|OD| = 12.8
|OD| / |PD| = 0.63
SN max = 3
For segments of SN = 1, 2, 3, 4:
No. of segments = 68, 36, 24, 0
Path length[µm] = 73.5, 44.1, 33.7, 0
Vector sum length [µm] = 16.3, 21.9, 23.1, 0
15
10
5
0
−5
−10
−15
−20
−15
−10
−5
0
5
10

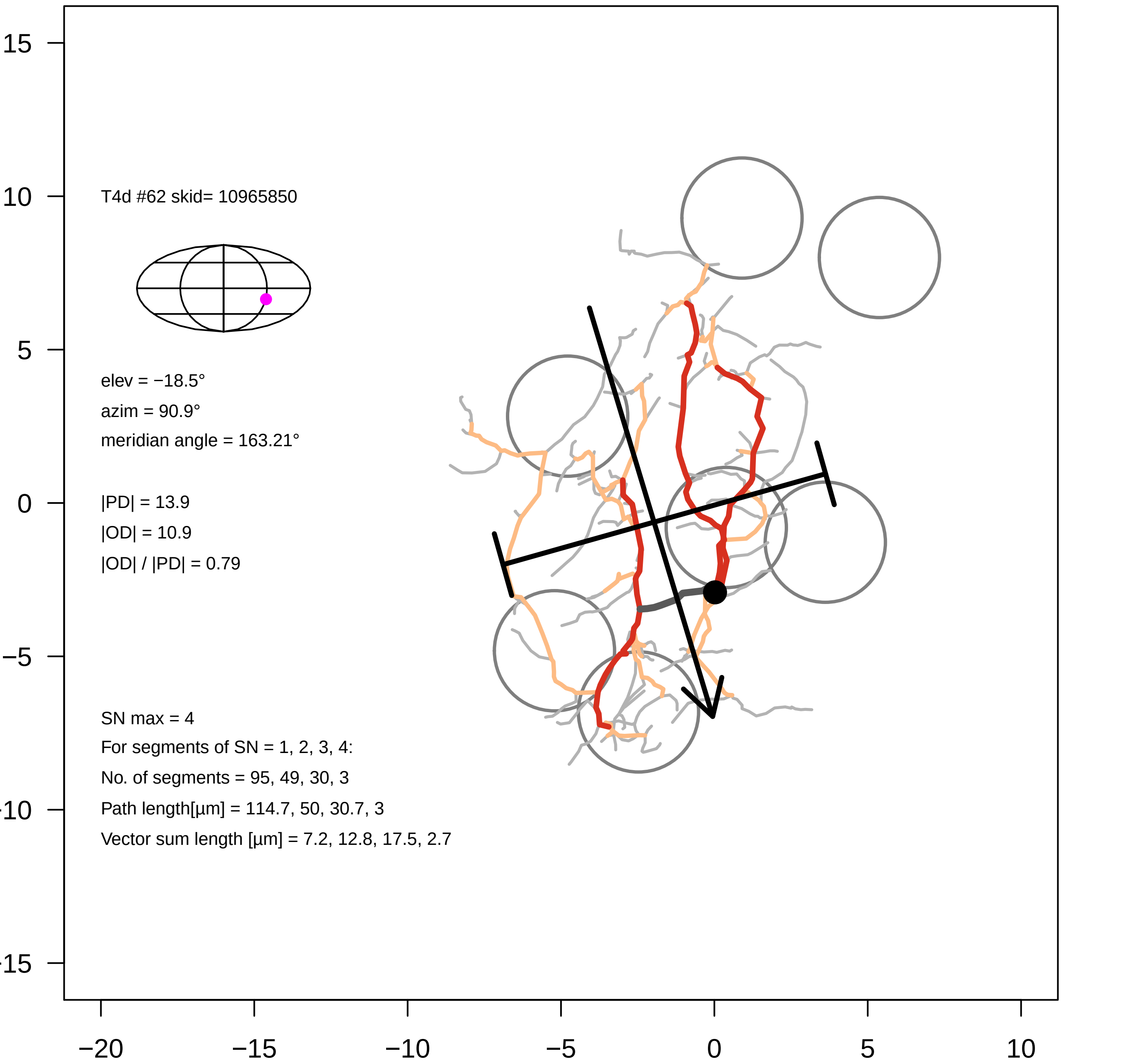
T4d #62 skid= 10965850
elev = −18.5°
azim = 90.9°
meridian angle = 163.21°
|PD| = 13.9
|OD| = 10.9
|OD| / |PD| = 0.79
SN max = 4
For segments of SN = 1, 2, 3, 4:
No. of segments = 95, 49, 30, 3
Path length[µm] = 114.7, 50, 30.7, 3
Vector sum length [µm] = 7.2, 12.8, 17.5, 2.7
15
10
5
0
−5
−10
−15
−20
−15
−10
−5
0
5
10

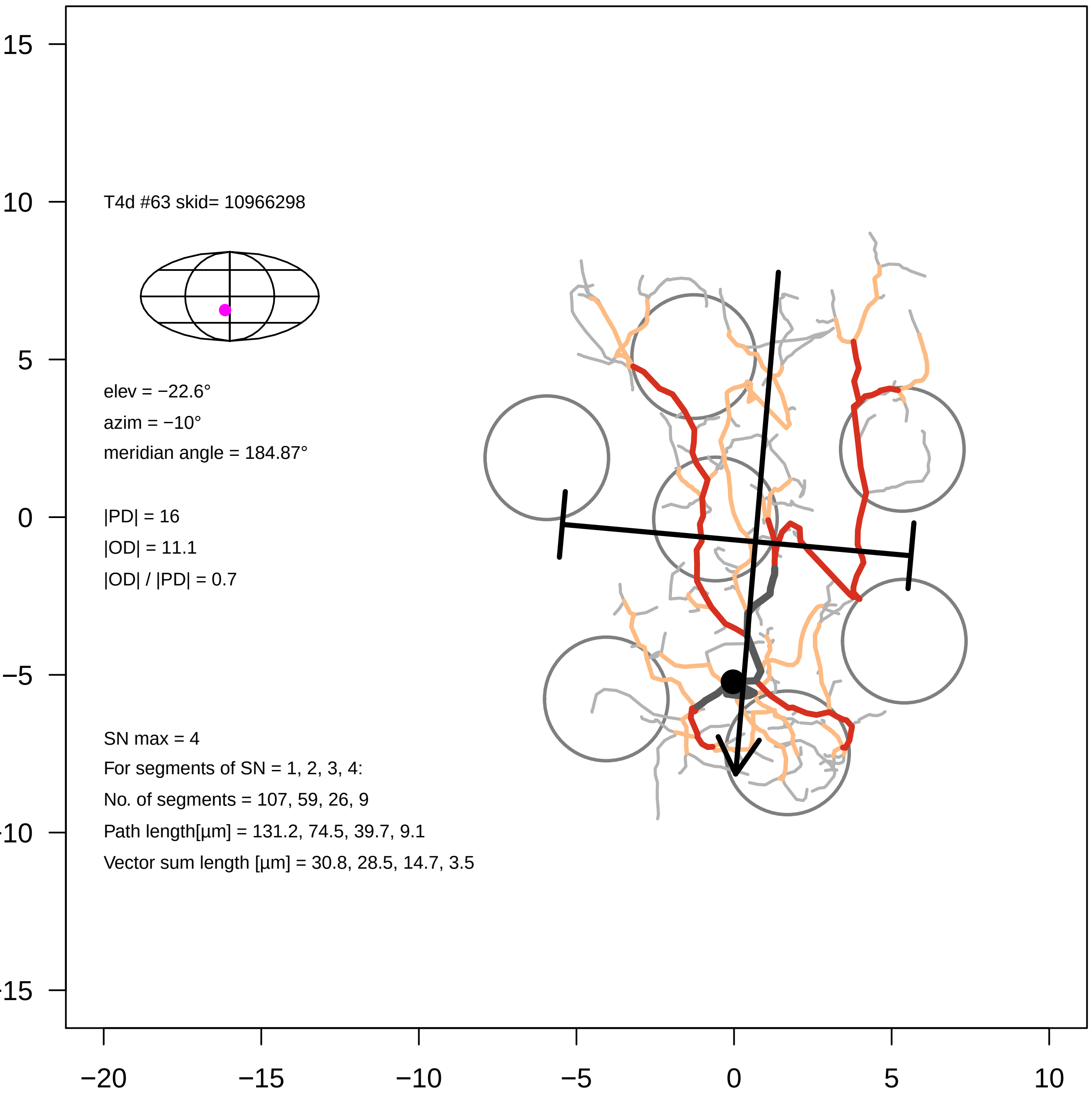
T4d #63 skid= 10966298
elev = −22.6°
azim = −10°
meridian angle = 184.87°
|PD| = 16
|OD| = 11.1
|OD| / |PD| = 0.7
SN max = 4
For segments of SN = 1, 2, 3, 4:
No. of segments = 107, 59, 26, 9
Path length[µm] = 131.2, 74.5, 39.7, 9.1
Vector sum length [µm] = 30.8, 28.5, 14.7, 3.5
15
10
5
0
−5
−10
−15
−20
−15
−10
−5
0
5
10

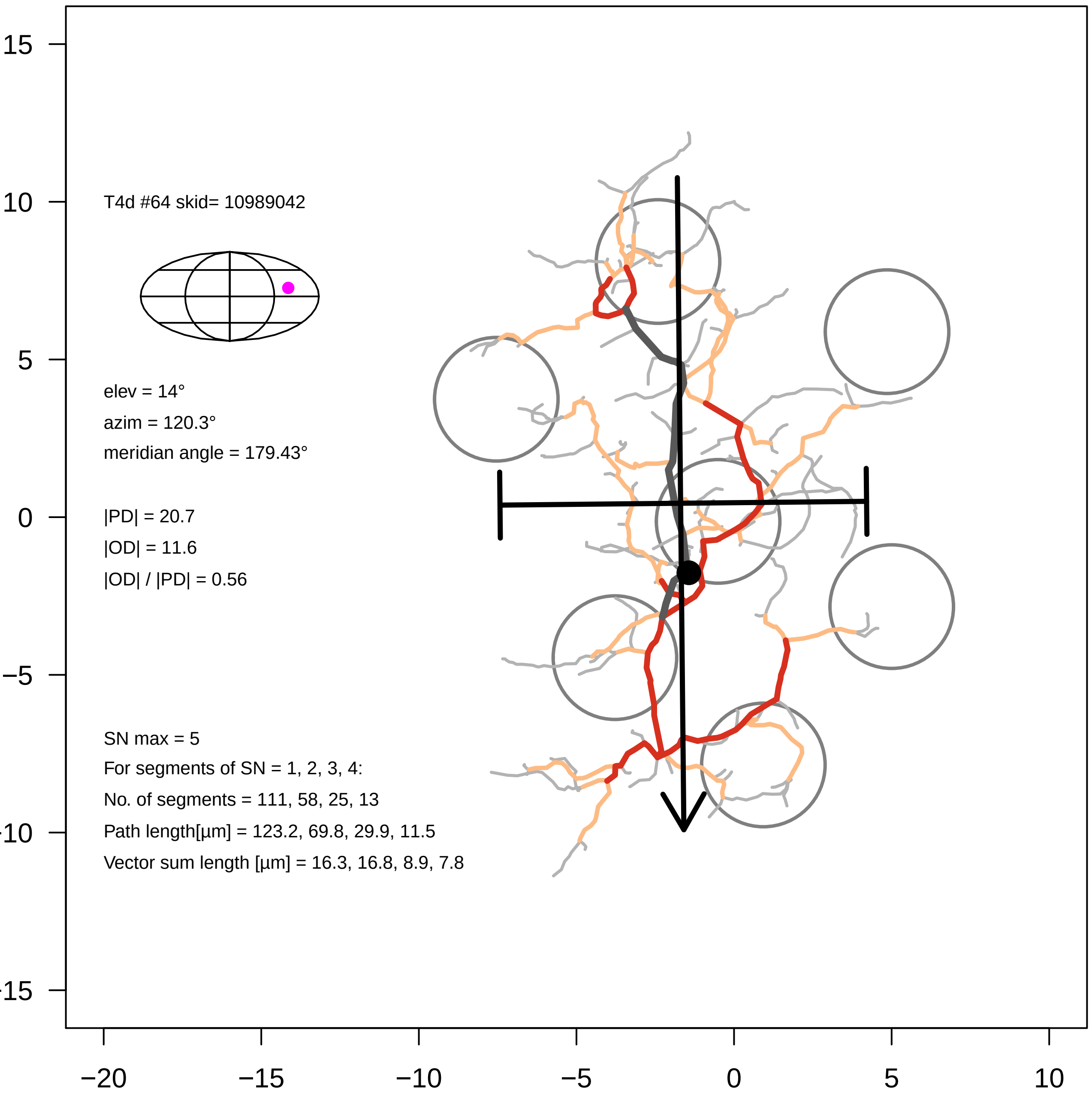

T4d #64 skid= 10989042
elev = 14°
azim = 120.3°
meridian angle = 179.43°
|PD| = 20.7
|OD| = 11.6
|OD| / |PD| = 0.56
SN max = 5
For segments of SN = 1, 2, 3, 4:
No. of segments = 111, 58, 25, 13
Path length[µm] = 123.2, 69.8, 29.9, 11.5
Vector sum length [µm] = 16.3, 16.8, 8.9, 7.8
15
10
5
0
−5
−10
−15
−20
−15
−10
−5
0
5
10

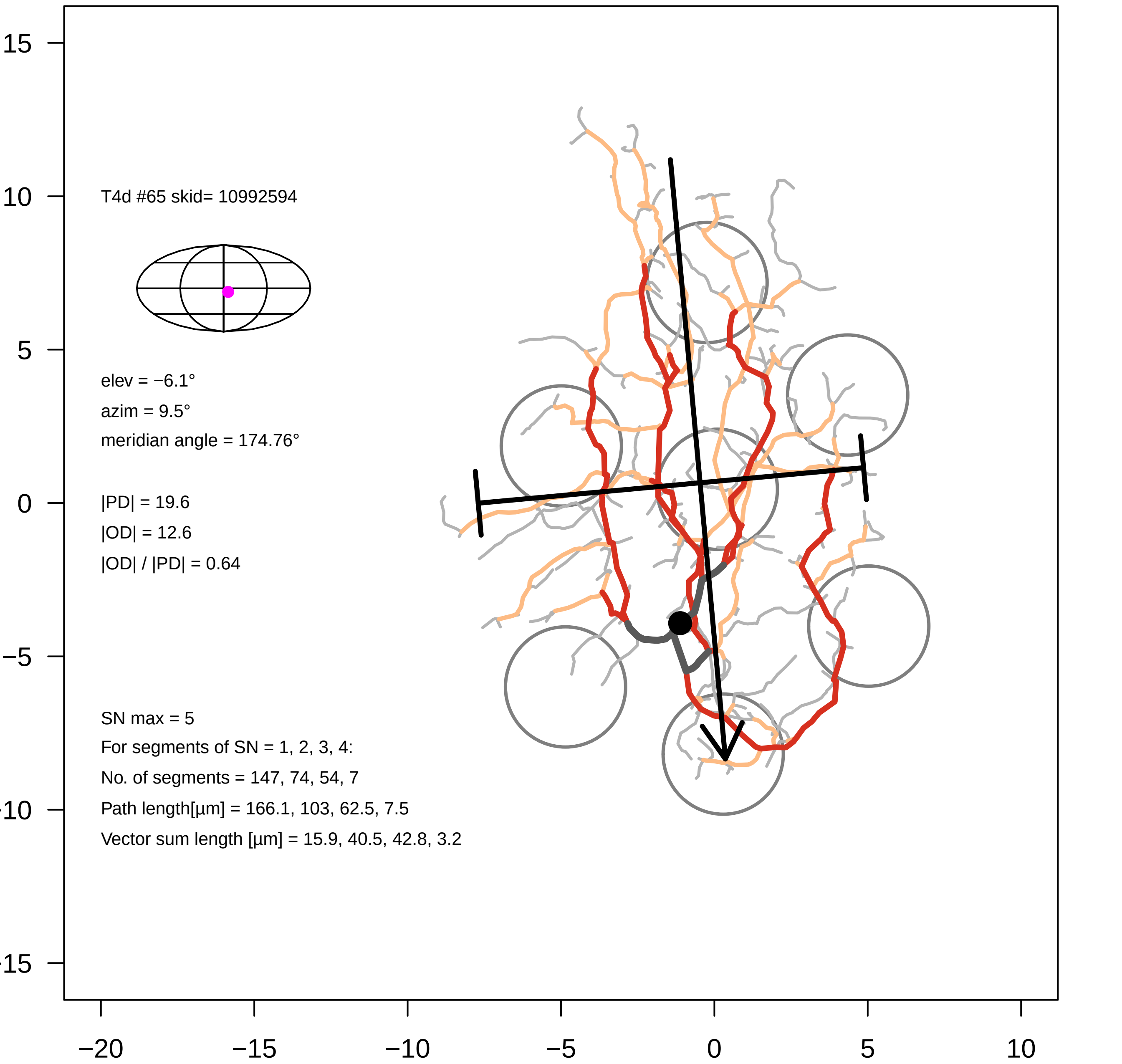

T4d #65 skid= 10992594
elev = −6.1°
azim = 9.5°
meridian angle = 174.76°
|PD| = 19.6
|OD| = 12.6
|OD| / |PD| = 0.64
SN max = 5
For segments of SN = 1, 2, 3, 4:
No. of segments = 147, 74, 54, 7
Path length[µm] = 166.1, 103, 62.5, 7.5
Vector sum length [µm] = 15.9, 40.5, 42.8, 3.2
15
10
5
0
−5
−10
−15
−20
−15
−10
−5
0
5
10

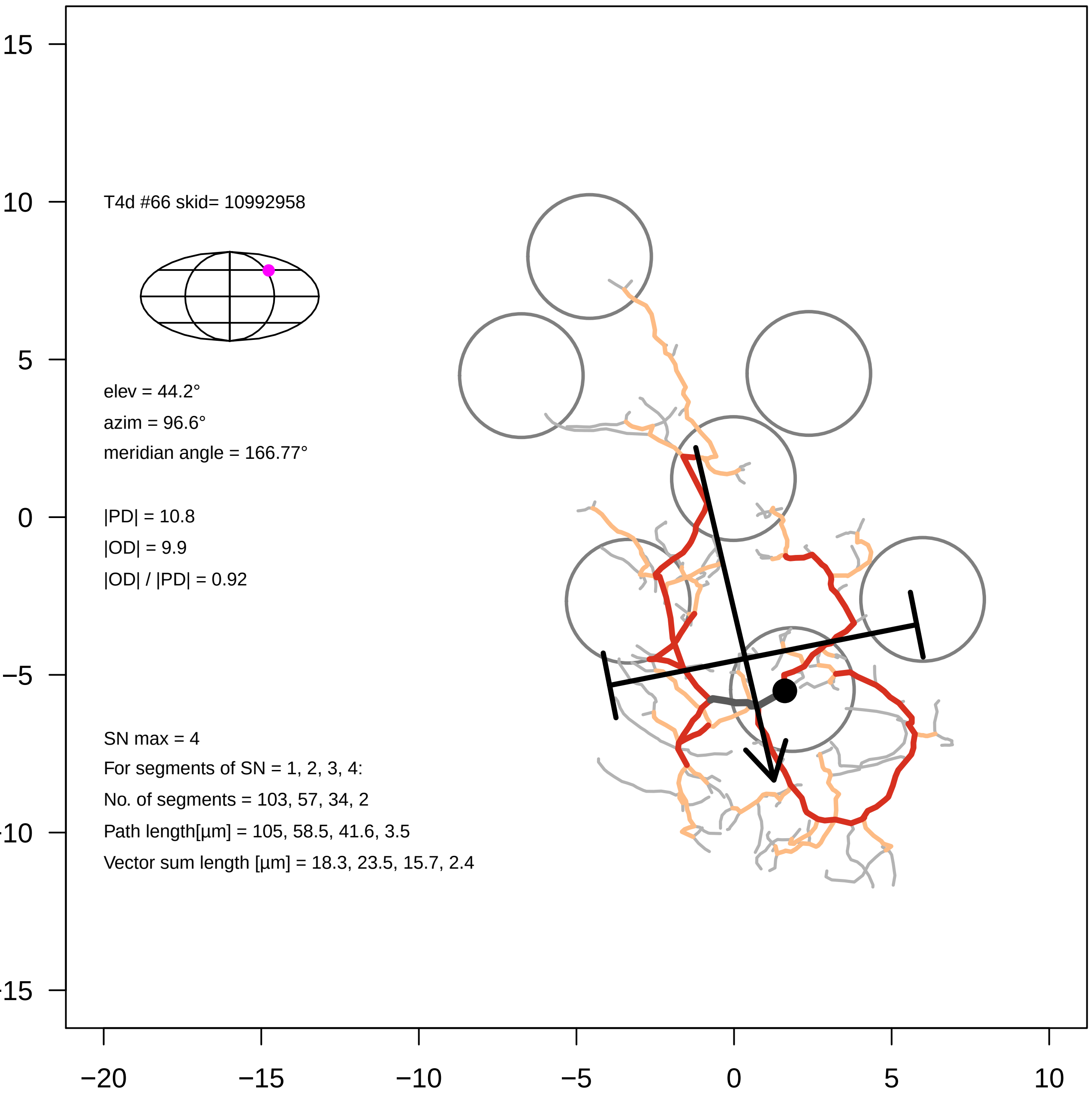

T4d #66 skid= 10992958
elev = 44.2°
azim = 96.6°
meridian angle = 166.77°
|PD| = 10.8
|OD| = 9.9
|OD| / |PD| = 0.92
SN max = 4
For segments of SN = 1, 2, 3, 4:
No. of segments = 103, 57, 34, 2
Path length[µm] = 105, 58.5, 41.6, 3.5
Vector sum length [µm] = 18.3, 23.5, 15.7, 2.4

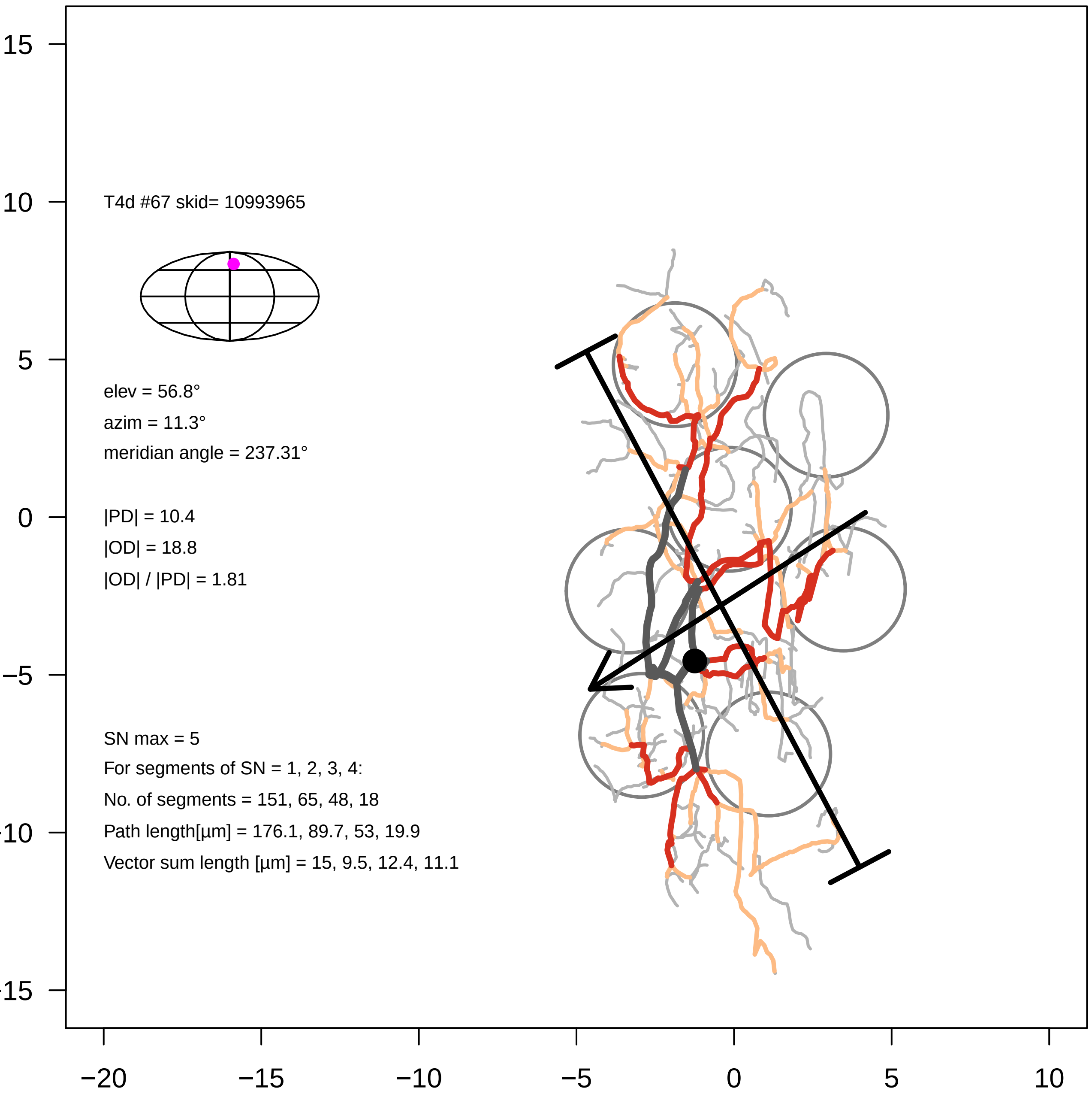
T4d #67 skid= 10993965
elev = 56.8°
azim = 11.3°
meridian angle = 237.31°
|PD| = 10.4
|OD| = 18.8
|OD| / |PD| = 1.81
SN max = 5
For segments of SN = 1, 2, 3, 4:
No. of segments = 151, 65, 48, 18
Path length[µm] = 176.1, 89.7, 53, 19.9
Vector sum length [µm] = 15, 9.5, 12.4, 11.1
15
10
5
0
−5
−10
−15
−20
−15
−10
−5
0
5
10

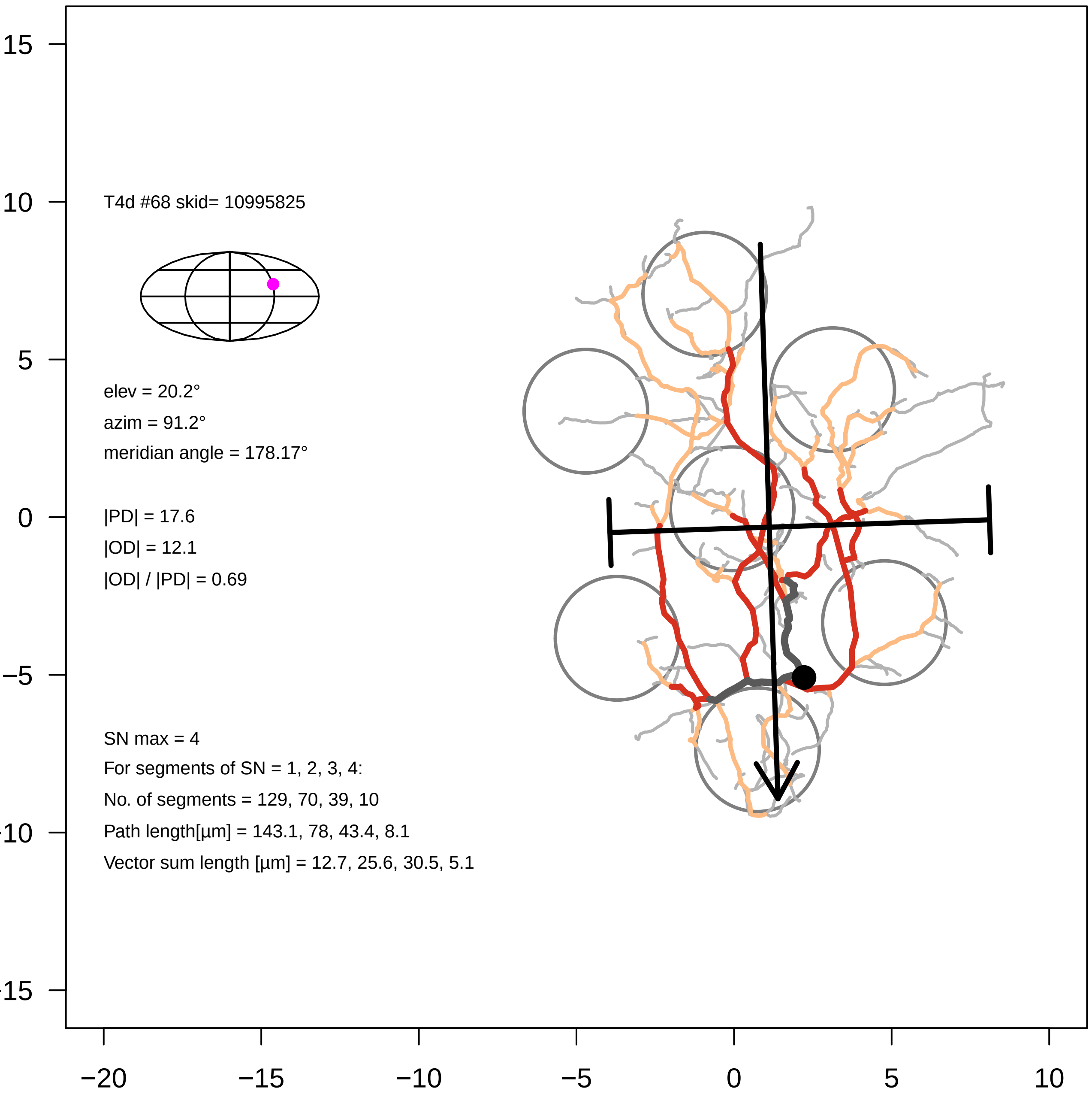

T4d #68 skid= 10995825
elev = 20.2°
azim = 91.2°
meridian angle = 178.17°
|PD| = 17.6
|OD| = 12.1
|OD| / |PD| = 0.69
SN max = 4
For segments of SN = 1, 2, 3, 4:
No. of segments = 129, 70, 39, 10
Path length[µm] = 143.1, 78, 43.4, 8.1
Vector sum length [µm] = 12.7, 25.6, 30.5, 5.1
15
10
5
0
−5
-10
-15
−20
−15
−10
−5
0
5
10

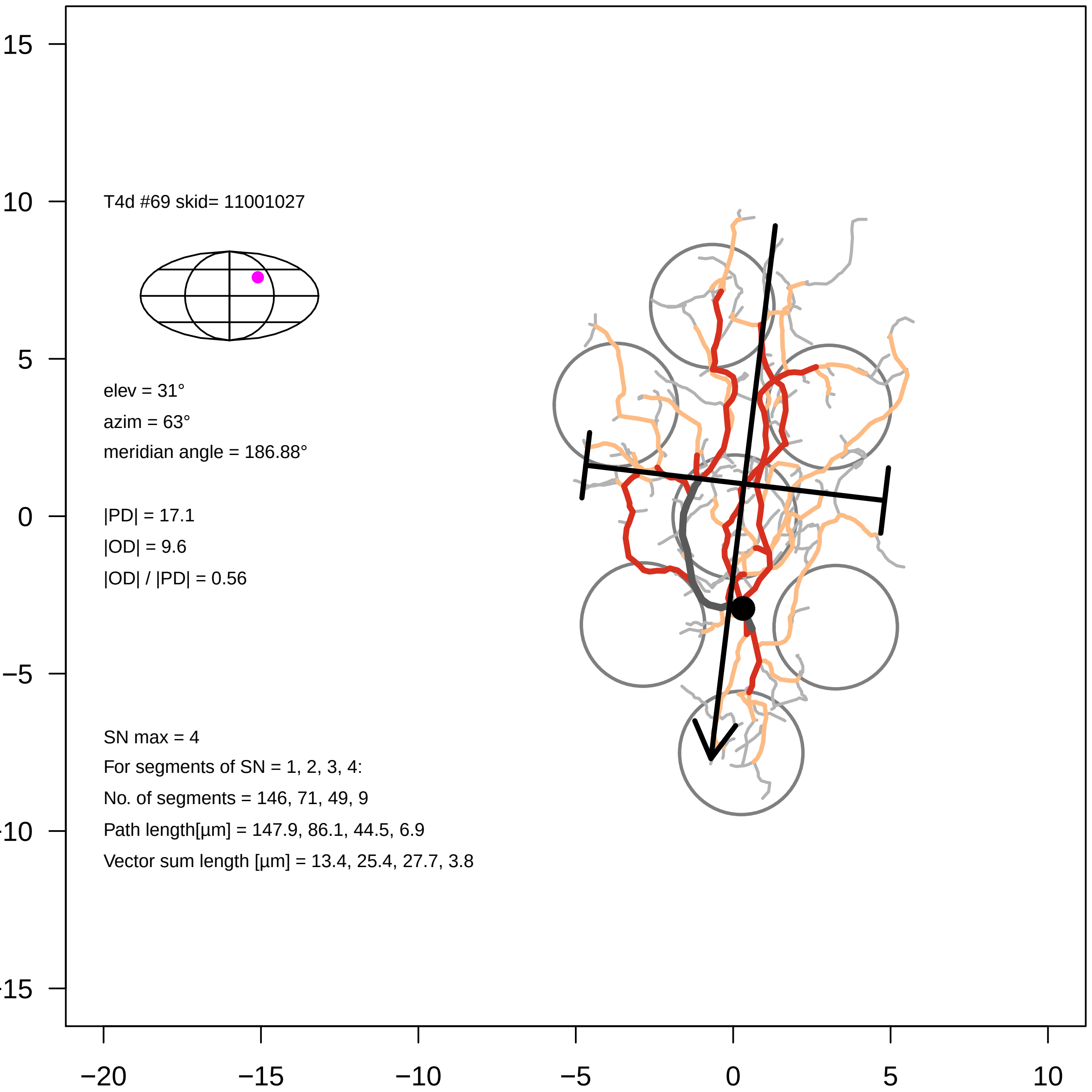
T4d #69 skid= 11001027
elev = 31°
azim = 63°
meridian angle = 186.88°
|PD| = 17.1
|OD| = 9.6
|OD| / |PD| = 0.56
SN max = 4
For segments of SN = 1, 2, 3, 4:
No. of segments = 146, 71, 49, 9
Path length[µm] = 147.9, 86.1, 44.5, 6.9
Vector sum length [µm] = 13.4, 25.4, 27.7, 3.8
15
10
5
0
−5
−10
−15
−20
−15
−10
−5
0
5
10

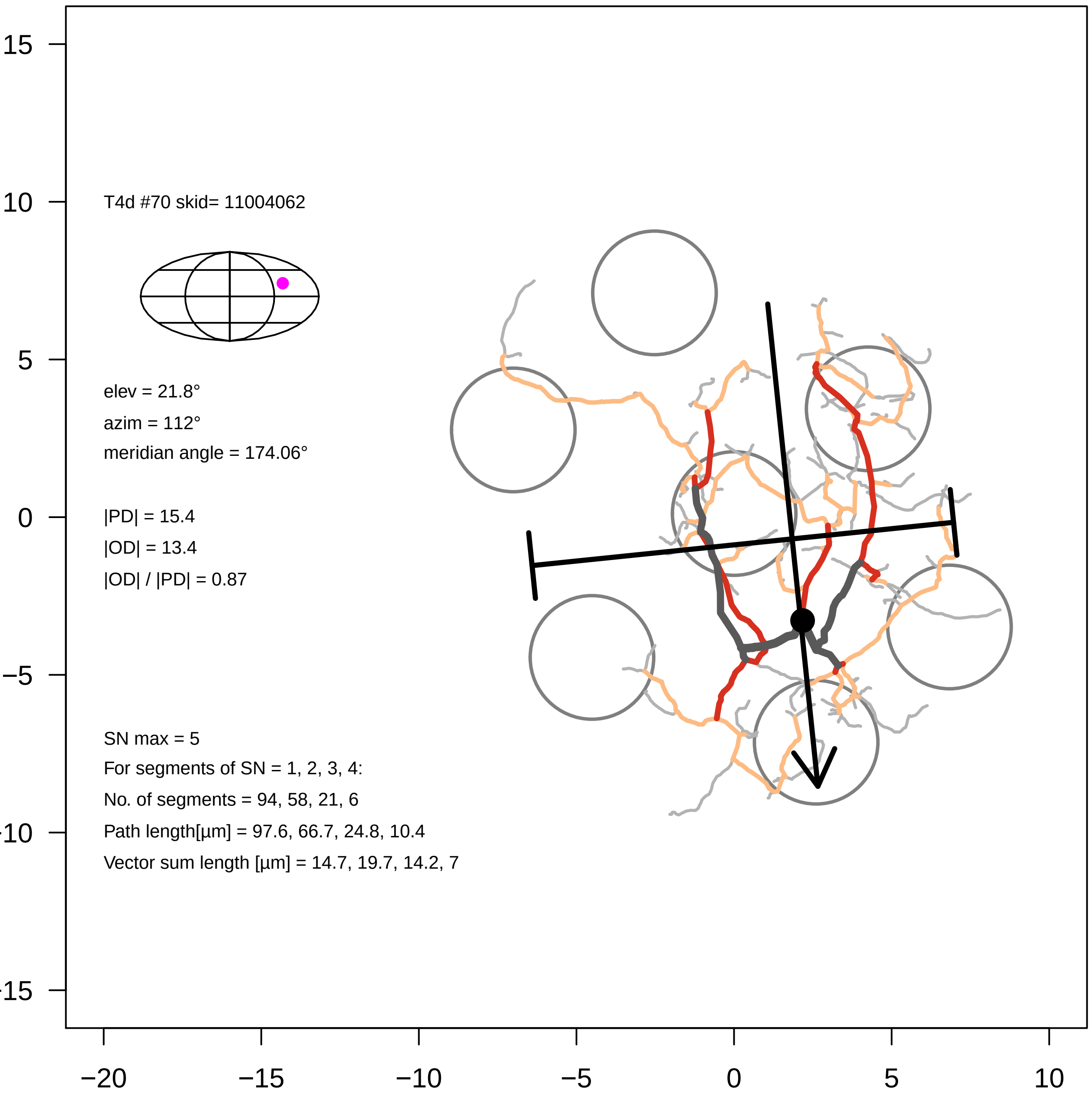

T4d #70 skid= 11004062
elev = 21.8°
azim = 112°
meridian angle = 174.06°
|PD| = 15.4
|OD| = 13.4
|OD| / |PD| = 0.87
SN max = 5
For segments of SN = 1, 2, 3, 4:
No. of segments = 94, 58, 21, 6
Path length[µm] = 97.6, 66.7, 24.8, 10.4
Vector sum length [µm] = 14.7, 19.7, 14.2, 7
15
10
5
0
−5
−10
−15
−20
−15
−10
−5
0
5
10

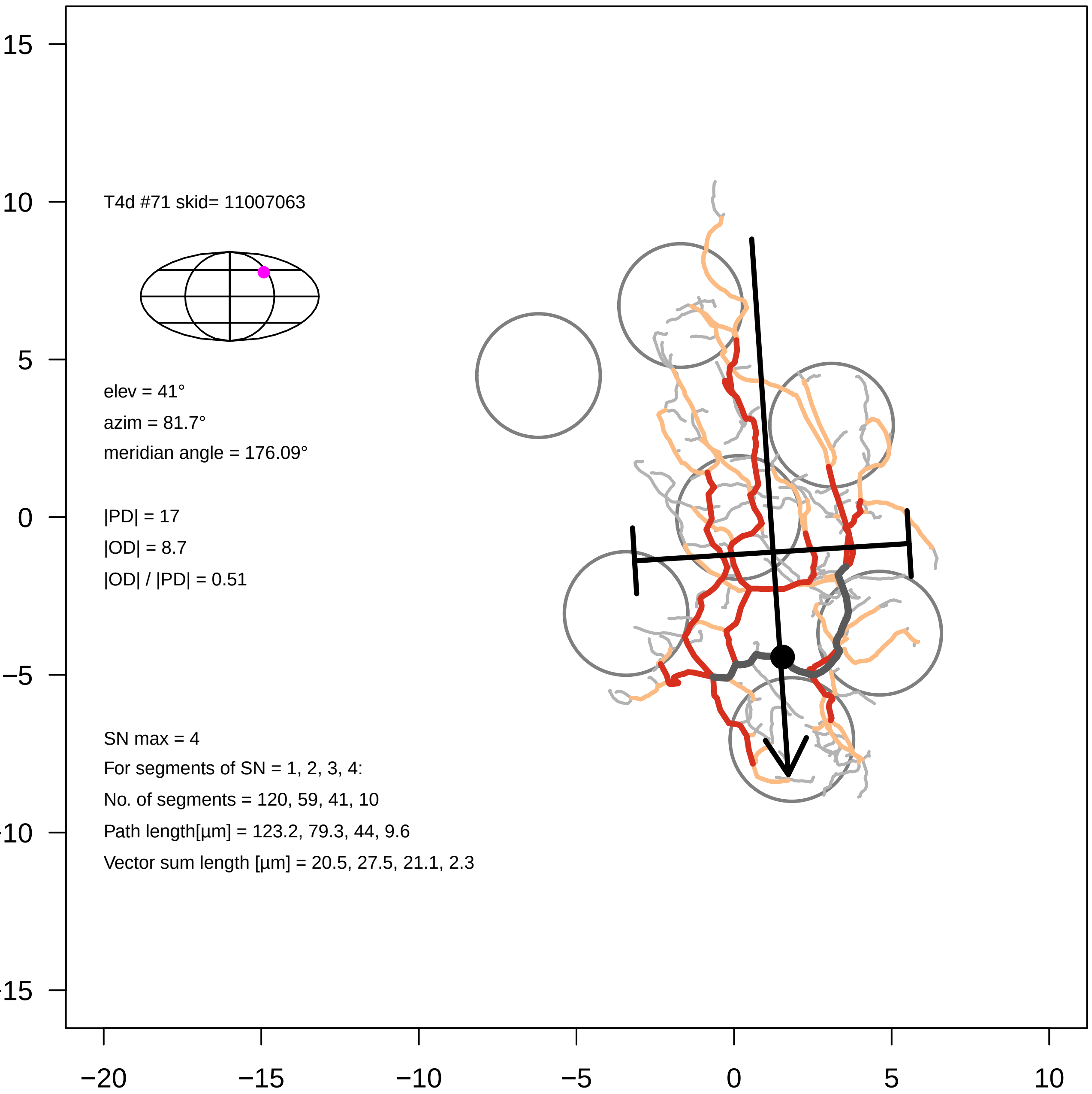

T4d #71 skid= 11007063
elev = 41°
azim = 81.7°
meridian angle = 176.09°
|PD| = 17
|OD| = 8.7
|OD| / |PD| = 0.51
SN max = 4
For segments of SN = 1, 2, 3, 4:
No. of segments = 120, 59, 41, 10
Path length[µm] = 123.2, 79.3, 44, 9.6
Vector sum length [µm] = 20.5, 27.5, 21.1, 2.3
15
10
5
0
−5
-10
-15
−20
−15
−10
−5
0
5
10

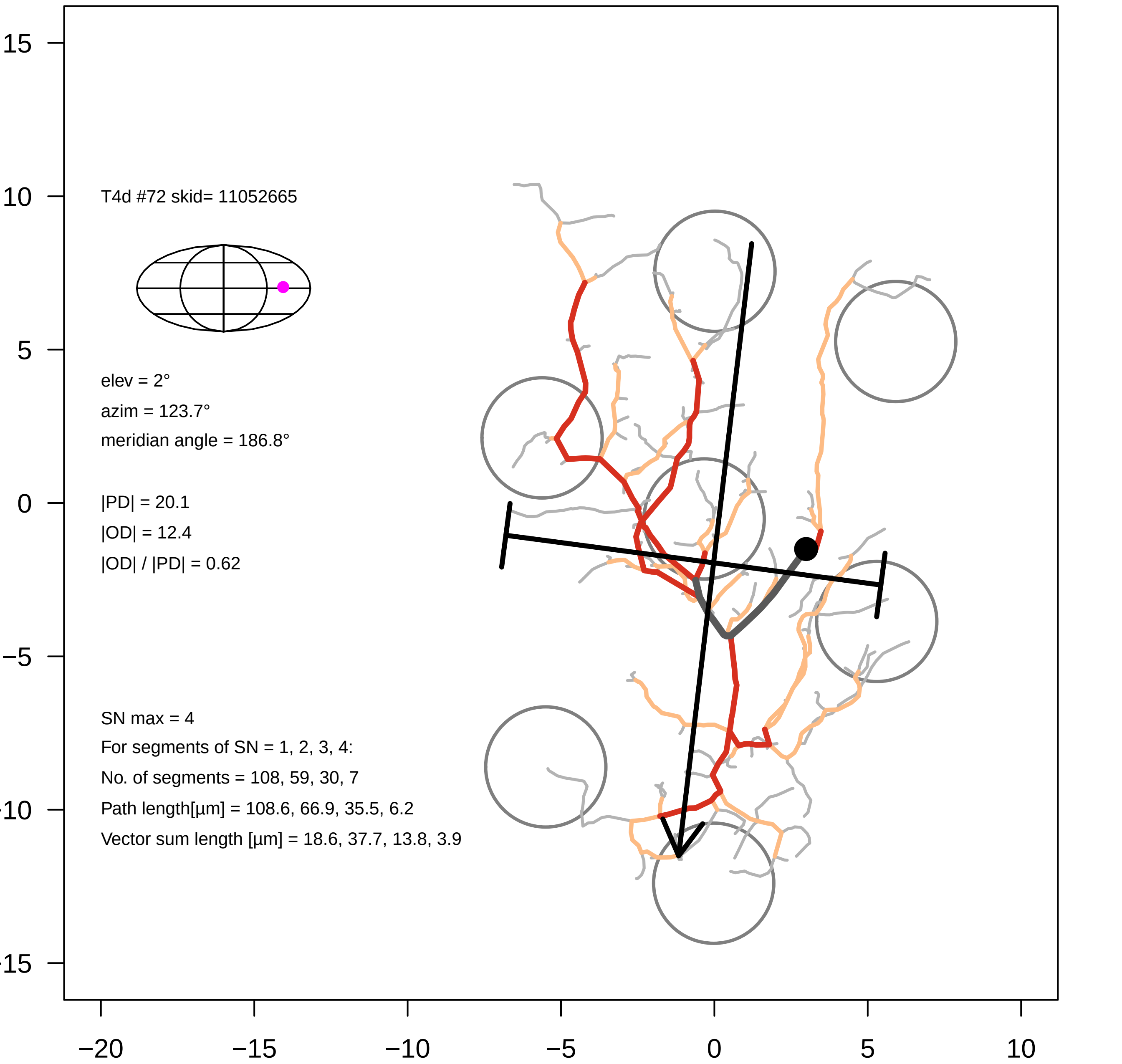
T4d #72 skid= 11052665
elev = 2°
azim = 123.7°
meridian angle = 186.8°
|PD| = 20.1
|OD| = 12.4
|OD| / |PD| = 0.62
SN max = 4
For segments of SN = 1, 2, 3, 4:
No. of segments = 108, 59, 30, 7
Path length[µm] = 108.6, 66.9, 35.5, 6.2
Vector sum length [µm] = 18.6, 37.7, 13.8, 3.9
15
10
5
0
−5
−10
−15
−20
−15
−10
−5
0
5
10

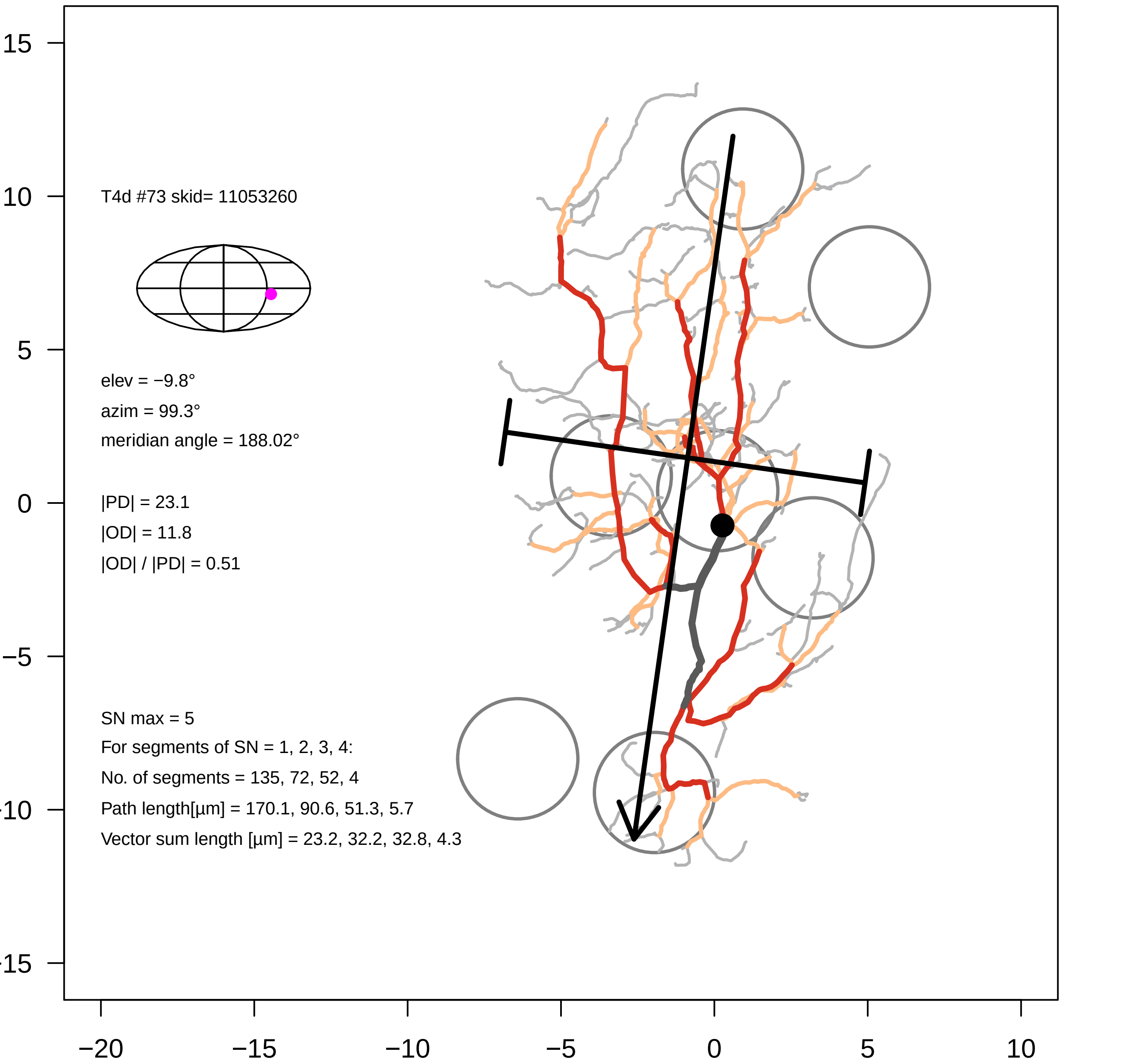

T4d #73 skid= 11053260
elev = −9.8°
azim = 99.3°
meridian angle = 188.02°
|PD| = 23.1
|OD| = 11.8
|OD| / |PD| = 0.51
SN max = 5
For segments of SN = 1, 2, 3, 4:
No. of segments = 135, 72, 52, 4
Path length[µm] = 170.1, 90.6, 51.3, 5.7
Vector sum length [µm] = 23.2, 32.2, 32.8, 4.3
−20
−15
−10
−5
0
5
10
15

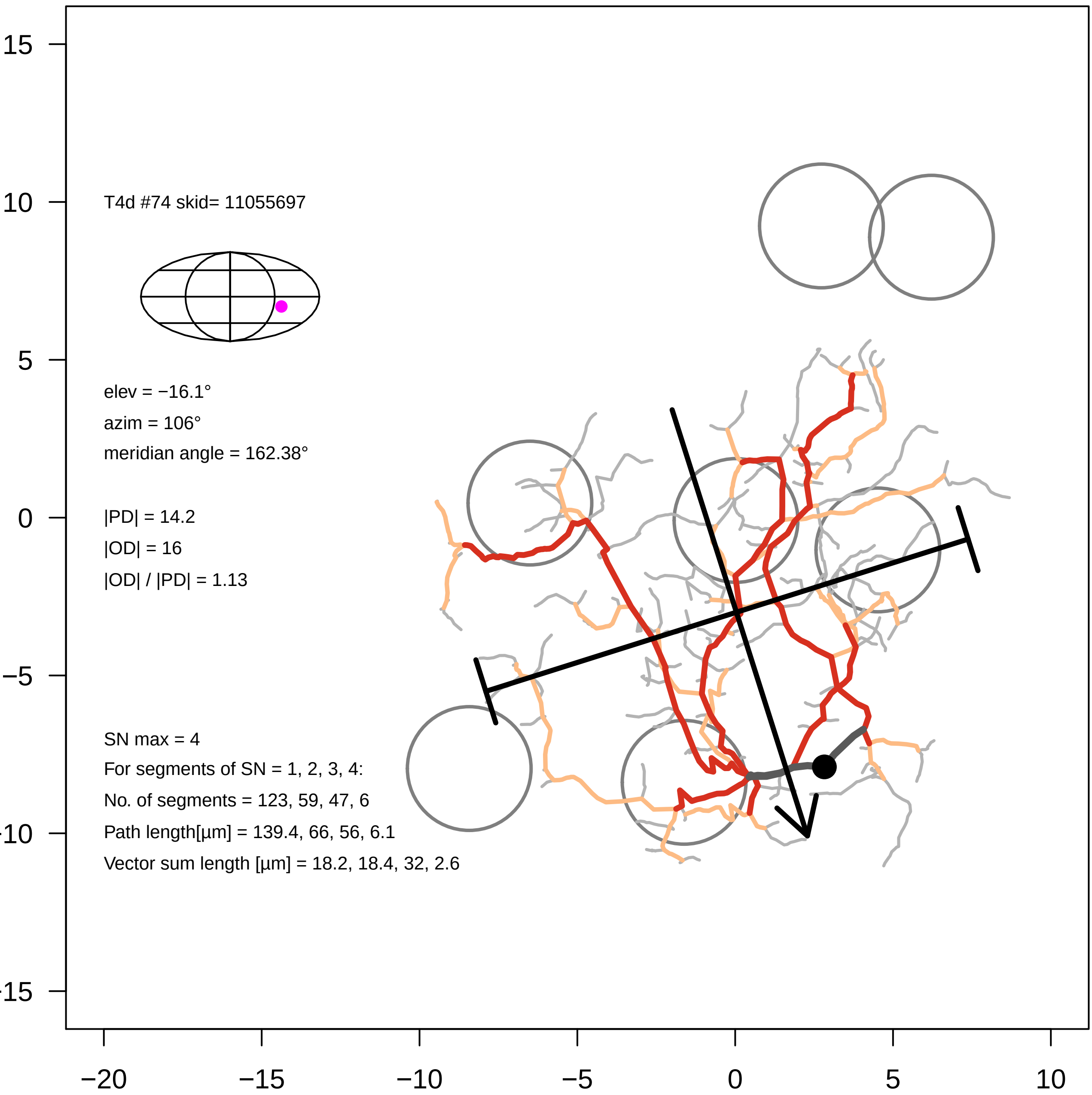

T4d #74 skid= 11055697
elev = −16.1°
azim = 106°
meridian angle = 162.38°
|PD| = 14.2
|OD| = 16
|OD| / |PD| = 1.13
SN max = 4
For segments of SN = 1, 2, 3, 4:
No. of segments = 123, 59, 47, 6
Path length[µm] = 139.4, 66, 56, 6.1
Vector sum length [µm] = 18.2, 18.4, 32, 2.6
15
10
5
0
−5
−10
−15
−20
−15
−10
−5
0
5
10

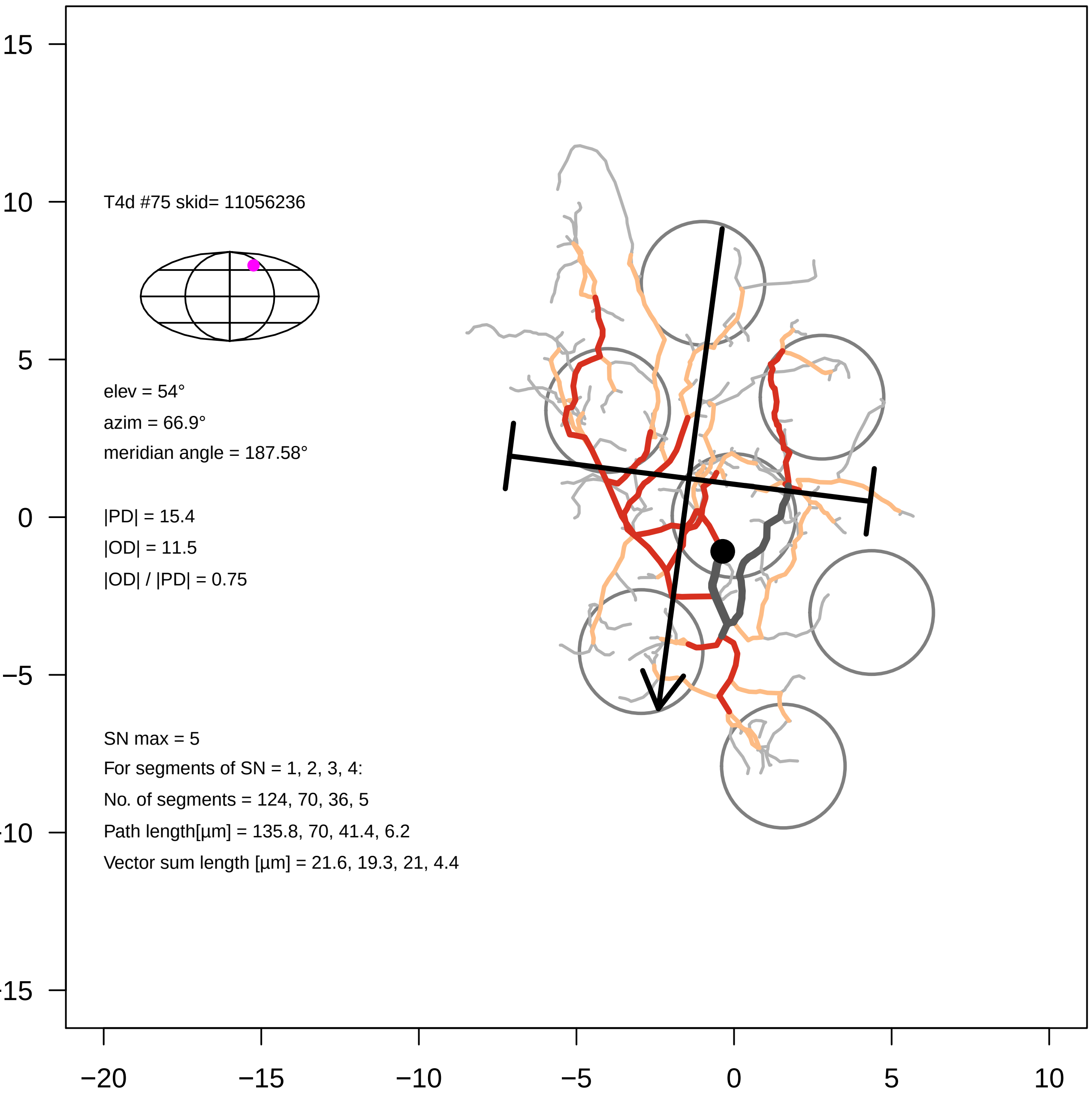

T4d #75 skid= 11056236
elev = 54°
azim = 66.9°
meridian angle = 187.58°
|PD| = 15.4
|OD| = 11.5
|OD| / |PD| = 0.75
SN max = 5
For segments of SN = 1, 2, 3, 4:
No. of segments = 124, 70, 36, 5
Path length[µm] = 135.8, 70, 41.4, 6.2
Vector sum length [µm] = 21.6, 19.3, 21, 4.4
15
10
5
0
−5
−20
−15
−10
−5
0
5
10

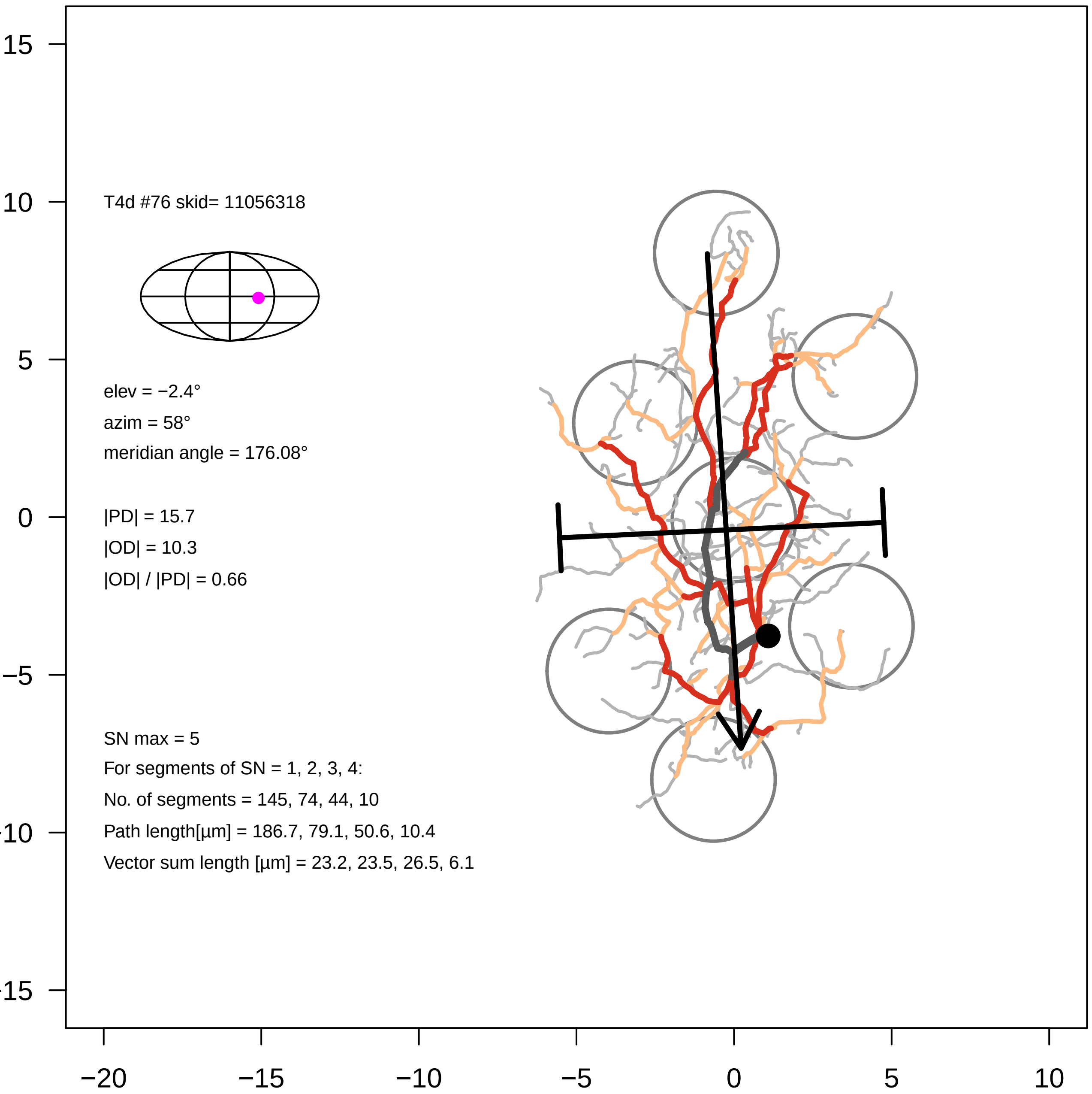

T4d #76 skid= 11056318
elev = −2.4°
azim = 58°
meridian angle = 176.08°
|PD| = 15.7
|OD| = 10.3
|OD| / |PD| = 0.66
SN max = 5
For segments of SN = 1, 2, 3, 4:
No. of segments = 145, 74, 44, 10
Path length[µm] = 186.7, 79.1, 50.6, 10.4
Vector sum length [µm] = 23.2, 23.5, 26.5, 6.1
15
10
5
0
−5
−10
−15
−20
−15
−10
−5
0
5
10

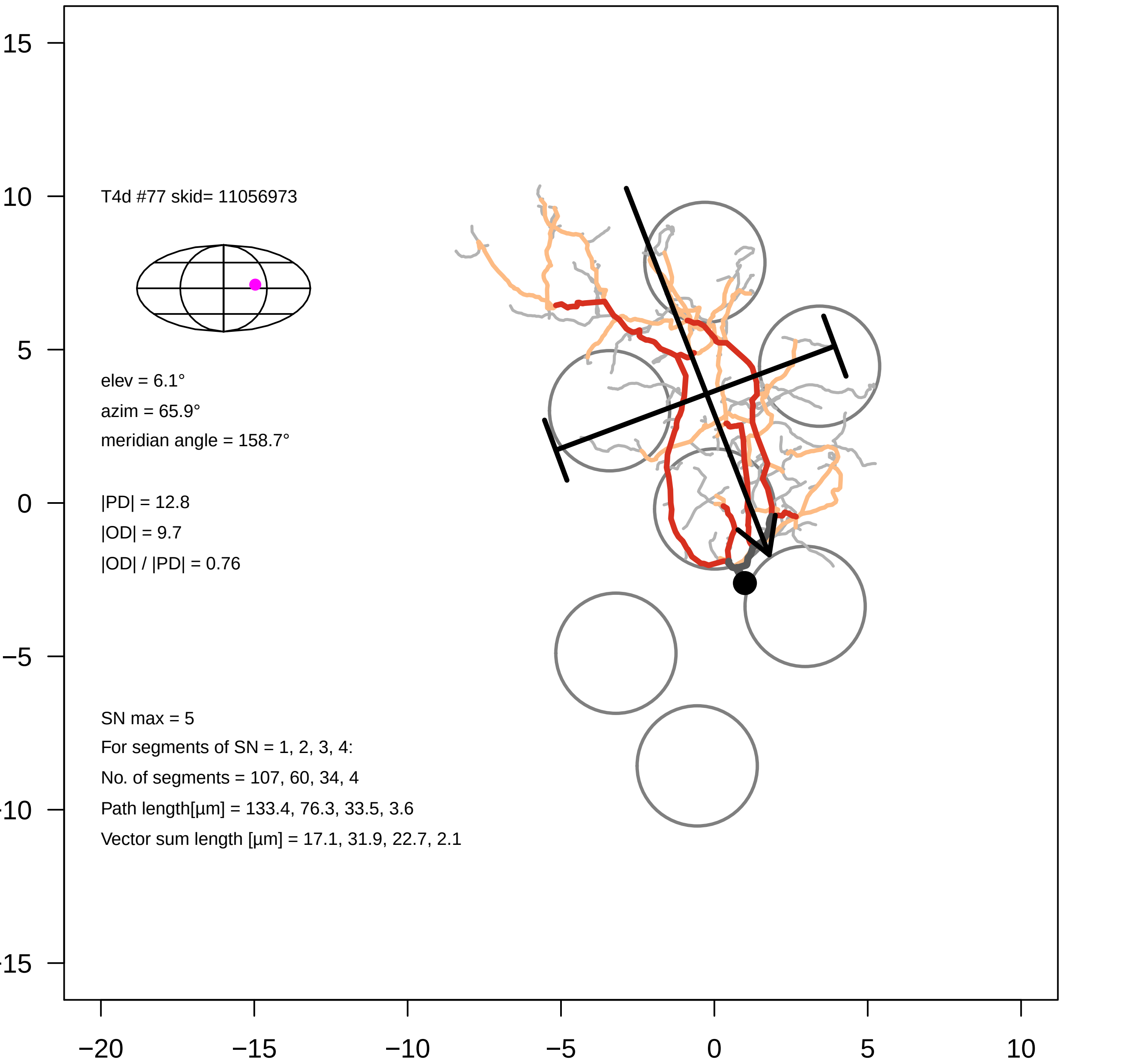

T4d #77 skid= 11056973
elev = 6.1°
azim = 65.9°
meridian angle = 158.7°
|PD| = 12.8
|OD| = 9.7
|OD| / |PD| = 0.76
SN max = 5
For segments of SN = 1, 2, 3, 4:
No. of segments = 107, 60, 34, 4
Path length[µm] = 133.4, 76.3, 33.5, 3.6
Vector sum length [µm] = 17.1, 31.9, 22.7, 2.1
15
10
5
0
−5
−10
−15
−20
−15
−10
−5
0
5
10

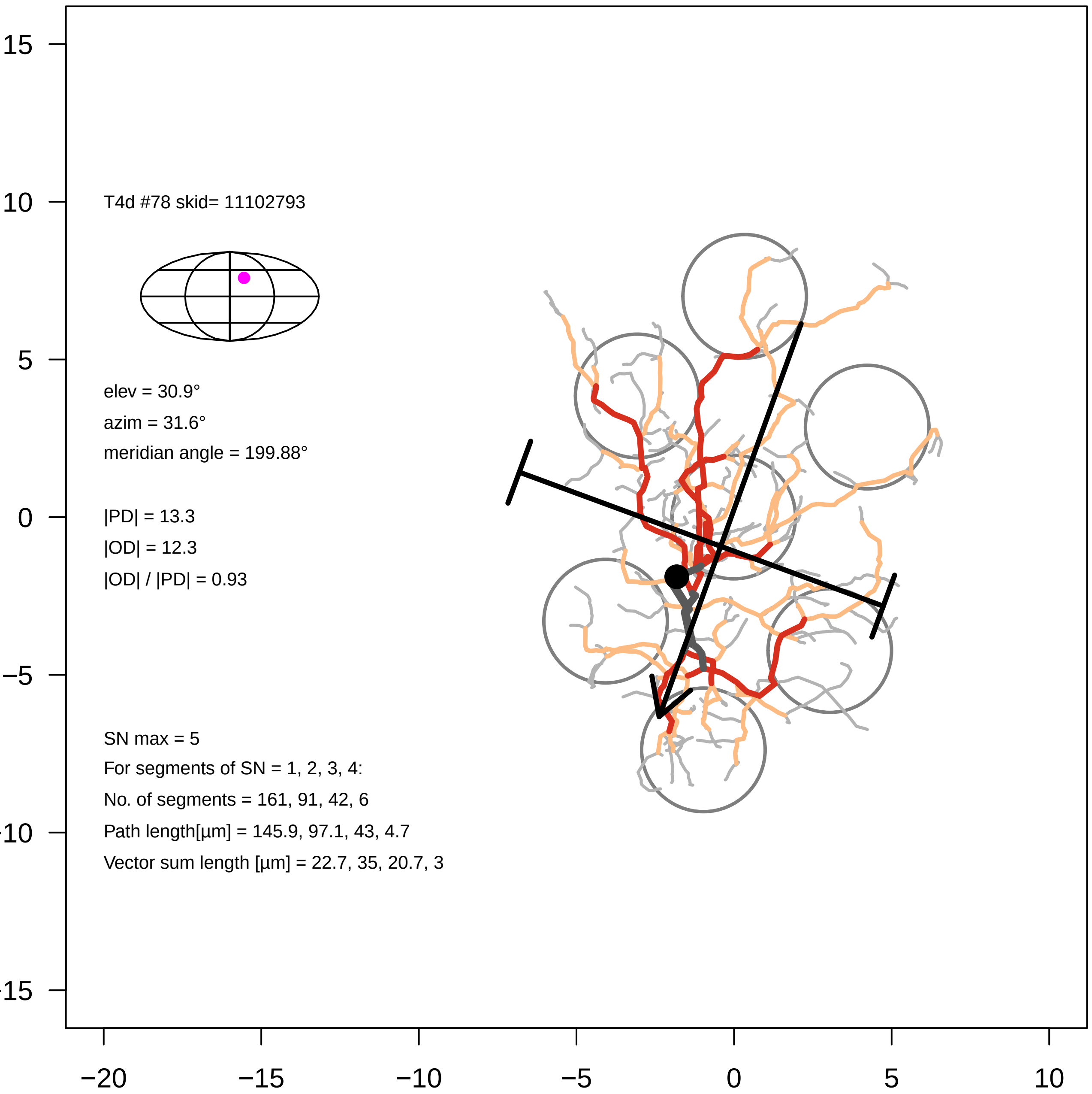
T4d #78 skid= 11102793
elev = 30.9°
azim = 31.6°
meridian angle = 199.88°
|PD| = 13.3
|OD| = 12.3
|OD| / |PD| = 0.93
SN max = 5
For segments of SN = 1, 2, 3, 4:
No. of segments = 161, 91, 42, 6
Path length[µm] = 145.9, 97.1, 43, 4.7
Vector sum length [µm] = 22.7, 35, 20.7, 3
15
10
5
0
-5
-10
-15
−20
−15
−10
−5
0
5
10

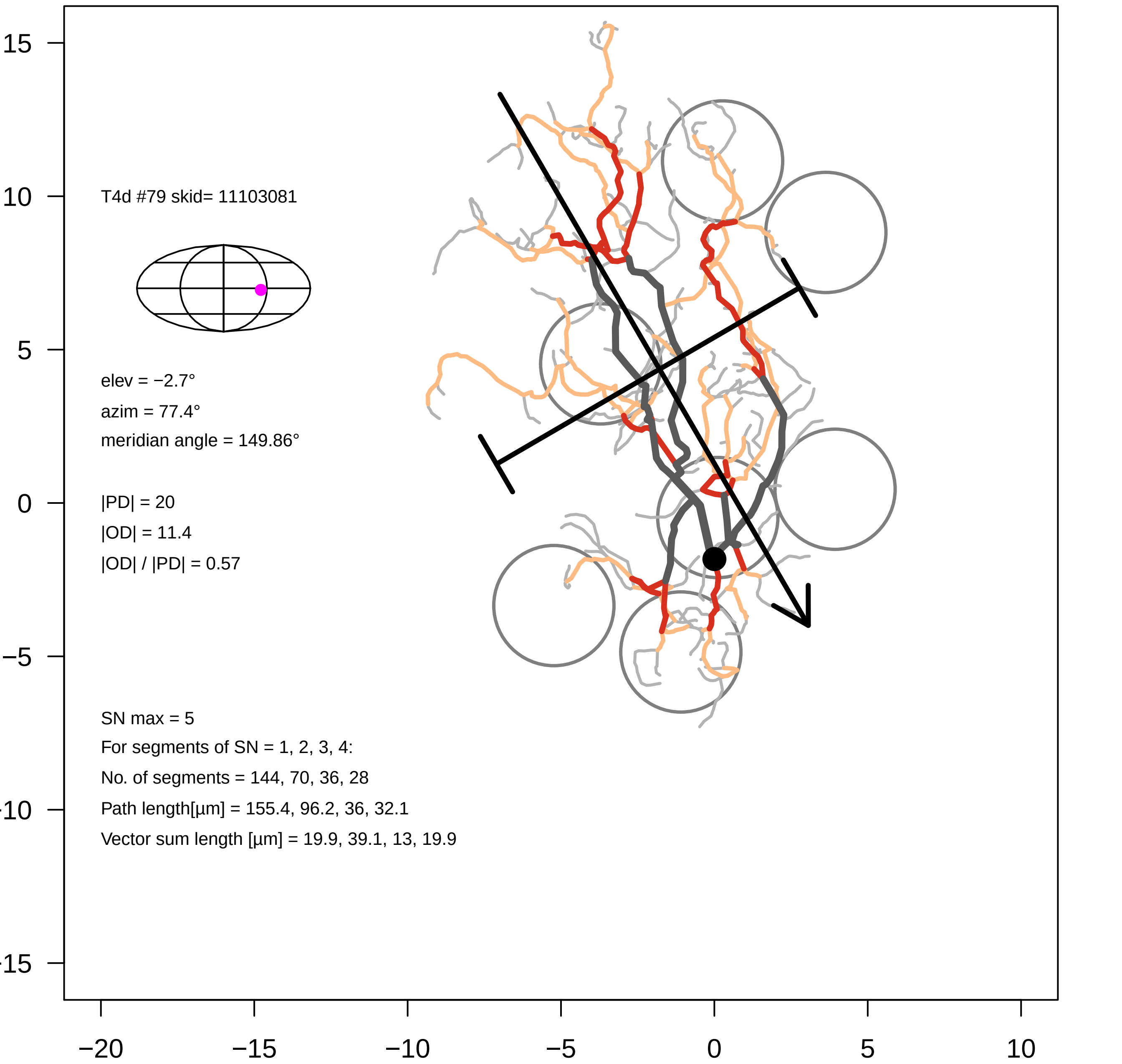
T4d #79 skid= 11103081
elev = −2.7°
azim = 77.4°
meridian angle = 149.86°
|PD| = 20
|OD| = 11.4
|OD| / |PD| = 0.57
SN max = 5
For segments of SN = 1, 2, 3, 4:
No. of segments = 144, 70, 36, 28
Path length[µm] = 155.4, 96.2, 36, 32.1
Vector sum length [µm] = 19.9, 39.1, 13, 19.9
15
10
5
0
−5
-10
-15
−20
−15
−10
−5
0
5
10

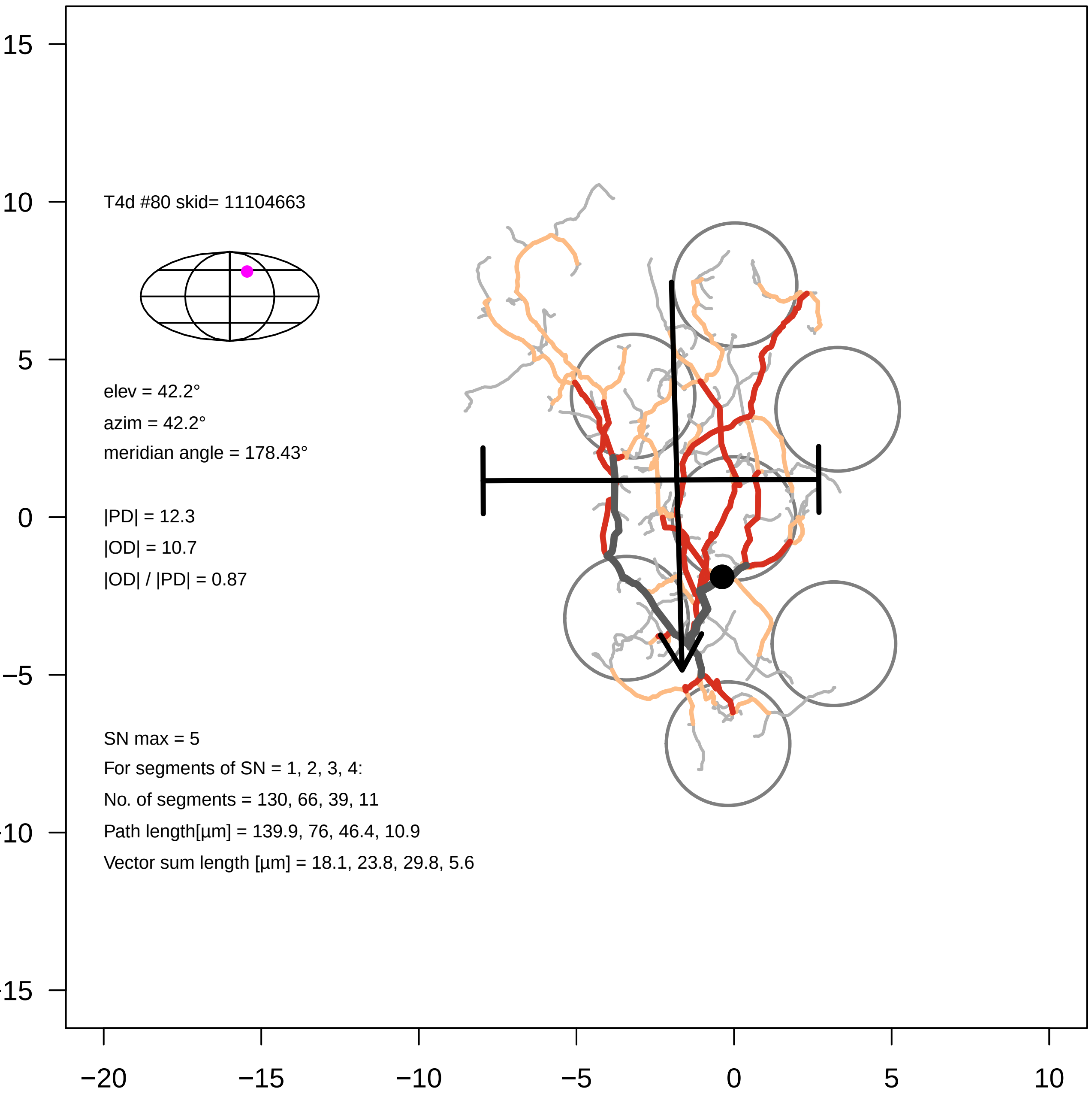

T4d #80 skid= 11104663
elev = 42.2°
azim = 42.2°
meridian angle = 178.43°
|PD| = 12.3
|OD| = 10.7
|OD| / |PD| = 0.87
SN max = 5
For segments of SN = 1, 2, 3, 4:
No. of segments = 130, 66, 39, 11
Path length[µm] = 139.9, 76, 46.4, 10.9
Vector sum length [µm] = 18.1, 23.8, 29.8, 5.6
15
10
5
0
−5
-10
-15
−20
−15
−10
−5
0
5
10

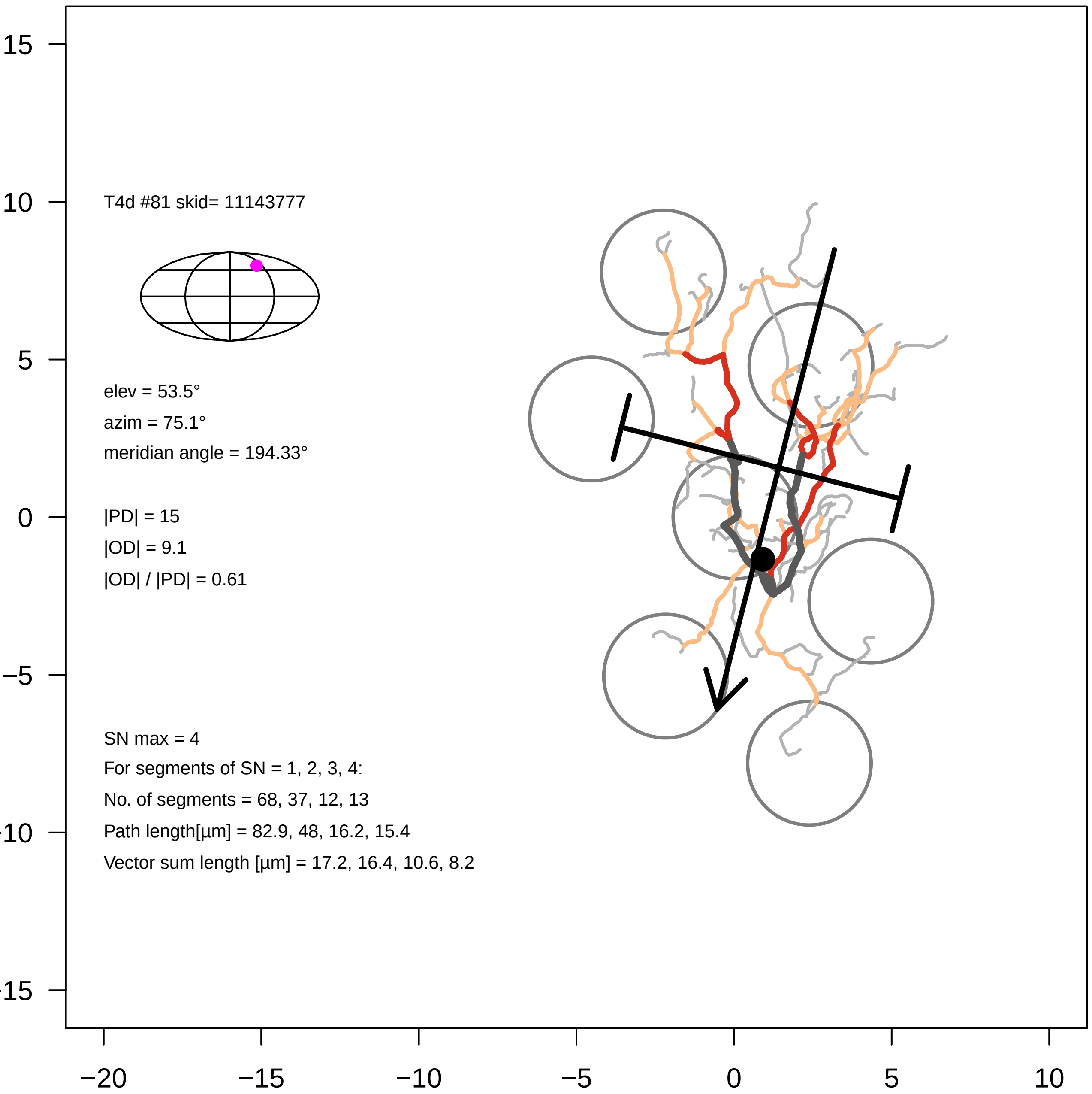
T4d #81 skid= 11143777
elev = 53.5°
azim = 75.1°
meridian angle = 194.33°
|PD| = 15
|OD| = 9.1
|OD| / |PD| = 0.61
SN max = 4
For segments of SN = 1, 2, 3, 4:
No. of segments = 68, 37, 12, 13
Path length[µm] = 82.9, 48, 16.2, 15.4
Vector sum length [µm] = 17.2, 16.4, 10.6, 8.2
15
10
5
0
−5
−10
−15
−20
−15
−10
−5
0
5
10

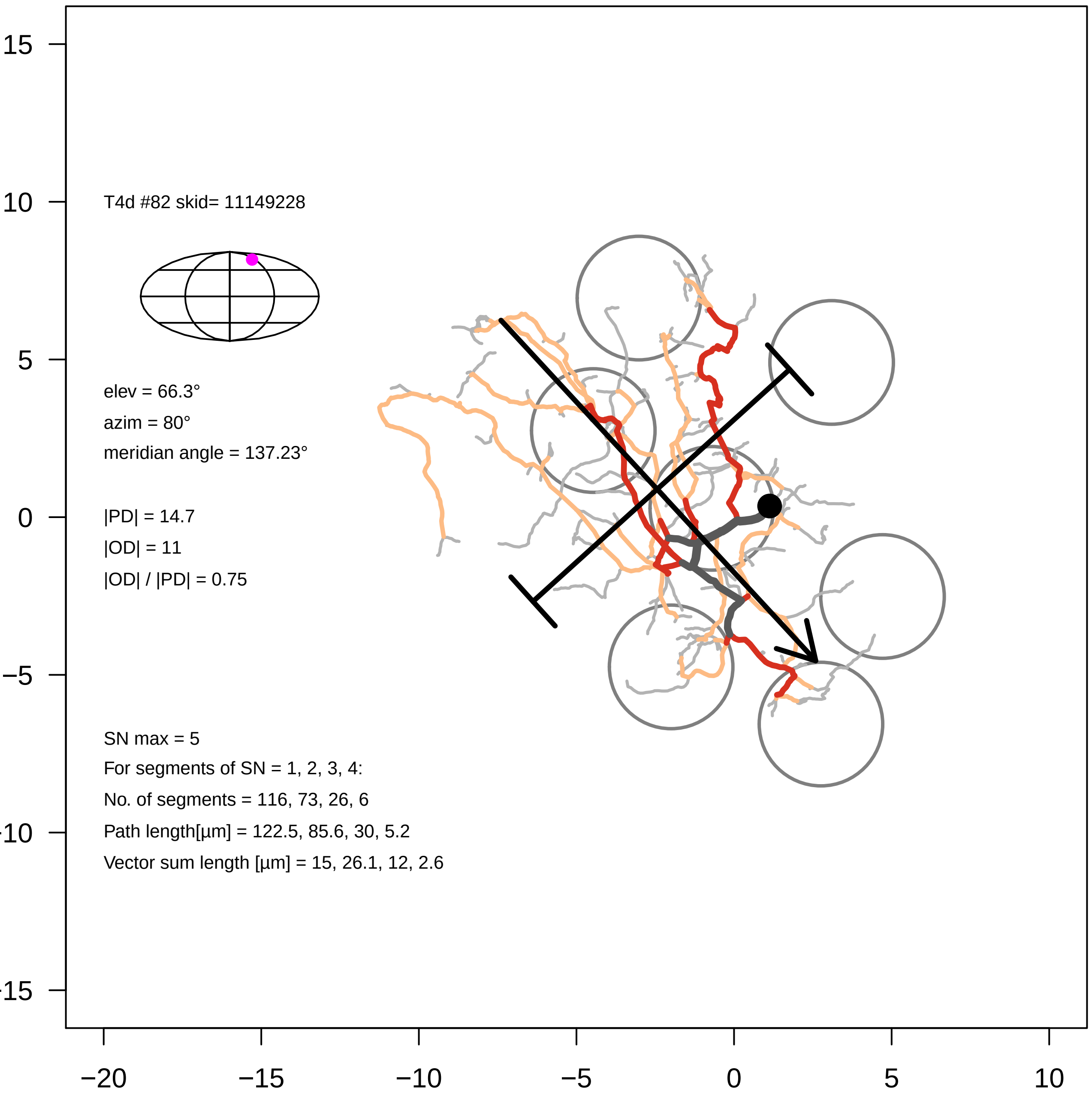
T4d #82 skid= 11149228
elev = 66.3°
azim = 80°
meridian angle = 137.23°
|PD| = 14.7
|OD| = 11
|OD| / |PD| = 0.75
SN max = 5
For segments of SN = 1, 2, 3, 4:
No. of segments = 116, 73, 26, 6
Path length[µm] = 122.5, 85.6, 30, 5.2
Vector sum length [µm] = 15, 26.1, 12, 2.6
15
10
5
0
−5
−10
−15
−20
−15
−10
−5
0
5
10

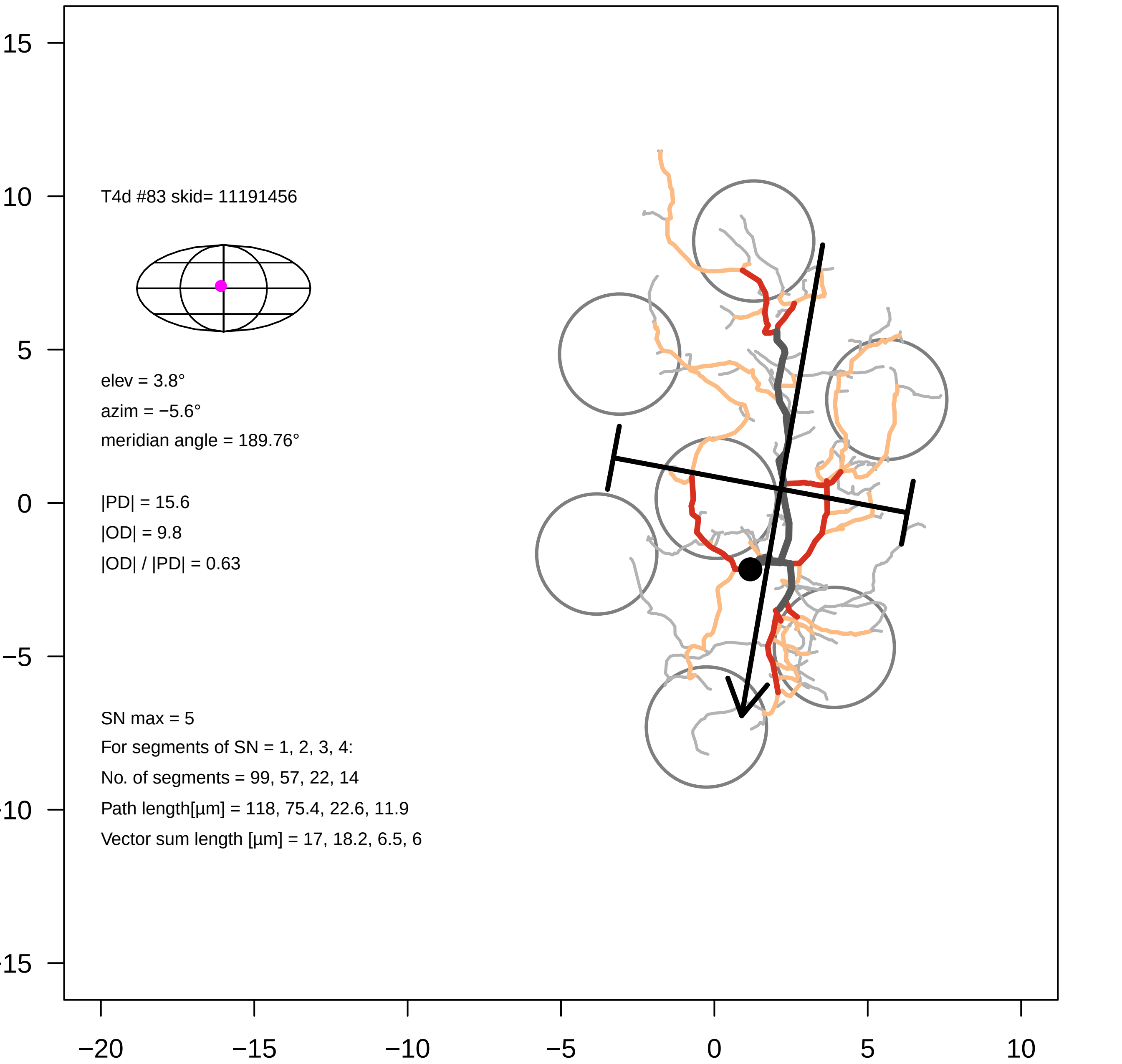

T4d #83 skid= 11191456
elev = 3.8°
azim = −5.6°
meridian angle = 189.76°
|PD| = 15.6
|OD| = 9.8
|OD| / |PD| = 0.63
SN max = 5
For segments of SN = 1, 2, 3, 4:
No. of segments = 99, 57, 22, 14
Path length[µm] = 118, 75.4, 22.6, 11.9
Vector sum length [µm] = 17, 18.2, 6.5, 6
15
10
5
0
−5
−10
−15
−20
−15
−10
−5
0
5
10

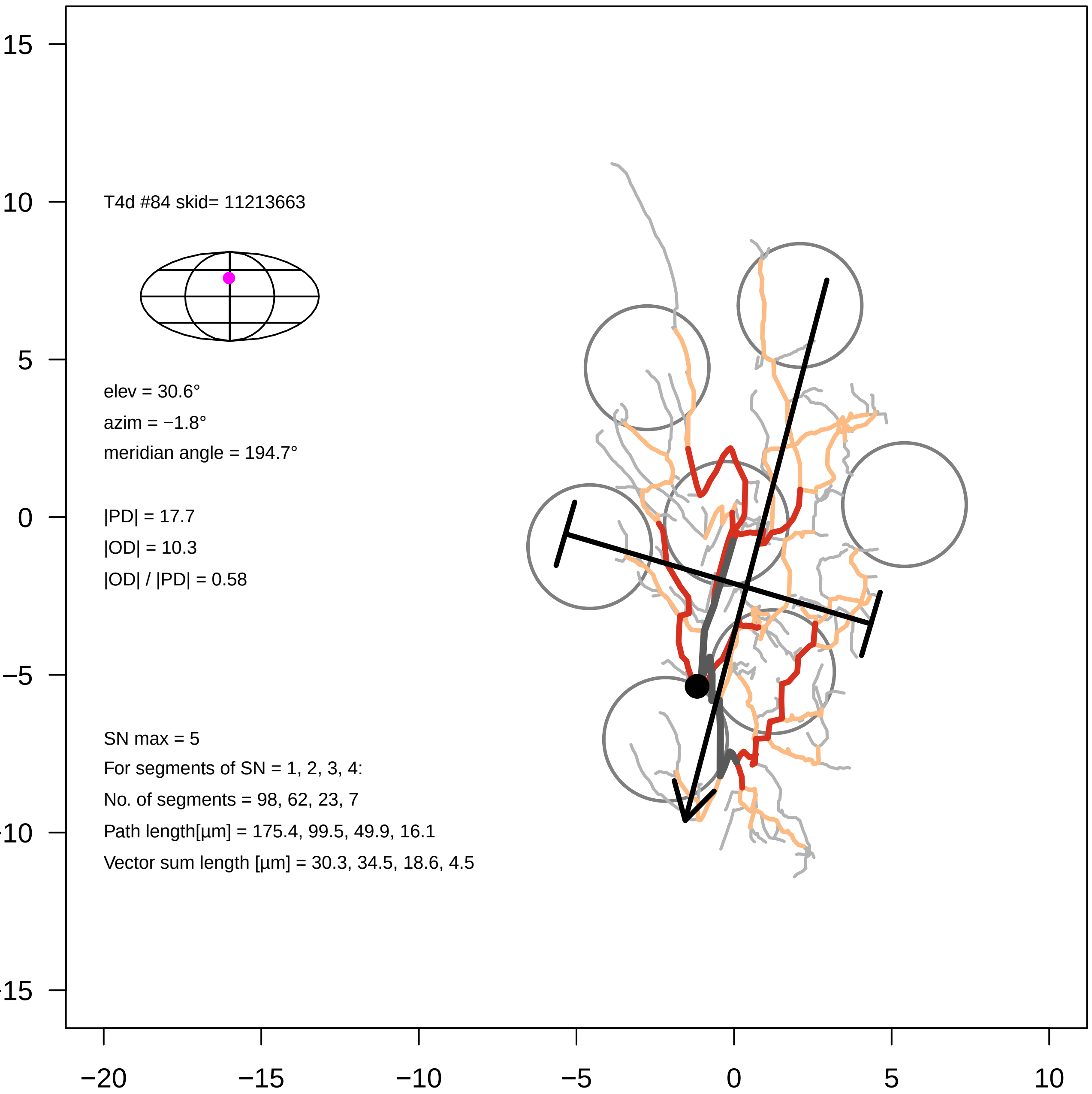
T4d #84 skid= 11213663
elev = 30.6°
azim = −1.8°
meridian angle = 194.7°
|PD| = 17.7
|OD| = 10.3
|OD| / |PD| = 0.58
SN max = 5
For segments of SN = 1, 2, 3, 4:
No. of segments = 98, 62, 23, 7
Path length[µm] = 175.4, 99.5, 49.9, 16.1
Vector sum length [µm] = 30.3, 34.5, 18.6, 4.5
15
10
5
0
−5
−10
−15
−20
−15
−10
−5
0
5
10

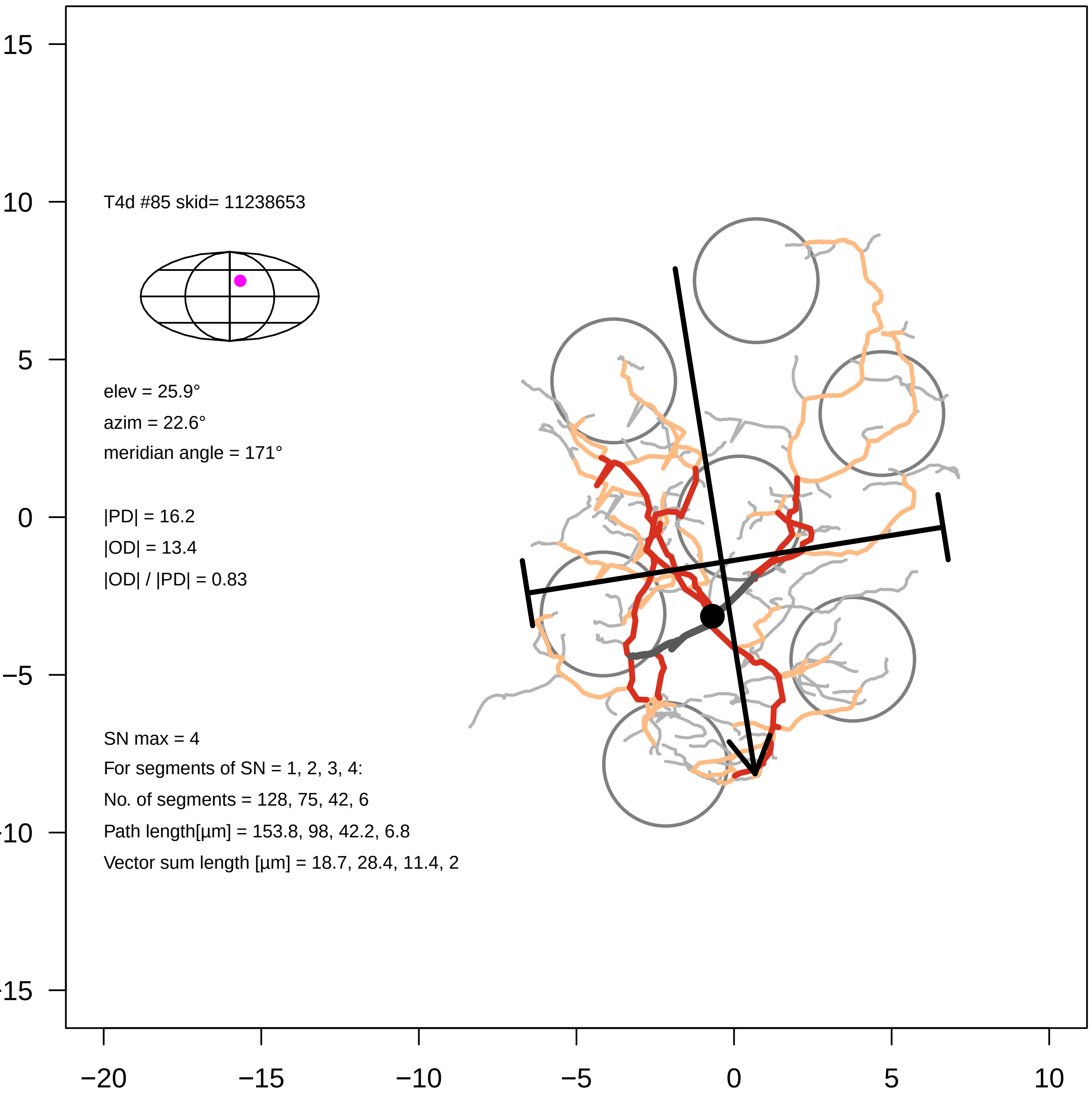
T4d #85 skid= 11238653
elev = 25.9°
azim = 22.6°
meridian angle = 171°
|PD| = 16.2
|OD| = 13.4
|OD| / |PD| = 0.83
SN max = 4
For segments of SN = 1, 2, 3, 4:
No. of segments = 128, 75, 42, 6
Path length[µm] = 153.8, 98, 42.2, 6.8
Vector sum length [µm] = 18.7, 28.4, 11.4, 2
15
10
5
0
−5
−10
−15
−20
−15
−10
−5
0
5
10

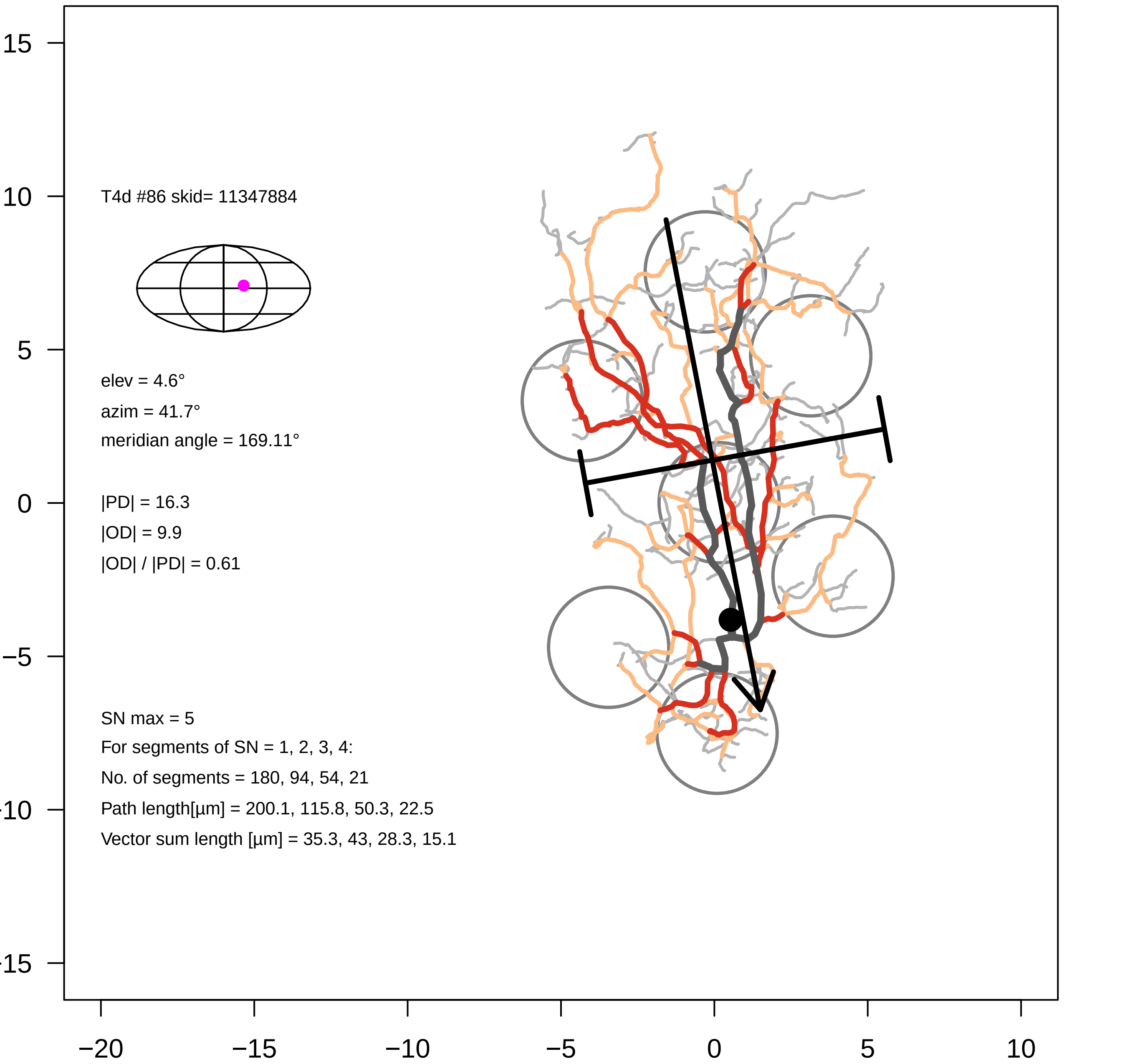

T4d #86 skid= 11347884
elev = 4.6°
azim = 41.7°
meridian angle = 169.11°
|PD| = 16.3
|OD| = 9.9
|OD| / |PD| = 0.61
SN max = 5
For segments of SN = 1, 2, 3, 4:
No. of segments = 180, 94, 54, 21
Path length[µm] = 200.1, 115.8, 50.3, 22.5
Vector sum length [µm] = 35.3, 43, 28.3, 15.1
15
10
5
0
−5
−10
−15
−20
−15
−10
−5
0
5
10

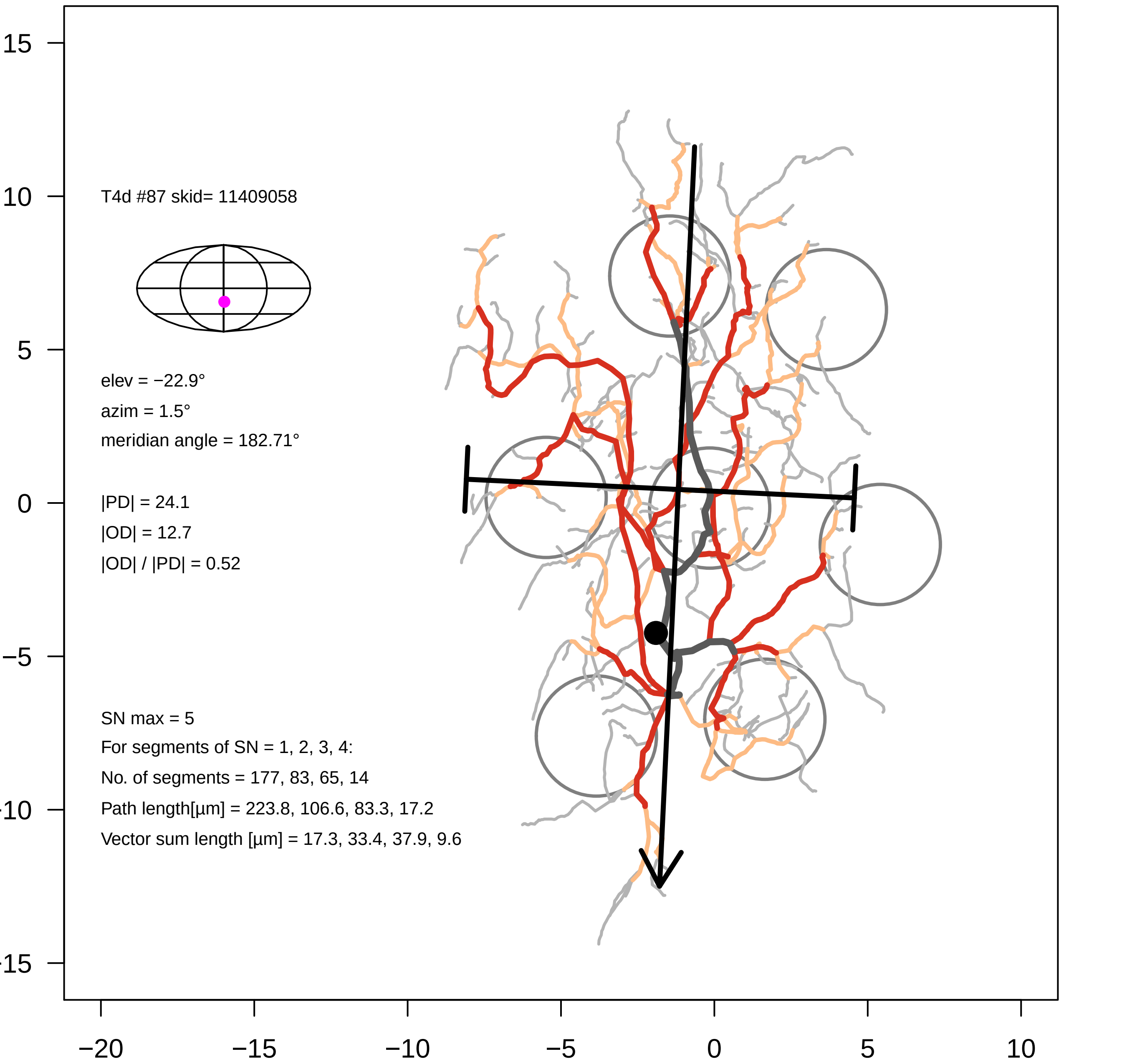
T4d #87 skid= 11409058
elev = −22.9°
azim = 1.5°
meridian angle = 182.71°
|PD| = 24.1
|OD| = 12.7
|OD| / |PD| = 0.52
SN max = 5
For segments of SN = 1, 2, 3, 4:
No. of segments = 177, 83, 65, 14
Path length[µm] = 223.8, 106.6, 83.3, 17.2
Vector sum length [µm] = 17.3, 33.4, 37.9, 9.6
15
10
5
0
−5
−10
−15
−20
−15
−10
−5
0
5
10

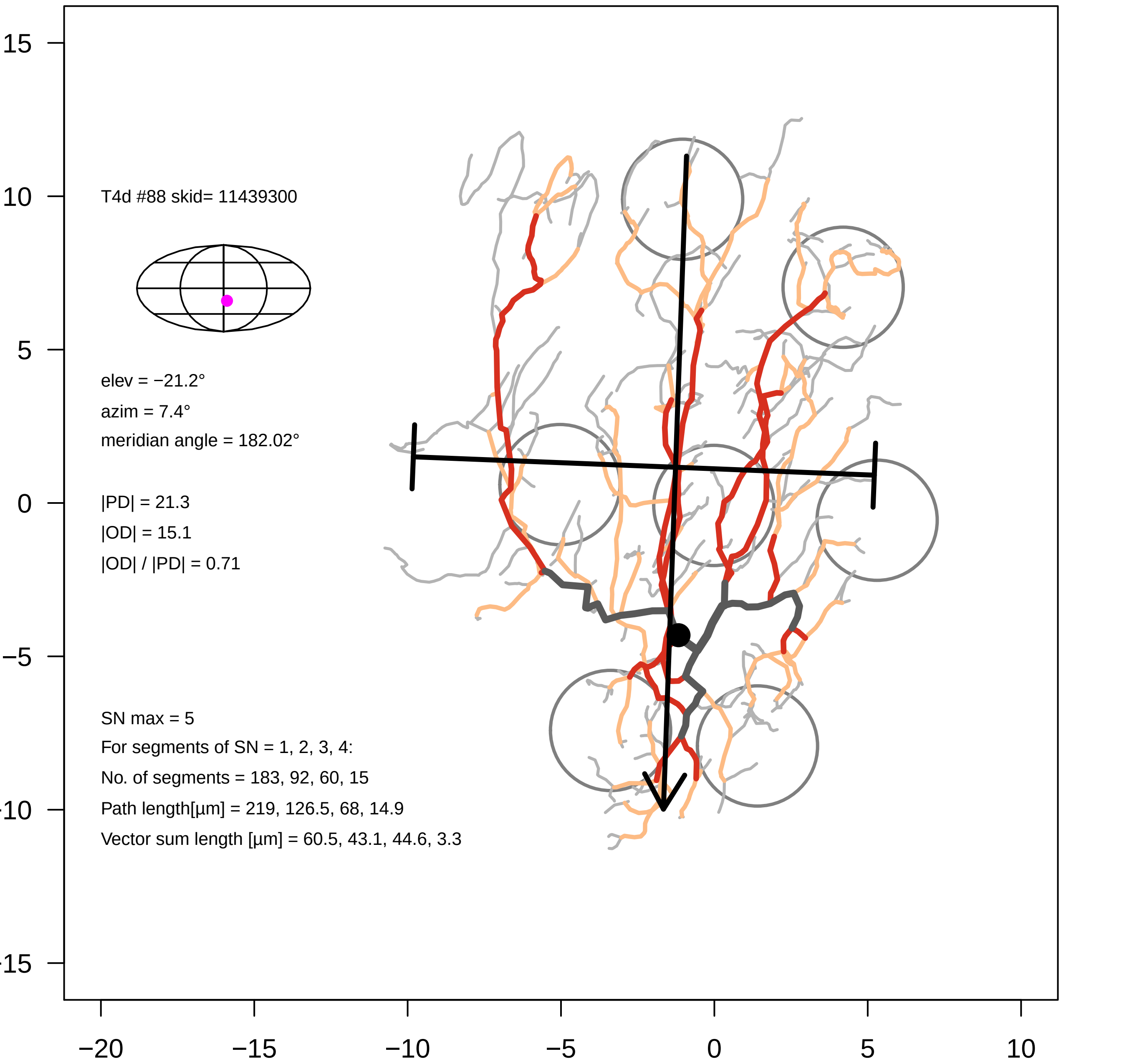

T4d #88 skid= 11439300
elev = −21.2°
azim = 7.4°
meridian angle = 182.02°
|PD| = 21.3
|OD| = 15.1
|OD| / |PD| = 0.71
SN max = 5
For segments of SN = 1, 2, 3, 4:
No. of segments = 183, 92, 60, 15
Path length[µm] = 219, 126.5, 68, 14.9
Vector sum length [µm] = 60.5, 43.1, 44.6, 3.3
15
10
5
0
−5
-10
-15
−20
−15
−10
−5
0
5
10

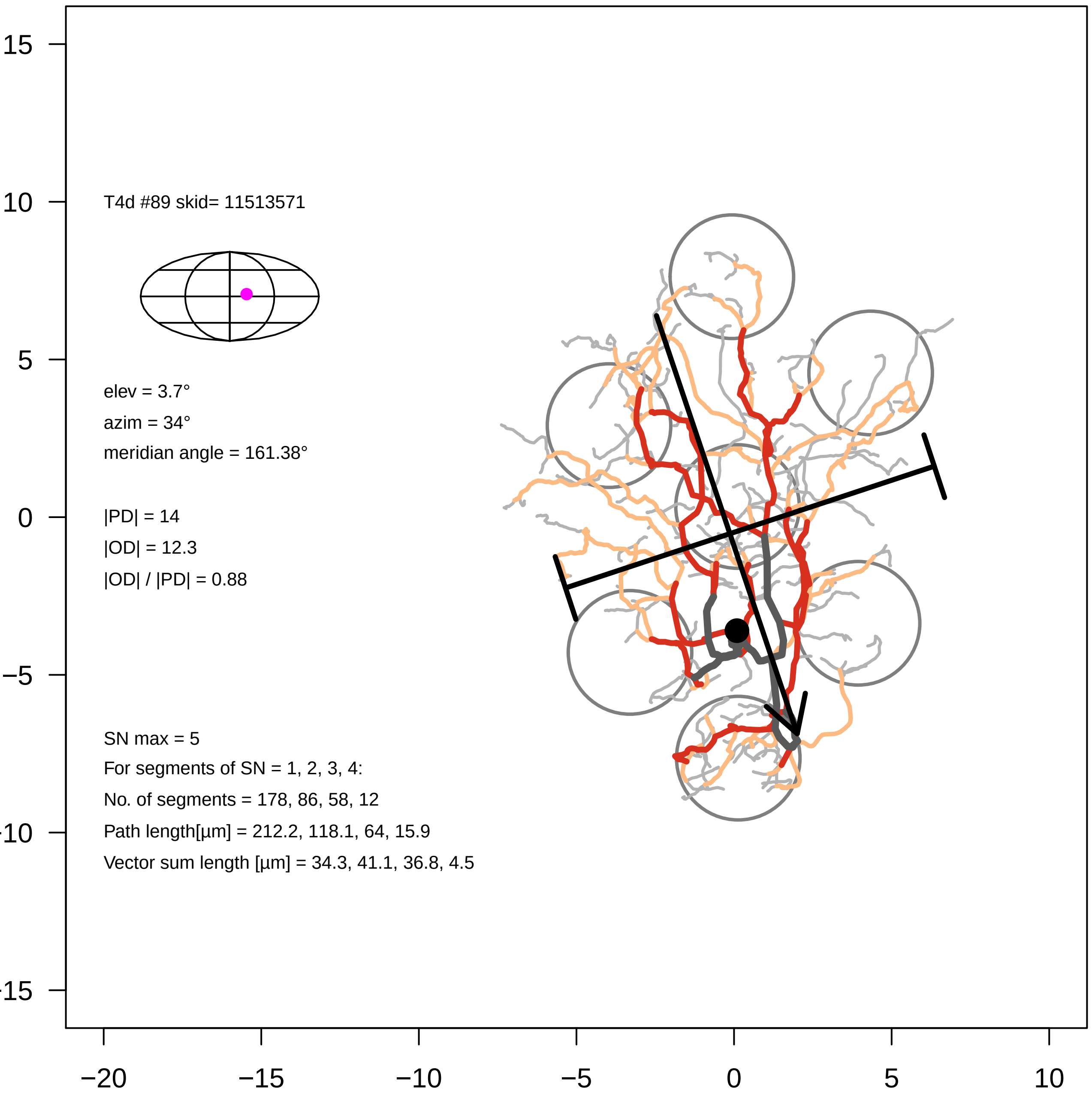

T4d #89 skid= 11513571
elev = 3.7°
azim = 34°
meridian angle = 161.38°
|PD| = 14
|OD| = 12.3
|OD| / |PD| = 0.88
SN max = 5
For segments of SN = 1, 2, 3, 4:
No. of segments = 178, 86, 58, 12
Path length[µm] = 212.2, 118.1, 64, 15.9
Vector sum length [µm] = 34.3, 41.1, 36.8, 4.5
15
10
5
0
−5
−10
−15
−20
−15
−10
−5
0
5
10

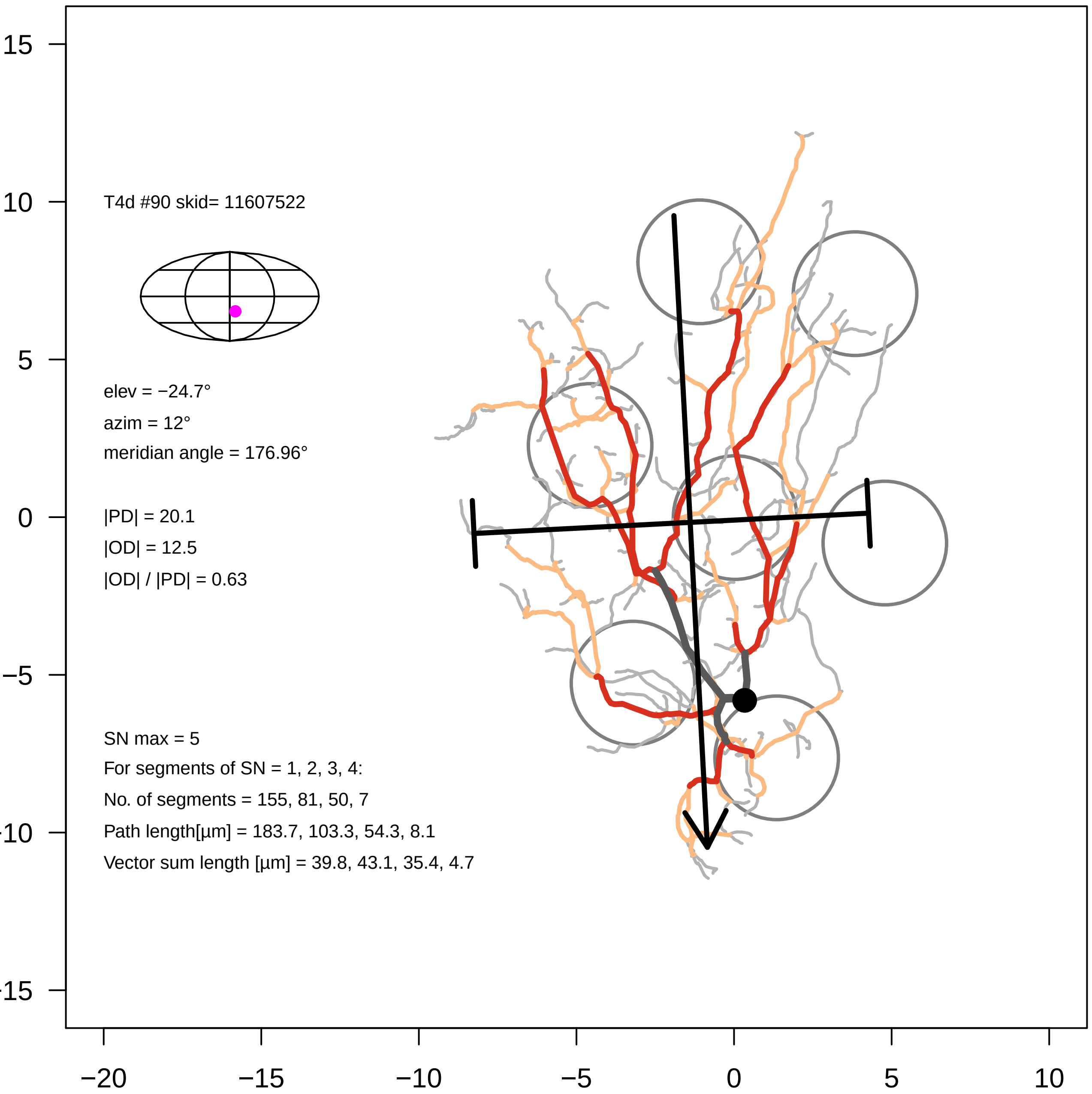

T4d #90 skid= 11607522
elev = −24.7°
azim = 12°
meridian angle = 176.96°
|PD| = 20.1
|OD| = 12.5
|OD| / |PD| = 0.63
SN max = 5
For segments of SN = 1, 2, 3, 4:
No. of segments = 155, 81, 50, 7
Path length[µm] = 183.7, 103.3, 54.3, 8.1
Vector sum length [µm] = 39.8, 43.1, 35.4, 4.7
15
10
5
0
−5
−10
−15
−20
−15
−10
−5
0
5
10

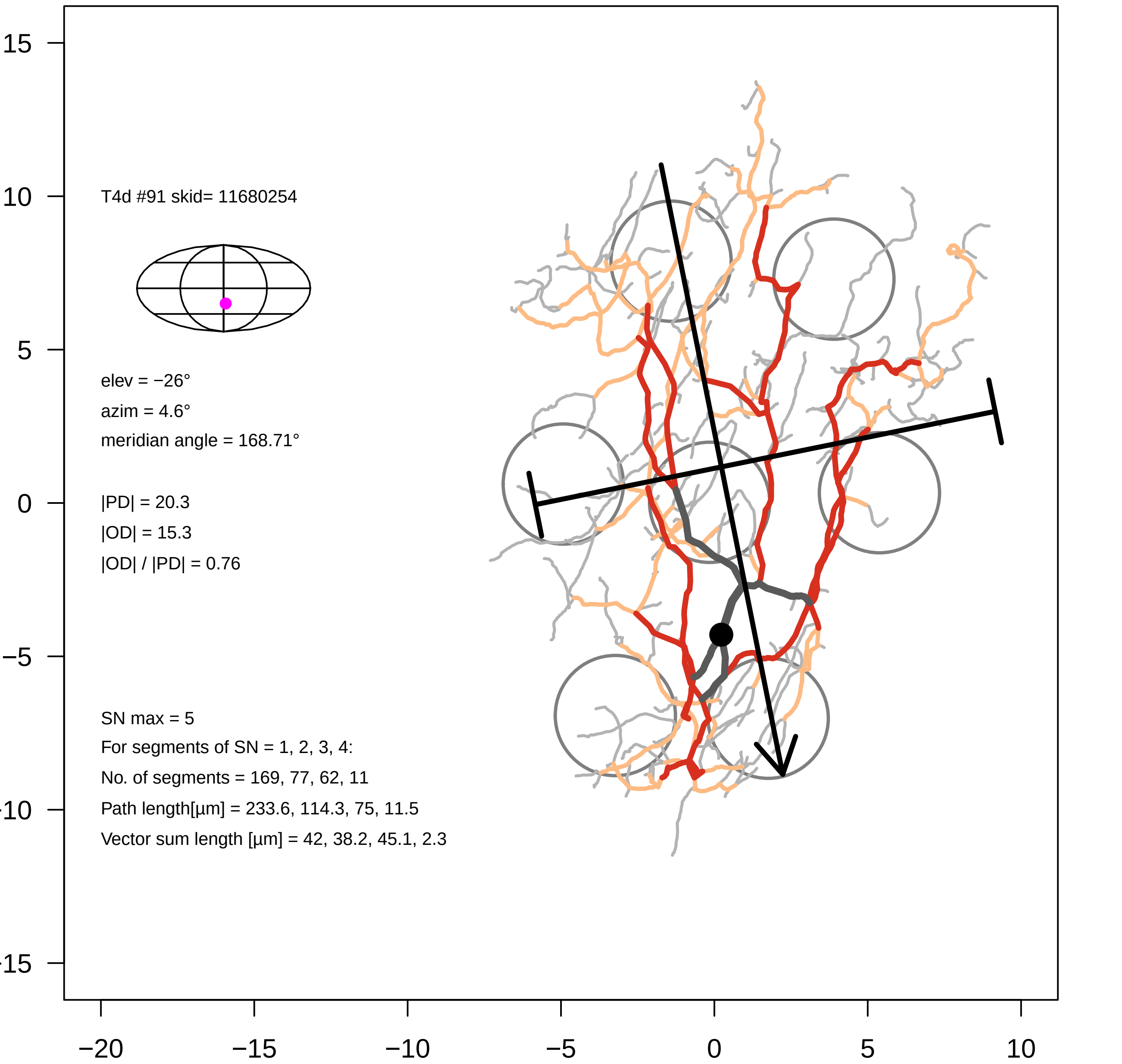

T4d #91 skid= 11680254
elev = −26°
azim = 4.6°
meridian angle = 168.71°
|PD| = 20.3
|OD| = 15.3
|OD| / |PD| = 0.76
SN max = 5
For segments of SN = 1, 2, 3, 4:
No. of segments = 169, 77, 62, 11
Path length[µm] = 233.6, 114.3, 75, 11.5
Vector sum length [µm] = 42, 38.2, 45.1, 2.3
15
10
5
0
−5
-10
-15
−20
−15
−10
−5
0
5
10

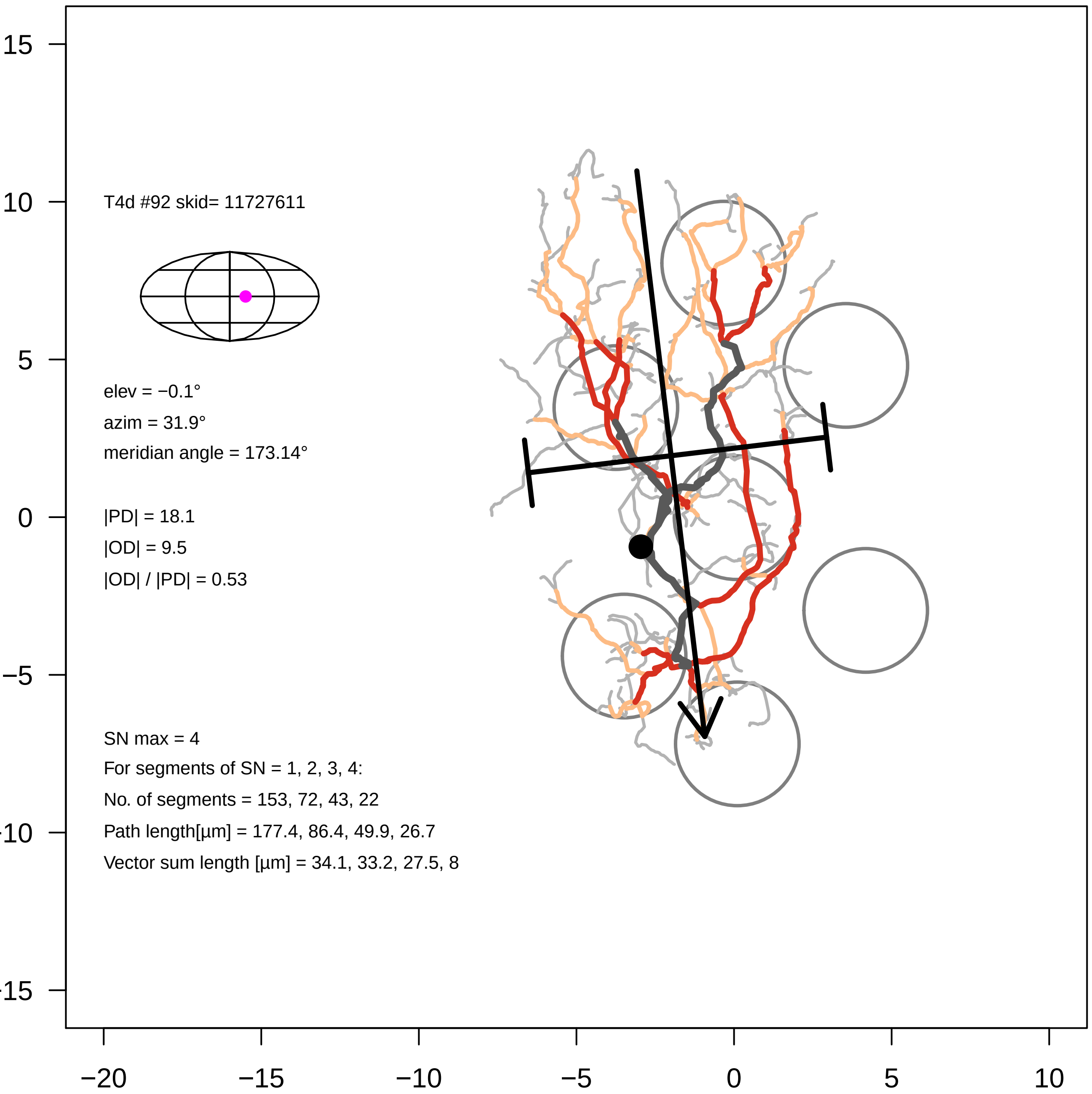

T4d #92 skid= 11727611
elev = −0.1°
azim = 31.9°
meridian angle = 173.14°
|PD| = 18.1
|OD| = 9.5
|OD| / |PD| = 0.53
SN max = 4
For segments of SN = 1, 2, 3, 4:
No. of segments = 153, 72, 43, 22
Path length[µm] = 177.4, 86.4, 49.9, 26.7
Vector sum length [µm] = 34.1, 33.2, 27.5, 8
15
10
5
0
−5
−10
−15
−20
−15
−10
−5
0
5
10

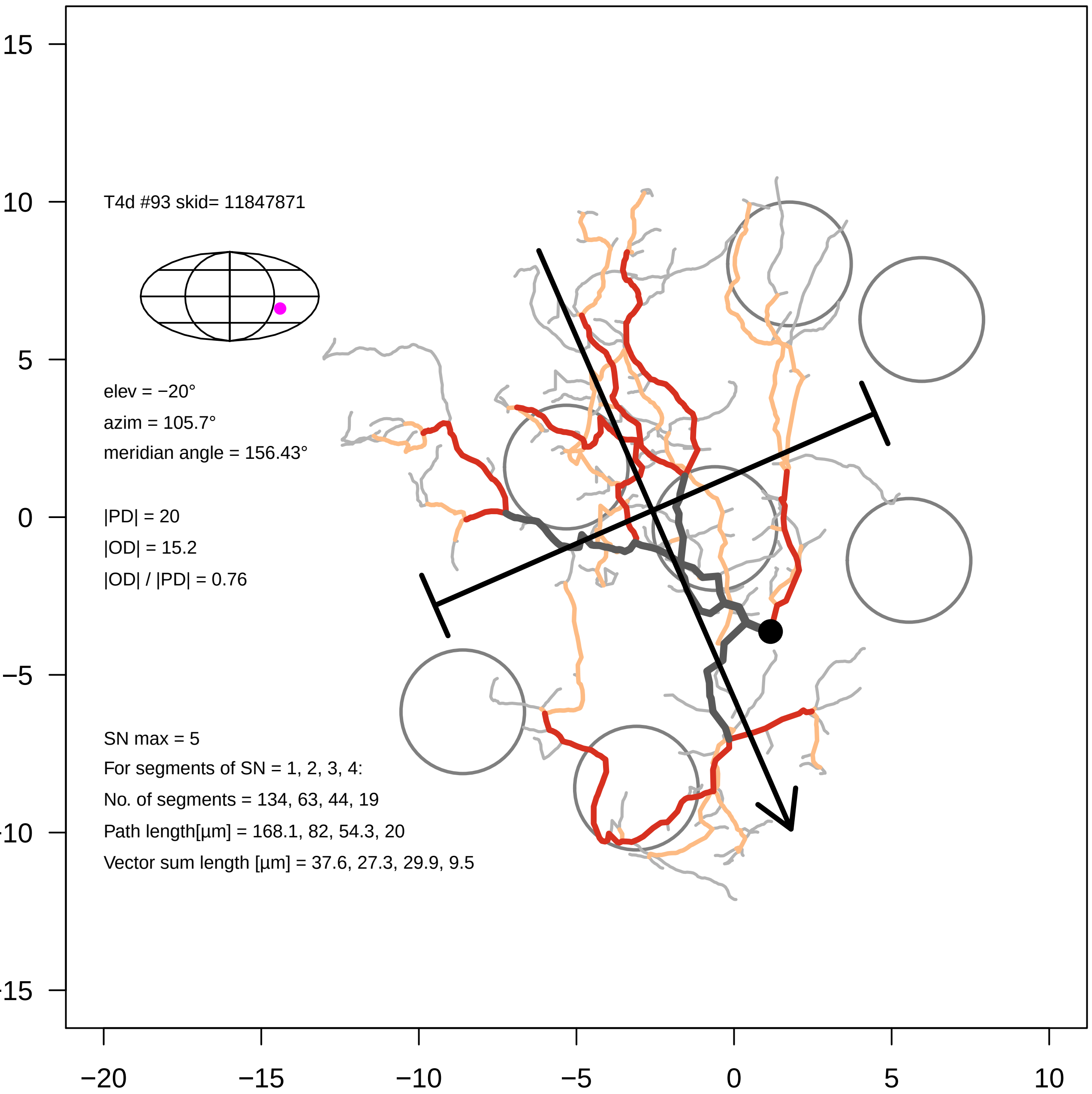
T4d #93 skid= 11847871
elev = −20°
azim = 105.7°
meridian angle = 156.43°
|PD| = 20
|OD| = 15.2
|OD| / |PD| = 0.76
SN max = 5
For segments of SN = 1, 2, 3, 4:
No. of segments = 134, 63, 44, 19
Path length[µm] = 168.1, 82, 54.3, 20
Vector sum length [µm] = 37.6, 27.3, 29.9, 9.5
15
10
5
0
−5
−10
−15
−20
−15
−10
−5
0
5
10

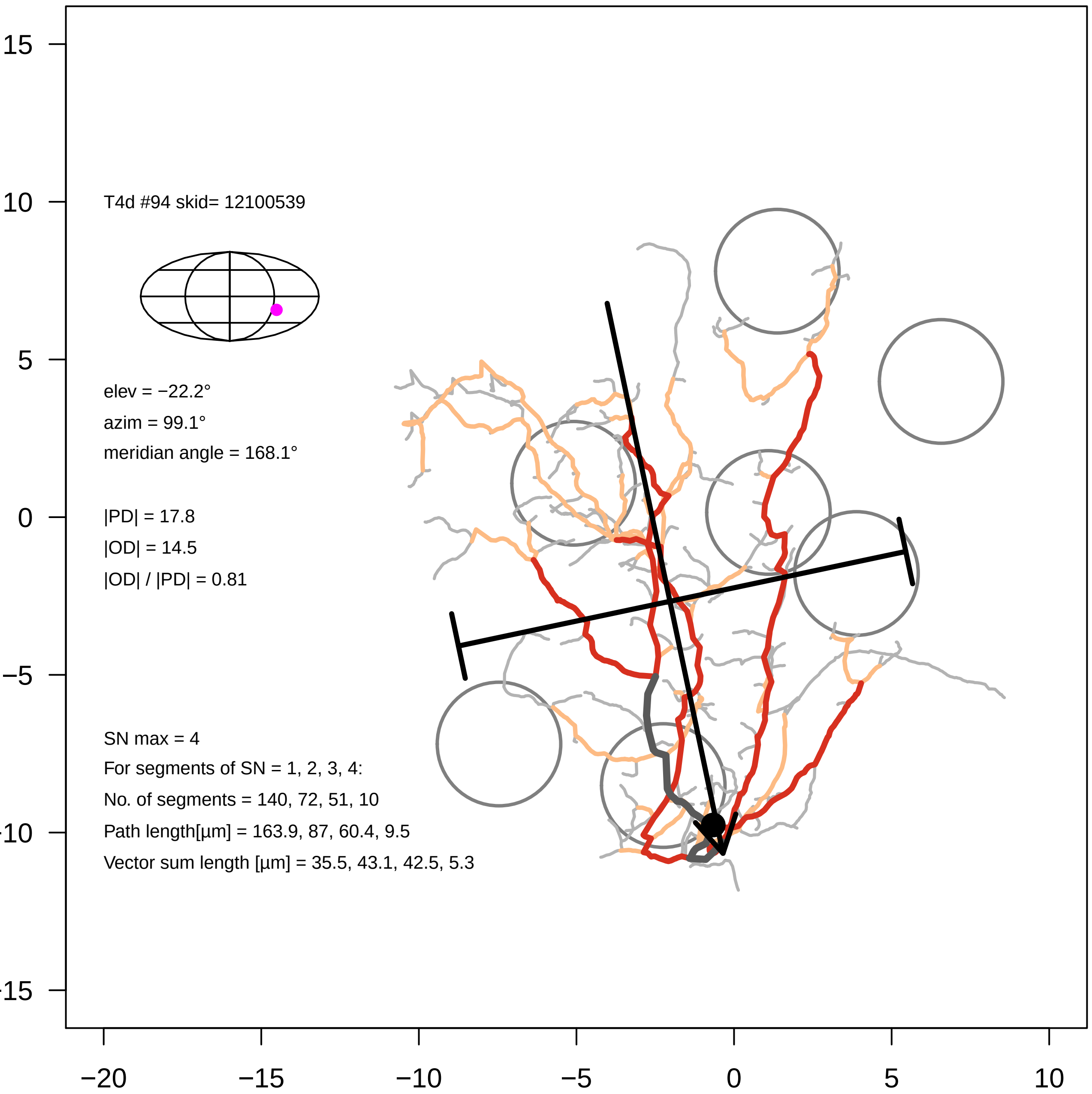

T4d #94 skid= 12100539
elev = −22.2°
azim = 99.1°
meridian angle = 168.1°
|PD| = 17.8
|OD| = 14.5
|OD| / |PD| = 0.81
SN max = 4
For segments of SN = 1, 2, 3, 4:
No. of segments = 140, 72, 51, 10
Path length[µm] = 163.9, 87, 60.4, 9.5
Vector sum length [µm] = 35.5, 43.1, 42.5, 5.3
15
10
5
0
−5
-10
-15
−20
−15
−10
−5
0
5
10

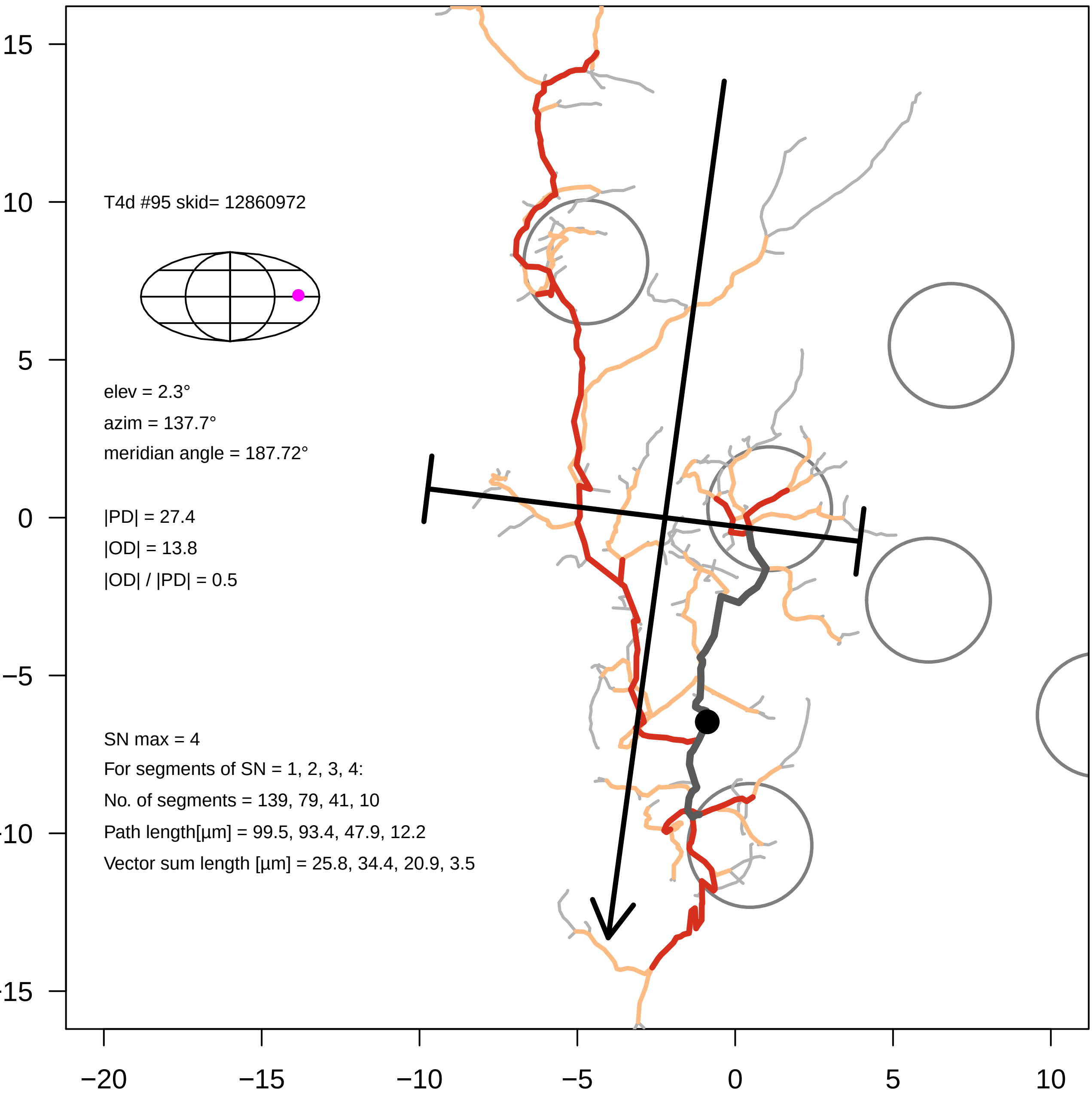
T4d #95 skid= 12860972
elev = 2.3°
azim = 137.7°
meridian angle = 187.72°
|PD| = 27.4
|OD| = 13.8
|OD| / |PD| = 0.5
SN max = 4
For segments of SN = 1, 2, 3, 4:
No. of segments = 139, 79, 41, 10
Path length[µm] = 99.5, 93.4, 47.9, 12.2
Vector sum length [µm] = 25.8, 34.4, 20.9, 3.5
15
10
5
0
−5
−10
−15
−20
−15
−10
−5
0
5
10

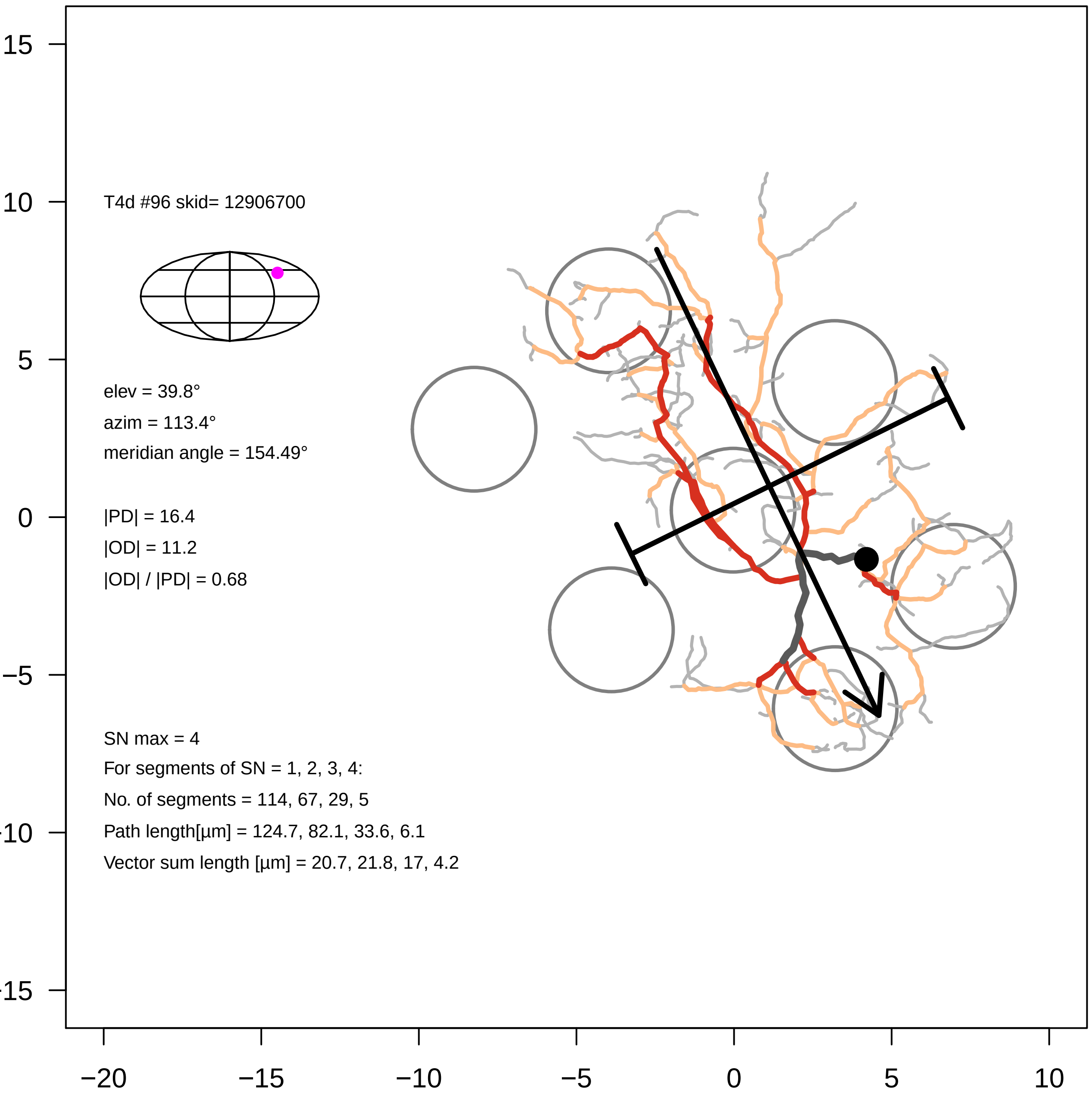
T4d #96 skid= 12906700
elev = 39.8°
azim = 113.4°
meridian angle = 154.49°
|PD| = 16.4
|OD| = 11.2
|OD| / |PD| = 0.68
SN max = 4
For segments of SN = 1, 2, 3, 4:
No. of segments = 114, 67, 29, 5
Path length[µm] = 124.7, 82.1, 33.6, 6.1
Vector sum length [µm] = 20.7, 21.8, 17, 4.2
15
10
5
0
−5
−10
−15
−20
−15
−10
−5
0
5
10

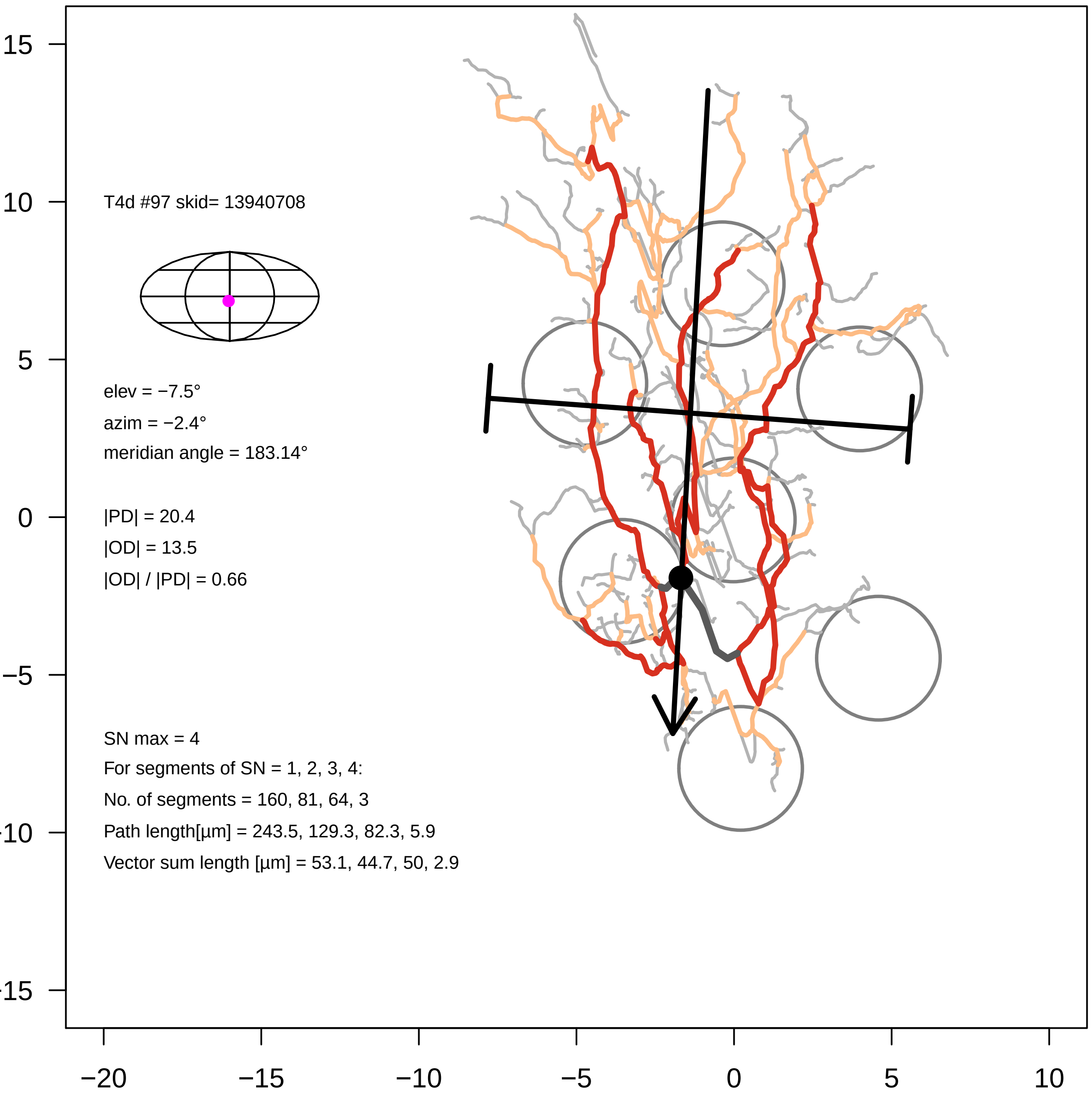
T4d #97 skid= 13940708
elev = −7.5°
azim = −2.4°
meridian angle = 183.14°
|PD| = 20.4
|OD| = 13.5
|OD| / |PD| = 0.66
SN max = 4
For segments of SN = 1, 2, 3, 4:
No. of segments = 160, 81, 64, 3
Path length[µm] = 243.5, 129.3, 82.3, 5.9
Vector sum length [µm] = 53.1, 44.7, 50, 2.9
15
10
5
0
−5
−10
−15
−20
−15
−10
−5
0
5
10

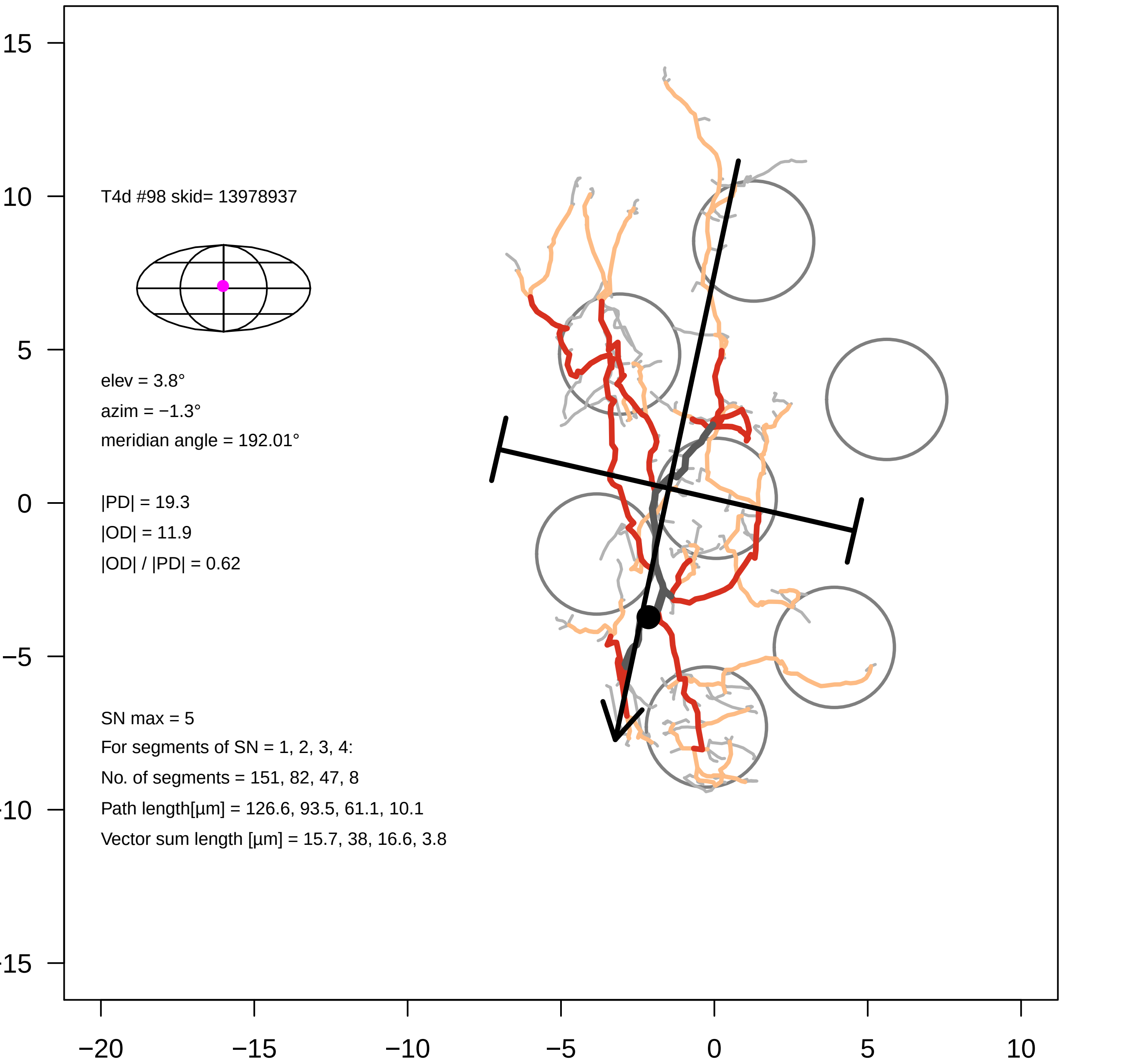

T4d #98 skid= 13978937
elev = 3.8°
azim = −1.3°
meridian angle = 192.01°
|PD| = 19.3
|OD| = 11.9
|OD| / |PD| = 0.62
SN max = 5
For segments of SN = 1, 2, 3, 4:
No. of segments = 151, 82, 47, 8
Path length[µm] = 126.6, 93.5, 61.1, 10.1
Vector sum length [µm] = 15.7, 38, 16.6, 3.8
15
10
5
0
−5
−15
−20
−15
−10
−5
0
5
10

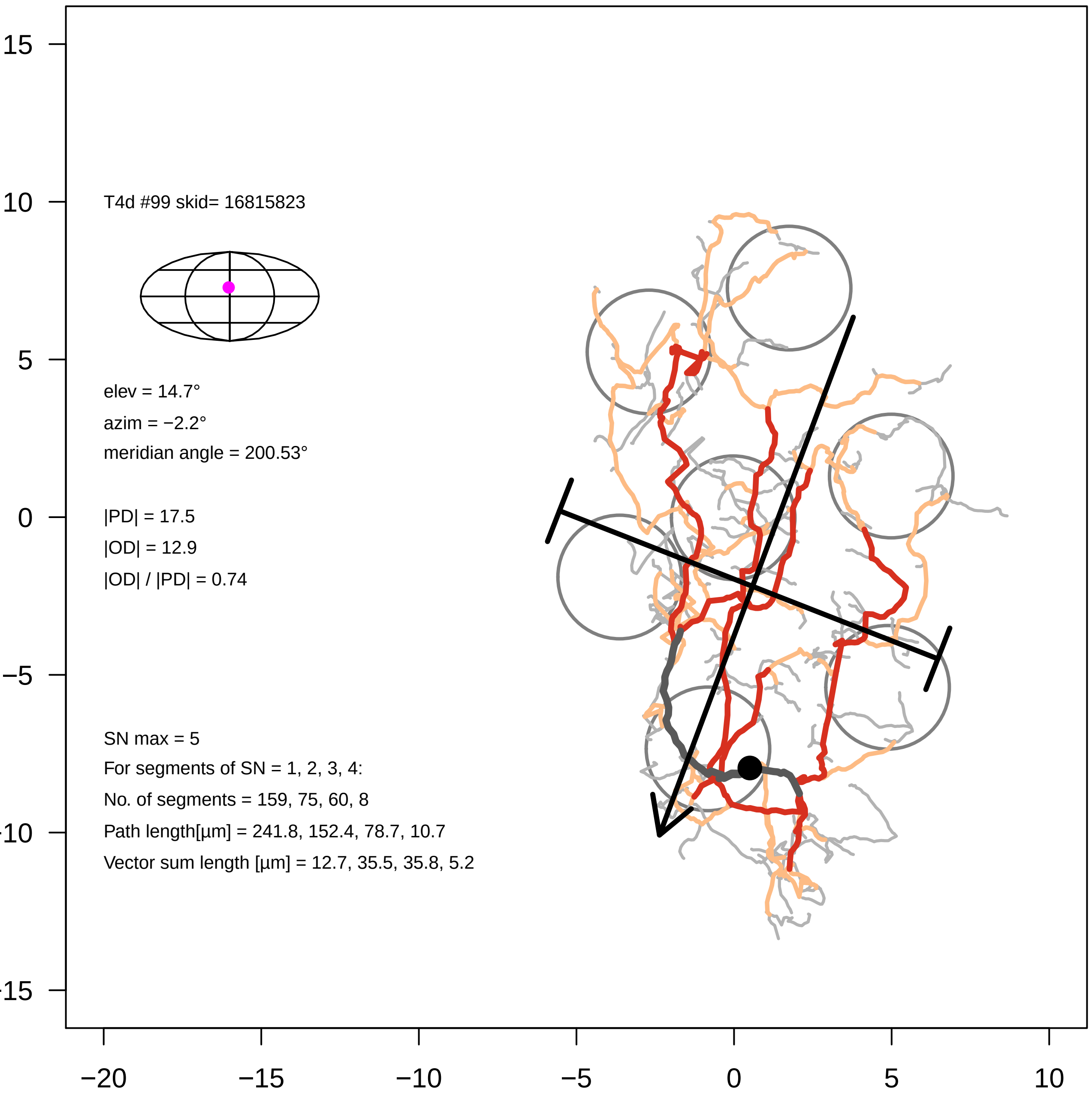

T4d #99 skid= 16815823
elev = 14.7°
azim = −2.2°
meridian angle = 200.53°
|PD| = 17.5
|OD| = 12.9
|OD| / |PD| = 0.74
SN max = 5
For segments of SN = 1, 2, 3, 4:
No. of segments = 159, 75, 60, 8
Path length[µm] = 241.8, 152.4, 78.7, 10.7
Vector sum length [µm] = 12.7, 35.5, 35.8, 5.2
15
10
5
0
−5
−10
−15
−20
−15
−10
−5
0
5
10

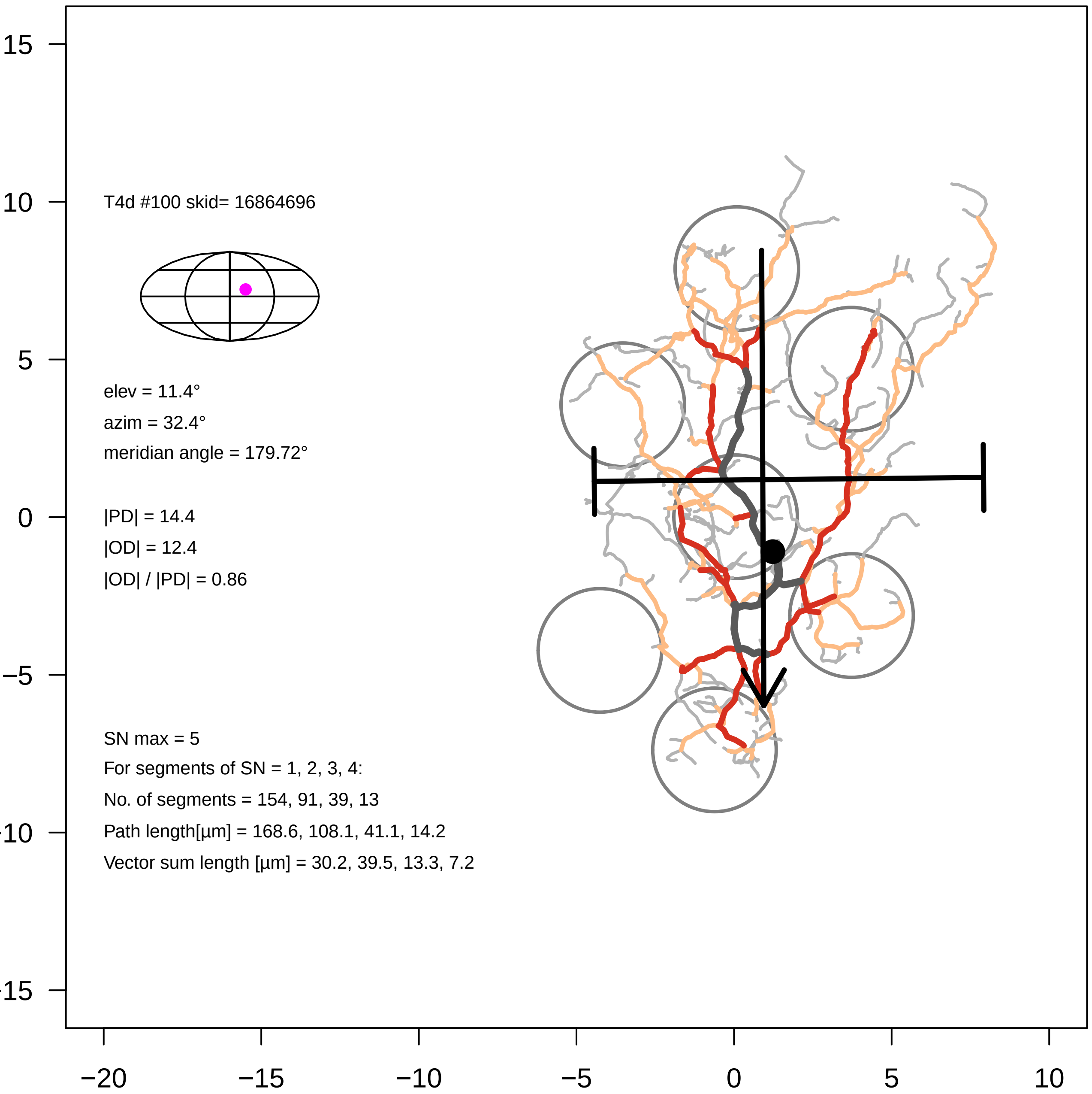

T4d #100 skid= 16864696
elev = 11.4°
azim = 32.4°
meridian angle = 179.72°
|PD| = 14.4
|OD| = 12.4
|OD| / |PD| = 0.86
SN max = 5
For segments of SN = 1, 2, 3, 4:
No. of segments = 154, 91, 39, 13
Path length[µm] = 168.6, 108.1, 41.1, 14.2
Vector sum length [µm] = 30.2, 39.5, 13.3, 7.2
15
10
5
0
−5
−10
−15
−20
−15
−10
−5
0
5
10

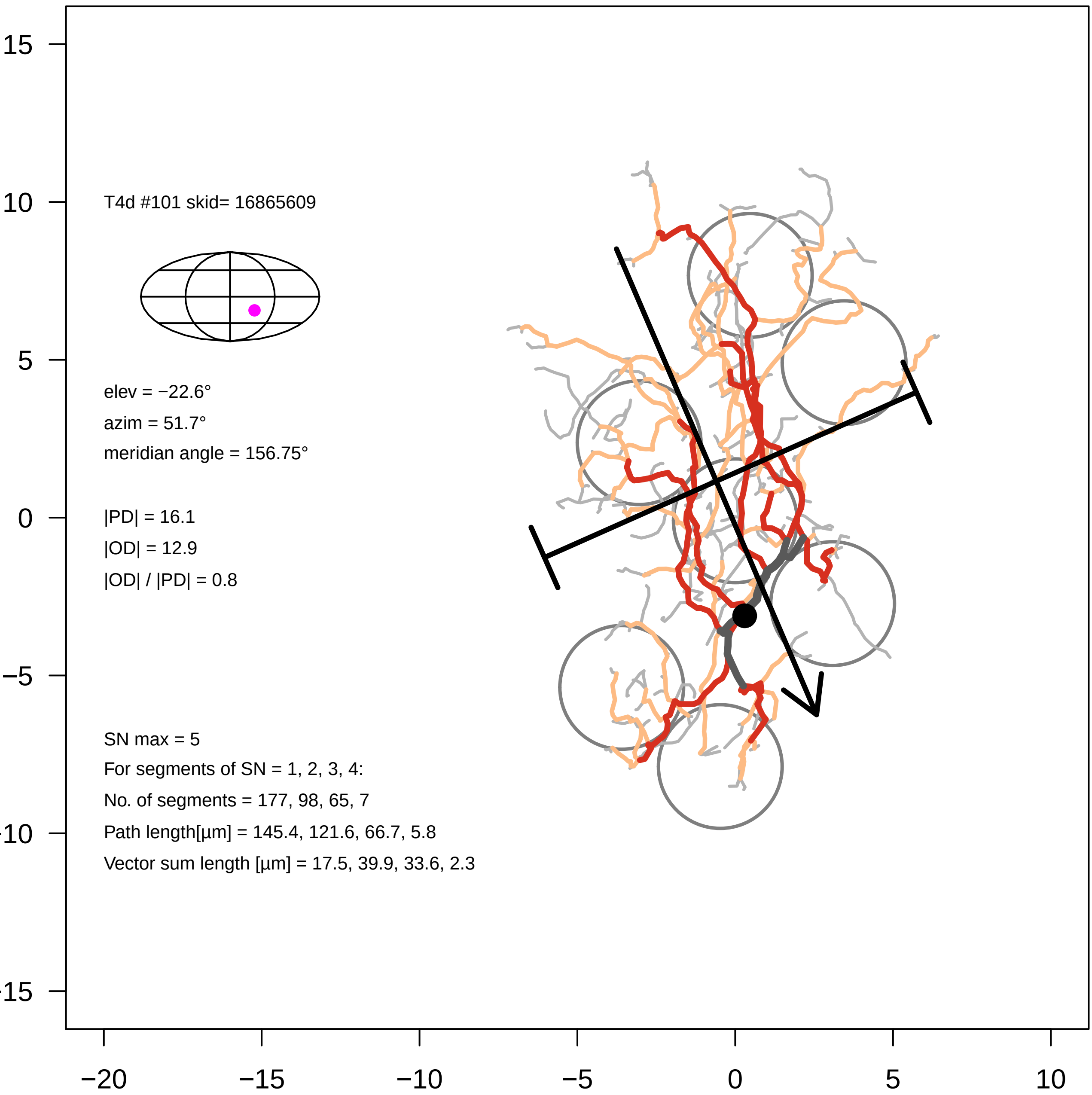

T4d #101 skid= 16865609
elev = −22.6°
azim = 51.7°
meridian angle = 156.75°
|PD| = 16.1
|OD| = 12.9
|OD| / |PD| = 0.8
SN max = 5
For segments of SN = 1, 2, 3, 4:
No. of segments = 177, 98, 65, 7
Path length[µm] = 145.4, 121.6, 66.7, 5.8
Vector sum length [µm] = 17.5, 39.9, 33.6, 2.3

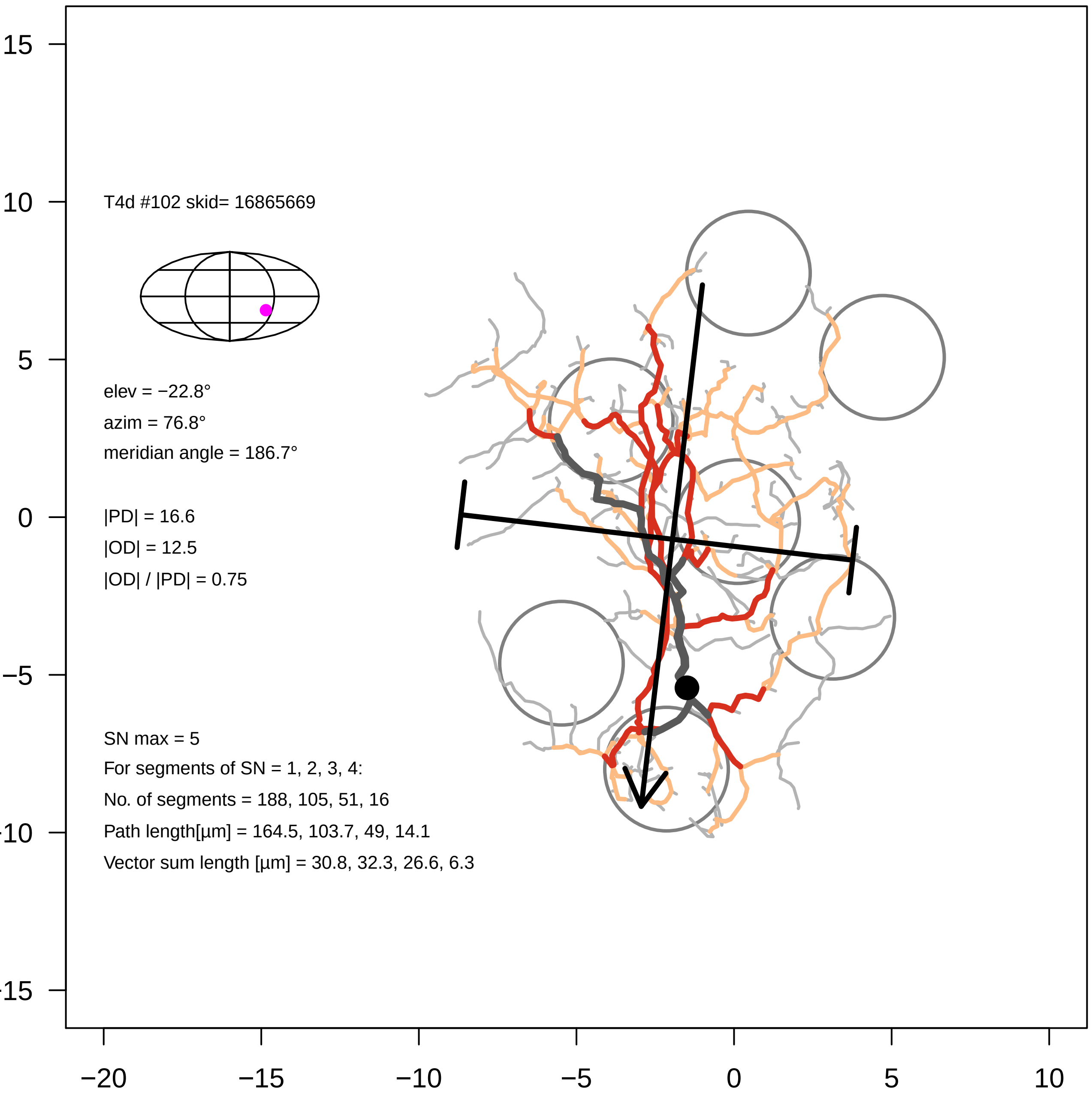
T4d #102 skid= 16865669
elev = −22.8°
azim = 76.8°
meridian angle = 186.7°
|PD| = 16.6
|OD| = 12.5
|OD| / |PD| = 0.75
SN max = 5
For segments of SN = 1, 2, 3, 4:
No. of segments = 188, 105, 51, 16
Path length[µm] = 164.5, 103.7, 49, 14.1
Vector sum length [µm] = 30.8, 32.3, 26.6, 6.3
15
10
5
0
−5
−10
−15
−20
−15
−10
−5
0
5
10

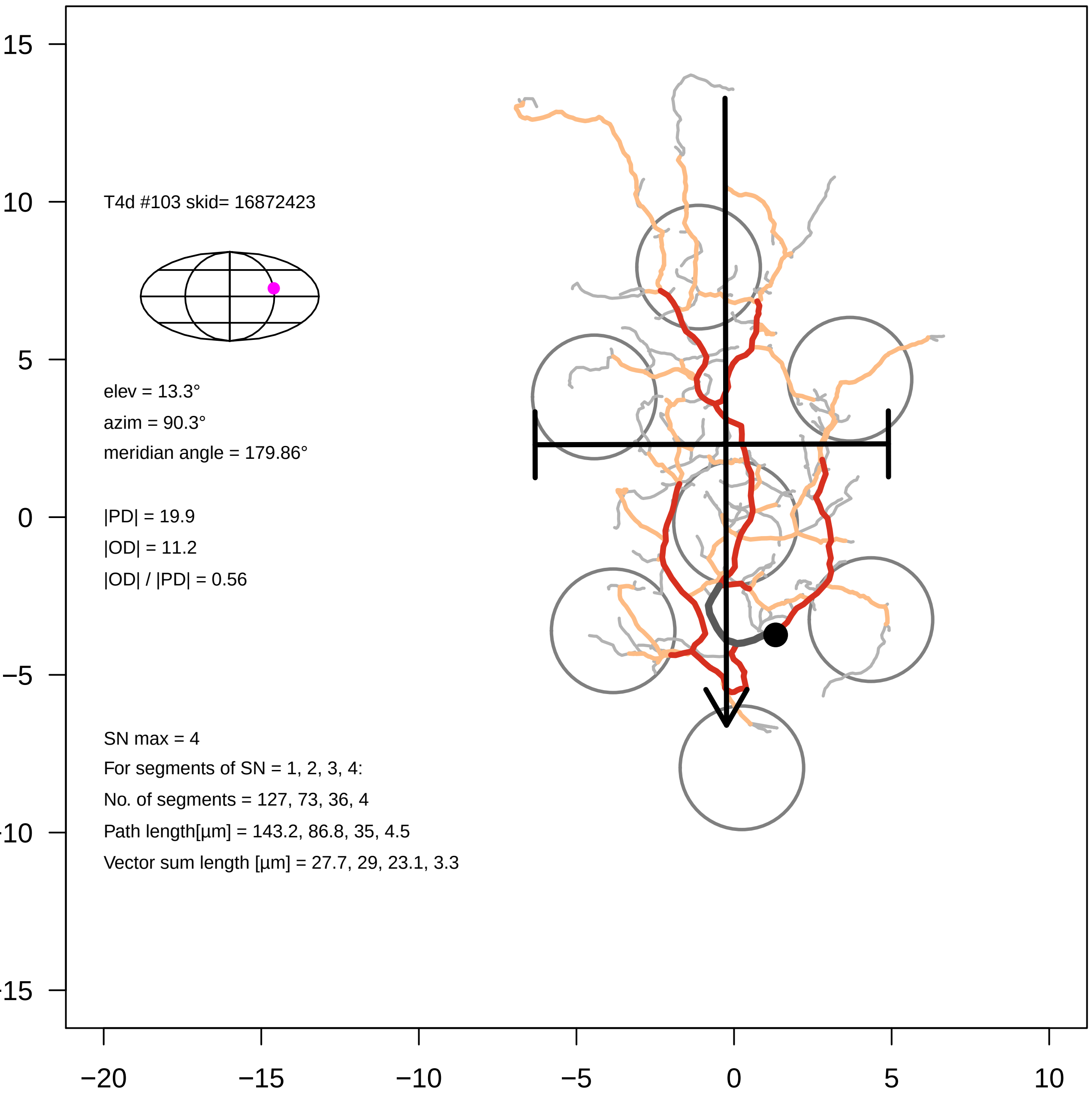

T4d #103 skid= 16872423
elev = 13.3°
azim = 90.3°
meridian angle = 179.86°
|PD| = 19.9
|OD| = 11.2
|OD| / |PD| = 0.56
SN max = 4
For segments of SN = 1, 2, 3, 4:
No. of segments = 127, 73, 36, 4
Path length[µm] = 143.2, 86.8, 35, 4.5
Vector sum length [µm] = 27.7, 29, 23.1, 3.3
15
10
5
0
−5
−10
−15
−20
−15
−10
−5
0
5
10

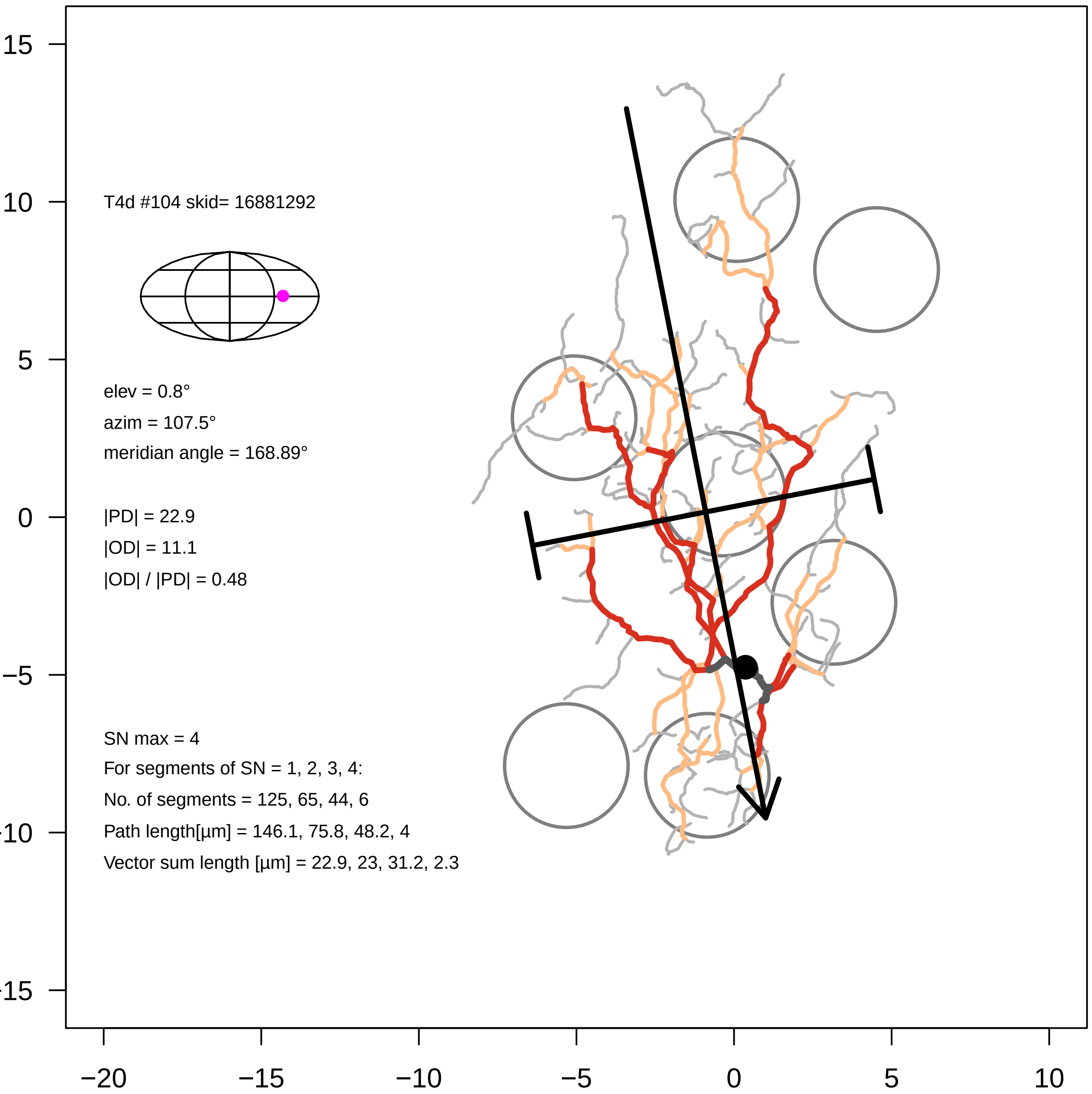

T4d #104 skid= 16881292
elev = 0.8°
azim = 107.5°
meridian angle = 168.89°
|PD| = 22.9
|OD| = 11.1
|OD| / |PD| = 0.48
SN max = 4
For segments of SN = 1, 2, 3, 4:
No. of segments = 125, 65, 44, 6
Path length[µm] = 146.1, 75.8, 48.2, 4
Vector sum length [µm] = 22.9, 23, 31.2, 2.3
15
10
5
0
−5
−20
−15
−10
−5
0
5
10

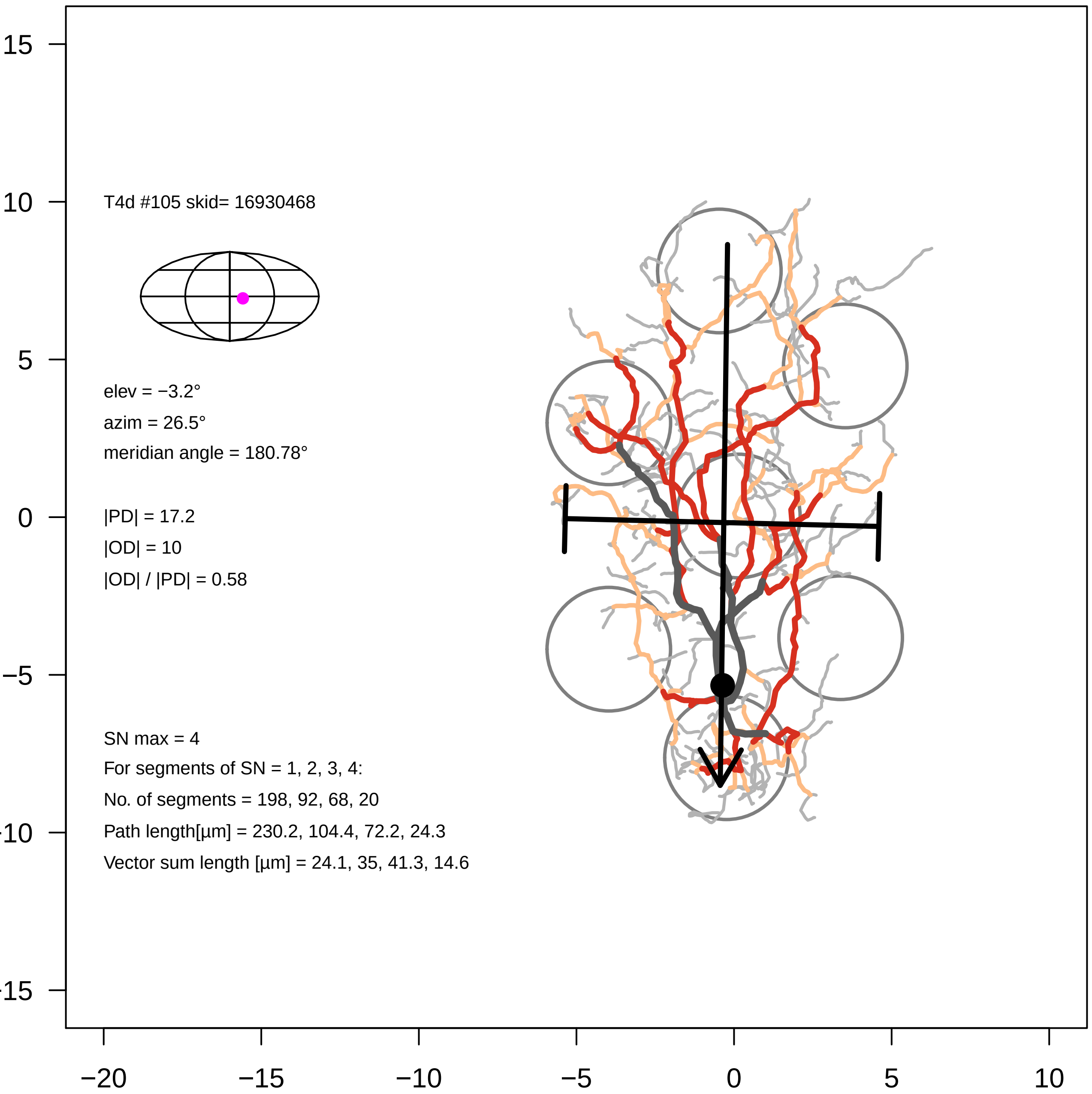
T4d #105 skid= 16930468
elev = −3.2°
azim = 26.5°
meridian angle = 180.78°
|PD| = 17.2
|OD| = 10
|OD| / |PD| = 0.58
SN max = 4
For segments of SN = 1, 2, 3, 4:
No. of segments = 198, 92, 68, 20
Path length[µm] = 230.2, 104.4, 72.2, 24.3
Vector sum length [µm] = 24.1, 35, 41.3, 14.6
15
10
5
0
−5
−10
−15
−20
−15
−10
−5
0
5
10

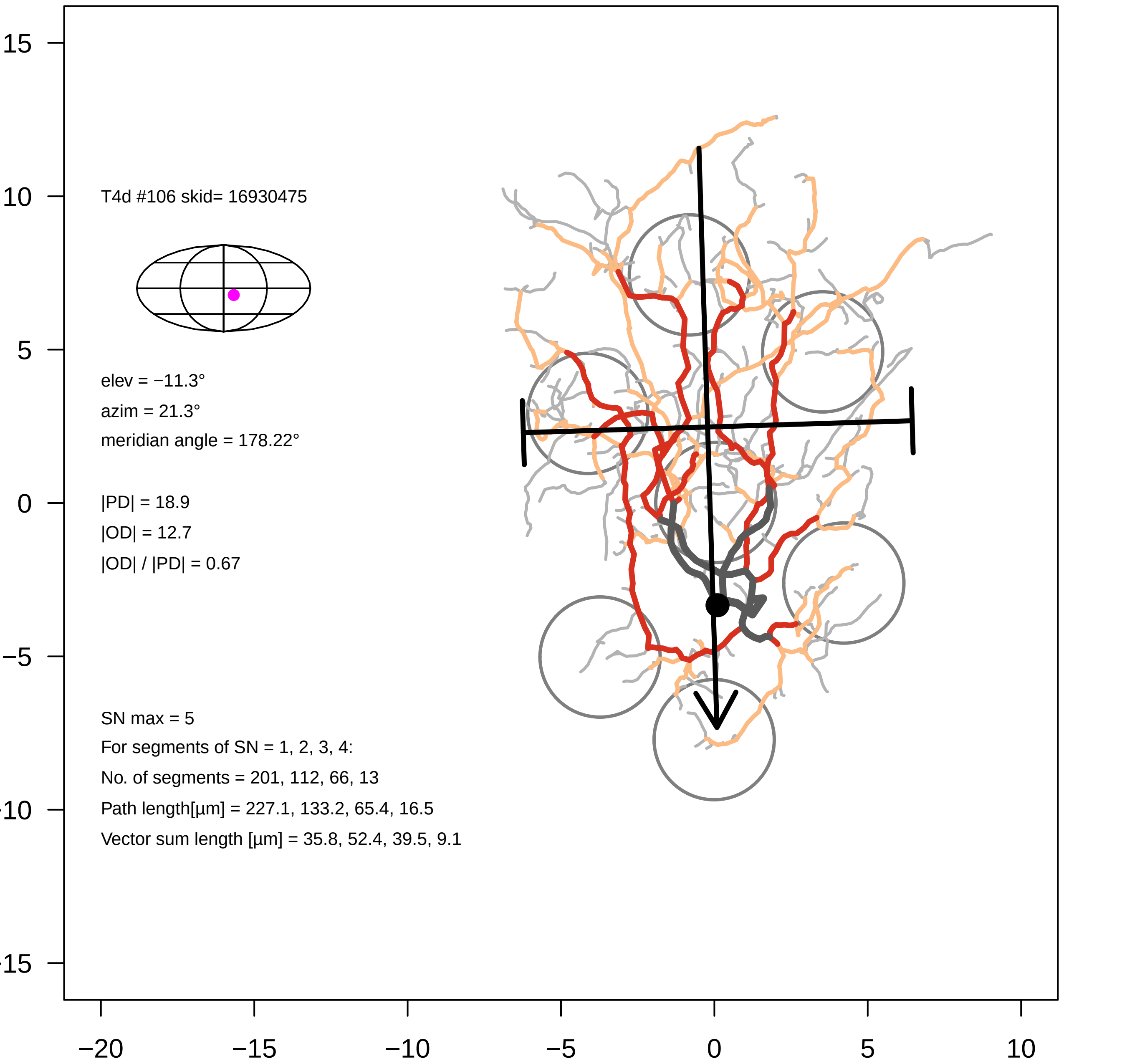
T4d #106 skid= 16930475
elev = −11.3°
azim = 21.3°
meridian angle = 178.22°
|PD| = 18.9
|OD| = 12.7
|OD| / |PD| = 0.67
SN max = 5
For segments of SN = 1, 2, 3, 4:
No. of segments = 201, 112, 66, 13
Path length[µm] = 227.1, 133.2, 65.4, 16.5
Vector sum length [µm] = 35.8, 52.4, 39.5, 9.1
15
10
5
0
−5
−10
−15
−20
−15
−10
−5
0
5
10

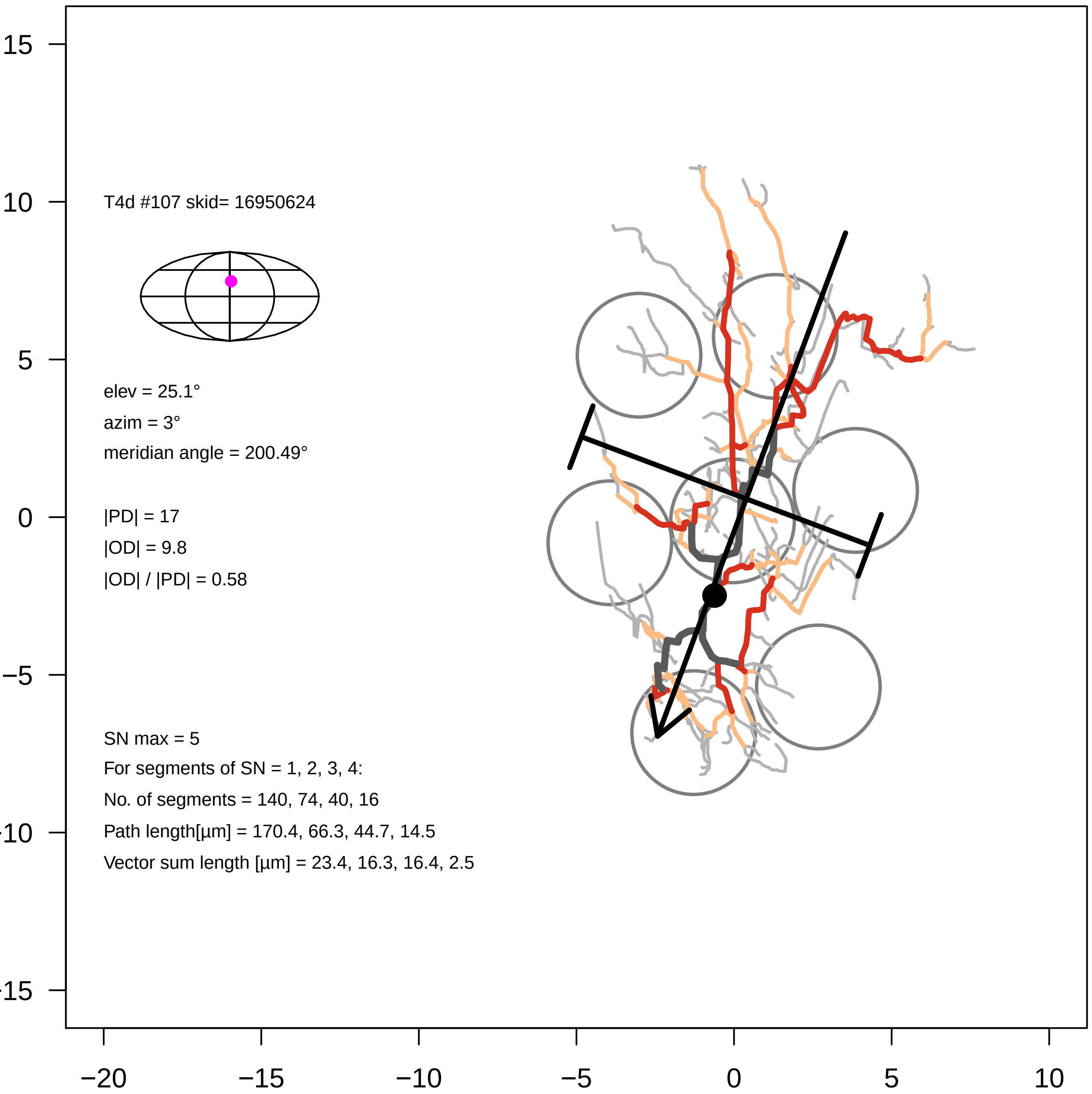

T4d #107 skid= 16950624
elev = 25.1°
azim = 3°
meridian angle = 200.49°
|PD| = 17
|OD| = 9.8
|OD| / |PD| = 0.58
SN max = 5
For segments of SN = 1, 2, 3, 4:
No. of segments = 140, 74, 40, 16
Path length[µm] = 170.4, 66.3, 44.7, 14.5
Vector sum length [µm] = 23.4, 16.3, 16.4, 2.5
15
10
5
0
−5
−10
−15
−20
−15
−10
−5
0
5
10

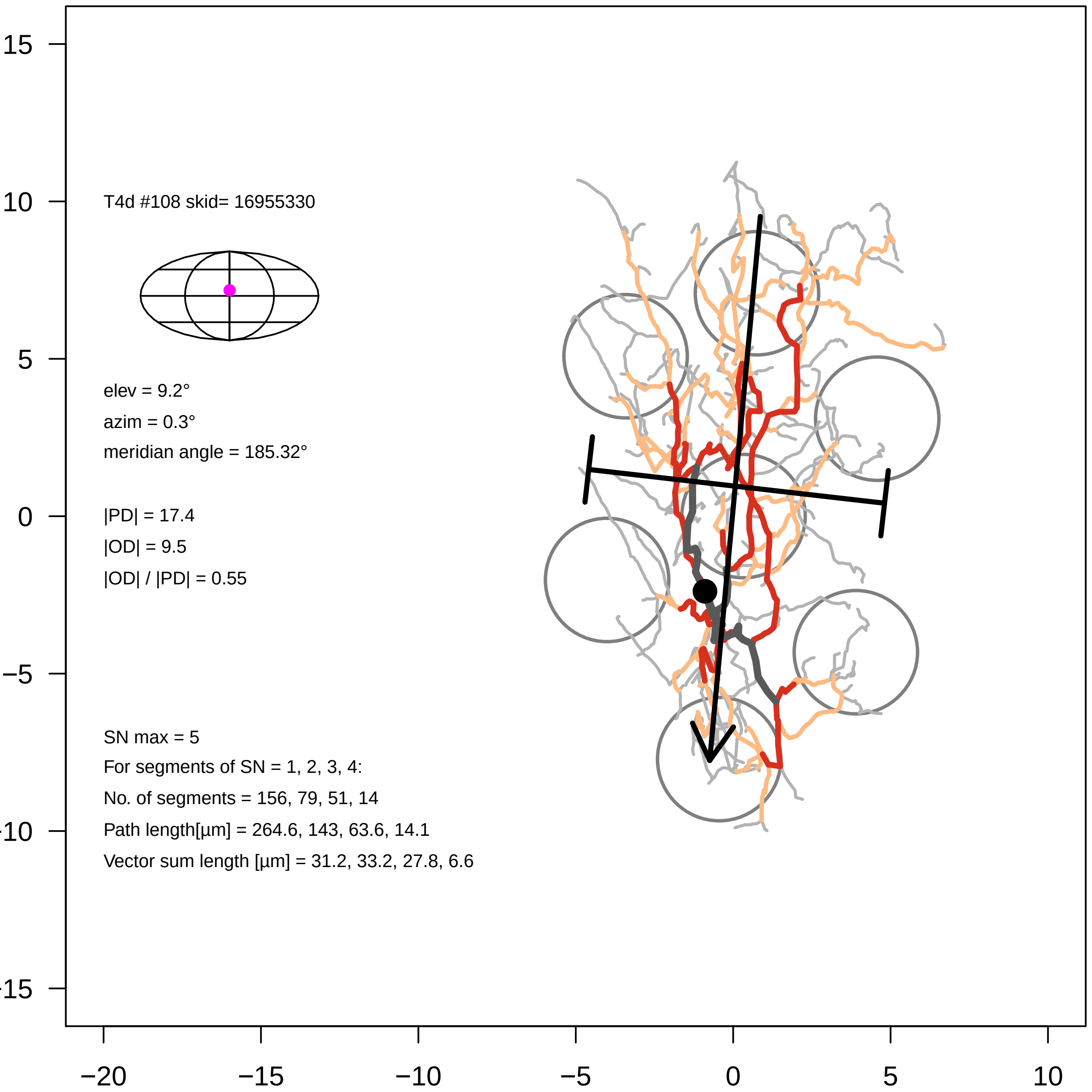

T4d #108 skid= 16955330
elev = 9.2°
azim = 0.3°
meridian angle = 185.32°
|PD| = 17.4
|OD| = 9.5
|OD| / |PD| = 0.55
SN max = 5
For segments of SN = 1, 2, 3, 4:
No. of segments = 156, 79, 51, 14
Path length[µm] = 264.6, 143, 63.6, 14.1
Vector sum length [µm] = 31.2, 33.2, 27.8, 6.6
15
10
5
0
−5
−10
−15
−20
−15
−10
−5
0
5
10

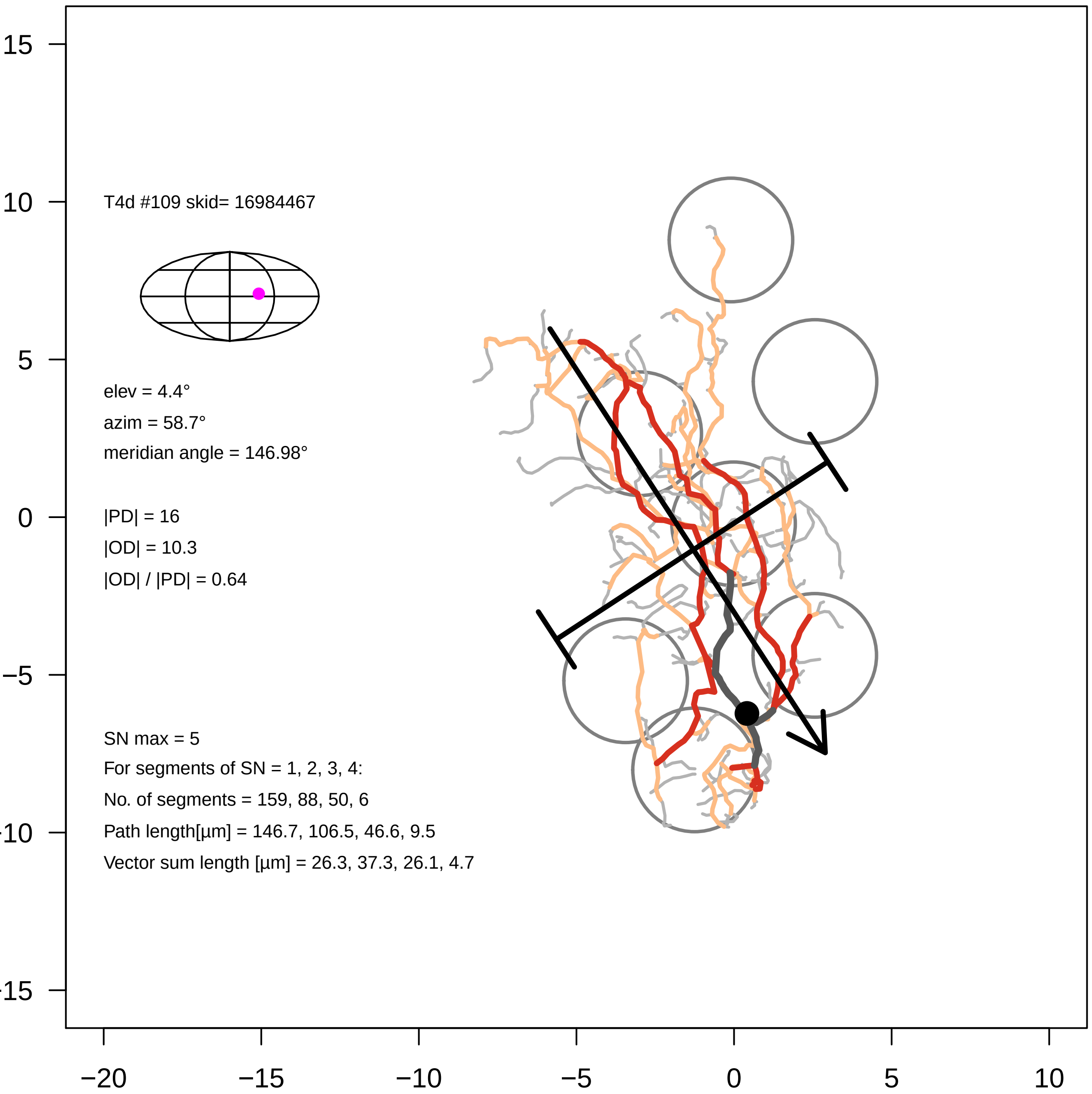

15
10
5
0
−5
−10
−15
−20
−15
−10
−5
0
5
10
T4d #109 skid= 16984467
elev = 4.4°
azim = 58.7°
meridian angle = 146.98°
|PD| = 16
|OD| = 10.3
|OD| / |PD| = 0.64
SN max = 5
For segments of SN = 1, 2, 3, 4:
No. of segments = 159, 88, 50, 6
Path length[µm] = 146.7, 106.5, 46.6, 9.5
Vector sum length [µm] = 26.3, 37.3, 26.1, 4.7

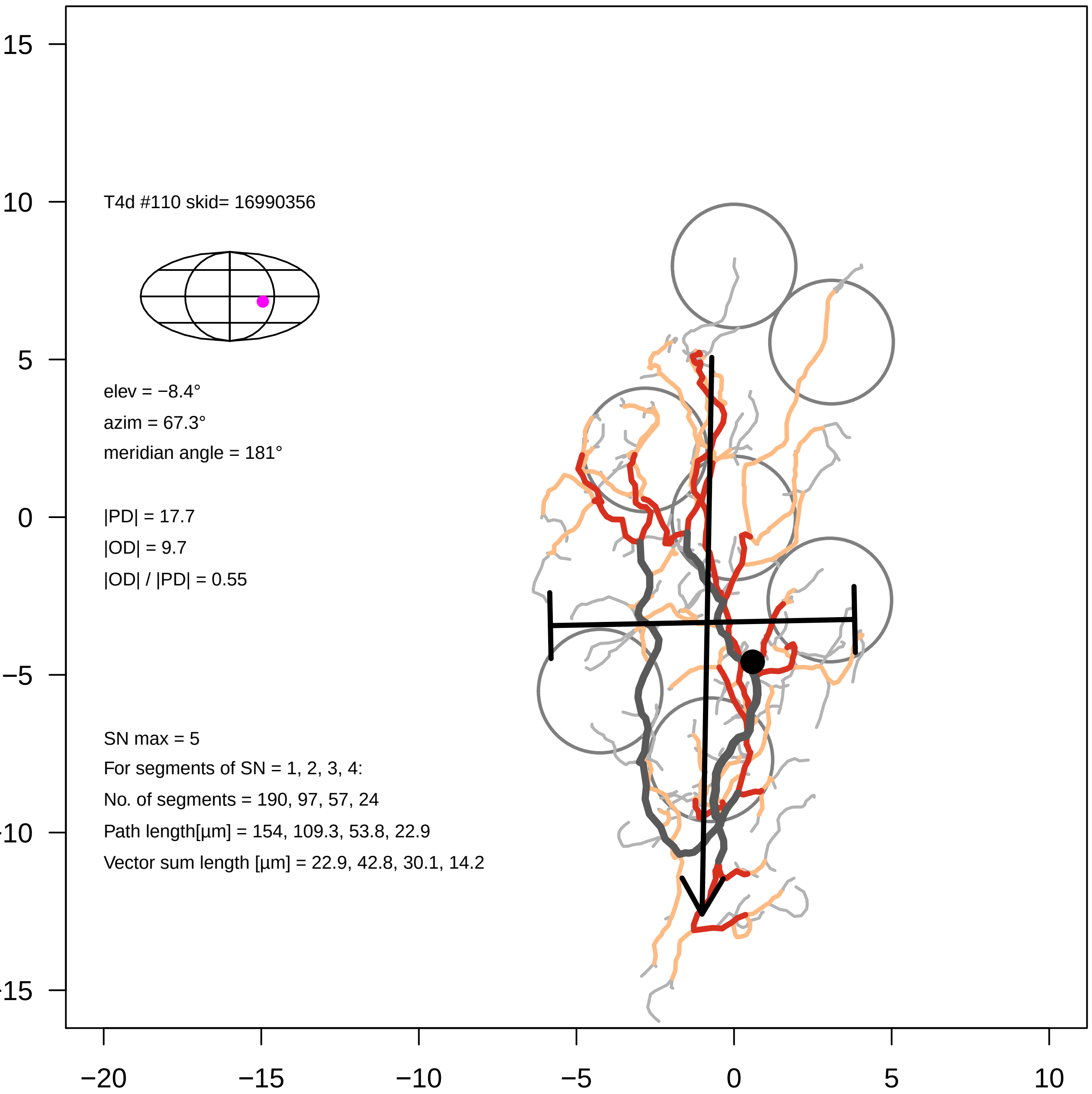

T4d #110 skid= 16990356
elev = −8.4°
azim = 67.3°
meridian angle = 181°
|PD| = 17.7
|OD| = 9.7
|OD| / |PD| = 0.55
SN max = 5
For segments of SN = 1, 2, 3, 4:
No. of segments = 190, 97, 57, 24
Path length[µm] = 154, 109.3, 53.8, 22.9
Vector sum length [µm] = 22.9, 42.8, 30.1, 14.2
15
10
5
0
−5
−10
−15
−20
−15
−10
−5
0
5
10

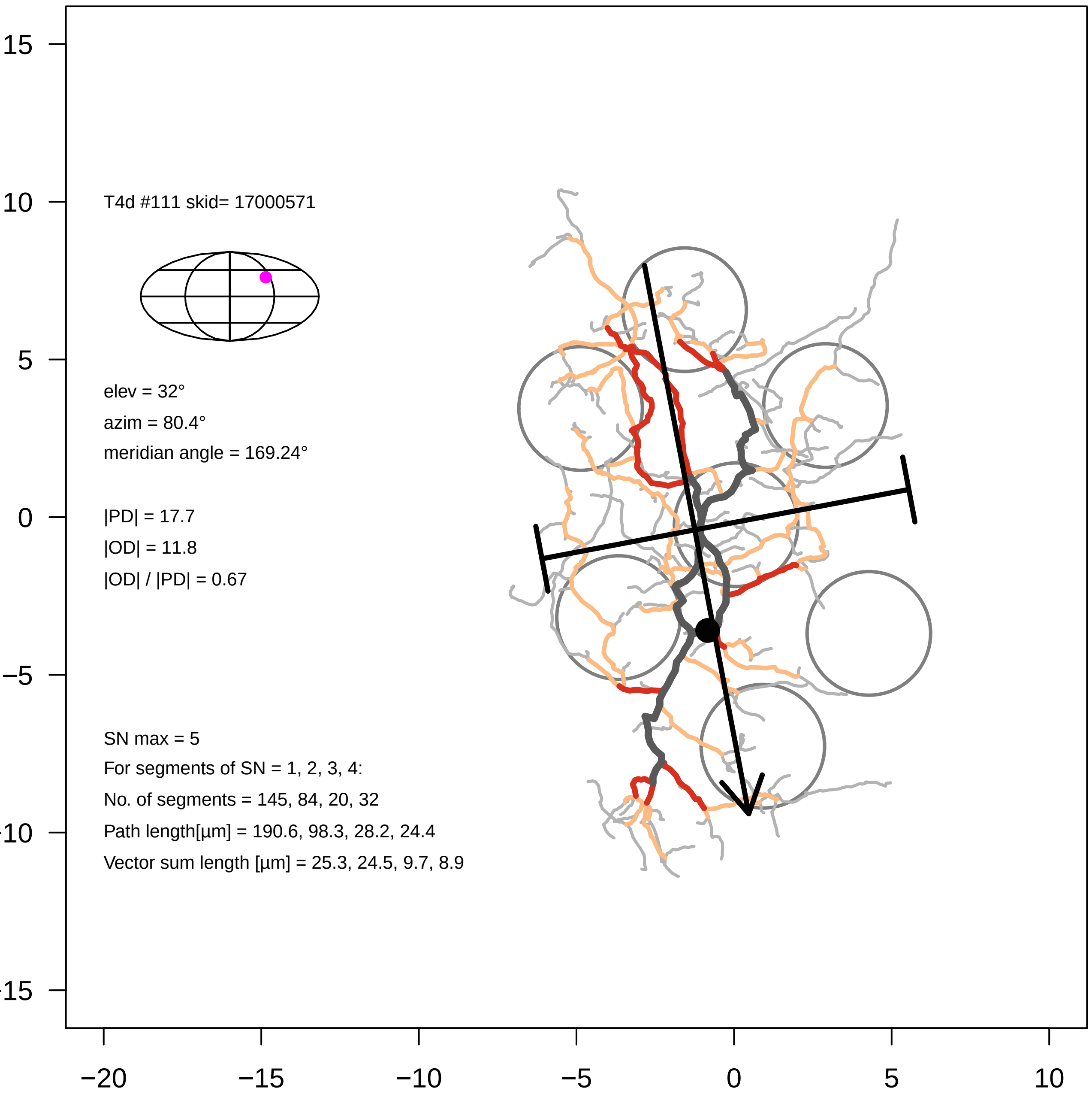

T4d #111 skid= 17000571
elev = 32°
azim = 80.4°
meridian angle = 169.24°
|PD| = 17.7
|OD| = 11.8
|OD| / |PD| = 0.67
SN max = 5
For segments of SN = 1, 2, 3, 4:
No. of segments = 145, 84, 20, 32
Path length[µm] = 190.6, 98.3, 28.2, 24.4
Vector sum length [µm] = 25.3, 24.5, 9.7, 8.9
15
10
5
0
−5
−10
−15
−20
−15
−10
−5
0
5
10

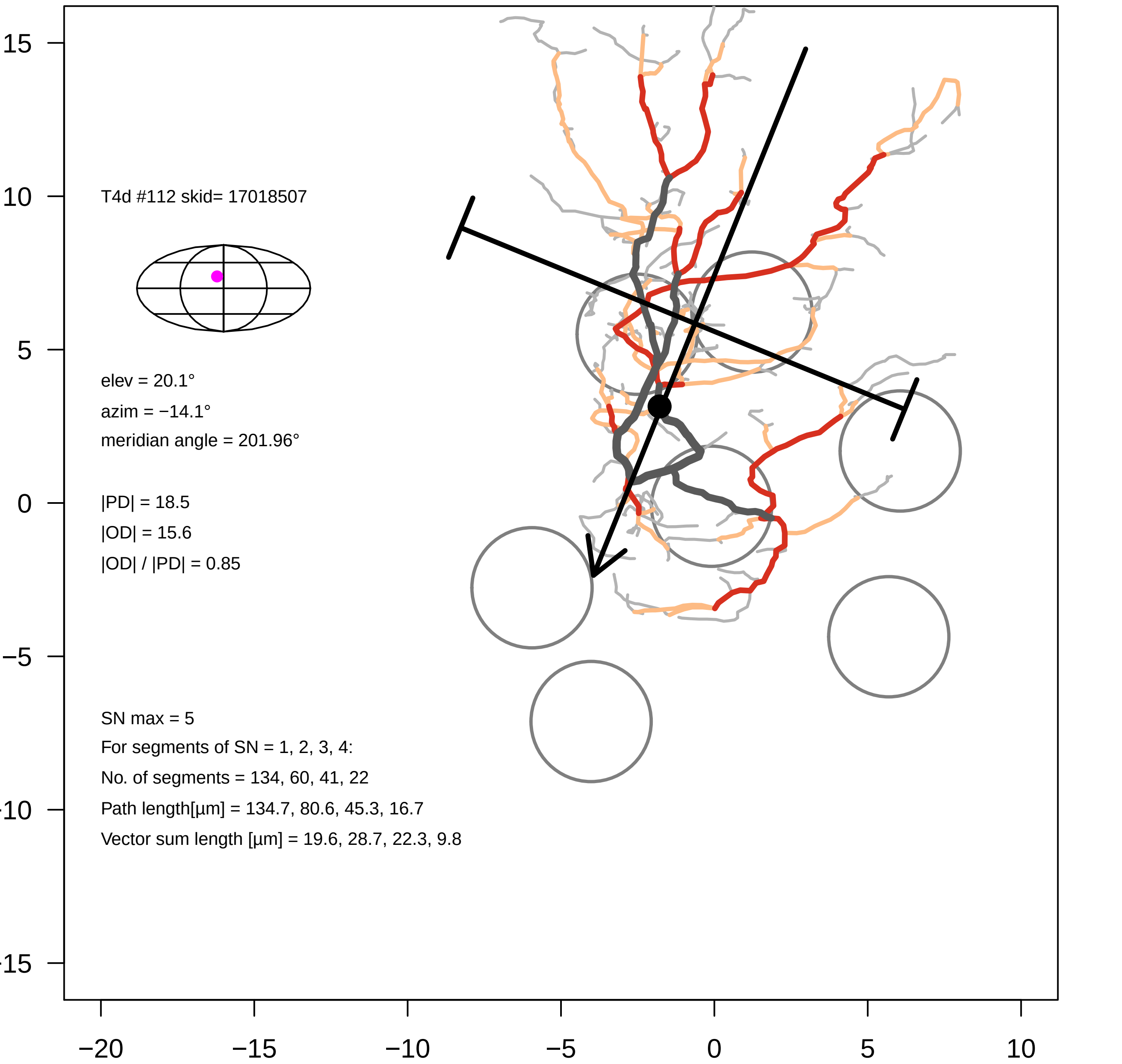

T4d #112 skid= 17018507
elev = 20.1°
azim = −14.1°
meridian angle = 201.96°
|PD| = 18.5
|OD| = 15.6
|OD| / |PD| = 0.85
SN max = 5
For segments of SN = 1, 2, 3, 4:
No. of segments = 134, 60, 41, 22
Path length[µm] = 134.7, 80.6, 45.3, 16.7
Vector sum length [µm] = 19.6, 28.7, 22.3, 9.8

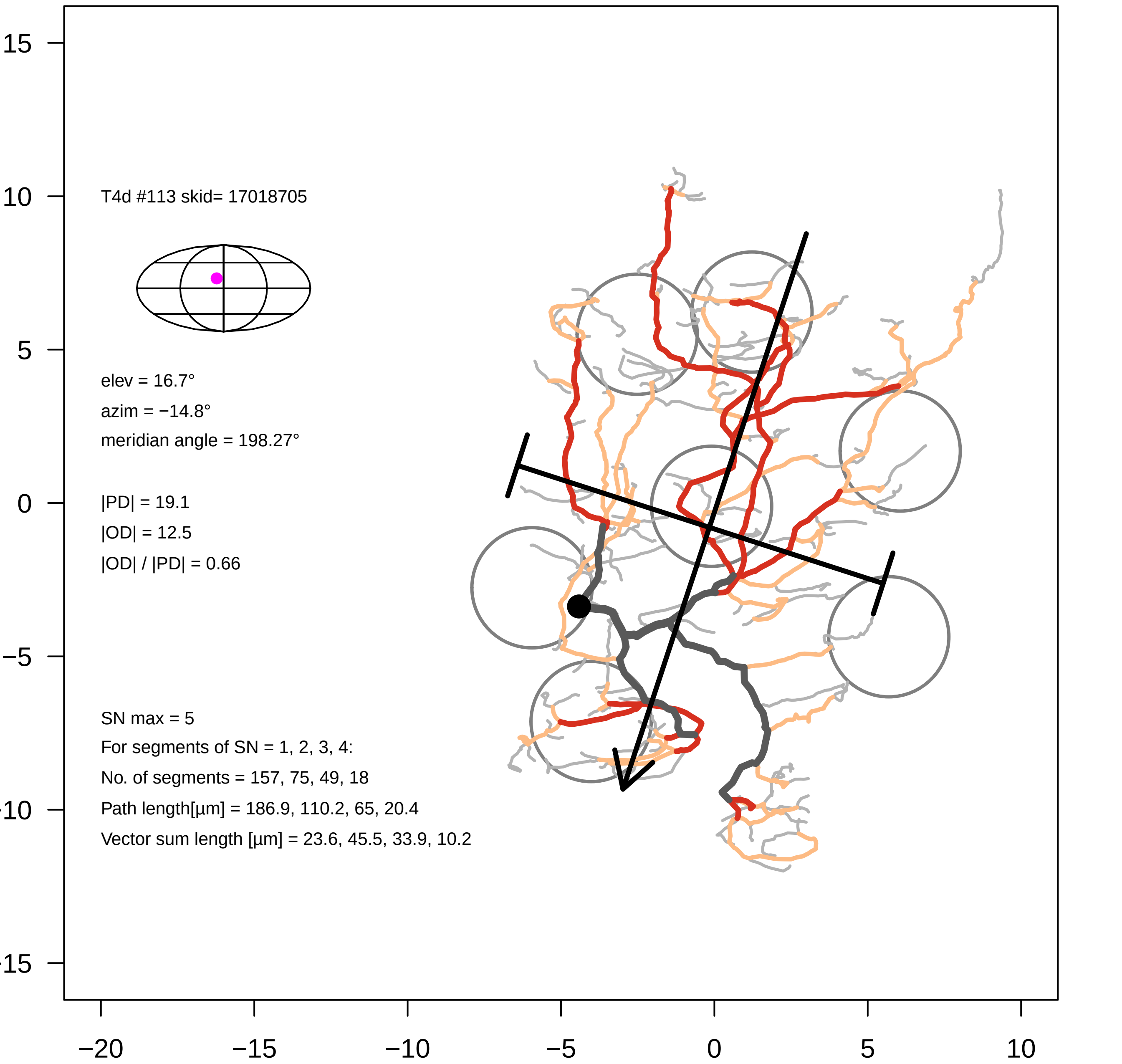

T4d #113 skid= 17018705
elev = 16.7°
azim = −14.8°
meridian angle = 198.27°
|PD| = 19.1
|OD| = 12.5
|OD| / |PD| = 0.66
SN max = 5
For segments of SN = 1, 2, 3, 4:
No. of segments = 157, 75, 49, 18
Path length[µm] = 186.9, 110.2, 65, 20.4
Vector sum length [µm] = 23.6, 45.5, 33.9, 10.2
15
10
5
0
−5
−15
−20
−15
−10
−5
0
5
10

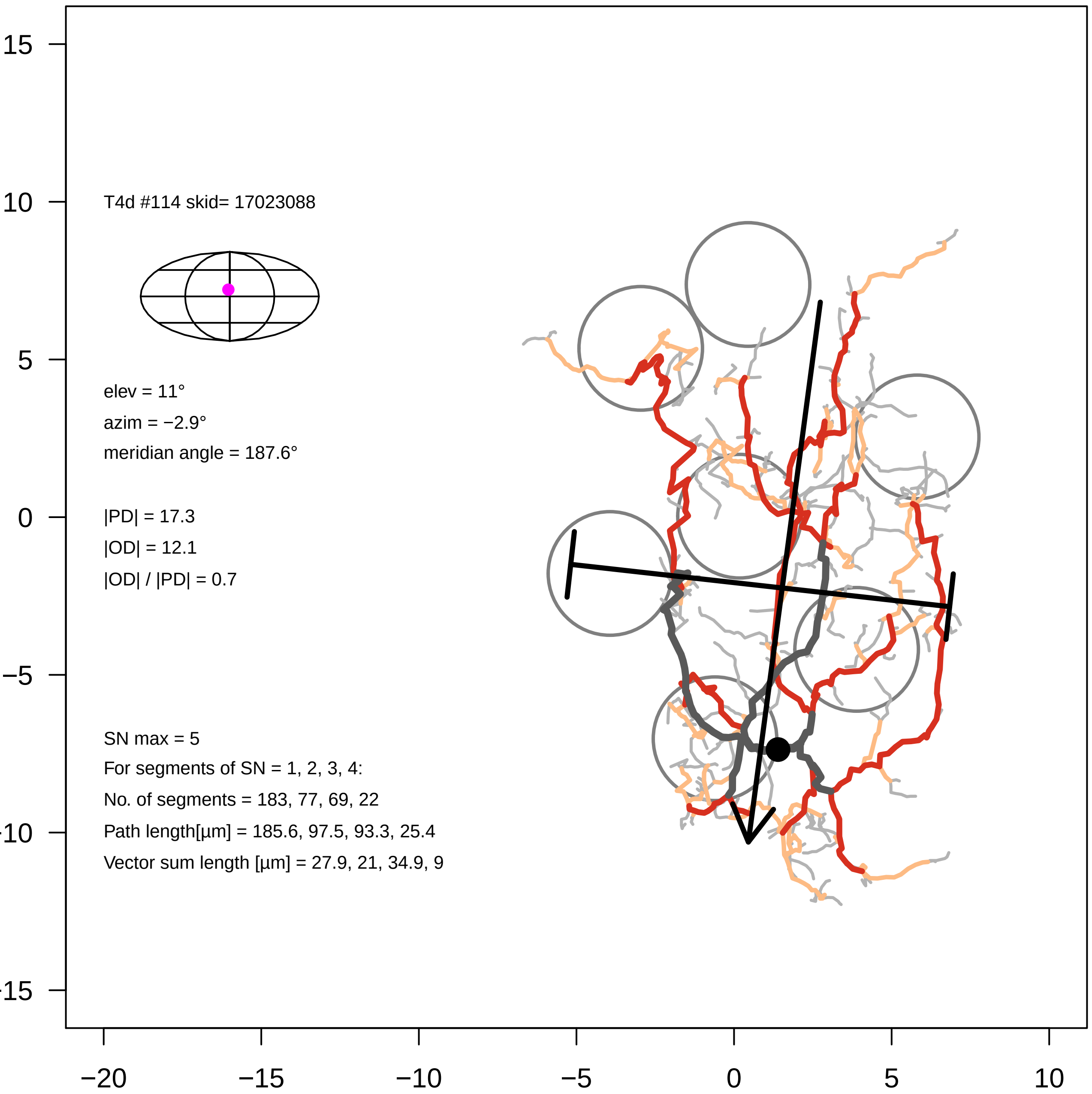
T4d #114 skid= 17023088
elev = 11°
azim = −2.9°
meridian angle = 187.6°
|PD| = 17.3
|OD| = 12.1
|OD| / |PD| = 0.7
SN max = 5
For segments of SN = 1, 2, 3, 4:
No. of segments = 183, 77, 69, 22
Path length[µm] = 185.6, 97.5, 93.3, 25.4
Vector sum length [µm] = 27.9, 21, 34.9, 9
15
10
5
0
−5
−20
−15
−10
−5
0
5
10
